# High Density Domain-Focused CRISPR Screens Reveal Novel Epigenetic Regulators of *HOX/MEIS* Activation in Acute Myeloid Leukemia

**DOI:** 10.1101/2022.12.12.519332

**Authors:** Karina Barbosa, Anagha Deshpande, Marlenne Perales, Ping Xiang, Rabi Murad, Anna Minkina, Neil Robertson, Fiorella Schischlik, Xue Lei, Younguk Sun, Adam Brown, Diana Amend, Irmela Jeremias, John G. Doench, R. Keith Humphries, Eytan Ruppin, Jay Shendure, Prashant Mali, Peter D Adams, Aniruddha J. Deshpande

## Abstract

Aberrant expression of stem-cell-associated genes is a common feature in acute myeloid leukemia (AML) and is linked to leukemic self-renewal and therapy resistance. Using AF10-rearranged leukemia as a prototypical example displaying a recurrent “*stemness*” network activated in AML, we screened for chromatin regulators that sustain aberrant activation of these networks. We deployed a CRISPR-Cas9 screen with a bespoke domain-focused library and identified several novel chromatin-modifying complexes as regulators of the TALE domain transcription factor MEIS1, a key leukemia stem cell (LSC)-associated gene. CRISPR droplet sequencing revealed that many of these MEIS1 regulators coordinately controlled the transcription of several AML oncogenes. In particular, we identified a novel role for the Tudor-domain containing chromatin reader protein SGF29 in the transcription of key AML oncogenes. Furthermore, SGF29 deletion impaired leukemogenesis in models representative of multiple AML subtypes. Our studies reveal a novel role for SGF29 as a non-oncogenic dependency in AML and identify the SGF29 Tudor domain as an attractive target for drug discovery.

## MAIN

Acute Myeloid Leukemia (AML) is a devastating form of blood cancer with a dismal survival rate ^1^. Current therapies are often associated with strong toxicity and undesirable side effects, highlighting the need for safer, more effective alternatives. A major challenge in the development of new drugs for AML is the heterogeneity of the disease at the molecular level ^2^. AML is composed of several morphologic and molecular subtypes, with distinct mutational profiles and clinical characteristics ^2^. However, there are molecular pathways that are commonly dysregulated across different AML subtypes. Studies have shown that a variety of upstream genetic alterations in AML, including mutations in *DNMT3* or nucleophosmin 1 (*NPM1*), and fusion products of several different chromosomal translocations result in the activation of the clustered homeobox (HOX) genes and their co-factors such as the three amino acid loop extension (TALE) homeobox gene *MEIS1* (reviewed in (1–4)). Specific *HOX* genes such as the *HOXA* and/or *HOXB* genes as well as *MEIS1* display significantly higher expression in a large proportion of AML samples compared to normal bone marrow cells ^3–6^. In particular, highly elevated expression of posterior *HOXA* or *B* genes and *MEIS1* can be observed in as many as two-thirds of AML with diverse mutational profiles ^7^. Studies conducted using both loss-of-function as well as gain-of-function approaches have demonstrated the importance of *HOX/MEIS* expression in leukemia pathogenesis in diverse subsets of AML including those bearing MLL-fusions (5,6), NUP98-fusions ^8^, and AF10-fusions ^9, 10^, to name a few. Importantly, ectopic *HOXA9* overexpression in murine hematopoietic stem and progenitor cells (HSPCs) causes a myeloproliferative phenotype in mice that can progress to AML upon *MEIS1*-co-expression^11, 12^. *MEIS1* is a critical co-factor of leukemia-associated HOXA transcription factors and is required for their full leukemogenic capability ^11, 12^. MEIS1 has also been shown to act as a rate-limiting regulator of leukemia stem cell (LSC) activity^13^. Taken together, there is compelling evidence implicating the *HOX/MEIS* oncoproteins as a key node integrating a variety of functionally distinct oncogenic insults in AML.

Thus, there is compelling preclinical evidence that pharmacological targeting of the *HOX/MEIS* pathway may yield therapeutic benefit in multiple, genetically heterogeneous AML sub-types. Since the HOX/MEIS proteins are DNA-binding transcription factors, they have proven difficult to target directly using traditional drug-development methods. We reasoned that a detailed and systematic identification of epigenetic modulators critical for sustaining HOX/MEIS-activation in AML would help identify novel nodes for targeted drug-discovery campaigns aimed at this clinically important pathway. To this end, we conducted pharmacological and CRISPR-based genetic screens using enhanced green fluorescence protein (eGFP) tagged *MEIS1* AML cells ^14^. Our screens identified several novel regulators of *HOX/MEIS* expression in AML and many of them were important for sustaining the expression of an array of leukemia oncogenes that drive AML including *BMI1, SATB1* and *MYC*. Most notably, our studies identified a novel role for the Tudor domain-containing chromatin reader protein SGF29 in AML. Our studies show that SGF29, through its Tudor-domain- is a critical epigenetic regulator required for maintaining transcription of key leukemia oncogenes and leukemogenesis of diverse AML subtypes.

## RESULTS

### A high-density, domain-focused CRISPR screen identifies epigenetic regulators of *MEIS1* expression

AML with the t(10;11)(p13;q14) translocation - which results in the *CALM-AF10* fusion gene - display highly elevated expression of several stem cell-associated factors including the *HOXA* cluster genes, *MEIS1* and *BMI1* ^15, 16^. Indeed, AML cell lines with CALM-AF10 fusions display some of the highest expression of these oncogenes (Supplementary Fig. S1A). We used a *CALM-AF10* positive AML cell line – U937 - where the enhanced green fluorescence protein (eGFP) is knocked into the endogenous *MEIS1* locus immediately upstream of the start codon^14^. Leveraging this system, we sought to comprehensively identify epigenetic regulators of *MEIS1* expression as a surrogate for identifying *HOX/MEIS* regulators. First, we validated these U937-eGFP-MEIS1 cells (henceforth termed UB3) by perturbation with sgRNAs targeting *DOT1L* (Fig. 1A). Using these cells, we conducted a chemical compound screen using high-throughput flow-cytometry in a 384 well format assaying for eGFP levels (see Supplementary Fig. S1C for schematic). Screening with a library of 265 small-molecule compounds targeting epigenetic regulators (Supplementary Table S1) revealed inhibitors of DOT1L and ENL, targets already established to regulate *HOX/MEIS* expression in AML ^17–19^ as “hits” - defined by >50% drop in eGFP-MEIS1 expression (Fig. 1B). Highly similar results were obtained in mouse MLL-AF9 transformed myeloid cells that we established from the bone marrow (BM) of a recently reported eGFP-MEIS1 transgenic mouse (Supplementary Fig. S1B). Since no novel MEIS1 regulators were identified in our small-molecule screens, we reasoned that this was likely due to limitations of the small-molecule library composition. Most families of poorly studied epigenetic readers, writers and eraser proteins are not represented in currently available chemical epigenetic inhibitor libraries, precluding the identification of their activities in phenotypic screens. Therefore, we sought to extend our investigation of *MEIS1* epigenetic regulators using a genetic screening approach. Towards this end, we developed a high-density CRISPR library targeting 645 epigenetic regulators including chromatin readers, writers of histone acetylation, methylation and other modifications, genes involved in chromatin remodeling, transcription elongation, and DNA & RNA modifications in addition to control genes and non-targeting sgRNA controls (Fig. 1C and Supplementary Table S2). Studies have shown that targeting exons encoding functional domains may generate a higher proportion of null mutations and result in greater phenotypic selection compared to sgRNAs targeting early constitutive exons^20^. Thus, we extracted the coding sequence corresponding to annotated protein domains available in Uniprot, for all our epigenetic CRISPR library genes (see Supplementary Methods). Then, we designed sgRNAs for the domain-encoding sequences on the Broad Institute Genetic Perturbation (GPP) Platform CRISPRko tool^21^. We incorporated the 5 top-scoring sgRNAs of each annotated domain of every gene (see Methods). In addition to domain-focused sgRNAs, we designed 5 sgRNAs based on the standard early exon-targeting criterion. We used the epigenetics CRISPR library to conduct an *in vitro* phenotypic enrichment screen in single-cloned lentiviral Cas9-expressing UB3 cells. We hypothesized that enrichments in low- or high- eGFP fractions would help identify MEIS1 positive or negative regulators, respectively. Thus, we sorted the top 25% and bottom 25% fractions based on eGFP levels using flow cytometry (FACS, see Fig. 1D for schematic). Normalized read counts of non-targeting controls and polymerase-encoding genes were uniformly distributed in both high- and low- eGFP fractions, indicating successful technical performance of controls (Supplementary Fig. S2A). Our analyses of the screen with the MAGeCKFlute pipeline (22,23) revealed several novel regulators, including AFF2, CSNK2A1, CSNK2B, SGF29/CCDC101, ENY2, TAF6, HDAC1, and LDB1, as well as most of the previously described epigenetic regulators of *MEIS1* or *HOX* gene expression in AML, such as DOT1L (24–26), MLLT1/ENL (18,19), MLLT10/AF10 (24,27), KAT7/HBO1(28,29), JADE3 (28) and KMT2A/MLL1 (Fig. 1E). Analysis of the top candidate hits using the STRING database (17) showed that multiple proteins from six chromatin complexes were identified (Fig. 1F). This reinforced the possibility that we identified distinct constituents of chromatin-modifying complexes with MEIS1 regulatory activity – offering independent, complementary nodes for therapeutic targeting.

**Figure 1.**
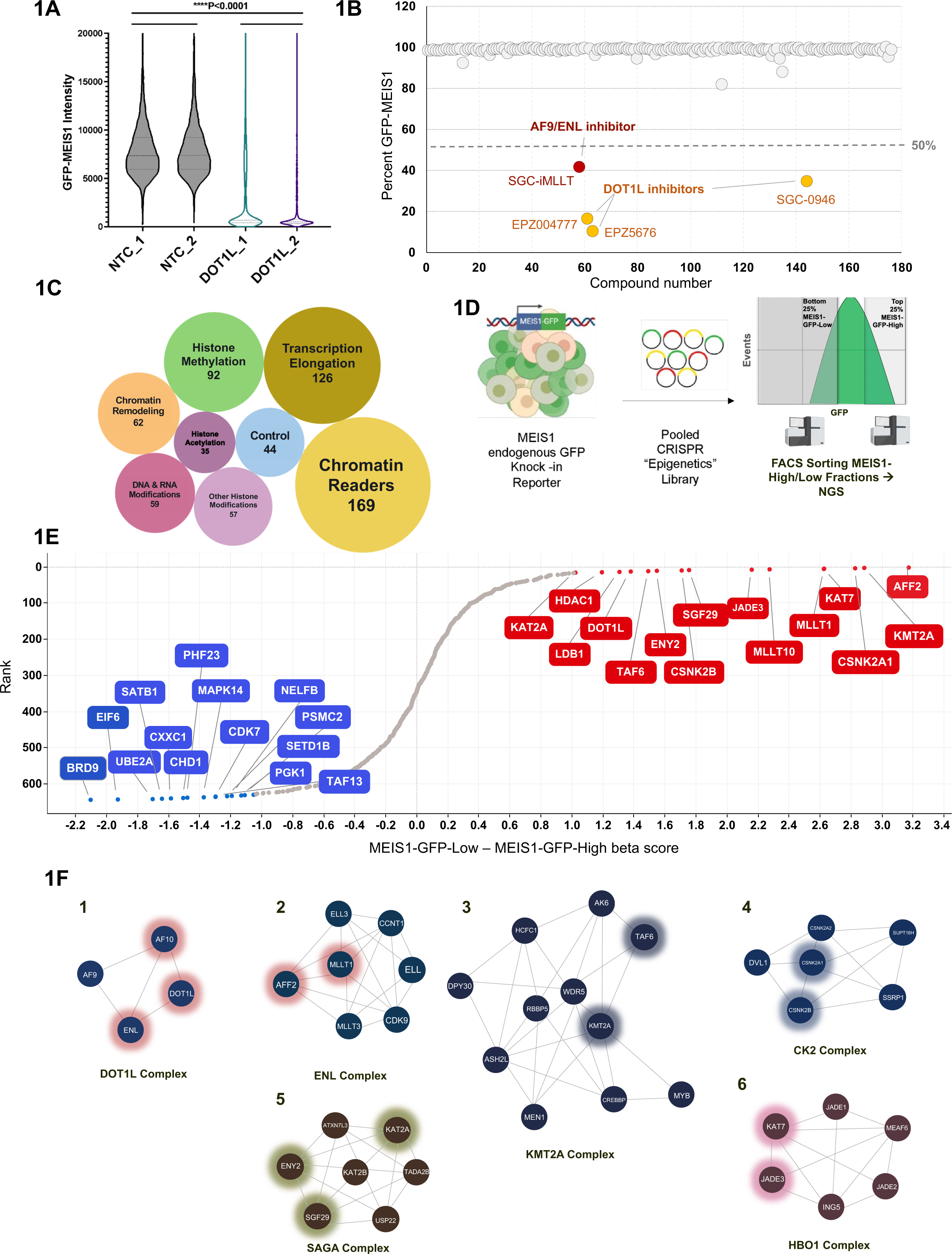
Screening for epigenetic modulators identifies novel MEIS1 regulators in AML cells. **(A)** GFP-MEIS1 signal in U937 GFP-MEIS1 AML upon DOT1L CRISPR knockout is plotted on the y-axis. Biological duplicates indicate independent sgRNAs. The y-axis indicates eGFP intensity values from single cells (n ≥ 5,000). (B) Small-molecule screen in human U937 GFP-MEIS1 AML cells; y-axis indicates the percentage GFP-Meis1 intensity compared to controls on day 5 of treatment with the small molecule library; x-axis indicates a compound number assigned to list the compounds in the library. (C) Bubble plot to illustrate the categories of chromatin modulators included in the library design; the numbers indicate the different genes under the category and the size of the bubble is proportional to the gene-set size. Controls refer to pan-essential genes. (D) Strategy for phenotypic pooled CRISPR screening of epigenetic regulators for MEIS1 expression in the U937 cell line with an eGFP-MEIS1 endogenous knock-in tag. (E) Sorted gene hits based on differential beta scores for the eGFP-MEIS1 low minus eGFP- MEIS1 high fractions. Beta scores were calculated using MAGeCKFlute. (F) Annotated protein complexes comprising top candidate hits identified in the epigenetic screen as identified using String database analysis are shown. Hits from the epigenetics CRISPR library screen are marked with a glowing hue within the protein complex.

### Competition assays reveal the role of MEIS1 regulators in AML cell growth

Next, we sought to test the requirement of our top candidate *MEIS1* regulators for AML cell growth. For this, we conducted arrayed CRISPR competition assays, first in UB3 cells (see Fig. 2A for schematic). Specifically, we cloned 2-3 sgRNAs for each of the top candidate hits (*SGF29, ENY2, AFF2, CSNK2A1, CSNK2B, MLLT1, KAT7,* and *JADE3),* in addition to *DOT1L* as a positive control, and non-targeting controls (NTC). For this, we used a plasmid vector co-expressing a blue fluorescence protein (BFP) and lentivirally transduced U937 cells at a ∼50% transduction rate. We then sampled the populations by flow cytometry for up to 14 days and observed a significant (P<0.05, n =3) and progressive decline in the percentage of BFP-positive cells targeted by all the candidate hits compared to NTCs over time (Fig. 2B). In addition, we observed a significant drop in eGFP fluorescence (P<0.05, n=3), for multiple sgRNAs targeting candidate hits, confirming a strong reduction in MEIS1 expression (Supplementary Fig. S2B). In sum, these results indicate our phenotypic screening strategy uncovered high-confidence *MEIS1* regulators that affect the growth of U937 cells. In order to rule out that this anti-proliferative effect was limited to the CALM-AF10 positive cell line U937, we tested some of the novel MEIS1 regulators, including *CSNK2A1*, *ENY2,* and *SGF29* and controls in a Cas9-expressing MLL-AF9 fusion-positive MOLM13 cell line (Supplementary Fig. S2C). CRISPR deletion of these genes also led to a progressive decline in the percentage of BFP-positive cells compared to NTC-transduced cells, indicating that growth of MOLM13 cells also depended on these genes (Supplementary Fig. S2C). Interestingly, among these genes, the deletion of SGF29 and ENY2 had the most detrimental effects on the growth of both U937 as well as MOLM13-cells (Fig. 2B and Supplementary Fig. S2C).

**Figure 2.**
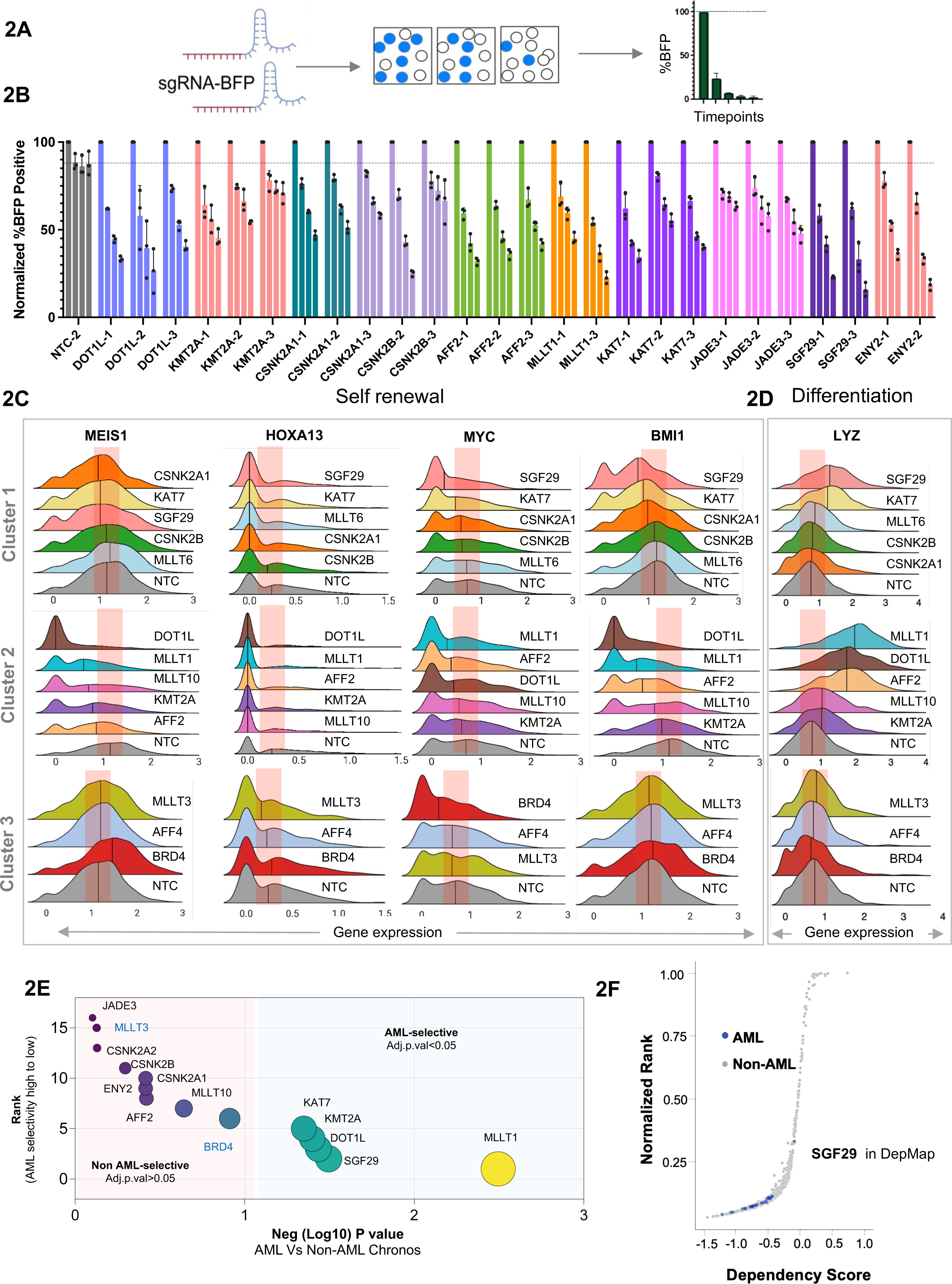
Requirement for MEIS1 regulators for transcriptomic changes and AML cell growth: **(A)** Schematic of competition assay with cells transduced with sgRNAs expressed in a BFP+ backbone. The proportion of BFP-positive cells is assessed over time using flow cytometry**. (B)** The y-axis indicates flow cytometry measurements of the percentage of U937 BFP+ cells over time, normalized to the baseline (time 0 or T”0”) measurement (n=3 technical replicates); the x-axis indicates the sgRNA number for non-targeting controls (NTC) or each gene (n=2). Each bar represents a timepoint measurement: baseline, day 8, day 11, and day 14. **(C)** Ridge plots for CROP-Seq perturbations. Knockout genes are indicated on the y-axis and RNA expression levels for the **(A)** self- renewal-associated or **(B)** differentiation-associated genes listed are on the x-axis. The median expression value is indicated. **(E)** A relative score of DepMap dependencies for AML (n=26) compared to non-AML cell lines (n=1080) is shown for candidate hits identified in our screen. Y-axis is the rank of AML selectivity, and the x-axis shows the negative log 10 p-values. (F) A sigmoid plot of DepMap data showing the dependency score (Chronos – x-axis), compared to the normalized dependency rank (y-axis) for SFG29 in 1,086 cancer cell lines.

### CRISPR droplet-sequencing reveals candidate hits regulate leukemia oncotranscriptome

Since we used MEIS1 as a surrogate reporter for the screen, we wanted to further evaluate the following: a) whether our candidate hits could regulate the expression not only of *MEIS1*, but also of other AML-associated oncogenes including the HOXA cluster genes, and b) which epigenetic regulators from our hits had the most pronounced effects on sustaining transcription of AML-promoting oncogenes. For this, we generated a targeted, small pooled CRISPR library containing 2-3 validated sgRNAs per candidate gene together with NTCs (a total of 29 sgRNAs) and performed a single cell RNA-sequencing experiment in the UB3 cell line. In our pool, we also included sgRNAs for control genes which are either chromatin readers that do not show HOX/MEIS-specific gene regulatory activity in U937 cells (*MLLT3*, *MLLT6*), or have non-selective transcriptional effects (*BRD4*). With these sgRNAs, we used CRISPR droplet sequencing (CROP-Seq) ^22^, which allows for matching the single-cell RNA-sequencing profile of each cell to the sgRNA expressed within it, and thus infer knockout signatures of each of the CRISPR-deleted genes. After sgRNA assignment, we performed unbiased clustering of whole transcriptome single-cell RNA-seq data using the Ward.D2 minimum variance method (see Supplemental Methods) revealing the relatedness of each of the perturbations (Supplementary Figs. S2D and S2E). While the transcriptomes of *CSNK2A1*-, *KMT2A*- and *MLLT10*- knockout cells clustered together, *DOT1L*-, *MLLT1*-, *AFF2*- and *SGF29*- deleted transcriptomes formed a distinct cluster, indicating transcriptional similarities. We then also performed clustering on the basis of *HOX/MEIS* expression to investigate transcriptional relatedness (Supplementary Fig. S2F, see Supplemental Methods). Our analysis identified 3 major clusters. One cluster comprises *DOT1L*, *MLLT1*, *MLLT10*, and *AFF2*. This possibly reflects the overlapping activities or proteins that have been described to interact in multi-protein complexes ^23–25^. Interestingly, in our studies, *AFF2*, but not the super-elongation (SEC) complex constituent *AFF4* was identified as a regulator of *HOX/MEIS* expression in U937 cells (Fig. 1E, Fig. S2F, and Supplementary Table S3).

Similarly, the deletion of *MLLT1*, but not its homolog *MLLT3*, modulated *HOX/MEIS* expression in the UB3 cells, consistent with a prior study in a KMT2A-rearranged AML (8). Another cluster consisted of the genes *KAT7*, *CSNK2A1*, *CSNK2B*, and *SGF29*, whose deletion also had potent effects on the downregulation of *MEIS1* and other leukemia oncogenes, such as *HOXA* cluster, *BMI1, SATB1*, *RNF220,* and *MSI2* (Fig. 2C, Fig. S2F, and Supplementary Table S3). Moreover, their deletion also led to a concomitant increase in the expression of myeloid differentiation-associated genes such as lysozyme (*LYZ*), *ELANE*, the cathepsins *CTSA* and *CTSD*, and *S100A8*/*A9* (Fig. 2D and Supplementary Fig. S2H). Cluster 3 consisted of the control genes we included in our CROP-Seq study, namely *AFF4* and the chromatin readers *BRD4* and *MLLT3*, whose deletion had no significant effects on *HOX/MEIS* or on differentiation-associated signatures in the setting of the UB3 cell line. Taken together, our studies showed that several epigenetic regulators identified in our eGFP-MEIS1 screen were important for sustaining key LSC signature genes and limiting the gene expression characteristic of a differentiation block in U937 cells.

### CRISPR assays earmark SGF29 as an AML-selective dependency

One of the challenges in developing epigenetic candidates as therapeutic targets is their potential for non-selective toxicity. One way of testing the selectivity of a candidate gene is to investigate the effect of its genetic deletion in the cancer dependency maps (DepMap) dataset, a database of genome-scale functional genetic screens across cancer cell lines ^26^. We evaluated the efficacy (the degree to which CRISPR knockout of the candidate gene reduces cell fitness in sensitive lines), and selectivity (the degree of differences in its essentiality across all cancer cell lines) of our top candidates using CRISPR and shRNA data available in Shiny DepMap ^27^. Our analyses showed that depletion of some of our hits, including SGF29, ENY2 and CSNK2A1, as well as previously characterized genes such as DOT1L, MLLT1, MLLT10 and KAT7 had a higher selectivity compared to pan-essential genes, such as PSMA4 or the broader transcriptional regulator BRD4 (Fig. S2I), indicating that these targets may have a better therapeutic index. Next, we assessed the lineage specificity of depleting our candidates in AML, by comparing the differences in median CRISPR knockout fitness scores (Chronos) between 26 AML cell lines and 1,060 non-AML cancer cell lines. This analysis showed that while depletion of some of the hits was highly AML-selective, that of others was not. Specifically, the most AML-selective genes were *MLLT1*, *SGF29*, *DOT1L*, *AFF2*, and *KMT2A* (ranked in decreasing order of AML selectivity as assessed by a Wilcox test of AML vs non-AML Chronos scores, Fig. 2F). Other chromatin regulators, namely *MLLT10*, *CSNK2A1*, *CSNK2B*, *JADE3*, and *ENY2* were not AML selective (with an FDR p-value cutoff <0.05). Of particular interest to us was the chromatin reader SGF29, in terms of its strong effect on leukemogenic gene expression programs, high AML selectivity across 1,086 cancer cell types (Fig. 2E and 2F) and hitherto uncharacterized role in leukemia.

### SGF29 deletion has marked anti-leukemia effects in cell line- and patient-derived AML models

Since our transcriptomic, arrayed CRISPR validation and computational studies showed that SGF29 was one of the strongest dependencies from our hits across AML cell lines, we sought to directly assess the impact of its deletion on human AML cell growth and leukemogenesis. Consistent with our flow cytometry-based *in vitro* competition assays (Fig. 2B and Supplementary Fig. S2C), we observed that SGF29 inactivation using validated CRISPR sgRNAs (Supplementary Fig. S3A) led to significantly increased retention of the Cell Trace Violet dye (see Methods) in both U937, as well as MOLM13 AML cell lines, indicating diminished proliferation compared to cells treated with NTCs (Figs. 3A and 3B). Furthermore, the antiproliferative effects of SGF29 deletion were accompanied by a significant increase in the proportion of cells in the G0/G1 fraction compared to SGF29 wild-type cells (Supplementary Fig. S3B). Importantly, SGF29- deleted cells showed increased uptake of fluorescence-labeled heat-inactivated *E. coli* bioparticles, indicative of functional myeloid differentiation of U937 cells (Fig. 3C and 3D). Next, we performed bulk RNA sequencing (RNA-seq) of MOLM13 and U937 cells with SGF29 CRISPR knockout compared to NTC cells. Our RNA-seq analysis showed that SGF29 deletion significantly decreased the expression of several *HOXA* cluster genes in addition to *MEIS1* in both cell lines (Figs. 3E & 3G), confirming its role in *HOX/MEIS* regulation in diverse AML subtypes. Further, there was a significant reduction in the expression of other oncogenes such as *MYC* (Fig. S3C). The expression of MYC target genes was highly enriched in NTC cells, compared to the SGF29-deleted counterparts, as assessed using gene-set enrichment analysis (GSEA^28^, Fig. S3D). We then tested the effects of SGF29 deletion on *in vivo* leukemogenesis using cell line-derived (CDX) xenograft models in NOD.Cg-*Rag1^tm1Mom^ Il2rg^tm1Wjl^*/SzJ (NRG) mice. In these experiments, we observed a significant increase in disease latency upon SGF29 knockout in Cas9-expressing U937 and MOLM13 cell lines (Figs. 3F & 3H).

**Figure 3.**
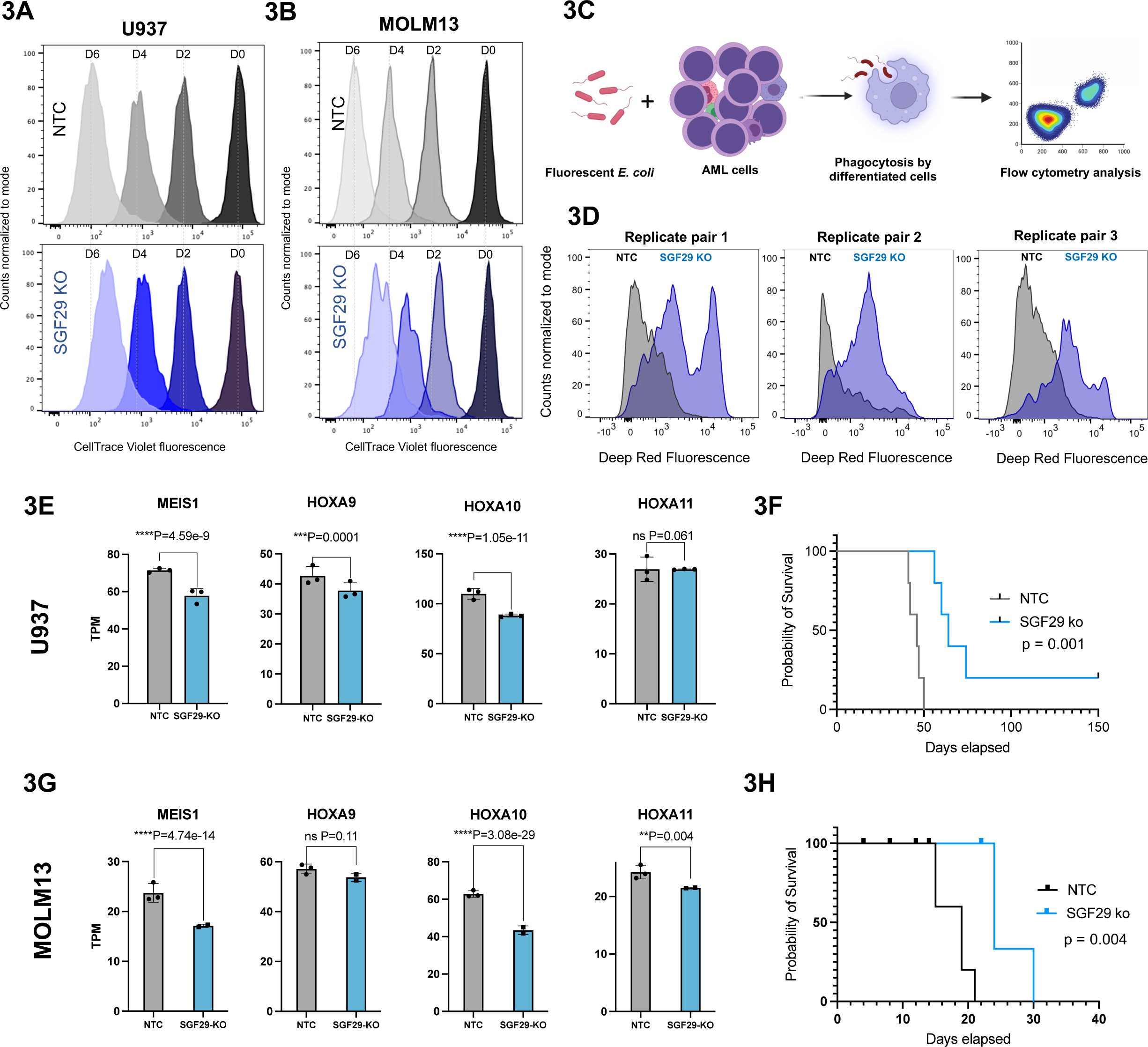
Proliferative and transcriptional effects of SGF29 loss in AML. Retention of the CellTrace™ Far Red dye in **(A)** U937 or **(B)** MOLM13 cells transduced with an SGF229 sgRNA (top) compared to non-targeting control (NTC), measured by flow cytometry over time. **(C)** Schematic for differentiation assessment via phagocytosis of fluorescently labeled E. coli bioparticles. **(D)** Histograms of U937 cells transduced with an NTC (grey) or an SGF29 sgRNA (violet) show the fluorescence intensity of engulfed pHrodo™ Red E. coli bioparticles measured at 9 days after staining. Effect of SGF29 deletion on the transcription of HOXA/MEIS genes in **(E)** U937 or **(G)** MOLM13 AML cell lines, compared to non-targeting control (NTC) is shown. TPM: transcripts per million, P val. *<0.05, **<0.01, ***<0.005. (F) Kaplan-Meier curves for NTC versus SGF29 knockout in **(F)** U937 and **(H)** MOLM13 AML cell lines and an MLL-AF10 patient-derived xenograft line.

### Genetic Sgf29 inactivation impairs the clonogenicity of transformed but not normal hematopoietic progenitors

Next, given the role of SGF29 in the transcription of self-renewal-associated genes, we wanted to test the effect of Sgf29 inactivation on the clonogenic capability of myeloid bone marrow (BM) cells transformed by distinct AML driver oncoproteins. We therefore performed methylcellulose-based colony forming unit (CFU) assays on murine bone marrow hematopoietic stem and progenitor cells (HSPCs) transformed using the CALM- AF10, MLL-AF10 or MLL-AF9 fusions. We observed a significant reduction in the number of colonies with a blast-like morphology in CALM-AF10 transformed cells upon *Sgf29* deletion using exon-targeting sgRNAs (Fig. S4A), compared to *Sgf29* intron-targeting controls (Figs. 4A-C). Interestingly, while there was a dramatic decrease in the number of immature blast-like colonies, there were modest effects on the number of differentiated colonies. Thus, most colonies in the Sgf29^-/-^ arm bore a differentiated morphology (Fig. 4B). The strong reduction in blast colony formation persisted for at least two rounds of replating (Supplementary Fig. S4B). At the cellular level, in contrast to the immature, myeloid blast-like cellular morphology observed in Wright-Giemsa-stained cytospins of cells with wild-type *Sgf29*, the *Sgf29* knockout cells resembled terminally differentiated, or differentiating myeloid lineage cells (Fig. 4C). Similar results were obtained from BM cells transformed using MLL-AF9 (Fig. 4D-F) and MLL-AF10 (Fig. 4G-I) fusion oncoproteins. *Sgf29* deleted cells also showed increased apoptosis compared to wild-type cells as measured by immunoblotting for cleaved PARP (Supplementary Fig. S4E). Next, tested the effect of *Sgf29* deletion on the colony-forming ability of normal HSPCs. In contrast to BM cells transformed using AML oncoproteins, CRISPR-mediated Sgf29 deletion in murine lineage negative, Sca-1 positive, Kit-positive (LSK) cells did not discernably alter either the number or type of colonies observed in CFU assays (Fig. 4J- K). Taken together, these data indicate that *Sgf29* inactivation influences the clonogenicity and differentiation of cells transformed by diverse AML oncoproteins, but not normal hematopoietic stem and progenitor cells (HSPCs).

**Figure 4.**
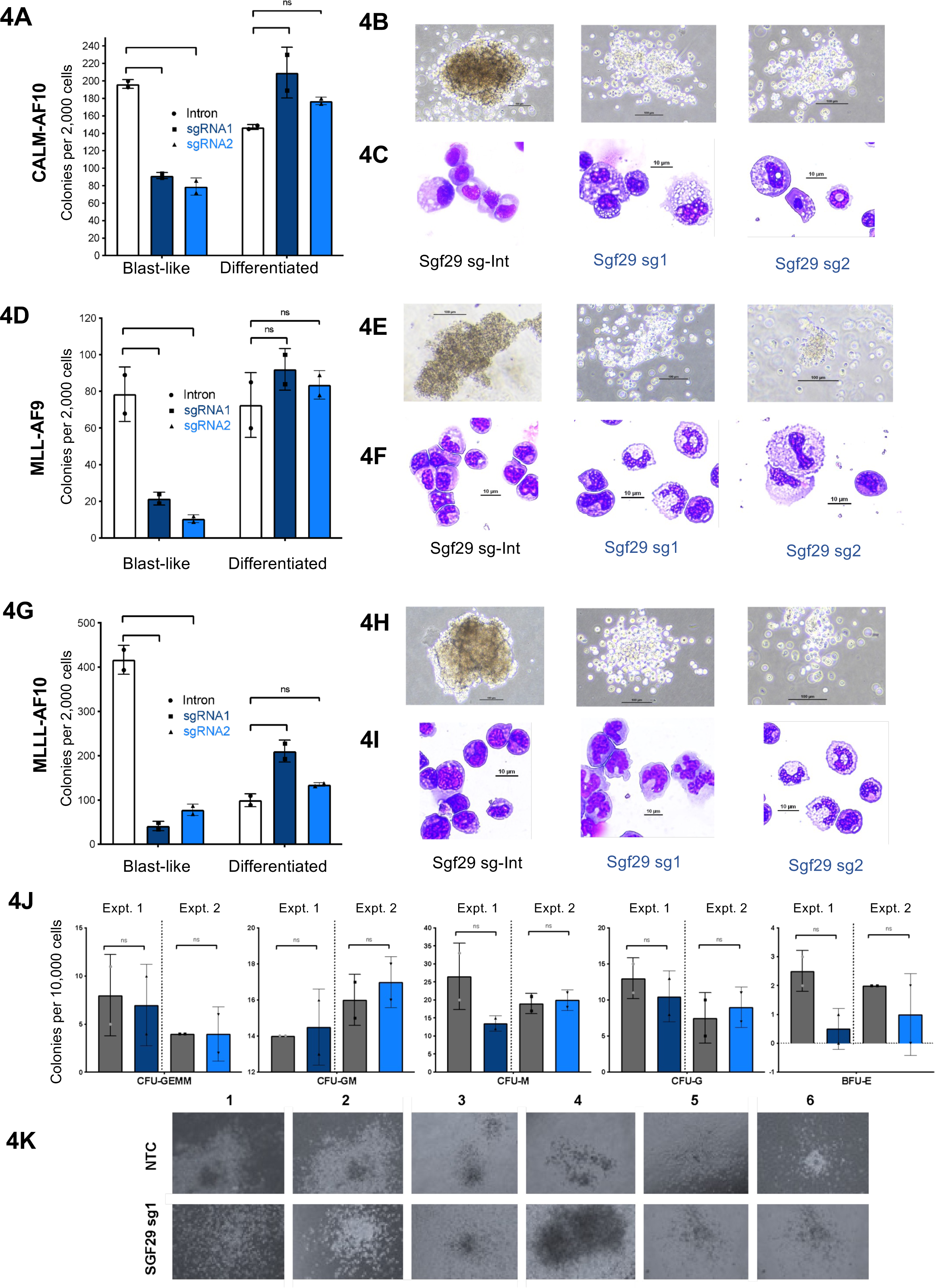
*Sgf29* deletion impairs the clonogenicity of transformed but not normal hematopoietic cells: (A) Number of colony-forming units (CFU) from CALM-AF10 transformed cells transduced with Sgf29 intron-targeting sgRNA (Sgf29-Int) or two independent Sgf29 exon-targeting sgRNAs are shown. CFUs per 2,000 plated cells at 1 week is plotted on the Y-axis, and colonies are divided into those with a blast-like or differentiated colony morphology. **(B)** Pictures of representative colonies with each of the labeled sgRNA-transduced cells are shown. Scale bar: 100 mm. **(C)** Wright-Giemsa- stained cytospins of representative cells from each of the CRISPR perturbations are shown. Scale bar: 10 mm. The same experiments as shown for CALM-AF10 transformed BM cells shown in (**A-C)** are shown for MLL-AF9-transformed BM cells in **(D-F)** and MLL- AF10-transformed cells in **(G-I). (J)** CFUs per 10,000 lineage negative, Sca-1 positive, Kit positive (LSK) cells with Sgf29 deletion (Sgf_sg1) compared to Sgf29 wildtype LSKs (non-targeting control; NTC) are shown. Y-axis shows the numbers of different types of colonies, with each plot indicating a separate kind of morphologically distinct CFU in this assay. P values: *= p<0.05, ** = P<0.01.

### The Tudor domain of SGF29 is essential for its role in myeloid transformation

The SGF29 protein harbors a carboxy (C)-terminal tandem Tudor domain, which is important for the recognition of H3K4 di/trimethylated chromatin modifications and recruitment of KAT2A (GCN5)-containing chromatin-modifying complexes^29–31^. In our epigenetic CRISPR library screen, we identified that top-performing sgRNAs for SGF29 mapped to its Tudor domain. Thus, we hypothesized that this domain may be required for the transcriptional activation of AML oncogenes including the *HOX/MEIS* genes, *BMI1* and *MYC*. To assess the dependency of the Tudor domain for SGF29 localization in the HOX/MEIS loci, we cloned a Flag-tagged SGF29 gene and generated a stably transduced U937 cell line. Similarly, in order to test the effect of the SGF29 Tudor domain we also generated U937 cells expressing the SGF29^D196R^ mutant, which has been shown to disrupt H3K4me3 binding using *in vitro* peptide binding assays ^29^. ChIP-sequencing (ChIP-seq) using a Flag antibody in the Flag-SGF29 U937 cells showed that the protein-occupied regions that were enriched for H3K4me3 marked active promoters and H3K27 acetylated enhancers (Figs. 5A&B). SGF29 occupied the promoter and/or enhancer regions of HOX/MEIS genes as well as other AML oncogenes whose expression was dependent on SGF29 (including *BMI1* and *MYC,* Fig. 5C). Interestingly, in contrast to the wildtype SGF29, binding of the SGF29^D196R^ mutant to these gene loci was almost obliterated (Fig. 5C). Next, we tested whether the Tudor domain is important for the clonogenicity of leukemia cells using the CFU assay. Using the MLL-AF9 murine model in which we observe a highly significant decrease in the formation of immature blast-like colonies upon endogenous Sgf29 deletion, we retrovirally overexpressed wildtype human SGF29 (impervious to the mouse anti-Sgf29 sgRNAs) or the SGF29^D196R^ mutant counterpart. CFU assays demonstrated that ectopic overexpression of wildtype, but not mutant SGF29, could completely rescue the loss of blast-like CFUs upon endogenous *Sgf29* deletion (Fig. 5D). Notably, ectopic expression of the SGF29^D196R^ mutant itself dramatically reduced the number of blast-like colonies even in MLL-AF9 cells with intact endogenous Sgf29 alleles, indicating that the SGF29^D196R^ mutant has dominant negative activity (Fig. 5D).

**Figure 5.**
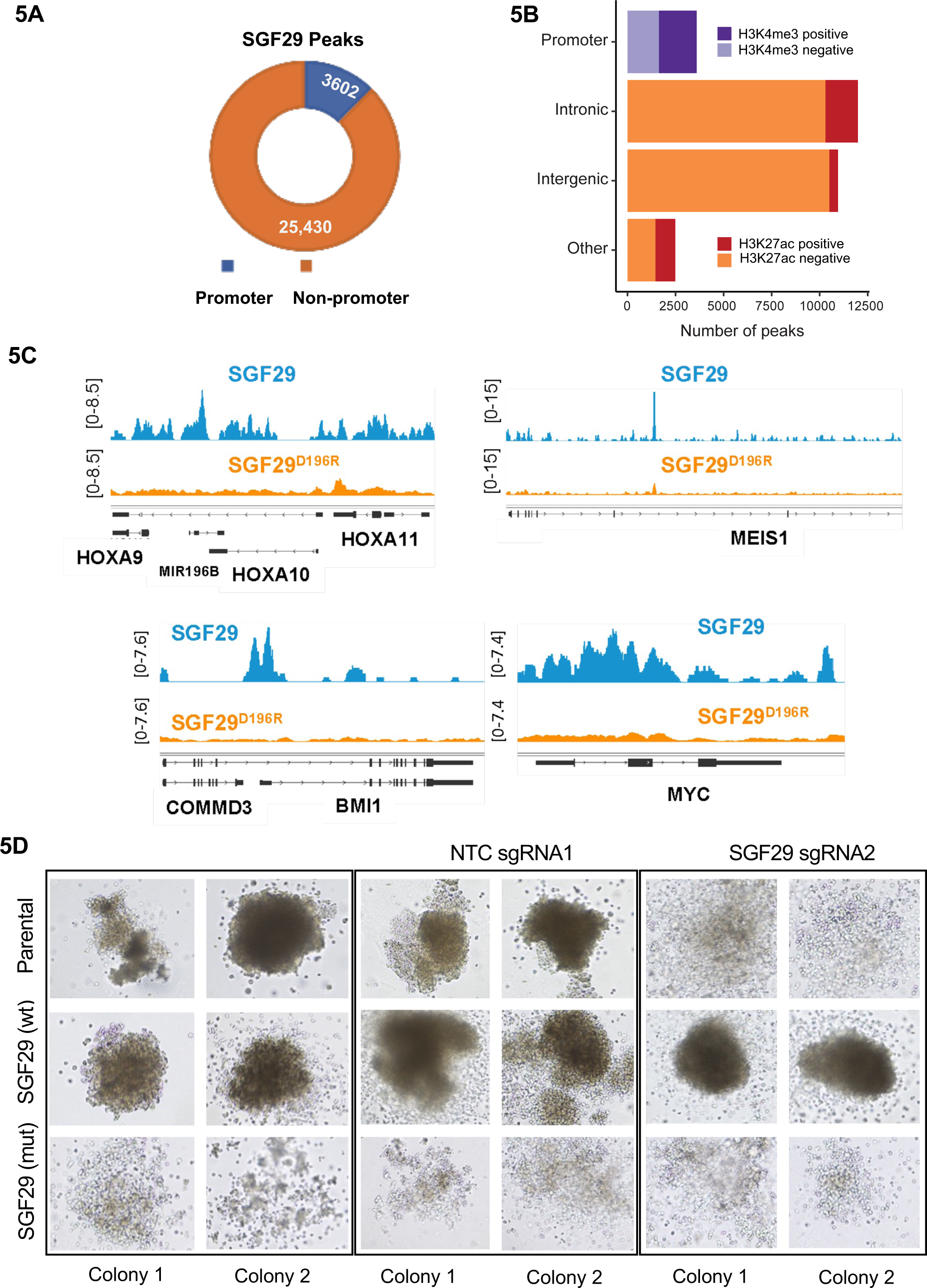
Evaluation of the Tudor domain of SGF29 activity in AML cells. (A) The proportion of SGF29 peaks in promoter regions and intergenic regions in U937 cells is shown in the Donut plot. **(B)** SGF29 peaks in promoter regions and their association with H3K4me3, and intergenic, intronic, and other regions marked by H3K27ac in U937 cells are shown in the bar graph as marked. X-axis shows number of peaks. **(C)** Genome tracks depicting normalized reads (y-axis) for wildtype or mutant SGF29 at the genomic locus of the HOXA/MEIS1, MYC, and BMI1 genes in U937 cells are shown. **(D)** Top panel: Representative images of CFUs from MLL-AF9 transformed bone marrow cells with a mock transduction (two top left colonies), non-targeting control (two top middle colonies), or Sgf29 sgRNA (two right colonies) are shown. Middle panel: Same cells as above, but with ectopic retroviral overexpression of wild-type human SGF29 impervious to SGF29 sgRNA. Lower panel: Same MLL-AF9 transformed cells as in top panel but with retroviral overexpression of the human SGF29D196R mutant is shown. Representative colonies are shown in bright field at 20x magnification.

### SGF29 regulates chromatin localization of proteins with established roles in AML pathogenesis

We sought to identify the molecular mechanism by which SGF29 elicits its effects on the transcription of leukemia oncogenes. SGF29 participates in distinct chromatin regulatory complexes, including the ADA2A-containing (ATAC) complex and the SPT3-TAF9- GCN5-acetyltransferase complex (STAGA) complex, the mammalian homolog of the yeast SAGA (SPT-ADA-GCN5-acetyltransferase complex). Both of these complexes harbor the KAT2A (GCN5) and KAT2B (PCAF) acetyltransferases with histone acetylating activity and prominent roles in transcriptional activation^32^. Thus, we sought to investigate which epigenetic regulators are evicted from chromatin upon SGF29 depletion in AML cells. For this, we performed chromatin enrichment proteomics (ChEP)^33^ (see Fig. 6A for schematic). This recently developed method allows an unbiased quantitative and qualitative assessment of the chromatin-associated proteome^33^. Our ChEP purification yielded substantial enrichment of the chromatin fraction as measured by the high proportional abundance of histones, histone modifying proteins, transcription factors, and other chromatin and nuclear-associated proteins in the enriched fractions (Fig. 6B and Supplementary Fig. S5A). *SGF29* deletion in U937 cells significantly decreased the abundance of key AML oncoproteins, as measured by the intensity of chromatin-associated peptides by mass spectrometry. Specifically, the levels of HOXA13 and SATB1 were significantly reduced in the chromatin fraction of SGF29-deleted cells in contrast to their wild-type counterparts. Of note, the levels of MEIS1 and MYC were undetectable in the chromatin fraction of SGF29 deleted cells in contrast to their SGF29 wild-type counterparts, where these proteins were highly abundant (Figs. 6C-D and Supplementary Fig. S5B). We also observed a significant decrease in the abundance of CDK4 and CDK6 proteins, involved in cell cycle regulation and known to play prominent roles in the proliferation of cancer cells, particularly in AML^34^. Most importantly, our ChEP analysis demonstrated that SGF29 deletion led to a significant decrease in the chromatin abundance of key components of the SAGA complex, the histone acetyltransferase KAT2A, as well as the transcriptional adaptor protein TADA3 (Fig. 6C and Supplementary Fig. S5B). These results indicate that SGF29 deletion may lead to the eviction of the SAGA complex from the chromatin fraction of U937 cells. We confirmed our results by immunoblotting using a KAT2A-specific antibody in different cellular fractions in the SGF29 wildtype or CRISPR-deleted AML cells. Our studies demonstrated that while KAT2A could be found mostly in the chromatin fraction in wildtype U937 cells, SGF29 deletion indeed led to an increased abundance of KAT2A in the cytoplasmic fraction (Fig. 6E).

**Figure 6.**
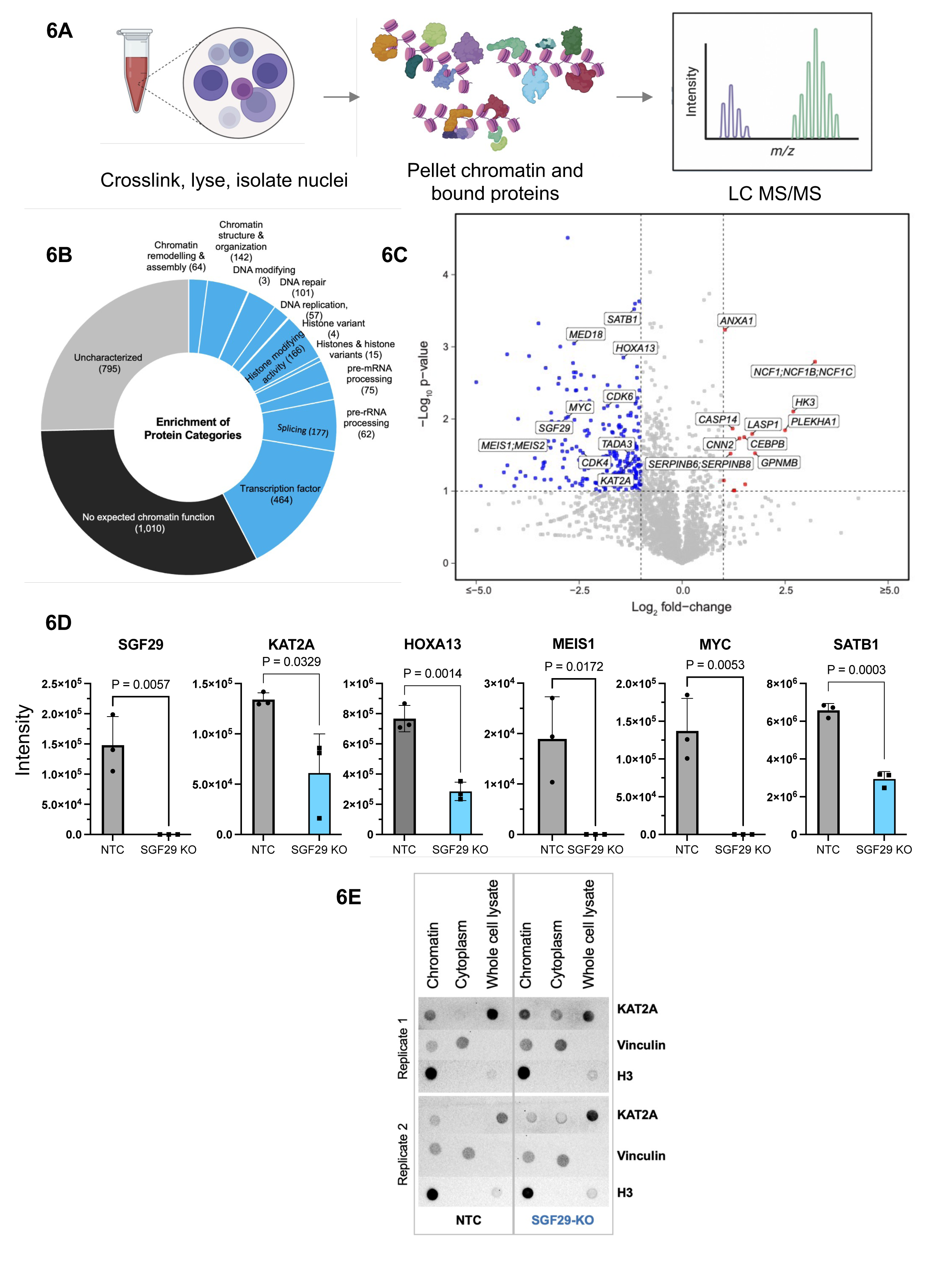
Chromatin enrichment proteomics reveals SGF29-dependent proteins in the chromatin proteome (A) Schematic for the ChEP sample preparation, as previously described ^33^ . **(B)** A donut plot showing the functional protein categories in proteomic analysis of the ChEP SGF29-deleted fraction across 4 samples. Proteins were annotated using the categories outlined in ^33^. The number of proteins per category is shown in parentheses. **(C)**Volcano plots depicting -log(10) of the adjusted P value on the y-axis and log(2) of fold change (LogFC) of all proteins in the ChEP fractions for both SGF29 knockout vs. NTC. Differentially abundant protein (Absolute FC >2, Adj. P value. < 0.05 are not shown; n=3 for every experimental condition. **(D)** Bar graphs depicting intensity values for self-renewal-associated proteins enriched by ChEP in the SGF29 knockout vs NTC experiment. **(E)** Dot blots for ChEP-enriched chromatin, cytoplasmic fraction, and whole-cell lysate for NTC and SGF29 knockout are shown. Vinculin and Histone H3 were included as cytoplasmic and chromatin protein controls respectively.

### SGF29 deletion diminishes H3K9Ac on the promoters of key leukemia oncogenes

Our results showing that SGF29 deletion decreased the abundance of the histone acetyltransferase KAT2A in the chromatin fraction led us to hypothesize that SGF29 recruits the chromatin activity of KAT2A on AML oncogene loci. KAT2A is the acetyltransferase subunit of the SAGA complex that plays important roles in leukemia and other cancers ^35^. We performed ChIP-seq for H3K9Ac, the major histone modification deposited by the SAGA complex, which is associated with transcriptional activation ^32^. Our analysis of the ChIP-seq data showed that upon SGF29 deletion, there were significant changes in H3K9Ac, with 10,834 peaks showing reduced acetylation and 3,119 peaks showing increased acetylation in SGF29 knockout compared to wildtype U937 cells (Fig. 7A, and Supplementary Fig. S6A). Importantly, there was a pronounced decrease in acetylation levels at the promoter of several AML oncogenes that were also transcriptionally downregulated in the SGF29 knockout RNA-seq data. This included *HOXA* cluster genes, *MYC*, *BMI1*, and *SATB1* – all bona fide AML oncogenes with established roles in augmenting leukemic self-renewal (Fig. 7B).

**Figure 7.**
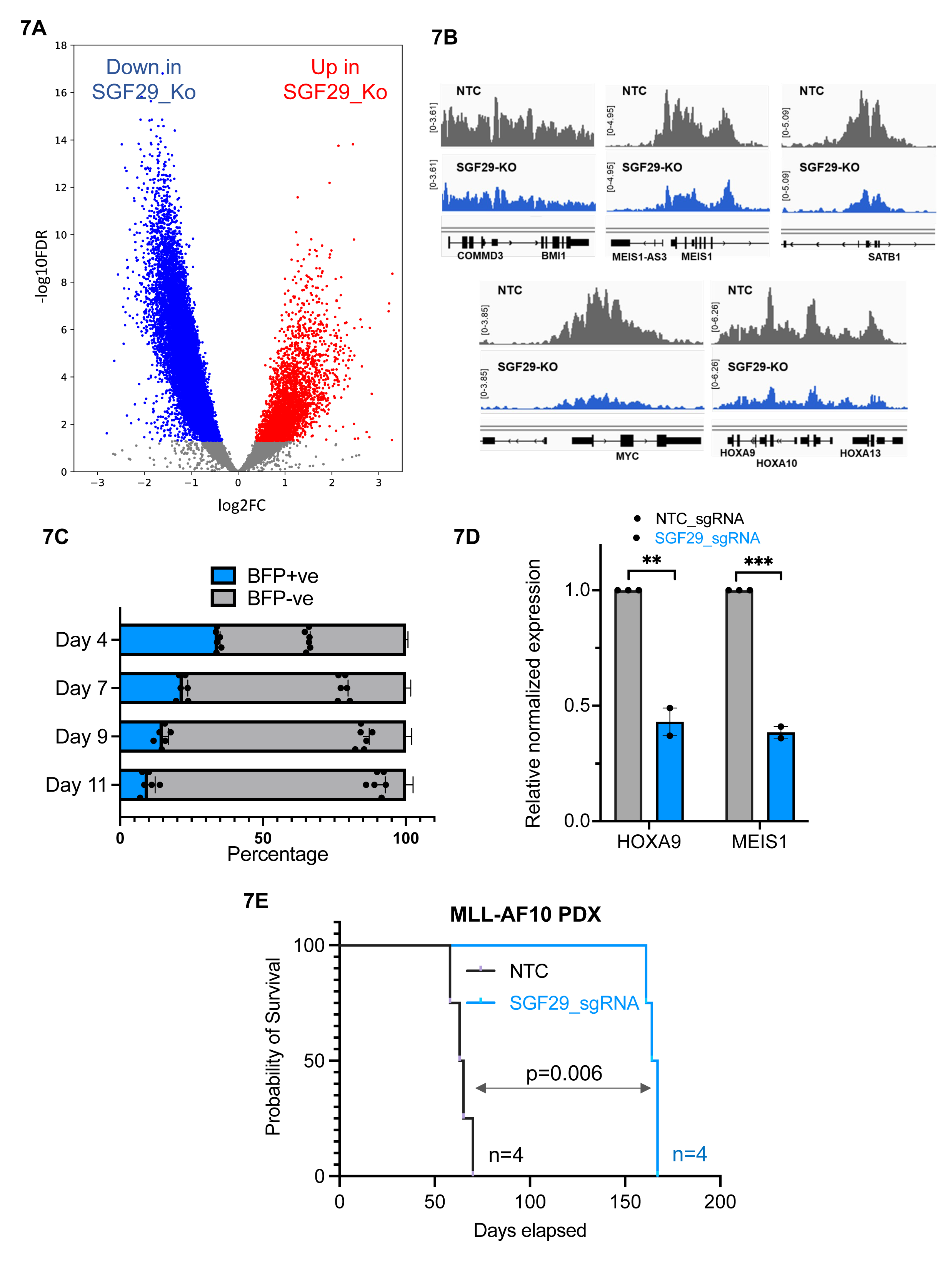
Effect on H3K9 acetylation and antileukemia activity of SGF29 in AML patient cells (A) Volcano plot indicating changes in H3K9 acetylation across the genome in UB3 cells after deletion of SGF29. Blue dots are the loci with decreased and red are the loci with increased acetylation. **(B)** Integrated genome viewer (IGV) tracks showing ChIP-seq peaks for H3K9ac on AML oncogene loci in U937 cells expressing non-targeting (grey), compared to SGF29 targeting sgRNAs (blue). **(C)** *In vitro* competition assay is shown with percentage of SGF29 sgRNA expressing BFP positive AML393 human AML cells (y-axis) is shown in blue bars, compared to control cells in grey. Progressive percentage of BFP is shown over time at indicated time points shown on the y-axis. **(D)** Relative expression of *HOXA9* and *MEIS1* in SGF29 deleted AML393 cells is plotted on the y-axis normalized to the SGF29 wildtype control (n=3). P values: **= p<0.05, *** = P<0.01. **(E)** Kaplan-Meier curves for NTC versus SGF29 deleted AML393 PDX (MLL-AF10 positive) are shown with the survival probability plotted on the y-axis.

### SGF29 deletion impairs *in vitro* and *in vivo* leukemogenesis in a patient-derived xenograft model of AML

We then wanted to assess the effect of SGF29 deletion in a human AML patient sample. For this, we used AML393 cells, a patient-derived xenograft expressing split-Cas9^36^. In these cells, SGF29 knockout using a BFP-co-expressing sgRNA led to a strong reduction in proliferative advantage in a competition assay compared to non-SGF29-sgRNA expressing cells (Fig. 7C). Quantitative real-time PCR (qRT-PCR) of SGF29-knockout compared to NTC transduced AML393 cells showed a significant reduction of *HOXA9* and *MEIS1* transcripts (Fig. 7D). We then injected the AML393 cells with SGF29 or NTC sgRNAs into NSG mice and monitored engraftment of human Cas9-GFP+ cells in peripheral blood by flow cytometry (data not shown). In comparison to the AML CDX models described in Fig. 3F&H, we observed more pronounced effects of SGF29 deletion in this human PDX model of AML. Whereas mice injected with NTC-targeting sgRNAs succumbed to AML with a median latency of 64 days, those injected with sgRNAs targeting SGF29 showed a median disease latency of 165 days post-injection (Fig. 7E). Taken together, our results demonstrate that SGF29 is important for sustaining critical transcriptional networks in leukemogenesis and may serve as an attractive therapeutic target in AML.

## DISCUSSION

In recent years, there has been increasing evidence that the expression of cancer-promoting oncogenes is sustained by specific chromatin modulators that are often essential and/or rate-limiting for oncogenesis. Thus, these epigenetic regulators may act as non-oncogene dependencies presenting attractive new therapeutic targets in cancer^6, 37–40^. The chromatin reader protein SGF29 was discovered in yeast as a component of the SAGA (Spt–Ada–Gcn5 acetyltransferase) complex ^48^ and is highly conserved across several species (reviewed in ^49, 50^). SGF29 was shown to contain two tandem Tudor domains at the C-terminus, which specifically bind di- and tri- methylated H3K4 residues ^29, 30^. In mammalian cells, SGF29 is a key component of diverse chromatin-modifying complexes with overlapping subunits. The Ada two-A-containing (ATAC) and the SAGA complexes are the most well-characterized and have distinct chromatin targets ^29^. As part of the enzymatic HAT module in SAGA (along with Gcn5, Ada3, and Ada2), SGF29 allows the complex to dock to pre-existing trimethyl marks at promoters and stimulates subsequent processive acetylation^51^. The SAGA complex is responsible for H3K9 acetylation, a modification that fine-tunes, rather than initiates, locus-specific transcriptional activity^52^. SAGA integrates multiple coactivator functions, but there are distinct genetic requirements for each module during gene regulation ^47^. Data from studies in murine MLL-AF9 leukemia suggest that ATAC is a generic requirement in cancer, whereas SAGA is selectively important in AML^35^. In *Saccharomyces cerevisiae,* Sgf29 is required for SAGA promoter recruitment and H3K9 acetylation *in vivo* ^29^ and our results show that this is also true in AML cells where the localization of human KAT2A is controlled by SGF29. This indicates that SGF29 may be rate-limiting for the oncogenic activities of SAGA in AML ^47^ and perhaps in other cancers where KAT2A activity is implicated in oncogenesis^48, 49^.

Interestingly, our results also show that SGF29 is an important regulator of the transcription as well as chromatin abundance of several leukemia-associated transcription factors in AML cells and plays an important role in sustaining the leukemogenesis of diverse AML oncoproteins. Our results indicate that SGF29 may regulate the differentiation block in AML, as its deletion promoted the expression of several differentiation-associated genes in both U937 and MOLM13 cells, enhanced phagocytic uptake, and led to a significant increase in the formation of colonies with a differentiated morphology from murine AMLs harboring distinct driver gene-fusions. One of the genes highly downregulated by SGF29 loss in AML was MYC, in agreement with observations in human hepatocellular carcinoma^50^. SGF29 has also recently been identified as a negative regulator of anti-tumor immunity in a mouse model of adenocarcinoma^51^, indicating that the protein may have cell-intrinsic as well as non-intrinsic tumor-promoting activities.

In this study, we demonstrate that a phenotypic pooled CRISPR screen based on the expression of *MEIS1* revealed multiple constituents of 6 chromatin-modifying complexes as regulators of a gene-expression program linked to AML oncogenesis. Among our top hits, we encountered “writers” and “readers” known to function in concert. For example, our top hits include the MYST complex gene KAT7, an acetyltransferase (writer), and its plant homeodomain (PHD)-containing acetylation reader, JADE3 (reader); or the methyltransferase DOT1L (writer) and AF10 or ENL (readers), or SGF29 (reader) and the acetyltransferase KAT2A (writer). These results are interesting because they not only validate the robustness of the screen but importantly, point to separate nodes for therapeutic targeting of the same complex. Several chromatin writers such as DOT1L, KAT2A, and KAT7 are known to be targeted to different gene loci by separate chromatin readers. Therefore, targeting the chromatin reader required for the specific localization of leukemia-promoting chromatin writers to oncogene loci may arguably present a better safety profile in terms of therapeutic targeting compared to targeting the chromatin writers themselves. The paradigm of inhibiting reader proteins began with the development of tool compounds targeting bromodomains^52^, which spurred translation to clinical candidates. Other efforts for reader targeting include PHD fingers, WD40 repeat domains, and Royal family methyl-lysine readers (MBT, chromodomain, Tudor domain, PWWP domain, and the YEATS domain ^53–58^. Royal family domains all share a structural aromatic cage that binds methyl-lysine side chains and for discrimination between lysine mono-, di-, and tri- methylation residues. These structural features comprise pockets of amenable size and hydrophobicity which may allow small-molecule binding. Our work provides a rationale for inhibitors targeting the chromatin reading activity of the Tudor domain, which is required for the chromatin occupancy of SGF29-associated complexes, including SAGA, and transcriptional coactivation of oncogenic gene expression programs. We believe that small-molecule inhibitors of the SGF29 Tudor domain will have potent anti-leukemia effects and may also be effective in other cancers driven by activated expression of the *HOX/MEIS* or *MYC* oncogenes.

## METHODS

### Animal Studies

All animal studies were approved and performed as per the guidelines of the SBP Medical Discovery Institutional Animal Care and Use Committee. 200 uL of 200,000 UB3 or MOLM13 cells expressing Cas9 and NTC or SGF29 targeting sgRNA were injected intravenously by tail vein injections into NOD/SCID mice (Jackson Laboratories, #005557). Mice were monitored every day and sacrificed when critically ill. For MLL-AF10 PDX studies, 150,000 cells expressing split Cas9 and NTC or SGF29 sgRNA were injected into NSG mice (Jackson Laboratories, #013062) by tail vein injections. Mice were bled every 2 to 3 weeks for checking AML cell engraftment by flow cytometry. Blood was collected by retro-orbital bleeding and red blood cells were lysed using 1X PharmLyse Buffer (BD Biosciences), then washed with 1X PBS, and analyzed on the LSR Fortessa. Animals were sacrificed when critically ill.

### Cell culture

Human leukemia cell lines U937 (kind gift from Dr. Daniel Tenen, Beth Israel Deaconess Medical Center) and MOLM13 (ACC-554, DSMZ), were cultured in RPMI-1640 medium supplemented with 2 mM L-glutamine and sodium pyruvate, 10% Fetal bovine serum and 50 U/ml Penicillin/Streptomycin (Thermo Fisher Scientific, Carlsbad, CA), 2 mM L-glutamine and incubated in 5% CO_2_ at 37°C. Murine leukemia cells were cultured in DMEM medium supplemented with 2 mM L-glutamine, 15% FBS and 50U/ml Penicillin/Streptomycin, in the presence of the following cytokines: 10 ng/ml murine interleukin 6 (mIL-6), 6 ng/ml murine interleukin 3 (mIL3) and 20 ng/ml murine stem cell factor (mSCF) (all from Peprotech, Rocky Hill, NJ), and incubated at 5% CO_2_ and 37°C. HEK293T cells were cultured in DMEM medium supplemented with 2 mM L-glutamine and sodium pyruvate, 10% FBS and 50 U/ml Penicillin/Streptomycin, and incubated in 5% CO_2_ at 37°C. MLL-AF10 patient-derived xenograft (PDX) cells were cultured in IMDM medium supplemented with 20% BIT9500 (STEMCELL Technologies), human cytokines, and StemRegenin 1 44 (SR1), and UM171 as described earlier^59^.

### CRISPR Screen data analysis

Fastq files from the epigenetics CRISPR screen were analyzed using the MAGeCK-VISPR MLE pipeline ^60^. Gene rankings were generated comparing the GFP -low and - high fractions to T0. Both experimental conditions were further compared using MAGeCKFlute ^61^. For subsequent functional hit-validation, the 15 genes with the greatest differences in beta scores in the GFP-Low minus GFP-High conditions were considered as candidate epigenetic regulator hits.

## Data availability

Sequencing data for RNA-seq, ChIP-seq, high-throughput CRISPR screens, and CROP-seq are deposited in the NCBI GEO under accession number: GSE217829. The sgRNA sequences used for the Human Epigenetics CRISPR Library are deposited in: https://github.com/kobarbosa/HOX-Manuscript.

## Code availability

The scripts used in this manuscript are available at: https://github.com/kobarbosa/HOX-Manuscript.

## Authors contributions

**K.B.** and **A.D**. contributed equally to the study. **KB.:** conceptualization, data curation, formal analysis, funding acquisition, investigation, methodology, project administration, software, supervision, validation, visualization, writing-original draft. **A.D.:** conceptualization, data curation, formal analysis, investigation, methodology, project administration, software, supervision, validation, visualization, writing-original draft. **M.P.:** investigation, visualization, validation**. P.X.:** resources. **R.K.H.:** resources. **R.M**: data curation, formal analysis, methodology, software. **A.M.:** data curation, formal analysis, methodology, software. **N.R.:** data curation, formal analysis, methodology, software. **F.S.:** data curation, formal analysis, methodology, software. **X.L.:** data curation, formal analysis, software**. Y.S.:** investigation. **D.A.:** resources. **I. J.:** resources. **A.B.:** resources, software. **J.D.:** resources, software. **P.M.:** resources**. E.R.:** conceptualization, resources, software. **J.S.:** conceptualization, resources. **P.D.A.:** conceptualization, resources. **A.J.D.:** conceptualization, resources, funding acquisition, investigation, methodology, project administration, supervision, visualization, writing-original draft.

## Acknowledgments

We would like to thank Adriana Charbono and Buddy Charbono for their invaluable assistance with mouse studies. Dr. Chih-Cheng Yang and Dr. Chun-Teng Huang from the Sanford Burnham Prebys Medical Discovery Institute (SBP) functional genomics core, Yoav Altman from the SBP Flow Cytometry Core, and Dr. Kang Liu from the Genomics Core for their excellent support. We would like to acknowledge the help of Dr. Philip Koeffler, Dr. Gery Sigal, Dr. Sarah Parker, and Dr. Aleks Scotland for their help with chromatin proteomics at Cedars Sinai.

This work was supported by National Institutes of Health (NIH) National Cancer Institute grants CA262746 and P30 CA030199, the Rally Foundation for Childhood Cancer Research and the Luke Tatsu Johnson Foundation grant (Award number 22IC33), an Emerging Scientist Award from the Children’s Cancer Research Fund (Award Number 20IC17), the V Foundation for Cancer Research (TVF) under award number DVP2019- 015, the Department of Defense Horizon Award grant W81XWH-20-1-0703.

## SUPPLEMENTARY FIGURE LEGENDS

**Fig. S1A:**
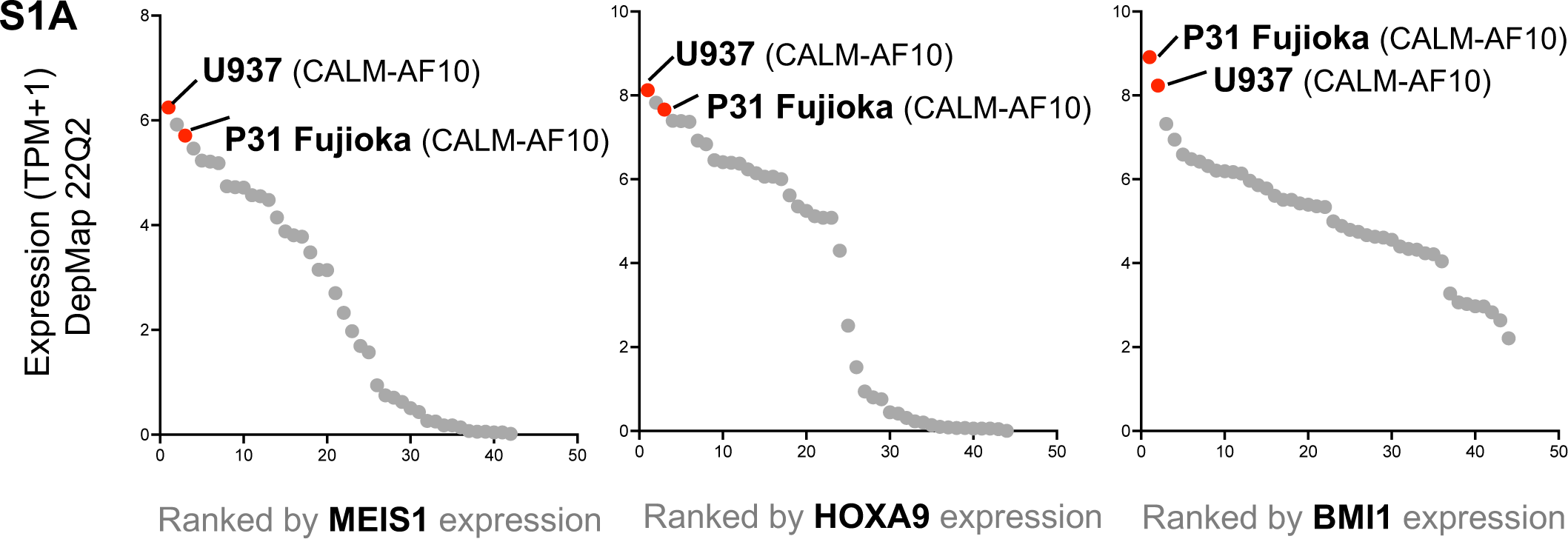
Expression of indicated genes (MEIS1, HOXA9 and BMI1) in 42 AML cell lines from the DepMap data ranked by expression (x-axis is rank, high to low expression), (y-axis is log transformed transcripts per million (TPM)+1) values.

**Fig. S1B:**
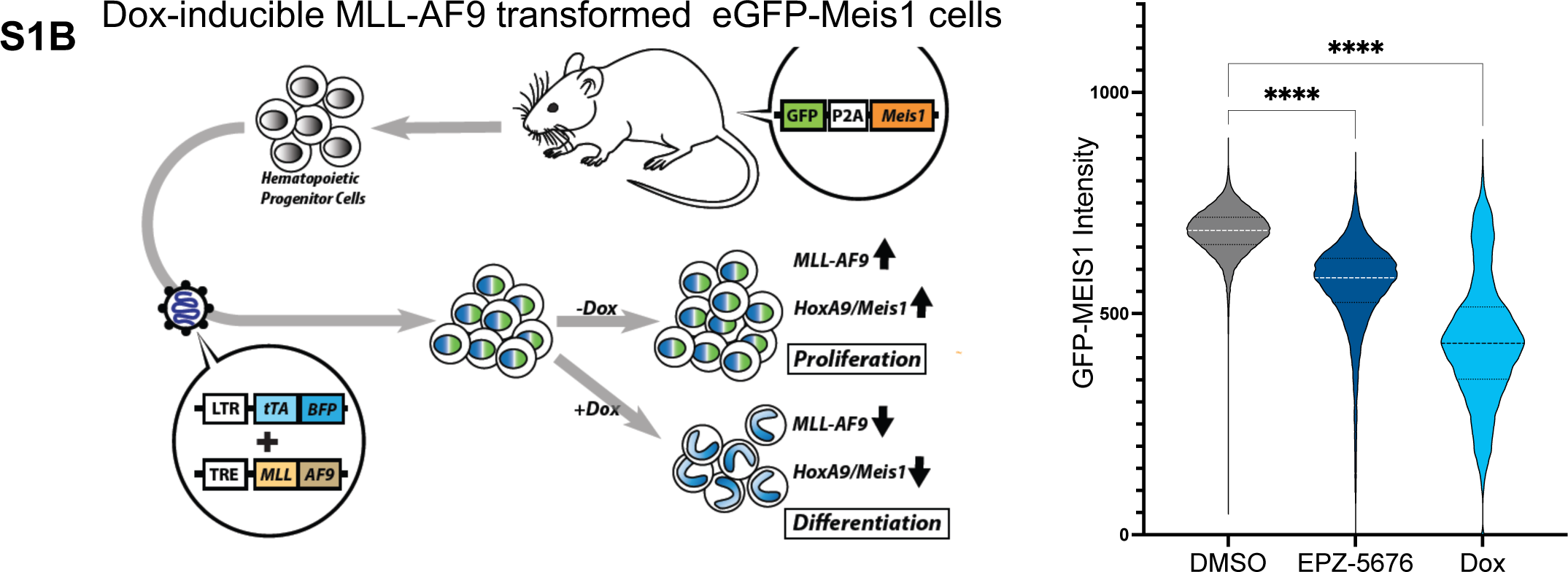
Schematic for generation of inducible Tet-Off MLL-AF9 mouse AML in the GFP- Meis1 background. GFP fluorescence intensity measured upon treatment with DMSO, prostaglandin D2, DOT1L inhibitor, or doxycycline in mouse Tet-Off MLL-AF9 AML (n=3)

**Fig S1C:**
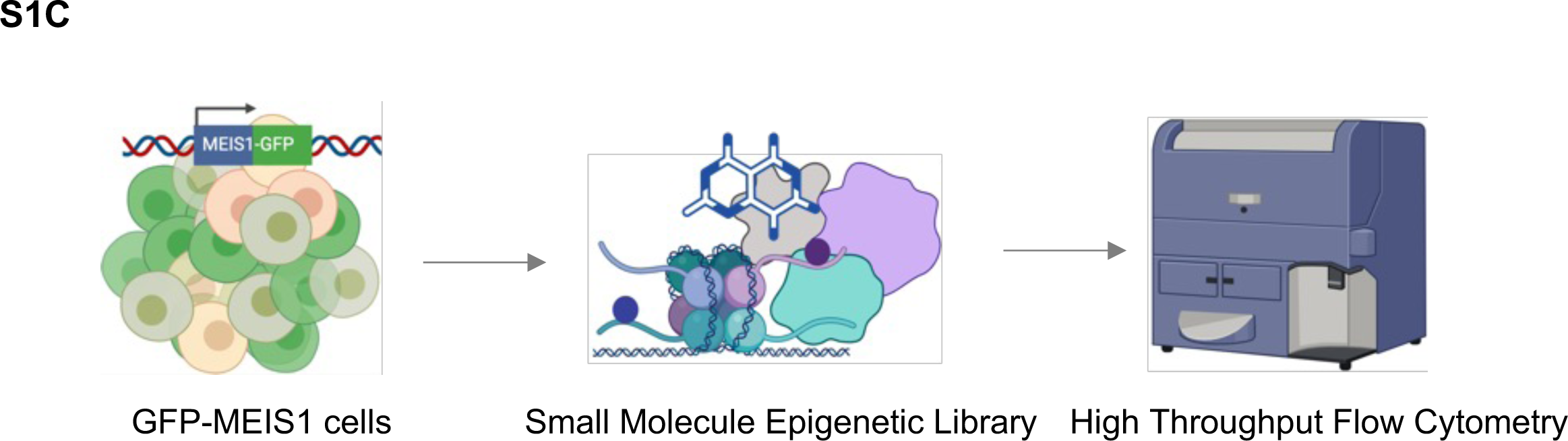
Schematic for phenotypic epigenetics compound screen.

**Fig. S2D:**
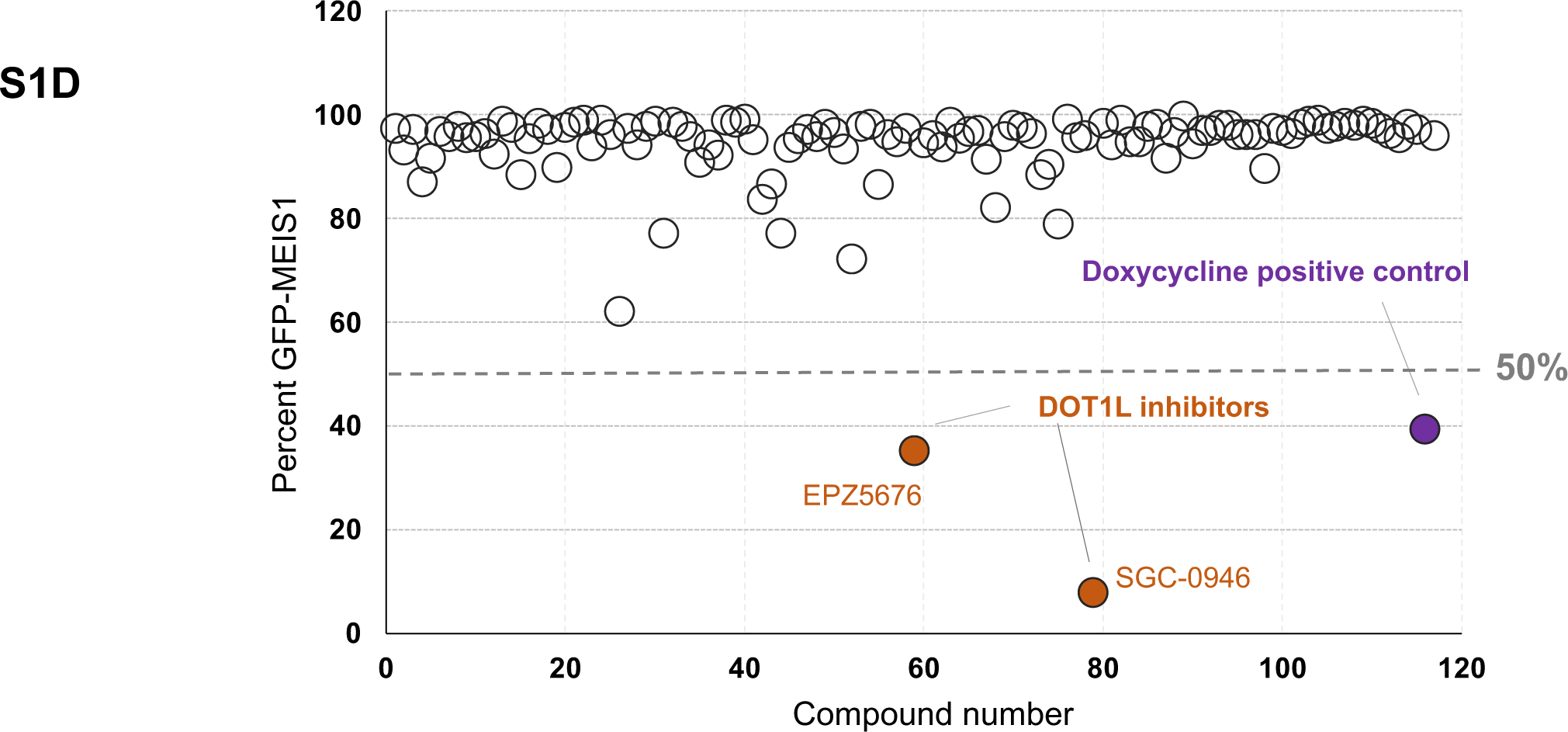
Screen results for Tet-Off mouse MLL-AF9 AML; the x-axis indicates % viability on day 5 after treatment with small molecules; the y-axis indicates the percentage of decrease in GFP-Meis1 signal compared to controls. Doxycyline was a positive control that shut off MLL-AF9 expression.

**Fig. S2A:**
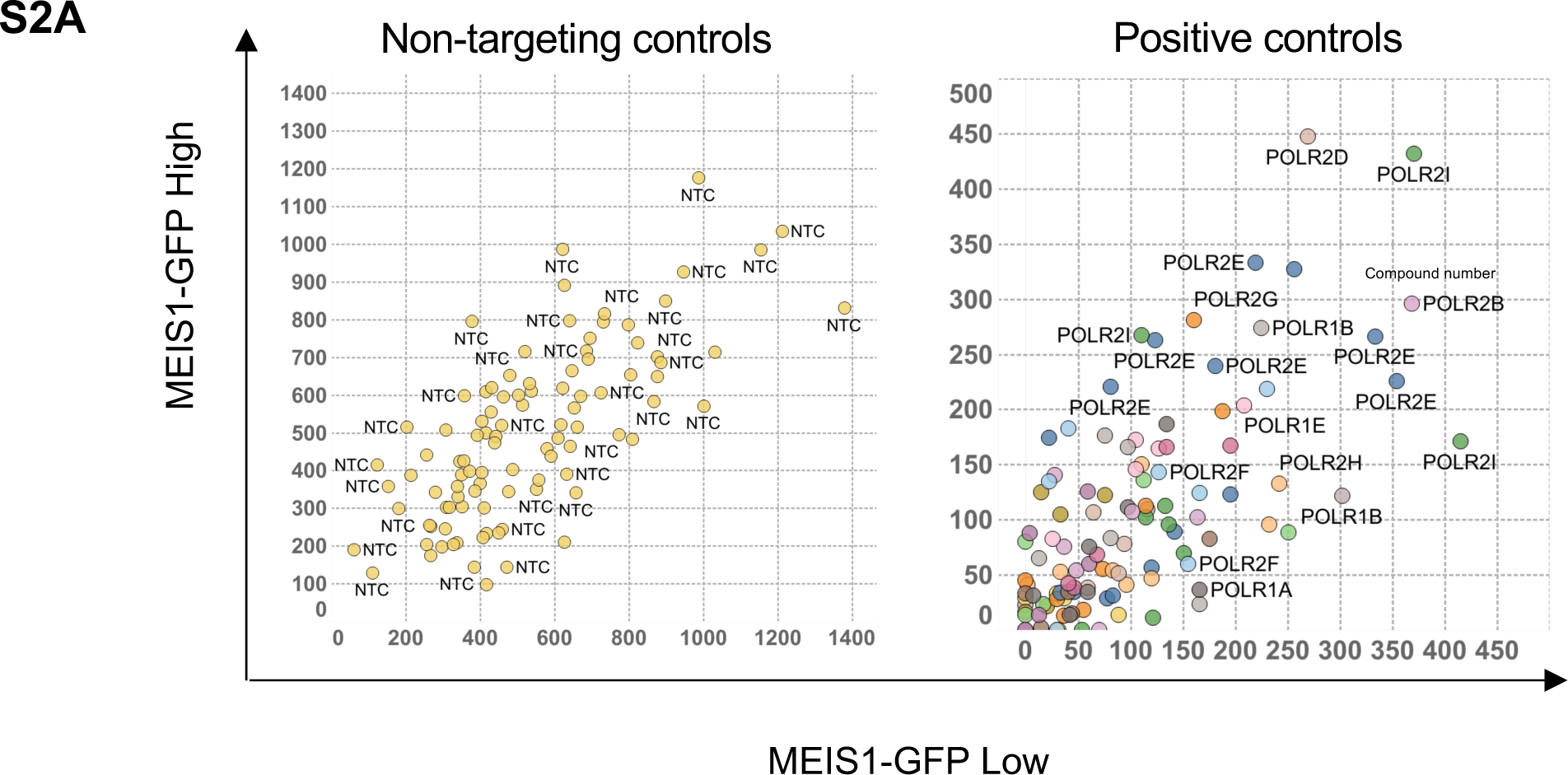
sgRNA normalized counts in each sorted fraction (low vs high). Panels depict counts for non-targeting controls and polymerase essential genes, as well as the known HOX/MEIS regulators DOT1L and ENL/MLLT1.

**Fig. S2B:**
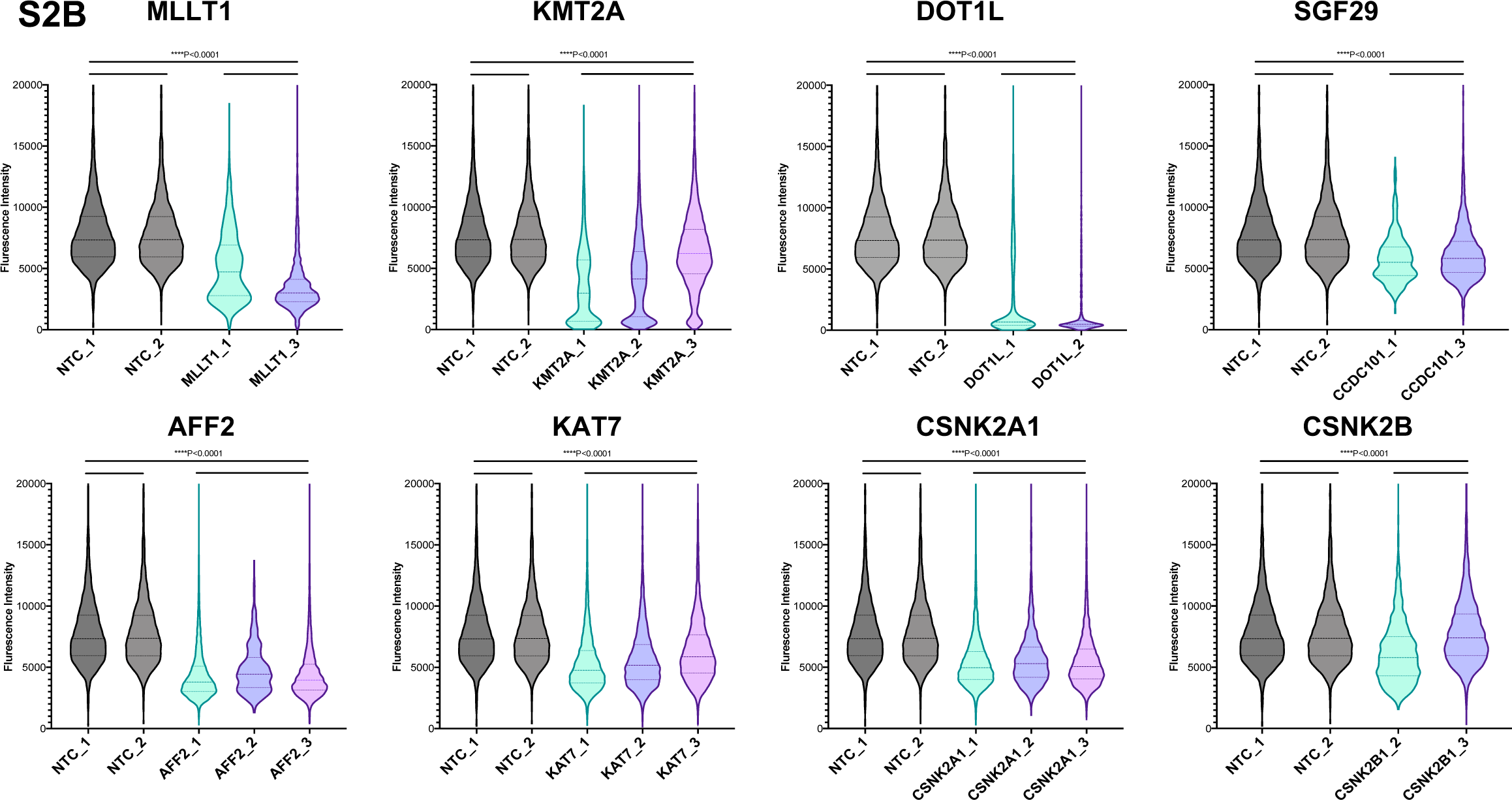
Violin plots show the GFP-MEIS fluorescence intensity values plotted on the y-axis from 2-3 unique sgRNAs targeting the indicated genes, compared to 2 non-targeting controls (NTC), measured 7 days after puromycin selection.

**Fig. S2C:**
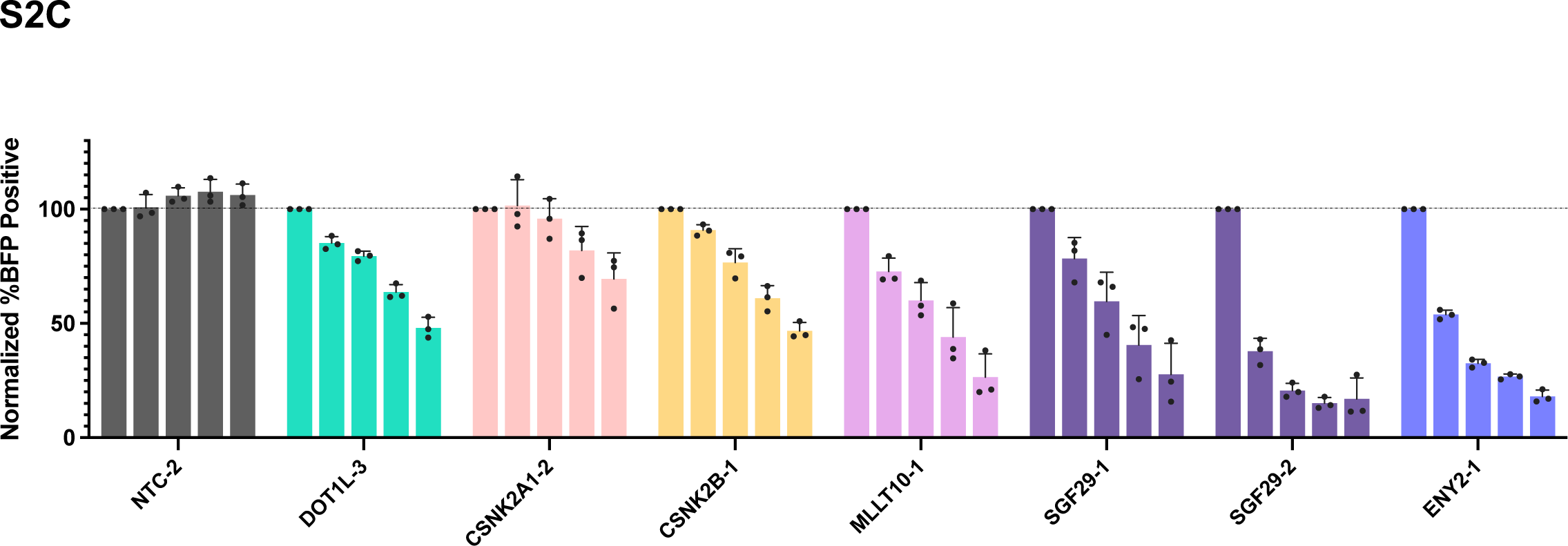
MOLM13 BFP+ cells over time, normalized to the baseline measurement; the x-axis indicates the sgRNA number for non-targeting controls (NTC) or each gene (n=3). Each bar represents a timepoint measurement: baseline, day 8, day 11, and day 14 for U937; baseline, day 6, day 12, day 18, and day 24 for MOLM13.

**Fig.S2D: Uniform.**
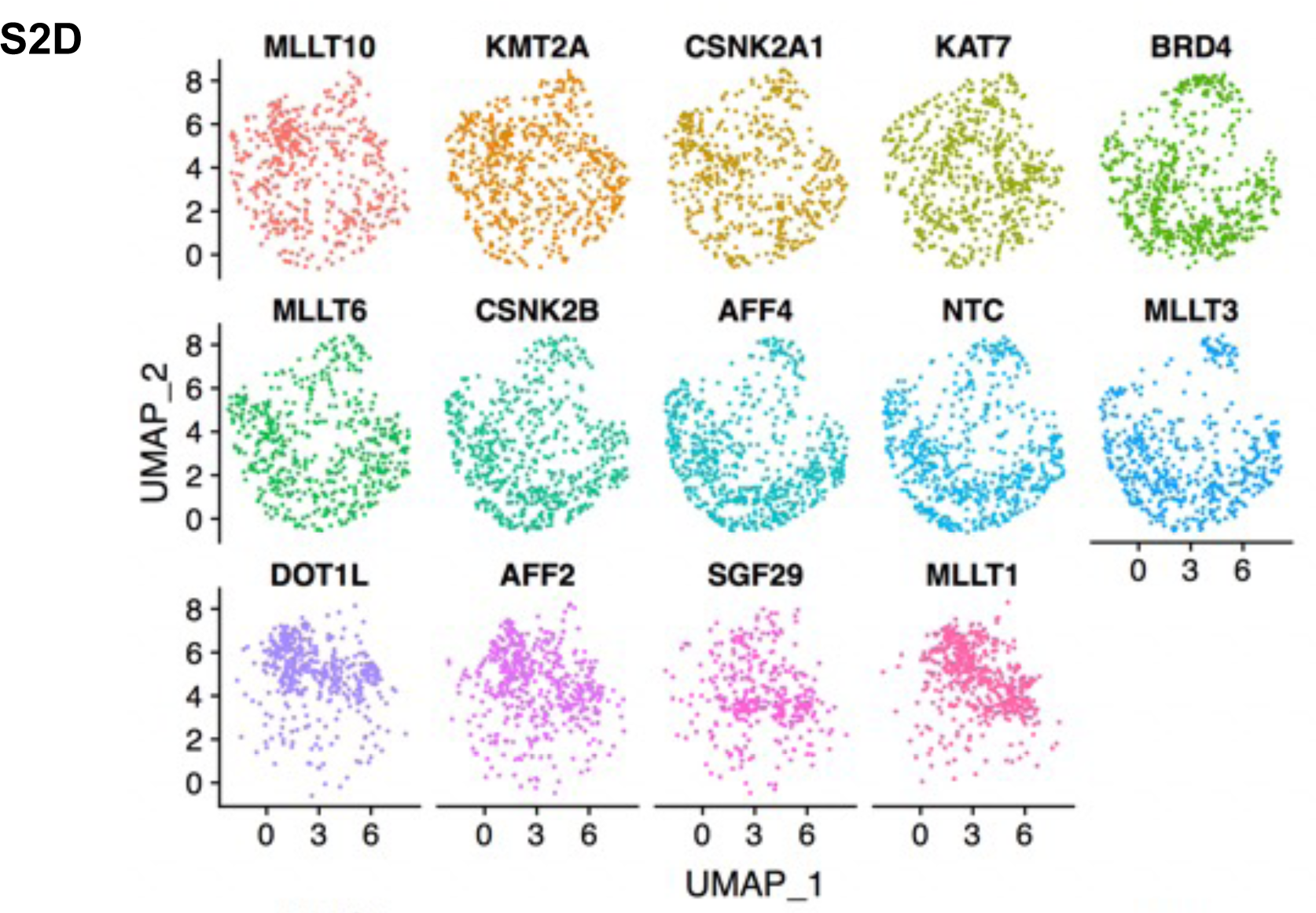
Manifold Approximation (UMAP) plots of synthetic gene expression profiles from CROP-Seq perturbations in U937 GFP-MEIS1 cells.

**Fig. S2E-F:**
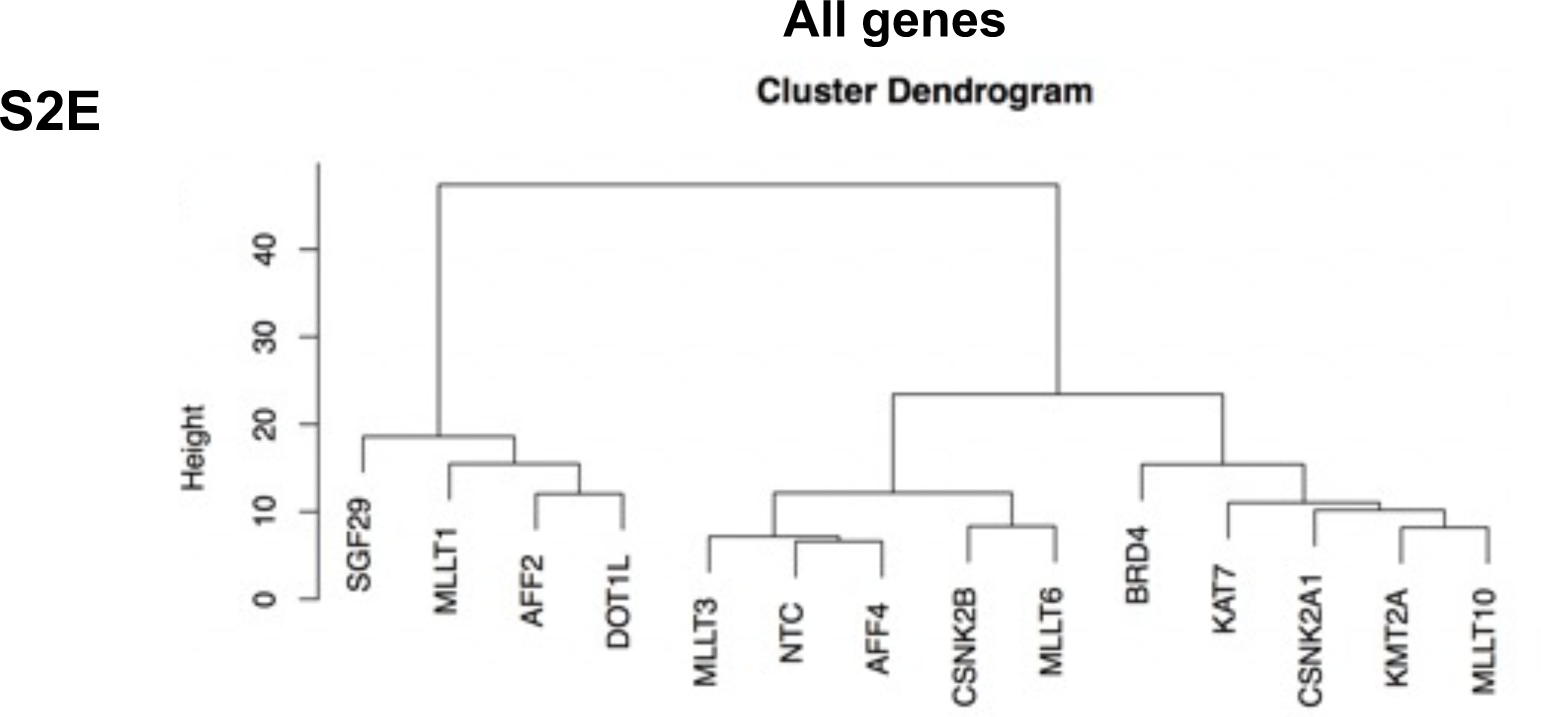
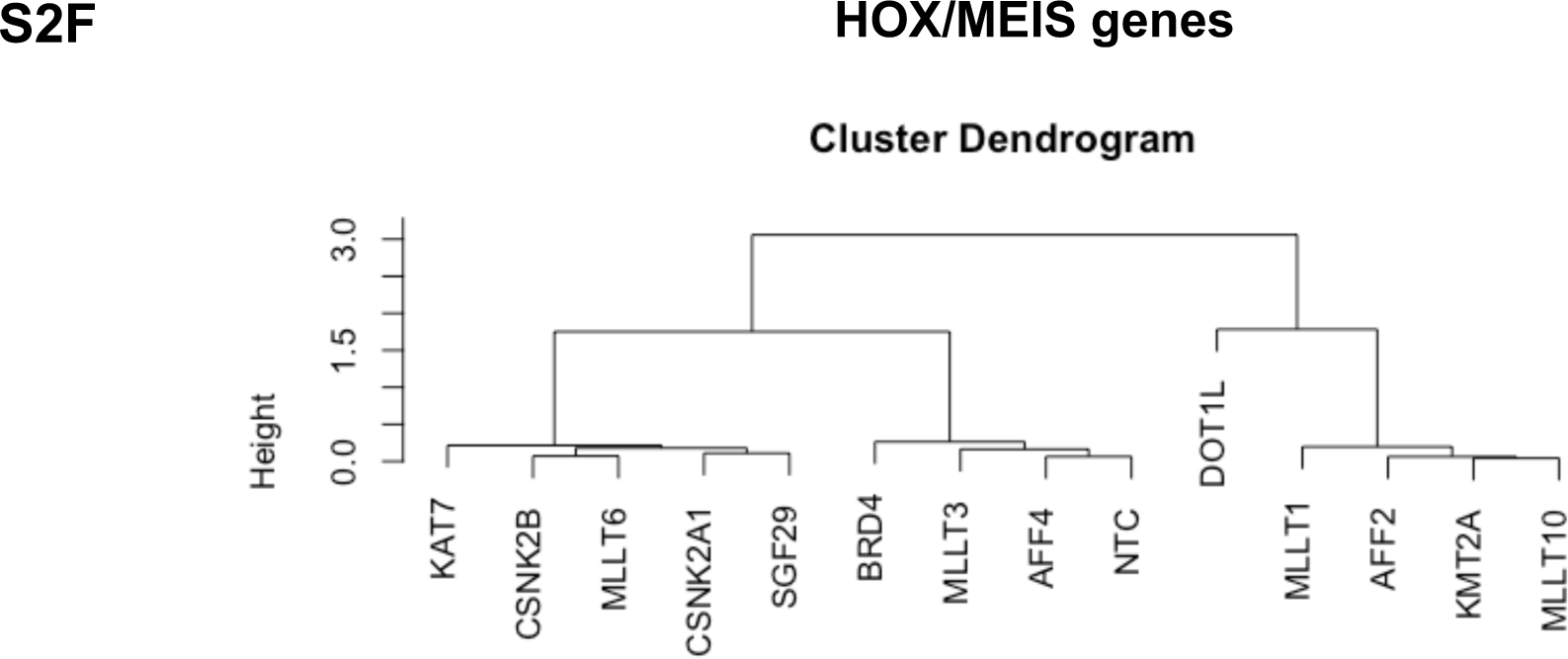
Hierarchical clustering dendrogram for gene expression of variable features: all genes (**Fig. S2E**) or HOXA-cluster genes and MEIS1 (**Fig. S2F**, see Methods). Branches corresponding to perturbed genes by CRISPR.

**Fig. S2G:**
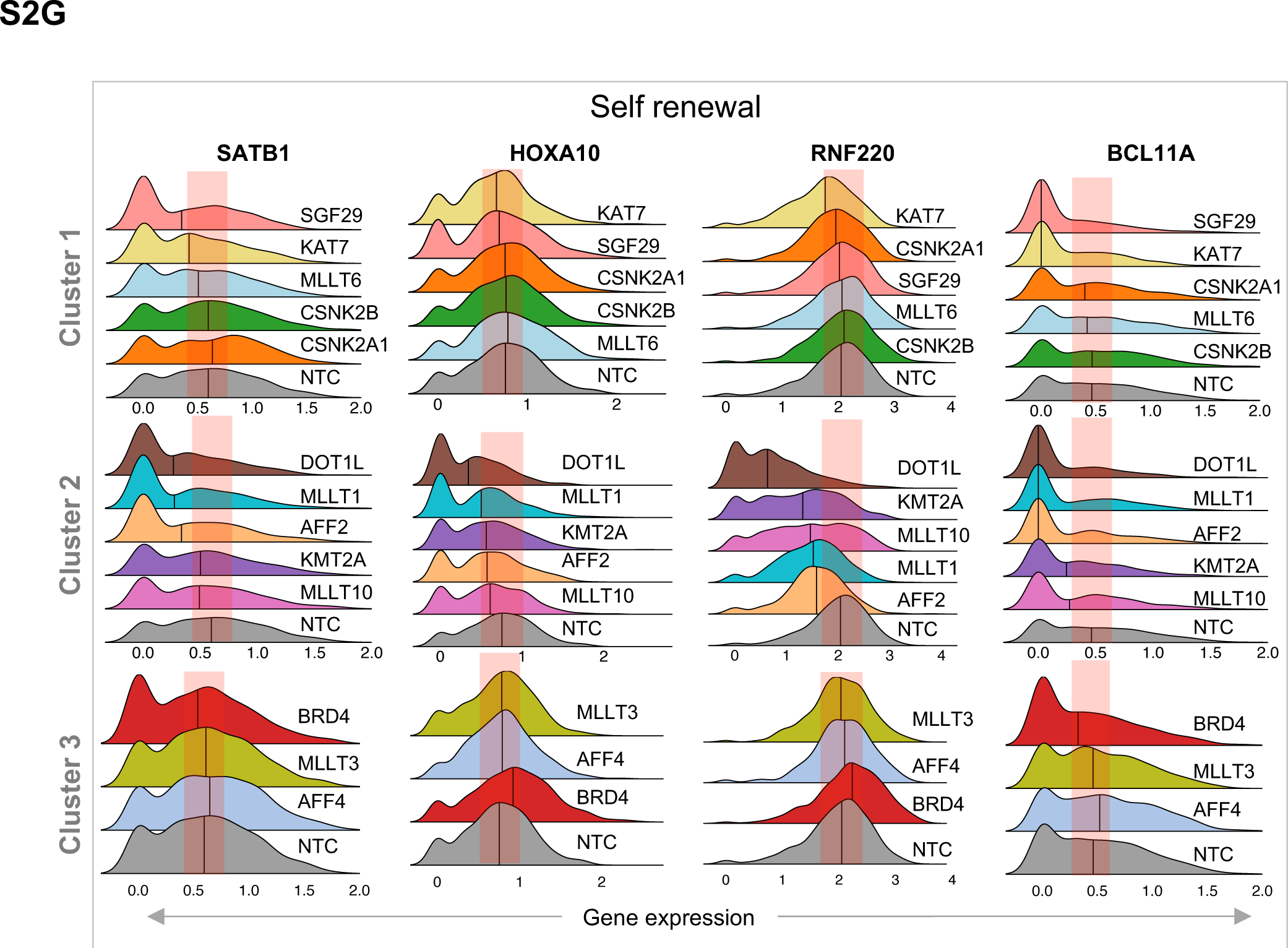
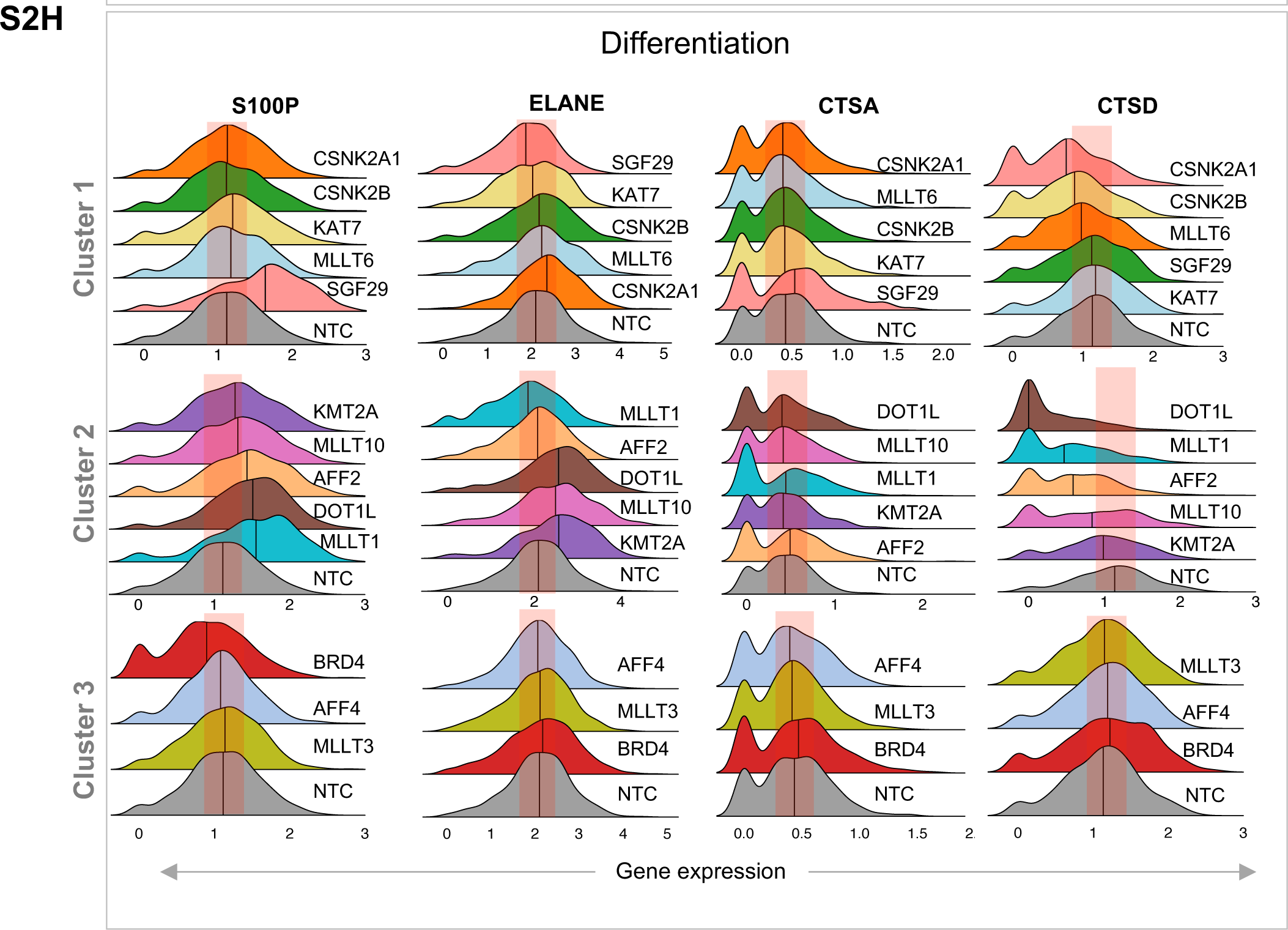
Ridge plots for CROP-Seq perturbations. Knockout genes are indicated on the y-axis and RNA expression levels for the (**Fig. S2G**) self-renewal-associated genes or (**Fig. S2H**) differentiation-associated genes are on the x-axis. The median expression value is indicated.

**Fig. S2I:**
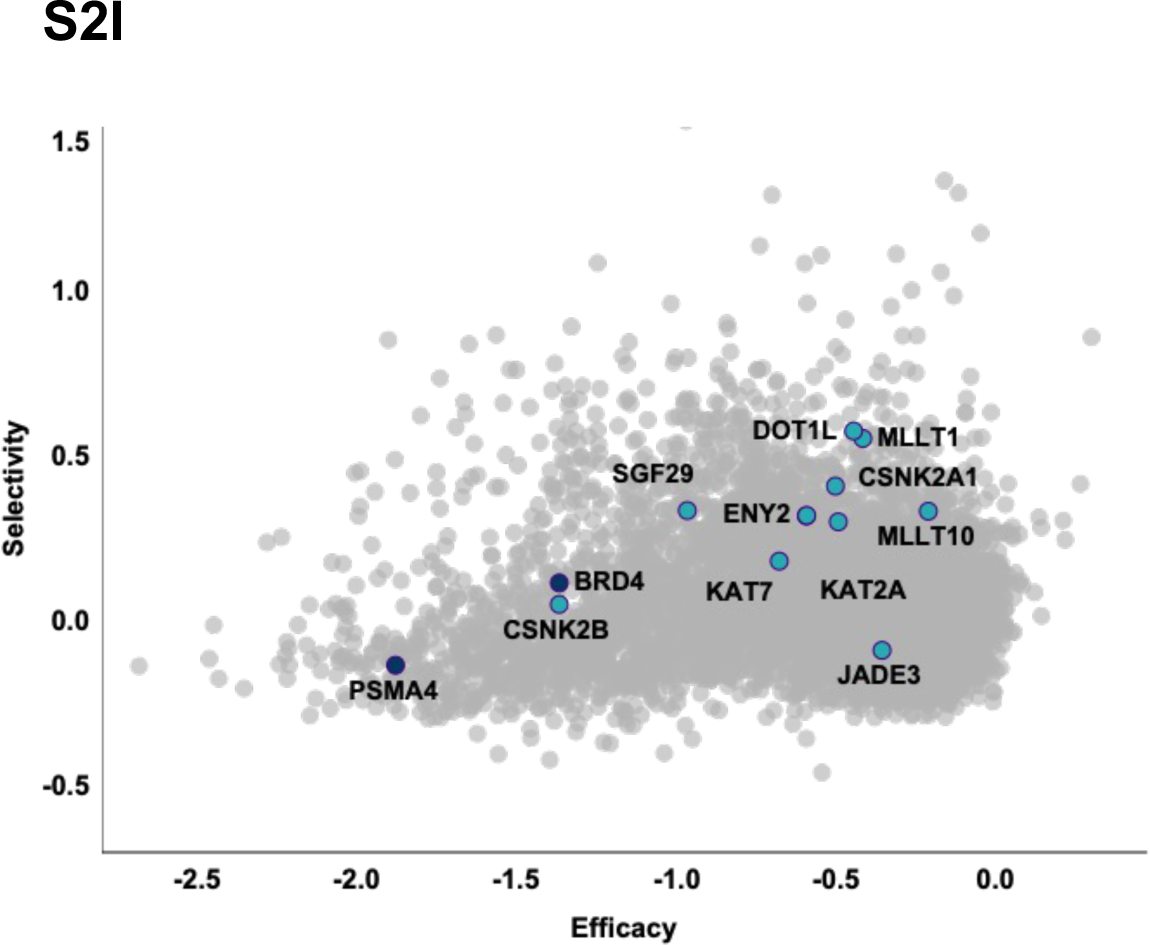
Selectivity (y-axis) and efficacy (x-axis) analysis for candidate hits in the DepMap data is shown in the scatter plot. PSME4 and BRD4 are included as examples of genes with high efficacy and low selectivity. Each dot in the plot is a gene from the DepMap data. More negative efficacy values indicate higher dependence or fitness score and higher values on the selectivity plot indicates higher differences between fitness scores of sensitive and insensitive cell lines.

**Fig. S3A:**
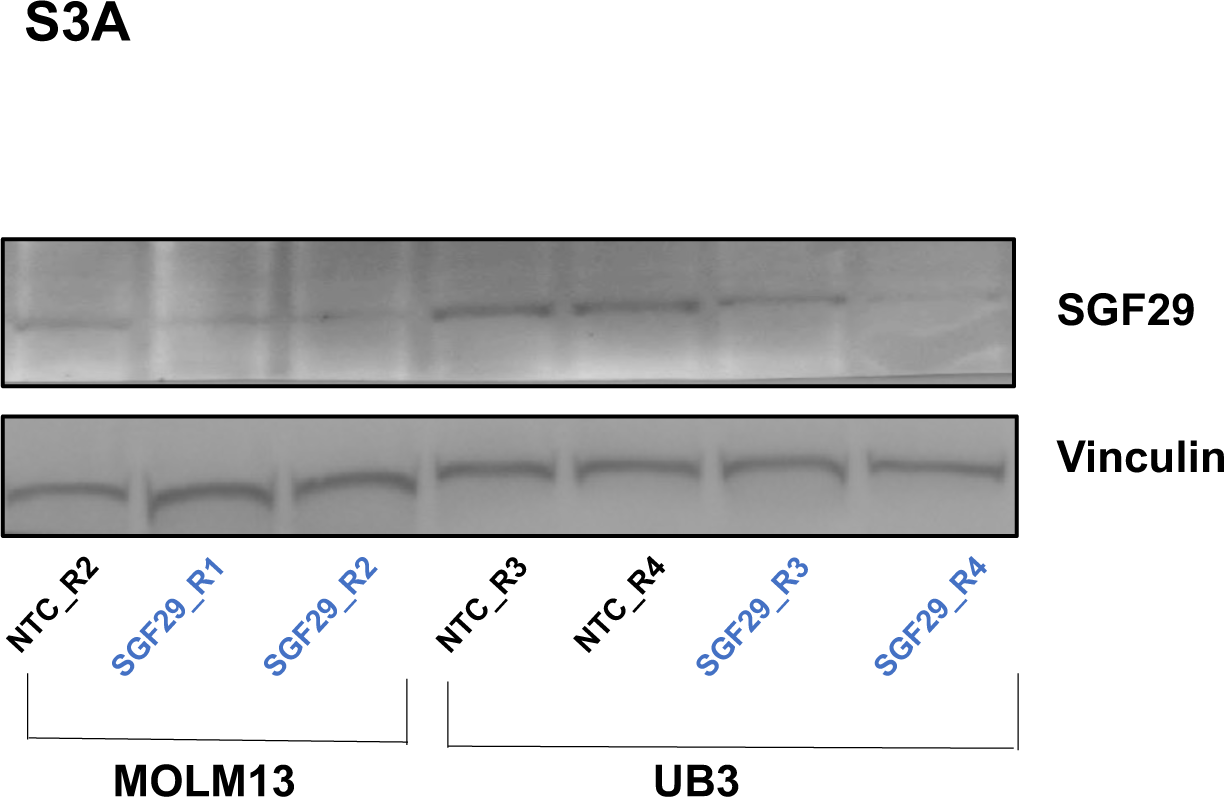
Western blotting for SGF29 in non-targeting control treated MOLM13 and U937 lysates compared to vinculin is shown as a loading control.

**Fig. S3B:**
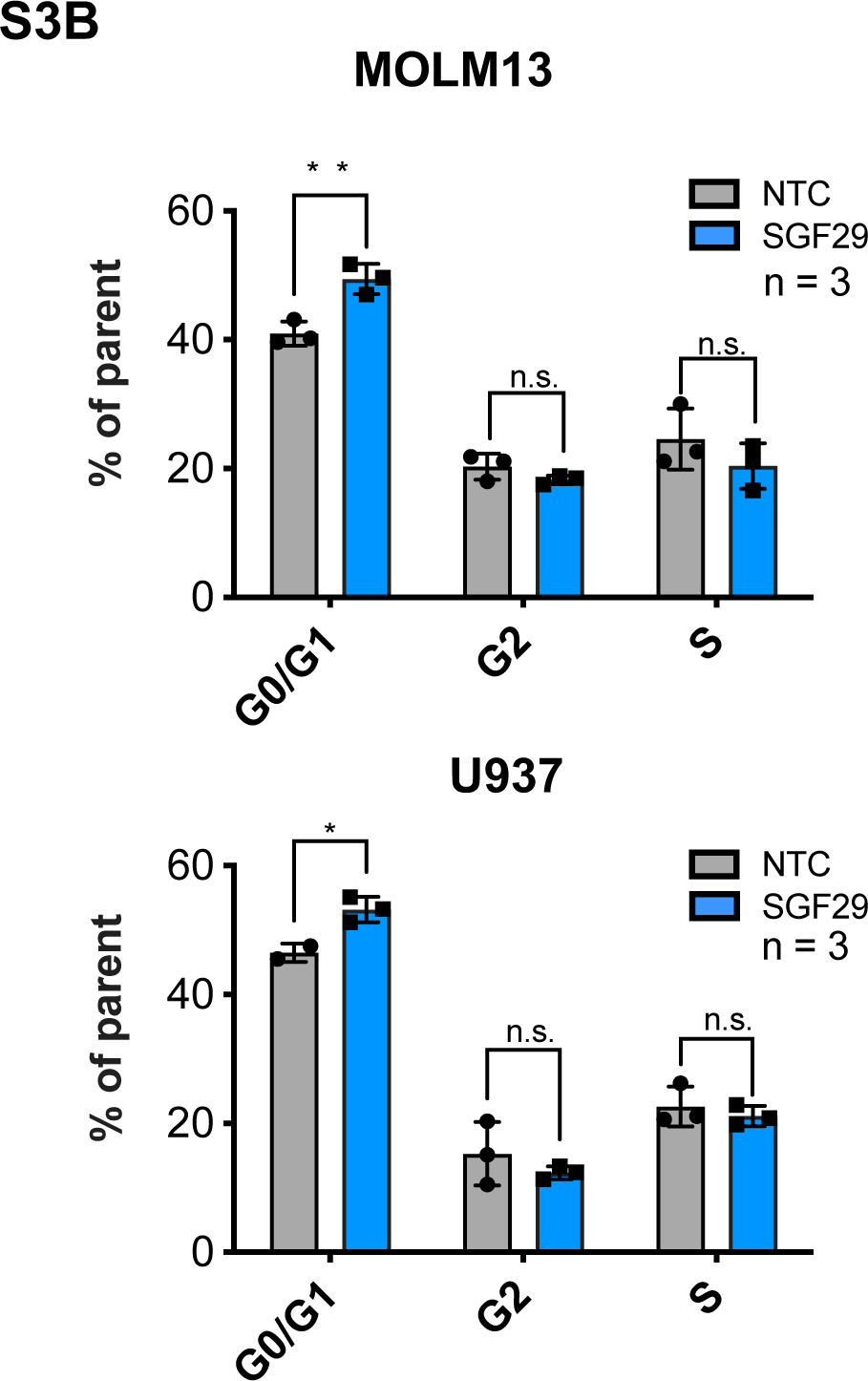
Cell cycle progression analysis by propidium iodide staining in human AML cells expressing NTC (blue) or SGF29 (light violset) sgRNAs (n=3).

**Fig. S3C:**
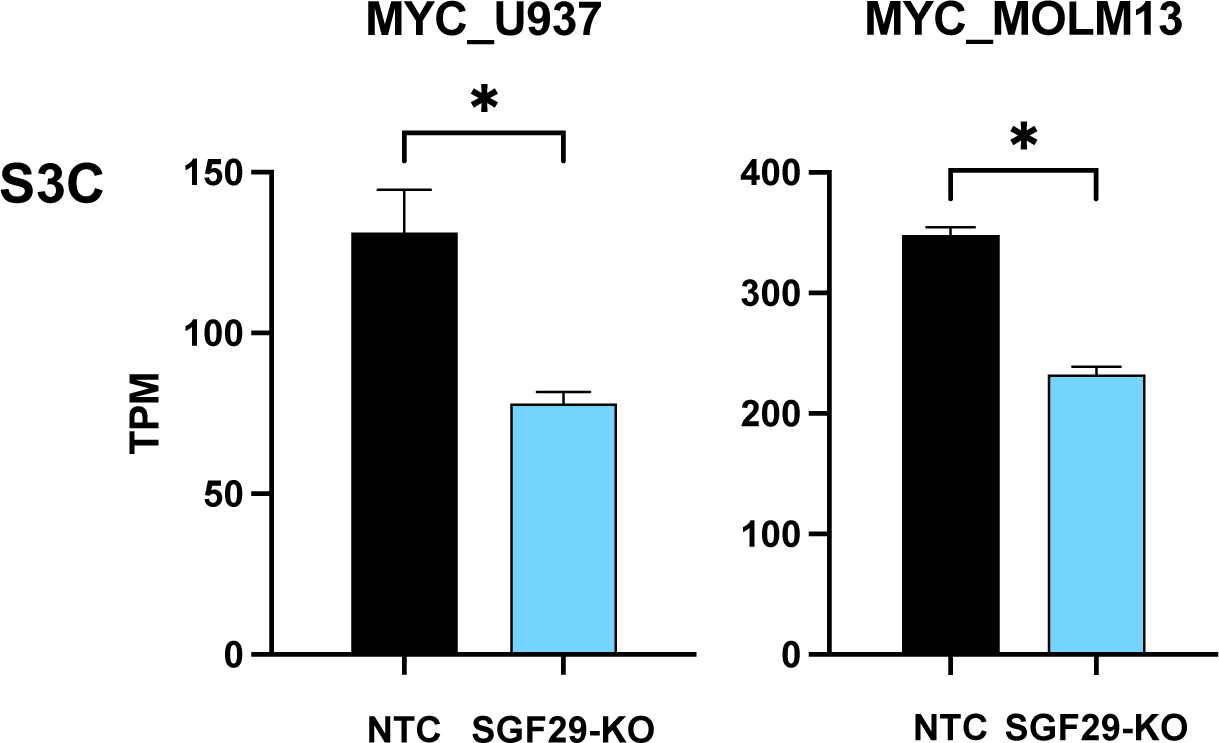
Differences in MYC transcript levels are shown as TPMs on the y-axis in U937 and MOLM13 cells with SGF29 deletion.

**Fig. S3D:**
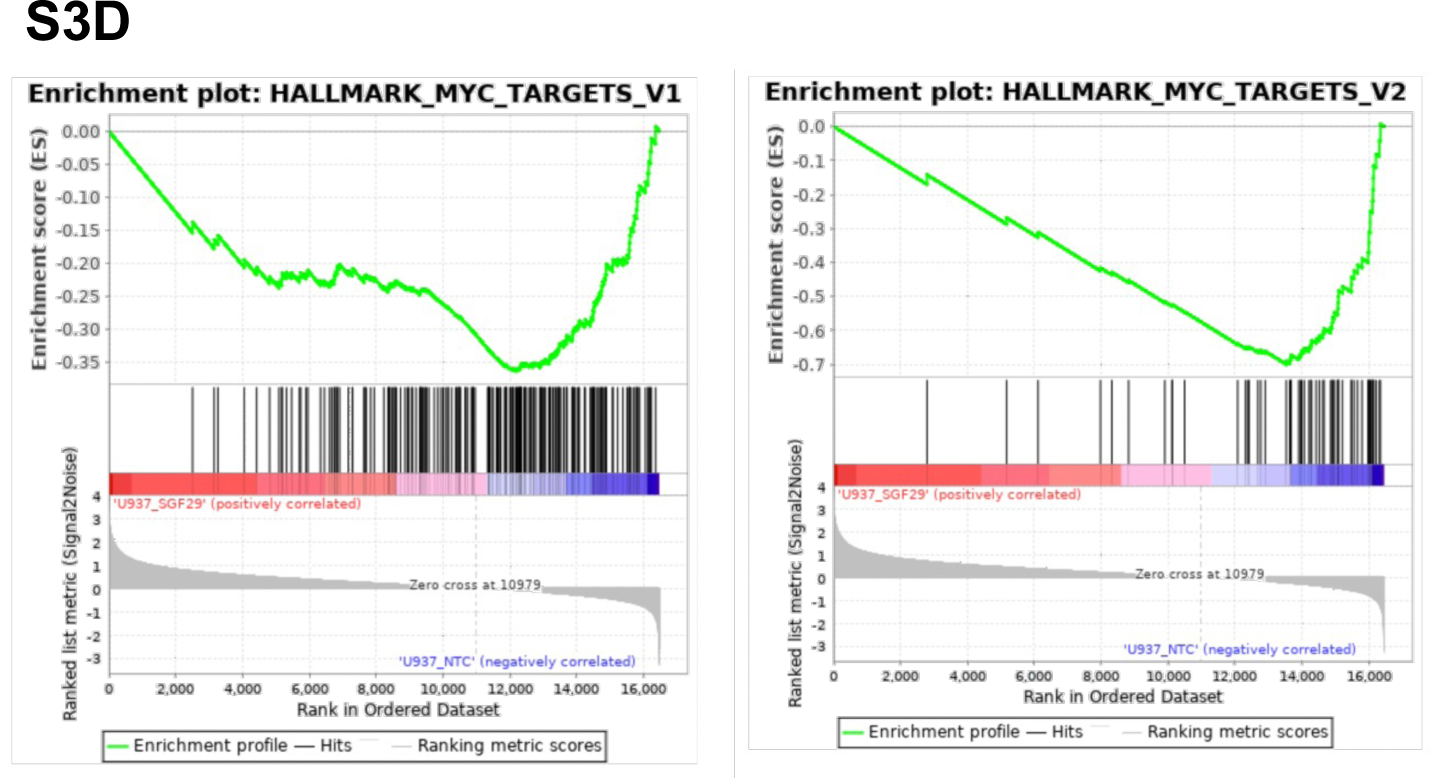
Hallmarks gene set enrichment analysis (GSEA) for the U937 cell line with SGF29 deletion compared to non-targeting control is shown.

**Fig. S4A:**
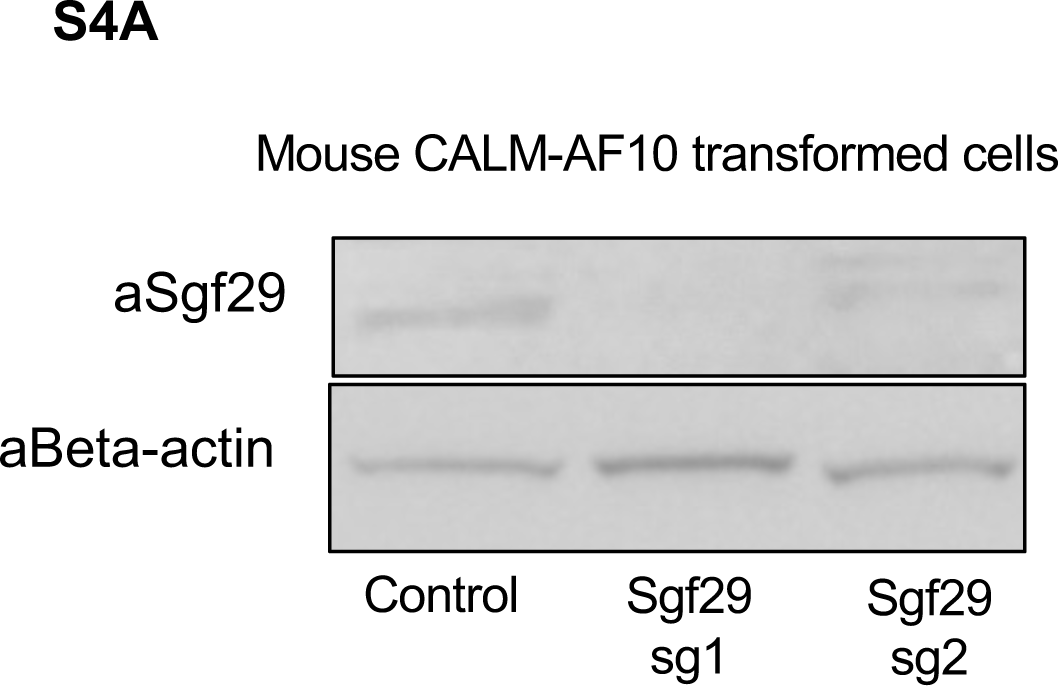
Western blot demonstrating loss of SGF29 protein in murine CALM-AF10 leukemia cells transduced with SGF29 sgRNA1 or sgRNA2. Beta Actin as a loading control.

**Fig. S4B:**
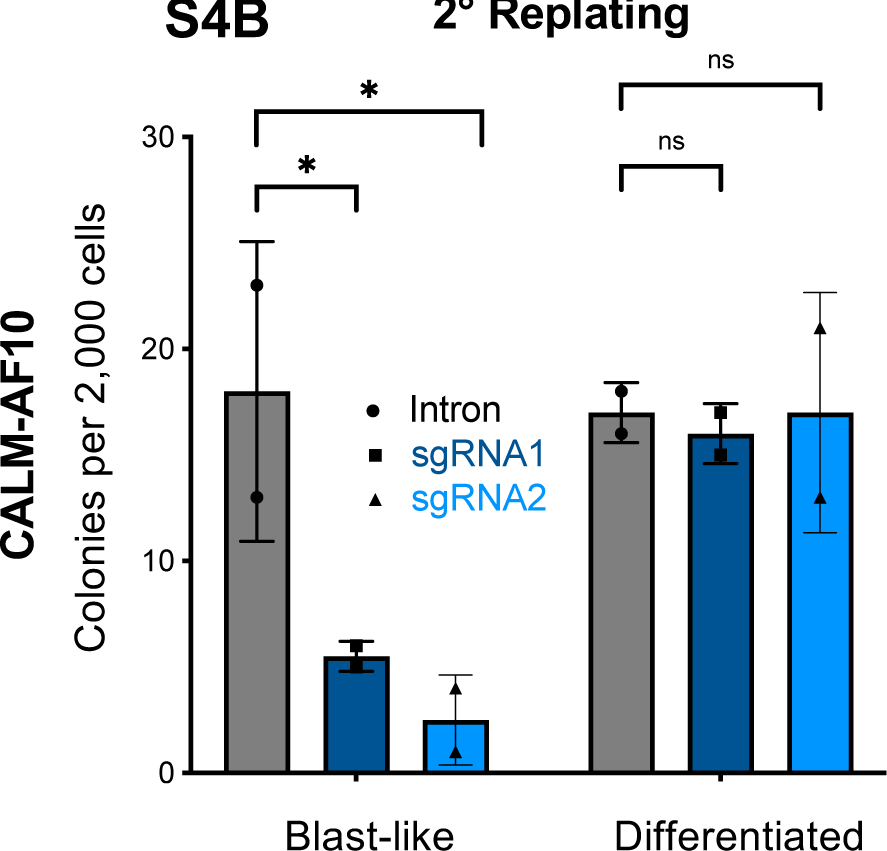
Colony forming unit (CFU) assay in murine CALM-AF10 leukemia. 2 nd week of CFU assay showing number of Blast-like as well as differentiated type colonies in cells transduced with intron targeting control or SGF29 targeting sgRNAs. Gray bar indicates cells with Intron whereas Blue bars indicate Sgf29 sgRNA expressing cells. (n=3); ns=non-significant; *=P<0.05.

**Fig. S4C:**
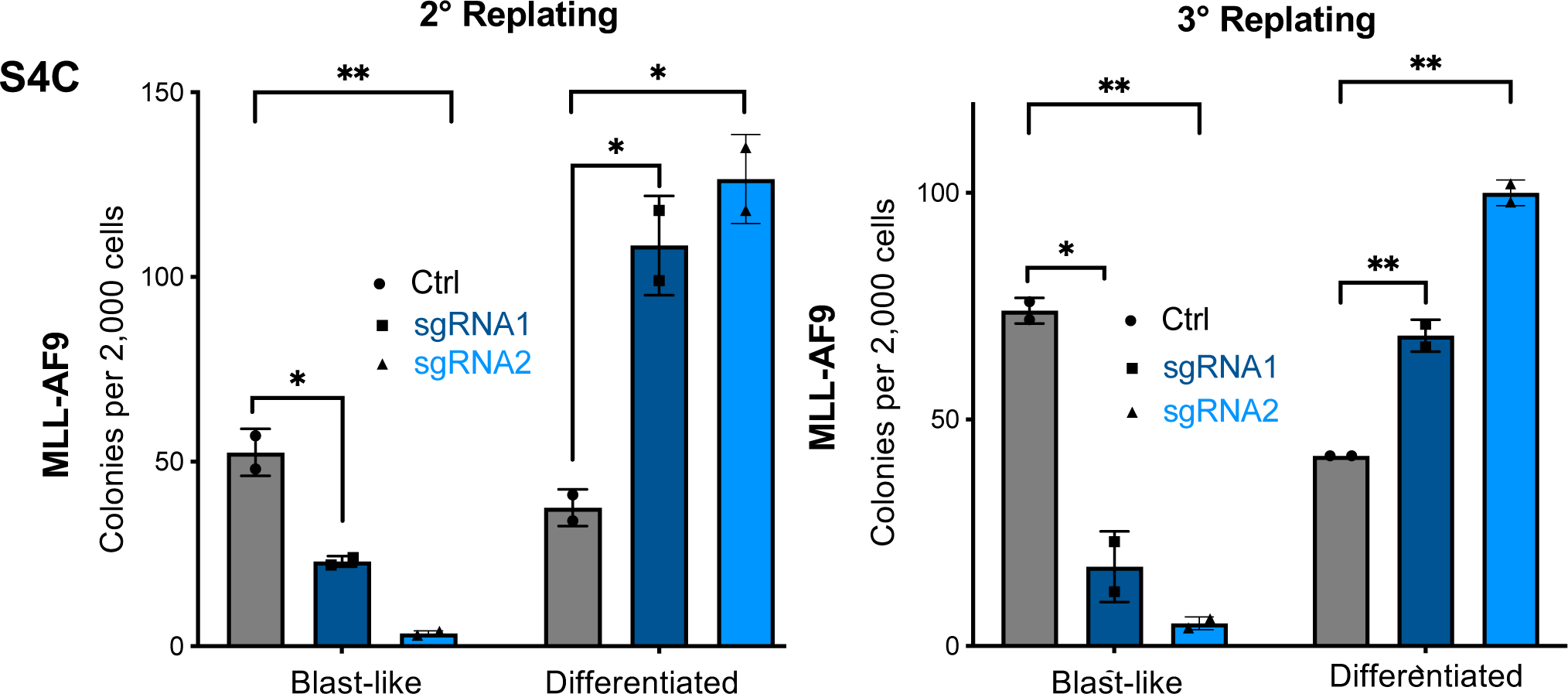
Colony forming unit (CFU) assay in murine MLL-AF9 leukemia cells showing second (left panel) and third (right panel) replating. Gray bars indicate the colonies from cells expressing Intron sgRNA. Blue bars indicate colonies from cells expressing sgRNAs targeting Sgf29. n=3); *=P<0.05; **=P<0.005.

**Fig. S4D:**
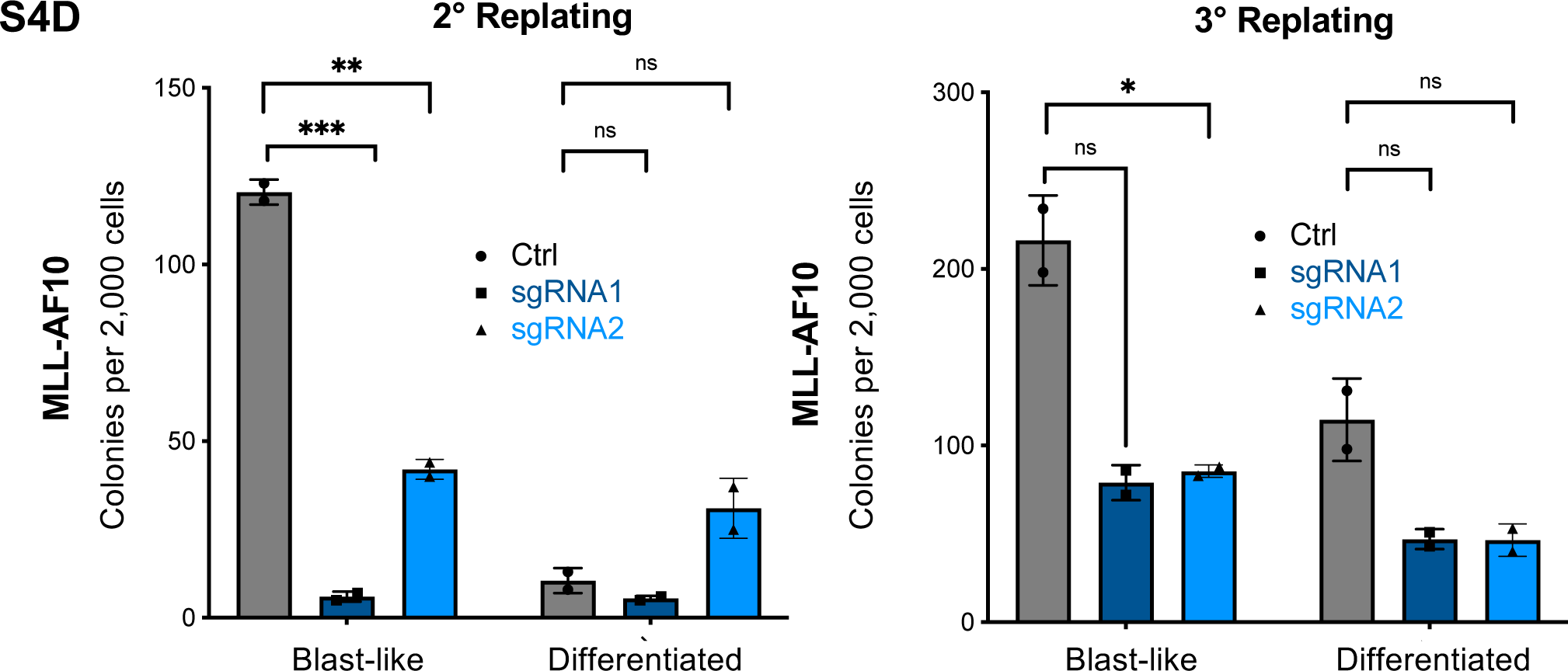
Colony forming unit (CFU) assay in murine MLL-AF10 transformed leukemia. The left panel indicates 2^nd^ week and the right panel indicates 3^rd^ week of the assay. Gray bars indicate the colonies from cells expressing Intron sgRNA. Blue bars indicate colonies from cells expressing sgRNAs targeting Sgf29. n=3); ns=non-significant; *=P<0.05;

**Fig. S4E:**
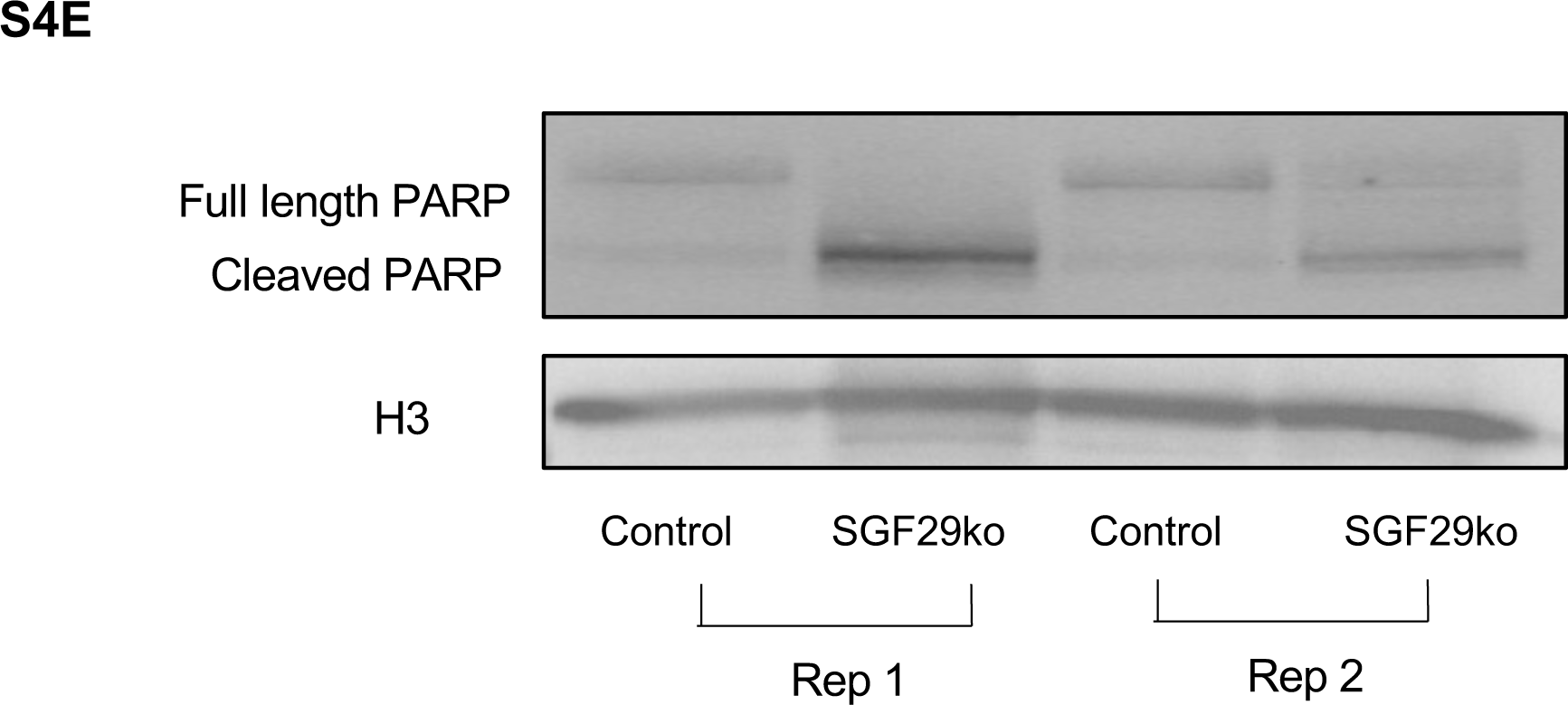
Effect of Sgf29 deletion on apoptosis in murine MLL-AF9 transformed leukemia. Immnoblot for PARP protein cleavage showing apoptosis in murine leukemia expressing Intron or Sgf29 targeting sgRNA. Two representative replicates are shown.

**Fig. S5A:**
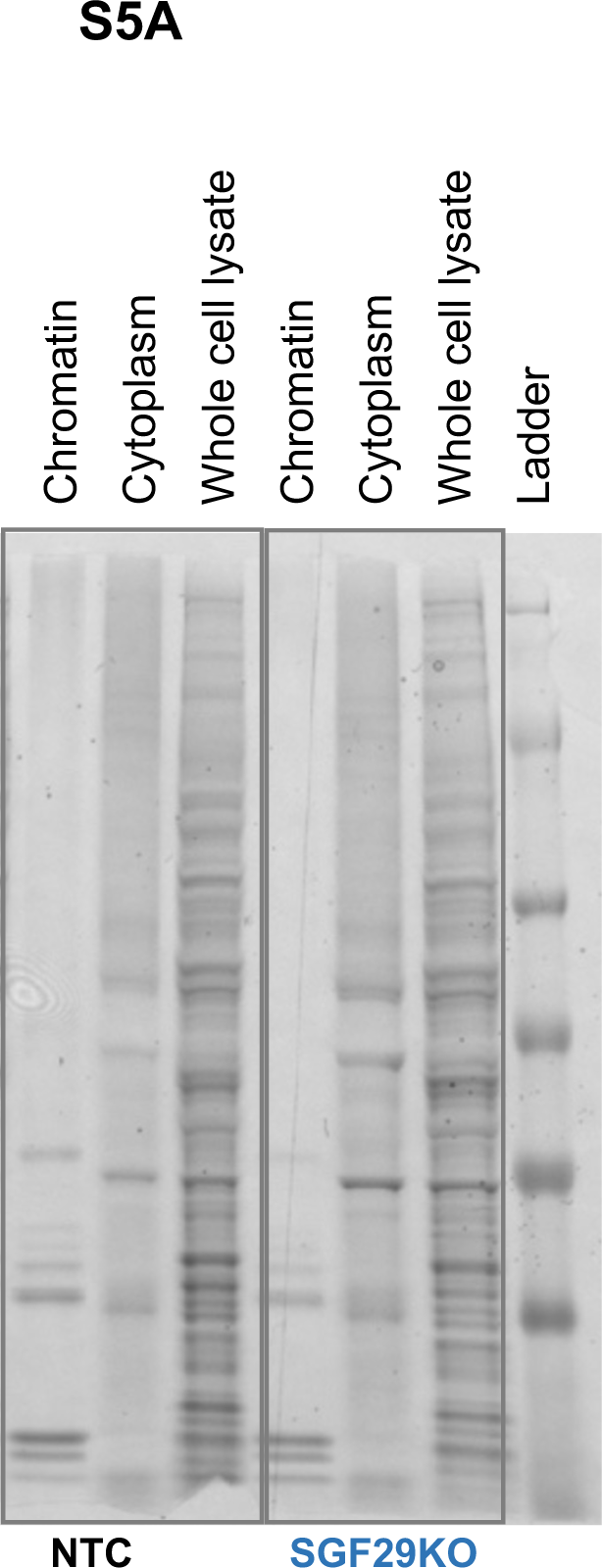
SDS-PAGE showing whole cell extract, cytoplasmic and ChEP-enriched fractions for NTC vs SGF29 knockout.

**Fig. S5B:**
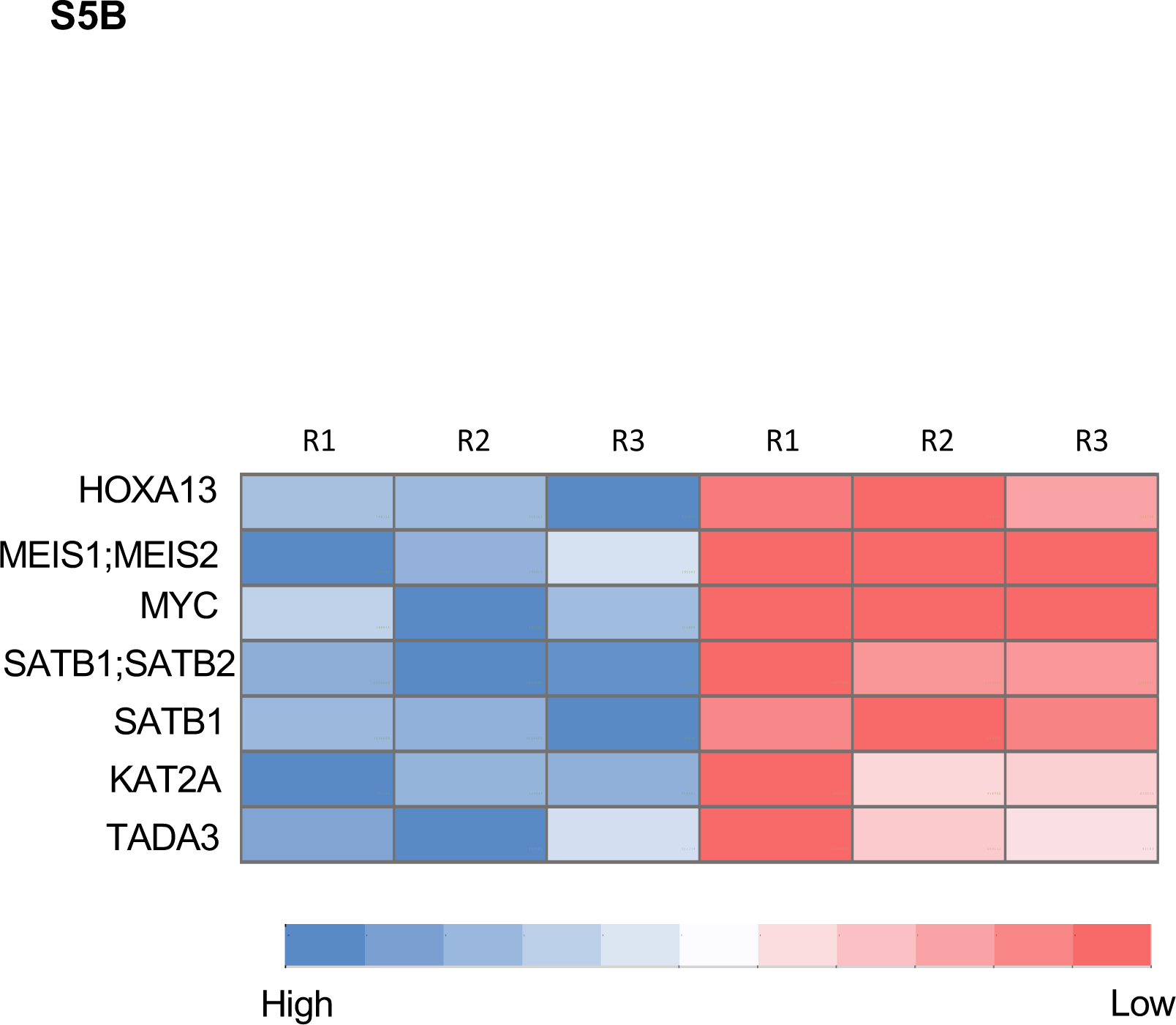
Heatmap of intensity changes in peptides for proteins evicted from the chromatin fraction of UB3 cells after SGF29 deletion in ChEP assay. R1, r2 and R3 are three independent replicates.

**Fig. S6A:**
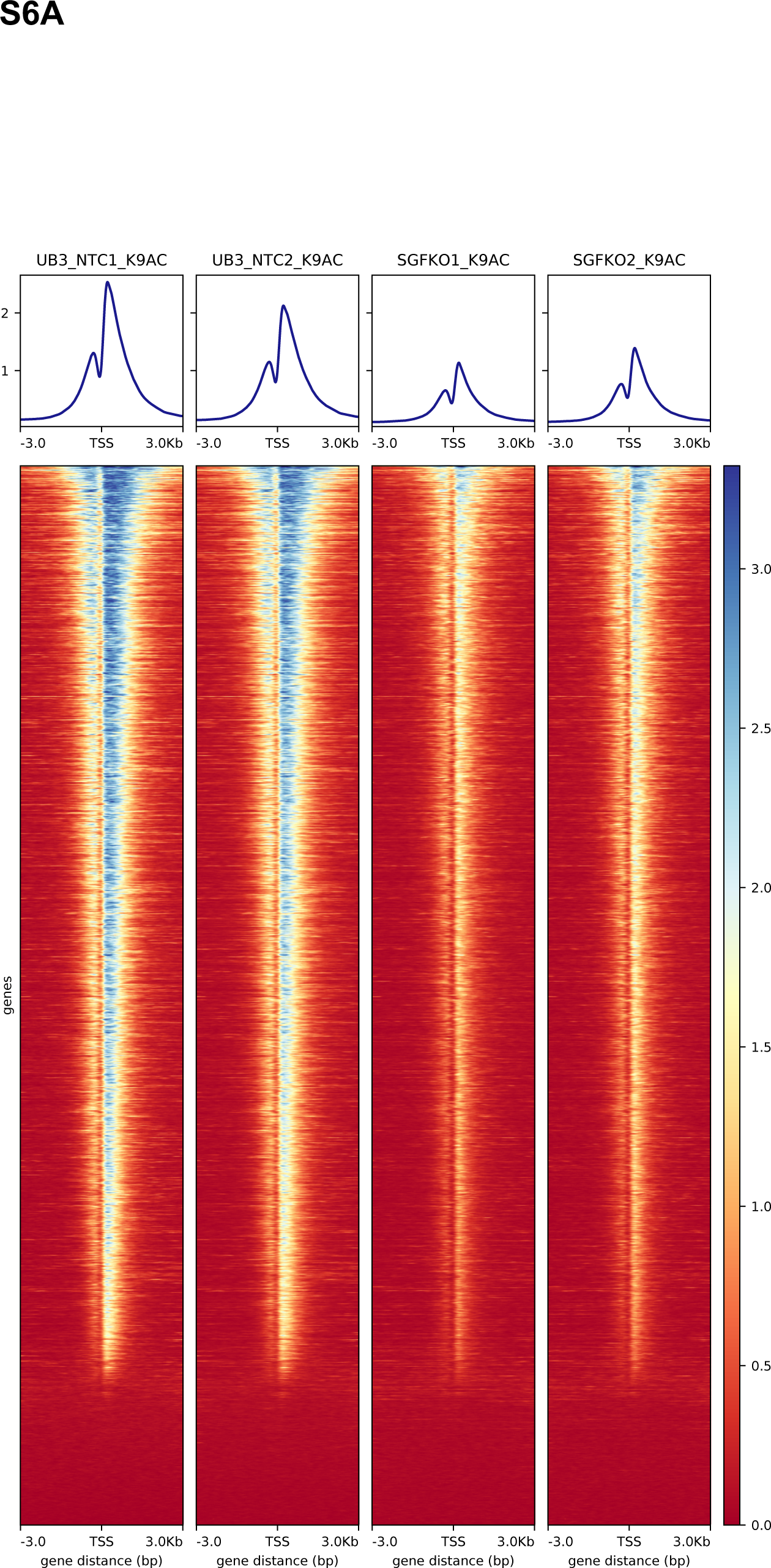
Heatmaps showing histone H3K9 acetylation (H3K9Ac) peaks centered around transcription start site (TSS) in UB3 cells transduced with non-targeting control (NTC) or SGF29 targeting sgRNA.

**Fig. S6B:** Volcano plot indicating changes in H3K9 acetylation across the genome in UB3 cells after deletion of SGF29. Blue dots are the loci with decreased and red are the loci with increased acetylation.

**Fig. S7A:**
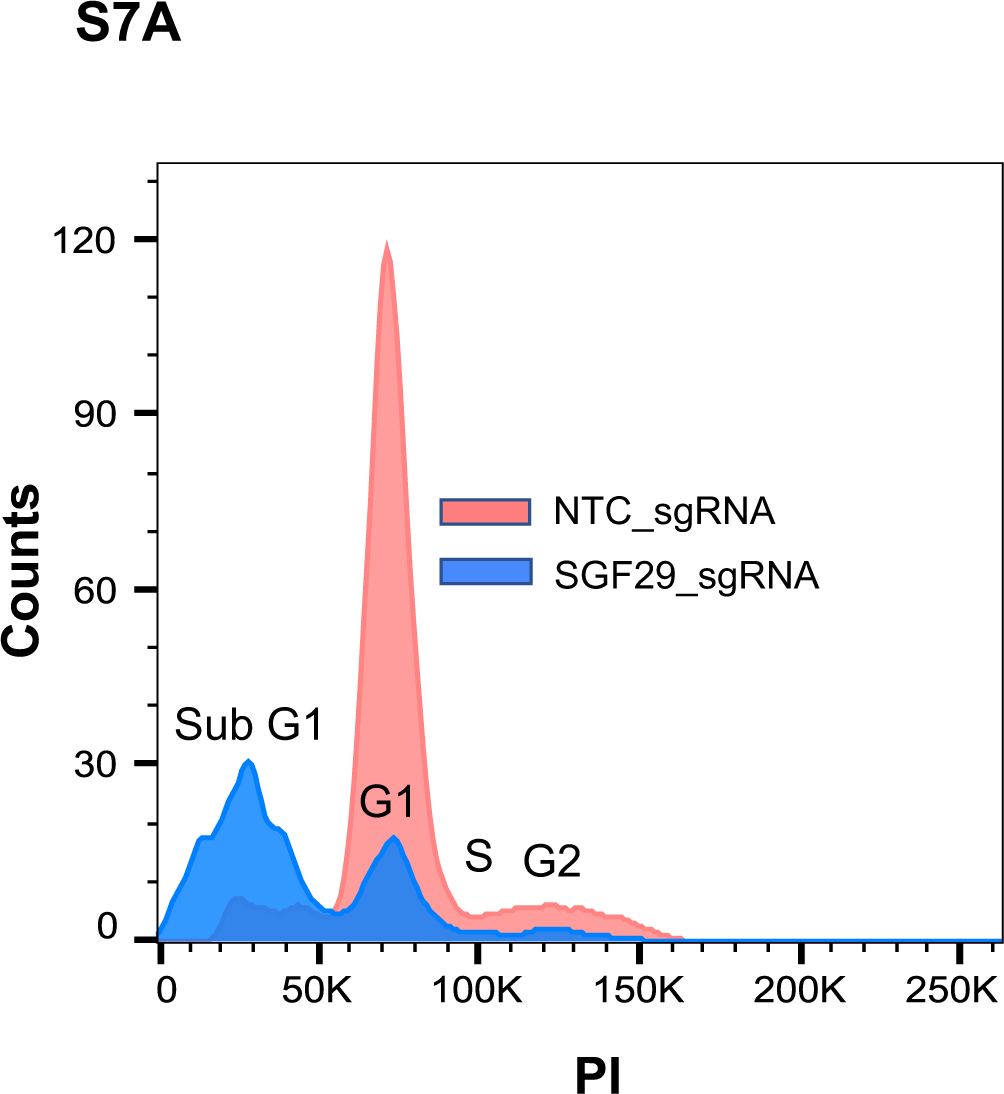
cell cycle analysis in MLL-AF10 PDX cells after deletion of SGF29. A representative sample showing changes in different phases of cell cycle upon loss of SGF29 is shown here. Orange represents cells expressing non-targeting control (NTC) sgRNA whereas blue indicates the cells with SGF29 ko.

**Fig. S8A:**
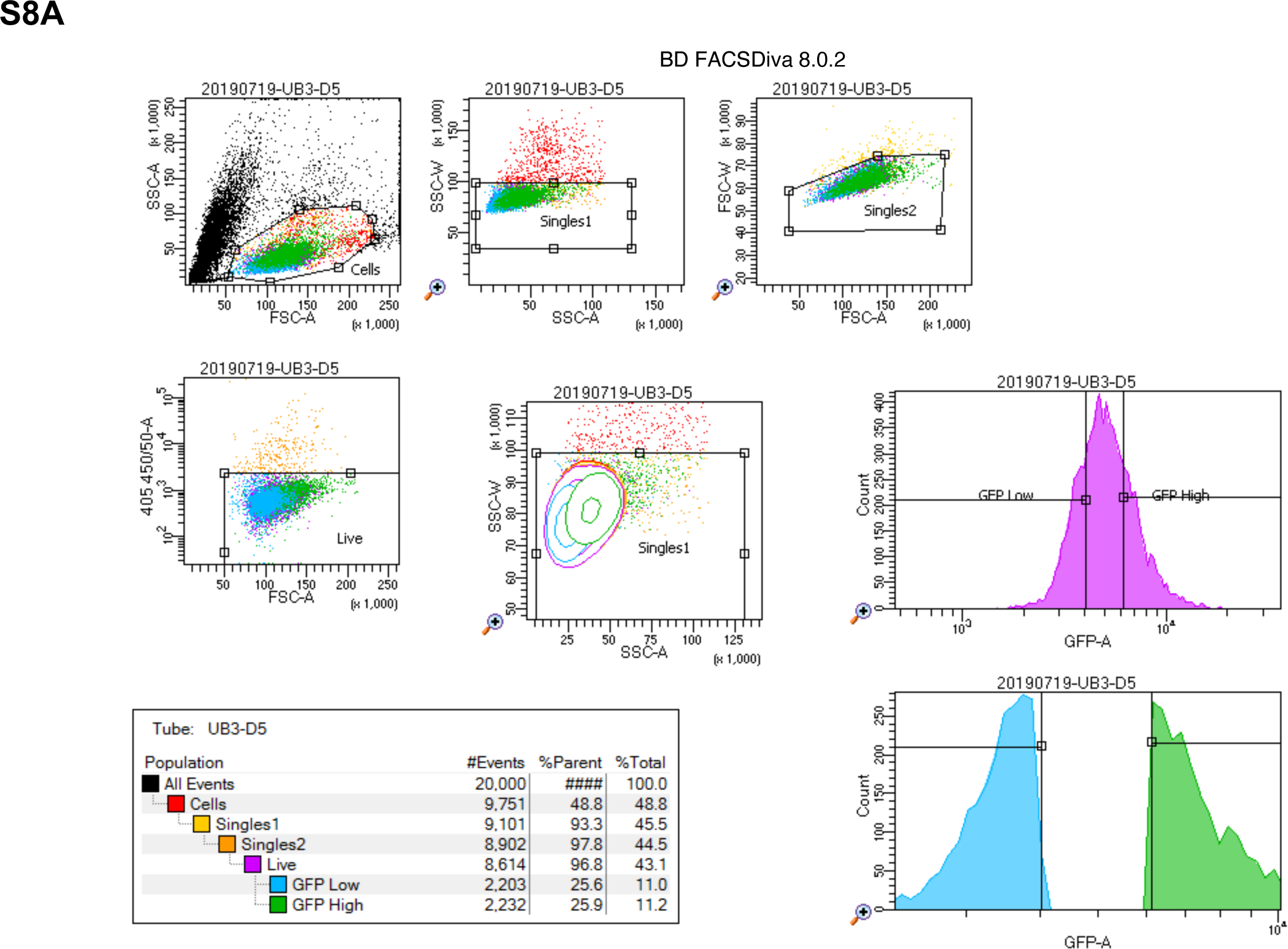
Schematic of FACS gating, A representative example of gating used for sorting UB3 high- and low- GFP fractions.

**Fig. S8B:**
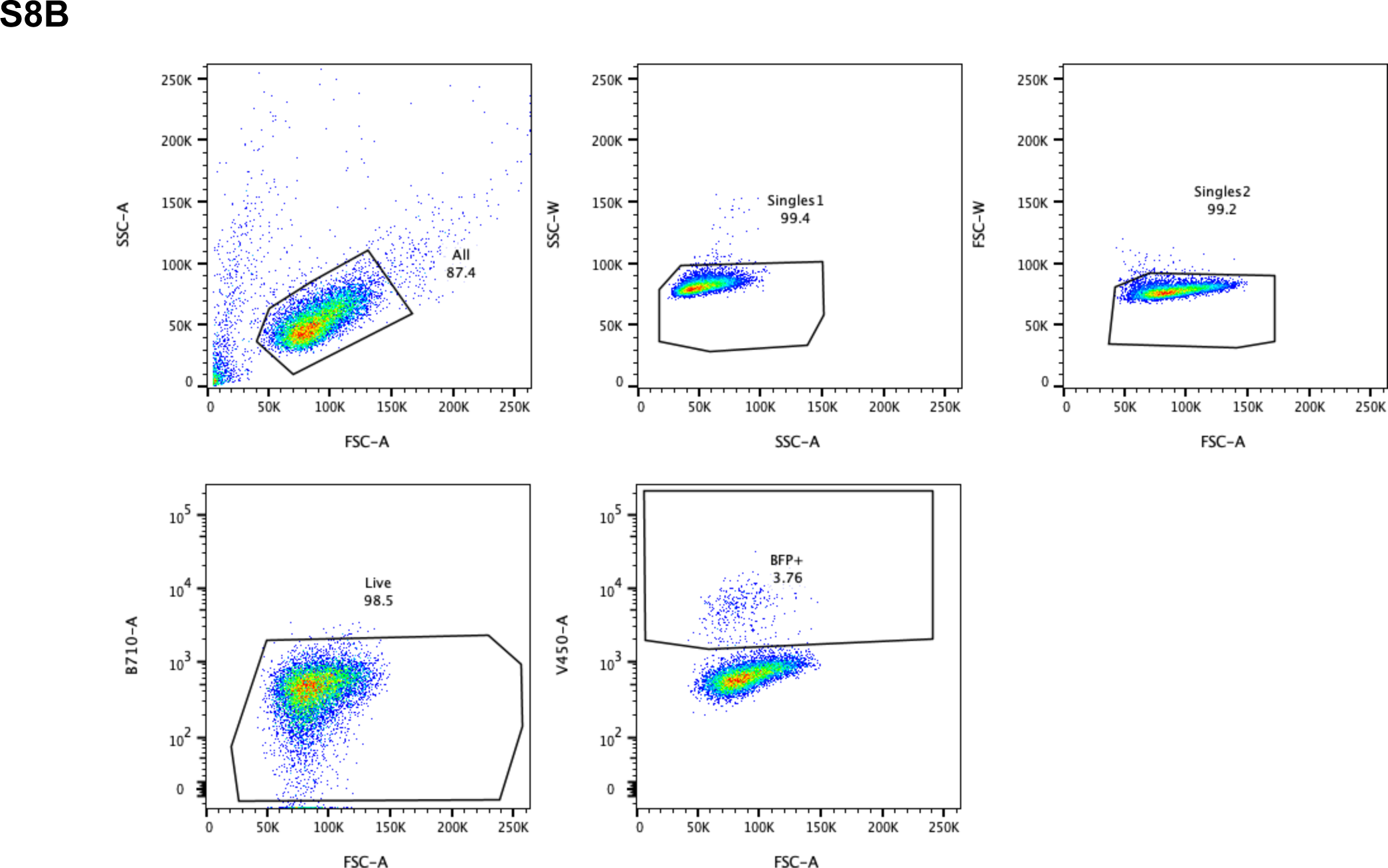
Gating strategy to analyze percentages of BFP positive cells expressing sgRNAs in BFP-puro.

## SUPPLEMENTARY METHODS

### Epigenetic sgRNA library design

Our custom sgRNA library was designed based on a manually curated list of known epigenetic regulators, assigned to the following categories: regulators of chromatin reading, chromatin remodeling, DNA and RNA modification, histone acetylation and methylation, transcriptional elongation and other histone modifications. The gene list was then used directly as the input for the Broad Institute Genetic Perturbation Platform (GPP) sgRNA designer tool ^210–213^ to obtain 5 sgRNAs targeting early constitutive exons. To design sgRNAs targeting functional protein domains, we retrieved the coding sequences for ENSEMBL-annotated protein domains (GRCh37) using a custom script (cds_domain_finder.py). Non-targeting control sgRNAs were included in the design and represented ∼5% of the total library size. The library sequences were synthesized on an array platform (Genscript, NJ) and then cloned into the lenti-guide Puro vector (Addgene: #52963, a kind gift from Dr. Feng Zhang, MIT) with the Gibson Assembly kit (New England Biolabs, NJ). The library plasmid was amplified using ElectroMAX™ Stbl4™ Competent Cells (Invitrogen, cat.11635018). The number of bacterial colonies was kept >500 times the number of sgRNA sequences in the library to maintain representation. To ensure the representation and identity of sgRNAs in the amplified pooled lentiviral plasmids, deep sequencing was performed on a HiSeq instrument (Illumina) and fastq files were analyzed using CRISPREssoCount ^214^. The sgRNA sequences in these libraries are deposited as indicated in the data availability section.

### Epigenetic CRISPR screen

The pooled epigenetics CRISPR library virus was produced using HEK293T cells transfected with Polyethylimine (PEI), pMD2.G, and psPAX2 as previously described ^215^. 30 million UB3 cells were transduced with the pooled epigenetics library lentivirus in RPMI medium supplemented with 10% fetal bovine serum, antibiotics, and 8 μg/ml polybrene. The medium was changed 24 h after transduction to remove polybrene and cells were plated in fresh culture medium. 48 h after transduction, puromycin was added at a concentration of 1 μg/ml to select for cells transduced with the sgRNA library and then removed after 72 h. The virus titer was measured by infection of cells with serially diluted virus. To ensure transduction of a single sgRNA per cell, the multiplicity of infection (MOI) was set to 0.3 ∼ 0.4. Adequate representation of sgRNAs during the screen was ensured by keeping >1000x cells in culture relative to the library size. 10 million cells per replicate were harvested 3 days after puromycin removal for an initial time point (T0). Cells were FACS-sorted 5 days later to collect the upper and lower quartiles on the basis of GFP-expression using the FACSAria II Instrument (BD Biosciences) at the Sanford Burnham Prebys Flow Cytometry Core. Genomic DNA extraction from the T0 and GFP -low and -high cells was performed using the Zymo QuickDNA™ Midiprep Plus Kit (Zymo Research, Cat. D4068), according to the manufacturer’s instructions. A two-step PCR-amplification for sequencing library preparation was conducted with TaKaRa Ex Taq™ Polymerase (TaKara, Cat. RR001) and custom primers to achieve adequate sequencing coverage in 1x75bp single-end reads, following published guidelines ^214^. Barcoded libraries were pooled and sequenced using an Illumina Hiseq 500.

### CRISPR competition experiments

To assess the candidate hits in independent assays, two top ranked individual sgRNAs from the sgRNA library were cloned for CSNK2A1, JADE3, AFF2, CCDC101, CSNK2B, DOT1L, ENY2, KAT7, KMT2A, MLLT1, and for non-targeting controls in the pKLV2-U6gRNA5(BbsI)- PGKpuro2ABFP-W vector (a gift from Kosuke Yusa, Addgene plasmid # 67974) ^218^ (sgRNA sequences provided in Table 2). Lentiviral supernatants were made from these constructs in a 96-well arrayed format. 10,000 UB3 cells were plated in a retronectin (Takara, #T202) coated flat bottom 96-well plate and transduced with the sgRNA viral supernatants by spinfection in polybrene-supplemented medium at an MOI of ∼0.5. Cells were maintained in culture in 96-well plate format and assayed for proliferation every 3 days. A sample >10% of the culture volume was stained with SytoxRed (1:1000) (Thermo Fisher, # S34859) in PBS and monitored for the percentage of blue fluorescent protein (BFP) expressing cells progressively up to 21 days using high-throughput flow-cytometry ^219^.

### Cell culture

Human leukemia cell lines U937 (Daniel Tenen, Beth Israel Deaconess Medical Center) and MOLM13 (ACC-554, DSMZ), were cultured in RPMI-1640 medium supplemented with 2 mM L-glutamine and sodium pyruvate, 10% Fetal bovine serum (FBS from… and 50 U/ml Penicillin/Streptomycin (Thermo Fisher Scientific, Carlsbad, CA), 2 mM L-glutamine and incubated in 5% CO_2_ at 37°C. Murine leukemia cells were cultured in DMEM medium supplemented with 2 mM L-glutamine, 15% FBS and 50U/ml Penicillin/Streptomycin, in the presence of following cytokines: 10 ng/ml murine interleukin 6 (mIL-6), 6 ng/ml murine interleukin 3 (mIL3) and 20 ng/ml murine stem cell factor (mSCF) (all from Peprotech, Rocky Hill, NJ), and incubated at 5% CO_2_ and 37°C. HEK293T cells were cultured in DMEM medium supplemented with 2 mM L-glutamine and sodium pyruvate, 10% FBS and 50 U/ml Penicillin/Streptomycin, and incubated in 5% CO_2_ at 37°C. MLL-AF10 patient derived xenograft (PDX) cells were cultured in IMDM medium supplemented with 20% BIT9500 (STEMCELL Technologies), human cytokines and StemRegenin 1 44 (SR1) and UM171 as described earlier(11).

### Generation of murine reporter leukemia

For mouse experiments, inducible MLL-AF9 murine leukemias were generated in lineage-depleted bone marrow cells of mice with a *GFP-Meis1* (12) or *CAG-Cas9* (13) background. Hematopoietic stem and progenitor cells (HSPCs) were isolated from bone marrow of C57Bl/6 mice using EasySep lineage depletion (STEMCELL Technologies) kit. HSPCs were cultured overnight in DMEM medium supplemented with 15% FBS and mSCF, mIL6 and mIL3 in a 37 °C incubator with 5% CO_2_. The following day, the cells were transduced with viral supernatants containing the MSCV-MLL-AF9-neo and the MSCV-rTTA-2A-BFP constructs by spinfection at 2500 rpm at 30°C for 90 min. 72 hr post-transduction, BFP+ve cells were sorted using the FACSAria II (BD Biosciences) flow cytometer (11). *In vitro* transformed BFP+ cells were then injected into sub-lethally irradiated C57Bl/6 recipient mice from the SBP Animal vivarium to generate primary leukemias.

### Small-molecule library composition and design

265 small-molecule inhibitors targeting diverse classes of epigenetic regulators were manually curated and purchased from various commercial sources by the Conrad Prebys Center for Chemical Genomics (Table S1).

### Flow-cytometry-based small-molecule screen

Compounds were dispensed with the Labcyte Echo 555 (Labcyte Inc., Sunnyvale, CA) acoustic droplet ejection dispenser to a final concentration of 0.11 µM and 10 µM, in technical triplicates. Human UB3 cells were then seeded in 384-well plates (PerkinElmer Health Sciences, Inc.) with a MicroClime® lid (Labcyte Inc.) at a density of 2,000 cells per well in 45 μl of culture medium using the Multidrop™ Combi Reagent Dispenser (ThermoFisher Scientific). Cells were incubated for 5 days at 37 °C, 5% CO_2_ and 95% humidity, and 5 μl of Sytox Red viability stain was then added to the culture volume using the Multidrop™ Combi Reagent Dispenser. High throughput flow cytometry was performed in the HTS unit of the Fortessa 14-color (BD) at the SBP Flow Cytometry Core. Flow cytometry data was analyzed with Flowjo version 10.0.4 (Flowjo LLC).

### Quantitative RT-PCR

The mRNA levels were measured by real time quantitative PCR (RT-qPCR) using standard protocols. RNA was extracted using the RNA extraction kit (Qiagen, Hilden, Germany) and was used to synthesize cDNA with the Protoscipt® II First Strand cDNA Synthesis Kit (New England Biolabs Inc., Beverly, MA). For PCR-based amplification, cDNA and primers were added to the SYBR Green PCR Master mix (ThermoFisher, Carlsbad, CA). Expression levels for *HOXA7,HOXA9, HOXA10, MEIS1, GAPDH and HPRT* were determined with the oligo sequences listed in Table S4 and measured on the Stratagene MX3000P (Agilent Technologies). Relative quantity of mRNA was determined by the ΔΔCT method as previously described ^150^ using HPRT as the internal reference.

### Colony formation assays

For murine leukemia colony forming cell (CFC) assays, CALM-AF10, MLL-AF9, and MLL-AF10 lines were transduced with retroviral particles encoding 2 sgRNAs targeting Sgf29 exon sequences and one sgRNA targeting an intronic sequence of Sgf29 as a control (sequences provided in Table S4), cloned in pMSCV-U6sgRNA(BbsI)-PGKpuro2ABFP (a gift from Sarah Teichmann, Addgene #102796). Cells were transduced by spinfection as described above and cultured for 48 hours in the 37 °C incubator with 5% CO_2_. Puromycin (2.5 μg/ml) was then added to the cultures for 48 hours to select for puromycin resistant cells. 1000 cells were plated in 35 mm dish, in duplicates, in 1mL Methylcellulose-based MethoCult M3234 medium (STEMCELL Technologies) supplemented with 10 ng/ml mIL6, 6 ng/ml mIL3 and 20 ng/ml mSCF (Peprotech) and colonies were scored on day 7. After scoring, colonies were washed with PBS, pooled, and counted for subsequent replating. A sample was taken for cell morphology assessment, performed by spinning 100,000 cells resuspended in 150 μl PBS onto glass slides, using the Cytospin 4 cytocentrifuge (Thermo Scientific) followed by Wright-Giemsa staining. Cells were replated every week for 3 weeks at the same cell number to test secondary and tertiary replating potential. For colony assays with Lin- Sca+ c-Kit+ (LSK) murine cells, bone marrow cells from Cas9 transgenic mice (13) were lineage-depleted using the EasySep lineage depletion kit (STEMCELL Technologies) to obtain a lineage negative (Lin-) population. Lin- cells were then stained with the following antibodies: FITC- conjugated Streptavidin (eBioscience), Pacific Blue-conjugated Sca-1 (Biolegend, San Diego, CA), and Alexa647-conjugated cKit (Biolegend, San Diego, CA) and sorted for the Lin- Sca-1+ cKit+ (LSK) population using the FACSAria II cytometer (BD Biosciences). Cells were kept in culture overnight and transduced the following day with retroviral particles with sgRNAs targeting Sgf29 and an intron control, as described above. The cells were counted and plated for colony formation in M3434 methylcellulose-based media (STEMCELL Technologies) at a concentration of 10,000 cells per ml, as described above. Colonies were scored on day 10.

### Dye-dilution experiments

To trace cell proliferation by flow cytometry, MOLM13 and U937 cells were transduced by spinfection with lentiviral particles containing sgRNAs targeting SGF29 or non-targeting controls (refer to Table S4), cloned in lentiGuide-Puro vector (a gift from Feng Zhang, Addgene #52963). Then, cells were incubated overnight and treated with 1 μg/ml puromycin for 3 days. Upon selection, the cells were washed and stained with the CellTrace™ Violet dye (ThermoFisher Scientific), according to the manufacturer’s instructions. Cells were maintained in culture in the dark and a sample (>10% of culture volume) was assayed every day for 6 days by flow cytometry.

### Viral preparation, transduction and selection

Lentivirus was made from the CROP-seq pooled DNA by transfection of the CROP-seq pool together with Polyethylimine (PEI), pMD2.G, and psPAX2 in HEK293T cells, as described previously ^215^. Viral supernatants were collected at 48 and 72 hours post-transfection, and then filtered (0.45 μm) and pooled. Viral supernatants were then concentrated by centrifugation (20,000 rpm, 2 hrs, 4° C) and used for transduction of UB3 cells by incubation with 0.8 μg/μl of polybrene overnight. Medium was changed after overnight incubation. At 48h post-transduction, puromycin (1μg/μl) was added to cell suspensions for 3 days of selection. Cells were pelleted for sequencing 3 days after removal of puromycin.

### Single cell sequencing

Two independently transduced samples were sequenced for the CROP-Seq experiment. Cell concentration and viability was assessed using a hemocytometer and viability for all samples was > 80%. A total of 18,000 cells per sample were loaded on a Chromium Single Cell Instrument (10x Genomics, Pleasanton, CA). RNAseq libraries were prepared using the Chromium Single Cell 3′ v3.1 Library, Gel Beads & Mutiplex Kit (10x Genomics). CDNA was PCR-amplified and purified using SPRIselect Reagent Kit (Beckman Coulter). Sequencing was performed on the Illumina NovaSeq using the S4 200 kit and PE100 run configuration at the University of California San Diego Center for Epigenomics. Cell Ranger Single-Cell Software Suite (v4.0.0; 10x Genomics) was used for sample demultiplexing, alignment, filtering, and UMI counting.

### CROP-seq data analysis

sgRNAs were detected in the fastq files via a custom script, which required a perfect match for either the 10bp upstream or 10bp downstream of the sgRNA, as well as the sgRNA sequence with at most 1 mismatch. If a partial sgRNA was located at the end of a sequence but at least 10bp matched a unique sgRNA (along with the correct neighboring sequence), the read was counted as well (CropSeq_sgRNA_from_ fastq.cpp).

10X fastq reads were trimmed using trim-galore (v0.6.6, flags -a AAAAAAAAAAAA --length 10) which relies on cutadapt (v2.5). Reads were aligned to the genome (hgGRCh38) using STAR (v2.7.3a). All multimapping reads were removed (samtools view -b -q 254), and reads were deduplicated by UMI using custom code (CropSeq_10x_Remove_Duplicates.cpp). Reads were then mapped to genes using bedtools v2.29.2. Reads spanning exon boundaries were split (-split flag used), and all reads were first mapped to exons (flags -f 0.95 -c 7 -o distinct) and then mapped to genes (-f 0.95 -c 4 -o distinct), such that a read mapping to overlapping genes is preferentially assigned to the gene in which it overlaps an exon. Cells with fewer than 10^3.5 (3162) UMIs or fewer than 200 expressed genes, or whose barcodes were not on the 10x barcode whitelist were removed. Cells with more than one gRNA were also removed. Finally, genes present in fewer than 3 cells were not included in downstream analyses. General processing and evaluation of sgRNA target gene expression, was performed in R (CROPSeq_HOX.R). All plots in Figure 24 were generated in R (CROPSeq_HOX.R). UMAP and ridgeplots were generated using the Seurat package. The dendrogram was generated using the R function dist(method = "euclidean") and hclust(method = "ward.D2"). The heatmap was generated using the pheatmap package.

### RNA-sequencing

U937-MEIS1-GFP and MOLM13 cells were plated at a density of 1 million cells per ml in T75 tissue culture flasks and transduced in triplicates with sgRNAs for non-targeting controls or SGF29. Cells were selected with puromycin as described above. 2 million cells were harvested at 7 days post-selection from every replicate and RNA was extracted using the RNeasy Mini kit (Qiagen, Hilden, Germany). RNA-seq Libraries were prepared using NEBNext Ultra II Directional RNA Library Prep Kit (New England Biolabs, Ipswich, MA) for Illumina as per the manufacturer’s protocol. For each sample, 900 ng of total RNA was amplified in 10 cycles of PCR amplification. Sequencing was performed to obtain 1x75bp reads on the Illumina NextSeq500 by the Genomics core at SBP Medical Discovery Institute. Next-generation sequencing data were demultiplexed and processed with the Illumina Basespace RNA-Seq Differential Expression Analysis workflow.

### RNA-sequencing data analysis

Raw reads were preprocessed by trimming Illumina Truseq adapters, polyA, and polyT sequences using cutadapt v2.3^226^ with parameters “cutadapt -j 4 -m 20 --interleaved -a AGATCGGAAGAGCACACGTCTGAACTCCAGTCAC -A AGATCGGAAGAGCGTCGTGTAGGGAAAGAGTGT Fastq1 Fastq2 | cutadapt --interleaved -j 4 -m 20 -a "A{100}" -A "A{100}" - | cutadapt -j 4 -m 20 -a "T{100}" -A "T{100}" -”. Trimmed reads were subsequently aligned to human genome version hg38 using STAR aligner v2.7.0d_0221 ^227^ with parameters according to ENCODE long RNA-seq pipeline (https://github.com/ENCODE-DCC/long-rna-seq-pipeline). Gene expression levels were quantified using RSEM v1.3.1 ^228^. Ensembl v84 gene annotations were used for the alignment and quantification steps. RNA-seq sequence, alignment, and quantification qualities were assessed using FastQC v0.11.5 (https://www.bioinformatics.babraham.ac.uk/projects/fastqc/) and MultiQC v1.8 ^229^. Lowly expressed genes were filtered out by retaining genes with estimated counts (from RSEM) ≥ number of samples times 5. Filtered estimated read counts from RSEM were used for differential expression comparisons using the Wald test implemented in the R Bioconductor package DESeq2 v1.22.2 based on generalized linear model and negative binomial distribution ^230^(Love, Huber et al. 2014). Genes with Benjamini-Hochberg corrected p-value < 0.05 and fold change ≥ 2.0 or ≤ 2.0 were selected as differentially expressed genes. Pathway analyses of differential expression comparisons were performed using Ingenuity Pathway Analysis (Qiagen, Redwood City, USA).

### Chromatin immunoprecipitation and sequencing (ChIP-Seq)

UB3 cells expressing Cas9 were transduced with sgRNAs for a non-targeting control or SGF29, as described for the dye-dilution experiment. Chromatin immunoprecipitation was performed as previously described ^150^. Cells were fixed with 1% formaldehyde and chromatin was sheared using a Bioruptor™ (Diagenode Inc, NJ) in 15 cycles (each one in high-setting for 30 s on and 30 s off), at 4°C. Chromatin was immunoprecipitated using antibodies for normal rabbit IgG (Cell Signaling Technology, Cat. No, 2729) and Histone H3K9ac (Abcam, ab4441). For SGF29 binding studies, U937 cells transduced with 3x Flag tagged SGF29 wt or D196R mutant viral supernatants were fixed with 1% formaldehyde for 15 minutes and chromatin was sheared as described earlier. Chromatin was immunoprecipitated using Anti-FLAG M2 affinity beads (Sigma Aldrich, #A2220). Library preparation on eluted DNA was performed using the NEBNext Ultra II DNA library prep kit for Illumina (E7645S and E7600S) as per the manufacturer’s protocol. DNA libraries were sequenced in paired-end 150 bp reads in a NovaSeq600 sequencer (Illumina, San Diego, CA) by Novogene Corporation (Sacramento, CA).

### ChIP-seq processing methods

Raw reads were processed with Cutadapt v2.3 to remove low-quality bases and TruSeq adapter contamination using parameters “-j 12 -m 25 -O 5 -q 15 -a AGATCGGAAGAGCACACGTCTGAACTCCAGTCAC”. Trimmed reads were first mapped to human chrM to remove reads mapping to it from downstream analysis using Bowtie2 v2.2.5 and parameters “--local”. Reads not mapped to chrM were aligned to human genome version hg38 using Bowtie2 “--very-sensitive”. Unmapped, multimapper, and PCR duplicate reads were removed using sambamba v0.7.1 and parameters ‘sambamba view -h -t 8 -f bam -F "[XS] == null and not unmapped and not duplicate"’. Genome coverage bigwig files were generated using deepTools bamCoverage v3.4.3 and parameters “--binSize 10 -p 12 --normalizeUsing RPGC -- effectiveGenomeSize 2913022398”. Peak-calling was performed using MACS2 v2.2.71 and parameters “--broad-cutoff 0.0001 -q 0.0001 –broad”. K4me3 and K27ac data in Fig. 5 were obtained from publicly available deposited files at: ENA biosample ID SAMEA2396211and GEO accession number GSM4083812, respectively.

### (ChEP)

Five million U937 cells were transduced in 4 replicates with concentrated lentiviral supernatants containing a non-targeting control or sgRNAs targeting CSNK2A1 or SGF29 in the pKLV2- U6gRNA5(BbsI)-PGKpuro-2ABFP backbone (a gift from Kosuke Yusa, Addgene #67974; see Table 2 for sequences). 48 hours after transduction, cells were treated with 1 μg/ml puromycin for 3 days. The cultures were grown for 7 days and counted for chromatin enrichment for proteomics procedure ^231^. Briefly, 100 million cells per replicate were collected by centrifugation for 5 minutes at 600 xg at room temperature and washed with pre-warmed PBS. Cells were then resuspended in 1% formaldehyde in pre-warmed PBS and incubated for 10 min at 37°C in a rotator. Crosslinking was halted by adding glycine to a final concentration of 0.25M and incubating for 5 min at room temperature in a rotator. Cells were then washed with pre-warmed PBS and pellets were collected by centrifugation at room temperature for 5 min at 600 xg and stored at -80 °C. Pellets were thawed and resuspended in 1 ml cold cell lysis buffer (25 mM Tris, 0.1% Triton X- 100, 85 mM KCl, and protease inhibitors), transferred to 2 ml tubes, and homogenized carefully with a 200 μl pipette tip. The samples were then centrifuged for 2300 xg for 5 min at 4°C and the supernatants containing the cytoplasmic fraction were recovered and stored at -80°. The nuclei pellets were resuspended in 500 μl cell lysis buffer containing 200μl/ml RNAse A and were incubated for 15 min at 37°C. Samples were then kept on ice and centrifuged at 2,300 xg for 10 min at 4 °C. The supernatants were then discarded, and pellets were resuspended in 500 μl SDS buffer (50 mM Tris, 10 mM EDTA, 4% SDS, and protease inhibitors) with 200 μl pipette tips and incubated for 10 min at room temperature. Samples were then centrifuged at 16,000 xg for 30 min at room temperature and the supernatants were discarded. The pellets were washed by resuspending in 500 μl SDS buffer with a 200 μl tip, then adding 1.5 ml urea buffer (10 mM Tris, 1 mM EDTA, and 8 M urea) and mixing by inversion several times. The tubes were then spun down at 16,000 xg for 25 min at room temperature and supernatants were discarded. Urea was washed out by resuspending in 500 μl SDS buffer with a 200 μl tip, then adding 1.5 ml SDS buffer and mixing by inversion several times. The tubes were spun down at 16,000 xg for 25 min at room temperature and the pellets were covered with storage buffer (10 mM Tris, 1mM EDTA, 25 mM NaCl, 10% glycerol, and protease inhibitors) and flicked to dislodge the pellets. The pellets were transferred to Bioruptor® shearing tubes (Diagenode Inc, Denville, NJ) and sonicated in a Bioruptor® (Diagenode Inc., Denville, NJ) for 15 min in 15 cycles (each 30 s on, 30 s off on high-intensity setting). The samples were centrifuged at 16,000 xg for 30 min at 4 °C and the supernatants were transferred to a new tube and stored at -80 °C for mass spectrometry analysis.

### LC-MS/MS analysis

For the chromatin-enriched for proteomics samples, 30 μg of each sample were processed using the ProTiFi S-trap™ digestion system. Mass spectrometry data were acquired on an Orbitrap Exploris 480 (ThermoFisher, Bremen, Germany) instrument at the Cedars Sinai Proteomics and Metabolomics Core in the Advanced Clinical Biosystems Research Institute. Desalted peptides were separated on an Ultimate 3000 ultra-high-pressure chromatography system with a 60- min gradient. Peptides were separated on a gradient of 1% B organic phase for 2 minutes, 1-5% B for 0.5 minutes, 5-9% B for 3.5 minutes, 9-27% B for 39 minutes, and 27-44% B for the final 15 minutes on a C18 column (15 cm length, 300 µm diameter) at a flow rate of 9.5 nL/min. Source parameters included spray voltage at 3 kV, capillary temp of 300 °C, and RF funnel level of 40%. MS1 resolutions were set to 60,000 and AGC was set to “standard” with ion transmission of 100ms. Mass range of 400-1000 and AGC target value for fragment spectra of 300% were used. Peptide ions were fragmented at a normalized collision energy of 30%. Fragmented ions were detected across 50 DIA windows of 12 Da. MS2 resolutions were set to 15,000 with an ion transmission time of 25 ms. All data were acquired in profile mode using positive polarity. The data were searched using the DIA-NN tool against a human in silico digested sequence reference library.

### Data analysis for ChEP

Protein peak data (mz/rt) were analyzed using MetaboAnalyst 5.0. Missing values were imputed by replacing them with 1/5^th^ of the minimum positive values of their corresponding variables. No data filtering was applied. Important features were selected based on fold change analysis with a threshold of 2 and t-test and P. values below 0.05.

### Statistical Analysis

Flow cytometry data were analyzed using FlowJo v10.7.1 (FlowJo Software, Tree Star, Ashland, OR). All statistical analyses were performed using GraphPad Prism 9 Software (San Diego).

### Western and dot blotting

Whole cells were lysed in cold RIPA buffer containing the Halt™ protease inhibitor cocktail for 30 min on ice (ThermoFisher Scientific, Carlsbad, CA). Protein supernatant was collected after 30 min of 12,000 rpm centrifugation at 4°C and stored at -80°C until use. Protein concentrations from whole cell lysates were determined using the Pierce™ BCA protein assay kit (ThermoFisher Scientific). For the immunoblotting, the following antibodies were used: KAT2A (Abcam, ab217876), Flag M2 (Sigma-Aldrich, F1804), Vinculin (Sigma-Aldrich, V9131), H3 (Abcam, ab1791), and SGF29 (Abcam, ab204367). Protein samples (50µg total protein) were resolved by SDS-PAGE using 4-12% Bis-Tris Bolt™ gels (Invitrogen, NW04120BOX) and transferred to iBlot2™ nitrocellulose membranes (ThermoFisher Scientific, IB23002). The membranes were blocked with 5% non-fat milk in Tris-buffered Saline containing 0.01% tween-20 (0.01% TBST). The membranes were probed with species-specific goat anti-rabbit (MilliporeSigma, AP307P) or goat anti-mouse (ThermoFisher Scientific, 31446) HRP-conjugated secondary antibody and detected with enhanced chemiluminescence (ECL) detection kit (Thermo Fisher, Carlsbad, CA).

### Cell Cycle Analysis

500,000 MLL-AF10 PDX cells expressing NTC or SGf29 sgRNA were washed with cold PBS and fixed with chilled 70% ethanol. Fixed cells were incubated at -20°C for 1 hr. Cells were then centrifuged at 2000 rpm for 3 min. at 4°C. Pellets were resuspended in PBS containing RNAse A and Propidium Iodide (PI) and incubated at 37°C for 30 min. in dark and analysed using flow cytometry.

## REFERENCES

1. Dohner, H., Weisdorf, D. J. & Bloomfield, C. D. Acute Myeloid Leukemia. N. Engl. J. Med. 373, 1136–1152 (2015).

2. Papaemmanuil, E., Döhner, H. & Campbell, P. J. Genomic classification in acute myeloid leukemia. The New England journal of medicine vol. 375 900–901 (2016).

3. Abramovich, C., Pineault, N., Ohta, H. & Humphries, R. K. Hox genes: from leukemia to hematopoietic stem cell expansion. Ann. N. Y. Acad. Sci. 1044, 109–116 (2005).

4. Argiropoulos, B. & Humphries, R. K. Hox genes in hematopoiesis and leukemogenesis. Oncogene 26, 6766–6776 (2007).

5. Alharbi, R. A., Pettengell, R., Pandha, H. S. & Morgan, R. The role of HOX genes in normal hematopoiesis and acute leukemia. Leukemia 27, 1000–1008 (2013).

6. Collins, C. T. & Hess, J. L. Deregulation of the HOXA9/MEIS1 axis in acute leukemia. Curr. Opin. Hematol. 23, 354–361 (2016).

7. Spencer, D. H. et al. Epigenomic analysis of the HOX gene loci reveals mechanisms that may control canonical expression patterns in AML and normal hematopoietic cells. Leukemia 29, 1279–1289 (2015).

8. Wang, G. G., Cai, L., Pasillas, M. P. & Kamps, M. P. NUP98-NSD1 links H3K36 methylation to Hox-A gene activation and leukaemogenesis. Nat. Cell Biol. 9, 804– 812 (2007).

9. Caudell, D., Zhang, Z., Chung, Y. J. & Aplan, P. D. Expression of a CALM-AF10 fusion gene leads to Hoxa cluster overexpression and acute leukemia in transgenic mice. Cancer Res. 67, 8022–8031 (2007).

10. DiMartino, J. F. et al. The AF10 leucine zipper is required for leukemic transformation of myeloid progenitors by MLL-AF10. Blood 99, 3780–3785 (2002).

11. Thorsteinsdottir, U. et al. Overexpression of the myeloid leukemia-associated Hoxa9 gene in bone marrow cells induces stem cell expansion. Blood 99, 121–129 (2002).

12. Kroon, E. et al. Hoxa9 transforms primary bone marrow cells through specific collaboration with Meis1a but not Pbx1b. EMBO J. 17, 3714–3725 (1998).

13. Wong, P., Iwasaki, M., Somervaille, T. C., So, C. W. & Cleary, M. L. Meis1 is an essential and rate-limiting regulator of MLL leukemia stem cell potential. Genes Dev. 21, 2762–2774 (2007).

14. Xiang, P. et al. Elucidating the importance and regulation of key enhancers for human MEIS1 expression. Leukemia (2022) doi:10.1038/s41375-022-01602-4.

15. Barbosa, K. et al. Acute myeloid leukemia driven by the CALM-AF10 fusion gene is dependent on BMI1. Exp. Hematol. 74, 42–51 e3 (2019).

16. The role of CALM-AF10 gene fusion in acute leukemia. Leukemia vol. 22 678–685 (2008).

17. Daigle, S. R. et al. Selective killing of mixed lineage leukemia cells by a potent small-molecule DOT1L inhibitor. Cancer Cell 20, 53–65 (2011).

18. Erb, M. A. et al. Transcription control by the ENL YEATS domain in acute leukaemia. Nature 543, 270–274 (2017).

19. Wan, L. et al. ENL links histone acetylation to oncogenic gene expression in acute myeloid leukaemia. Nature 543, 265–269 (2017).

20. Shi, J. et al. Discovery of cancer drug targets by CRISPR-Cas9 screening of protein domains. Nat. Biotechnol. 33, 661–667 (2015).

21. Doench, J. G. et al. Optimized sgRNA design to maximize activity and minimize off- target effects of CRISPR-Cas9. Nat. Biotechnol. 34, 184–191 (2016).

22. Datlinger, P. et al. Pooled CRISPR screening with single-cell transcriptome readout. Nat. Methods 14, 297–301 (2017).

23. Okada, Y. et al. hDOT1L links histone methylation to leukemogenesis. Cell 121, 167– 178 (2005).

24. Mueller, D. et al. A role for the MLL fusion partner ENL in transcriptional elongation and chromatin modification. Blood 110, 4445–4454 (2007).

25. Luo, Z. et al. The super elongation complex family of RNA polymerase II elongation factors: gene target specificity and transcriptional output. Mol. Cell. Biol. 32, 2608– 2617 (2012).

26. Tsherniak, A. et al. Defining a Cancer Dependency Map. Cell 170, 564–576 e16 (2017).

27. Shimada, K., Bachman, J. A., Muhlich, J. L. & Mitchison, T. J. shinyDepMap, a tool to identify targetable cancer genes and their functional connections from Cancer Dependency Map data. Elife 10, (2021).

28. Subramanian, A. et al. Gene set enrichment analysis: a knowledge-based approach for interpreting genome-wide expression profiles. Proc. Natl. Acad. Sci. U. S. A. 102, 15545–15550 (2005).

29. Bian, C. et al. Sgf29 binds histone H3K4me2/3 and is required for SAGA complex recruitment and histone H3 acetylation. EMBO J. 30, 2829–2842 (2011).

30. Vermeulen, M. et al. Quantitative interaction proteomics and genome-wide profiling of epigenetic histone marks and their readers. Cell 142, 967–980 (2010).

31. Vosnakis, N. et al. Coactivators and general transcription factors have two distinct dynamic populations dependent on transcription. EMBO J. 36, 2710–2725 (2017).

32. Espinola-Lopez, J. M. & Tan, S. The Ada2/Ada3/Gcn5/Sgf29 histone acetyltransferase module. Biochim. Biophys. Acta Gene Regul. Mech. 1864, 194629 (2021).

33. Kustatscher, G., Wills, K. L. H., Furlan, C. & Rappsilber, J. Chromatin enrichment for proteomics. Nat. Protoc. 9, 2090–2099 (2014).

34. Goel, S., Bergholz, J. S. & Zhao, J. J. Targeting CDK4 and CDK6 in cancer. Nat. Rev. Cancer 22, 356–372 (2022).

35. Domingues, A. F. et al. Loss of Kat2a enhances transcriptional noise and depletes acute myeloid leukemia stem-like cells. Elife 9, (2020).

36. Liu, W.-H., Völse, K., Senft, D. & Jeremias, I. A reporter system for enriching CRISPR/Cas9 knockout cells in technically challenging settings like patient models. Sci. Rep. 11, 12649 (2021).

37. Rau, R. E. et al. DOT1L as a therapeutic target for the treatment of DNMT3A-mutant acute myeloid leukemia. Blood 128, 971–981 (2016).

38. Bernt, K. M. et al. MLL-rearranged leukemia is dependent on aberrant H3K79 methylation by DOT1L. Cancer Cell 20, 66–78 (2011).

39. Deshpande, A. J. et al. AF10 Regulates Progressive H3K79 Methylation and HOX Gene Expression in Diverse AML Subtypes. Cancer Cell (2014) doi:10.1016/j.ccell.2014.10.009.

40. Deshpande, A. J. et al. Leukemic transformation by the MLL-AF6 fusion oncogene requires the H3K79 methyltransferase Dot1l. Blood (2013) doi:10.1182/blood-2012-11-465120.

41. Sanders, S. L., Jennings, J., Canutescu, A., Link, A. J. & Weil, P. A. Proteomics of the eukaryotic transcription machinery: identification of proteins associated with components of yeast TFIID by multidimensional mass spectrometry. Mol. Cell. Biol. 22, 4723–4738 (2002).

42. Baker, S. P. & Grant, P. A. The SAGA continues: expanding the cellular role of a transcriptional co-activator complex. Oncogene 26, 5329–5340 (2007).

43. Rodríguez-Navarro, S. Insights into SAGA function during gene expression. EMBO reports vol. 10 843–850 Preprint at https://doi.org/10.1038/embor.2009.168 (2009).

44. Ringel, A. E., Cieniewicz, A. M., Taverna, S. D. & Wolberger, C. Nucleosome competition reveals processive acetylation by the SAGA HAT module. Proc. Natl. Acad. Sci. U. S. A. 112, E5461–70 (2015).

45. Balasubramanian, R., Pray-Grant, M. G., Selleck, W., Grant, P. A. & Tan, S. Role of the Ada2 and Ada3 transcriptional coactivators in histone acetylation. J. Biol. Chem. 277, 7989–7995 (2002).

46. Soffers, J. H. M. & Workman, J. L. The SAGA chromatin-modifying complex: the sum of its parts is greater than the whole. Genes Dev. 34, 1287–1303 (2020).

47. Tzelepis, K. et al. A CRISPR Dropout Screen Identifies Genetic Vulnerabilities and Therapeutic Targets in Acute Myeloid Leukemia. Cell Rep. 17, 1193–1205 (2016).

48. Han, X. & Chen, J. KAT2A affects tumor metabolic reprogramming in colon cancer progression through epigenetic activation of E2F1. Hum. Cell 35, 1140–1158 (2022).

49. Arede, L. et al. KAT2A complexes ATAC and SAGA play unique roles in cell maintenance and identity in hematopoiesis and leukemia. Blood Adv. 6, 165–180 (2022).

50. Kurabe, N. et al. Deregulated expression of a novel component of TFTC/STAGA histone acetyltransferase complexes, rat SGF29, in hepatocellular carcinoma: possible implication for the oncogenic potential of c-Myc. Oncogene 26, 5626–5634 (2007).

51. Long, L. et al. CRISPR screens unveil signal hubs for nutrient licensing of T cell immunity. Nature 600, 308–313 (2021).

52. Filippakopoulos, P. et al. Selective inhibition of BET bromodomains. Nature 468, 1067–1073 (2010).

53. Cipriano, A., Sbardella, G. & Ciulli, A. Targeting epigenetic reader domains by chemical biology. Curr. Opin. Chem. Biol. 57, 82–94 (2020).

54. Arrowsmith, C. H. & Schapira, M. Targeting non-bromodomain chromatin readers. Nat. Struct. Mol. Biol. 26, 863–869 (2019).

55. Mio, C., Bulotta, S., Russo, D. & Damante, G. Reading cancer: Chromatin readers as druggable targets for cancer treatment. Cancers (Basel) 11, 61 (2019).

56. Liu, Y. et al. Small-molecule inhibition of the acyl-lysine reader ENL as a strategy against acute myeloid leukemia. Cancer Discov. (2022) doi:10.1158/2159-8290.CD-21-1307.

57. Asiaban, J. N. et al. Cell-Based Ligand Discovery for the ENL YEATS Domain. ACS Chem. Biol. 15, 895–903 (2020).

58. Garnar-Wortzel, L., et al. Chemical inhibition of ENL/AF9 YEATS domains in acute leukemia. 2020.12.01.406694 (2020).

59. Chen, B.-R. et al. A JAK/STAT-mediated inflammatory signaling cascade drives oncogenesis in AF10-rearranged AML. Blood 137, 3403–3415 (2021).

60. Li, W. et al. Quality control, modeling, and visualization of CRISPR screens with MAGeCK-VISPR. Genome Biol. 16, 281 (2015).

61. Wang, B. et al. Integrative analysis of pooled CRISPR genetic screens using MAGeCKFlute. Nat. Protoc. 14, 756–780 (2019).

## REFERENCES

1. Doench JG, Fusi N, Sullender M, Hegde M, Vaimberg EW, Donovan KF, et al. Optimized sgRNA design to maximize activity and minimize off-target effects of CRISPR-Cas9. Nat Biotechnol. Nature Publishing Group; 2016;34:184–91.

2. Sanson KR, Hanna RE, Hegde M, Donovan KF, Strand C, Sullender ME, et al. Optimized libraries for CRISPR-Cas9 genetic screens with multiple modalities. Nat Commun. Nature Publishing Group; 2018;9:1–15.

3. Kim HK, Min S, Song M, Jung S, Choi JW, Kim Y, et al. Deep learning improves prediction of CRISPR-Cpf1 guide RNA activity. Nat Biotechnol. 2018;36:239–41.

4. DeWeirdt PC, Sanson KR, Sangree AK, Hegde M, Hanna RE, Feeley MN, et al. Optimization of AsCas12a for combinatorial genetic screens in human cells. Nat Biotechnol [Internet]. 2020; Available from: https://www.ncbi.nlm.nih.gov/pubmed/32661438

5. Canver MC, Haeussler M, Bauer DE, Orkin SH, Sanjana NE, Shalem O, et al. Integrated design, execution, and analysis of arrayed and pooled CRISPR genome-editing experiments. Nat Protoc. 2018;13:946–86.

6. Joung J, Konermann S, Gootenberg JS, Abudayyeh OO, Platt RJ, Brigham MD, et al. Genome-scale CRISPR-Cas9 Knockout and Transcriptional Activation Screening. Nat Protoc. 2017;12:828–63.

7. Li W, Köster J, Xu H, Chen CH, Xiao T, Liu JS, et al. Quality control, modeling, and visualization of CRISPR screens with MAGeCK-VISPR. Genome Biol. 2015;16:281.

8. Wang B, Wang M, Zhang W, Xiao T, Chen CH, Wu A, et al. Integrative analysis of pooled CRISPR genetic screens using MAGeCKFlute. Nat Protoc. 2019;14:756–80.

9. Tzelepis K, Koike-Yusa H, De Braekeleer E, Li Y, Metzakopian E, Dovey OM, et al. A CRISPR Dropout Screen Identifies Genetic Vulnerabilities and Therapeutic Targets in Acute Myeloid Leukemia. Cell Rep. 2016;17:1193–205.

10. Deshpande A, Chen BR, Zhao L, Saddoris K, Kerr M, Zhu N, et al. Investigation of Genetic Dependencies Using CRISPR-Cas9-based Competition Assays. J Vis Exp [Internet]. 2019; Available from: https://www.ncbi.nlm.nih.gov/pubmed/30663717

11. Chen B-R, Deshpande A, Barbosa K, Kleppe M, Lei X, Yeddula N, et al. A JAK/STAT-mediated inflammatory signaling cascade drives oncogenesis in AF10-rearranged AML. Blood. American Society of Hematology; 2021;137:3403–15.

12. Xiang P, Wei W, Hofs N, Clemans-Gibbon J, Maetzig T, Lai CK, et al. A knock-in mouse strain facilitates dynamic tracking and enrichment of MEIS1. Blood Adv. 2017;1:2225–35.

13. Platt RJ, Chen S, Zhou Y, Yim MJ, Swiech L, Kempton HR, et al. CRISPR-Cas9 knockin mice for genome editing and cancer modeling. Cell. Elsevier BV; 2014;159:440–55.

14. Deshpande AJ, Deshpande A, Sinha AU, Chen L, Chang J, Cihan A, et al. AF10 Regulates Progressive H3K79 Methylation and HOX Gene Expression in Diverse AML Subtypes. Cancer Cell [Internet]. 2014; Available from: http://dx.doi.org/10.1016/j.ccell.2014.10.009

15. Datlinger P, Rendeiro AF, Schmidl C, Krausgruber T, Traxler P, Klughammer J, et al. Pooled CRISPR screening with single-cell transcriptome readout. Nat Methods. 2017;14:297–301.

16. Martin M. Cutadapt removes adapter sequences from high-throughput sequencing reads. EMBnet J. EMBnet Stichting; 2011;17:10.

17. Dobin A, Davis CA, Schlesinger F, Drenkow J, Zaleski C, Jha S, et al. STAR: ultrafast universal RNA-seq aligner. Bioinformatics. Oxford University Press (OUP); 2013;29:15–21.

18. Li B, Dewey CN. RSEM: accurate transcript quantification from RNA-Seq data with or without a reference genome. BMC Bioinformatics. Springer Science and Business Media LLC; 2011;12:323.

19. Ewels P, Magnusson M, Lundin S, Käller M. MultiQC: summarize analysis results for multiple tools and samples in a single report. Bioinformatics. Oxford University Press (OUP); 2016;32:3047–8.

20. Love MI, Huber W, Anders S. Moderated estimation of fold change and dispersion for RNA-seq data with DESeq2. Genome Biol. Springer Science and Business Media LLC; 2014;15:550.

21. Kustatscher G, Wills KLH, Furlan C, Rappsilber J. Chromatin enrichment for proteomics. Nat Protoc. Springer Science and Business Media LLC; 2014;9:2090–9.

