## Supplementary Tables for "High Density Domain-Focused CRISPR Screens Reveal Novel Epigenetic Regulators of *HOX/MEIS* Activation in Acute Myeloid Leukemia"

**TABLE S1**

| Molecule name | Drug classification | CAS Number |
|---|---|---|
| B2 | SIRT Inhibitor | 115687-05-3 |
| Valproic acid | HDAC Inhibitor | 99-66-1 |
| Piceatannol | SIRT Activator | 10083-24-6 |
| Resveratrol | SIRT Activator | 501-36-0 |
| Suramin·6Na | SIRT Inhibitor | 129-46-4 |
| Triacetylresveratrol | SIRT Activator | 42206-94-0 |
| Phenylbutyrate·Na | HDAC Inhibitor | 1716-12-7 |
| NSC-3852 | HDAC Inhibitor | 3565-26-2 |
| Nicotinamide | SIRT Inhibitor | 98-92-0 |
| BML-266 | SIRT Inhibitor | 96969-83-4 |
| AGK2 | SIRT Inhibitor | 304896-28-4 |
| BIX-01294 | HMT Inhibitor | 935693-62-2 |
| SAHA | HDAC Inhibitor | 149647-78-9 |
| Anacardic acid | HAT Inhibitor | 16611-84-0 |
| 5-Aza-2'-deoxycytidine | DNMT Inhibitor | 2353-33-5 |
| M-344 | HDAC Inhibitor | 251456-60-7 |
| ITSA-1 | HDAC inhibitor | 200626-61-5 |
| Scriptaid | HDAC Inhibitor | 287383-59-9 |
| EX-527 | SIRT Inhibitor | 49843-98-3 |
| Salermide | SIRT Inhibitor | 1105698-15-4 |
| CI-994 | HDAC Inhibitor | 112522-64-2 |
| BML-210 | HDAC Inhibitor | 537034-17-6 |
| Tranylcypromine hemisulfate | KDM Inhibitor | 13492-01-8 ($H_2SO_4$) |
| Trichostatin A | HDAC Inhibitor | 58880-19-6 |
| 2,4-Pyridinedicarboxylic Acid | HDM Inhibitor | 499-80-9 |
| Garcinol | HAT Inhibitor | 78824-30-3 |
| Splitomicin | SIRT Inhibitor | 3/9/90 |
| Apicidin | HDAC Inhibitor | 183506-66-3 |
| Suberoyl bis-hydroxamic acid | HDAC Inhibitor | 38937-66-5 |
| Nullscript | Scriptaid Neg control | 300816-11-9 |
| Zebularine | DNMT Inhibitor | 10/6/90 |
| Isonicotinamide | Nicotinamide antagonist | 1453-82-3 |
| Fluoro-SAHA | HDAC Inhibitor | 149648-08-8 |

| | | |
|---|---|---|
| Valproic acid hydroxamate | HDAC Inhibitor | 106132-78-9 |
| MC-1293 | HDAC Inhibitor | 117378-93-5 |
| Butyrolactone 3 | HAT inhibitor | 778649-18-6 |
| CTPB | HAT inhibitor | 586976-24-1 |
| Oxamflatin | HDAC Inhibitor | 151720-43-3 |
| Sirtinol | SIRT Inhibitor | 410536-97-9 |
| BML-278 | SIRT Activator | 120533-76-8 |
| NCH-51 | HDAC Inhibitor | 848354-66-5 |
| Aminoresveratrol sulfate | SIRT Activator | 1224713-76-1 |
| BML-281 | HDAC Inhibitor | 1045792-66-2 |
| Droxinostat | HDAC Inhibitor | 99873-43-5 |
| Azacitidine | Nucleoside analog | 320-67-2 |
| INO-1001 (3-Aminobenzamide) | PARP Inhibitor | 3544-24-9 |
| 2-Methoxyestradiol (2-MeOE2) | HIF Inhibitor | 362-07-2 |
| Procainamide HCl | DNMT Inhibitor | 614-39-1 |
| Quercetin | PI3K inhibitror | 117-39-5 |
| AG-490 (Tyrphostin B42) | JAK Inhibitor | 133550-30-8 |
| RG108 | DNMT Inhibitor | 48208-26-0 |
| WHI-P154 | JAK Inhibitor | 211555-04-3 |
| JNJ-7706621 | Aurora Kinase Inhibitor | 443797-96-4 |
| PJ34 | PARP Inhibitor | 344458-19-1 |
| WP1066 | JAK Inhibitor | 857064-38-1 |
| Entinostat (MS-275) | HDAC Inhibitor | 209783-80-2 |
| Mocetinostat (MGCD0103) | HDAC Inhibitor | 726169-73-9 |
| Belinostat (PXD101) | HDAC Inhibitor | 414864-00-9 |
| Panobinostat (LBH589) | HDAC Inhibitor | 404950-80-7 |
| Entacapone | Methyltransferase inhibitor | 130929-57-6 |
| Alisertib (MLN8237) | Aurora Kinase Inhibitor | 1028486-01-2 |
| Romidepsin (FK228, Depsipeptide) | HDAC Inhibitor | 128517-07-7 |
| S-Ruxolitinib (INCB018424) | JAK Inhibitor | 941678-49-5 |

| | | |
|---|---|---|
| ZM 447439 | Aurora Kinase Inhibitor | 331771-20-1 |
| VX-680 (Tozasertib, MK-0457) | Aurora Kinase Inhibitor | 639089-54-6 |
| Danusertib (PHA-739358) | Aurora Kinase Inhibitor | 827318-97-8 |
| AT9283 | JAK Inhibitor | 896466-04-9 |
| Barasertib (AZD1152-HQPA) | Aurora Kinase Inhibitor | 722544-51-6 |
| SNS-314 Mesylate | Aurora Kinase Inhibitor | 1146618-41-8 |
| CYC116 | Aurora Kinase Inhibitor | 693228-63-6 |
| ENMD-2076 | Aurora Kinase Inhibitor | 1291074-87-7 |
| Aurora A Inhibitor I | Aurora Kinase Inhibitor | 1158838-45-9 |
| PHA-680632 | Aurora Kinase Inhibitor | 398493-79-3 |
| CCT129202 | Aurora Kinase Inhibitor | 942947-93-5 |
| Hesperadin | Aurora Kinase Inhibitor | 422513-13-1 |
| NVP-BSK805 2HCl | JAK Inhibitor | 1092499-93-8 (free base) |
| KW-2449 | Aurora Kinase Inhibitor | 1000669-72-6 |
| LY2784544 | JAK Inhibitor | 1229236-86-5 |
| AZ 960 | JAK Inhibitor | 905586-69-8 |
| CYT387 | JAK Inhibitor | 1056634-68-4 |
| Tofacitinib (CP-690550,Tasocitinib) | JAK Inhibitor | 540737-29-9 |
| TAK-901 | JAK Inhibitor | 934541-31-8 |
| TG101209 | JAK Inhibitor | 936091-14-4 |
| AMG-900 | Aurora Kinase Inhibitor | 945595-80-2 |
| MLN8054 | Aurora Kinase Inhibitor | 869363-13-3 |
| Baricitinib (LY3009104, INCB028050) | JAK Inhibitor | 1187594-09-7 |
| TG101348 (SAR302503) | JAK Inhibitor | 936091-26-8 |
| MK-5108 (VX-689) | Aurora Kinase Inhibitor | 1010085-13-8 |
| CCT137690 | Aurora Kinase Inhibitor | 1095382-05-0 |
| CEP-33779 | JAK Inhibitor | 1257704-57-6 |
| FG-4592 | HIF Inhibitor | 808118-40-3 |
| CUDC-907 | HDAC Inhibitor | 1339928-25-4 |
| Olaparib (AZD2281, Ku-0059436) | PARP Inhibitor | 763113-22-0 |
| IOX2 | HIF Inhibitor | 931398-72-0 |

| | | |
|---|---|---|
| Veliparib (ABT-888) | PARP Inhibitor | 912444-00-9 |
| AR-42 | HDAC Inhibitor | 935881-37-1 |
| Iniparib (BSI-201) | PARP Inhibitor | 160003-66-7 |
| PCI-24781 (Abexinostat) | HDAC Inhibitor | 783355-60-2 |
| LAQ824 (Dacinostat) | HDAC Inhibitor | 404951-53-7 |
| Quisinostat (JNJ-26481585) | HDAC Inhibitor | 875320-29-9 |
| Rucaparib (AG-014699,PF-01367338) | PARP Inhibitor | 459868-92-9 |
| SRT1720 | SIRT Activator | 1001645-58-4 |
| CUDC-101 | HDAC Inhibitor | 1012054-59-9 |
| MC1568 | HDAC Inhibitor | 852475-26-4 |
| Pracinostat (SB939) | HDAC Inhibitor | 929016-96-6 |
| Givinostat (ITF2357) | HDAC Inhibitor | 732302-99-7 |
| AG-14361 | PARP Inhibitor | 328543-09-5 |
| SGI-1776 free base | Pim Kinase Inhibitor | 1025065-69-3 |
| Tubastatin A HCl | HDAC Inhibitor | 1310693-92-5 |
| PCI-34051 | HDAC Inhibitor | 950762-95-5 |
| PFI-1 (PF-6405761) | BET Inhibitor | 1403764-72-6 |
| Sodium Phenylbutyrate | HDAC Inhibitor | 1716-12-7 |
| AZD2461 | PARP Inhibitor | 1174043-16-3 |
| Resminostat | HDAC Inhibitor | 864814-88-0 |
| I-BET151 (GSK1210151A) | BET Inhibitor | 1300031-49-5 |
| AZD1480 | JAK Inhibitor | 935666-88-9 |
| XL019 | JAK Inhibitor | 945755-56-6 |
| Tubacin | HDAC Inhibitor | 537049-40-4 |
| ZM 39923 HCl | JAK Inhibitor | 1021868-92-7 |
| 3-Deazaneplanocin A (DZNeP) | Nucleoside analog | 120964-45-6 |
| SMI-4a | Pim Kinase Inhibitor | 438190-29-5 |
| (+)-JQ1 | BET Inhibitor | 1268524-70-4 |
| BMN 673 | PARP Inhibitor | 1207456-01-6 |

| | | |
|---|---|---|
| Pacritinib (SB1518) | JAK Inhibitor | 937272-79-2 |
| Rocilinostat (ACY-1215) | HDAC Inhibitor | 1316214-52-4 |
| UPF 1069 | PARP Inhibitor | 1048371-03-4 |
| EPZ5676 | DOT1L Inhibitor | 1380288-87-8 |
| GSK J4 HCl | HDM Inhibitor | 1797983-09-5 |
| EPZ004777 | DOT1L Inhibitor | 1338466-77-5 |
| Bromosporine | BET Inhibitor | 1619994-69-2 |
| Lomeguatrib | Methyltransferase inhibitor | 192441-08-0 |
| I-BET-762 | BET Inhibitor | 1260907-17-2 |
| RGFP966 | HDAC Inhibitor | 1396841-57-8 |
| SGC 0946 | DOT1L Inhibitor | 1561178-17-3 |
| SGI-1027 | DNMT Inhibitor | 1020149-73-8 |
| EPZ-6438 | EZH Inhibitor | 1403254-99-8 |
| RVX-208 | BET Inhibitor | 1044870-39-4 |
| MM-102 | MLL-complex Inhibitor | 1417329-24-8 |
| RG2833 (RGFP109) | HDAC Inhibitor | 1215493-56-3 |
| SGC-CBP30 | CBP Inhibitor | 1613695-14-9 |
| ME0328 | PARP Inhibitor | 1445251-22-8 |
| UNC669 | MBT inhibitor | 1314241-44-5 |
| OTX015 | BET Inhibitor | 202590-98-5 |
| Nexturastat A | HDAC Inhibitor | 1403783-31-2 |
| OG-L002 | LSD1 Inhibitor | 1357302-64-7 |
| C646 | HAT Inhibitor | 328968-36-1 |
| UNC1215 | MBT antagonist | 1415800-43-9 |
| IOX1 | HDM Inhibitor | 5852-78-8 |
| AZD1208 | Pim Kinase Inhibitor | 1204144-28-4 |
| CX-6258 HCl | Pim Kinase Inhibitor | 1353859-00-3 |
| CPI-203 | BET Inhibitor | 1446144-04-2 |
| TMP269 | HDAC Inhibitor | 1314890-29-3 |
| Filgotinib (GLPG0634) | JAK Inhibitor | 1206161-97-8 |
| Isoliquiritigenin | Methyltransferase inhibitor | 961-29-5 |
| Ellagic Acid | CK2 inhibitor | 476-66-4 |
| Sodium Butyrate | HDAC Inhibitor | 156-54-7 |
| Etoposide | Topoisomerase inhibitor | 33419-42-0 |
| Tenovin-1 | SIRT Inhibitor | 380315-80-0 |
| Gemcitabine | Nucleoside analog | 95058-81-4 |
| CPTH2 (hydrochloride) | HAT Inhibitor | 357649-93-5 |
| UNC0638 | G9a/GLP Inhibitor | 1255580-76-7 |
| Phthalazinone pyrazole | Aurora Kinase Inhibitor | 88048-62-7 |

| | | |
|---|---|---|
| 4-iodo-SAHA | HDAC Inhibitor | 1219807-87-0 |
| UNC0321 (trifluoroacetate salt) | G9a/GLP Inhibitor | 1238673-32-9 |
| (-)-Neplanocin A | SAM inhibitor | 72877-50-0 |
| Cl-Amidine (trifluoroacetate) | Methyltransferase inhibitor | 913723-61-2 |
| F-Amidine (trifluoroacetate salt) | Methylation inhibitor | 877617-46-4 |
| JGB1741 | SIRT Inhibitor | 1256375-38-8 |
| CCG-100602 | RhoA/C SRF Inhibitor | 1207113-88-9 |
| CAY10669 | HAT Inhibitor | 1243583-88-1 |
| Delphinidin (chloride) | CBP Inhibitor | 528-53-0 |
| MI-2 (hydrochloride) | MLL-complex Inhibitor | 1271738-62-5 |
| MI-nc (hydrochloride) | MLL-complex Inhibitor | 1359873-45-2 |
| Octyl-.alpha.-ketoglutarate | Metabolite | 876150-14-0 |
| Daminozide | HDM Inhibitor | 1596-84-5 |
| GSK-J1 (sodium salt) | HDM Inhibitor | 1373422-53-7 |
| GSK-J2 (sodium salt) | Negative control | 1394854-52-4 |
| GSK-J5 (hydrochloride) | Demethylase inhibitor | 1797983-32-4 |
| HC Toxin | HDAC Inhibitor | 83209-65-8 |
| (+)-Abscisic Acid | HDAC inhibitor | 21293-29-8 |
| 4-pentynoyl-Coenzyme A (trifluoroacetate salt) | KAT Inhibitor | 50347-32-5 |
| coumarin-SAHA | HDAC Inhibitor | 1260635-77-5 |
| SAHA-BPyne | HDAC Inhibitor | 930772-88-6 |
| UNC0631 | G9a/GLP Inhibitor | 1320288-19-4 |
| UNC0646 | G9a/GLP Inhibitor | 1320288-17-2 |
| GSK4112? | #N/A | 1216744-19-2 |
| Lestaurtinib | JAK Inhibitor | 111358-88-4 |
| Tenovin-6 | Sirtuin inhibitor | 1011557-82-6 |
| Chaetocin | G9a/GLP Inhibitor | 28097-03-2 |
| CBHA | HDAC Inhibitor | 174664-65-4 |
| Mirin | HDR inhibitor | 299953-00-7 |

| | | |
|---|---|---|
| 6-Thioguanine | Nucleoside analog | 154-42-7 |
| SIRT1/2 Inhibitor IV | HDAC Inhibitor | 14513-15-6 |
| CAY10591 | SIRT Activator | 839699-72-8 |
| S-Adenosylhomocysteine | Methyltransferase inhibitor | 979-92-0 |
| HNHA | HDAC Inhibitor | 926908-04-5 |
| 2-Hydroxyglutaric Acid (sodium salt) | Methyltransferase inhibitor | 40951-21-1 |
| 3,3'-Diindolylmethane | HDAC inhibitor | 5/4/68 |
| S-(5'-Adenosyl)-L-methionine chloride (hydrochloride) | Methyltransferase inhibitor | 86867-01-8 |
| Pimelic Diphenylamide 106 | HDAC Inhibitor | 937039-45-7 |
| 2',3',5'-triacetyl-5-Azacytidine | Nucleoside analog | 10302-78-0 |
| UNC0224 | G9a/GLP Inhibitor | 1197196-48-7 |
| Sinefungin | Nucleoside analog | 58944-73-3 |
| Pyroxamide | HDAC Inhibitor | 382180-17-8 |
| WDR5-0103 | MLL-complex Inhibitor | 890190-22-4 |
| AMI-1 (sodium salt) | PRMT Inhibitor | 20324-87-2 |
| GSK343 | EZH Inhibitor | 1346704-33-3 |
| I-CBP112 (hydrochloride) | CBP Inhibitor | 1640282-31-0 |
| UNC1999 | EZH Inhibitor | 1431612-23-5 |
| PFI-3 | SMARCA4 Inhibitor | 1819363-80-8 |
| 2,4-DPD | HIF Inhibitor | 41438-38-4 |
| DMOG | HIF Inhibitor | 89464-63-1 |
| CAY10398 | HDAC Inhibitor | 193551-00-7 |
| RSC-133 | HDAC Inhibitor | 1418131-46-0 |
| N-Oxalylglycine | HDM Inhibitor | 5262-39-5 |
| Chidamide | HDAC Inhibitor | 743420-02-2 |
| EPZ005687 | EZH Inhibitor | 1396772-26-1 |
| AK-7 | SIRT Inhibitor | 420831-40-9 |
| UNC0642 | G9a/GLP Inhibitor | 1481677-78-4 |
| (R)-PFI-2 (hydrochloride) | Methyltransferase inhibitor | 1627607-87-7 |
| HPOB | HDAC Inhibitor | 1429651-50-2 |

| | | |
|---|---|---|
| 2-hexyl-4-Pentynoic Acid | HDAC Inhibitor | 96017-59-3 |
| JIB-04 | HDM Inhibitor | 199596-05-9 |
| CAY10683 | HDAC Inhibitor | 1477949-42-0 |
| GSK 126 | EZH Inhibitor | 1346574-57-9 |
| MS-436 | BET Inhibitor | 1395084-25-9 |
| 5-Methylcytidine | Nucleoside analog | 2140-61-6 |
| AGK7 | Negative Control | 304896-21-7 |
| 5-Methyl-2'-deoxycytidine | Nucleoside analog | 838-07-3 |
| B32B3 | Histone phosphorylation inhibitor | 294193-86-5 |
| GSK-LSD1 (hydrochloride) | LSD1 Inhibitor | 1431368-48-7 |
| AZ 505 | KMT Inhibitor | 1035227-43-0 |
| BRD73954 | HDAC Inhibitor | 1440209-96-0 |
| CPI-360 | EZH Inhibitor | 1802175-06-9 |
| Remodelin | Acetyltranserase inhibitor | 1622921-15-6 |
| UNC0379 | KMT Inhibitor | 1620401-82-2 |
| GSK2801 | BAZ2 Inhibitor | 1619994-68-1 |
| CPI-169 | EZH Inhibitor | 1450655-76-1 |
| ORY-1001 (RG-6016) | LSD1 Inhibitor | 1431326-61-2 |
| SP2509 | LSD1 Inhibitor | 1423715-09-6 |
| EI1 | EZH Inhibitor | 1418308-27-6 |
| BRD4770 | G9a/GLP Inhibitor | 1374601-40-7 |
| GSK503 | EZH Inhibitor | 1346572-63-1 |
| GSK1324726A (I-BET726) | BET Inhibitor | 1300031-52-0 |
| MI-3 (Menin-MLL Inhibitor) | MLL-complex Inhibitor | 1271738-59-0 |
| MG149 | HAT Inhibitor | 1243583-85-8 |
| ML324 | HDM Inhibitor | 1222800-79-4 |
| OF-1 | BRPF Inhibitor | 919973-83-4 |
| 4SC-202 | HDAC Inhibitor | 910462-43-0 |
| NI-57 | BRPF Inhibitor | 1883548-89-7 |
| MS023 hydrochloride | PRMT Inhibitor | 1831110-54-3 |
| OICR-9429 | MLL-complex Inhibitor | 1801787-56-3 |
| LLY-507 | KMT Inhibitor | 1793053-37-8 |
| I-BRD9 | BET Inhibitor | 1714146-59-4 |
| SGC707 | PRMT Inhibitor | 1687736-54-4 |
| BAZ2-ICR | BAZ2 Inhibitor | 1665195-94-7 |
| A-366 | G9a/GLP Inhibitor | 1527503-11-2 |
| MS049 hydrochloride | PRMT Inhibitor | 1502816-23-0 |
| PFI-4 | BRPF Inhibitor | 900305-37-5 |

| | | |
|---|---|---|
| A-196 | Methyltransferase inhibitor | 1982372-88-2 |
| (+/-)-JQ1 | BET Inhibitor | 1268524-69-1 |
| NM1-100 | YEATS Domain inhibitor | |
| TM-7 | YEATS Domain inhibitor | |
| SK99-53 | YEATS Domain inhibitor | |
| SGC-IMLLT | YEATS Domain inhibitor | |

**TABLE S2**

| Library | Gene | Annotation | sgRNA Sequence |
|---|---|---|---|
| Human | AARS2 | Control | AGACACAGCGTACCGCGTGG |
| Human | AARS2 | Control | CCAACTTCTACGCAGAACAG |
| Human | AARS2 | Control | TAGGACGACTCATTGAAGTG |
| Human | AARS2 | Control | GAGACTGTGTCCACAATCAG |
| Human | AARS2 | Control | CCACTGATACCACCCGCACA |
| Human | ACTL6A | Chromatin Remodeling | ACTGCAATTCCAGTCCACGA |
| Human | ACTL6A | Chromatin Remodeling | ACCACCATACCAATAGCTGT |
| Human | ACTL6A | Chromatin Remodeling | GATGGCTATGTCCTTCAACA |
| Human | ACTL6A | Chromatin Remodeling | TGCCAAGACCTCGTAACCTG |
| Human | ACTL6A | Chromatin Remodeling | AGCTCATCTACTAACCAAAG |
| Human | ACTL6B | Chromatin Remodeling | GTGCACTACGAGATGCCCAA |
| Human | ACTL6B | Chromatin Remodeling | ATGCCAGCTTTGCAAACGGG |
| Human | ACTL6B | Chromatin Remodeling | CACTCACTGTTCATCGTAGG |
| Human | ACTL6B | Chromatin Remodeling | ACACTCACTGTTCATCGTAG |
| Human | ACTL6B | Chromatin Remodeling | ACCTTACATGATCGCAGCCA |
| Human | ACTR5 | Chromatin Remodeling | CCAAGCGGAAGTCAACCTCG |
| Human | ACTR5 | Chromatin Remodeling | ACATCGTGTTGCCGCCAGTG |
| Human | ACTR5 | Chromatin Remodeling | TGAGCACAGCTACATCGCTG |
| Human | ACTR5 | Chromatin Remodeling | TATGCTGCCAAGTTAAACAC |
| Human | ACTR5 | Chromatin Remodeling | AGCGGAAGTCAACCTCGAGG |
| Human | ACTR8 | Chromatin Remodeling | CAGCCCGCTCTCGTAACATG |
| Human | ACTR8 | Chromatin Remodeling | GGCCGGCAACGATTCCGAGG |
| Human | ACTR8 | Chromatin Remodeling | GTCTCCGGCCATAATGACCC |
| Human | ACTR8 | Chromatin Remodeling | TGGATACAACACAGGAACTG |
| Human | ACTR8 | Chromatin Remodeling | AGGCTTTGTCTGGCATACGG |
| Human | ADRM1 | Transcription Elongation | CCGCTTATCCGGAGTCACGG |
| Human | ADRM1 | Transcription Elongation | GGTATTCTGGATCTCATCCG |
| Human | ADRM1 | Transcription Elongation | GCTTGAACTCACAGTCGTCA |
| Human | ADRM1 | Transcription Elongation | GCTTGGAAAGAGCGCGCCTG |
| Human | ADRM1 | Transcription Elongation | TCTTTAACGTGTTCTAGATG |
| Human | AFF2 | Transcription Elongation | AAAGTCGAATAGATCCATAG |
| Human | AFF2 | Transcription Elongation | ATCACTAGCAATGTGTTACG |
| Human | AFF2 | Transcription Elongation | GAGCTTCATGACCCACCAAG |
| Human | AFF2 | Transcription Elongation | GTACTCTAGAGAGTAAGTCA |
| Human | AFF2 | Transcription Elongation | ACAGTCCTTGGCGAACGTAG |
| Human | AFF3 | Transcription Elongation | GAGCAATCAGTACTACAACC |
| Human | AFF3 | Transcription Elongation | CAGCTTAGGAGACTCATCAG |
| Human | AFF3 | Transcription Elongation | TCTGTCCAAAGCACAGACCG |
| Human | AFF3 | Transcription Elongation | GATGCTGACGATGGTCGACG |

| | | | |
|---|---|---|---|
| Human | AFF3 | Transcription Elongation | CAGATAGAAATGCATTACGG |
| Human | AFF4 | Transcription Elongation | TGGGAATGATCACCATAGCA |
| Human | AFF4 | Transcription Elongation | AAAGAAACAGAGCCGCCCAA |
| Human | AFF4 | Transcription Elongation | GGTGTATGAATAGCCGTTAG |
| Human | AFF4 | Transcription Elongation | ATTGTGATAAGACAATGCCG |
| Human | AFF4 | Transcription Elongation | TCCACCGTCTCTGAATACAT |
| Human | AFG3L2 | Control | GAATGAGACTCACTCTAGCA |
| Human | AFG3L2 | Control | CTTCATCGATGAAATCGATG |
| Human | AFG3L2 | Control | ATTTGTTACAACTCAAGCAA |
| Human | AFG3L2 | Control | TGACATTCCATGGGACGACA |
| Human | AFG3L2 | Control | TGTCGGAGAAACCACTGCCA |
| Human | AIRE | Chromatin Readers | AGGCGTTACCTCGCTCAGAA |
| Human | AIRE | Chromatin Readers | GAGATGGTACGGACGTGCTG |
| Human | AIRE | Chromatin Readers | GTGGCGGTACCAAAGCCTTG |
| Human | AIRE | Chromatin Readers | AGGAGGTAAGAGGTCCACCT |
| Human | AIRE | Chromatin Readers | CCACCTGAAACTTGTCCTCG |
| Human | AKAP1 | Chromatin Readers | ATTCGATACGTGGACTACGG |
| Human | AKAP1 | Chromatin Readers | TTCCAGGCTGAGAGTAACAG |
| Human | AKAP1 | Chromatin Readers | CGAAGTGGAGATTCGATACG |
| Human | AKAP1 | Chromatin Readers | GGTGTTAGGAAGAATGCCCG |
| Human | AKAP1 | Chromatin Readers | GTCCGGCAACTCTTCACTGG |
| Human | ALKBH5 | DNA and RNA Modifications | ATCAACGACTACCAGCCCGG |
| Human | ALKBH5 | DNA and RNA Modifications | GGACAAGATTAGATGCACCC |
| Human | ALKBH5 | DNA and RNA Modifications | CCTGTACAACGAGCACACGG |
| Human | ALKBH5 | DNA and RNA Modifications | TCCCGGGACAACTATAAGGC |
| Human | ALKBH5 | DNA and RNA Modifications | TGGACTTGAGCTTCTCACGC |
| Human | ALYREF | Control | TTCCTTGCTAGCGTAAACAG |
| Human | ALYREF | Control | CGTCACAGATTGACGCACAG |
| Human | ALYREF | Control | CACAGATTGACGCACAGCGG |
| Human | ALYREF | Control | CAGCAGACGTGCACTTTGAG |
| Human | ALYREF | Control | CTTGCTAGCGTAAACAGAGG |
| Human | ARID1A | Chromatin Remodeling | CAGCAGAACTCTCACGACCA |
| Human | ARID1A | Chromatin Remodeling | TTACACTCGCCAACATCTCG |
| Human | ARID1A | Chromatin Remodeling | GATATCCCGCCGAATCATGG |
| Human | ARID1A | Chromatin Remodeling | CAGACACATAGAGGCGATAG |
| Human | ARID1A | Chromatin Remodeling | ATATCCCGCCGAATCATGGG |
| Human | ARID1B | Chromatin Remodeling | TTGAGTGCAAGATCGAACGT |
| Human | ARID1B | Chromatin Remodeling | CAGTGCGCGATGGTACACAG |
| Human | ARID1B | Chromatin Remodeling | TGAGTGCAAGATCGAACGTG |
| Human | ARID1B | Chromatin Remodeling | TGCCAATTGGATACCGCTGT |
| Human | ARID1B | Chromatin Remodeling | AAGTTGCTTCCGTTCCCGTG |
| Human | ARID2 | Chromatin Remodeling | GTAAGCCAGCCAGCTCAACA |

| | | | |
|---|---|---|---|
| Human | ARID2 | Chromatin Remodeling | GCAGTCTCCATTACACACAG |
| Human | ARID2 | Chromatin Remodeling | AGAAGTTGTTACATACTGTG |
| Human | ARID2 | Chromatin Remodeling | TTGATGTCCCTGTATAACAG |
| Human | ARID2 | Chromatin Remodeling | TTGTTCAGAATCATAGTACA |
| Human | ARID4A | Chromatin Readers | GATGCTAGTTGGTATACCGT |
| Human | ARID4A | Chromatin Readers | ATTGCAAAGAAGACGAACAG |
| Human | ARID4A | Chromatin Readers | AAGTTATGTTGGACTACAGG |
| Human | ARID4A | Chromatin Readers | AGATGCTAGTTGGTATACCG |
| Human | ARID4A | Chromatin Readers | AAGGTTTCAGGCCCGATCAG |
| Human | ARID4B | Chromatin Readers | AAGTAGCTTCCATTGATCGG |
| Human | ARID4B | Chromatin Readers | TTACTATCGACATTGACTGG |
| Human | ARID4B | Chromatin Readers | CAGGCATGAAAGTCCAAGTG |
| Human | ARID4B | Chromatin Readers | GATGCGAGTTGGTACACTGT |
| Human | ARID4B | Chromatin Readers | ATAGTACCAAATCTCCCGCA |
| Human | ARIH1 | Control | GATACTGTGAGAGTCGACGA |
| Human | ARIH1 | Control | ATACTGTGAGAGTCGACGAA |
| Human | ARIH1 | Control | ACATGAGTGTGGCACGACAC |
| Human | ARIH1 | Control | TCACTTAACAGCTTTAACTG |
| Human | ARIH1 | Control | GATATTGGACTTTAACAACA |
| Human | ASH1L | Histone Methylation | CCTATGACCAACGTTCAAGT |
| Human | ASH1L | Histone Methylation | CTGTCTTATAGTAACCAGTG |
| Human | ASH1L | Histone Methylation | ACCTCGGTGGACTAAAGTGG |
| Human | ASH1L | Histone Methylation | GTTCATTATGATAGAACCGA |
| Human | ASH1L | Histone Methylation | GACAGAGCATTACTTAATGG |
| Human | ASXL1 | Chromatin Readers | GAATCAGTGCGTATACAGCG |
| Human | ASXL1 | Chromatin Readers | CTTTGAAGGTCACCTCACGG |
| Human | ASXL1 | Chromatin Readers | ATCTGACGTACACTTTCCAG |
| Human | ASXL1 | Chromatin Readers | GAGTCATTGCAGCAGAACGT |
| Human | ASXL1 | Chromatin Readers | AAAGCCATGATCATGTGCCA |
| Human | ASXL2 | Chromatin Readers | ACAGCCCTTTCTCAATAGAG |
| Human | ASXL2 | Chromatin Readers | TGATCAACAAGAGTTGCTAG |
| Human | ASXL2 | Chromatin Readers | TTGGAGGAGATGGATATGCT |
| Human | ASXL2 | Chromatin Readers | GCACCTGTCGATCTACCTCT |
| Human | ASXL2 | Chromatin Readers | TGACATTGACGTTGAGACAC |
| Human | ASXL3 | Chromatin Readers | GAACCCGAGTGAGTAACCAG |
| Human | ASXL3 | Chromatin Readers | AGGTAAGTCAGACATCGGTG |
| Human | ASXL3 | Chromatin Readers | GTTCTGCTGGAAGCTACACG |
| Human | ASXL3 | Chromatin Readers | TCAGCTCTTATCGTACAACC |
| Human | ASXL3 | Chromatin Readers | CTGCTAATCACAGACACCTG |
| Human | ATAD2 | Chromatin Readers | CATCACATGGTAGTAAGACG |
| Human | ATAD2 | Chromatin Readers | TAGTTGCATTGGATCAACAT |
| Human | ATAD2 | Chromatin Readers | CTATAGATCCTGCTTTACGA |

| | | | |
|---|---|---|---|
| Human | ATAD2 | Chromatin Readers | AGATACCGATTATCTTCCGC |
| Human | ATAD2 | Chromatin Readers | ATCTTAAAGCACGTGTCCGG |
| Human | ATAD2B | Chromatin Readers | AGACTATGAGAACCATACGG |
| Human | ATAD2B | Chromatin Readers | TCACGATCAACTATAAGAGG |
| Human | ATAD2B | Chromatin Readers | AAGTAGAACGACTTCGAATG |
| Human | ATAD2B | Chromatin Readers | TGGACAACATTAACATCCGA |
| Human | ATAD2B | Chromatin Readers | AGAGCAGAGGACTTAGCTAG |
| Human | ATAT1 | Histone Acetylation | CATGGAGTTCCCGTTCGATG |
| Human | ATAT1 | Histone Acetylation | TGATGCACTAGTGATAGGAG |
| Human | ATAT1 | Histone Acetylation | AGGGTCGGTCAATTGCCAGT |
| Human | ATAT1 | Histone Acetylation | TGAGAATATAAACAACATGG |
| Human | ATAT1 | Histone Acetylation | TATCGACTCTCCTCAGAGCG |
| Human | ATP6V0B | Control | GAAGAGGTTAGACAGGCCTA |
| Human | ATP6V0B | Control | TTTGGGCTTCCGCTTTGATG |
| Human | ATP6V0B | Control | CACTCCTCCACCAATGATGG |
| Human | ATP6V0B | Control | GCTAGCACTGCTCTACTCCG |
| Human | ATP6V0B | Control | CTTTGTAAAGATTCTCATCG |
| Human | ATRIP | Control | TGTACATGTACATCACATCA |
| Human | ATRIP | Control | GCACCATCATTACCTGACAT |
| Human | ATRIP | Control | GGGAGACGACTGCTCCGCTG |
| Human | ATRIP | Control | TATGACCTCAGCCCTAAGGG |
| Human | ATRIP | Control | CGGTGATGTTGCACAGACAG |
| Human | AURKB | Other Histone Modifications | CTATCGCCGCATCGTCAAGG |
| Human | AURKB | Other Histone Modifications | TCTTTCCGGAGGACTCGCTG |
| Human | AURKB | Other Histone Modifications | CTTACCGTCTGTCGGCCGTA |
| Human | AURKB | Other Histone Modifications | ATGCTCTAATGTACTGCCAT |
| Human | AURKB | Other Histone Modifications | TGACGAGCAGCGAACAGCCA |
| Human | AXIN1 | Transcription Elongation | AGCCGGCATTGACATAATAG |
| Human | AXIN1 | Transcription Elongation | GGAGCCCTGTGACTCGAACG |
| Human | AXIN1 | Transcription Elongation | CGAACTTCTGAGGCTCCACG |
| Human | AXIN1 | Transcription Elongation | ATATTTATTTGGAATATACG |
| Human | AXIN1 | Transcription Elongation | GAGCGACGAGTTTGACTGTG |
| Human | BANP | Chromatin Remodeling | CACTGCAATCTGAACCACAT |
| Human | BANP | Chromatin Remodeling | AGACGTGCAACAAAGTGCGA |
| Human | BANP | Chromatin Remodeling | GCGTCGGCAGAACACCATTG |
| Human | BANP | Chromatin Remodeling | TGGCGGTCAAGAGCTTCTCG |
| Human | BANP | Chromatin Remodeling | CAATCTGAACCACATCGGCC |
| Human | BAZ1A | Chromatin Readers | TATCACGCTACGATCCAAGG |
| Human | BAZ1A | Chromatin Readers | CAGTGCATAAGCCAAATCGG |
| Human | BAZ1A | Chromatin Readers | TGGTGTCAAATCATACACTG |
| Human | BAZ1A | Chromatin Readers | AGCAAACCAACATATAGTCG |
| Human | BAZ1A | Chromatin Readers | GGACCAAGTTAGCACACCAC |

| | | | |
|---|---|---|---|
| Human | BAZ1B | Other Histone Modifications | AGACCTCGAAAGACCATCCG |
| Human | BAZ1B | Other Histone Modifications | GCTGAGGTTTACAACTGCCG |
| Human | BAZ1B | Other Histone Modifications | ATGTTTATGAGGTTTACTAG |
| Human | BAZ1B | Other Histone Modifications | AGAAGCCACACTCTACCAGG |
| Human | BAZ1B | Other Histone Modifications | ACTTTGTTGGATATGGCCAA |
| Human | BAZ2A | Chromatin Readers | ACGGTACCCAGAGTCCCATG |
| Human | BAZ2A | Chromatin Readers | GCGCCAGACACGAATGCGAG |
| Human | BAZ2A | Chromatin Readers | CACTGGGCAATGTCAGACCA |
| Human | BAZ2A | Chromatin Readers | TGAATGCTAGATCCCAACAT |
| Human | BAZ2A | Chromatin Readers | AGTGTCAGACTACTATCCAA |
| Human | BAZ2B | Chromatin Readers | AGCTCTACAATCAACCCATG |
| Human | BAZ2B | Chromatin Readers | AGGGCAATGGAAGGTCGTAG |
| Human | BAZ2B | Chromatin Readers | TGGTTACAGCTATATCTAGG |
| Human | BAZ2B | Chromatin Readers | GTAGCATATTATGCTCCATG |
| Human | BAZ2B | Chromatin Readers | GTCTGGGATTCAGACGCAAG |
| Human | BPTF | Chromatin Readers | ACTGCTATCCTGAATAGACA |
| Human | BPTF | Chromatin Readers | GTGATGACTCAAATCATCAG |
| Human | BPTF | Chromatin Readers | AATTGGTGGAGTATTGATTG |
| Human | BPTF | Chromatin Readers | CAAATCGGAGAAGTCCAACG |
| Human | BPTF | Chromatin Readers | GGACGTATTCGAGTCTGTGA |
| Human | BRD1 | Chromatin Readers | AAGACAGATGACGACCGCTG |
| Human | BRD1 | Chromatin Readers | TGTTCTATAGAGCCGCGGTG |
| Human | BRD1 | Chromatin Readers | GCAGCAGTCTCTGATCGACG |
| Human | BRD1 | Chromatin Readers | GGTCAGCGTTTCTCGCGTAG |
| Human | BRD1 | Chromatin Readers | CCATGGACTTTGCCACAATG |
| Human | BRD2 | Chromatin Readers | CCAGCTGCAATACCTACACA |
| Human | BRD2 | Chromatin Readers | GCGGAAACGTACTTATTGTG |
| Human | BRD2 | Chromatin Readers | CCTGCCTACGTAAGAAACCC |
| Human | BRD2 | Chromatin Readers | TGTGCATATAATCCAAGCCA |
| Human | BRD2 | Chromatin Readers | ACAGTGGTAGGTATCTCAGG |
| Human | BRD3 | Chromatin Readers | CATCACTGCAAACGTCACGT |
| Human | BRD3 | Chromatin Readers | TGAAGGTACACAGCAAGTGG |
| Human | BRD3 | Chromatin Readers | CGAGAGTACCCAGACGCACA |
| Human | BRD3 | Chromatin Readers | ACCTTGACGACAGGCGGCGT |
| Human | BRD3 | Chromatin Readers | TCTCTCCAGTTCCCGCAAAG |
| Human | BRD4 | Chromatin Readers | TCTTCCTCCGACTCATACGT |
| Human | BRD4 | Chromatin Readers | ATCGTCCCTTCGTAGCAAGA |
| Human | BRD4 | Chromatin Readers | GGAAGGGAGTAGCGGCGTAG |
| Human | BRD4 | Chromatin Readers | AGTCGAACTGTCACTGTCCG |
| Human | BRD4 | Chromatin Readers | TCTTGGCAAACATCTCCTTG |
| Human | BRD7 | Chromatin Readers | TCGGACAAACACCTCTACGA |
| Human | BRD7 | Chromatin Readers | GAAGTCACCGAACTCTCCAC |

| | | | |
|---|---|---|---|
| Human | BRD7 | Chromatin Readers | GTCGGACAAACACCTCTACG |
| Human | BRD7 | Chromatin Readers | AAGAGACAGATCATGTTCGG |
| Human | BRD7 | Chromatin Readers | GAGACAGATCATGTTCGGAG |
| Human | BRD8 | Chromatin Readers | TTTGCAATGACATTGCAACG |
| Human | BRD8 | Chromatin Readers | AGGAGGTGATTATCCACTTG |
| Human | BRD8 | Chromatin Readers | GGTGGCACAAGTTTAACTGG |
| Human | BRD8 | Chromatin Readers | TGTTTGCATAATCAACTGCG |
| Human | BRD8 | Chromatin Readers | TAAGGGTCGGATTCGCACCA |
| Human | BRD9 | Chromatin Readers | GAGTGGCGCTCGTCCTACGA |
| Human | BRD9 | Chromatin Readers | ATCCACAGAAACGTCGGGAT |
| Human | BRD9 | Chromatin Readers | AGATACCGTGTACTACAAGT |
| Human | BRD9 | Chromatin Readers | AGGGAGCACTGTGACACGGA |
| Human | BRD9 | Chromatin Readers | AGAGAGGGAGCACTGTGACA |
| Human | BRDT | Chromatin Readers | TCTCCCTTGAACGTGGTACA |
| Human | BRDT | Chromatin Readers | CAACTCCAGTTCACAAACTG |
| Human | BRDT | Chromatin Readers | AGGTGTGGTCTGATGAACAA |
| Human | BRDT | Chromatin Readers | ATCTATTTCTCCCTTGAACG |
| Human | BRDT | Chromatin Readers | AACTCCCTGGAGATAAACTT |
| Human | BRIX1 | Control | TAAGCTATTTGTGATTAACG |
| Human | BRIX1 | Control | GAAACCAATAGAAATACAGT |
| Human | BRIX1 | Control | ACTAGATCTTTAGTACACCA |
| Human | BRIX1 | Control | AAGGCACCTGCTTTAGAATG |
| Human | BRIX1 | Control | GTGGCGTGCCGCTTAGCTGG |
| Human | BRPF1 | Chromatin Readers | TTCTAGATGTGACAGCATGG |
| Human | BRPF1 | Chromatin Readers | GCCACCATACGAGTGCCCGG |
| Human | BRPF1 | Chromatin Readers | CCAACCGCCTGACCATCCAA |
| Human | BRPF1 | Chromatin Readers | TGCTACATTTGCAAACAACG |
| Human | BRPF1 | Chromatin Readers | GCACAATCCACAGCACGAGA |
| Human | BRPF3 | Chromatin Readers | AGAACCAGTCAACTTGAGTG |
| Human | BRPF3 | Chromatin Readers | GCGACCCACACCACTAAGTG |
| Human | BRPF3 | Chromatin Readers | CAGGTGCGGCCACTGCTAGG |
| Human | BRPF3 | Chromatin Readers | TGGCGTCCCATACATCCCTG |
| Human | BRPF3 | Chromatin Readers | TCCAAGAAACCCTCATCCAA |
| Human | BRWD1 | Chromatin Readers | TTCCTAGTGGAGATACATCG |
| Human | BRWD1 | Chromatin Readers | AGAATACAATGATTGCTGCG |
| Human | BRWD1 | Chromatin Readers | ACAAACCTAAATATGCCTAG |
| Human | BRWD1 | Chromatin Readers | TCTGTAGCAATCTGACTGCG |
| Human | BRWD1 | Chromatin Readers | GGTATGTACACAAATCAACA |
| Human | BRWD3 | Chromatin Readers | TAAGAGGACGATAATCCGTG |
| Human | BRWD3 | Chromatin Readers | CCCTCCCAATCTAAGCGGCG |
| Human | BRWD3 | Chromatin Readers | TACAATCAAACACCGCACCA |
| Human | BRWD3 | Chromatin Readers | AGAGCATACTCTGTTAATGG |

| | | | |
|---|---|---|---|
| Human | BRWD3 | Chromatin Readers | GCTGTGGGCGTTCTTTACGC |
| Human | BTBD18 | Transcription Elongation | TTGTAGCTTAGTGCACATGG |
| Human | BTBD18 | Transcription Elongation | TCTTGTAGCTTAGTGCACAT |
| Human | BTBD18 | Transcription Elongation | GATTGTCAATGGGACTTGCT |
| Human | BTBD18 | Transcription Elongation | CTGCAGTCAATAGCCAACAT |
| Human | BTBD18 | Transcription Elongation | CTTGTAGCTTAGTGCACATG |
| Human | CARM1 | Histone Methylation | TAGAGCTGTTCATCCGTGAA |
| Human | CARM1 | Histone Methylation | CTGCTCAAGTCATAGGCGGT |
| Human | CARM1 | Histone Methylation | TTGAAGAGCATGTAGCCCAT |
| Human | CARM1 | Histone Methylation | TGTCTCTGCCATAGATACAC |
| Human | CARM1 | Histone Methylation | CCATGCACTACGGGAGCTAG |
| Human | CBX1 | Chromatin Readers | TCGAGTGGTAAAGGGCAAAG |
| Human | CBX1 | Chromatin Readers | GTTTCAGTGAGGACAACACA |
| Human | CBX1 | Chromatin Readers | CAGAAACTCAGCAATGAGGT |
| Human | CBX1 | Chromatin Readers | AAACCTCGAGCAAAGCCTCG |
| Human | CBX1 | Chromatin Readers | TTTCAGTGAGGACAACACAT |
| Human | CBX2 | Chromatin Readers | TGTCAAGAATGGCATGCCCG |
| Human | CBX2 | Chromatin Readers | GCTTTGCAGACATAACAGCT |
| Human | CBX2 | Chromatin Readers | AGTGACTTAGATGCTAAGAG |
| Human | CBX2 | Chromatin Readers | ACAGTGACTTAGATGCTAAG |
| Human | CBX2 | Chromatin Readers | TGACAAATACGTGCTCGATG |
| Human | CBX3 | Chromatin Readers | AGAGCCTGAAGAATTTGTCG |
| Human | CBX3 | Chromatin Readers | AAACAGGAAAGATTCAGATG |
| Human | CBX3 | Chromatin Readers | AAACAGGCTGACAAACCAAG |
| Human | CBX3 | Chromatin Readers | TTTCTTAACTCTCAGAAAGC |
| Human | CBX3 | Chromatin Readers | AAACCAAGAGGATTTGCCAG |
| Human | CBX4 | Chromatin Readers | GGGCTGCGGATCGCTAACCC |
| Human | CBX4 | Chromatin Readers | CGGTCAGGACATTGGAACGA |
| Human | CBX4 | Chromatin Readers | CCTTGCAAGATATAACACGT |
| Human | CBX4 | Chromatin Readers | ACTCCTTGAAAGTAACGGTG |
| Human | CBX4 | Chromatin Readers | GAAGCCCGAGACACCGGCGT |
| Human | CBX5 | Chromatin Readers | CAGAGCAATGATATCGCTCG |
| Human | CBX5 | Chromatin Readers | TAGGAACATTAAATCACCAC |
| Human | CBX5 | Chromatin Readers | AGAAGTCAGAAAGTAACAAG |
| Human | CBX5 | Chromatin Readers | GACATGGCATGCATATCCTG |
| Human | CBX5 | Chromatin Readers | TAAATTCAGAAATTAGCTCA |
| Human | CBX6 | Chromatin Readers | TGTGGCGGATCTGTGTACGC |
| Human | CBX6 | Chromatin Readers | TCTTGCTCTTGCCTATAACG |
| Human | CBX6 | Chromatin Readers | CCGAATCCATCATCAAACGG |
| Human | CBX6 | Chromatin Readers | ATGCACATCACTGATGCGGA |
| Human | CBX6 | Chromatin Readers | AACCTGAAGGTGATCGACAA |
| Human | CBX7 | Chromatin Readers | AGGAGAGAGACCGAGCATCG |

| | | | |
|---|---|---|---|
| Human | CBX7 | Chromatin Readers | GCTGCCGAGTGGGCACGTCA |
| Human | CBX7 | Chromatin Readers | GCTTCTTCCGAGACCGCAGT |
| Human | CBX7 | Chromatin Readers | TGATTCAGAACTTCCCACTG |
| Human | CBX7 | Chromatin Readers | TCACGGTCACCTCACTTGAG |
| Human | CBX8 | Chromatin Readers | GACTCGGACCTGGTGCAGTG |
| Human | CBX8 | Chromatin Readers | TCCCTTTAGGTACAGCACAT |
| Human | CBX8 | Chromatin Readers | AAGGAAAGTAACACGGACCA |
| Human | CBX8 | Chromatin Readers | GGGACAAACCCATGTTTCGA |
| Human | CBX8 | Chromatin Readers | GTTTCGAAGTGACTCAGCCA |
| Human | CCAR2 | Transcription Elongation | TTACCTTCGAACTATAAGCG |
| Human | CCAR2 | Transcription Elongation | AATGGCAGCCTGATTAACGT |
| Human | CCAR2 | Transcription Elongation | AGTAACCAATAAGACGCTGG |
| Human | CCAR2 | Transcription Elongation | TGCACCGGCGAGACTTAGAG |
| Human | CCAR2 | Transcription Elongation | TGAAGTAACCAATAAGACGC |
| Human | CCNH | Transcription Elongation | GGGCTGGAATTACTATGGAA |
| Human | CCNH | Transcription Elongation | GCAGTCAGGGCAATTTGGGA |
| Human | CCNH | Transcription Elongation | AGTTTACCTTTAAGTCGATG |
| Human | CCNH | Transcription Elongation | GTCCAAGAGGACTCTCCCGG |
| Human | CCNH | Transcription Elongation | GATTCTCCAATATGGGATAG |
| Human | CCNK | Transcription Elongation | ACGGCTCGCTTAACCTGTCG |
| Human | CCNK | Transcription Elongation | TGAACCGAGCGCCCTCTCGG |
| Human | CCNK | Transcription Elongation | CCGACAGGTTAAGCGAGCCG |
| Human | CCNK | Transcription Elongation | CAGTCCATGATGAAGACCGA |
| Human | CCNK | Transcription Elongation | CCTCGCTGCTGCCTTAGGTG |
| Human | CCNT1 | Transcription Elongation | GAGGAGATTCCACCAGACCG |
| Human | CCNT1 | Transcription Elongation | GTTTCTAGCAGCTAAAGTGG |
| Human | CCNT1 | Transcription Elongation | TGTGTACTGCAGGCTAAATG |
| Human | CCNT1 | Transcription Elongation | TGATAAGCACAATTCTGTAG |
| Human | CCNT1 | Transcription Elongation | ACTTTACTACATGAGTATGT |
| Human | CCNT2 | Transcription Elongation | CAGGGAGCAGTAAACACCAT |
| Human | CCNT2 | Transcription Elongation | AACAGTGGGTGGTATTCCGT |
| Human | CCNT2 | Transcription Elongation | GAGTAGTGGAAATTGGCCCA |
| Human | CCNT2 | Transcription Elongation | TGAAGAAGATTCGAAACTGG |
| Human | CCNT2 | Transcription Elongation | GGATATATACTGCTTAACAG |
| Human | CD3EAP | Transcription Elongation | AGCGCCTCCAAGGAGAAACG |
| Human | CD3EAP | Transcription Elongation | GCGCCTCCAAGGAGAAACGA |
| Human | CD3EAP | Transcription Elongation | CAAGACCACTCACCATTCTG |
| Human | CD3EAP | Transcription Elongation | GGTCGCCAGAAATGGATGTG |
| Human | CD3EAP | Transcription Elongation | CGGGTCCAGATACGGAGCTG |
| Human | CDC6 | DNA and RNA Modifications | TTTGCGACAGACTTTACTGT |
| Human | CDC6 | DNA and RNA Modifications | GGTGGGAAGTTCAACAGCTG |
| Human | CDC6 | DNA and RNA Modifications | ATAGCGTGTACAATACATCC |

| | | | |
|---|---|---|---|
| Human | CDC6 | DNA and RNA Modifications | CCTTGTTCTCCACCAAAGCA |
| Human | CDC6 | DNA and RNA Modifications | TCTGATTTCTGGTATAAGGT |
| Human | CDC73 | Transcription Elongation | ACCCAGAGAGTACTACACAT |
| Human | CDC73 | Transcription Elongation | ACATAACTGCCCTTAAACAG |
| Human | CDC73 | Transcription Elongation | TTCTACTTACAGCTGCACGT |
| Human | CDC73 | Transcription Elongation | GGTCTTCAGCGATCTACTCA |
| Human | CDC73 | Transcription Elongation | AGGCTGTTTGGTGCGCAAAG |
| Human | CDK1 | Other Histone Modifications | ATTTATACCAAATCCTACAG |
| Human | CDK1 | Other Histone Modifications | AATCCATGTACTGACCAGGA |
| Human | CDK1 | Other Histone Modifications | ACCCTTATACACAACTCCAT |
| Human | CDK1 | Other Histone Modifications | GAATCCATGTACTGACCAGG |
| Human | CDK1 | Other Histone Modifications | ACATTTCCCAAATGGAAACC |
| Human | CDK12 | Transcription Elongation | TGACACTAGATCCTAGTAAG |
| Human | CDK12 | Transcription Elongation | AAGTTCAAAGCGTTCGAATG |
| Human | CDK12 | Transcription Elongation | AACCGAAGAAGCAATATCGA |
| Human | CDK12 | Transcription Elongation | CCAGTCCCTATGGTCGAAGG |
| Human | CDK12 | Transcription Elongation | GACAAACAGAAAGCGACTGG |
| Human | CDK13 | Transcription Elongation | GAGTCTTCGAAACCGTTGGG |
| Human | CDK13 | Transcription Elongation | GTAGAATTGCCAATTCACTG |
| Human | CDK13 | Transcription Elongation | GGTTCCTTGTAGGCCGAAGG |
| Human | CDK13 | Transcription Elongation | AAACTACGTCGTTCTAGTGG |
| Human | CDK13 | Transcription Elongation | GCCACCTCTTCAGGTAACGA |
| Human | CDK2 | Other Histone Modifications | CATGGGTGTAAGTACGAACA |
| Human | CDK2 | Other Histone Modifications | TGTTCGTACTTACACCCATG |
| Human | CDK2 | Other Histone Modifications | TTCAGGAGCTCGGTACCACA |
| Human | CDK2 | Other Histone Modifications | TCTGCTTATTAACACAGAGG |
| Human | CDK2 | Other Histone Modifications | TGCACTACGACCCTAACAAG |
| Human | CDK4 | Transcription Elongation | AGCCACTGGCTCATATCGAG |
| Human | CDK4 | Transcription Elongation | CCAGATGGCACTTACACCCG |
| Human | CDK4 | Transcription Elongation | TACCTCTCGATATGAGCCAG |
| Human | CDK4 | Transcription Elongation | GTCCACATATGCAACACCTG |
| Human | CDK4 | Transcription Elongation | TGCCACATCCCGAACTGACC |
| Human | CDK6 | Transcription Elongation | GCCCGCGACTTGAAGAACGG |
| Human | CDK6 | Transcription Elongation | TGACCAGCAGTACGAATGCG |
| Human | CDK6 | Transcription Elongation | CCAGCAGTACGAATGCGTGG |
| Human | CDK6 | Transcription Elongation | AACACTCCAGAGATCCACGG |
| Human | CDK6 | Transcription Elongation | GGAAACTATAGATGCGGGCA |
| Human | CDK7 | Transcription Elongation | AGCTCCAAATAGTAACTCGG |
| Human | CDK7 | Transcription Elongation | GATGTGTATAAGCTCTATTG |
| Human | CDK7 | Transcription Elongation | ACTTCACGTCCAGAGCCATC |
| Human | CDK7 | Transcription Elongation | ATACACATCAGGTTGTAACC |
| Human | CDK7 | Transcription Elongation | ATCTCTGGCCTTGTAAACGG |

| | | | |
|---|---|---|---|
| Human | CDK8 | Transcription Elongation | AATATTTCACTGTCGACAAG |
| Human | CDK8 | Transcription Elongation | GGTCATGATAAGTCCACCTG |
| Human | CDK8 | Transcription Elongation | AGTACTCACATCAGACACAT |
| Human | CDK8 | Transcription Elongation | CAAGACAGCAGTCACACACA |
| Human | CDK8 | Transcription Elongation | GCAACACCCAGTTAGCATGC |
| Human | CDK9 | Transcription Elongation | TGCGTATGGGTCACGCACAT |
| Human | CDK9 | Transcription Elongation | TGGTGCCAAGTACTCGAACA |
| Human | CDK9 | Transcription Elongation | GGCAAAGCAGTACGACTCGG |
| Human | CDK9 | Transcription Elongation | CGATGAGGTCCAGTGCGTAT |
| Human | CDK9 | Transcription Elongation | CCAGAGTGTCACCACACGGT |
| Human | CDY1 | Chromatin Readers | ATGCACCTAGTCCAATCGCA |
| Human | CDY1 | Chromatin Readers | CTTAATATTACAGCGAACGA |
| Human | CDY1 | Chromatin Readers | GCGAAAGCTGACAGCAAGGG |
| Human | CDY1 | Chromatin Readers | GACGTGACAAAGGGTCCCGG |
| Human | CDY1 | Chromatin Readers | TCTGCACCAGGACGTGACAA |
| Human | CDY2A | Chromatin Readers | GCGAAAGCTGACAGCACGGG |
| Human | CDY2A | Chromatin Readers | AATATTACTGATGACAGCAG |
| Human | CDY2A | Chromatin Readers | GATGGATGCACCTAGTCCAA |
| Human | CDY2A | Chromatin Readers | ATGCACCTAGTCCAATGGCA |
| Human | CDY2A | Chromatin Readers | TCTGCACCAGGATGTGACAA |
| Human | CDYL | Chromatin Readers | ACATACAACTGGATTGCACG |
| Human | CDYL | Chromatin Readers | GCCCAAACCACATCGCAAAG |
| Human | CDYL | Chromatin Readers | CGACTTCAACAGACGCCACA |
| Human | CDYL | Chromatin Readers | GAATAACTCACTAAATCCAG |
| Human | CDYL | Chromatin Readers | CTGCGGTCCTGCTCTTAACG |
| Human | CDYL2 | Chromatin Readers | GGTTTCGAGACATCGTTGTG |
| Human | CDYL2 | Chromatin Readers | TGTCAGGGCATTGTTATCCG |
| Human | CDYL2 | Chromatin Readers | CCAGTTTAAGAAGCCTATCG |
| Human | CDYL2 | Chromatin Readers | GGCTCTGCTCTGACCAACGG |
| Human | CDYL2 | Chromatin Readers | TGACGTGAATCACGCTACAC |
| Human | CECR2 | Chromatin Readers | TCATTGTAGATCGAGCGAAG |
| Human | CECR2 | Chromatin Readers | AGGTGGTCAGACATCCACCT |
| Human | CECR2 | Chromatin Readers | GGGCGGTCGCATGTACCCTG |
| Human | CECR2 | Chromatin Readers | AGGTTTATACTGTACCAAGG |
| Human | CECR2 | Chromatin Readers | AGGCCAGGCATATACAGCGG |
| Human | CENPB | Chromatin Remodeling | GTACCATAGACTGGTCTCGG |
| Human | CENPB | Chromatin Remodeling | AAGCCCTCGAGCTTGTCGTA |
| Human | CENPB | Chromatin Remodeling | TCTCCCTACGACAAGCTCGA |
| Human | CENPB | Chromatin Remodeling | CATAGCCGCCTGCTTTCGTG |
| Human | CENPB | Chromatin Remodeling | TCTGCACGCGGTCATCAATG |
| Human | CENPC | Chromatin Remodeling | ATCAACTTCGTATCATCGGG |
| Human | CENPC | Chromatin Remodeling | GCGTCCATCTGATTGGTGGG |

| | | | |
|---|---|---|---|
| Human | CENPC | Chromatin Remodeling | GAGATCCTTTGCAGCCAACG |
| Human | CENPC | Chromatin Remodeling | GAAGGTTCTGGAGGTATCGT |
| Human | CENPC | Chromatin Remodeling | TATAAACACTTACCCACAGG |
| Human | CENPE | Chromatin Remodeling | TTGTGTGGTAGGATTAACCT |
| Human | CENPE | Chromatin Remodeling | AACAGAACAGTCCACAACCA |
| Human | CENPE | Chromatin Remodeling | AATTACACCCTCTCAATGCA |
| Human | CENPE | Chromatin Remodeling | GTAGCTCTTGATGTAACATG |
| Human | CENPE | Chromatin Remodeling | CGGCCGCACTCGCACGCAGA |
| Human | CENPF | Chromatin Remodeling | GAAACTGAGAGTTCGCATTG |
| Human | CENPF | Chromatin Remodeling | ATACGACCAGGCGTCAACCA |
| Human | CENPF | Chromatin Remodeling | AGAAAGGTGAGTTCGCATTG |
| Human | CENPF | Chromatin Remodeling | TATCCTGCGAAGAACAACCA |
| Human | CENPF | Chromatin Remodeling | TTCTTTGCCGAGACTCAGTG |
| Human | CENPH | Chromatin Remodeling | GAGTAAAGTCAATTGGGCAG |
| Human | CENPH | Chromatin Remodeling | CGAGCCCGCGGACTCCGGAG |
| Human | CENPH | Chromatin Remodeling | TCGTCGGCGTCTTGCATCTG |
| Human | CENPH | Chromatin Remodeling | GCAAGACGCCGACGAGCCCG |
| Human | CENPH | Chromatin Remodeling | ATTAATAATGAAATCACAGC |
| Human | CENPI | Chromatin Remodeling | GGAACTGTGACCGACAACAA |
| Human | CENPI | Chromatin Remodeling | TGGATAGCCACCCTACATAG |
| Human | CENPI | Chromatin Remodeling | CTCTGAAGCACACTACACTG |
| Human | CENPI | Chromatin Remodeling | AACTGATCCACTATGTAGGG |
| Human | CENPI | Chromatin Remodeling | GAAACAATACCACTAATGGA |
| Human | CENPJ | Chromatin Remodeling | GACTAACCAGAGCACTTCCG |
| Human | CENPJ | Chromatin Remodeling | TAGTCATTGTAACCGCAGTG |
| Human | CENPJ | Chromatin Remodeling | ATCATCAGCTCCGATGTAGG |
| Human | CENPJ | Chromatin Remodeling | CCTGTGCAGCCAGTATCGCA |
| Human | CENPJ | Chromatin Remodeling | GGTCCGGGTAGACATATGAT |
| Human | CENPK | Chromatin Remodeling | ATGTCGAAAGGGAACAACGG |
| Human | CENPK | Chromatin Remodeling | CCGGATAGTACTACAGATGT |
| Human | CENPK | Chromatin Remodeling | GTCTTCTAGAAACTCGCCCA |
| Human | CENPK | Chromatin Remodeling | GAATCAGGAGGATCTAGATC |
| Human | CENPK | Chromatin Remodeling | TAACCGCTGAACTCAGTCAA |
| Human | CENPL | Chromatin Remodeling | TGAAACACGAACTAATCTTG |
| Human | CENPL | Chromatin Remodeling | CATTCCTTCTGCATAAACAG |
| Human | CENPL | Chromatin Remodeling | TTCTTTCGCAATACATCCAG |
| Human | CENPL | Chromatin Remodeling | GATGCACTAGGAGTTGACTC |
| Human | CENPL | Chromatin Remodeling | AGAAACGATTAGAATCGGTC |
| Human | CENPM | Chromatin Remodeling | AACACGATCAGGTCAATTCG |
| Human | CENPM | Chromatin Remodeling | CAGCATTCACCGGCACACCG |
| Human | CENPM | Chromatin Remodeling | CGTACTTACCACCTGTGGCG |
| Human | CENPM | Chromatin Remodeling | AGAGGAGTCCCTGCGCCATG |

| | | | |
|---|---|---|---|
| Human | CENPM | Chromatin Remodeling | GGCTACTTACACCTTCAGCT |
| Human | CENPN | Chromatin Remodeling | CCAGTACAAACCTACCTACG |
| Human | CENPN | Chromatin Remodeling | CTGGGAGTAGTACACCACGT |
| Human | CENPN | Chromatin Remodeling | GCACTTGATCCATCTGTGTG |
| Human | CENPN | Chromatin Remodeling | TGTATTGCGCCTCAGCATGG |
| Human | CENPN | Chromatin Remodeling | AGTCTGGATTCGAATTGCCT |
| Human | CENPO | Chromatin Remodeling | AATGACAGCAAGTTACACAG |
| Human | CENPO | Chromatin Remodeling | CCAGATGGCGAGTCCAAAGG |
| Human | CENPO | Chromatin Remodeling | ACATGTCACGGTGACGTCAG |
| Human | CENPO | Chromatin Remodeling | CTTCTTACCTTGACATGTCA |
| Human | CENPO | Chromatin Remodeling | TACTTACAAACTGGATCCAG |
| Human | CENPP | Chromatin Remodeling | GCGCACGTTTAAACATCTCA |
| Human | CENPP | Chromatin Remodeling | CTCAGCTTCCGAACCCTGGT |
| Human | CENPP | Chromatin Remodeling | ATACACCAATTCAATTTGGA |
| Human | CENPP | Chromatin Remodeling | CTCACCAAAGTCCCACAGCG |
| Human | CENPP | Chromatin Remodeling | ATGCCGTGTACCTCTCGGAG |
| Human | CENPQ | Chromatin Remodeling | ATGCATCAAATAAATAGTAG |
| Human | CENPQ | Chromatin Remodeling | AGAAGGTCTGGCATTACTAC |
| Human | CENPQ | Chromatin Remodeling | AGACCACAGAGTTAATGACT |
| Human | CENPQ | Chromatin Remodeling | GAGACCACAGAGTTAATGAC |
| Human | CENPQ | Chromatin Remodeling | TGTGTTTAGATCAAGATGTC |
| Human | CENPT | Chromatin Remodeling | CAGGGAGTGATACACGTTGG |
| Human | CENPT | Chromatin Remodeling | ACTTGGAGGAACAGACACCT |
| Human | CENPT | Chromatin Remodeling | AGACTGTCAGTGTTTCAGCA |
| Human | CENPT | Chromatin Remodeling | TCTGCCCACAGACATTGTGT |
| Human | CENPT | Chromatin Remodeling | GAGTTGCCACTGTATGCACA |
| Human | CENPV | Chromatin Remodeling | GCGGCAGAAGCTTACCTCCG |
| Human | CENPV | Chromatin Remodeling | CGGCAGAAGCTTACCTCCGA |
| Human | CENPV | Chromatin Remodeling | TCGGCGTCCAACCTGGACCT |
| Human | CENPV | Chromatin Remodeling | AGCCGCGGCTTCTCCGACGG |
| Human | CENPV | Chromatin Remodeling | GGCCGGGAGCAAGAGCCAGG |
| Human | CENPW | Chromatin Remodeling | ACTCGCTTTAGAAAGCCACG |
| Human | CENPW | Chromatin Remodeling | AGACTCGCTTTAGAAAGCCA |
| Human | CENPW | Chromatin Remodeling | CATTAACAAGGAGCATGTAC |
| Human | CENPW | Chromatin Remodeling | ACTCGCACAAGCGTTTGTCC |
| Human | CENPW | Chromatin Remodeling | TGTCGACCATAGTCTCCCAG |
| Human | CHAF1A | DNA and RNA Modifications | TCCTCAGACAGGTACCCATG |
| Human | CHAF1A | DNA and RNA Modifications | TTCCGGGTGGAAACGTGCGT |
| Human | CHAF1A | DNA and RNA Modifications | CCAAAGAGGCCTCTAAATCG |
| Human | CHAF1A | DNA and RNA Modifications | GAAAGCGGTCCGACGCCGAG |
| Human | CHAF1A | DNA and RNA Modifications | GATGACATGTCAGACGATCA |
| Human | CHD1 | Chromatin Readers | TTAATTCGCCTAAGAGAACG |

| | | | |
|---|---|---|---|
| Human | CHD1 | Chromatin Readers | TTCTCCCGAGGAATAGTCCG |
| Human | CHD1 | Chromatin Readers | TTCCGATGACTCATCAAGTG |
| Human | CHD1 | Chromatin Readers | GAGACAGATTGATTCATCTG |
| Human | CHD1 | Chromatin Readers | CATCAAGCCTCATCTAATAG |
| Human | CHD2 | Chromatin Readers | AAGAAAGACGACGTGACTGG |
| Human | CHD2 | Chromatin Readers | GTCTTACATCCACAGCACAT |
| Human | CHD2 | Chromatin Readers | GTGGCAACAACAATCCACCA |
| Human | CHD2 | Chromatin Readers | AAGGGACGACTATAAGAAAG |
| Human | CHD2 | Chromatin Readers | ACAACCTGCATATTTAGGAG |
| Human | CHD3 | Chromatin Readers | TTTCCACTCACTGTGCATCG |
| Human | CHD3 | Chromatin Readers | AGCGCTTAGAATCGGCGCTG |
| Human | CHD3 | Chromatin Readers | GTAGCCATGGCTATGCACGG |
| Human | CHD3 | Chromatin Readers | ACGGTCCAATGGGCGACGAG |
| Human | CHD3 | Chromatin Readers | ACCATATGACCAGTCCACGT |
| Human | CHD4 | Chromatin Readers | CAGTCACTCACCGTACAACG |
| Human | CHD4 | Chromatin Readers | TACTTACTGGAACCTCTCGG |
| Human | CHD4 | Chromatin Readers | CAATGGCCTTATCATCGTAG |
| Human | CHD4 | Chromatin Readers | GAGTACCTGGATGCTACAGG |
| Human | CHD4 | Chromatin Readers | TGATGTTCCGAAACTATCAG |
| Human | CHD5 | Chromatin Readers | CGTTCTCCCGAATCACCGAG |
| Human | CHD5 | Chromatin Readers | GTGGACGACGACAATCCCGG |
| Human | CHD5 | Chromatin Readers | CCAGAGCCGTAGTCAAAGGG |
| Human | CHD5 | Chromatin Readers | TGGGCTTCAATGCCCGACAG |
| Human | CHD5 | Chromatin Readers | CCGAGGCCCGAGTCACGAAG |
| Human | CHD6 | Chromatin Readers | AACAGCTCCAAGTCGAACGG |
| Human | CHD6 | Chromatin Readers | CTAGGCGGTTAATTATCACG |
| Human | CHD6 | Chromatin Readers | CAGGACATCAACCGAAAGGG |
| Human | CHD6 | Chromatin Readers | CCGAAAGGGCGGCACCAATG |
| Human | CHD6 | Chromatin Readers | CCTAGATAATATCTGCCACG |
| Human | CHD7 | Chromatin Readers | GTTCTTCGCATCGCCTCCGG |
| Human | CHD7 | Chromatin Readers | GGATCTGATCACACCCACAG |
| Human | CHD7 | Chromatin Readers | TGGGCAGCTATATGGCACGT |
| Human | CHD7 | Chromatin Readers | AGGTTCAGGGATCCACACAT |
| Human | CHD7 | Chromatin Readers | TAATGGTGTGGGTTCGACGT |
| Human | CHD8 | Chromatin Readers | CTTTAATCCAGACTACGTAG |
| Human | CHD8 | Chromatin Readers | CTGTCTTCTACACTACCGTG |
| Human | CHD8 | Chromatin Readers | CACGGCGGGAGAAACAACAA |
| Human | CHD8 | Chromatin Readers | CATACCTCGAGTCTTAAATG |
| Human | CHD8 | Chromatin Readers | ACTGTCTTCTACACTACCGT |
| Human | CHD9 | Chromatin Readers | GGGAGAGAGAATTTCGTACG |
| Human | CHD9 | Chromatin Readers | TAGCCATTGGAACGGTCACA |
| Human | CHD9 | Chromatin Readers | TTAGATGCATTAGTCATCTG |

| | | | |
|---|---|---|---|
| Human | CHD9 | Chromatin Readers | AATGTTGACTCATCAAAGTG |
| Human | CHD9 | Chromatin Readers | ACAGAAATAATCTCAACCCA |
| Human | CHEK1 | Other Histone Modifications | ACACCACCTGAAGTGACTCG |
| Human | CHEK1 | Other Histone Modifications | TGGTATTGGAATAACTCACA |
| Human | CHEK1 | Other Histone Modifications | AGCATATGATCAGGACATGT |
| Human | CHEK1 | Other Histone Modifications | CCAATTTATCAATGTATGAG |
| Human | CHEK1 | Other Histone Modifications | TCTTCCATCAACTCATGGCA |
| Human | CHMP2A | Control | TCTCTAGTTTCTGTCGCTCG |
| Human | CHMP2A | Control | ACTCAAGTCCAACAACTCGA |
| Human | CHMP2A | Control | AGGAACGGCTTAAGAACCTG |
| Human | CHMP2A | Control | AACGGCTTAAGAACCTGCGG |
| Human | CHMP2A | Control | AGACGCCAGAGGAGCTACTG |
| Human | CHP1 | Chromatin Readers | AGAGGCGAGTGATTTGACTG |
| Human | CHP1 | Chromatin Readers | TTCGTCCCGCAGTAACGTGG |
| Human | CHP1 | Chromatin Readers | AACTTGCCATCAACCCACTG |
| Human | CHP1 | Chromatin Readers | CAGGTGCTACGCATGATGGT |
| Human | CHP1 | Chromatin Readers | ACTTGCCATCAACCCACTGG |
| Human | CHRAC1 | Chromatin Remodeling | GCCACCTATTCCTACAGACA |
| Human | CHRAC1 | Chromatin Remodeling | TATTCCTACAGACACGGCAG |
| Human | CHRAC1 | Chromatin Remodeling | GTCGTGGGTAAAGACAAGGG |
| Human | CHRAC1 | Chromatin Remodeling | AAGATGGCGGACGTGGTCGT |
| Human | CHRAC1 | Chromatin Remodeling | GTCTGTAGGAATAGGTGGCT |
| Human | CLOCK | Histone Acetylation | GAACTATACAACGCACACAT |
| Human | CLOCK | Histone Acetylation | GGCAAATACCCTATTATGGG |
| Human | CLOCK | Histone Acetylation | TTCTGTTGTCACATGCTGCG |
| Human | CLOCK | Histone Acetylation | AAACATACCTTGATGAACTG |
| Human | CLOCK | Histone Acetylation | AGGCTATGATTACTATCATG |
| Human | COPZ1 | Control | GGAGAGTCATACCTGAGACA |
| Human | COPZ1 | Control | ACACGGTCTGCTCCGTAAGG |
| Human | COPZ1 | Control | GCTTTGACAGTATACAGGGA |
| Human | COPZ1 | Control | GATGGCTTTGACAGTATACA |
| Human | COPZ1 | Control | ATGACCTACTGTCAGTCCGA |
| Human | CREBBP | Histone Acetylation | CAACTGTCGGAGCTTCTACG |
| Human | CREBBP | Histone Acetylation | AGTACCGAGAAATGTTACGG |
| Human | CREBBP | Histone Acetylation | GATGAGCTGATACCCAATGG |
| Human | CREBBP | Histone Acetylation | TAATTAATCAGGCTTCACAA |
| Human | CREBBP | Histone Acetylation | GGCTCGGCAGATCGAGCGTG |
| Human | CSNK2A1 | Other Histone Modifications | TGAGGATAGCCAAGGTTCTG |
| Human | CSNK2A1 | Other Histone Modifications | CTAGTTGGTGAGGATAGCCA |
| Human | CSNK2A1 | Other Histone Modifications | AGTCACATGTGGTGGAATGG |
| Human | CSNK2A1 | Other Histone Modifications | CACCTACTATACTCACCACA |
| Human | CSNK2A1 | Other Histone Modifications | GATTGATCATGAGCACAGAA |

| | | | |
|---|---|---|---|
| Human | CSNK2A2 | Other Histone Modifications | AGTTTACCTGATAGTCCACG |
| Human | CSNK2A2 | Other Histone Modifications | ACAATGTTCGTGTAGCCTCA |
| Human | CSNK2A2 | Other Histone Modifications | CCCAGGATATCGTTGAAGTG |
| Human | CSNK2A2 | Other Histone Modifications | TGGTCCCTTGAAGTACCTTG |
| Human | CSNK2A2 | Other Histone Modifications | CAGACATTCACGGAAACGCT |
| Human | CSNK2B | Other Histone Modifications | CGCTACATCCTTACCAACCG |
| Human | CSNK2B | Other Histone Modifications | CTAGAGCTTGTCGATAGTGA |
| Human | CSNK2B | Other Histone Modifications | TGTACACATCCATGCACTTG |
| Human | CSNK2B | Other Histone Modifications | TGGTTCTCACAGTACACACG |
| Human | CSNK2B | Other Histone Modifications | GTGTACACATCCATGCACTT |
| Human | CTDP1 | Transcription Elongation | AGTCGTAGATCCGAACCTCG |
| Human | CTDP1 | Transcription Elongation | GGATCTACGACTCCAACACG |
| Human | CTDP1 | Transcription Elongation | AAATCATTCTCGAGGCACTG |
| Human | CTDP1 | Transcription Elongation | TCGGATCTACGACTCCAACA |
| Human | CTDP1 | Transcription Elongation | TGGGAGCGAAGATCCTCACT |
| Human | CTR9 | Transcription Elongation | TCAGATCACGAATCGGAGAG |
| Human | CTR9 | Transcription Elongation | GTTGCTAGGATCAATAGTAT |
| Human | CTR9 | Transcription Elongation | AGAAGCTATTACCATCTGAA |
| Human | CTR9 | Transcription Elongation | TGCCGAATCAGATCACGAAT |
| Human | CTR9 | Transcription Elongation | TGACTTCGATCAGTTACCGG |
| Human | CUL1 | Control | TCGCTTCATAAACAACAACG |
| Human | CUL1 | Control | GCATCCAGCAGGTGTACACA |
| Human | CUL1 | Control | GTTCGCCGTGAATGTGACGA |
| Human | CUL1 | Control | AGTATCGAGCCAGCAACTCA |
| Human | CUL1 | Control | GAGACACACATTCATAATCA |
| Human | CXXC1 | Chromatin Readers | AGAAGGAGGAGCGATACAAG |
| Human | CXXC1 | Chromatin Readers | GGCTGCGGAGAGGATTTGTG |
| Human | CXXC1 | Chromatin Readers | CCAAGCTAGAGATTCGCTAT |
| Human | CXXC1 | Chromatin Readers | TCAGGGCCAGCAATCCCGCG |
| Human | CXXC1 | Chromatin Readers | TGTGGCGCAAGGCAACACGT |
| Human | DAPK3 | Other Histone Modifications | CGTGAACTACGACTTCGACG |
| Human | DAPK3 | Other Histone Modifications | TCGCGCACAAGATCGAGGCG |
| Human | DAPK3 | Other Histone Modifications | CTCCTTCCTAGGCGATCCGG |
| Human | DAPK3 | Other Histone Modifications | GCGGCCAGTTTGCGATCGTG |
| Human | DAPK3 | Other Histone Modifications | GGAGGACCATTATGAGATGG |
| Human | DCTN2 | Control | TGATCAAGCGGAGTTCGATG |
| Human | DCTN2 | Control | GTAGGGTACCCACTTAGCCA |
| Human | DCTN2 | Control | ACTTAGCCAGGGCGCCATCG |
| Human | DCTN2 | Control | GGAGCTGACAAGCACAAGTG |
| Human | DCTN2 | Control | GCAACAAAGAACAGCAAAGG |
| Human | DDOST | Control | CAGGTCGGACTCACCAATGT |
| Human | DDOST | Control | CCCAGTATCCACATGCGGTG |

| | | | |
|---|---|---|---|
| Human | DDOST | Control | GAACCCTTACCTTCTACCGA |
| Human | DDOST | Control | CCTGTCTCTCATAAAGTATG |
| Human | DDOST | Control | GTGCGGGATTGAGTTTGACG |
| Human | DDX39B | Transcription Elongation | AGTTATGGCAGAGAACGATG |
| Human | DDX39B | Transcription Elongation | CTCAAAGCCACAGTCGACAA |
| Human | DDX39B | Transcription Elongation | CCTGCCAAGAAGGATGTCAA |
| Human | DDX39B | Transcription Elongation | AGTCACGAAAGCCAGAGCTG |
| Human | DDX39B | Transcription Elongation | CAGGACATCCATTCCCAGAA |
| Human | DHPS | Control | TAAGTGCGGGACTAAACACA |
| Human | DHPS | Control | GTGCGCGGTAATTCACACCG |
| Human | DHPS | Control | GTGCTGCACAAGGTAGCGAA |
| Human | DHPS | Control | CCTTGTGCAGCACAACATGG |
| Human | DHPS | Control | TGAGGTTGGATGTATATCCC |
| Human | DIDO1 | Chromatin Readers | GCCATCGCTGTCACTATCGG |
| Human | DIDO1 | Chromatin Readers | AGCTGAAGCACCACTCGATG |
| Human | DIDO1 | Chromatin Readers | GCAAGAGTTTGGATGGAACA |
| Human | DIDO1 | Chromatin Readers | GGCTTAGCTAGTATGGACTT |
| Human | DIDO1 | Chromatin Readers | CTGCGCGGAGTGAAGCACTC |
| Human | DIMT1 | Control | AACAGCATTATCGATAAGGT |
| Human | DIMT1 | Control | CTTCAGACCACCGCCCAAGG |
| Human | DIMT1 | Control | GAAGTTCAGCTACTAGCCTT |
| Human | DIMT1 | Control | ACAGGACTCATGTTCAACAC |
| Human | DIMT1 | Control | ACTTCCAGCACTACATCAGT |
| Human | DMAP1 | DNA and RNA Modifications | ATGCTGGGCACGAACGACGG |
| Human | DMAP1 | DNA and RNA Modifications | GGACATTATCAACCCGGACA |
| Human | DMAP1 | DNA and RNA Modifications | TTGATGCTGGGCACGAACGA |
| Human | DMAP1 | DNA and RNA Modifications | ACGGGACATTCTAGAACTCG |
| Human | DMAP1 | DNA and RNA Modifications | GCTGAGCCCGACACCTACGG |
| Human | DNA2 | DNA and RNA Modifications | AATTGGTACGGTTCTCCATG |
| Human | DNA2 | DNA and RNA Modifications | GGGTTAACATTAGACACCAA |
| Human | DNA2 | DNA and RNA Modifications | GGAGGATAGCAATTTAGTGA |
| Human | DNA2 | DNA and RNA Modifications | CTAAGGTAGGATATAAACTG |
| Human | DNA2 | DNA and RNA Modifications | ACTGTGAAGCAGTGATGACC |
| Human | DNMT1 | DNA and RNA Modifications | CACCTGTGTCCCATCGACAC |
| Human | DNMT1 | DNA and RNA Modifications | CCTCCCAAGTGCATTCAGTG |
| Human | DNMT1 | DNA and RNA Modifications | CCTCTATGGAAGGCTCGAGT |
| Human | DNMT1 | DNA and RNA Modifications | CCCATGGTACCTACCGCCTG |
| Human | DNMT1 | DNA and RNA Modifications | CAGGTCCTCATAGGTCGAGT |
| Human | DNMT3A | Chromatin Readers | GCGTACCAGTACGACGACGA |
| Human | DNMT3A | Chromatin Readers | CGGAGACGGCAAATTCTCAG |
| Human | DNMT3A | Chromatin Readers | TGTGAGGACTCCATCACGGT |
| Human | DNMT3A | Chromatin Readers | AGGTCAAACTCCATAAAGCA |

| | | | |
|---|---|---|---|
| Human | DNMT3A | Chromatin Readers | CGATGACGAGCCAGAGTACG |
| Human | DNMT3B | Chromatin Readers | AGTCGAACTCGATCAAACAG |
| Human | DNMT3B | Chromatin Readers | GGCCTTCCAAGACACCACCA |
| Human | DNMT3B | Chromatin Readers | AGACTCGATCCTCGTCAACG |
| Human | DNMT3B | Chromatin Readers | ACATTTGAGAGATCGTTGCA |
| Human | DNMT3B | Chromatin Readers | CCCAACAACACGCAACCAGG |
| Human | DNMT3L | DNA and RNA Modifications | TGTTCGAAACCGTGCCTGTG |
| Human | DNMT3L | DNA and RNA Modifications | AGATCCCTCCCTCAAACAGA |
| Human | DNMT3L | DNA and RNA Modifications | GTAGCACACCCAGTTGCTCA |
| Human | DNMT3L | DNA and RNA Modifications | GTCATCGTCGTACAGGAAGA |
| Human | DNMT3L | DNA and RNA Modifications | GGTGCACGCCATGAGCAACT |
| Human | DOLK | Control | CCGTATGGGACCGATACTCG |
| Human | DOLK | Control | TTGGACTAGTCATGAAGGAG |
| Human | DOLK | Control | AGCCGCCACTGTATGCCTGG |
| Human | DOLK | Control | CTTCTACGTCCAATACAAGT |
| Human | DOLK | Control | CCGATACTCGTGGTGCGCCG |
| Human | DOT1L | Histone Methylation | GGGAGCGAATCGCCAACACG |
| Human | DOT1L | Histone Methylation | GCAGCAGGTCTACAACCACT |
| Human | DOT1L | Histone Methylation | GTCCACAAACAGGTCGTCGT |
| Human | DOT1L | Histone Methylation | GCTGGAGGACTCATCCAAGG |
| Human | DOT1L | Histone Methylation | GAGATGGCTGTAATCTCCAG |
| Human | DPF1 | Chromatin Readers | GAGGCACAAAGGAAACACAC |
| Human | DPF1 | Chromatin Readers | GTTTCCGCATGACCTCGAGG |
| Human | DPF1 | Chromatin Readers | CCTGCGAGTACAAGATCGGT |
| Human | DPF1 | Chromatin Readers | GTACGTGTAAATCTGTCCCG |
| Human | DPF1 | Chromatin Readers | CTGCACACAGTAGAGCCTCG |
| Human | DPF2 | Chromatin Readers | GAAGATACTCCCAAGCGTCG |
| Human | DPF2 | Chromatin Readers | ATAGATGGGAAGGAAAGTCG |
| Human | DPF2 | Chromatin Readers | GATACACACGTCATTCTCGG |
| Human | DPF2 | Chromatin Readers | CAAGAAGACGGGACAACCCG |
| Human | DPF2 | Chromatin Readers | AAGAAGATACTCCCAAGCGT |
| Human | DPF3 | Chromatin Readers | CAAGTAGGCACTCACGCCTG |
| Human | DPF3 | Chromatin Readers | GTGAACCCATCCTTCTTCAG |
| Human | DPF3 | Chromatin Readers | AAGCGAAAGAACAGGACTAG |
| Human | DPF3 | Chromatin Readers | AGCGCTACAAGAACCGACCG |
| Human | DPF3 | Chromatin Readers | ACAGCCGTGAGTTGTAACTC |
| Human | DSN1 | Chromatin Remodeling | GATGAATCAAGGCGTTTCAG |
| Human | DSN1 | Chromatin Remodeling | AGGCTAAAGAGATATTGTCC |
| Human | DSN1 | Chromatin Remodeling | GTATGAAAGAAACGAACCGG |
| Human | DSN1 | Chromatin Remodeling | AACGCCTTGATTCATCTCCA |
| Human | DSN1 | Chromatin Remodeling | GGAATCAAGTCTCAGTCCTG |
| Human | DYNLRB1 | Control | AGGAAGGTGAGATCGTTCTG |

| | | | |
|---|---|---|---|
| Human | DYNLRB1 | Control | AAGAGAGTGGCTTATTCGGT |
| Human | DYNLRB1 | Control | CTTGGAGCGAATTCGAAGGA |
| Human | DYNLRB1 | Control | AATGAAATTATGGTTGCACC |
| Human | DYNLRB1 | Control | TCAGGATGAAGCTGTGCATG |
| Human | EAF1 | Transcription Elongation | ACCGCGAGGAACATTGCCTG |
| Human | EAF1 | Transcription Elongation | GCCTCAGGCAATGTTCCTCG |
| Human | EAF1 | Transcription Elongation | ACCAGCAGCCCTACAACAGT |
| Human | EAF1 | Transcription Elongation | ACAACGAATAGTGTGGAAGG |
| Human | EAF1 | Transcription Elongation | CCCTTGAACACAGTCATGGG |
| Human | EAF2 | Transcription Elongation | AATAGATTTCGAGACAACAG |
| Human | EAF2 | Transcription Elongation | CGAGCGGGTTCTCAAGTTAG |
| Human | EAF2 | Transcription Elongation | ACTTGAGAACCCGCTCGCGA |
| Human | EAF2 | Transcription Elongation | CACACCTAGACCGTCGCGAG |
| Human | EAF2 | Transcription Elongation | ATCCTACCATGACACAGTAC |
| Human | EAPP | Transcription Elongation | GGTTGAAGAGCCTAGCGACG |
| Human | EAPP | Transcription Elongation | AAGTTGCAACAGCTCCGACA |
| Human | EAPP | Transcription Elongation | TAGGCTCTTCAACCGCGTAG |
| Human | EAPP | Transcription Elongation | TGAAGTGGCAGTCTACGACA |
| Human | EAPP | Transcription Elongation | AAGTTTACCAGTTCCCAGAG |
| Human | EHMT1 | Histone Methylation | GGTACTGGACCACTTCGTAG |
| Human | EHMT1 | Histone Methylation | GCTGGCGTTATTGACTCGAG |
| Human | EHMT1 | Histone Methylation | TTGCATTCGAAGATCAAGGG |
| Human | EHMT1 | Histone Methylation | GAGTCGAGAGATCACCACAC |
| Human | EHMT1 | Histone Methylation | AGGTTTACTGCATCGACGCG |
| Human | EHMT2 | Histone Methylation | TTGAAGCGCAAACCACACGT |
| Human | EHMT2 | Histone Methylation | CCATGCTGTCAACTACCATG |
| Human | EHMT2 | Histone Methylation | CTCCAGGTGGTTGTTCACCA |
| Human | EHMT2 | Histone Methylation | CGGACAGGTACAACTGCCGA |
| Human | EHMT2 | Histone Methylation | ATCCGCATGCTACTGACGCG |
| Human | EIF2B1 | Control | GAGGACGCCATGGACGACAA |
| Human | EIF2B1 | Control | GGCACTTACGGAGTATTCCA |
| Human | EIF2B1 | Control | AGTCACAGCCTGATTTGTCA |
| Human | EIF2B1 | Control | CAGAGGAGTCCACACCACAC |
| Human | EIF2B1 | Control | ACCTCAAAGAGGAGCATCCG |
| Human | EIF5B | Control | TCTGAGTTAACTCTACAAGA |
| Human | EIF5B | Control | TTGATAGACTCAATTGTCTG |
| Human | EIF5B | Control | TACTTGGGCATGTGGACACA |
| Human | EIF5B | Control | TGGTATCACACAACAAATTG |
| Human | EIF5B | Control | CAGGTCCTAACATAGAAAGT |
| Human | EIF6 | Control | CGATAGACGCGTGCACCACG |
| Human | EIF6 | Control | GGTCCACCCAGACTTGGACA |
| Human | EIF6 | Control | TGTCCCTACAGTGTGTTCGA |

| | | | |
|---|---|---|---|
| Human | EIF6 | Control | CAGGAATGCTTACCACAAGG |
| Human | EIF6 | Control | TCCGATCGCTACCAGACAGT |
| Human | ELL | Transcription Elongation | TGAGTGAAGTGCGATATCCG |
| Human | ELL | Transcription Elongation | GCCGTCCTTAGCACTCATGT |
| Human | ELL | Transcription Elongation | GTAGTCAGGCGTCTCCGACG |
| Human | ELL | Transcription Elongation | GGCACTCGCCAAGTTGATGG |
| Human | ELL | Transcription Elongation | TCATCGCCGAGTACGACCAG |
| Human | ELL2 | Transcription Elongation | CTTTAGTAGAACGGAACCAG |
| Human | ELL2 | Transcription Elongation | CTATGTTTATGACATACCTG |
| Human | ELL2 | Transcription Elongation | ACTTTAGTAGAACGGAACCA |
| Human | ELL2 | Transcription Elongation | GACACGAGAAAGAATGACCC |
| Human | ELL2 | Transcription Elongation | AATCCTTTAAGGTATATGAG |
| Human | ELL3 | Transcription Elongation | GATCCACTGGCAAGCAACCA |
| Human | ELL3 | Transcription Elongation | CAGGATGAGACCTTTAATGT |
| Human | ELL3 | Transcription Elongation | AGGATCCTCAAGGGAGCACA |
| Human | ELL3 | Transcription Elongation | TGGCCTTGGAAAGCAATCAC |
| Human | ELL3 | Transcription Elongation | CCTGAGAGTTGGCAGAACAC |
| Human | ELOA | Transcription Elongation | CGTGGCGAGAGATGTACCTG |
| Human | ELOA | Transcription Elongation | GTTAAAGCTGATGCTACTGG |
| Human | ELOA | Transcription Elongation | TGGTCGACGTACATCTGATG |
| Human | ELOA | Transcription Elongation | GACCCGAAGAGTATGAGTCG |
| Human | ELOA | Transcription Elongation | CTCAGTTCAGAAATTACCCA |
| Human | ELOA2 | Transcription Elongation | ACGAATATCCGATCTGCACG |
| Human | ELOA2 | Transcription Elongation | GATCAGGGAGAAATCATGCG |
| Human | ELOA2 | Transcription Elongation | ATGGAGTCAGAATCCGAAAG |
| Human | ELOA2 | Transcription Elongation | GAGAGACAGACGAATTACGG |
| Human | ELOA2 | Transcription Elongation | CTCTTCACCAAAGTTCCGGG |
| Human | ELOA3B | Transcription Elongation | CGGGTTCAAGAACCGAGTAG |
| Human | ELOA3B | Transcription Elongation | CATGACCGCAGACATCCTGG |
| Human | ELOA3B | Transcription Elongation | GTCTGCGGTCATGGGCAAGG |
| Human | ELOA3B | Transcription Elongation | AGTCTCCGCCAGGATGTCTG |
| Human | ELOA3B | Transcription Elongation | AACGGGTTCAAGAACCGAGT |
| Human | ELOA3D | Transcription Elongation | CTCCGCGCCAACGCTCCAGG |
| Human | ELOA3D | Transcription Elongation | CGGGTTCAAGAGCCGAGTAG |
| Human | ELOA3D | Transcription Elongation | AACGGGTTCAAGAGCCGAGT |
| Human | ELOA3D | Transcription Elongation | TTCCTCCTGGAGCGTTGGCG |
| Human | ELOA3D | Transcription Elongation | TTGTCTTTCTCTGTGCGGTA |
| Human | ELOB | Transcription Elongation | TTTGACAGCATGTTCAACGA |
| Human | ELOB | Transcription Elongation | GTTTGACAGCATGTTCAACG |
| Human | ELOB | Transcription Elongation | ATGCTGTCAAACCAGGACAC |
| Human | ELOB | Transcription Elongation | CACAGGTCCACCCACGCTGG |
| Human | ELOB | Transcription Elongation | GTTGAACATGCTGTCAAACC |

| | | | |
|---|---|---|---|
| Human | ELOC | Transcription Elongation | AATTGCACCTGAAATTGCAC |
| Human | ELOC | Transcription Elongation | AGGTCCTTCACAGCCACCAT |
| Human | ELOC | Transcription Elongation | ACTTTCGATAGCACATGTGA |
| Human | ELOC | Transcription Elongation | TGGGAATTCAGGAATCTCGG |
| Human | ELOC | Transcription Elongation | TTACTTTACTTTAGATGGAG |
| Human | ELOF1 | Transcription Elongation | TGATTGGATAGACGCCTGCG |
| Human | ELOF1 | Transcription Elongation | CTCTTGTACCGTGTGCCTAG |
| Human | ELOF1 | Transcription Elongation | TCACATCACAGGATTTCTCG |
| Human | ELOF1 | Transcription Elongation | CTGGAATTCCTCTAGGCACA |
| Human | ELOF1 | Transcription Elongation | CACGGTACAAGAGATGACTC |
| Human | ELP1 | Transcription Elongation | TCTCTCTCCGAACTGAACAG |
| Human | ELP1 | Transcription Elongation | CAGCAATCATGTGTCCCATG |
| Human | ELP1 | Transcription Elongation | GGAGCACATGTATGAGCCAG |
| Human | ELP1 | Transcription Elongation | AAATGGAAACTAATTATCAG |
| Human | ELP1 | Transcription Elongation | TGGACGACTGACCGGAGCGT |
| Human | ELP2 | Transcription Elongation | CAGTTTGACAGTCACGCAGA |
| Human | ELP2 | Transcription Elongation | CCAGTACCAATATTAGCATG |
| Human | ELP2 | Transcription Elongation | AATTTCATGCCAAGTCACCT |
| Human | ELP2 | Transcription Elongation | TCCAAAGGTGCAACGCTCCG |
| Human | ELP2 | Transcription Elongation | TATTTGATAATCCCAATGCA |
| Human | ELP3 | Histone Acetylation | CAGATTACAAGGCCCGTACA |
| Human | ELP3 | Histone Acetylation | TATTCCACCCAGTCTTACAC |
| Human | ELP3 | Histone Acetylation | TCCGAGCCACCAATTCAACC |
| Human | ELP3 | Histone Acetylation | GTCCCACGAATCACCAGGGT |
| Human | ELP3 | Histone Acetylation | TACTTCCAACAATATTTACG |
| Human | ELP4 | Transcription Elongation | GCCAGCGTGACCAACGACAG |
| Human | ELP4 | Transcription Elongation | ATCTGGCAGAATCCGCCAAG |
| Human | ELP4 | Transcription Elongation | CCCGCAATGGACACCAGTCG |
| Human | ELP4 | Transcription Elongation | CAGATCAAGTTATTAAGCCG |
| Human | ELP4 | Transcription Elongation | AAGGAAGCTATTCACCATTG |
| Human | ENY2 | Transcription Elongation | ACACGTTACTGTTGATGACT |
| Human | ENY2 | Transcription Elongation | TCTAATTCAATCTTAAAGGC |
| Human | ENY2 | Transcription Elongation | ATTAATTGAATGTGGCTGGA |
| Human | ENY2 | Transcription Elongation | CAGTTGAAGGCACACTGTAA |
| Human | ENY2 | Transcription Elongation | AAAGGCTGGCATGCTGAGCA |
| Human | EP300 | Histone Acetylation | GGTACGACTAGGTACAGGCG |
| Human | EP300 | Histone Acetylation | ATGCAACAGCCTTCCAACCA |
| Human | EP300 | Histone Acetylation | AGGCCCAAATGCTTCGCAGG |
| Human | EP300 | Histone Acetylation | TTCTTCATTGTGCGACAGTG |
| Human | EP300 | Histone Acetylation | ATGGTGAACCATAAGGATTG |
| Human | EPC2 | Chromatin Remodeling | GATGTATAGGATTTGCAAGG |
| Human | EPC2 | Chromatin Remodeling | AAGTCGCAGTGAGGTAGCCA |

| | | | |
|---|---|---|---|
| Human | EPC2 | Chromatin Remodeling | TGGGATTAAGTGGTTTGTAG |
| Human | EPC2 | Chromatin Remodeling | ACCTGTTCCAAGTCGCAGTG |
| Human | EPC2 | Chromatin Remodeling | GGATGATAAAGCCATATTGG |
| Human | ERCC2 | Transcription Elongation | TCCTTGCTCGATACTCAGTG |
| Human | ERCC2 | Transcription Elongation | GCGTACCTGCTCATACCAGG |
| Human | ERCC2 | Transcription Elongation | GTAGCTATAAACCACCACAT |
| Human | ERCC2 | Transcription Elongation | TCCTCACTGAGTATCGAGCA |
| Human | ERCC2 | Transcription Elongation | TTTGGCGTAGGTGCTGACAA |
| Human | ERCC3 | Transcription Elongation | ACATGGAGTACCACTCATCG |
| Human | ERCC3 | Transcription Elongation | TCTACGGACCTACGTCTCAG |
| Human | ERCC3 | Transcription Elongation | AAATGGCTTACTCAACCAAG |
| Human | ERCC3 | Transcription Elongation | CTACGGACCTACGTCTCAGG |
| Human | ERCC3 | Transcription Elongation | AGGGCATGCTTACCACCCAG |
| Human | ERCC6 | DNA and RNA Modifications | AGACAGAATGATCCGATGAG |
| Human | ERCC6 | DNA and RNA Modifications | ACTGATTACGAGATACAATG |
| Human | ERCC6 | DNA and RNA Modifications | GGGTGAAGGAATTTCACACG |
| Human | ERCC6 | DNA and RNA Modifications | AGAATTGCCACTCTGAACGG |
| Human | ERCC6 | DNA and RNA Modifications | CCCAGACTGGAACCCAAGCA |
| Human | ESCO1 | DNA and RNA Modifications | AAGGCTACAACAATTAACAG |
| Human | ESCO1 | DNA and RNA Modifications | TGGAAACACGCATGAGTACA |
| Human | ESCO1 | DNA and RNA Modifications | TTAGGCAAAGAATGATTTCG |
| Human | ESCO1 | DNA and RNA Modifications | AATTGCGGAACATATCCAAT |
| Human | ESCO1 | DNA and RNA Modifications | AGTAGTTGGCTGCCTAATTG |
| Human | ESCO2 | DNA and RNA Modifications | ATTATATACGAGGAAATTAG |
| Human | ESCO2 | DNA and RNA Modifications | AGTGATCTATAAGCCAATTG |
| Human | ESCO2 | DNA and RNA Modifications | AAACCACAAGTTACACTCCA |
| Human | ESCO2 | DNA and RNA Modifications | ACTGAAGATTGGATAGACAG |
| Human | ESCO2 | DNA and RNA Modifications | ATAGCAAAGCTTGGATCATG |
| Human | EYA1 | Other Histone Modifications | CCTACGCCAACAGATATGGG |
| Human | EYA1 | Other Histone Modifications | AGTCCAAGGGAAACTGAAGT |
| Human | EYA1 | Other Histone Modifications | GATTGCGTCGAGGTTCAGAT |
| Human | EYA1 | Other Histone Modifications | GATTCTGATCGATTGCGTCG |
| Human | EYA1 | Other Histone Modifications | ACAGCTTCAACGACAGCCGA |
| Human | EZH1 | Histone Methylation | CCCACCTCAACTCTGCGTAG |
| Human | EZH1 | Histone Methylation | GGGTCTTACAGCGACAGCCA |
| Human | EZH1 | Histone Methylation | ACCATTCTCAGGGAACATTG |
| Human | EZH1 | Histone Methylation | GACGACCAGAGCACTTGGAG |
| Human | EZH1 | Histone Methylation | TCACCTTCTTAAGTCCACGC |
| Human | EZH2 | Histone Methylation | CAAGGGCACGAACTGTCACA |
| Human | EZH2 | Histone Methylation | TGTCATAGTAAGTGCCAATG |
| Human | EZH2 | Histone Methylation | CAACACCCAACACTTATAAG |
| Human | EZH2 | Histone Methylation | TGGTAACACTGTGGTCCACA |

| | | | |
|---|---|---|---|
| Human | EZH2 | Histone Methylation | GCTCTAGACAACAAACCTTG |
| Human | FEN1 | DNA and RNA Modifications | TGAGGAGCGAATCCGCAGTG |
| Human | FEN1 | DNA and RNA Modifications | TCAGGTGTCGCATTAGCACA |
| Human | FEN1 | DNA and RNA Modifications | GACTACTGTGAGAGTATCCG |
| Human | FEN1 | DNA and RNA Modifications | CTACCGCACCATTCGCATGA |
| Human | FEN1 | DNA and RNA Modifications | ACTCTCTTCAGCTAAGCGCA |
| Human | FMR1 | Chromatin Readers | TCGTCCACGTAATCCAAGAG |
| Human | FMR1 | Chromatin Readers | GTAGACCTTACAGAAATAGG |
| Human | FMR1 | Chromatin Readers | GTTGGTGGTTAGCTAAAGTG |
| Human | FMR1 | Chromatin Readers | TAGGGTACTCCATTCACGAG |
| Human | FMR1 | Chromatin Readers | CGTAATCCAAGAGAGGCTAA |
| Human | FNTB | Control | CAGCTTACCTTGGTACAAGG |
| Human | FNTB | Control | GGAGCTTAAGGATGAGACAT |
| Human | FNTB | Control | GTACAACATTGGACCAGACA |
| Human | FNTB | Control | CAGGACCACATCATGCAACA |
| Human | FNTB | Control | CCACAGCTTACCTTGGTACA |
| Human | FOXA1 | DNA and RNA Modifications | AACATGTCCTATGCCAACCC |
| Human | FOXA1 | DNA and RNA Modifications | TGGACGGCGCGTACGCCATG |
| Human | FOXA1 | DNA and RNA Modifications | GGACATGTTGAAGGACGCCG |
| Human | FOXA1 | DNA and RNA Modifications | CAGCTACTACGCAGACACGC |
| Human | FOXA1 | DNA and RNA Modifications | TACACACCTTGGTAGTACGC |
| Human | FOXA2 | DNA and RNA Modifications | ATGAACATGTCGTCGTACGT |
| Human | FOXA2 | DNA and RNA Modifications | CAGATACCTCCTACTACCAG |
| Human | FOXA2 | DNA and RNA Modifications | GAACAGGTGATGCACTACCC |
| Human | FOXA2 | DNA and RNA Modifications | TCCGTGAGCAACATGAACGC |
| Human | FOXA2 | DNA and RNA Modifications | CGTGCCAGGAGCACAAGCGA |
| Human | FOXA3 | DNA and RNA Modifications | GCACCTCCCAAACTGGACGT |
| Human | FOXA3 | DNA and RNA Modifications | CACCTCCCAAACTGGACGTG |
| Human | FOXA3 | DNA and RNA Modifications | AGTCGTTGAAAGACAGCGAG |
| Human | FOXA3 | DNA and RNA Modifications | CCACTGGTAGATTTCACTCA |
| Human | FOXA3 | DNA and RNA Modifications | CGAGTGGAGCTACTACCCGG |
| Human | FOXO1 | DNA and RNA Modifications | TAGCATTTGAGCTAGTTCGA |
| Human | FOXO1 | DNA and RNA Modifications | GAGTTGGACTGGCTAAACTC |
| Human | FOXO1 | DNA and RNA Modifications | GGTGGCGCAAACGAGTAGCA |
| Human | FOXO1 | DNA and RNA Modifications | TCGTTGTCTTGACACTGTGT |
| Human | FOXO1 | DNA and RNA Modifications | GGAGTTTAGCCAGTCCAACT |
| Human | FOXO3 | DNA and RNA Modifications | CCTGCCATATCAGTCAGCCG |
| Human | FOXO3 | DNA and RNA Modifications | CGTGCACGGTGGAACTGCCA |
| Human | FOXO3 | DNA and RNA Modifications | CAGAGTGAGCCGTTTGTCCG |
| Human | FOXO3 | DNA and RNA Modifications | GTTGCTGACAGAATTCGACA |
| Human | FOXO3 | DNA and RNA Modifications | CCCGCTCTCTCCGCTCGAAG |
| Human | FOXO4 | DNA and RNA Modifications | GTTCATCAAGGTTCACAACG |

| | | | |
|---|---|---|---|
| Human | FOXO4 | DNA and RNA Modifications | GCCCGTGGCCAGCTTACTGG |
| Human | FOXO4 | DNA and RNA Modifications | GGGATCTAGGTCTATGATCG |
| Human | FOXO4 | DNA and RNA Modifications | GAGACCACTCCGAGATAGCA |
| Human | FOXO4 | DNA and RNA Modifications | GTAACAGGTCCTCGGAAGGG |
| Human | FOXP1 | DNA and RNA Modifications | AGAGGAGGAGACACATGTCG |
| Human | FOXP1 | DNA and RNA Modifications | TGAACCAGTAAACGAGGACA |
| Human | FOXP1 | DNA and RNA Modifications | TGCTTACTTCCGACGCAACG |
| Human | FOXP1 | DNA and RNA Modifications | AGCAACCACTTACTAGAGTG |
| Human | FOXP1 | DNA and RNA Modifications | GACCGCCGCACTCTAGTAAG |
| Human | FOXP2 | DNA and RNA Modifications | CCAATGTGGGAGCCATACGA |
| Human | FOXP2 | DNA and RNA Modifications | AGTCCAGGATTACTTAGCAA |
| Human | FOXP2 | DNA and RNA Modifications | TTGCTACAGCTCGTCACATG |
| Human | FOXP2 | DNA and RNA Modifications | GCTCCATAGCCTAAACTGGT |
| Human | FOXP2 | DNA and RNA Modifications | TCAGTGTCGAGTGCAAATGC |
| Human | FOXP3 | DNA and RNA Modifications | ACCCAGGCATCATCCGACAA |
| Human | FOXP3 | DNA and RNA Modifications | CCCACCCACAGGGATCAACG |
| Human | FOXP3 | DNA and RNA Modifications | AGCGTGGCGTAGGTGAAAGG |
| Human | FOXP3 | DNA and RNA Modifications | CACTCAATGAGATCTACCAC |
| Human | FOXP3 | DNA and RNA Modifications | GAGGGTGCCACCATGACTAG |
| Human | FOXP4 | DNA and RNA Modifications | ACGAGCGGGAGTATCAGAAG |
| Human | FOXP4 | DNA and RNA Modifications | TGCAGCCCAACCAAGCCTCG |
| Human | FOXP4 | DNA and RNA Modifications | CATTAAGTGCTCCATAGCTG |
| Human | FOXP4 | DNA and RNA Modifications | GCACAAGTGCTTCGTCCGCG |
| Human | FOXP4 | DNA and RNA Modifications | AGCCGACCTGATTGTCTCCG |
| Human | FTO | DNA and RNA Modifications | AAATCTGCAGACATACACAA |
| Human | FTO | DNA and RNA Modifications | TTTCATTCCTTTGTTCCACG |
| Human | FTO | DNA and RNA Modifications | TCATGGGACATAGAGACACC |
| Human | FTO | DNA and RNA Modifications | TGCAGCCTGGATTACCAATG |
| Human | FTO | DNA and RNA Modifications | TCTGTGAGGTCAAACGGCAG |
| Human | FXR2 | Chromatin Readers | CATAGGTTCGAGTCGCATGG |
| Human | FXR2 | Chromatin Readers | CATACTACCCAACTCCAAGG |
| Human | FXR2 | Chromatin Readers | TCCGATTACCACGGTTACGG |
| Human | FXR2 | Chromatin Readers | TCCCATAACTGTCTCACCAT |
| Human | FXR2 | Chromatin Readers | CATGCGACTCGAACCTATGG |
| Human | FYN | Other Histone Modifications | TGAACTCTTCGTCTCATACG |
| Human | FYN | Other Histone Modifications | GATTGTGAACCTCCCGTACA |
| Human | FYN | Other Histone Modifications | AACAACTTCCACGCAGCCGG |
| Human | FYN | Other Histone Modifications | TGGATACTACATTACCACCC |
| Human | FYN | Other Histone Modifications | TGCAGGCATGAACAACCGGG |
| Human | G2E3 | Chromatin Readers | TGATAACTTCGTTTACATCG |
| Human | G2E3 | Chromatin Readers | AACCAAGCGTTCTTACAACA |
| Human | G2E3 | Chromatin Readers | GCCATTTCTTTAGTTCACGG |

| | | | |
|---|---|---|---|
| Human | G2E3 | Chromatin Readers | AAATTACCCAGACAGTCACC |
| Human | G2E3 | Chromatin Readers | TAGAGACAAGTTCTTTGACA |
| Human | G6PD | Control | CTTGAAGGTGAGGATAACGC |
| Human | G6PD | Control | AGAGGTGCAGGCCAACAATG |
| Human | G6PD | Control | GTATGAGGGCACCTACAAGT |
| Human | G6PD | Control | GATCCGCGTGCAGCCCAACG |
| Human | G6PD | Control | TGCCCGTTCCCGCCTCACAG |
| Human | GGPS1 | Control | AACAGATTGATGCACGTGGT |
| Human | GGPS1 | Control | ACTCCGACGTGGCTTTCCAG |
| Human | GGPS1 | Control | GGACTGTTTGGATTAGCAGT |
| Human | GGPS1 | Control | TTACTAAGGCTACTAGCTCA |
| Human | GGPS1 | Control | TATAAACAGATTGATGCACG |
| Human | GINS4 | Control | CTGGGACAGGACTCTGATGG |
| Human | GINS4 | Control | CCTGGGACAGGACTCTGATG |
| Human | GINS4 | Control | CCTGTTTACCTGCCCGAAAG |
| Human | GINS4 | Control | AGAGAAGATGACCGAAGAAG |
| Human | GINS4 | Control | ACCGAAGAAGTGGATTTCCT |
| Human | GLE1 | Control | GCCCTACGCAGTTCCGACAA |
| Human | GLE1 | Control | GTTCAGATGCCATCTCCGAG |
| Human | GLE1 | Control | TGGTACTTCAGTCCTCACGG |
| Human | GLE1 | Control | AGCCAGGTCGACCTTAAGCA |
| Human | GLE1 | Control | ATGGTACTTCAGTCCTCACG |
| Human | GLYR1 | Chromatin Readers | TCATGGCCACTATTGCCGAG |
| Human | GLYR1 | Chromatin Readers | GAATCGACGTAATTCCAGTG |
| Human | GLYR1 | Chromatin Readers | CTACGTGGACATGTCAACAG |
| Human | GLYR1 | Chromatin Readers | AGGTTGAGACGACTTCAGCG |
| Human | GLYR1 | Chromatin Readers | GTGTATGTAGGCTCGGTACA |
| Human | GMPPB | Control | GGAACCCTCCAAGTACGGTG |
| Human | GMPPB | Control | GGCTTACCCACTGACCCACG |
| Human | GMPPB | Control | CACCTGCGACATGTAGCTCA |
| Human | GMPPB | Control | AGCCTCGTATCATCATGTGA |
| Human | GMPPB | Control | ACCACCACACCGTACTTGGA |
| Human | GPN3 | Control | TCTCTGCGATTCGGTCCCAA |
| Human | GPN3 | Control | CTGCGATTCGGTCCCAACGG |
| Human | GPN3 | Control | AAACTGACTAAAGCTATATG |
| Human | GPN3 | Control | AGACATCCGGGAACTGATCG |
| Human | GPN3 | Control | TTCACAGTGCTGGACCATGG |
| Human | GTF2A1 | Transcription Elongation | CTCGGCAAATACAAACACCG |
| Human | GTF2A1 | Transcription Elongation | GAATCTGGAACAATAACTTG |
| Human | GTF2A1 | Transcription Elongation | CCATCAACTTGTAAGACCAA |
| Human | GTF2A1 | Transcription Elongation | GTGTGTGGAAATGGCGAACT |
| Human | GTF2A1 | Transcription Elongation | ACATCATCTTCACTATTGAG |

| | | | |
|---|---|---|---|
| Human | GTF2A2 | Transcription Elongation | ACAGATTCTGCGATAATGTG |
| Human | GTF2A2 | Transcription Elongation | GAATGATGTTGAATTCAGAG |
| Human | GTF2A2 | Transcription Elongation | TAAATGCAGCACTGGCTCAG |
| Human | GTF2A2 | Transcription Elongation | TAAGGCTATAAATGCAGCAC |
| Human | GTF2A2 | Transcription Elongation | CAGGAACAGAGTCAATTTCA |
| Human | GTF2B | Transcription Elongation | TGTGGATTTGATTACAACTG |
| Human | GTF2B | Transcription Elongation | ACATATAGCCCGTAAAGCTG |
| Human | GTF2B | Transcription Elongation | AGTTGTAATCAAATCCACAC |
| Human | GTF2B | Transcription Elongation | CTGTGGTAGTTTGTCCACTG |
| Human | GTF2B | Transcription Elongation | GATATGATCTGTCCTGAATG |
| Human | GTF2E1 | Transcription Elongation | TGAGAGAGATTCGACCAACC |
| Human | GTF2E1 | Transcription Elongation | TTAGCCAAGTATGTGATCCG |
| Human | GTF2E1 | Transcription Elongation | ACATTTGAAGGAAGCCCGGT |
| Human | GTF2E1 | Transcription Elongation | ATGCTGCTGGAACTTCAGTG |
| Human | GTF2E1 | Transcription Elongation | GAGAGAAAGCACTGTCCAAG |
| Human | GTF2E2 | Transcription Elongation | GAAGAATATCTGAAGCGACA |
| Human | GTF2E2 | Transcription Elongation | TGATAGACACGGCATCAGCG |
| Human | GTF2E2 | Transcription Elongation | TAAGATTGTGAATTACATGA |
| Human | GTF2E2 | Transcription Elongation | TGAAGAATATCTGAAGCGAC |
| Human | GTF2E2 | Transcription Elongation | GAAACAATGGCTAATGACTG |
| Human | GTF2F1 | Transcription Elongation | ACAAAGTCAACTTTGCTACG |
| Human | GTF2F1 | Transcription Elongation | GAGGTGGACTACATGTCAGA |
| Human | GTF2F1 | Transcription Elongation | GGCAAGACAACACCCAACAG |
| Human | GTF2F1 | Transcription Elongation | GCAAGACAACACCCAACAGC |
| Human | GTF2F1 | Transcription Elongation | GCTGCCAGCAAACTCGAGCA |
| Human | GTF2F2 | Transcription Elongation | CCAGAGTACAGACACTATCA |
| Human | GTF2F2 | Transcription Elongation | CAGACCCATGGCCGAGCGCG |
| Human | GTF2F2 | Transcription Elongation | GCATATAGATCGAATATGAA |
| Human | GTF2F2 | Transcription Elongation | CACAGGAGTGTGGCTAGTCA |
| Human | GTF2F2 | Transcription Elongation | GGACATCACAAAGCAACCTG |
| Human | GTF2H1 | Transcription Elongation | GACGCCGTGACTGCCATGTG |
| Human | GTF2H1 | Transcription Elongation | TTATACACCCAAGTTAACTC |
| Human | GTF2H1 | Transcription Elongation | ATATTTAGGACCTATCCAGC |
| Human | GTF2H1 | Transcription Elongation | TTCAGCTGGTCCTACATGCA |
| Human | GTF2H1 | Transcription Elongation | TCACCTACCTGAGTTAACTT |
| Human | GTF2H2 | Transcription Elongation | AACGTCTTTGAAGAAAGCTG |
| Human | GTF2H2 | Transcription Elongation | AGTCTGCATAGCTATGCTTA |
| Human | GTF2H2 | Transcription Elongation | AGTACTTTGCCCGACACTGT |
| Human | GTF2H2 | Transcription Elongation | CATCTACTACCACATAAAGG |
| Human | GTF2H2 | Transcription Elongation | AAGCCAAAGAACTTACATGT |
| Human | GTF2H3 | Transcription Elongation | ACTCGCACGTAGTACAAATG |
| Human | GTF2H3 | Transcription Elongation | CAGGCTTGTGACATCACGGG |

| | | | |
|---|---|---|---|
| Human | GTF2H3 | Transcription Elongation | GATGCCAACCCAATTTGGTG |
| Human | GTF2H3 | Transcription Elongation | CCTGTAGTCAACATGAACTG |
| Human | GTF2H3 | Transcription Elongation | CAAATGCATAGATGCCGTGA |
| Human | GTF2H4 | Transcription Elongation | CTGGTTATACAGGACACCTG |
| Human | GTF2H4 | Transcription Elongation | TACCGACTGTATGCCTACAC |
| Human | GTF2H4 | Transcription Elongation | GACAGACTCCGGTTCACTGA |
| Human | GTF2H4 | Transcription Elongation | TTTGTCTCTAGTACTGAACC |
| Human | GTF2H4 | Transcription Elongation | TGTTCGAGAACTCGGCCAAG |
| Human | GTF2H5 | Transcription Elongation | TTGGATGAGTCCAATGCCCT |
| Human | GTF2H5 | Transcription Elongation | TGGATGAGTCCAATGCCCTG |
| Human | GTF2H5 | Transcription Elongation | AGCAGAAACTGCTTCATGGC |
| Human | GTF2H5 | Transcription Elongation | AATGTCCTCCAGGAGCGAGT |
| Human | GTF2H5 | Transcription Elongation | AACATGGTCAACGTCTTGAA |
| Human | GTF3C4 | Histone Acetylation | GGATAGGGTGTTCAACCCTG |
| Human | GTF3C4 | Histone Acetylation | AACAACGAATGCCGGGACGT |
| Human | GTF3C4 | Histone Acetylation | TGAAAGTCTGACTGACCGTG |
| Human | GTF3C4 | Histone Acetylation | ATGAAAGTCTGACTGACCGT |
| Human | GTF3C4 | Histone Acetylation | CTTGTGGAAACCCACCCATG |
| Human | GUK1 | Control | TCTCTCTGCTAGATACCACG |
| Human | GUK1 | Control | GCAGCGCAACACTGAAACCG |
| Human | GUK1 | Control | AGATTACTACTTTGTAACCA |
| Human | GUK1 | Control | TAGATACCACGAGGAACCCG |
| Human | GUK1 | Control | ACATCAAGGCCACCGATCTG |
| Human | HAT1 | Histone Acetylation | TGATCGTAGCTCGGAAATGG |
| Human | HAT1 | Histone Acetylation | CAAGTCGTTCAATAACACGC |
| Human | HAT1 | Histone Acetylation | GCTACGCTCTTTGCGACCGT |
| Human | HAT1 | Histone Acetylation | ACTGCCAATTACCTTACACG |
| Human | HAT1 | Histone Acetylation | ATTCTAGCAACACGCTAGAA |
| Human | HAUS3 | Control | CATCAGCAAGAATATCAGTG |
| Human | HAUS3 | Control | AGATTATCTGAAAGATATTG |
| Human | HAUS3 | Control | TGTGATACTTTGTATCAAGG |
| Human | HAUS3 | Control | CCAGTTATTGAATATGCCAG |
| Human | HAUS3 | Control | TGAGGAGAGACGACTAGAGA |
| Human | HDAC1 | Histone Acetylation | CATCCGTCCAGATAACATGT |
| Human | HDAC1 | Histone Acetylation | ACTCGTCATCAATCCCGTCT |
| Human | HDAC1 | Histone Acetylation | TCATATTGGAAGGACTGATG |
| Human | HDAC1 | Histone Acetylation | TGAGTCATGCGGATTCGGTG |
| Human | HDAC1 | Histone Acetylation | GCACCGGGCAACGTTACGAA |
| Human | HDAC10 | Histone Acetylation | AGTCAGATGCAGACGCAGTG |
| Human | HDAC10 | Histone Acetylation | GACCGCGCTTGTGTACCATG |
| Human | HDAC10 | Histone Acetylation | GCACATCCCAGTCCACGACG |
| Human | HDAC10 | Histone Acetylation | AGGATTTGACTCAGCCATCG |

| | | | |
|---|---|---|---|
| Human | HDAC10 | Histone Acetylation | CCGCAGCCCTGGATCGCCTG |
| Human | HDAC11 | Histone Acetylation | CAGGCACCGACATCCTCGAG |
| Human | HDAC11 | Histone Acetylation | TGCATTGTATACCACCACGT |
| Human | HDAC11 | Histone Acetylation | CGAGGCTGGGCCATCAACGT |
| Human | HDAC11 | Histone Acetylation | CAACCGCCACATCTACCCAG |
| Human | HDAC11 | Histone Acetylation | GCAGGCACCGACATCCTCGA |
| Human | HDAC2 | Histone Acetylation | GATGTATCAACCTAGTGCTG |
| Human | HDAC2 | Histone Acetylation | CTTGATTGTGAGATTCCCAA |
| Human | HDAC2 | Histone Acetylation | CAGTTCGAGCATCAGACAAG |
| Human | HDAC2 | Histone Acetylation | TCATTATCTGGTGATAGACT |
| Human | HDAC2 | Histone Acetylation | TACAACAGATCGTGTAATGA |
| Human | HDAC3 | Histone Acetylation | ACCTGGAGCACAATGCACGT |
| Human | HDAC3 | Histone Acetylation | GGGCAACTTCCACTACGGTG |
| Human | HDAC3 | Histone Acetylation | TGTCAATGTAGAGCACCCGA |
| Human | HDAC3 | Histone Acetylation | AGTCTTAATGCCTTCAACGT |
| Human | HDAC3 | Histone Acetylation | TATGTCAACGACATTGTGAT |
| Human | HDAC4 | Histone Acetylation | GCATCAGCGTGTCATACACG |
| Human | HDAC4 | Histone Acetylation | ACTTACCCGTACCAGTAGCG |
| Human | HDAC4 | Histone Acetylation | CTTACCCGTACCAGTAGCGA |
| Human | HDAC4 | Histone Acetylation | GCAGTGGTTCAGATTCCGGT |
| Human | HDAC4 | Histone Acetylation | AGAAGTTCCCATCGTCGTAG |
| Human | HDAC5 | Histone Acetylation | CCCTCATAGGACATTCACCA |
| Human | HDAC5 | Histone Acetylation | CACATCAGGTGAGAACTCGT |
| Human | HDAC5 | Histone Acetylation | CTACAGCAGAAGTTGAACGT |
| Human | HDAC5 | Histone Acetylation | TGGCTGTGGATTCCTCGGCG |
| Human | HDAC5 | Histone Acetylation | TTCAACAGCATCAAACCCGG |
| Human | HDAC6 | Histone Acetylation | TGTGCTGAGTTCCATTACCG |
| Human | HDAC6 | Histone Acetylation | AATGGAAGAAGACCTAATCG |
| Human | HDAC6 | Histone Acetylation | AGGACACGCAGCGATCTAGG |
| Human | HDAC6 | Histone Acetylation | CAGTGCTACAGTCTCGCACA |
| Human | HDAC6 | Histone Acetylation | GCTGATTCGATGCTGATGCA |
| Human | HDAC7 | Histone Acetylation | TGCAGTCGGTCCACTCTGAG |
| Human | HDAC7 | Histone Acetylation | ACGACGGCAACTTCTTCCCG |
| Human | HDAC7 | Histone Acetylation | AACATGGTAGCCACCCAGTG |
| Human | HDAC7 | Histone Acetylation | AGTCCTTAATGACCACCGAG |
| Human | HDAC7 | Histone Acetylation | GTTACCTGTAGGGAATGCCG |
| Human | HDAC8 | Histone Acetylation | ATGCTGGACATACTTGACCG |
| Human | HDAC8 | Histone Acetylation | ATTTGAGCGTATTCTCTACG |
| Human | HDAC8 | Histone Acetylation | TGGGAATATTACGATTGCGA |
| Human | HDAC8 | Histone Acetylation | CGGGACTATAGATATAAACC |
| Human | HDAC8 | Histone Acetylation | AGCATCGAGCCGTGTTGGCA |
| Human | HDAC9 | Histone Acetylation | AGCTTTGATCCAATGATGTG |

| | | | |
|---|---|---|---|
| Human | HDAC9 | Histone Acetylation | GTCACTTTGTACCCTCCTAG |
| Human | HDAC9 | Histone Acetylation | TCCCTGTAGGATGTTCACCA |
| Human | HDAC9 | Histone Acetylation | AACAGCATGAGAACTTGACA |
| Human | HDAC9 | Histone Acetylation | TTTCCCTCTAAAGTAACATG |
| Human | HDGF | Chromatin Readers | CTCTCCTAAACGTCCCAAGG |
| Human | HDGF | Chromatin Readers | GCAGAAGGAGTACAAATGCG |
| Human | HDGF | Chromatin Readers | TCTCCTTGGATTCCTCGTAA |
| Human | HDGF | Chromatin Readers | CGAGAACAACCCTACTGTCA |
| Human | HDGF | Chromatin Readers | GGGCCTCTCAACCTCCAAGG |
| Human | HDGFL1 | Chromatin Readers | GGACGACGACAAGCCCACTG |
| Human | HDGFL1 | Chromatin Readers | GAACCAGCTATGTCGGCCTA |
| Human | HDGFL1 | Chromatin Readers | CATGCCCATGTACAAGAGCG |
| Human | HDGFL1 | Chromatin Readers | GACAAGCCGACCCACGCTGG |
| Human | HDGFL1 | Chromatin Readers | GGCATGCCCATGTACAAGAG |
| Human | HIF1AN | Histone Methylation | TGTGGTTGGTTACGAAACAG |
| Human | HIF1AN | Histone Methylation | GGATGTAAAGAACATCACCA |
| Human | HIF1AN | Histone Methylation | TAGAGTCATTACTAAATGGG |
| Human | HIF1AN | Histone Methylation | CAAACGCTCAATGACACTGT |
| Human | HIF1AN | Histone Methylation | AACTGGATTAATAAGCAACA |
| Human | HJURP | DNA and RNA Modifications | TCCAGACCAACAATACGACA |
| Human | HJURP | DNA and RNA Modifications | CTGACCTACGAGACGCCACA |
| Human | HJURP | DNA and RNA Modifications | TGGCGTCTCGTAGGTCAGCG |
| Human | HJURP | DNA and RNA Modifications | ACCATCAGTGACCTGTACGC |
| Human | HJURP | DNA and RNA Modifications | CATCACGAGGGATGGCACGA |
| Human | HLTF | Chromatin Remodeling | TCACGTGCTAATTCTTCTGG |
| Human | HLTF | Chromatin Remodeling | GTTGGACTACGCTATTACAC |
| Human | HLTF | Chromatin Remodeling | GAAAGAAAGTTGGATATGAG |
| Human | HLTF | Chromatin Remodeling | TGGAAGTTTGAGAGGTCATG |
| Human | HLTF | Chromatin Remodeling | TGTGCACAATGTGTTATCAC |
| Human | HMGA1 | DNA and RNA Modifications | AGCGCTGGTAGGGAGTCAGG |
| Human | HMGA1 | DNA and RNA Modifications | GCCCACCTCACAGGAGAAGG |
| Human | HMGA1 | DNA and RNA Modifications | CATCTCGCAGGAGTCCTCGG |
| Human | HMGA1 | DNA and RNA Modifications | ACTGCCCACCTCACAGGAGA |
| Human | HMGA1 | DNA and RNA Modifications | GGGTCACTGCTCCTCCTCCG |
| Human | HMGN1 | Transcription Elongation | AACTAAAGAAGACTTACCTG |
| Human | HMGN1 | Transcription Elongation | CGCCACGATGCCCAAGAGGA |
| Human | HMGN1 | Transcription Elongation | CCGCAGGTCAGCTCCGCCGA |
| Human | HMGN1 | Transcription Elongation | TTTCTTTCTCCAAGCCCAAG |
| Human | HMGN1 | Transcription Elongation | CCGCCGCCACGATGCCCAAG |
| Human | HMGN4 | DNA and RNA Modifications | AGCCACAGAGGAGATCAGCT |
| Human | HMGN4 | DNA and RNA Modifications | AGGTGAAGGATGAGCCACAG |
| Human | HMGN4 | DNA and RNA Modifications | CTCCAGTCCCAGAAAGCGGA |

| | | | |
|---|---|---|---|
| Human | HMGN4 | DNA and RNA Modifications | AACCGAGCTGATCTCCTCTG |
| Human | HMGN4 | DNA and RNA Modifications | AGCTTCCCAAAGGGAGAAAG |
| Human | HNRNPU | Transcription Elongation | CGTGGCGGATATAATAGGAG |
| Human | HNRNPU | Transcription Elongation | ATCGAGTTAGAGGACCAAAG |
| Human | HNRNPU | Transcription Elongation | CGTGGTAGTTACTCAAACAG |
| Human | HNRNPU | Transcription Elongation | GGCAAGAACCAGTTTAACAG |
| Human | HNRNPU | Transcription Elongation | GGAGGATTCAATATGCGTGG |
| Human | HR | Histone Methylation | GAGCAGTTTGAATGTCCACG |
| Human | HR | Histone Methylation | GTTCGCCCAAAGTCAGACGA |
| Human | HR | Histone Methylation | GGGTCAAGTTTGATATCCGG |
| Human | HR | Histone Methylation | AGTTCGCCCAAAGTCAGACG |
| Human | HR | Histone Methylation | GAGTTTAAGCCAAACAACCC |
| Human | HSPBAP1 | Histone Methylation | TTACATGTAGTTTCAAACTG |
| Human | HSPBAP1 | Histone Methylation | GAATACCAAGTTACAACCAT |
| Human | HSPBAP1 | Histone Methylation | AGCCGCAACGATCACAGGAG |
| Human | HSPBAP1 | Histone Methylation | GGAAGAGGATCACCTAGCCC |
| Human | HSPBAP1 | Histone Methylation | GATGTGAAATGGTCTGACTT |
| Human | HTATSF1 | Transcription Elongation | ACTTCGAACACTCTACTCGA |
| Human | HTATSF1 | Transcription Elongation | TCTCGAGAAAGATTTGGACG |
| Human | HTATSF1 | Transcription Elongation | AATGCTCCTGAGGCCAACAG |
| Human | HTATSF1 | Transcription Elongation | AGACAGAATCTGAACGCCTA |
| Human | HTATSF1 | Transcription Elongation | GTGGCAAAGTTTCAACTGAA |
| Human | ICE1 | Transcription Elongation | AGAGCAAAGTTCTAACTGCG |
| Human | ICE1 | Transcription Elongation | GCTTTCATAAATACTGACCG |
| Human | ICE1 | Transcription Elongation | AGACCTAAGTGATAACACTT |
| Human | ICE1 | Transcription Elongation | GCAGCGTGGCCAGATGCACA |
| Human | ICE1 | Transcription Elongation | TGTTACACAGGCATTCGAGA |
| Human | ICE2 | Transcription Elongation | TAATCATGGACCAACGTACA |
| Human | ICE2 | Transcription Elongation | AGAATTCTATACTCTCCACG |
| Human | ICE2 | Transcription Elongation | TCCAGATTTCGGCCAACTAG |
| Human | ICE2 | Transcription Elongation | GACCAACGTACAAGGAACAG |
| Human | ICE2 | Transcription Elongation | GACTAGTTAGAGCTATCTGA |
| Human | ING1 | Chromatin Readers | CATGTTGAGTCCTGCCAACG |
| Human | ING1 | Chromatin Readers | TGGTTGCACAGACAGTACGT |
| Human | ING1 | Chromatin Readers | CTTCAGTCGCGAGACAGACG |
| Human | ING1 | Chromatin Readers | AGCTGGTGGAGAACCGCACG |
| Human | ING1 | Chromatin Readers | GGTTCCACTTCTCGTGCGTG |
| Human | ING2 | Chromatin Readers | GCAGCAGCAACTGTACTCGT |
| Human | ING2 | Chromatin Readers | ATGCAACCAAGTGTCTTATG |
| Human | ING2 | Chromatin Readers | CACTTACCTATAAACCAAAG |
| Human | ING2 | Chromatin Readers | TTCACTTACCTATAAACCAA |
| Human | ING2 | Chromatin Readers | ACGTTCCTCTGCATGTCGTG |

| | | | |
|---|---|---|---|
| Human | ING3 | Chromatin Readers | AGAGGCATCTGACGTAAGGG |
| Human | ING3 | Chromatin Readers | TCTCGCGCATTTCCGTGAAG |
| Human | ING3 | Chromatin Readers | TTCATTGCAGCAGTGCACTG |
| Human | ING3 | Chromatin Readers | CTTTCGACCACTCCGTGAGT |
| Human | ING3 | Chromatin Readers | GAATATGATCTGTTGTCGTA |
| Human | ING4 | Chromatin Readers | GGACCTAGACCAAAGAACAG |
| Human | ING4 | Chromatin Readers | GGCACTACTCATATACTCAG |
| Human | ING4 | Chromatin Readers | GGGCTGACAACCAAGCCTCG |
| Human | ING4 | Chromatin Readers | GCATATCCAACACATCAGAG |
| Human | ING4 | Chromatin Readers | CAAGGAATTTGGTGACGACA |
| Human | ING5 | Chromatin Readers | GACCTTACCACGAAACCCAA |
| Human | ING5 | Chromatin Readers | AAGTTCCTCTGAAGTTCGCA |
| Human | ING5 | Chromatin Readers | GGCTGCAGAGTACATCTCCA |
| Human | ING5 | Chromatin Readers | CTGGACCAGAGGACGGAAGG |
| Human | ING5 | Chromatin Readers | ACCTTCCCTGCGAACTTCAG |
| Human | INO80 | Chromatin Remodeling | TCCGTCTCAGATACCGTGAA |
| Human | INO80 | Chromatin Remodeling | AAATAAGTATAGTAAAGACA |
| Human | INO80 | Chromatin Remodeling | GGTAATAACCACATGGAAGG |
| Human | INO80 | Chromatin Remodeling | TCTCCAAGTACTGAAGATAG |
| Human | INO80 | Chromatin Remodeling | GATCTCGGAGAGGCGAGACA |
| Human | INO80B | Chromatin Remodeling | CTGTGCGGGAGCAAGCGTAG |
| Human | INO80B | Chromatin Remodeling | GCTGCAGTAGCGCACCATGG |
| Human | INO80B | Chromatin Remodeling | CCTCCTGATAGGTCCCGAAG |
| Human | INO80B | Chromatin Remodeling | CAGGCACGGTACTGCTCAAG |
| Human | INO80B | Chromatin Remodeling | ACCTCATGAGTAAGCTGTGG |
| Human | INO80C | Chromatin Remodeling | GGCGGCGCAAATTCCAATTG |
| Human | INO80C | Chromatin Remodeling | TAGCCGGTGACGACGTCAGA |
| Human | INO80C | Chromatin Remodeling | TTAGGATCGTTCAGTTGCCA |
| Human | INO80C | Chromatin Remodeling | TTAGCTGGCTTAAAGGATGG |
| Human | INO80C | Chromatin Remodeling | GTGCCCTCTGAGTTTAGCAC |
| Human | INO80D | Chromatin Remodeling | AATACAATCCTTGAGCCGAG |
| Human | INO80D | Chromatin Remodeling | GGCATCTTCAAGGCCAACGT |
| Human | INO80D | Chromatin Remodeling | TGCAAGCAGAGGCGACTCAA |
| Human | INO80D | Chromatin Remodeling | GGCAGAAGTAAGGCCAACGT |
| Human | INO80D | Chromatin Remodeling | TGTGGTCTCTGAGTTCGAAG |
| Human | INO80E | Chromatin Remodeling | ATCATCAGATAACAGCGAGA |
| Human | INO80E | Chromatin Remodeling | CAGATAACAGCGAGACGGAG |
| Human | INO80E | Chromatin Remodeling | TCAGATAACAGCGAGACGGA |
| Human | INO80E | Chromatin Remodeling | ACTTCTGCAGTACGAGAACG |
| Human | INO80E | Chromatin Remodeling | GGTAGTCAGAAGGGAATGGG |
| Human | INTS12 | Chromatin Readers | AACTTACCTACAAACAACGC |
| Human | INTS12 | Chromatin Readers | CACAATACCACACCAGGCGA |

| | | | |
|---|---|---|---|
| Human | INTS12 | Chromatin Readers | GGAACTAAACTACTTGGACT |
| Human | INTS12 | Chromatin Readers | GTTTCCTCGTCAGTAACTAG |
| Human | INTS12 | Chromatin Readers | ATGAAATCAGACATCACTGA |
| Human | IWS1 | Transcription Elongation | ATGGACTCGGAATATTACAG |
| Human | IWS1 | Transcription Elongation | CTCTCAGAGTCACTGACTCG |
| Human | IWS1 | Transcription Elongation | CTTTCTGAGTCACTGATACG |
| Human | IWS1 | Transcription Elongation | TAATAAAGGTGCCACCATCG |
| Human | IWS1 | Transcription Elongation | ATCCCGTTCATCCTGTACTG |
| Human | JADE1 | Chromatin Readers | GGTCCACAGATGTGTCCCAG |
| Human | JADE1 | Chromatin Readers | CAGCTCACATAGGAAACCCG |
| Human | JADE1 | Chromatin Readers | CCACCCGTAGCGGAACCAAG |
| Human | JADE1 | Chromatin Readers | GCTATGAAGCCCACCCGTAG |
| Human | JADE1 | Chromatin Readers | CACCCGTAGCGGAACCAAGT |
| Human | JADE2 | Chromatin Readers | ACTGAACAATGGGCACCGCG |
| Human | JADE2 | Chromatin Readers | TATATTAGCAGACAACGATG |
| Human | JADE2 | Chromatin Readers | AGGAGGCCAGTACGCCCATG |
| Human | JADE2 | Chromatin Readers | CAGATGTGGGCTCATTACGT |
| Human | JADE2 | Chromatin Readers | CGTGAACCAGGCCCGTCCTG |
| Human | JADE3 | Chromatin Readers | CTGCCCATGATAGCACTCCG |
| Human | JADE3 | Chromatin Readers | TCAGCATTGCTTGTCCTGAG |
| Human | JADE3 | Chromatin Readers | AAGGAAAGATAGCTCAGACA |
| Human | JADE3 | Chromatin Readers | GCTGCCCTGACTGACCAAGT |
| Human | JADE3 | Chromatin Readers | GAGCAGTGAATATACTTCCG |
| Human | JAK2 | Other Histone Modifications | CTGCCACTGCAATACCAACG |
| Human | JAK2 | Other Histone Modifications | TAGGGATCTAGCTCTTCGAG |
| Human | JAK2 | Other Histone Modifications | TATCGGCATGGAATATCTCG |
| Human | JAK2 | Other Histone Modifications | AATGAAGAGTACAACCTCAG |
| Human | JAK2 | Other Histone Modifications | TCTTCAGGAGAGAATACCAT |
| Human | JARID2 | Histone Methylation | AGGGACTGACCTCATCGTAG |
| Human | JARID2 | Histone Methylation | CTGAGCCGACCGATGCAGAG |
| Human | JARID2 | Histone Methylation | ACCGACTCGATGTAGATGAG |
| Human | JARID2 | Histone Methylation | TGAGATTGTGTGGTTGACAG |
| Human | JARID2 | Histone Methylation | TTGGTCTCGAGACCAGCATG |
| Human | JMJD1C | Histone Methylation | CAGCTGGAAAGCTACGTGTG |
| Human | JMJD1C | Histone Methylation | TTGGCATTACATCACGACGC |
| Human | JMJD1C | Histone Methylation | GAGAGCCTTGAAGATACACG |
| Human | JMJD1C | Histone Methylation | TTATCCATATTGCATTCCTG |
| Human | JMJD1C | Histone Methylation | CAATCTTTCAATACTGAACG |
| Human | JMJD4 | Histone Methylation | GAAGATGTCAGCATGGAACG |
| Human | JMJD4 | Histone Methylation | CAGTCGGACGAGAAGTACAC |
| Human | JMJD4 | Histone Methylation | GGACGAGAAGTACACAGGCA |
| Human | JMJD4 | Histone Methylation | CTGGAAAGAGTACATACAGG |

| | | | |
|---|---|---|---|
| Human | JMJD4 | Histone Methylation | ACTACCGCTTTGTCTACGCG |
| Human | JMJD6 | Histone Methylation | GCTGTCGGAAGCTATCCCTG |
| Human | JMJD6 | Histone Methylation | TTAATTCCCTCAGGATAACG |
| Human | JMJD6 | Histone Methylation | CGGAACCAGAAGTTCAAGTG |
| Human | JMJD6 | Histone Methylation | CCGTCTTGTGCCATACCACA |
| Human | JMJD6 | Histone Methylation | AAGTGACCCGAGACGAAGGA |
| Human | JMJD7 | Histone Methylation | GTGCTAGGTCTGGCGCCAAG |
| Human | JMJD7 | Histone Methylation | GCTGGAAGTAACTATACTTG |
| Human | JMJD7 | Histone Methylation | GTCATACCAGAAATTCACTG |
| Human | JMJD7 | Histone Methylation | CCTTGGCGCCAGACCTAGCA |
| Human | JMJD7 | Histone Methylation | AGTGAATTTCTGGTATGACA |
| Human | JMJD8 | Histone Methylation | TAGATCACTTCTGAGTACCC |
| Human | JMJD8 | Histone Methylation | GTTCCCAGCAGGCCAAATGG |
| Human | JMJD8 | Histone Methylation | GCAGGGACTCACGGACAACT |
| Human | JMJD8 | Histone Methylation | TGAGCGTAGCATGCCACCAG |
| Human | JMJD8 | Histone Methylation | GCCCGGATGGTACACTCCAG |
| Human | KAT2A | Histone Acetylation | AGATCCGCAAGGTCTACCCG |
| Human | KAT2A | Histone Acetylation | ATGAGATAAACCGACTGCTG |
| Human | KAT2A | Histone Acetylation | CAAGGTCAATTACACCAGGT |
| Human | KAT2A | Histone Acetylation | GAAGCCGCTACTACGTGACC |
| Human | KAT2A | Histone Acetylation | GTAGCCGATGGCGTACTCGT |
| Human | KAT2B | Histone Acetylation | ACCTCGGTACGAAACCACAC |
| Human | KAT2B | Histone Acetylation | TGTTAATGGTATTACCCATG |
| Human | KAT2B | Histone Acetylation | ACACGTAGTACCTATTCTTG |
| Human | KAT2B | Histone Acetylation | GCGAGCGCGTACCTTGCAGG |
| Human | KAT2B | Histone Acetylation | GGGTGTAATTGAATTTCACG |
| Human | KAT5 | Histone Acetylation | GATTGATGGACGTAAGAACA |
| Human | KAT5 | Histone Acetylation | CTGAGGGTACCGTTGAGCGG |
| Human | KAT5 | Histone Acetylation | ACCAGGCTGCCAGTCATGCG |
| Human | KAT5 | Histone Acetylation | GCAGCCCAACCACCGCTCAA |
| Human | KAT5 | Histone Acetylation | ATGAATGGGTGACGCATGAG |
| Human | KAT6A | Histone Acetylation | CATCCTGCGAAGACAAACGA |
| Human | KAT6A | Histone Acetylation | GGAAGTCCAGCATTCGTAGG |
| Human | KAT6A | Histone Acetylation | TGAGAGTGGTCTGATACGGA |
| Human | KAT6A | Histone Acetylation | AGCAGTTACGACTCCACGAT |
| Human | KAT6A | Histone Acetylation | TACGGAAGTCCAGCATTCGT |
| Human | KAT6B | Histone Acetylation | TTACGACGCATACGTTTGGG |
| Human | KAT6B | Histone Acetylation | TGTCAACTAGAGCTAACAGT |
| Human | KAT6B | Histone Acetylation | TGAAGAGAGAATCTGCCATG |
| Human | KAT6B | Histone Acetylation | ATGTAATGATCTCCGCAATG |
| Human | KAT6B | Histone Acetylation | TGATCGACAAGAGAGATGGC |
| Human | KAT7 | Histone Acetylation | AGAGCCAAACGATACTCCGC |

| | | | |
|---|---|---|---|
| Human | KAT7 | Histone Acetylation | TGATGAGTGGATAGCCAAAG |
| Human | KAT7 | Histone Acetylation | CTGAGAGGTTATGATACAGT |
| Human | KAT7 | Histone Acetylation | ATGAACGAGTCTGCCGAAGA |
| Human | KAT7 | Histone Acetylation | TCTCATCGTGAGATACATTG |
| Human | KAT8 | Histone Acetylation | GTCTCGAGTGAACGACCAGG |
| Human | KAT8 | Histone Acetylation | CCATACTTACAGTTAACCGG |
| Human | KAT8 | Histone Acetylation | GGAATTCTATGTACACTACG |
| Human | KAT8 | Histone Acetylation | TCTCGAGTGAACGACCAGGA |
| Human | KAT8 | Histone Acetylation | TATGTGGACAAGATCCACAT |
| Human | KDM1A | Histone Methylation | TGGAATAGCAGAGACTCCGG |
| Human | KDM1A | Histone Methylation | CTAAATAACTGTGAACTCGG |
| Human | KDM1A | Histone Methylation | TGTATACCACACCTTGCATA |
| Human | KDM1A | Histone Methylation | TGAGAAGTCATCCGGTCATG |
| Human | KDM1A | Histone Methylation | TTTCTGAAACAGGATCGTGT |
| Human | KDM1B | Histone Methylation | ATGTAATGGCAACTCCACGG |
| Human | KDM1B | Histone Methylation | ATAGCCAAGACTTATCGATG |
| Human | KDM1B | Histone Methylation | GGAGAGATGATACCATCGGG |
| Human | KDM1B | Histone Methylation | GAAATCAGTCATCATTATCG |
| Human | KDM1B | Histone Methylation | CAACACTGGAGTTCTCAGCG |
| Human | KDM2A | Histone Methylation | GTATGCGAGTGTGCAAGACG |
| Human | KDM2A | Histone Methylation | GCGAATTATGAGGCGAAGCG |
| Human | KDM2A | Histone Methylation | GGCAGGGAGTCGTCGCATGG |
| Human | KDM2A | Histone Methylation | GGCCTCACAGGTGTTAACCG |
| Human | KDM2A | Histone Methylation | AATGTAGAGTATATTCAGCG |
| Human | KDM2B | Histone Methylation | CCAGCGATACGACGAGAACG |
| Human | KDM2B | Histone Methylation | TGTCCGAAGTGTAACCACGC |
| Human | KDM2B | Histone Methylation | TGACGTGGTTACAGTAACTG |
| Human | KDM2B | Histone Methylation | GCTGCGGATCTACGAGATCG |
| Human | KDM2B | Histone Methylation | GCCACACACAAGGCACACGG |
| Human | KDM3A | Histone Methylation | TGGCTGGCCGACCTAACCAG |
| Human | KDM3A | Histone Methylation | TGCATTTGAAACATCCGATG |
| Human | KDM3A | Histone Methylation | TGTCTTGGGTCCTTACTAGG |
| Human | KDM3A | Histone Methylation | CATTCTGTAAGAGCGAAATG |
| Human | KDM3A | Histone Methylation | TTATCACAAAGCCAATAGTG |
| Human | KDM3B | Histone Methylation | GGATGGGTCATGCATCAATG |
| Human | KDM3B | Histone Methylation | GGTACTCAAGACAATTGACG |
| Human | KDM3B | Histone Methylation | CACATTTGCGACAAACCCAG |
| Human | KDM3B | Histone Methylation | GTAGAGTGGTGAGATCAAGT |
| Human | KDM3B | Histone Methylation | TTCGAGTCTCCATTGCACAA |
| Human | KDM4A | Histone Methylation | TGAGCCTGACCACTTCACAC |
| Human | KDM4A | Histone Methylation | TTGCCACAGCACATGAAACG |
| Human | KDM4A | Histone Methylation | ATATTTCTTCAGCATTAACG |

| | | | |
|---|---|---|---|
| Human | KDM4A | Histone Methylation | AGCACCGAATAGGGACAAAG |
| Human | KDM4A | Histone Methylation | CTGTGGGAGAGCCATGCACG |
| Human | KDM4B | Histone Methylation | ATGTCATCATACGTCTGCCG |
| Human | KDM4B | Histone Methylation | CGTGATTGAGCGCCACCCTG |
| Human | KDM4B | Histone Methylation | CGGCACCGATGACGCGACAG |
| Human | KDM4B | Histone Methylation | CAGATCACGTGGATCCACCT |
| Human | KDM4B | Histone Methylation | TGCCTCAAGCACAAGTCGGG |
| Human | KDM4C | Histone Methylation | AGATGAAGTCGTTACATCGG |
| Human | KDM4C | Histone Methylation | GCAAGAGTATAATGCAACAG |
| Human | KDM4C | Histone Methylation | GGTCATCTGTGACTGAGTCG |
| Human | KDM4C | Histone Methylation | AAACGGTCATCACGAAGCAT |
| Human | KDM4C | Histone Methylation | TTCTCATAGCCACTAGACAA |
| Human | KDM4D | Histone Methylation | AACCTGAACGCTATGACCTG |
| Human | KDM4D | Histone Methylation | ATAATCAATCCATCGCGGAG |
| Human | KDM4D | Histone Methylation | CCATGGTTTCAACTGCGCAG |
| Human | KDM4D | Histone Methylation | AAATCGGTGAATTATAGATG |
| Human | KDM4D | Histone Methylation | CCACTAACTGCAGATCGGCG |
| Human | KDM5A | Histone Methylation | GTGTCCTAAATGTGTCGCCG |
| Human | KDM5A | Histone Methylation | CGAGCTTGGCTGATGACACA |
| Human | KDM5A | Histone Methylation | GGCACAGTCCTATGACACTT |
| Human | KDM5A | Histone Methylation | ACAGACTGAGGTACCTTGGG |
| Human | KDM5A | Histone Methylation | GAGCTACAAACTACCAATGG |
| Human | KDM5B | Histone Methylation | GCGAACTCGGATACGCTGAA |
| Human | KDM5B | Histone Methylation | GCAGTGGGCTCACATATCAG |
| Human | KDM5B | Histone Methylation | GAGTTCACATTGAATCATAG |
| Human | KDM5B | Histone Methylation | AACTCGGATACGCTGAAGGG |
| Human | KDM5B | Histone Methylation | AGGCTATTCAATGAATGTCG |
| Human | KDM5C | Histone Methylation | AGATTCCCAATGTAGAACGG |
| Human | KDM5C | Histone Methylation | CCAAGGCTACAACTTTGCCG |
| Human | KDM5C | Histone Methylation | GTGCCCAAAGTGTGTCATGG |
| Human | KDM5C | Histone Methylation | AGTACAGACTGTTCCAACAC |
| Human | KDM5C | Histone Methylation | TGAGGGTGTTGGGATTCATG |
| Human | KDM5D | Histone Methylation | TAAGGCCCGACATGGAACCG |
| Human | KDM5D | Histone Methylation | CACTGTGTATGCGCTCACGA |
| Human | KDM5D | Histone Methylation | GGCAGGAACTCGTCACACCC |
| Human | KDM5D | Histone Methylation | GGTGGAGGCCTATCAAGCTG |
| Human | KDM5D | Histone Methylation | TTGCCAAGTATGCTCCCGTG |
| Human | KDM6A | Histone Methylation | CCTAGCAATTCAGTAACACA |
| Human | KDM6A | Histone Methylation | AGGATTCATAGAGAGTGCCT |
| Human | KDM6A | Histone Methylation | TCTGGTATTCAGAATCAGAA |
| Human | KDM6A | Histone Methylation | GGTCAGAGTTCACATTCGGC |
| Human | KDM6A | Histone Methylation | TGTAGCACATCAAGAACGCT |

| | | | |
|---|---|---|---|
| Human | KDM6B | Histone Methylation | TGGCCCTGGAACGATACGAG |
| Human | KDM6B | Histone Methylation | GCAGTCGGAAACCGTTCTTG |
| Human | KDM6B | Histone Methylation | GAAACCGAAGATCAACACTG |
| Human | KDM6B | Histone Methylation | GGCAACATGCTGAGCCACGT |
| Human | KDM6B | Histone Methylation | CACAGCGCCCTTCGATACGG |
| Human | KDM7A | Histone Methylation | GTATCGCTATGATAAACCAG |
| Human | KDM7A | Histone Methylation | AAGTTCCACCGAAGTCAATG |
| Human | KDM7A | Histone Methylation | CATCCTAGAGCTCCACACAA |
| Human | KDM7A | Histone Methylation | CATCTAGTACTTACTCAAGG |
| Human | KDM7A | Histone Methylation | GTCACACAAAGAAACGTGCA |
| Human | KDM8 | Histone Methylation | GAACTCGTTGACCGTCATGA |
| Human | KDM8 | Histone Methylation | TCCTGGGAGAAGCTCAACAC |
| Human | KDM8 | Histone Methylation | TGGTTCGAGGTACACAGATG |
| Human | KDM8 | Histone Methylation | TGAACTCGTTGACCGTCATG |
| Human | KDM8 | Histone Methylation | CAGAGAAGGCACTTTATGGG |
| Human | KIAA2026 | Chromatin Readers | CATGTACCTCAACAAGGCGT |
| Human | KIAA2026 | Chromatin Readers | TGATGTACCACTAGGAACTG |
| Human | KIAA2026 | Chromatin Readers | TTGGCAGTGGAGGATTACTG |
| Human | KIAA2026 | Chromatin Readers | TGGAGCAGAACTGTAACTGG |
| Human | KIAA2026 | Chromatin Readers | TTGTGGTGGAACAATAAACC |
| Human | KMT2A | Histone Methylation | AAGATCAGTAGCGGTCCCGG |
| Human | KMT2A | Histone Methylation | ATCCTCTATAAACCGCCGAG |
| Human | KMT2A | Histone Methylation | AGAAAGGACGTCGATCGAGG |
| Human | KMT2A | Histone Methylation | GTACAAATTGTACGACGGAG |
| Human | KMT2A | Histone Methylation | GGATGTGAGACAGCAACCCA |
| Human | KMT2B | Histone Methylation | GGCCTCAGTTAAGGAACCGA |
| Human | KMT2B | Histone Methylation | CGATTCGTAGATCTTCAGGT |
| Human | KMT2B | Histone Methylation | ATTCTGGAGTATCGGCCATG |
| Human | KMT2B | Histone Methylation | AAGCGGGAGAAGTTCTACGA |
| Human | KMT2B | Histone Methylation | TGCCGTCGGTTCCTTAACTG |
| Human | KMT2C | Histone Methylation | ATTCAGACATTAGGCCATCG |
| Human | KMT2C | Histone Methylation | CAAGATTCTACTATAACCCA |
| Human | KMT2C | Histone Methylation | GCACACGGTCTAGTTCTCAG |
| Human | KMT2C | Histone Methylation | CTAGTGACCACTCCACACAA |
| Human | KMT2C | Histone Methylation | TTCGTGTGCACCTAATTGTG |
| Human | KMT2D | Histone Methylation | ATTGATGCTACGTTGACCGG |
| Human | KMT2D | Histone Methylation | GGTGGAAATTCCCGCCAACG |
| Human | KMT2D | Histone Methylation | CCAAGTCATCTCAGTACCGG |
| Human | KMT2D | Histone Methylation | CTTCCCTATGGGACTCAACG |
| Human | KMT2D | Histone Methylation | CCAAGCTCAGTGCTCGACGG |
| Human | KMT2E | Histone Methylation | GGATCTGTAGTTATGCGTAG |
| Human | KMT2E | Histone Methylation | GTGAGTTGGTGTAGCATACG |

| | | | |
|---|---|---|---|
| Human | KMT2E | Histone Methylation | GGGCAGTGCTACTACAACGC |
| Human | KMT2E | Histone Methylation | AATGCAGTATATAATTCCAC |
| Human | KMT2E | Histone Methylation | CATGAGCATAGTGATCCCAT |
| Human | KNTC1 | Chromatin Remodeling | AATCAATTGGACGAGACAGG |
| Human | KNTC1 | Chromatin Remodeling | ACTCTCCTTCATCAGTCACG |
| Human | KNTC1 | Chromatin Remodeling | GTGTACCTACCATGGACTCG |
| Human | KNTC1 | Chromatin Remodeling | TCTCTATAGTCAACCCAGAG |
| Human | KNTC1 | Chromatin Remodeling | GTTGACTATAGAGAGTAAGC |
| Human | L3MBTL1 | Chromatin Readers | CGCCAAGTGGACCATCGATG |
| Human | L3MBTL1 | Chromatin Readers | GGAGGACCATCGGATAAAGG |
| Human | L3MBTL1 | Chromatin Readers | GAATCTTTCGATACTTCGGA |
| Human | L3MBTL1 | Chromatin Readers | AGACAAGGGACGGGTTCATG |
| Human | L3MBTL1 | Chromatin Readers | GTATAAGCCGCTGAACATGG |
| Human | L3MBTL2 | Chromatin Readers | CTTCTCACCTCAATACTCCG |
| Human | L3MBTL2 | Chromatin Readers | TGAGCTCACACCAGCCGACG |
| Human | L3MBTL2 | Chromatin Readers | ATGCACCACTCGTTTCACCG |
| Human | L3MBTL2 | Chromatin Readers | CTCATCAACTCACTCCGTGG |
| Human | L3MBTL2 | Chromatin Readers | GGTCCCACTCTATGACCAGT |
| Human | L3MBTL3 | Chromatin Readers | GGAAATGACGGACTAGCCGG |
| Human | L3MBTL3 | Chromatin Readers | AGCAGTTGGTACCATCCAAG |
| Human | L3MBTL3 | Chromatin Readers | GATTCGGCTGTACTAAAGCA |
| Human | L3MBTL3 | Chromatin Readers | ACATGGTGGATGCTCAACCC |
| Human | L3MBTL3 | Chromatin Readers | ATACCAACTCGAAAGCCCGA |
| Human | L3MBTL4 | Chromatin Readers | CTGTGATGTCCCGAATAACG |
| Human | L3MBTL4 | Chromatin Readers | CGAGGAATAGGCCATATCCG |
| Human | L3MBTL4 | Chromatin Readers | CAGAATACCGATGGATGTCG |
| Human | L3MBTL4 | Chromatin Readers | AGCCCTGATATCCACCCGAT |
| Human | L3MBTL4 | Chromatin Readers | GTGACATCACACCATCCGAT |
| Human | LBR | Chromatin Readers | GCAGGCGACAGGTCATTCCG |
| Human | LBR | Chromatin Readers | GCATGAATCCAAATCCATCG |
| Human | LBR | Chromatin Readers | ATACCCATCCAATCAATCCG |
| Human | LBR | Chromatin Readers | GTGACCTTGATCGACTCCCT |
| Human | LBR | Chromatin Readers | GGAAGCGTTGTTGACGACCA |
| Human | LDB1 | Transcription Elongation | GTGGTCGGAGACTCACTCGG |
| Human | LDB1 | Transcription Elongation | CACTCGGAGGTAGTTGAGAG |
| Human | LDB1 | Transcription Elongation | GGAGTGTGACAATCTCTGGT |
| Human | LDB1 | Transcription Elongation | TCATGTTTGACGACATGATG |
| Human | LDB1 | Transcription Elongation | CGTGACATGAGCTCTTGCAT |
| Human | LEO1 | Transcription Elongation | GGAACCTGATCGATACGATG |
| Human | LEO1 | Transcription Elongation | GCAATGCTCGGATAGTCAAG |
| Human | LEO1 | Transcription Elongation | TGAGAGACAACAGCTATCTG |
| Human | LEO1 | Transcription Elongation | GCGAAGCTGAGCGTAAAGGT |

| | | | |
|---|---|---|---|
| Human | LEO1 | Transcription Elongation | GTGGCACTGTCCGTAGAGTG |
| Human | LMNA | DNA and RNA Modifications | GCTGCGCAACAAGTCCAATG |
| Human | LMNA | DNA and RNA Modifications | CCGTCATGAGACCCGACTGG |
| Human | LMNA | DNA and RNA Modifications | CGACTGGTGGAGATTGACAA |
| Human | LMNA | DNA and RNA Modifications | AGTTGATGAGAGCCGTACGC |
| Human | LMNA | DNA and RNA Modifications | AGTTTAAGGAGCTGAAAGCG |
| Human | LMNB1 | DNA and RNA Modifications | GTGTAGGATCAACCAATGGG |
| Human | LMNB1 | DNA and RNA Modifications | TGTACGTACACTACGACTTG |
| Human | LMNB1 | DNA and RNA Modifications | CTGGTCTCGTTAATCTCCTG |
| Human | LMNB1 | DNA and RNA Modifications | AAAGGCTCTCAATTCTCATG |
| Human | LMNB1 | DNA and RNA Modifications | TGGAAGAACCAGAACTCGTG |
| Human | LMNB2 | DNA and RNA Modifications | GCTCAAGAACAACTCGGACA |
| Human | LMNB2 | DNA and RNA Modifications | CTCGATGCTGACGCTACCCG |
| Human | LMNB2 | DNA and RNA Modifications | GAGGGAGGGCGAGCTTACGG |
| Human | LMNB2 | DNA and RNA Modifications | GCTGCCCTCAGCGACAAGCG |
| Human | LMNB2 | DNA and RNA Modifications | ACCTACCGTGACCATCTGGC |
| Human | MAPK14 | Other Histone Modifications | TGATGAAATGACAGGCTACG |
| Human | MAPK14 | Other Histone Modifications | GGGACCTCCTTATAGATGAG |
| Human | MAPK14 | Other Histone Modifications | CTCAGATAAGAGAATTACAG |
| Human | MAPK14 | Other Histone Modifications | CAGGCTACGTGGCCACTAGG |
| Human | MAPK14 | Other Histone Modifications | AGCTCCTGCCGGTAGAACGT |
| Human | MAPK3 | Other Histone Modifications | GCAGTTGCAGTACATCGGCG |
| Human | MAPK3 | Other Histone Modifications | CCACGTGCGCAAGACTCGCG |
| Human | MAPK3 | Other Histone Modifications | AGTAGGTCTGATGTTCGAAG |
| Human | MAPK3 | Other Histone Modifications | GAATTGTATCATCAACATGA |
| Human | MAPK3 | Other Histone Modifications | GTACTATGACCCGACGGATG |
| Human | MAPK8 | Other Histone Modifications | TCGCTACTACAGAGCACCCG |
| Human | MAPK8 | Other Histone Modifications | TAGTGGATTTATGGTCTGTG |
| Human | MAPK8 | Other Histone Modifications | TGTTGTGTTCTGAGTCAGCT |
| Human | MAPK8 | Other Histone Modifications | GGGAACACACAATAGAAGAG |
| Human | MAPK8 | Other Histone Modifications | AGAATCAGACTCATGCCAAG |
| Human | MAPK9 | Other Histone Modifications | AGTACCGTGTCACCACGTAA |
| Human | MAPK9 | Other Histone Modifications | TATCATCGACAGCCTTCAAG |
| Human | MAPK9 | Other Histone Modifications | CAGCAGTCTTGATGCCTCGA |
| Human | MAPK9 | Other Histone Modifications | CGTGGTGACACGGTACTACC |
| Human | MAPK9 | Other Histone Modifications | TAGTGATTGATCCTGACAAG |
| Human | MBD3 | DNA and RNA Modifications | TGGTGTTGAGCCATACGCCG |
| Human | MBD3 | DNA and RNA Modifications | TCAGCGCAGGCCTTGTCCAG |
| Human | MBD3 | DNA and RNA Modifications | CACTAGCACCATGCCCATCA |
| Human | MBD3 | DNA and RNA Modifications | GCGCTACGACTCCTCCAACC |
| Human | MBD3 | DNA and RNA Modifications | CGACTCCTCCAACCAGGTCA |
| Human | MBD4 | DNA and RNA Modifications | ACTTACCGGAGGTCATTCGG |

| | | | |
|---|---|---|---|
| Human | MBD4 | DNA and RNA Modifications | TAGGCGCTCACTAGAGGTGA |
| Human | MBD4 | DNA and RNA Modifications | TGCCGTAAGTCTGTCCCATG |
| Human | MBD4 | DNA and RNA Modifications | CCAAACTGAGCAGAAGCGAT |
| Human | MBD4 | DNA and RNA Modifications | CTTACCGGAGGTCATTCGGC |
| Human | MBD5 | Chromatin Readers | GAACCATGTAGCATTACTGA |
| Human | MBD5 | Chromatin Readers | CTTGAGTAGCAATAGTACCC |
| Human | MBD5 | Chromatin Readers | ATCCTCGGTTGCAACTCACA |
| Human | MBD5 | Chromatin Readers | CACCTTGTGAGTTGCAACCG |
| Human | MBD5 | Chromatin Readers | GAGGCAAAGAGTGTGACGGA |
| Human | MBD6 | DNA and RNA Modifications | AATGCTCCCTCATACAACTG |
| Human | MBD6 | DNA and RNA Modifications | GTAGTCAGAAAGTCTCGTCG |
| Human | MBD6 | DNA and RNA Modifications | AGTTGTATGAGGGAGCATTG |
| Human | MBD6 | DNA and RNA Modifications | AGCAATCGAAATACAGTAGG |
| Human | MBD6 | DNA and RNA Modifications | TCAGCACACCAAATAAACTG |
| Human | MBTD1 | Chromatin Readers | GTGGCTACACATATTAAACG |
| Human | MBTD1 | Chromatin Readers | TACATCAAACAAGAGCCATG |
| Human | MBTD1 | Chromatin Readers | GAGTAGAAGTGGTTGACAAG |
| Human | MBTD1 | Chromatin Readers | ATTGAACTTACTCCACCCAG |
| Human | MBTD1 | Chromatin Readers | GTGTCGAACACGAGTAGCAG |
| Human | MCM2 | DNA and RNA Modifications | CATCAATGAGACACACTCCT |
| Human | MCM2 | DNA and RNA Modifications | CATGCCCAACACGTATGGCG |
| Human | MCM2 | DNA and RNA Modifications | TCCTCGGTCAGCCAGAACCA |
| Human | MCM2 | DNA and RNA Modifications | CGGTGTCCCTCACCACACAC |
| Human | MCM2 | DNA and RNA Modifications | GGACAGAACCAGCATCCATG |
| Human | MCM3 | DNA and RNA Modifications | CAGTATGAGGAGTTCTACGT |
| Human | MCM3 | DNA and RNA Modifications | GGCATAGGTAGCATCAATGG |
| Human | MCM3 | DNA and RNA Modifications | GAAACGTAAGAAGCGAAGTG |
| Human | MCM3 | DNA and RNA Modifications | ATATCCACAGCACTACCCAA |
| Human | MCM3 | DNA and RNA Modifications | GCAGGAGGTAAGAGTCCGCG |
| Human | MCM4 | DNA and RNA Modifications | TGTGACACAAGCTCGGAGTG |
| Human | MCM4 | DNA and RNA Modifications | GGAAACCAGACATTTATGAG |
| Human | MCM4 | DNA and RNA Modifications | CGCACTGTACTACCAGAGCG |
| Human | MCM4 | DNA and RNA Modifications | CTTTCATTACGTACGCAGTG |
| Human | MCM4 | DNA and RNA Modifications | GGTAGTACAGTGCGACCAGG |
| Human | MCM5 | DNA and RNA Modifications | CTCGGCGAGTAAGTCCATCA |
| Human | MCM5 | DNA and RNA Modifications | TGTGTGACAAGGTCGTCCCT |
| Human | MCM5 | DNA and RNA Modifications | TTCCCAGACGTGTATACCTG |
| Human | MCM5 | DNA and RNA Modifications | CCTGATGGACTTACTCGCCG |
| Human | MCM5 | DNA and RNA Modifications | GGACAAGGCAGCATCCAACG |
| Human | MCM6 | DNA and RNA Modifications | GAATAAATCAATCATCCGTG |
| Human | MCM6 | DNA and RNA Modifications | CTAAGCAAATATGGACCTCG |
| Human | MCM6 | DNA and RNA Modifications | TGTTGCGCCAACCAACCCAA |

| | | | |
|---|---|---|---|
| Human | MCM6 | DNA and RNA Modifications | AGAAGGGACCTCTCTTCGAG |
| Human | MCM6 | DNA and RNA Modifications | ACAGGCTGGTACAAAGATTG |
| Human | MCM7 | DNA and RNA Modifications | ACCAGCACCGTGATACTACG |
| Human | MCM7 | DNA and RNA Modifications | AAGCCCAACTACTTACTGCG |
| Human | MCM7 | DNA and RNA Modifications | GTGCTGAGAGACTCCGTGAG |
| Human | MCM7 | DNA and RNA Modifications | CCAGCACCGTGATACTACGA |
| Human | MCM7 | DNA and RNA Modifications | AAATTGAGTCCACCAACTCG |
| Human | MECOM | Histone Methylation | AATATGAGTCATGCCAACCC |
| Human | MECOM | Histone Methylation | AACCTTGATAACGTCATACG |
| Human | MECOM | Histone Methylation | ACTATGGCGCCGGATATCCA |
| Human | MECOM | Histone Methylation | CACCAGATGTCACATGACAG |
| Human | MECOM | Histone Methylation | AGCCGGAAGGAAACAGACCA |
| Human | METTL14 | DNA and RNA Modifications | TGTGAAGCGTAGCACAGACG |
| Human | METTL14 | DNA and RNA Modifications | AAATCTAAATCTGACCGAGG |
| Human | METTL14 | DNA and RNA Modifications | AGAGATAGTACAATTCGACC |
| Human | METTL14 | DNA and RNA Modifications | AATGTCTTCAACAATTATCG |
| Human | METTL14 | DNA and RNA Modifications | TAGACCTCAGAATTTCATCA |
| Human | METTL3 | DNA and RNA Modifications | ATTCTGTGACTATGGAACCA |
| Human | METTL3 | DNA and RNA Modifications | GAGTTGATTGAGGTAAAGCG |
| Human | METTL3 | DNA and RNA Modifications | AGTTCCATGTGAATATCCCA |
| Human | METTL3 | DNA and RNA Modifications | AGAGAATGTCTAAACCTCTG |
| Human | METTL3 | DNA and RNA Modifications | ATGTTAAGGCCAGATCAGAG |
| Human | MIS12 | Chromatin Remodeling | TGTGGATCCAATGACCTACG |
| Human | MIS12 | Chromatin Remodeling | TAAGGAGACACCTTATAGTG |
| Human | MIS12 | Chromatin Remodeling | GACTGAATTATGTACTAAGC |
| Human | MIS12 | Chromatin Remodeling | TGCATTTGCGAATCTGCACT |
| Human | MIS12 | Chromatin Remodeling | TAGGGAGAGTTTAGTATCCC |
| Human | MIS18A | Chromatin Remodeling | CATGAGCGAAGACGCGTCGG |
| Human | MIS18A | Chromatin Remodeling | TTGAGACTTTGTGCTGCGCG |
| Human | MIS18A | Chromatin Remodeling | TGGCAGGCGTTCGGTCACTG |
| Human | MIS18A | Chromatin Remodeling | CAGCTCAGCGAGTCGCCCAG |
| Human | MIS18A | Chromatin Remodeling | CCTTGAGACTTTGTGCTGCG |
| Human | MIS18BP1 | Chromatin Remodeling | TCAATCAACTATGCTAACAG |
| Human | MIS18BP1 | Chromatin Remodeling | TTCATATAACAATACCTCGG |
| Human | MIS18BP1 | Chromatin Remodeling | AGAATGCCAGAGGAAATACA |
| Human | MIS18BP1 | Chromatin Remodeling | AAAGCAAACTATGAATCACC |
| Human | MIS18BP1 | Chromatin Remodeling | ATGAGGTATAATCTGTCCGC |
| Human | MLLT1 | Chromatin Readers | GAGCTTGTACCGGAACTCCG |
| Human | MLLT1 | Chromatin Readers | CTGCCGGCAAGACCAACCCG |
| Human | MLLT1 | Chromatin Readers | CAGCCATGGACAATCAGGTG |
| Human | MLLT1 | Chromatin Readers | AAGAAGACCAAACCATCCCA |
| Human | MLLT1 | Chromatin Readers | GTGGAATTGTGGGTAACATG |

| | | | |
|---|---|---|---|
| Human | MLLT10 | Chromatin Readers | ACCTTATTATAACGATCATG |
| Human | MLLT10 | Chromatin Readers | TTCAAGTACCCACTGGACCG |
| Human | MLLT10 | Chromatin Readers | GCCCGCTTATATCCTGCTGT |
| Human | MLLT10 | Chromatin Readers | TTGTACCTCTGGAATATACA |
| Human | MLLT10 | Chromatin Readers | ATACAGGGCACAAACCACAT |
| Human | MLLT3 | Chromatin Readers | GACTGGATGGTGTTCGTACG |
| Human | MLLT3 | Chromatin Readers | CAGCGGAGGTGATTCACTGG |
| Human | MLLT3 | Chromatin Readers | AACTTACTCACCATCACCAG |
| Human | MLLT3 | Chromatin Readers | AAAGAATGGTGAATGTGACA |
| Human | MLLT3 | Chromatin Readers | GTACGAACACCATCCAGTCG |
| Human | MLLT6 | Chromatin Readers | CGTTGGCAAATTGCACCTCG |
| Human | MLLT6 | Chromatin Readers | GCTGTGCCCACACAAAGACG |
| Human | MLLT6 | Chromatin Readers | TGCGTATGTTCGGACGAGAG |
| Human | MLLT6 | Chromatin Readers | GCCTCATGATCGCTTCAACA |
| Human | MLLT6 | Chromatin Readers | TCTTTCCAGACCGCTGACAA |
| Human | MNAT1 | Transcription Elongation | TATTTGGTGGTCTTACACCG |
| Human | MNAT1 | Transcription Elongation | TGAGTGGAGTACCACACTCA |
| Human | MNAT1 | Transcription Elongation | CACTACAGGATGCATTCAGT |
| Human | MNAT1 | Transcription Elongation | GAAACCGCCATGGACGATCA |
| Human | MNAT1 | Transcription Elongation | GTAGATTTACTGTTTGTGAG |
| Human | MORF4L1 | Chromatin Readers | GAAGGGAGCAGTATGCAGAG |
| Human | MORF4L1 | Chromatin Readers | TTTCTTCCGAGGAGGCTGAG |
| Human | MORF4L1 | Chromatin Readers | AAATACTTCATACATTACAG |
| Human | MORF4L1 | Chromatin Readers | CCGAAGCCTAAATTCCAGGA |
| Human | MORF4L1 | Chromatin Readers | CCCGAAGCCTAAATTCCAGG |
| Human | MPHOSPH8 | Chromatin Readers | GTACGACTCGTAATTGAATG |
| Human | MPHOSPH8 | Chromatin Readers | AAACCAACACAGGATTCCAG |
| Human | MPHOSPH8 | Chromatin Readers | ACTACAGCTTGTACACTACA |
| Human | MPHOSPH8 | Chromatin Readers | TTGTGATGAGGAGTCGCAGG |
| Human | MPHOSPH8 | Chromatin Readers | GTTTATTGGGACCTGCAACA |
| Human | MRPL20 | Control | CTGGATCCGAAAGTAGCGGT |
| Human | MRPL20 | Control | CCTCCTGGATCCGAAAGTAG |
| Human | MRPL20 | Control | GCCAGTAGGAGGCGACACGA |
| Human | MRPL20 | Control | AATACAGCAACAGTCCTCAG |
| Human | MRPL20 | Control | GCTCAACAGGAAAGTCCTAG |
| Human | MSH6 | Chromatin Readers | ATCACCACCTCCACTAACGT |
| Human | MSH6 | Chromatin Readers | AAGGCGAAGAACCTCAACGG |
| Human | MSH6 | Chromatin Readers | TGCCTGGCTAACTATAGTCG |
| Human | MSH6 | Chromatin Readers | CCTGGCTAACTATAGTCGAG |
| Human | MSH6 | Chromatin Readers | GTAGGACCATAGTCTATTGG |
| Human | MSL3 | Chromatin Readers | AAGAACTAATGAAATAAACG |
| Human | MSL3 | Chromatin Readers | ATTGTTACTACATTAACAGG |

| | | | |
|---|---|---|---|
| Human | MSL3 | Chromatin Readers | TGCACGTTCATGTTGGCATG |
| Human | MSL3 | Chromatin Readers | CAGGCACAAGCTTCCAGGAG |
| Human | MSL3 | Chromatin Readers | TGGCATGTGGCATAACGTGA |
| Human | MST1 | Other Histone Modifications | AGGCACCACATACCATTCAG |
| Human | MST1 | Other Histone Modifications | CACAGCCAATACCACCACTG |
| Human | MST1 | Other Histone Modifications | ACTCGCGCCCTGACTCCGTG |
| Human | MST1 | Other Histone Modifications | AATGACTTCCAAGTGCTCCG |
| Human | MST1 | Other Histone Modifications | GAGAGTGAGATGTGCACTGA |
| Human | MTF2 | Chromatin Readers | ATGCCTGGAAATGCTAGACG |
| Human | MTF2 | Chromatin Readers | GCTGCAGGTAGAATAGCATG |
| Human | MTF2 | Chromatin Readers | AACTGGGATAGATTGCACCC |
| Human | MTF2 | Chromatin Readers | ACCACTGAAGTGACCGAATG |
| Human | MTF2 | Chromatin Readers | ATCACACTGGAATCAATATG |
| Human | MUM1 | Chromatin Readers | AAATACGCTCTTACCTAACG |
| Human | MUM1 | Chromatin Readers | ATTCCTACTTCAAACGAACG |
| Human | MUM1 | Chromatin Readers | CTGCCCAACACGTCTCCGCG |
| Human | MUM1 | Chromatin Readers | GCAAGTGTGCTATACATCGA |
| Human | MUM1 | Chromatin Readers | GGACGAGGTGGACTACAAGA |
| Human | NAE1 | Control | ATGGACTAGTTGGTTATATG |
| Human | NAE1 | Control | TAGCTAAATATTTAGCACAG |
| Human | NAE1 | Control | TCAAAGAAGCAGTATCGGCA |
| Human | NAE1 | Control | CACAGCACTAAATACAACTC |
| Human | NAE1 | Control | CAATCCATGGAGTATGACTG |
| Human | NAF1 | Control | CCCTCAAACGACTGTAGCGG |
| Human | NAF1 | Control | TGGAGGAAGTGATATACAAG |
| Human | NAF1 | Control | AGATACCCTCGATCTTGCCA |
| Human | NAF1 | Control | TCAAGTATTATTGAACAACT |
| Human | NAF1 | Control | GAGTTATAGAAATGCTGAGG |
| Human | NARFL | Control | CTGTGTCCTCACAACAGGTG |
| Human | NARFL | Control | GGAGCACGTGTTCCGGCACG |
| Human | NARFL | Control | GCATACGCAGTACCACGCCG |
| Human | NARFL | Control | GCCAGCGATGCTCTAGACTG |
| Human | NARFL | Control | TGGTGCAGAGGCTCAAACGA |
| Human | NCBP1 | Transcription Elongation | CGGCACAGCGACGAGAACGA |
| Human | NCBP1 | Transcription Elongation | CTTGAACTACCACATAGTTG |
| Human | NCBP1 | Transcription Elongation | TTCAGTGTACCCAATGCCAA |
| Human | NCBP1 | Transcription Elongation | TACATCACATCAATGTGAGG |
| Human | NCBP1 | Transcription Elongation | GTGTACATCACATCAATGTG |
| Human | NCBP2 | Transcription Elongation | ATCGGCAGGACTACGATGCT |
| Human | NCBP2 | Transcription Elongation | ATATTACTCACGCGCAGATG |
| Human | NCBP2 | Transcription Elongation | AGGGCAGGCAATACGGCCGT |
| Human | NCBP2 | Transcription Elongation | CGTCCCATTTATGTACCGCA |

| | | | |
|---|---|---|---|
| Human | NCBP2 | Transcription Elongation | TGGGACGCAGGCTTTAAGGA |
| Human | NCOA1 | Histone Acetylation | GAGATACTAGGATTGACCGA |
| Human | NCOA1 | Histone Acetylation | TTGACCGAGGGATTTACTCG |
| Human | NCOA1 | Histone Acetylation | CTGGACTACCAGTTCAAATG |
| Human | NCOA1 | Histone Acetylation | TTTGAAACAGTGATGACTCG |
| Human | NCOA1 | Histone Acetylation | GCAAACTCTAAACAGACCTC |
| Human | NCOA3 | Histone Acetylation | TCAGCCACGAGCTATGATGG |
| Human | NCOA3 | Histone Acetylation | TTTGGACTCCCACGATTACG |
| Human | NCOA3 | Histone Acetylation | CCTGTAACAAATGATCGACA |
| Human | NCOA3 | Histone Acetylation | AGGACTAGACACTCGACCAA |
| Human | NCOA3 | Histone Acetylation | GCAGCAGCATCTAACCAACT |
| Human | NDC80 | Chromatin Remodeling | GGGAAATACTCACGTTTGAG |
| Human | NDC80 | Chromatin Remodeling | GAAATTAAATGGTCTCAATG |
| Human | NDC80 | Chromatin Remodeling | TGTCAGGAAGTTCGTATGAG |
| Human | NDC80 | Chromatin Remodeling | GGATCCCGGAATAGTCAACT |
| Human | NDC80 | Chromatin Remodeling | TCATACATGGCCTCACATTG |
| Human | NDUFAB1 | Control | ACCTGCGCGAGCACTAAGGC |
| Human | NDUFAB1 | Control | ATGCCCTCTAACGTCAAAGG |
| Human | NDUFAB1 | Control | GAGGCATGTCGCTATACTGG |
| Human | NDUFAB1 | Control | TGAAAGGACACGAGACGCCA |
| Human | NDUFAB1 | Control | TGGCCATGATAATCTCCACT |
| Human | NELFA | Transcription Elongation | TCCATGCGGGAGAGCGACAC |
| Human | NELFA | Transcription Elongation | CGGGCGCCCATGTACAACAG |
| Human | NELFA | Transcription Elongation | AAGAACCTGTCCCTCACGGT |
| Human | NELFA | Transcription Elongation | GGAGAAGGAAGACTCTCGGT |
| Human | NELFA | Transcription Elongation | GGTGTCTTCATAAGATGCCG |
| Human | NELFB | Transcription Elongation | GCTCATCGAATACCGACTGG |
| Human | NELFB | Transcription Elongation | TGTCAGCCATCGCTTCGGAG |
| Human | NELFB | Transcription Elongation | TGGACCTGCACGGGACGCCG |
| Human | NELFB | Transcription Elongation | TGGACGCCTGCATCCGAGAG |
| Human | NELFB | Transcription Elongation | CTGCATCCGAGAGCGGTTCG |
| Human | NELFE | Transcription Elongation | AAGAATCCCATTCTTACCGA |
| Human | NELFE | Transcription Elongation | TCTTCCTCGCTCAGTCCGGG |
| Human | NELFE | Transcription Elongation | TCATACAGAGATTTCCTCTG |
| Human | NELFE | Transcription Elongation | AGCGTTCTCGAACCCTTGAG |
| Human | NELFE | Transcription Elongation | ACAGCCCTAAGGGTTGCCAC |
| Human | NFRKB | Chromatin Remodeling | AGCTGTAGTCACAACAACAG |
| Human | NFRKB | Chromatin Remodeling | AACCAGCTAAGTCTAGCTCG |
| Human | NFRKB | Chromatin Remodeling | GTGGCATACTGGAGTCACTG |
| Human | NFRKB | Chromatin Remodeling | AGTATTGTACTCACACCCGA |
| Human | NFRKB | Chromatin Remodeling | CTGGGACGTGGACACAACCG |
| Human | NOC4L | Control | CCACACCTTGTACTTCCGTG |

| | | | |
|---|---|---|---|
| Human | NOC4L | Control | GAAAGGCAGGACCATGAGCA |
| Human | NOC4L | Control | AGGAAGTCGATCATGAGCGT |
| Human | NOC4L | Control | TGGGCTGGGATAAACTCCAG |
| Human | NOC4L | Control | GTACTTGACGTGAAAGACAG |
| Human | NOP16 | Control | TCTGCCGTACCGATTTAGCG |
| Human | NOP16 | Control | TGTAGCGTACATAGTCAATG |
| Human | NOP16 | Control | GGGACCACGCTAAATCGGTA |
| Human | NOP16 | Control | GAACCGGAATGCTCGACGGA |
| Human | NOP16 | Control | GTCTGAACCGGAATGCTCGA |
| Human | NSD1 | Histone Methylation | GCCCTATCGGCAGTACTACG |
| Human | NSD1 | Histone Methylation | AATTCAAGAGACGCCCATGG |
| Human | NSD1 | Histone Methylation | GAGGAAAGCCTTCTAGCCCG |
| Human | NSD1 | Histone Methylation | TATGCATGATAGTAAGACGA |
| Human | NSD1 | Histone Methylation | AAGCACATAAAGATGAACGG |
| Human | NSL1 | Chromatin Remodeling | TTGCCTGCATTAATTGAACA |
| Human | NSL1 | Chromatin Remodeling | CGCACCCGAAAGTCTTCTCG |
| Human | NSL1 | Chromatin Remodeling | GCATTAATTGAACAAGGAGA |
| Human | NSL1 | Chromatin Remodeling | TTGTTCAATTAATGCAGGCA |
| Human | NSL1 | Chromatin Remodeling | CCAGTGGATGTACAACAGGG |
| Human | NSMCE1 | Control | GTCCTCCAACTTATCTACGG |
| Human | NSMCE1 | Control | GATGACCCATGGCGTGCTAG |
| Human | NSMCE1 | Control | TGCAGATCTTCACCGCGTCG |
| Human | NSMCE1 | Control | CACGCACTTGGGATCTCGTG |
| Human | NSMCE1 | Control | TCTCTTTATCTCAATATACA |
| Human | NSUN2 | DNA and RNA Modifications | GCTGCGGATTGCAACACGCG |
| Human | NSUN2 | DNA and RNA Modifications | ACTTACAGGTTATTAACACG |
| Human | NSUN2 | DNA and RNA Modifications | TGCTCGTCCATCAAGCCAAG |
| Human | NSUN2 | DNA and RNA Modifications | GAGCTCAAGATCGTGCCCGA |
| Human | NSUN2 | DNA and RNA Modifications | CAGCTGCGGATTGCAACACG |
| Human | NUF2 | Chromatin Remodeling | GCTTTGGAGAAATACCACGA |
| Human | NUF2 | Chromatin Remodeling | ACAGTTAAACGCCGCACACC |
| Human | NUF2 | Chromatin Remodeling | TGAGTAACTAAATTGCTGAA |
| Human | NUF2 | Chromatin Remodeling | AATAGTATATGGAATTCGAC |
| Human | NUF2 | Chromatin Remodeling | ATTAGCCAGTATCTTAAAGG |
| Human | NUP62 | Control | CGAGCACCTGAACACGTCCG |
| Human | NUP62 | Control | CAGTTGCTATTGACGCAAAG |
| Human | NUP62 | Control | GAGCCAATGTTGAAACCGGA |
| Human | NUP62 | Control | GCAGTGGATCGACCAGAACT |
| Human | NUP62 | Control | TGAGCCAATGTTGAAACCGG |
| Human | OIP5 | Chromatin Remodeling | CTGCGTATCCCACTCCATGG |
| Human | OIP5 | Chromatin Remodeling | GGACCTGTCGCGGTCCCTCG |
| Human | OIP5 | Chromatin Remodeling | GTGAACCTTCAATGCCAACT |

| | | | |
|---|---|---|---|
| Human | OIP5 | Chromatin Remodeling | CCTTCAATGCCAACTAGGAA |
| Human | OIP5 | Chromatin Remodeling | CAGCATGGGTAGAATACAGA |
| Human | ORC6 | Control | GCTTTAGTGCCGGCGCCATG |
| Human | ORC6 | Control | TACAGCTAAACTGTACAGCT |
| Human | ORC6 | Control | GACCTACTTACCCTGTCCAA |
| Human | ORC6 | Control | ACCTACTTACCCTGTCCAAG |
| Human | ORC6 | Control | GAGGCCGACACACTTCACCC |
| Human | PAF1 | Transcription Elongation | CCATCCAGACCCAGGCCCAG |
| Human | PAF1 | Transcription Elongation | CATCTCCAATGAGAAGCCTG |
| Human | PAF1 | Transcription Elongation | AGACGTTGAAGAAACGAAAG |
| Human | PAF1 | Transcription Elongation | TGTGAAGCAGCAGTTTACCG |
| Human | PAF1 | Transcription Elongation | GAAACGAAAGCGGGACCAGG |
| Human | PAK2 | Other Histone Modifications | GATTTCGTATGATCCGGTCG |
| Human | PAK2 | Other Histone Modifications | TTTCGTATGATCCGGTCGCG |
| Human | PAK2 | Other Histone Modifications | TGTCTCCACAGATTTACACA |
| Human | PAK2 | Other Histone Modifications | CCAGTATGGCGTTCCGACCA |
| Human | PAK2 | Other Histone Modifications | ACAGACCGTGTGTAAATCTG |
| Human | PARP1 | Other Histone Modifications | TACCGATCACCGTACCCACA |
| Human | PARP1 | Other Histone Modifications | CGATGCCTATTACTGCACTG |
| Human | PARP1 | Other Histone Modifications | TTCATAACCGAAGATTGCTG |
| Human | PARP1 | Other Histone Modifications | ATACTCGTTATATAGTAGAG |
| Human | PARP1 | Other Histone Modifications | AGCTAGGCATGATTGACCGC |
| Human | PARP10 | Other Histone Modifications | ATGGACTCACACGAACGACG |
| Human | PARP10 | Other Histone Modifications | GGTGCTGTACCACGGCACGA |
| Human | PARP10 | Other Histone Modifications | CACGCCCTTCCCGTAGACCG |
| Human | PARP10 | Other Histone Modifications | GAACACCGCCTTATGGCCAT |
| Human | PARP10 | Other Histone Modifications | TTCAAGGCCTGTGTCCAACG |
| Human | PARP11 | Other Histone Modifications | GACACGTCAGATACCCAGTG |
| Human | PARP11 | Other Histone Modifications | GCAAAGATGACATAAAGCAT |
| Human | PARP11 | Other Histone Modifications | GCTAACTGAACACTGACTGT |
| Human | PARP11 | Other Histone Modifications | TGGACACGTCAGATACCCAG |
| Human | PARP11 | Other Histone Modifications | GTATTCACATTCTCCCAGTG |
| Human | PARP14 | Other Histone Modifications | AATGATTATATAATCACCGG |
| Human | PARP14 | Other Histone Modifications | TAATAGGTACACACACCTCG |
| Human | PARP14 | Other Histone Modifications | ACCACGTGATCCATGCAGTG |
| Human | PARP14 | Other Histone Modifications | AACGGAGTTGACAACAACAT |
| Human | PARP14 | Other Histone Modifications | GGTGCTGGAAGCTTGATCAG |
| Human | PARP15 | Other Histone Modifications | TGAACAGAGGAAGTACACAT |
| Human | PARP15 | Other Histone Modifications | GAAAGTTCAGTGATGCAGAG |
| Human | PARP15 | Other Histone Modifications | GCTCCTCACCATACTCCGGG |
| Human | PARP15 | Other Histone Modifications | GTATATACAAATGACGATGA |
| Human | PARP15 | Other Histone Modifications | TCCGTGCATAAGTTCTCTGG |

| | | | |
|---|---|---|---|
| Human | PARP16 | Other Histone Modifications | GTGACCACGAAGTACTTGGG |
| Human | PARP16 | Other Histone Modifications | GCAGACATCCTTGTTCGGAG |
| Human | PARP16 | Other Histone Modifications | ATATCATGACGGTAAACCAA |
| Human | PARP16 | Other Histone Modifications | CAAAGTCCTTACAGTCGCCG |
| Human | PARP16 | Other Histone Modifications | GGTGCAGGAACAGGCGTGTG |
| Human | PARP2 | Other Histone Modifications | TCAGCGTTCGAATTCCATGG |
| Human | PARP2 | Other Histone Modifications | TCAAGGTAAACATAGCACCA |
| Human | PARP2 | Other Histone Modifications | CGAATTCCATGGCGGCGCGG |
| Human | PARP2 | Other Histone Modifications | GTACCGCATACGGACCTGGT |
| Human | PARP2 | Other Histone Modifications | ATGGTATAGTCGCTGTGTGT |
| Human | PARP3 | Other Histone Modifications | GTGGACATGTTGGATCCACG |
| Human | PARP3 | Other Histone Modifications | AGAGAAGCGCATAATCCGCG |
| Human | PARP3 | Other Histone Modifications | TGCCCGGGTGAGACACAAAG |
| Human | PARP3 | Other Histone Modifications | CAGAGAGCACCATATCAACA |
| Human | PARP3 | Other Histone Modifications | AGCAAGCAACAGATTGCACG |
| Human | PARP4 | Other Histone Modifications | TGTTCCGTGACCCTCGAGCA |
| Human | PARP4 | Other Histone Modifications | ATGCCCAAAGAAGTGAACCT |
| Human | PARP4 | Other Histone Modifications | GAGTCGATCATTGAGAAACA |
| Human | PARP4 | Other Histone Modifications | TAAGTCCATACACTTTCCGA |
| Human | PARP4 | Other Histone Modifications | TGGGACAAGACGCCGACTGT |
| Human | PARP6 | Other Histone Modifications | TGTCAAACTACCTCTCAGCA |
| Human | PARP6 | Other Histone Modifications | AGACCCGATCCATTCAGTCA |
| Human | PARP6 | Other Histone Modifications | ATATGGAACTATCGATGACG |
| Human | PARP6 | Other Histone Modifications | TTGACTTGATTACCTGAACG |
| Human | PARP6 | Other Histone Modifications | TCTCACCACTAGGACATCAA |
| Human | PARP8 | Other Histone Modifications | AGCTCCCGTTCACATACGGT |
| Human | PARP8 | Other Histone Modifications | TTTGTGTTGTGCTCACATCG |
| Human | PARP8 | Other Histone Modifications | TGCATTCCAGTACGAGACCG |
| Human | PARP8 | Other Histone Modifications | TTTGATTACCAATTACTCGG |
| Human | PARP8 | Other Histone Modifications | CACTTCCATACATTGCACCA |
| Human | PARP9 | Other Histone Modifications | TGTTCATAACCACCTCAATG |
| Human | PARP9 | Other Histone Modifications | GATGTAGAGATTAATCAACA |
| Human | PARP9 | Other Histone Modifications | TTACCGTCTGCCATTCAATG |
| Human | PARP9 | Other Histone Modifications | AGAATGTACTCGACACCTTG |
| Human | PARP9 | Other Histone Modifications | TGTACTCGACACCTTGCGGT |
| Human | PBRM1 | Chromatin Readers | AGGATGAAGTTCTTACCCGG |
| Human | PBRM1 | Chromatin Readers | AGGAGAGACGATATCGAGGA |
| Human | PBRM1 | Chromatin Readers | AAGTATTGGAACGAGCAAGA |
| Human | PBRM1 | Chromatin Readers | ATACATTGAAGGACTCAGTG |
| Human | PBRM1 | Chromatin Readers | ACACCCAGACTACTCTTTCG |
| Human | PCID2 | Transcription Elongation | AGTCGTGGGTTTGCAACATG |
| Human | PCID2 | Transcription Elongation | GTAACATACAAATACTACGT |

| | | | |
|---|---|---|---|
| Human | PCID2 | Transcription Elongation | CGTGTACTTTCAACACACCG |
| Human | PCID2 | Transcription Elongation | CCAGGTGCACTTATGCAGTG |
| Human | PCID2 | Transcription Elongation | TCTGGCTAACTTGATATACA |
| Human | PGK1 | Control | TTGCTGTGCCAAATGGAACA |
| Human | PGK1 | Control | AGTGACAAAGTCAACAGGCA |
| Human | PGK1 | Control | GGAGCTGAACTACTTTGCAA |
| Human | PGK1 | Control | AGTCAGCCATGTGAGCACTG |
| Human | PGK1 | Control | GCTTTCTAACAAGCTGACGC |
| Human | PHF1 | Histone Methylation | GGAGACAAACTACGGCACAG |
| Human | PHF1 | Histone Methylation | CGATCGCAAAGACACACTGG |
| Human | PHF1 | Histone Methylation | CAGTCCCAGGAGACAAACTA |
| Human | PHF1 | Histone Methylation | CAGGCCGTACTCTCCGAGCA |
| Human | PHF1 | Histone Methylation | ATGAGGCAGTCATCGAGTGG |
| Human | PHF10 | Chromatin Readers | AGGTTATCCAGGTACCTCAA |
| Human | PHF10 | Chromatin Readers | ATTTAATAGCAACTTAAACC |
| Human | PHF10 | Chromatin Readers | TTTGCCAACAGAGCGAACAA |
| Human | PHF10 | Chromatin Readers | CCCTTCAGATACATTGCCAG |
| Human | PHF10 | Chromatin Readers | TTTGACTTGCAGACACATGT |
| Human | PHF11 | Chromatin Readers | GGTACACACCGGTGGGAAGG |
| Human | PHF11 | Chromatin Readers | TTCCCGATCATCGCTCAAAG |
| Human | PHF11 | Chromatin Readers | CTGCAGCACAGTCATGGCCC |
| Human | PHF11 | Chromatin Readers | GAGGCGCTCCTCCTTCCCAC |
| Human | PHF11 | Chromatin Readers | TGTACATGATTGCCTGAGAG |
| Human | PHF12 | Chromatin Readers | GTACGACTTGGACACATCAG |
| Human | PHF12 | Chromatin Readers | GAATCACATCGAACATGTGG |
| Human | PHF12 | Chromatin Readers | AATTATTCCAAATCAACCCA |
| Human | PHF12 | Chromatin Readers | TTCGCAGCATGTCGTCAAAG |
| Human | PHF12 | Chromatin Readers | CACTACCAGGCCTCTCACAC |
| Human | PHF13 | Chromatin Readers | AGTGGCACAAGATCCACTCG |
| Human | PHF13 | Chromatin Readers | AAACGGACTATCGTCCGGCA |
| Human | PHF13 | Chromatin Readers | GTGGCACAAGATCCACTCGA |
| Human | PHF13 | Chromatin Readers | ACTCATTACACTCGATCATG |
| Human | PHF13 | Chromatin Readers | TCCGCCGTTCCAACCGCTCG |
| Human | PHF14 | Chromatin Readers | ATGGAACCAAACGATCAAGG |
| Human | PHF14 | Chromatin Readers | ATTACGACCAGTAACACTAA |
| Human | PHF14 | Chromatin Readers | CAGATGGAACCAAACGATCA |
| Human | PHF14 | Chromatin Readers | GAACTGTGTCCTAATCAGGA |
| Human | PHF14 | Chromatin Readers | TGCGGAAAGCAGAACTCATG |
| Human | PHF19 | Chromatin Readers | AATTATCTTCGAAAGTCACG |
| Human | PHF19 | Chromatin Readers | TTGATGTCCAAACTGACGGA |
| Human | PHF19 | Chromatin Readers | TGGTCCTCGATCTGTCACTG |
| Human | PHF19 | Chromatin Readers | CAAACATCATGGGCTCATTG |

| | | | |
|---|---|---|---|
| Human | PHF19 | Chromatin Readers | CCAGTGCGAAGATGCAGCGT |
| Human | PHF2 | Histone Methylation | GTCACACACCTCATCTGACG |
| Human | PHF2 | Histone Methylation | ACGTCACTGACATAGAACGT |
| Human | PHF2 | Histone Methylation | GGATGAGTATGAGTACGTGT |
| Human | PHF2 | Histone Methylation | GTCACTGACATAGAACGTGG |
| Human | PHF2 | Histone Methylation | AGTGACGTCGAGAACTACGT |
| Human | PHF20 | Chromatin Readers | AGAACCACTCATCATAACGA |
| Human | PHF20 | Chromatin Readers | AAGCATCCACCTAACAGACG |
| Human | PHF20 | Chromatin Readers | TAGGATTCCTCCACAGAACG |
| Human | PHF20 | Chromatin Readers | ACACTGACAAAGACTTATCG |
| Human | PHF20 | Chromatin Readers | CATAGAAGACATTGACTACG |
| Human | PHF20L1 | Chromatin Readers | TTACCGTAATGAATGTCCCA |
| Human | PHF20L1 | Chromatin Readers | ACTGGTTACACGATCCAGCT |
| Human | PHF20L1 | Chromatin Readers | AAACAGCTCCTAATTGACAT |
| Human | PHF20L1 | Chromatin Readers | TACAACCTATCAGTACCCAA |
| Human | PHF20L1 | Chromatin Readers | GCCATGCCCGAGGATGCTAA |
| Human | PHF21A | Chromatin Readers | GCATTGTGATAGCTACTCCA |
| Human | PHF21A | Chromatin Readers | GCACAGCGAACTGTAACCAG |
| Human | PHF21A | Chromatin Readers | CACAGCGAACTGTAACCAGG |
| Human | PHF21A | Chromatin Readers | TGACCTTACTAGACGTCTGG |
| Human | PHF21A | Chromatin Readers | CAGAGTCTACAGGTTTGGAG |
| Human | PHF21B | Chromatin Readers | GGTGACATAAGAGTGCACGA |
| Human | PHF21B | Chromatin Readers | CAGGCCTAGCGCTACCATGA |
| Human | PHF21B | Chromatin Readers | GGGACACGTACCTTCCAGCA |
| Human | PHF21B | Chromatin Readers | GTTACCACGGAACATTTGGA |
| Human | PHF21B | Chromatin Readers | TCAAGAAGCAGCTCCACGAA |
| Human | PHF23 | Chromatin Readers | ACCGAAAGAACCGAAAGTTG |
| Human | PHF23 | Chromatin Readers | GAAACGGCGGAGAACAATTG |
| Human | PHF23 | Chromatin Readers | GGATGGTTCGAAGATCTGAG |
| Human | PHF23 | Chromatin Readers | GTCGAAAGCCCTTTGCAGGG |
| Human | PHF23 | Chromatin Readers | GATGGTTCGAAGATCTGAGG |
| Human | PHF3 | Chromatin Readers | ACTGTGCACCCATTTCGAAG |
| Human | PHF3 | Chromatin Readers | ACCTTGTTGCTAATACAGCG |
| Human | PHF3 | Chromatin Readers | TTACCTTGTGCCATTAAACG |
| Human | PHF3 | Chromatin Readers | GGAGAAAGATCATCTACAGG |
| Human | PHF3 | Chromatin Readers | CACTGGTCAAGTATATGACC |
| Human | PHF5A | Chromatin Readers | GCCCAAAGATTGTCAATCTG |
| Human | PHF5A | Chromatin Readers | TGTGATCTGTGGAGGACCTG |
| Human | PHF5A | Chromatin Readers | ATATGTGATGAGTGTAACTA |
| Human | PHF5A | Chromatin Readers | GGGCGCTGTGTGATCTGTGG |
| Human | PHF5A | Chromatin Readers | TGTAACTATGGATCTTACCA |
| Human | PHF6 | Chromatin Readers | GTACTTCAGGAGATTAAACG |

| | | | |
|---|---|---|---|
| Human | PHF6 | Chromatin Readers | CTTTATCATGCAATGCACAG |
| Human | PHF6 | Chromatin Readers | GGCAGCGCACCATAAGTGCA |
| Human | PHF6 | Chromatin Readers | TCACATCCAATAGTAGCACC |
| Human | PHF6 | Chromatin Readers | ATACGAGAGAAACCTTCACA |
| Human | PHF7 | Chromatin Readers | CAACACAGAACATCCAACAT |
| Human | PHF7 | Chromatin Readers | TCTAGTAAGCTGCCTCAGAG |
| Human | PHF7 | Chromatin Readers | AGCTTACTAGATAAGATCTG |
| Human | PHF7 | Chromatin Readers | CTCATTCTGTGTGCTACATG |
| Human | PHF7 | Chromatin Readers | CTTACTGTTAGATCTAAGAG |
| Human | PHF8 | Histone Methylation | CGGAGCTCTCTGACGAACGT |
| Human | PHF8 | Histone Methylation | TCAGAGCAAGGCTTCGCACG |
| Human | PHF8 | Histone Methylation | TGTCTCAATAGAGGCTACCC |
| Human | PHF8 | Histone Methylation | CTTTCCCACATACCAACAGA |
| Human | PHF8 | Histone Methylation | CATGCAAGACTTCACAGTTG |
| Human | PHIP | Chromatin Readers | GTTCATTCACCACCAAACGT |
| Human | PHIP | Chromatin Readers | GGTAGCTTGGAGTCGAAGGG |
| Human | PHIP | Chromatin Readers | TGGTGGTGAATGAACCTCTG |
| Human | PHIP | Chromatin Readers | TCATTTGAACTCCAGGCCGA |
| Human | PHIP | Chromatin Readers | AACGTAGGACTAAGACGTAG |
| Human | PHRF1 | Chromatin Readers | TCTGCTTGAGATCTTAGACG |
| Human | PHRF1 | Chromatin Readers | CCACTCGCTCTCGAATCGCG |
| Human | PHRF1 | Chromatin Readers | GATGGTTCGTGTGATCTCAG |
| Human | PHRF1 | Chromatin Readers | AGATCAGCGTCCAGACCTCG |
| Human | PHRF1 | Chromatin Readers | TGTCTAGAAGAAAGCAGCGT |
| Human | PKM | Other Histone Modifications | TCATGGCAAAGTTCACCCGG |
| Human | PKM | Other Histone Modifications | AACCTTACAGATAGCTCGTG |
| Human | PKM | Other Histone Modifications | GTGGTGAATCAATGTCCAGG |
| Human | PKM | Other Histone Modifications | GTTTGCGTCATTCATCCGCA |
| Human | PKM | Other Histone Modifications | TGGGTGGTGAATCAATGTCC |
| Human | PKN1 | Other Histone Modifications | CGACGTCAGCAACTTCGACG |
| Human | PKN1 | Other Histone Modifications | TGAACATCGATGTCGCCACG |
| Human | PKN1 | Other Histone Modifications | CTGCTGCAGTGACATATCGG |
| Human | PKN1 | Other Histone Modifications | CCACTTCCGAGTGGAGCACG |
| Human | PKN1 | Other Histone Modifications | GAACATCGATGTCGCCACGT |
| Human | POLE3 | Chromatin Remodeling | CGTCCTTACAGCATATAGGC |
| Human | POLE3 | Chromatin Remodeling | CGTGATCACCAGGATCATCA |
| Human | POLE3 | Chromatin Remodeling | CCGTCCTTACAGCATATAGG |
| Human | POLE3 | Chromatin Remodeling | GTACAGCACGAAGACGCTGG |
| Human | POLE3 | Chromatin Remodeling | TTCCAGAGCTTCTTTCAATG |
| Human | POLR1A | Transcription Elongation | TGACTACCAGTACGACACCG |
| Human | POLR1A | Transcription Elongation | CATACGTGTTGGCTATGGCG |
| Human | POLR1A | Transcription Elongation | CCTGATACTTACTTGCAGTG |

| | | | |
|---|---|---|---|
| Human | POLR1A | Transcription Elongation | GCTCATAAATATAATCCCAG |
| Human | POLR1A | Transcription Elongation | TTCAGCCGAATACATCCCGA |
| Human | POLR1B | Transcription Elongation | ACAATTTCTATGGCACCGAG |
| Human | POLR1B | Transcription Elongation | GTAAGAAATCACAGCAACGA |
| Human | POLR1B | Transcription Elongation | TGGCATTGGTACATGTTCCG |
| Human | POLR1B | Transcription Elongation | GATGCCACTATATAACCTCT |
| Human | POLR1B | Transcription Elongation | ATAAATTTGCCAGTCGCCAT |
| Human | POLR1C | Transcription Elongation | CAGTCTCTGTTGAGTCAACG |
| Human | POLR1C | Transcription Elongation | GGTTCACGTACAGTTCGTTG |
| Human | POLR1C | Transcription Elongation | AGTCTCTGTTGAGTCAACGG |
| Human | POLR1C | Transcription Elongation | TCGACGAATTCTGCTAGCTG |
| Human | POLR1C | Transcription Elongation | TGCTGAAGGTATCCAGCCGG |
| Human | POLR1E | Transcription Elongation | AGAGACTATCATTGATACGA |
| Human | POLR1E | Transcription Elongation | ATGAGCTCGAAATTTGATGA |
| Human | POLR1E | Transcription Elongation | AAGAGCTTCATACTCCGCAG |
| Human | POLR1E | Transcription Elongation | TGGAGATAGCCAAAGCCATG |
| Human | POLR1E | Transcription Elongation | GAGACTATCATTGATACGAA |
| Human | POLR2A | Transcription Elongation | CAAATACCCAGAGACGACTG |
| Human | POLR2A | Transcription Elongation | CTTCAAGAACTAGTGCGCAG |
| Human | POLR2A | Transcription Elongation | GCCGAAACTCCTCTGCCATG |
| Human | POLR2A | Transcription Elongation | CATGCGGCGGGAACACAACG |
| Human | POLR2A | Transcription Elongation | TAGGTAGGAGATGTGGACCA |
| Human | POLR2B | Transcription Elongation | GTTTGGCAATATGTACGACG |
| Human | POLR2B | Transcription Elongation | GACCAATAGTCATACGAGAG |
| Human | POLR2B | Transcription Elongation | AATATGTACGACGCGGATGA |
| Human | POLR2B | Transcription Elongation | AAGATTCACTCTCGTGCTAG |
| Human | POLR2B | Transcription Elongation | ATAGACCGTGTAGTCACAAG |
| Human | POLR2C | Transcription Elongation | GGTGGCCAATTCGATTCGGA |
| Human | POLR2C | Transcription Elongation | ACAATGTCATCACTAATGAG |
| Human | POLR2C | Transcription Elongation | TTCATTGCACCGCACATCGA |
| Human | POLR2C | Transcription Elongation | ACGCCAACCAGCCTACCGTG |
| Human | POLR2C | Transcription Elongation | ACGGTAGGCTGGTTGGCGTA |
| Human | POLR2D | Transcription Elongation | AGTCCAAGGCTCTAATCCCA |
| Human | POLR2D | Transcription Elongation | TAGCACTCACCTACGAACAC |
| Human | POLR2D | Transcription Elongation | AGACCATTGCCAGTGTTCGT |
| Human | POLR2D | Transcription Elongation | TCACCTACGAACACTGGCAA |
| Human | POLR2D | Transcription Elongation | TAACCTTGGGATTAGAGCCT |
| Human | POLR2E | Transcription Elongation | GGAGTTCAAAGCCCAATCTG |
| Human | POLR2E | Transcription Elongation | TGAGCACGTCGTCATGACCA |
| Human | POLR2E | Transcription Elongation | TCCCAAAGTAGCGCGCCACA |
| Human | POLR2E | Transcription Elongation | CCTGGGTCACCAGATAGCCA |
| Human | POLR2E | Transcription Elongation | CATCACACGGGCTCTCATCG |

| | | | |
|---|---|---|---|
| Human | POLR2F | Transcription Elongation | GGGTGTCATGTCAGACAACG |
| Human | POLR2F | Transcription Elongation | GTACTTGGTCATGTATGGTG |
| Human | POLR2F | Transcription Elongation | GAAGATGACTCCAGCTCAGT |
| Human | POLR2F | Transcription Elongation | GGTAACGGCGAATGATGATG |
| Human | POLR2F | Transcription Elongation | TGTGGTGATTCGCTTCTGGT |
| Human | POLR2G | Transcription Elongation | ACCCAAGTAATCGTCCATCA |
| Human | POLR2G | Transcription Elongation | AGCACCAATATTGTCAATGG |
| Human | POLR2G | Transcription Elongation | AATGTCATTCTTGTCCACAC |
| Human | POLR2G | Transcription Elongation | AGATCCGCTTAAAGATTGTG |
| Human | POLR2G | Transcription Elongation | ACCAGCACCAATATTGTCAA |
| Human | POLR2H | Transcription Elongation | CAACCTGCATGGATTCGAGG |
| Human | POLR2H | Transcription Elongation | TCTGCGTACGTGTCCTATGG |
| Human | POLR2H | Transcription Elongation | AGTCCACCTCGAATCCATGC |
| Human | POLR2H | Transcription Elongation | CAACAACCTGCATGGATTCG |
| Human | POLR2H | Transcription Elongation | CTGGAAGGCCTATCATCAGT |
| Human | POLR2I | Transcription Elongation | CAACAAGATCACGCACGAAG |
| Human | POLR2I | Transcription Elongation | GAACCGCATTCTGCTCTACG |
| Human | POLR2I | Transcription Elongation | GAGGTCACTCGGTCCAGCGG |
| Human | POLR2I | Transcription Elongation | GAGGAGGTCACTCGGTCCAG |
| Human | POLR2I | Transcription Elongation | CCTCGGTCCGCGGCAACGTG |
| Human | POLR2J | Transcription Elongation | AGCAGGAAGGAATTGAGTAG |
| Human | POLR2J | Transcription Elongation | CCGAAAGCGCTCCTCCAGCA |
| Human | POLR2J | Transcription Elongation | CATCCGAGTGCAGACCACGC |
| Human | POLR2J | Transcription Elongation | GAAGAGCAAGAACGACTCGA |
| Human | POLR2J | Transcription Elongation | CAAGCAGGAAGGAATTGAGT |
| Human | POLR2K | Transcription Elongation | CCAATCAGATGCAGAGAATG |
| Human | POLR2K | Transcription Elongation | CAACCAATGATATATATCTG |
| Human | POLR2K | Transcription Elongation | GCTAACAATGGACACCCAGA |
| Human | POLR2K | Transcription Elongation | CCAATGATATATATCTGTGG |
| Human | POLR2K | Transcription Elongation | ACAGAATAATGTACAAGAAA |
| Human | POLR2L | Transcription Elongation | TGCAGGCCGAGTACACCGAG |
| Human | POLR2L | Transcription Elongation | CAAGATCGTCGGCAACAAGT |
| Human | POLR2L | Transcription Elongation | GCAAGATCGTCGGCAACAAG |
| Human | POLR2L | Transcription Elongation | GATCGTCGGCAACAAGTGGG |
| Human | POLR2L | Transcription Elongation | CCTGTACGCTGCTTCACTTG |
| Human | POP5 | Control | CGTACGGGACACGATCGCCA |
| Human | POP5 | Control | TGGCGATCGTGTCCCGTACG |
| Human | POP5 | Control | TTCAACACATTACATGTGGG |
| Human | POP5 | Control | AGTTCTGAGCAGCCTCGTAC |
| Human | POP5 | Control | TGCTCAGAACTCGGTCATCG |
| Human | POU4F1 | Transcription Elongation | GCTGTTCATCGTGTGGTACG |
| Human | POU4F1 | Transcription Elongation | GCCCGAGAAGCGCTCCCTCG |

| | | | |
|---|---|---|---|
| Human | POU4F1 | Transcription Elongation | ACGGCACGCTGTTCATCGTG |
| Human | POU4F1 | Transcription Elongation | AATTCTCTGCCACTTACTGA |
| Human | POU4F1 | Transcription Elongation | GTGGACATCGCCGTGTCCCA |
| Human | PPP2R1A | Control | TGCAGTATATGCCATCCGCG |
| Human | PPP2R1A | Control | TCACAGCACTGGACACTCGG |
| Human | PPP2R1A | Control | CGTCTATGAGCACCGCGATG |
| Human | PPP2R1A | Control | TGCAGAAGAGCGTAGTCATG |
| Human | PPP2R1A | Control | CATAGACGAACTCCGCAATG |
| Human | PRC1 | Control | CGAACACCTAGCAAGCGGCG |
| Human | PRC1 | Control | AATCGATTTACAAACCGAGG |
| Human | PRC1 | Control | GCGGTTACAAAGAACTGAGG |
| Human | PRC1 | Control | GCACTGTCAATATCATAGTG |
| Human | PRC1 | Control | TCCGCCATGAGGAGAAGGTG |
| Human | PRDM1 | Histone Methylation | GGACGCGTTCAAGTAAGCGT |
| Human | PRDM1 | Histone Methylation | CATTACAATTCATGCCGTAG |
| Human | PRDM1 | Histone Methylation | TTTGGACAGATCTATTCCAG |
| Human | PRDM1 | Histone Methylation | GTGTAGATTTCACCTATTAG |
| Human | PRDM1 | Histone Methylation | CCTGAAAGGGAACTGCGCTG |
| Human | PRDM10 | Histone Methylation | GCTCCGAGCTTATAAATCGG |
| Human | PRDM10 | Histone Methylation | CAGACTACCGAACGCCACAA |
| Human | PRDM10 | Histone Methylation | CTGGTTATGAACTGGCAATG |
| Human | PRDM10 | Histone Methylation | AGACTACCGAACGCCACAAG |
| Human | PRDM10 | Histone Methylation | TGTGGAGTCGCAGTTTATCG |
| Human | PRDM11 | Histone Methylation | TGACGTGCGATGTGTAAACG |
| Human | PRDM11 | Histone Methylation | GATAACCCAGAAGACCTGAG |
| Human | PRDM11 | Histone Methylation | GCTGCGGGTCTGGTACAGCG |
| Human | PRDM11 | Histone Methylation | TGTGTCAGACACAAACACCG |
| Human | PRDM11 | Histone Methylation | GTGGATGAATGCCCAAACCA |
| Human | PRDM12 | Histone Methylation | GATGTCGGAGGTGATAACCT |
| Human | PRDM12 | Histone Methylation | CCGCGCGTACCTCCCACATG |
| Human | PRDM12 | Histone Methylation | ACTCACACAACACCTTCCTG |
| Human | PRDM12 | Histone Methylation | GGCAGATGACGCATCGCATG |
| Human | PRDM12 | Histone Methylation | GTTACGTGCACACTTGATGT |
| Human | PRDM13 | Histone Methylation | GGCCTTTACAGTCTACAACG |
| Human | PRDM13 | Histone Methylation | CATTGCGAGACGTCCAGCCA |
| Human | PRDM13 | Histone Methylation | CCGCCGAAGTGCAATTTGAG |
| Human | PRDM13 | Histone Methylation | GGTGTGCCTTAAGGCTGTTG |
| Human | PRDM13 | Histone Methylation | CCGTTGTAGACTGTAAAGGC |
| Human | PRDM14 | Histone Methylation | AAGTGTCAGGTCCACAACAC |
| Human | PRDM14 | Histone Methylation | TGCTCGGTTCCAGTTCACGG |
| Human | PRDM14 | Histone Methylation | CATCACAGAATTGTCTCCGT |
| Human | PRDM14 | Histone Methylation | TGTCCGGCGTTCACACAAGG |

| | | | |
|---|---|---|---|
| Human | PRDM14 | Histone Methylation | ATTGGTGGCGACAACGAGAG |
| Human | PRDM15 | Histone Methylation | AGTGGCAGAGATCATTACCG |
| Human | PRDM15 | Histone Methylation | GGACTCAGCTTCTCTAAACG |
| Human | PRDM15 | Histone Methylation | ACTCAGCTTCTCTAAACGCG |
| Human | PRDM15 | Histone Methylation | GTGACCTTTGATACCGTCAG |
| Human | PRDM15 | Histone Methylation | GAGATTGGTAAACTGAGTCG |
| Human | PRDM16 | Histone Methylation | CTGGCTCAAGTACATCCGTG |
| Human | PRDM16 | Histone Methylation | CCTTGCACTTGATCTGCGTG |
| Human | PRDM16 | Histone Methylation | CACCAGATGTCCCACGACAG |
| Human | PRDM16 | Histone Methylation | AAGAACCATTACACGCCGGG |
| Human | PRDM16 | Histone Methylation | AAATGCTCCAGACTCCGACG |
| Human | PRDM2 | Histone Methylation | GTGGCAACAGTTAATACCGG |
| Human | PRDM2 | Histone Methylation | AGTGGCAACAGTTAATACCG |
| Human | PRDM2 | Histone Methylation | AGGTGTGGACTCAACAGTAG |
| Human | PRDM2 | Histone Methylation | CCTCAAGTAGGTTTAAGAGG |
| Human | PRDM2 | Histone Methylation | ACACTTTGTGCAGCGAATGG |
| Human | PRDM4 | Histone Methylation | AAGATCTAGCAGACTCTGTG |
| Human | PRDM4 | Histone Methylation | CATTGACCCGACTTGCAATG |
| Human | PRDM4 | Histone Methylation | GAGGGAAACAGATTCTAGGG |
| Human | PRDM4 | Histone Methylation | GTCCACTTTATGGGTCACAT |
| Human | PRDM4 | Histone Methylation | CCATGTTCGGGACAGTCCGA |
| Human | PRDM5 | Histone Methylation | TGAAGATATAGAAACAGACA |
| Human | PRDM5 | Histone Methylation | ATTAGCTGTCAGCTACACCA |
| Human | PRDM5 | Histone Methylation | GCACATTCGTACTCACACCA |
| Human | PRDM5 | Histone Methylation | GGCCCGCAGAGTGCGAAAGG |
| Human | PRDM5 | Histone Methylation | GAGGCCTGGATGAGCACAAG |
| Human | PRDM6 | Histone Methylation | TGAGGTATATCCGATGTGCA |
| Human | PRDM6 | Histone Methylation | GCCTGTATAGATATCCCTAG |
| Human | PRDM6 | Histone Methylation | TAGGGATATCTATACAGGCT |
| Human | PRDM6 | Histone Methylation | GCTTTAAGACTTTCACCCAG |
| Human | PRDM6 | Histone Methylation | CCTGGGCAATGCATTGTAAG |
| Human | PRDM7 | Histone Methylation | TGGGCTTGGAGTATGGAACG |
| Human | PRDM7 | Histone Methylation | CCGATGAAGAATGGACACCT |
| Human | PRDM7 | Histone Methylation | TATAATGCACTGATTACTGT |
| Human | PRDM7 | Histone Methylation | TCTCCAACCTAGAACTCAGG |
| Human | PRDM7 | Histone Methylation | GACGAAGAGGCAGCCAACAG |
| Human | PRDM8 | Histone Methylation | CTTTCCGAGTCAATTATGCG |
| Human | PRDM8 | Histone Methylation | GGAACTCAAACTGGAAACGT |
| Human | PRDM8 | Histone Methylation | CATATGGTACACCAGGTCGG |
| Human | PRDM8 | Histone Methylation | CACCGCTAGAGGGTCCGCGG |
| Human | PRDM8 | Histone Methylation | GGAGGCGACTCCCGCGCCGT |
| Human | PRDM9 | Histone Methylation | AAAGGGTCATGCATACAAAG |

| | | | |
|---|---|---|---|
| Human | PRDM9 | Histone Methylation | CAGAGGTAATCATCATCCTG |
| Human | PRDM9 | Histone Methylation | GCAAATTTACTGAATGCAAG |
| Human | PRDM9 | Histone Methylation | CAAGCCTCTGCTATCACCAG |
| Human | PRDM9 | Histone Methylation | GAGTGAGGAGGACTGACTGC |
| Human | PRKAA1 | Other Histone Modifications | GCAACTATCGATCTTGCCAA |
| Human | PRKAA1 | Other Histone Modifications | CATATTATTTGCGTGTACGA |
| Human | PRKAA1 | Other Histone Modifications | TCTTTACAACAGAAATCACC |
| Human | PRKAA1 | Other Histone Modifications | GAAGATTCGGAGCCTTGATG |
| Human | PRKAA1 | Other Histone Modifications | GAGTCTACAGTTATACCAAG |
| Human | PRKCB | Other Histone Modifications | CTGTAAGAAGAACAGACCGA |
| Human | PRKCB | Other Histone Modifications | TGACGTGGAGTGCACTATGG |
| Human | PRKCB | Other Histone Modifications | CATCCAGGCCCACATCGACA |
| Human | PRKCB | Other Histone Modifications | CAGAGGGCCAAGATCAGTCA |
| Human | PRKCB | Other Histone Modifications | ATTGACCAATCAGAATTCGA |
| Human | PRKCD | Other Histone Modifications | AGGAACCTCAATCTTCCAGG |
| Human | PRKCD | Other Histone Modifications | TTCCCAACGATGAACCGCCG |
| Human | PRKCD | Other Histone Modifications | TGGCCCTTACGTACGTGGCA |
| Human | PRKCD | Other Histone Modifications | AGCCGACCATGTATCCTGAG |
| Human | PRKCD | Other Histone Modifications | TCTCTCGGGCAGACAACAGT |
| Human | PRMT1 | Histone Methylation | AAAGCCAACAAGTTAGACCA |
| Human | PRMT1 | Histone Methylation | GGACATCTATACCGTCAAGG |
| Human | PRMT1 | Histone Methylation | GATGGCCGTCACATACAGCG |
| Human | PRMT1 | Histone Methylation | GGCCGCGAACTGCATCATGG |
| Human | PRMT1 | Histone Methylation | ACCGACTACCGGATGCGCTG |
| Human | PRMT2 | Histone Methylation | TGATAAGGATTATCGTAGCA |
| Human | PRMT2 | Histone Methylation | GCTGCGCTTCGACATCAGGA |
| Human | PRMT2 | Histone Methylation | TCTTCTGTGCACACTATGCG |
| Human | PRMT2 | Histone Methylation | ACATTCCGGCAAACCATGTG |
| Human | PRMT2 | Histone Methylation | CCACGTCTCCTGTATGGACA |
| Human | PRMT3 | Histone Methylation | ACACCTACCTTAATACCACA |
| Human | PRMT3 | Histone Methylation | GAATTCATGTACTCAACTGT |
| Human | PRMT3 | Histone Methylation | AATCTGAGATAGACGTCGTA |
| Human | PRMT3 | Histone Methylation | TGAGAAGAATTGCCACAACA |
| Human | PRMT3 | Histone Methylation | TTATGCAAAGAACAAATACT |
| Human | PRMT5 | Histone Methylation | GGTACTGAGAGTATTTGATG |
| Human | PRMT5 | Histone Methylation | TGCACCAACTACACACACAG |
| Human | PRMT5 | Histone Methylation | CTAGTGTGTCAGCTATTTCG |
| Human | PRMT5 | Histone Methylation | CAATGACATGATTAGATGGG |
| Human | PRMT5 | Histone Methylation | ATGAACTCCCTCTTGAAACG |
| Human | PRMT6 | Histone Methylation | GTGGCCCATGAGACAGCGCG |
| Human | PRMT6 | Histone Methylation | TGGATGCCATCGTGAGCGAG |
| Human | PRMT6 | Histone Methylation | GGCAAGACGGTACTGGACGT |

| | | | |
|---|---|---|---|
| Human | PRMT6 | Histone Methylation | CTCGGACGTTTCGGTCCACG |
| Human | PRMT6 | Histone Methylation | TCCTCCACGCGCGAACCAAG |
| Human | PRMT7 | Histone Methylation | ACCATACGCAGTAGTCATCG |
| Human | PRMT7 | Histone Methylation | TCATTGATCTCTCCAAACCG |
| Human | PRMT7 | Histone Methylation | CAGAAGTAGAGGTTGTGCCA |
| Human | PRMT7 | Histone Methylation | TATCAACAAGCATTCCACCG |
| Human | PRMT7 | Histone Methylation | CTGCATAGCTGACAGTCCGT |
| Human | PRMT8 | Histone Methylation | AGACCGGGCAGCTTTGTACG |
| Human | PRMT8 | Histone Methylation | CCAAGTCTACTGTGAAATCG |
| Human | PRMT8 | Histone Methylation | GTCGAAGTAATAATCTCTCG |
| Human | PRMT8 | Histone Methylation | ACATGATGGACGCATTGCAC |
| Human | PRMT8 | Histone Methylation | AGATTACCTCACTGTCCGGA |
| Human | PSIP1 | Chromatin Readers | GGATGTGAACAGATGCATTG |
| Human | PSIP1 | Chromatin Readers | GGGTCAAAGACTCTAAATGG |
| Human | PSIP1 | Chromatin Readers | CTTACTCGAGCTGGCCAATG |
| Human | PSIP1 | Chromatin Readers | ACTCACATGTCACTCTCTGA |
| Human | PSIP1 | Chromatin Readers | CAGCCTAAGAAGGATGAAGA |
| Human | PSMA4 | Control | GACCCTAGTGGAAATTACGG |
| Human | PSMA4 | Control | CAAAGTACTAAATAAGACCA |
| Human | PSMA4 | Control | GCTGTAACCAACTGCTCACA |
| Human | PSMA4 | Control | TGACCCTAGTGGAAATTACG |
| Human | PSMA4 | Control | ATTGGCTGGGATAAGCACTA |
| Human | PSMC2 | Control | CGATCAGCGGAAGACCAAAG |
| Human | PSMC2 | Control | TCTTCCGCTGATCGGCACCG |
| Human | PSMC2 | Control | AAGGACGACAAGCCCATCCG |
| Human | PSMC2 | Control | ATTACCTCGGTGCCGATCAG |
| Human | PSMC2 | Control | TGTACAGAAATACGTCGGTG |
| Human | PWWP2B | Chromatin Readers | GGACCATCAAGCGCACGCGG |
| Human | PWWP2B | Chromatin Readers | ATAGACAAGAACGACGTAGT |
| Human | PWWP2B | Chromatin Readers | TCGAAGTAGGGAGGGTACGG |
| Human | PWWP2B | Chromatin Readers | GTATCGGAAAGCTATAACCG |
| Human | PWWP2B | Chromatin Readers | CTGATGTCAAGAACACGCGC |
| Human | PYGO1 | Chromatin Readers | AAAGATGAACGTACCATGCG |
| Human | PYGO1 | Chromatin Readers | TTGTGGAATTTGTACAAACG |
| Human | PYGO1 | Chromatin Readers | AAATTCCACAAGGATACACT |
| Human | PYGO1 | Chromatin Readers | GGAGCATACTCAGACAATGG |
| Human | PYGO1 | Chromatin Readers | GTGTCATCCATATTCAAGTG |
| Human | PYGO2 | Chromatin Readers | CATGGGTACACCAAGCCTGG |
| Human | PYGO2 | Chromatin Readers | CCAGTCTGTCTACATCCGTG |
| Human | PYGO2 | Chromatin Readers | CCAGGCTTGGTGTACCCATG |
| Human | PYGO2 | Chromatin Readers | TCATGCCTGTGCACTCACGG |
| Human | PYGO2 | Chromatin Readers | GCAACCAGGTGATCCACCAT |

| | | | |
|---|---|---|---|
| Human | RAD51 | Chromatin Remodeling | CTATAGCTTCCCATTGACCG |
| Human | RAD51 | Chromatin Remodeling | AGTGGTAATCACTAATCAGG |
| Human | RAD51 | Chromatin Remodeling | ATACCTAGATTCTACCATCA |
| Human | RAD51 | Chromatin Remodeling | CAACTGAATTCCACCAAAGG |
| Human | RAD51 | Chromatin Remodeling | TGTTTGGAGAATTCCGAACT |
| Human | RAD54B | Chromatin Remodeling | TATGTTCAGATGTCTTGGTG |
| Human | RAD54B | Chromatin Remodeling | TATCTGGTCGTGGCATAACG |
| Human | RAD54B | Chromatin Remodeling | GCAGCACCAAGTCAGTTGCA |
| Human | RAD54B | Chromatin Remodeling | GCTCCTATACATATTAGATG |
| Human | RAD54B | Chromatin Remodeling | GACAAACACCAATCTCTCAA |
| Human | RAD54L | Chromatin Remodeling | GACGGCTGTTGACACAACGT |
| Human | RAD54L | Chromatin Remodeling | TGGACATCGTGCCATCCAGG |
| Human | RAD54L | Chromatin Remodeling | GAAGGATTCATGAACCAGCG |
| Human | RAD54L | Chromatin Remodeling | AACCACTGCACTGATAAGTG |
| Human | RAD54L | Chromatin Remodeling | ATTACAAAGCTTCTTTAGCG |
| Human | RAG1 | Chromatin Remodeling | CAGTGGACACCATTGCAAAG |
| Human | RAG1 | Chromatin Remodeling | CATGTGAGGTTTACTTCCCG |
| Human | RAG1 | Chromatin Remodeling | TACCTTCTCAGCATTCCGAA |
| Human | RAG1 | Chromatin Remodeling | ATTTGGAGGTGTACAACCAG |
| Human | RAG1 | Chromatin Remodeling | AGATGCGGCAGAGATGTCGA |
| Human | RAG2 | Chromatin Readers | TTAATGTGGTGTACAGCCGA |
| Human | RAG2 | Chromatin Readers | CATCAATACATCATCCATGG |
| Human | RAG2 | Chromatin Readers | GTGATGTGGATATCAACACT |
| Human | RAG2 | Chromatin Readers | ATTCACAGCTGGGCTACCCA |
| Human | RAG2 | Chromatin Readers | TTGATATCCACATCACAAGT |
| Human | RAI1 | Chromatin Readers | GCCGCCGACAAGTACCACCG |
| Human | RAI1 | Chromatin Readers | GATGTGTACTCGTTCTCTCA |
| Human | RAI1 | Chromatin Readers | CGGAGCCGTCGGGTGCACCG |
| Human | RAI1 | Chromatin Readers | CCAGCTCTGGACCTTTGACT |
| Human | RAI1 | Chromatin Readers | GTTGACAACAGTGCATATGG |
| Human | RECQL5 | Transcription Elongation | ACAGCTCACCGAGTACACAT |
| Human | RECQL5 | Transcription Elongation | CCTGTCCCTAGGTATGACGA |
| Human | RECQL5 | Transcription Elongation | TTTACAGGAGAGTGCGACCA |
| Human | RECQL5 | Transcription Elongation | GAGTCAGCAAGTGTGAGAGG |
| Human | RECQL5 | Transcription Elongation | TCGTGGAACTCACTCATCAG |
| Human | RNF168 | Transcription Elongation | AAGAAATTCTCTCGTCAACG |
| Human | RNF168 | Transcription Elongation | GCGTAAGTGATACTCATCTG |
| Human | RNF168 | Transcription Elongation | GCACCACAGGCACATAACCA |
| Human | RNF168 | Transcription Elongation | GATACCCGGCGGCGACAGAA |
| Human | RNF168 | Transcription Elongation | TCGTCTGCTCAGTAAACCTG |
| Human | RNF17 | Chromatin Readers | GCCACGATACCACAGAGTAT |
| Human | RNF17 | Chromatin Readers | AGAGTGTTCTGGTCTCACAT |

| | | | |
|---|---|---|---|
| Human | RNF17 | Chromatin Readers | GGTCTTCCTACCAGCGAATG |
| Human | RNF17 | Chromatin Readers | AGATGATAGTGGAGTCAGCG |
| Human | RNF17 | Chromatin Readers | TAGACAACGCTACTACCCAA |
| Human | RNF2 | Other Histone Modifications | CTTGGATAAGTGATCAACAG |
| Human | RNF2 | Other Histone Modifications | CCCTTGACTAGACTGCAGCG |
| Human | RNF2 | Other Histone Modifications | GCTTCATACTCATCACGACT |
| Human | RNF2 | Other Histone Modifications | AAGATGACAGTGCACAGACG |
| Human | RNF2 | Other Histone Modifications | GGTGCGTAATAAAGTTCCAT |
| Human | RNF20 | Other Histone Modifications | GCATCGCACCATGTCTCAGG |
| Human | RNF20 | Other Histone Modifications | GTCTAGTACTGAGGACCCGA |
| Human | RNF20 | Other Histone Modifications | GGAGGGCACTACCACTACGC |
| Human | RNF20 | Other Histone Modifications | TATTTAAGGCCCTATAAACA |
| Human | RNF20 | Other Histone Modifications | TCGGTTGACAATCAATAGTG |
| Human | RNF40 | Other Histone Modifications | AATGGGCTCGATAAGAGTCG |
| Human | RNF40 | Other Histone Modifications | GCAGAAGAAGCTACGCACAG |
| Human | RNF40 | Other Histone Modifications | GGGCTCGATAAGAGTCGTGG |
| Human | RNF40 | Other Histone Modifications | GTAGATACGATGGAAGTCGT |
| Human | RNF40 | Other Histone Modifications | GCTCCAGGATAAAGTGACAT |
| Human | RNF8 | Transcription Elongation | TGAGCCAAGTAAGACCACAG |
| Human | RNF8 | Transcription Elongation | TAATGTCCTTCCGACAAATG |
| Human | RNF8 | Transcription Elongation | CAAAGAATTAGAGCAGACCA |
| Human | RNF8 | Transcription Elongation | AGAGCTAAATCGCAGCAAGA |
| Human | RNF8 | Transcription Elongation | TTGAGCCAAGTAAGACCACA |
| Human | RPH3A | Chromatin Readers | ACAGTCCTCACATACTACAC |
| Human | RPH3A | Chromatin Readers | CAGGAGGTGCAGCTTAACGT |
| Human | RPH3A | Chromatin Readers | TAGCCTAATCACTTGACACG |
| Human | RPH3A | Chromatin Readers | CGGTGATGCCGTGATACACG |
| Human | RPH3A | Chromatin Readers | GTTCTCCACACAGTATGCAG |
| Human | RPS6KA3 | Other Histone Modifications | CCAGAAGTAGTTAATCGTCG |
| Human | RPS6KA3 | Other Histone Modifications | ACATCCTTGGATCGTCCACT |
| Human | RPS6KA3 | Other Histone Modifications | TGAATATCTTCACGCACAAG |
| Human | RPS6KA3 | Other Histone Modifications | AGCTGATGTGCATTAGCACT |
| Human | RPS6KA3 | Other Histone Modifications | ACAATACCAACTAAACAGAC |
| Human | RPS6KA4 | Other Histone Modifications | GTACGCCGACGACACGCCCG |
| Human | RPS6KA4 | Other Histone Modifications | TCATCGACTTCGGGTTCGCG |
| Human | RPS6KA4 | Other Histone Modifications | AAATCATCCGTAGCAAGACG |
| Human | RPS6KA4 | Other Histone Modifications | TGAAGCTCACGGCCGACACG |
| Human | RPS6KA4 | Other Histone Modifications | CGCCACGGGCCCGATCCGAG |
| Human | RPS6KA5 | Other Histone Modifications | GGCACCAGATATTGTCAGAG |
| Human | RPS6KA5 | Other Histone Modifications | CCACTCCAACATCATGCATG |
| Human | RPS6KA5 | Other Histone Modifications | ATGGCACCAGATATTGTCAG |
| Human | RPS6KA5 | Other Histone Modifications | CGGACTCAGTAGCTTAGGCA |

| | | | |
|---|---|---|---|
| Human | RPS6KA5 | Other Histone Modifications | AAGTTTGACGTGTACCAGCG |
| Human | RSF1 | Chromatin Readers | AAGCGGTCAGTTCGAAAGCG |
| Human | RSF1 | Chromatin Readers | TCCCAGAAGAGTTTGCGACG |
| Human | RSF1 | Chromatin Readers | GACATCAAAGAAGCCGATGG |
| Human | RSF1 | Chromatin Readers | TTGTAGAAACTCGGCGAAGG |
| Human | RSF1 | Chromatin Readers | TGTTACAGACATAATCAACA |
| Human | RTF1 | Transcription Elongation | TGTCCGAGTCGATCACGACG |
| Human | RTF1 | Transcription Elongation | AGATGGATCCCTTTACTCGG |
| Human | RTF1 | Transcription Elongation | AGGCCGCGTCGTGATCGACT |
| Human | RTF1 | Transcription Elongation | CACGGCATAAGCTAGAACGC |
| Human | RTF1 | Transcription Elongation | CTCACATTAGAAACGATGGT |
| Human | SART3 | Other Histone Modifications | ACTTACGCTCCGTATGTCTG |
| Human | SART3 | Other Histone Modifications | GACAAGGCAGTTCTCGACCA |
| Human | SART3 | Other Histone Modifications | TTTCCGAGGTTACTGCTACG |
| Human | SART3 | Other Histone Modifications | TGGGATAGCATCATGACCAG |
| Human | SART3 | Other Histone Modifications | CTGCGACATACAAGACCATG |
| Human | SATB1 | Chromatin Remodeling | TATTCATAGATCTACTGACA |
| Human | SATB1 | Chromatin Remodeling | TAGGTGTTGATACGAGCCCA |
| Human | SATB1 | Chromatin Remodeling | TTCATAGATCTACTGACAGG |
| Human | SATB1 | Chromatin Remodeling | GCTGGAACATGGATAATGTG |
| Human | SATB1 | Chromatin Remodeling | TTCCGGTTTAGAGGTCGATG |
| Human | SCMH1 | Chromatin Readers | CTGTCTTGGAACATCAACCA |
| Human | SCMH1 | Chromatin Readers | TACCTGGTAGGTACCTGTCG |
| Human | SCMH1 | Chromatin Readers | CAAACAGATTGTCACTACGA |
| Human | SCMH1 | Chromatin Readers | CTGAGGGATAGCCTACCTTG |
| Human | SCMH1 | Chromatin Readers | CTTCTCAGTATTCACATCGG |
| Human | SCML2 | Chromatin Readers | AAATCGCTGACCAGAGACCG |
| Human | SCML2 | Chromatin Readers | AGGTTACGGTTACGACTGGA |
| Human | SCML2 | Chromatin Readers | ACTTTGAAATCATTCACAGG |
| Human | SCML2 | Chromatin Readers | ATACTGAAGTGGCATTGCGA |
| Human | SCML2 | Chromatin Readers | TCATCACATCACTCTTGAGT |
| Human | SETD1A | Histone Methylation | GAGATCTCCGACGACGACCG |
| Human | SETD1A | Histone Methylation | ACCTGCGGCTTACGTACGAG |
| Human | SETD1A | Histone Methylation | CGTATGCTCTATATGCACAG |
| Human | SETD1A | Histone Methylation | TTGGACCTGACATGGCAACG |
| Human | SETD1A | Histone Methylation | ATCAGAACTACGAAACTGAG |
| Human | SETD1B | Histone Methylation | CAGATCGATGAGTTCTACGT |
| Human | SETD1B | Histone Methylation | TTATGTACACAATTCTCCCG |
| Human | SETD1B | Histone Methylation | AAGAGCCACATTCACGACTG |
| Human | SETD1B | Histone Methylation | GCCACCTAAGAAGCGCCATG |
| Human | SETD1B | Histone Methylation | GCAGCACATTAACGTCAATG |
| Human | SETD2 | Histone Methylation | ATGAACTGGGATTCCGACGA |

| | | | |
|---|---|---|---|
| Human | SETD2 | Histone Methylation | TTAAAGAACCAGTTGATACG |
| Human | SETD2 | Histone Methylation | TGACCTATTAGGAGTCTTCG |
| Human | SETD2 | Histone Methylation | TCTTCCACCAGGAATAAAGG |
| Human | SETD2 | Histone Methylation | ACTGTCAGGTTGCTGATACG |
| Human | SETD3 | Histone Methylation | CCTGGTCTTGAGAAACCTCG |
| Human | SETD3 | Histone Methylation | GATCACTACTGGTTACAACC |
| Human | SETD3 | Histone Methylation | AATATGAAGAGAGTAACCTT |
| Human | SETD3 | Histone Methylation | GATATGTGTAACCACACCAA |
| Human | SETD3 | Histone Methylation | GGAATACTATCGCCAACAGA |
| Human | SETD4 | Histone Methylation | AAATTAGAACGACTTCACGT |
| Human | SETD4 | Histone Methylation | ACAGCCGTTGATTATCGTGA |
| Human | SETD4 | Histone Methylation | AACAGCCGTTGATTATCGTG |
| Human | SETD4 | Histone Methylation | AGTGATTCGAAGCTACTTAG |
| Human | SETD4 | Histone Methylation | ACGGCCCTCACGATAATCAA |
| Human | SETD5 | Histone Methylation | GTGACAAGGATCCACGAGAG |
| Human | SETD5 | Histone Methylation | GTGTTGGTGTGAATACCCGG |
| Human | SETD5 | Histone Methylation | GTGCCCTTTACGTATCACAA |
| Human | SETD5 | Histone Methylation | AGGTTATTAGACTTCATCGG |
| Human | SETD5 | Histone Methylation | TCAGTAGAACGACTCCGAGA |
| Human | SETD6 | Histone Methylation | CTGACCTATAGGCCATCACA |
| Human | SETD6 | Histone Methylation | TTCCCAAGCTCAAAGCATCG |
| Human | SETD6 | Histone Methylation | TCAGATGGTGACAGTTCGTG |
| Human | SETD6 | Histone Methylation | TCAGCCGTTTACCTTTAGTG |
| Human | SETD6 | Histone Methylation | CGCCAATCTAGAATACTCTG |
| Human | SETD7 | Histone Methylation | AGGCACAGTACTTGGATACG |
| Human | SETD7 | Histone Methylation | AGTTCGAATTACACACCAAG |
| Human | SETD7 | Histone Methylation | GGATAGCGACGACGAGATGG |
| Human | SETD7 | Histone Methylation | GACGATGACGGATTACCGCA |
| Human | SETD7 | Histone Methylation | ATGAAGTCGACTTATCAAAG |
| Human | SETDB1 | Histone Methylation | AGATGTGAGTGGATCTATCG |
| Human | SETDB1 | Histone Methylation | AAGGAAAGAGTCTACTGTCG |
| Human | SETDB1 | Histone Methylation | AGGTCCAATGAAGCGTCAAG |
| Human | SETDB1 | Histone Methylation | CGTTACTATATGACTTCCGG |
| Human | SETDB1 | Histone Methylation | GGAACTGGGTATCTCTATGG |
| Human | SETDB2 | Histone Methylation | ATAGAGAAGAAACTCCACCA |
| Human | SETDB2 | Histone Methylation | GGAAATCTTCTGCCAATGTG |
| Human | SETDB2 | Histone Methylation | AGCAAGAACAGAGCTAACAT |
| Human | SETDB2 | Histone Methylation | ATGTAACCAGGCGACCACAT |
| Human | SETDB2 | Histone Methylation | TGGAAGGAGTCTACGAAACG |
| Human | SETMAR | Histone Methylation | GATCGAGAAATGGTTCGTTG |
| Human | SETMAR | Histone Methylation | CTGCAAGCGATTGACCCTTG |
| Human | SETMAR | Histone Methylation | CAATTACATTATAGCCATCA |

| | | | |
|---|---|---|---|
| Human | SETMAR | Histone Methylation | GAGCAGCTGGATGTCGCGTG |
| Human | SETMAR | Histone Methylation | AGTGGTCACTGCATCGGCAC |
| Human | SFMBT1 | Chromatin Readers | ACTGTCGGCGTGGCACACAA |
| Human | SFMBT1 | Chromatin Readers | TGCGTCCTGAGAACCACGCA |
| Human | SFMBT1 | Chromatin Readers | CTGCATTGATAAGTAAAGTG |
| Human | SFMBT1 | Chromatin Readers | TCCTACAGCTCCGTAATGGG |
| Human | SFMBT1 | Chromatin Readers | TTTCTACAGTATATAAACAG |
| Human | SFMBT2 | Chromatin Readers | CTTAATGAACCTCACCACGT |
| Human | SFMBT2 | Chromatin Readers | CCAGACGGACACCTCGTCGG |
| Human | SFMBT2 | Chromatin Readers | ACACTGAATCCTATGACCAG |
| Human | SFMBT2 | Chromatin Readers | AGTGGCTAATAAGAACAACC |
| Human | SFMBT2 | Chromatin Readers | GACTACGTCACACCAGAAGT |
| Human | SFN | Chromatin Readers | GGCCAAGACCACTTTCGACG |
| Human | SFN | Chromatin Readers | AGCCCGGTCAGCCTACCAGG |
| Human | SFN | Chromatin Readers | AGCCATGGCCTCGTCGAAAG |
| Human | SFN | Chromatin Readers | AGCCTTCATGAAAGGCGCCG |
| Human | SFN | Chromatin Readers | GAGAGCCAGTCTGATCCAGA |
| Human | SHH | Transcription Elongation | GCACAGCGTGACCCTAAGCG |
| Human | SHH | Transcription Elongation | AGTGGCCAGGAGTGAAACTG |
| Human | SHH | Transcription Elongation | GGTGCTGGCCTCGTGCTACG |
| Human | SHH | Transcription Elongation | GTGAAACTGCGGGTGACCGA |
| Human | SHH | Transcription Elongation | TCTTCTACGTGATCGAGACG |
| Human | SHPRH | Chromatin Readers | TCAATAACATTACGAGATGG |
| Human | SHPRH | Chromatin Readers | GTTGAGACCCTACCAAAGAG |
| Human | SHPRH | Chromatin Readers | GCTTCATTGGAATATGCATG |
| Human | SHPRH | Chromatin Readers | TGTGTGATGTACAGATGTGT |
| Human | SHPRH | Chromatin Readers | TTTGGTAGGGTCTCAACACA |
| Human | SIN3A | DNA and RNA Modifications | CAGCCATCATCACCCAACAG |
| Human | SIN3A | DNA and RNA Modifications | TGAATCCCATTATCAGATCG |
| Human | SIN3A | DNA and RNA Modifications | CAGAGGACTATATGTATCGG |
| Human | SIN3A | DNA and RNA Modifications | CTTGCAGAACAATCAACCTG |
| Human | SIN3A | DNA and RNA Modifications | CGGATCCGGAAGTGTCAACG |
| Human | SIRT1 | Histone Acetylation | CTCTGAGCCATACCTATCCG |
| Human | SIRT1 | Histone Acetylation | TCGTACAAGTTGTCGGCCAG |
| Human | SIRT1 | Histone Acetylation | CAAAGGAGCAGATTAGTAGG |
| Human | SIRT1 | Histone Acetylation | GAATTGTGTCATAGGTTAGG |
| Human | SIRT1 | Histone Acetylation | AAGACCCAATAACAATGAGG |
| Human | SIRT2 | Histone Acetylation | TGGATGGAGAGCGAAAGTCG |
| Human | SIRT2 | Histone Acetylation | TGGCTGGGTGAATGCGACCA |
| Human | SIRT2 | Histone Acetylation | GGAGAAGGGTTACTTACATG |
| Human | SIRT2 | Histone Acetylation | CCAGCATAGATGCCCAGTCG |
| Human | SIRT2 | Histone Acetylation | CCCTCGCCTGCTCATCAACA |

| | | | |
|---|---|---|---|
| Human | SIRT3 | Histone Acetylation | AATGTCGGGCTTCACAACGC |
| Human | SIRT3 | Histone Acetylation | TCACGGCGTGGAAAGCCTAG |
| Human | SIRT3 | Histone Acetylation | TCTACACGCAGAACATCGAT |
| Human | SIRT3 | Histone Acetylation | GCCGGCATCAGCACACCCAG |
| Human | SIRT3 | Histone Acetylation | GCGCGGAACATGGCGTTCTG |
| Human | SIRT4 | Histone Acetylation | TACTGGGCGAGAAACTTCGT |
| Human | SIRT4 | Histone Acetylation | CAGGAATCTCCACCGAATCG |
| Human | SIRT4 | Histone Acetylation | TGGCACAAATAACCCAATGG |
| Human | SIRT4 | Histone Acetylation | ATGGCCTCCACATTGAACGC |
| Human | SIRT4 | Histone Acetylation | GCCAAGTCATCCGACCGTGT |
| Human | SIRT5 | Histone Acetylation | CCACTAGACATAAATCACAG |
| Human | SIRT5 | Histone Acetylation | GGGAGTTCTACCACTACCGG |
| Human | SIRT5 | Histone Acetylation | GCTGGGAAATCAATCGACTT |
| Human | SIRT5 | Histone Acetylation | GGAGTTCTACCACTACCGGC |
| Human | SIRT5 | Histone Acetylation | TCTCCAAACCACACGACGTG |
| Human | SIRT6 | Histone Acetylation | GTACGTCCGAGACACAGTCG |
| Human | SIRT6 | Histone Acetylation | GAGGATGTCGGTGAATTACG |
| Human | SIRT6 | Histone Acetylation | CATCCATGGCTACGTTGACG |
| Human | SIRT6 | Histone Acetylation | TGTCGGTGAATTACGCGGCG |
| Human | SIRT6 | Histone Acetylation | GATGTCGGTGAATTACGCGG |
| Human | SIRT7 | Histone Acetylation | GCGTCTATCCCAGACTACCG |
| Human | SIRT7 | Histone Acetylation | GGAAGTGTGATGACGTCATG |
| Human | SIRT7 | Histone Acetylation | TTGGTCGTCTACACAGGCGC |
| Human | SIRT7 | Histone Acetylation | AAATACTTGGTCGTCTACAC |
| Human | SIRT7 | Histone Acetylation | TGCACAGCGACTTCCGACTG |
| Human | SKA1 | Chromatin Remodeling | TAATTGTTCCAGATCTGACG |
| Human | SKA1 | Chromatin Remodeling | GAGGCTATCAGAGGTCCGAG |
| Human | SKA1 | Chromatin Remodeling | ACACTGCCGGAGGCTATCAG |
| Human | SKA1 | Chromatin Remodeling | CTCGGACCTCTGATAGCCTC |
| Human | SKA1 | Chromatin Remodeling | AAGTAACAGTAACCCAGAGC |
| Human | SKA2 | Chromatin Remodeling | TCATCCTGATTCAGCAAGTG |
| Human | SKA2 | Chromatin Remodeling | TCAACTATTCAACATGGAGG |
| Human | SKA2 | Chromatin Remodeling | GGTCGATAAGCTGGAACTGA |
| Human | SKA2 | Chromatin Remodeling | ACTATTCAACATGGAGGCGG |
| Human | SKA2 | Chromatin Remodeling | CAACTGAAAGCATTACCTCC |
| Human | SKA3 | Chromatin Remodeling | TCTGAAAGTTGTGGACTACG |
| Human | SKA3 | Chromatin Remodeling | TTCTCATAAGAAGAAATCGT |
| Human | SKA3 | Chromatin Remodeling | ACAGGCAGTGAACAACTATA |
| Human | SKA3 | Chromatin Remodeling | AGGCCATAGATACAGAATCC |
| Human | SKA3 | Chromatin Remodeling | ATACAACTGCTTGCAACAGG |
| Human | SMARCA2 | Chromatin Readers | CTCCCAGTCCTACTACACCG |
| Human | SMARCA2 | Chromatin Readers | ACACCTAGGCTATTCAAATG |

| | | | |
|---|---|---|---|
| Human | SMARCA2 | Chromatin Readers | TCATCCTCCTCCATTAAACG |
| Human | SMARCA2 | Chromatin Readers | CTGAACTGTATCGGGCCTCA |
| Human | SMARCA2 | Chromatin Readers | GAATCTTAGCCGATGAAATG |
| Human | SMARCA4 | Chromatin Readers | GTGCCCGTCAGGAACCACGA |
| Human | SMARCA4 | Chromatin Readers | TCGTCCTTGATGATCCACGA |
| Human | SMARCA4 | Chromatin Readers | ACGTCTCTGAGGTGGACGCG |
| Human | SMARCA4 | Chromatin Readers | GCAGCAGACAGACGAGTACG |
| Human | SMARCA4 | Chromatin Readers | AAGAAGCTCAAATTTACCCG |
| Human | SMARCC1 | Chromatin Readers | CAGGCCTAGCTGTTTATCGA |
| Human | SMARCC1 | Chromatin Readers | GGATGATTGGAACAAAGTGT |
| Human | SMARCC1 | Chromatin Readers | CCTAGCTGTTTATCGACGGA |
| Human | SMARCC1 | Chromatin Readers | TGTTAAAGGAGGAACTGTAG |
| Human | SMARCC1 | Chromatin Readers | CCTTCCGTCGATAAACAGCT |
| Human | SMARCC2 | Chromatin Readers | TATGCCTATTACCTGCACCA |
| Human | SMARCC2 | Chromatin Readers | AGGATGTAGAGGGTTCGCCA |
| Human | SMARCC2 | Chromatin Readers | GGTACGACCAGTCATGAAGA |
| Human | SMARCC2 | Chromatin Readers | ATGAGGATGAGAACAGTACG |
| Human | SMARCC2 | Chromatin Readers | CTCACCGGCAGTTTAGTGAG |
| Human | SMARCD1 | Chromatin Remodeling | GAAACGGCTAGATATCCAAG |
| Human | SMARCD1 | Chromatin Remodeling | AGTCATTGATGAAACCCTGA |
| Human | SMARCD1 | Chromatin Remodeling | GATTGCTACTCTAGACAACA |
| Human | SMARCD1 | Chromatin Remodeling | TGACAAACTCCCGCTCGTGA |
| Human | SMARCD1 | Chromatin Remodeling | CTGGTGGCATAAGCAAGGCA |
| Human | SMARCD2 | Chromatin Remodeling | GGGAGTGCACACGCAGACGA |
| Human | SMARCD2 | Chromatin Remodeling | GAATCTCGGAGAAACGGAGT |
| Human | SMARCD2 | Chromatin Remodeling | ACCAGACCATTGCTCGCAAG |
| Human | SMARCD2 | Chromatin Remodeling | AGCCTGTTACGACATCGATG |
| Human | SMARCD2 | Chromatin Remodeling | CCAGCAGCCTTGCCAATCGG |
| Human | SMARCD3 | Chromatin Remodeling | GCAGTAGAAGTAGCGACTGA |
| Human | SMARCD3 | Chromatin Remodeling | CTCTTGGTCTTACCTCAACG |
| Human | SMARCD3 | Chromatin Remodeling | GGTCTTCACATACTGCCACA |
| Human | SMARCD3 | Chromatin Remodeling | CCCTGACAACCACCTCGTTG |
| Human | SMARCD3 | Chromatin Remodeling | GCTGCAGGACTCCCATGACA |
| Human | SMARCE1 | Chromatin Remodeling | AGACGACGAGAACATTCCGA |
| Human | SMARCE1 | Chromatin Remodeling | TATGTAAGCAAGGTACGCGG |
| Human | SMARCE1 | Chromatin Remodeling | TTATGTAAGCAAGGTACGCG |
| Human | SMARCE1 | Chromatin Remodeling | GGAGAGCAACAGTGCAACAG |
| Human | SMARCE1 | Chromatin Remodeling | ATTTATGTAAGCAAGGTACG |
| Human | SMC1B | Chromatin Remodeling | GCTGATCGGCATCTATCCTG |
| Human | SMC1B | Chromatin Remodeling | ACAGTCTGATCAAGAAATCG |
| Human | SMC1B | Chromatin Remodeling | ATAGATGCCGATCAGCGCGT |
| Human | SMC1B | Chromatin Remodeling | ACATTTGCAAGGATTATCCG |

| | | | |
|---|---|---|---|
| Human | SMC1B | Chromatin Remodeling | GCCCTAGACAATACTAACAT |
| Human | SMN1 | Chromatin Readers | TTAATTTCATGGTACATGAG |
| Human | SMN1 | Chromatin Readers | AGAGAATCTGGACATATGGG |
| Human | SMN1 | Chromatin Readers | GATTCCGTGCTGTTCCGGCG |
| Human | SMN1 | Chromatin Readers | GCAACCGTCTTCTGACCAAA |
| Human | SMN1 | Chromatin Readers | TGTGCCGCGCCGGAACAGCA |
| Human | SMNDC1 | Chromatin Readers | ACAGGCTTGAGGTTCAACAG |
| Human | SMNDC1 | Chromatin Readers | GTGTTATGAAGCGGAGATTG |
| Human | SMNDC1 | Chromatin Readers | ACCAGCAAAGGTGATTGCAG |
| Human | SMNDC1 | Chromatin Readers | GAAAGTAGGAGACAAGTGTA |
| Human | SMNDC1 | Chromatin Readers | AGCATTGCCATAACCAGCAA |
| Human | SMYD1 | Histone Methylation | ATAGACAAGGCTCGTTCCGA |
| Human | SMYD1 | Histone Methylation | ACGTTAGTGAAGAACGCAAG |
| Human | SMYD1 | Histone Methylation | CTGGCATGCTGGTAACATTG |
| Human | SMYD1 | Histone Methylation | TGAGCGGGCTTATTCCGCAG |
| Human | SMYD1 | Histone Methylation | AATGTTACCAGCATGCCAGT |
| Human | SMYD2 | Histone Methylation | AAGCTAGGGAGACTCTACAT |
| Human | SMYD2 | Histone Methylation | CCTGTGCAAAGAGTACTACG |
| Human | SMYD2 | Histone Methylation | GTGCTCACGGTCAACGAGCG |
| Human | SMYD2 | Histone Methylation | GTTAGTCTTACAGTCTCCGA |
| Human | SMYD2 | Histone Methylation | GAACATTCCAGCTTGTGCAT |
| Human | SMYD3 | Histone Methylation | TGTACGCCAAGGGATCCGAG |
| Human | SMYD3 | Histone Methylation | TTGCACACCGTGTACGCCAA |
| Human | SMYD3 | Histone Methylation | GGTCCGAGTACCATAGAACA |
| Human | SMYD3 | Histone Methylation | AGGAAGCCATCCCGTCAGAG |
| Human | SMYD3 | Histone Methylation | AGAAGTCGAACGGAGTCTGG |
| Human | SMYD4 | Histone Methylation | TCTACAGAGGAGTCTCTACG |
| Human | SMYD4 | Histone Methylation | GCATAACTGCATCCGTCACA |
| Human | SMYD4 | Histone Methylation | GCATCCGTCACACGGAACTG |
| Human | SMYD4 | Histone Methylation | TACAGGCCATCCCAACTGAG |
| Human | SMYD4 | Histone Methylation | TTAACAGAGCACAGACACAT |
| Human | SMYD5 | Histone Methylation | AATGCACTTTATCGCTACCG |
| Human | SMYD5 | Histone Methylation | GTGGGTAGTGAATACTCCTG |
| Human | SMYD5 | Histone Methylation | GCTACTTGGACTGCTGTCAG |
| Human | SMYD5 | Histone Methylation | AGCCACTGAGCAATACCACC |
| Human | SMYD5 | Histone Methylation | CTTCCACGGAGACCCGCGCG |
| Human | SND1 | Chromatin Readers | CACTCAGTGCCTGCTCAACG |
| Human | SND1 | Chromatin Readers | CAGATATATCTGCAACACGG |
| Human | SND1 | Chromatin Readers | CAATGATGGCGCACCCTGAG |
| Human | SND1 | Chromatin Readers | CAGGTGATAGGGTACCCAGG |
| Human | SND1 | Chromatin Readers | CTTGGAATCTGCAAACTGCA |
| Human | SOX10 | Transcription Elongation | CCATGTTGGACATTACCTCG |

| | | | |
|---|---|---|---|
| Human | SOX10 | Transcription Elongation | ACGTTGCCGAAGTCGATGTG |
| Human | SOX10 | Transcription Elongation | TCACATCGACTTCGGCAACG |
| Human | SOX10 | Transcription Elongation | CCACGAGGTAATGTCCAACA |
| Human | SOX10 | Transcription Elongation | GGGAAGCCTCACATCGACTT |
| Human | SP100 | Chromatin Readers | TGATGAGATCACGATCACGG |
| Human | SP100 | Chromatin Readers | TGTGAGGTGTGCAACAAATG |
| Human | SP100 | Chromatin Readers | TCTCATCTTTCATAACCACA |
| Human | SP100 | Chromatin Readers | AGGAATCTGAAGTCCTGATG |
| Human | SP100 | Chromatin Readers | AGGATGTTCACGGAAGACCA |
| Human | SP110 | Chromatin Readers | GATTACGGAAATGTACACGG |
| Human | SP110 | Chromatin Readers | ATGTATCTAAGACCCTGGAG |
| Human | SP110 | Chromatin Readers | TGCCCGAAAGTCGAGATCAG |
| Human | SP110 | Chromatin Readers | AGACGCCTAGTACACCACGA |
| Human | SP110 | Chromatin Readers | ATTCCGAATGCACTTCACTG |
| Human | SP140 | Chromatin Readers | GCACCCAGCCTACTACCAGG |
| Human | SP140 | Chromatin Readers | AGATGAAGGAGCGGTCTCGG |
| Human | SP140 | Chromatin Readers | AAGCAACGGGATGATAGATG |
| Human | SP140 | Chromatin Readers | CTAGCTCCCTAGCAAGACGT |
| Human | SP140 | Chromatin Readers | TTACTGCCTTATGATACAGA |
| Human | SP140L | Chromatin Readers | TTTGGAAGCACTGTTCAGCG |
| Human | SP140L | Chromatin Readers | TGTACACCACTCTTTGTACA |
| Human | SP140L | Chromatin Readers | AAGGAAAGTGACCAAGCATG |
| Human | SP140L | Chromatin Readers | TGATGAGTTCCCGATCGCGG |
| Human | SP140L | Chromatin Readers | TTGTGATGAGTTCCCGATCG |
| Human | SPC24 | Chromatin Remodeling | GCAGAACCTACACGGCCGAG |
| Human | SPC24 | Chromatin Remodeling | CGGCTGCTGGAAACGCAAGA |
| Human | SPC24 | Chromatin Remodeling | GGAGTCTGGTGGACACCGAG |
| Human | SPC24 | Chromatin Remodeling | CCAGAGCCTTCTCAATGCGA |
| Human | SPC24 | Chromatin Remodeling | GGCTCACCTCCTCTATGTCG |
| Human | SPC25 | Chromatin Remodeling | TGTATGTTAATTATAAACCG |
| Human | SPC25 | Chromatin Remodeling | GGGACTAAGAGATACCTACA |
| Human | SPC25 | Chromatin Remodeling | ATTGTCCTACATAATGGTAG |
| Human | SPC25 | Chromatin Remodeling | CTTACATCTCAATGAAGCAA |
| Human | SPC25 | Chromatin Remodeling | CCTTACATCTCAATGAAGCA |
| Human | SSRP1 | DNA and RNA Modifications | CGAATGTCATAACGACCACG |
| Human | SSRP1 | DNA and RNA Modifications | ACCACGAGCAAAGTTGACAA |
| Human | SSRP1 | DNA and RNA Modifications | TACAGAATGATGGTCGACTG |
| Human | SSRP1 | DNA and RNA Modifications | CACTGGGACACATTGCTGAG |
| Human | SSRP1 | DNA and RNA Modifications | GTTCAACGACGTCTATCAGG |
| Human | STK31 | Chromatin Readers | ACCAACCATCATGTTCACCG |
| Human | STK31 | Chromatin Readers | GATGGTCCCATACTACCCTA |
| Human | STK31 | Chromatin Readers | TTTGCTTTAAACCGTGAACA |

| | | | |
|---|---|---|---|
| Human | STK31 | Chromatin Readers | TTAACCTCAGAACGTACCAA |
| Human | STK31 | Chromatin Readers | CACAGTACAAGCTAAGTACA |
| Human | SUPT16H | Transcription Elongation | TACAACAATGGCATCAACGT |
| Human | SUPT16H | Transcription Elongation | AACACCCACTGATACAACAA |
| Human | SUPT16H | Transcription Elongation | GAATGTCAGTACCTTGATTG |
| Human | SUPT16H | Transcription Elongation | ATATACCATCGCTGTAAAGG |
| Human | SUPT16H | Transcription Elongation | GAAGGAATTAAGACATGGTG |
| Human | SUPT4H1 | Transcription Elongation | TTATGATTATTATTACCTTG |
| Human | SUPT4H1 | Transcription Elongation | TGAAGCGGCTTACCCATCAA |
| Human | SUPT4H1 | Transcription Elongation | AATGAAGGGTAACCGAGAGA |
| Human | SUPT4H1 | Transcription Elongation | CTCTCAGGTAACTTTAAGCC |
| Human | SUPT4H1 | Transcription Elongation | GATGCATATCTACAAATGAA |
| Human | SUPT5H | Transcription Elongation | CCTCACCATCAATGCTCAGT |
| Human | SUPT5H | Transcription Elongation | AAATACGCCAAGTCATCTGT |
| Human | SUPT5H | Transcription Elongation | GCATGACCTCGACCTATGGG |
| Human | SUPT5H | Transcription Elongation | GTGAAGTTGGTCACATCACG |
| Human | SUPT5H | Transcription Elongation | TGTGAAGGGCTACATCTACG |
| Human | SUPT6H | Transcription Elongation | GCTGTGATTACCTAGACCGA |
| Human | SUPT6H | Transcription Elongation | TTCATCGGAACTGCACACCT |
| Human | SUPT6H | Transcription Elongation | GGAATACGATGAATCAGCCG |
| Human | SUPT6H | Transcription Elongation | CAATTTCCCTGAACTAAGCG |
| Human | SUPT6H | Transcription Elongation | GTCGGATAATCACATCACCC |
| Human | SUV39H1 | Histone Methylation | TCATAGACAACCTTGACGAG |
| Human | SUV39H1 | Histone Methylation | GTTCCTCTTAGAGATACCGA |
| Human | SUV39H1 | Histone Methylation | TTTGATTACAACATGCAAGG |
| Human | SUV39H1 | Histone Methylation | GGCCAGATCTACGACCGTCA |
| Human | SUV39H1 | Histone Methylation | TGTACACGAAGGCCCGCGGA |
| Human | SUV39H2 | Histone Methylation | CCATCTATGAATGCAACTCA |
| Human | SUV39H2 | Histone Methylation | TTGTGAAGAAGGCTAAACAA |
| Human | SUV39H2 | Histone Methylation | TTAATGTAATAGAAGTCTGA |
| Human | SUV39H2 | Histone Methylation | CATAGATGGGAGTACCAGGT |
| Human | SUV39H2 | Histone Methylation | CTTCACAATGTACTCAGCAA |
| Human | TAF1 | Histone Acetylation | TCTGGTATATGGACGCTGGG |
| Human | TAF1 | Histone Acetylation | CTGTTACCAGACATTGACTG |
| Human | TAF1 | Histone Acetylation | TATTATTATCCCAAGCAACA |
| Human | TAF1 | Histone Acetylation | GTGGTGCCATTGGACACATG |
| Human | TAF1 | Histone Acetylation | GGCATCTCGAGACATACTGA |
| Human | TAF10 | Transcription Elongation | GGTTTACGTACTGCCGAGCG |
| Human | TAF10 | Transcription Elongation | GAAGATTACACGCCTACGGT |
| Human | TAF10 | Transcription Elongation | GGAAGATTACACGCCTACGG |
| Human | TAF10 | Transcription Elongation | CCATGGTTAGAGTGTACTTG |
| Human | TAF10 | Transcription Elongation | GCTGGAAGATTACACGCCTA |

| | | | |
|---|---|---|---|
| Human | TAF11 | Transcription Elongation | GAGTCGCCCTCCGACAAAGG |
| Human | TAF11 | Transcription Elongation | CACTACAACCCAAACATATG |
| Human | TAF11 | Transcription Elongation | CGACACCGATGGAATCCCAG |
| Human | TAF11 | Transcription Elongation | CTTTGTCGGAGGGCGACTCG |
| Human | TAF11 | Transcription Elongation | ACTACAACCCAAACATATGA |
| Human | TAF12 | Transcription Elongation | AGAGGTTGATTAGGGCTGAG |
| Human | TAF12 | Transcription Elongation | TCGCAAGTCTAGCACCCTGG |
| Human | TAF12 | Transcription Elongation | CAGAAGCTCACAAACAGGTG |
| Human | TAF12 | Transcription Elongation | ACTGCAGTGGTAAAGATACC |
| Human | TAF12 | Transcription Elongation | GAGAGGTTGATTAGGGCTGA |
| Human | TAF13 | Transcription Elongation | TTCTTTGTAATAGACTCACA |
| Human | TAF13 | Transcription Elongation | TCTTGATTCGAAAGGACCCA |
| Human | TAF13 | Transcription Elongation | ATGTATGATGTATGGCTTTG |
| Human | TAF13 | Transcription Elongation | GGACCCAAGGAAGTTTGCCA |
| Human | TAF13 | Transcription Elongation | GAATCCTTATACTGAGTCAG |
| Human | TAF15 | Transcription Elongation | CAGCAACCATATAATAACCA |
| Human | TAF15 | Transcription Elongation | AGCAGTCCTATCATTCACAA |
| Human | TAF15 | Transcription Elongation | TGTGTCTACAGATCAAGTTG |
| Human | TAF15 | Transcription Elongation | CATACCTACTCACATCACGA |
| Human | TAF15 | Transcription Elongation | TCTACCTTGCAGATTTCCGG |
| Human | TAF1A | Transcription Elongation | AAGAAGAACACCGTAAACTG |
| Human | TAF1A | Transcription Elongation | TTGCGTGGGTTCAAGAAGAG |
| Human | TAF1A | Transcription Elongation | CTTCTTGAACCCACGCAAGG |
| Human | TAF1A | Transcription Elongation | CAAACATACGTAGAAACTGT |
| Human | TAF1A | Transcription Elongation | ACTCTTCTTGAACCCACGCA |
| Human | TAF1B | Transcription Elongation | GCATAGCTTAACTTGCCACG |
| Human | TAF1B | Transcription Elongation | AAATTATATGGACGTGACAG |
| Human | TAF1B | Transcription Elongation | TCAGTCTTGGCCTGACTACG |
| Human | TAF1B | Transcription Elongation | TATTGGATGACAGTTTCGAG |
| Human | TAF1B | Transcription Elongation | ACAGAAACAATTTAGCACAC |
| Human | TAF1C | Transcription Elongation | AGACGAGATGAAGAGTAGGG |
| Human | TAF1C | Transcription Elongation | TCTTCATCTCGTCTGTACCC |
| Human | TAF1C | Transcription Elongation | TGGAGCCTAAGATCCAGTGG |
| Human | TAF1C | Transcription Elongation | TGGCCATGTGGATCCCTCAG |
| Human | TAF1C | Transcription Elongation | ACAGACGAGATGAAGAGTAG |
| Human | TAF1D | Transcription Elongation | CTGTGGAACTTGCAAATCGA |
| Human | TAF1D | Transcription Elongation | AGTATACACTCAGTCTTACA |
| Human | TAF1D | Transcription Elongation | ATATAGCAAGCTGTTGCAAG |
| Human | TAF1D | Transcription Elongation | CCACAGCATCAGATGTCACA |
| Human | TAF1D | Transcription Elongation | GAGCAATTGATGGATAAATC |
| Human | TAF2 | Transcription Elongation | TCCTGGAACTCAGAAATACG |
| Human | TAF2 | Transcription Elongation | CATGACTGGAGGTTACGGTG |

| | | | |
|---|---|---|---|
| Human | TAF2 | Transcription Elongation | CAGAGTATGAATCAACACAA |
| Human | TAF2 | Transcription Elongation | CTAACCTTACATAGGGTACA |
| Human | TAF2 | Transcription Elongation | GCGGGCTCTACACCAGTCAG |
| Human | TAF3 | Chromatin Readers | GAATGTGAGTCGTGACCTTG |
| Human | TAF3 | Chromatin Readers | GAAATGACGGAATTTGGTGT |
| Human | TAF3 | Chromatin Readers | GGTGTAACAAGCCTGACGAT |
| Human | TAF3 | Chromatin Readers | TCAGCTTCAGGACTATCCAG |
| Human | TAF3 | Chromatin Readers | GCAGTCATGATTCCAACACA |
| Human | TAF4 | Transcription Elongation | CACGCGACAAAGAATCACGC |
| Human | TAF4 | Transcription Elongation | AATTGGTGGGCACGCTAACG |
| Human | TAF4 | Transcription Elongation | TCACGCGACAAAGAATCACG |
| Human | TAF4 | Transcription Elongation | AGAGGAAAGTGGACTGTCCG |
| Human | TAF4 | Transcription Elongation | TCATCGGGCCAACTATGCAA |
| Human | TAF4B | Transcription Elongation | GCAACACAGGAACGACTACG |
| Human | TAF4B | Transcription Elongation | GGCACAAATAGGAACTACTG |
| Human | TAF4B | Transcription Elongation | AGGAGTAACTGTGACGACAC |
| Human | TAF4B | Transcription Elongation | TGTGGCAAACACAGTGACCA |
| Human | TAF4B | Transcription Elongation | AGCAGGAAACTGGATTGTCG |
| Human | TAF5 | Transcription Elongation | TCTATCTGTTGCTTACTACG |
| Human | TAF5 | Transcription Elongation | TAAGACTAGATAATACTCGT |
| Human | TAF5 | Transcription Elongation | GGCTGAACCTGAGACGACCG |
| Human | TAF5 | Transcription Elongation | AGTGTTCTTACCGAGCTACT |
| Human | TAF5 | Transcription Elongation | AGTGGCTGTAGTAAAGTCAT |
| Human | TAF6 | Transcription Elongation | GAGTGTCCAGAAGTACATCG |
| Human | TAF6 | Transcription Elongation | GAGATAAAGGTACTGAACCG |
| Human | TAF6 | Transcription Elongation | GAGCCATAACGAGTCGTCCA |
| Human | TAF6 | Transcription Elongation | GACAACTCGTGGATGCTCCG |
| Human | TAF6 | Transcription Elongation | CCTGTGCGATCTCTTTGATG |
| Human | TAF7 | Transcription Elongation | ACAGTAGAGGCATATTCTGG |
| Human | TAF7 | Transcription Elongation | TGAAGCTGTTAGTACTCGGT |
| Human | TAF7 | Transcription Elongation | GCGTCATGGAATCGTCAGAG |
| Human | TAF7 | Transcription Elongation | CTGAAGCTGTTAGTACTCGG |
| Human | TAF7 | Transcription Elongation | GCAACTCAGCTCTTTGCAAG |
| Human | TAF9 | Transcription Elongation | ACTGAACCAACACTTAACCG |
| Human | TAF9 | Transcription Elongation | CGGGAAGAATAACAGTCCCG |
| Human | TAF9 | Transcription Elongation | CTGAACCAACACTTAACCGC |
| Human | TAF9 | Transcription Elongation | TGGCTTGATCAATGGCAAAG |
| Human | TAF9 | Transcription Elongation | AATTGTGGTCACATATCGGA |
| Human | TAF9B | Transcription Elongation | TGGGACAGACACCGTTTGTG |
| Human | TAF9B | Transcription Elongation | GAATCACAGAGTATGAACCA |
| Human | TAF9B | Transcription Elongation | CTGTGACGAAACCATGTTGG |
| Human | TAF9B | Transcription Elongation | TTTGGGACAGACACCGTTTG |

| | | | |
|---|---|---|---|
| Human | TAF9B | Transcription Elongation | ACAGCACCAACACTTAATCG |
| Human | TBP | Transcription Elongation | GGTTTGCTGCGGTAATCATG |
| Human | TBP | Transcription Elongation | AAACGCCGAATATAATCCCA |
| Human | TBP | Transcription Elongation | GATAAGAGAGCCACGAACCA |
| Human | TBP | Transcription Elongation | GTTTCGGGCACGAAGTGCAA |
| Human | TBP | Transcription Elongation | TTTACAACCAAGATTCACTG |
| Human | TCEA1 | Transcription Elongation | GCGTATTTCAGTACTTCCAG |
| Human | TCEA1 | Transcription Elongation | ACATACCTGCAGTAATTCCA |
| Human | TCEA1 | Transcription Elongation | CAAGCACTTCTGATTCTGTG |
| Human | TCEA1 | Transcription Elongation | GCTGCAGCTCTTCGAACAGG |
| Human | TCEA1 | Transcription Elongation | TTCTGTGCGGTTGAAGTGTA |
| Human | TCEA2 | Transcription Elongation | TGGAGATACGACTCCGTACA |
| Human | TCEA2 | Transcription Elongation | GACAGACATCCCGACTCGGG |
| Human | TCEA2 | Transcription Elongation | TGAGCTGAAGGAGATCCGTA |
| Human | TCEA2 | Transcription Elongation | TGAAGTATAAGAACCGTGTA |
| Human | TCEA2 | Transcription Elongation | GTTGACAGACATCCCGACTC |
| Human | TCF19 | Chromatin Readers | CTCCACCACAACGCAATCGG |
| Human | TCF19 | Chromatin Readers | ACTTCCTTCAGAAATCGACG |
| Human | TCF19 | Chromatin Readers | GAGGACTCCACTTACCCAGT |
| Human | TCF19 | Chromatin Readers | ACAAGTACGAGTCAAGCCTC |
| Human | TCF19 | Chromatin Readers | GATTTCCTCCGATTGCGTTG |
| Human | TCF20 | Chromatin Readers | TTCCACAAGGGACCCAACAG |
| Human | TCF20 | Chromatin Readers | GCACTCAAACTACTCAACCC |
| Human | TCF20 | Chromatin Readers | ATGGCAGAGTCATTAGATGG |
| Human | TCF20 | Chromatin Readers | ACACGGGTAATGGTATCGGA |
| Human | TCF20 | Chromatin Readers | AGAGTGGCTGCATAATCTTG |
| Human | TDRD1 | Chromatin Readers | TGGCGATTTAACACTCATCG |
| Human | TDRD1 | Chromatin Readers | CAGTACTGCTCCATTAACAG |
| Human | TDRD1 | Chromatin Readers | CCTTGTCCATAAGTAATCCA |
| Human | TDRD1 | Chromatin Readers | AGCACAACATATATCACCAA |
| Human | TDRD1 | Chromatin Readers | TTGGTATCGTGCTTTAGTCA |
| Human | TDRD10 | Chromatin Readers | AGGCTGCTGGATCATATCAG |
| Human | TDRD10 | Chromatin Readers | ACTCATCCATAGCGTCCGTG |
| Human | TDRD10 | Chromatin Readers | GCGAGTCCCGCGCATAACGG |
| Human | TDRD10 | Chromatin Readers | AACAGTTTATCAGAGAGTTG |
| Human | TDRD10 | Chromatin Readers | GAGGGCTCCACCGTTATGCG |
| Human | TDRD12 | Chromatin Readers | TAAACGATTCTACTACAACT |
| Human | TDRD12 | Chromatin Readers | ATGCTATTTAGCAACTACCC |
| Human | TDRD12 | Chromatin Readers | ACACAATGTCAGTACTGTCT |
| Human | TDRD12 | Chromatin Readers | AGACGTAATTGATTTGACAA |
| Human | TDRD12 | Chromatin Readers | ATTGTGAGGAGCTAAAGTGC |
| Human | TDRD3 | Chromatin Readers | ACTTTATTGGGAAGACAACA |

| | | | |
|---|---|---|---|
| Human | TDRD3 | Chromatin Readers | GTGTGTATCTCATGTCCAAG |
| Human | TDRD3 | Chromatin Readers | CAAACTCTGGAGTTCCGTAG |
| Human | TDRD3 | Chromatin Readers | CTGCGATTACAGATGACTGA |
| Human | TDRD3 | Chromatin Readers | CTGTCATACCCGAAGAATGG |
| Human | TDRD5 | Chromatin Readers | TTATACACGACATCCAGTGG |
| Human | TDRD5 | Chromatin Readers | GGGAACTGGTGTCTAATGGT |
| Human | TDRD5 | Chromatin Readers | AAACAAATCAGAGCTCAACT |
| Human | TDRD5 | Chromatin Readers | CCAGGCATGGCATTCCAGTG |
| Human | TDRD5 | Chromatin Readers | GCTTGCAGTAAATAGATGGG |
| Human | TDRD6 | Chromatin Readers | GAGCAATGAATGCACAACCC |
| Human | TDRD6 | Chromatin Readers | TTGGATCTAAATGTGTTGTG |
| Human | TDRD6 | Chromatin Readers | AGCGAACAGTAAACAGACAG |
| Human | TDRD6 | Chromatin Readers | TTGTATGATGAGTTGATCTG |
| Human | TDRD6 | Chromatin Readers | GATGTATTCTTAGTTGACCG |
| Human | TDRD7 | Chromatin Readers | CAACCACCTTCACTAGATCA |
| Human | TDRD7 | Chromatin Readers | GCTGGCCGTAAATGCCGAGG |
| Human | TDRD7 | Chromatin Readers | GTCCACATGCCAATAGATTG |
| Human | TDRD7 | Chromatin Readers | TGGCGATTAATACTGCGATG |
| Human | TDRD7 | Chromatin Readers | TAATACTGCGATGAGGGTCA |
| Human | TDRD9 | Chromatin Readers | GTTTGACGTTCAATTCAGCG |
| Human | TDRD9 | Chromatin Readers | GGATGTGTACCAGTACTCAG |
| Human | TDRD9 | Chromatin Readers | GCAGTGTGACATGCATCCCA |
| Human | TDRD9 | Chromatin Readers | AGGCGGAGCGCTGAACGTAG |
| Human | TDRD9 | Chromatin Readers | GTGTTAAAGGAGACTTGCAT |
| Human | TDRKH | Chromatin Readers | CTAGAGATCTTGGCTACCAG |
| Human | TDRKH | Chromatin Readers | GGGCTAGAATTAGTACACAA |
| Human | TDRKH | Chromatin Readers | TTGCCTATATCCTATACCGC |
| Human | TDRKH | Chromatin Readers | GACAGGTGCTCGGATTGATG |
| Human | TDRKH | Chromatin Readers | CAATAGAATGTAGTCTGGCA |
| Human | TERF2 | DNA and RNA Modifications | TGATGATTAAGGATCGCTGG |
| Human | TERF2 | DNA and RNA Modifications | GACTGTAGAAGAAAGCGAGT |
| Human | TERF2 | DNA and RNA Modifications | GATGCTGGGAGAGCTTGAGT |
| Human | TERF2 | DNA and RNA Modifications | GGGTTATGCAGTGTCTGTCG |
| Human | TERF2 | DNA and RNA Modifications | ACATTGATAGCTGATTCCAG |
| Human | TET1 | DNA and RNA Modifications | CCAACTACGAAAGACAACCA |
| Human | TET1 | DNA and RNA Modifications | GGAGTGTGTCCTGAGCCCAA |
| Human | TET1 | DNA and RNA Modifications | TCCTGCCAGCTAACACTCAG |
| Human | TET1 | DNA and RNA Modifications | GAGGAATAACACCCAAAGAG |
| Human | TET1 | DNA and RNA Modifications | AATGACGCAACAGCCTCATG |
| Human | TET2 | DNA and RNA Modifications | TGGAGAAAGACGTAACTTCG |
| Human | TET2 | DNA and RNA Modifications | TCTGCCCTGAGGTATGCGAT |
| Human | TET2 | DNA and RNA Modifications | CAGAAATGTACTGAGACACA |

| | | | |
|---|---|---|---|
| Human | TET2 | DNA and RNA Modifications | GATGCACAGGCCAATTAAGG |
| Human | TET2 | DNA and RNA Modifications | CTCACCCATCGCATACCTCA |
| Human | TET3 | DNA and RNA Modifications | GTCCTCCTATGCCTACACGA |
| Human | TET3 | DNA and RNA Modifications | GTGACTCCAAGAAGCCACTG |
| Human | TET3 | DNA and RNA Modifications | CCTCTACAATGGGTGCACCG |
| Human | TET3 | DNA and RNA Modifications | ACATCCAGTCAGTATTCAAG |
| Human | TET3 | DNA and RNA Modifications | CTTCAAGTACAGCGGCAACG |
| Human | TFPT | Chromatin Remodeling | GAAGTGGAGTTTGTGTCAGG |
| Human | TFPT | Chromatin Remodeling | AGGATGTGGCCACCAAATAG |
| Human | TFPT | Chromatin Remodeling | AGGAGTCCAGCACTCTCATG |
| Human | TFPT | Chromatin Remodeling | GCGCTGCCGGGAGATCGAGC |
| Human | TFPT | Chromatin Remodeling | GATTCCAGTTGGGTTTCTCG |
| Human | THOC1 | Transcription Elongation | ACGAGGAAGCTCCAACAACG |
| Human | THOC1 | Transcription Elongation | GAGCCCTGAGACTATTAGCA |
| Human | THOC1 | Transcription Elongation | TCTTCAGTTTGCCCGAAGCG |
| Human | THOC1 | Transcription Elongation | TATGGTAATAAAGCTAGCCA |
| Human | THOC1 | Transcription Elongation | TCTCCGTGCTAATAGTCTCA |
| Human | THOC5 | Transcription Elongation | GGTCGAGCATGACAGACAGT |
| Human | THOC5 | Transcription Elongation | TCAGTTGGACGACAAACGCA |
| Human | THOC5 | Transcription Elongation | AGGTCGAGCATGACAGACAG |
| Human | THOC5 | Transcription Elongation | AGGAAGAGATCAGACCCACG |
| Human | THOC5 | Transcription Elongation | CCTGAGTTGCTTGTATCCTG |
| Human | TLR3 | DNA and RNA Modifications | ACGACTGATGCTCCGAAGGG |
| Human | TLR3 | DNA and RNA Modifications | CCAGTTCAGAAAGAACGGAT |
| Human | TLR3 | DNA and RNA Modifications | ACTGGGAACCCATGATAGTG |
| Human | TLR3 | DNA and RNA Modifications | CATAGCTTCTCTGTAAGGCT |
| Human | TLR3 | DNA and RNA Modifications | CTCTGGAAACACGCAAACCC |
| Human | TLR7 | DNA and RNA Modifications | CTGGTGGGTTAACCATACGG |
| Human | TLR7 | DNA and RNA Modifications | CAGCTACTAGAGATACCGCA |
| Human | TLR7 | DNA and RNA Modifications | AGACCCAGCTGTGACCGAGT |
| Human | TLR7 | DNA and RNA Modifications | AAGGAATAGTCACCTCCGTA |
| Human | TLR7 | DNA and RNA Modifications | TCCAAAGAACCATGTGATCG |
| Human | TP53BP1 | Chromatin Readers | ATGACCACTTGGCTACAACA |
| Human | TP53BP1 | Chromatin Readers | GCTTACACCGCACATCAAGT |
| Human | TP53BP1 | Chromatin Readers | TCATGTGACGATGTAAGACA |
| Human | TP53BP1 | Chromatin Readers | CGATGTAAGACATGGCCATG |
| Human | TP53BP1 | Chromatin Readers | ACTTTCACAGAAGCTTTGTG |
| Human | TRDMT1 | DNA and RNA Modifications | TCCCACATACCGAACTCTGG |
| Human | TRDMT1 | DNA and RNA Modifications | GAAGCTACATAGAAGGGACA |
| Human | TRDMT1 | DNA and RNA Modifications | CAATAATAATACCTGACCAG |
| Human | TRDMT1 | DNA and RNA Modifications | GGCATGCACCACGCGCTGAG |
| Human | TRDMT1 | DNA and RNA Modifications | AGGTCCGTGTGCTTTACCAA |

| | | | |
|---|---|---|---|
| Human | TRIM24 | Chromatin Readers | ATGCATCACACCTTGCACGA |
| Human | TRIM24 | Chromatin Readers | CTGCTGCACTAGTAATCGTG |
| Human | TRIM24 | Chromatin Readers | CTCTACACACAAAGTCCCAG |
| Human | TRIM24 | Chromatin Readers | CACTAGGTGAACGTATTGGG |
| Human | TRIM24 | Chromatin Readers | GCACTAGGTGAACGTATTGG |
| Human | TRIM28 | Chromatin Readers | TGAAACTTCATCTCGCCATG |
| Human | TRIM28 | Chromatin Readers | CTTCCCAGGCAGTACCACTG |
| Human | TRIM28 | Chromatin Readers | CCAGCGGGTGAAGTACACCA |
| Human | TRIM28 | Chromatin Readers | AGGCGTTCAAGGCTCACTCG |
| Human | TRIM28 | Chromatin Readers | GAGAGCGCCTGCGACCCGAG |
| Human | TRIM33 | Chromatin Readers | GTGTTGTGTACCGATACTAC |
| Human | TRIM33 | Chromatin Readers | GTTTGCACACTGATCAATCG |
| Human | TRIM33 | Chromatin Readers | AGTGTGCAAACAATGCAAAG |
| Human | TRIM33 | Chromatin Readers | CTCAGCGTGCCCATCCCGGG |
| Human | TRIM33 | Chromatin Readers | GGCAACTCCAGTGCCCACTG |
| Human | TRIM37 | Other Histone Modifications | GGAAACACTTACCGGCAATG |
| Human | TRIM37 | Other Histone Modifications | TTGAGGGCTAATGCTAAAGG |
| Human | TRIM37 | Other Histone Modifications | TACTGGAACCCACATAACCA |
| Human | TRIM37 | Other Histone Modifications | CATTTGCGAGTAAGTCCAAA |
| Human | TRIM37 | Other Histone Modifications | TAGTAAACTACTGGTAGCAG |
| Human | TRIM66 | Chromatin Readers | GGAAGGTGTGACTACACAGG |
| Human | TRIM66 | Chromatin Readers | TGAAGGTGGACAAATGTCCA |
| Human | TRIM66 | Chromatin Readers | GAGTCGCTCCAGTCGCACAT |
| Human | TRIM66 | Chromatin Readers | TTCCTGCTGATCATCGAGTG |
| Human | TRIM66 | Chromatin Readers | ATTATTACCAGATTATCAAG |
| Human | TSFM | Transcription Elongation | ACTTACTTTACTGTATGCAG |
| Human | TSFM | Transcription Elongation | AAACACAACTGTATTAGTAG |
| Human | TSFM | Transcription Elongation | TGTCGGTAGTAGACTTTGTG |
| Human | TSFM | Transcription Elongation | TAGGGTCTGACAATGCATCA |
| Human | TSFM | Transcription Elongation | TGCGCGTGTTTCTGGTCGCG |
| Human | TWISTNB | Transcription Elongation | CATGTACTAAACAGCCAATG |
| Human | TWISTNB | Transcription Elongation | TAAGTCGGCAACTCTAGGCA |
| Human | TWISTNB | Transcription Elongation | CAAACCATGGAGATAAACAT |
| Human | TWISTNB | Transcription Elongation | CAGTCGCTACTCATGCCTGG |
| Human | TWISTNB | Transcription Elongation | AGACCCAGAGACATATGAAG |
| Human | UBE2A | Other Histone Modifications | CAACATAATGGTGTGGAACG |
| Human | UBE2A | Other Histone Modifications | CGCTCGCGACATGTCCACCC |
| Human | UBE2A | Other Histone Modifications | TCTCTTACCATCCTCAAACG |
| Human | UBE2A | Other Histone Modifications | TGGACATACTTCAGAACCGT |
| Human | UBE2A | Other Histone Modifications | CATCATAGGTTGGACTCCAA |
| Human | UBE2B | Other Histone Modifications | CTGGATTTCCAGGTTACAAG |
| Human | UBE2B | Other Histone Modifications | GGTGCGCCACTGACACCCAC |

| | | | |
|---|---|---|---|
| Human | UBE2B | Other Histone Modifications | AGGTGGGTCCTCTTGTAACC |
| Human | UBE2B | Other Histone Modifications | GCTGCGGAGCATGTCGACCC |
| Human | UBE2B | Other Histone Modifications | TTGGACTCCATCGATTCTGA |
| Human | UBE2E1 | Other Histone Modifications | TATGTCGGATGACGATTCGA |
| Human | UBE2E1 | Other Histone Modifications | GAAGTGAGCAGATAGAAAGG |
| Human | UBE2E1 | Other Histone Modifications | CCCTCATACACGGATCCTGG |
| Human | UBE2E1 | Other Histone Modifications | GAATACACCACCCTCATACA |
| Human | UBE2E1 | Other Histone Modifications | AATGGAGATCAACCATTCTA |
| Human | UBR1 | Other Histone Modifications | AAGACAGTAGTACAATCGTG |
| Human | UBR1 | Other Histone Modifications | ACCAACACCAGACAACGTAA |
| Human | UBR1 | Other Histone Modifications | ATTCTAACTTGTGGACCGAA |
| Human | UBR1 | Other Histone Modifications | AATGTCACAAAGCTGTGATG |
| Human | UBR1 | Other Histone Modifications | GATTCTAACTTGTGGACCGA |
| Human | UBR2 | Other Histone Modifications | AGAGGGTGTTCTATCAACAT |
| Human | UBR2 | Other Histone Modifications | CAACACATTATGGGAACACG |
| Human | UBR2 | Other Histone Modifications | GATCCAATTACACGTCAAGT |
| Human | UBR2 | Other Histone Modifications | CAAGTAAGCGAGAAACTGGG |
| Human | UBR2 | Other Histone Modifications | GTGTCATATCTGAAGCCACA |
| Human | UBR3 | Other Histone Modifications | TCGGCGCACAGCGACATGCA |
| Human | UBR3 | Other Histone Modifications | TGTTGAAGTCGTGTCCGGTG |
| Human | UBR3 | Other Histone Modifications | AACAGTACAACTAATCCAGT |
| Human | UBR3 | Other Histone Modifications | AACTGGCCAATCCAGACAGG |
| Human | UBR3 | Other Histone Modifications | GAACTTGAATTGATTCATCG |
| Human | UBR5 | Other Histone Modifications | TTGTTGTCTAAGAACGACGA |
| Human | UBR5 | Other Histone Modifications | GAATAACTCGAGTCTGCCGG |
| Human | UBR5 | Other Histone Modifications | TTGACGGGCGCTAAGCAAGG |
| Human | UBR5 | Other Histone Modifications | TTGATCCTTCAAACTTACGC |
| Human | UBR5 | Other Histone Modifications | TTTCTCCCAACAATCACAGT |
| Human | UBR7 | Chromatin Readers | ACGTCGACCAACGATACCAC |
| Human | UBR7 | Chromatin Readers | TCAGAACTAGAGCCGGCACA |
| Human | UBR7 | Chromatin Readers | CATGAATAGAGTCCAGCAAG |
| Human | UBR7 | Chromatin Readers | CTGAGGATGATGGATTGGTG |
| Human | UBR7 | Chromatin Readers | AGTCTTCACAGACTACGCAC |
| Human | UBTF | Transcription Elongation | CATGTGCAAGCTCAAATGGG |
| Human | UBTF | Transcription Elongation | AAGAGCATGACCAAGCTGCG |
| Human | UBTF | Transcription Elongation | CCGGAGGAACACTCACCTCG |
| Human | UBTF | Transcription Elongation | TCTTCTCGGAGGAGAAACGG |
| Human | UBTF | Transcription Elongation | AAGATGAACATGGCCGACAC |
| Human | UHRF1 | Chromatin Readers | TGCTCGGGACACGAACATGG |
| Human | UHRF1 | Chromatin Readers | GCGGGAACTCTACGCCAACG |
| Human | UHRF1 | Chromatin Readers | TCAGACAAGTCCTCCACCCA |
| Human | UHRF1 | Chromatin Readers | TGTGGGATGAGACGGAATTG |

| | | | |
|---|---|---|---|
| Human | UHRF1 | Chromatin Readers | CCATACCCTCTTCGACTACG |
| Human | UHRF2 | Chromatin Readers | CTAAAGTAAAGAAAGCTCCG |
| Human | UHRF2 | Chromatin Readers | TTTCCCACCACATACACGAC |
| Human | UHRF2 | Chromatin Readers | AAAGCCAGGATCAATAAGAC |
| Human | UHRF2 | Chromatin Readers | CTCGCGCAGCTCCTCAATCG |
| Human | UHRF2 | Chromatin Readers | TTAGGTGGACTATTAGAACA |
| Human | UNG | DNA and RNA Modifications | TCCAACTGCTCGGCACTCAG |
| Human | UNG | DNA and RNA Modifications | GATGAGCCGTCTGTAGTACA |
| Human | UNG | DNA and RNA Modifications | GCCAACTCTCATAAGGAGCG |
| Human | UNG | DNA and RNA Modifications | CTTGATGGGCACGAACCGTG |
| Human | UNG | DNA and RNA Modifications | ATGGACCTAATCAAGCTCAC |
| Human | USP15 | Other Histone Modifications | CCAAGTTACTTAGGCCACAG |
| Human | USP15 | Other Histone Modifications | GCAATCAGATTATAGCGGCA |
| Human | USP15 | Other Histone Modifications | AAGGTGTTCCTTAAGTGACT |
| Human | USP15 | Other Histone Modifications | TGAACCAGCGACTATCGACT |
| Human | USP15 | Other Histone Modifications | ACTTGCAAAGGTCAACTCAC |
| Human | USP22 | Other Histone Modifications | GCCATTGATCTGATGTACGG |
| Human | USP22 | Other Histone Modifications | CATGGAAATAATCGCCAAGG |
| Human | USP22 | Other Histone Modifications | TTCACCAGACCAGAGCACTT |
| Human | USP22 | Other Histone Modifications | GCGTGGTGGTTCCCGACACG |
| Human | USP22 | Other Histone Modifications | GCATATTCACGAGCATGCGA |
| Human | USP7 | Other Histone Modifications | ACCATACCCAAATTATTCCG |
| Human | USP7 | Other Histone Modifications | GGCAGTAGAACAGCTCGATG |
| Human | USP7 | Other Histone Modifications | GGCAACCTTTCAGTTCACTG |
| Human | USP7 | Other Histone Modifications | GAGTGATGGACACAACACCG |
| Human | USP7 | Other Histone Modifications | AGTGTAGCGACTCCTCTTTG |
| Human | UTY | Histone Methylation | TCAGAACTTCTGTTGAACTG |
| Human | UTY | Histone Methylation | AGTAGAAGTAGACCAAACCA |
| Human | UTY | Histone Methylation | ATCCTAATTCTGGCCAATCG |
| Human | UTY | Histone Methylation | GTCTGTTAGCCTGACAGTCG |
| Human | UTY | Histone Methylation | CAGAATGGTATGCCCAACAT |
| Human | WAC | Other Histone Modifications | GCTGCTCAGCTCTCTACACA |
| Human | WAC | Other Histone Modifications | TGCAGGCGTTAATGAACACG |
| Human | WAC | Other Histone Modifications | CGCGAAGAAGCGCATAACAT |
| Human | WAC | Other Histone Modifications | AGTACTCCAGTAGTTAAGCA |
| Human | WAC | Other Histone Modifications | TCCACTGGCAAACCCACTAG |
| Human | WDR61 | Transcription Elongation | CCTGTTGAAGGCGTGATGAG |
| Human | WDR61 | Transcription Elongation | GAATATTCTTTGGACACGAG |
| Human | WDR61 | Transcription Elongation | CTGGCCACAGGAACTCATGT |
| Human | WDR61 | Transcription Elongation | AAGTCCATAGATGCAGGACC |
| Human | WDR61 | Transcription Elongation | GAATGCAACGTTCAGCACCC |
| Human | WEE1 | Other Histone Modifications | TCATCAACAGAGCCCGCCAA |

| | | | |
|---|---|---|---|
| Human | WEE1 | Other Histone Modifications | CCATGAAGAGAGAACTACCC |
| Human | WEE1 | Other Histone Modifications | AGTTTGCTCTCCAAAGCTCG |
| Human | WEE1 | Other Histone Modifications | TCCAGCTCTGTTAAACTCCG |
| Human | WEE1 | Other Histone Modifications | AATGAATATACCTCAAGCCT |
| Human | WHSC1L1 | Histone Methylation | GCTTCCATTACGATGCACAA |
| Human | WHSC1L1 | Histone Methylation | AGGGCCTTAAACATGACTTG |
| Human | WHSC1L1 | Histone Methylation | GGATCACTTGAAACCATACA |
| Human | WHSC1L1 | Histone Methylation | GGGCCTTAAACATGACTTGG |
| Human | WHSC1L1 | Histone Methylation | GCTGGCTTGCAGTTACAGCG |
| Human | WTAP | DNA and RNA Modifications | TTACGTAAATCAACTCAGTG |
| Human | WTAP | DNA and RNA Modifications | ACGTAAATCAACTCAGTGCG |
| Human | WTAP | DNA and RNA Modifications | GCATATGTACAAGCTTTGGA |
| Human | WTAP | DNA and RNA Modifications | TGAAAGTGTAGACTCTCCCA |
| Human | WTAP | DNA and RNA Modifications | GGGAAAGGTAATCGAACTGT |
| Human | YEATS4 | Chromatin Readers | TCAGACACCAATGCAATGCT |
| Human | YEATS4 | Chromatin Readers | CAGCTTTAGCAAATGATACA |
| Human | YEATS4 | Chromatin Readers | CAGACACCAATGCAATGCTG |
| Human | YEATS4 | Chromatin Readers | ATTGTTGCATCATTGCTGTT |
| Human | YEATS4 | Chromatin Readers | AGCTATGGCAATCCTTTAAG |
| Human | YTHDF1 | DNA and RNA Modifications | AGTTTCAAAGCCGACCTCGT |
| Human | YTHDF1 | DNA and RNA Modifications | TTGGCACCTGGGATAACAAG |
| Human | YTHDF1 | DNA and RNA Modifications | TAGACCACGGAGCCTCATTG |
| Human | YTHDF1 | DNA and RNA Modifications | CGCCGCGTTTCTGTTGCGTG |
| Human | YTHDF1 | DNA and RNA Modifications | ATTGGCACCTGGGATAACAA |
| Human | YTHDF2 | DNA and RNA Modifications | TGGGTAGCACCTCGGAACCG |
| Human | YTHDF2 | DNA and RNA Modifications | GAATACTATAGACCAAGGGA |
| Human | YTHDF2 | DNA and RNA Modifications | GTCCATTACTAGTAACATCG |
| Human | YTHDF2 | DNA and RNA Modifications | AGACCAAGAATGGCATTGCA |
| Human | YTHDF2 | DNA and RNA Modifications | GGCATGAATACTATAGACCA |
| Human | YTHDF3 | DNA and RNA Modifications | TGGTGTATTTAGTCAACCTG |
| Human | YTHDF3 | DNA and RNA Modifications | CAACCGAAACTTAAACCCAA |
| Human | YTHDF3 | DNA and RNA Modifications | ATGGTGTATTTAGTCAACCT |
| Human | YTHDF3 | DNA and RNA Modifications | ACACTGAAGAGTAAATAGAG |
| Human | YTHDF3 | DNA and RNA Modifications | AGAGGAGGAAGCCATGCGTA |
| Human | YWHAB | Chromatin Readers | CTGCTGGGAGTTCGACACAG |
| Human | YWHAB | Chromatin Readers | TGAAGGCAGTCACAGAACAG |
| Human | YWHAB | Chromatin Readers | AGAGTACCGTGAGAAGATAG |
| Human | YWHAB | Chromatin Readers | AGGGAGACGAAGGAGACGCT |
| Human | YWHAB | Chromatin Readers | CTCTGTTGCCTACAAGAATG |
| Human | YWHAE | Chromatin Readers | CTATGGACTTCAGACATGCA |
| Human | YWHAE | Chromatin Readers | GGCTGCGGAGAACAGCCTAG |
| Human | YWHAE | Chromatin Readers | TGCCACAGGAAACGACAGGA |

| | | | |
|---|---|---|---|
| Human | YWHAE | Chromatin Readers | AATTTGCCACAGGAAACGAC |
| Human | YWHAE | Chromatin Readers | GAGCAGGCTGAGCGATACGA |
| Human | YWHAG | Chromatin Readers | CGAGCTTGACACCCTCAACG |
| Human | YWHAG | Chromatin Readers | TTTCGTTCCTCATTCGACAG |
| Human | YWHAG | Chromatin Readers | GAATGAGCCACTGTCGAATG |
| Human | YWHAG | Chromatin Readers | GGCCTACAAGAACGTTGTGG |
| Human | YWHAG | Chromatin Readers | GAGCCAGGCCTAATCGGATG |
| Human | YWHAH | Chromatin Readers | TTACTACCGCTACTTAGCAG |
| Human | YWHAH | Chromatin Readers | TGAGCTGGACACACTAAACG |
| Human | YWHAH | Chromatin Readers | CTGGATCTCATAGTAGAACA |
| Human | YWHAH | Chromatin Readers | TTTCGATCTTCATTGGAGAG |
| Human | YWHAH | Chromatin Readers | AAGCGACCTCTGCTAAGTAG |
| Human | YWHAQ | Chromatin Readers | CCACACACCCAATCCGCCTG |
| Human | YWHAQ | Chromatin Readers | GAAACGATAGATAATTCCCA |
| Human | YWHAQ | Chromatin Readers | GCCTACAAGAACGTGGTCGG |
| Human | YWHAQ | Chromatin Readers | CACCACACGCAACTTCAGCA |
| Human | YWHAQ | Chromatin Readers | TACCTTGCTGAAGTTGCGTG |
| Human | YWHAZ | Chromatin Readers | TTGTGGACATCGGATACCCA |
| Human | YWHAZ | Chromatin Readers | TGTGACTGATCGACAATCCC |
| Human | YWHAZ | Chromatin Readers | TCTCTGCTTGTGAAGCATTG |
| Human | YWHAZ | Chromatin Readers | ACCTTTCTTGTCATCACCAG |
| Human | YWHAZ | Chromatin Readers | GCCGCTGGTGATGACAAGAA |
| Human | YY1 | DNA and RNA Modifications | CGACTACATTGAACAAACGC |
| Human | YY1 | DNA and RNA Modifications | AGATATTGACCATGAGACAG |
| Human | YY1 | DNA and RNA Modifications | TGAACAAACGCTGGTCACCG |
| Human | YY1 | DNA and RNA Modifications | GGAGACCATCGAGACCACAG |
| Human | YY1 | DNA and RNA Modifications | TCTTATTACAACCATCGAAG |
| Human | ZBTB33 | DNA and RNA Modifications | AGCATGAAATTCATCACACA |
| Human | ZBTB33 | DNA and RNA Modifications | TTTGCCGAGTAATAACACAG |
| Human | ZBTB33 | DNA and RNA Modifications | AACTCCTTGAATGAGCAACG |
| Human | ZBTB33 | DNA and RNA Modifications | GGCGAGTTATTGCTAGCACT |
| Human | ZBTB33 | DNA and RNA Modifications | GCATGAAATTCATCACACAG |
| Human | ZBTB38 | DNA and RNA Modifications | ACAGAATACATAGGGACCCG |
| Human | ZBTB38 | DNA and RNA Modifications | AGGATTCCACTGACAAACCG |
| Human | ZBTB38 | DNA and RNA Modifications | GGATAACATACAAACCGGTG |
| Human | ZBTB38 | DNA and RNA Modifications | GCAGCTTCACAAGCCAACCC |
| Human | ZBTB38 | DNA and RNA Modifications | CAGAACTGGATTGCGCCGTG |
| Human | ZBTB4 | DNA and RNA Modifications | ACAGCAATGTACACTCGTGG |
| Human | ZBTB4 | DNA and RNA Modifications | GACACAGCAATGTACACTCG |
| Human | ZBTB4 | DNA and RNA Modifications | ACCGCACGAAGCATGAAGTG |
| Human | ZBTB4 | DNA and RNA Modifications | GGTCACGTAGGAACGCTCGC |
| Human | ZBTB4 | DNA and RNA Modifications | CAAACACTTTCTCACAATAG |

| | | | |
|---|---|---|---|
| Human | ZCWPW1 | Chromatin Readers | AGTATCTACCCAATCAGACA |
| Human | ZCWPW1 | Chromatin Readers | GTGTCATAATGCCCTACCAG |
| Human | ZCWPW1 | Chromatin Readers | CATACCCAAGCACTCCTGAA |
| Human | ZCWPW1 | Chromatin Readers | GGGCCTATCAGCGTGTAAGG |
| Human | ZCWPW1 | Chromatin Readers | TCTGATTGGGTAGATACTAC |
| Human | ZCWPW2 | Chromatin Readers | GAAATATGTAACTTATGACC |
| Human | ZCWPW2 | Chromatin Readers | TATGTAACTTATGACCCGGA |
| Human | ZCWPW2 | Chromatin Readers | TGGTTCATCATGATCAACCT |
| Human | ZCWPW2 | Chromatin Readers | TTTATCCATGATCTTGAATG |
| Human | ZCWPW2 | Chromatin Readers | CATGTATGTAAACAAAGTGT |
| Human | ZGPAT | Chromatin Readers | CGGCTTCAGAGACTTGTGAG |
| Human | ZGPAT | Chromatin Readers | TGAGTACCAGGCTTTCCGGG |
| Human | ZGPAT | Chromatin Readers | ATGGGCTATGAGTTTGGCAA |
| Human | ZGPAT | Chromatin Readers | AGAGACCGTTCCTAAAGCAG |
| Human | ZGPAT | Chromatin Readers | AGAAGATCGAGCGAACCCAG |
| Human | ZMYND11 | Chromatin Readers | ATAAACACCCGATGTACAGG |
| Human | ZMYND11 | Chromatin Readers | TCTCTTTAGGGTGCATACCG |
| Human | ZMYND11 | Chromatin Readers | ATTGTCGAAACTCTAACAGT |
| Human | ZMYND11 | Chromatin Readers | GGACGTTCCCACCATTCAAG |
| Human | ZMYND11 | Chromatin Readers | AGGTATATGTCTCGAGTCCA |
| Human | ZMYND8 | Chromatin Readers | CACTTAGCGTGATAAACCCG |
| Human | ZMYND8 | Chromatin Readers | TATCCGACAAGGAAATCCGG |
| Human | ZMYND8 | Chromatin Readers | TTTCAGTCAGAACTAAGACA |
| Human | ZMYND8 | Chromatin Readers | ACACTTAGCGTGATAAACCC |
| Human | ZMYND8 | Chromatin Readers | TAAGTGTCTGAGACTGACAT |
| Human | ZNF326 | Transcription Elongation | AGAGTCAAGGCTATTCCGGT |
| Human | ZNF326 | Transcription Elongation | GAACTGAACTATGTAAAGCA |
| Human | ZNF326 | Transcription Elongation | AGACTCATAAGGTCCAAACC |
| Human | ZNF326 | Transcription Elongation | TATGGCATGGACAATCACAG |
| Human | ZNF326 | Transcription Elongation | CCCGAACAAAGCCGCTTCGG |
| Human | ZNRD1 | Transcription Elongation | GGCTTCAACATCAACGTTCG |
| Human | ZNRD1 | Transcription Elongation | AGATGCGTTCAGCCGATGAA |
| Human | ZNRD1 | Transcription Elongation | TTGCCTGTGCAGACTTTGAG |
| Human | ZNRD1 | Transcription Elongation | ACGGTCACCTGTATTCGCTG |
| Human | ZNRD1 | Transcription Elongation | TTCCTTCATGACCACATCGA |
| Human | ZW10 | Chromatin Remodeling | GCAATCTGAACTGATGCCCG |
| Human | ZW10 | Chromatin Remodeling | GACTCACTGATACGGCATGT |
| Human | ZW10 | Chromatin Remodeling | AGTGCCATCACAAAGAATGG |
| Human | ZW10 | Chromatin Remodeling | CGGTTGATACGTGAAGATCC |
| Human | ZW10 | Chromatin Remodeling | GATATGTTGGTACAACATCA |
| Human | ZWILCH | Chromatin Remodeling | CTTGGTATGAAATCACACCT |
| Human | ZWILCH | Chromatin Remodeling | AAATGACATAGTATTCATAG |

| | | | |
|---|---|---|---|
| Human | ZWILCH | Chromatin Remodeling | TCAGTCTTATCATGGAACCA |
| Human | ZWILCH | Chromatin Remodeling | ACGTGGTGATATACAGCCAT |
| Human | ZWILCH | Chromatin Remodeling | CAAGTGCAGTTGATCAGCAA |
| Human | EAF1 | Transcription Elongation | TCAACTGCAAGTCATTTCTG |
| Human | EAF1 | Transcription Elongation | CAACTGCAAGTCATTTCTGA |
| Human | EAF1 | Transcription Elongation | GATGACGATAGCTCCAGCAG |
| Human | EAF1 | Transcription Elongation | ATGAGCTGGTTGCTTCCTTG |
| Human | EAF1 | Transcription Elongation | GGAGCTCTGAATGGCATAGG |
| Human | EAF1 | Transcription Elongation | GCTCCAACGAAGCCTCCAGT |
| Human | EAF1 | Transcription Elongation | GGTGGCTGTGACGTCTGTGG |
| Human | KAT8 | Histone Acetylation | ACTACGTGGGCTTTAACCGG |
| Human | KAT8 | Histone Acetylation | GAATTCTATGTACACTACGT |
| Human | KAT8 | Histone Acetylation | GCTCATAACTGAAAGCGATG |
| Human | KAT8 | Histone Acetylation | TAAGAAACCACCCATCACAG |
| Human | KAT8 | Histone Acetylation | ACTTGGGTCAGTGCCAGTGG |
| Human | KAT8 | Histone Acetylation | ACGGAGTCCACTGTGATGGG |
| Human | KAT8 | Histone Acetylation | GGACTCCGTCTGCCTCAAGT |
| Human | KAT8 | Histone Acetylation | GCTTGACTTGCTTGTGCTTG |
| Human | KAT8 | Histone Acetylation | CTTGACTTGCTTGTGCTTGG |
| Human | KAT8 | Histone Acetylation | TTGACTTGCTTGTGCTTGGG |
| Human | KAT8 | Histone Acetylation | GGGTGCCCACTTGAGGCAGA |
| Human | BRD8 | Chromatin Readers | CAGCATTGTGCAGAGGCCTA |
| Human | BRD8 | Chromatin Readers | TAGGCCTCTGCACAATGCTG |
| Human | BRD8 | Chromatin Readers | GCTGAAATTCAGCTGTGCTT |
| Human | BRD8 | Chromatin Readers | GCTACCACAGCATTGTGCAG |
| Human | BRD8 | Chromatin Readers | AACAGGCTGCAGGAAGACAT |
| Human | BRD8 | Chromatin Readers | AGAGAAATCTCTCTAAGGGT |
| Human | BRD8 | Chromatin Readers | ACATCAGCATCAGGTCTCGC |
| Human | BRD8 | Chromatin Readers | GTCTCGCAGGAATTGGGCCA |
| Human | BRD8 | Chromatin Readers | AAACCCAGGAGAGATATAGA |
| Human | BRD8 | Chromatin Readers | ACTAATTCAAGCTGGACACA |
| Human | BRD8 | Chromatin Readers | TAGCCGTCTATATCTCTCCT |
| Human | BRD8 | Chromatin Readers | GATGCAGAACTAATTCAAGC |
| Human | BRD8 | Chromatin Readers | AAAGAAATTGGAAGAAGAGG |
| Human | GTF2H5 | Transcription Elongation | GTACAGCAGAAACTGCTTCA |
| Human | BRD7 | Chromatin Readers | TCAGTCCATAGAAGAACTAA |
| Human | BRD7 | Chromatin Readers | TTAATGATCATGGAGTAGCC |
| Human | BRD7 | Chromatin Readers | CAGATTGCTCAGGGAACTCC |
| Human | BRD7 | Chromatin Readers | GAGTTCCCTGAGCAATCTGG |
| Human | BRD7 | Chromatin Readers | CTGGAGTTCCCTGAGCAATC |
| Human | BRD7 | Chromatin Readers | TGAGACCACCAGATTGCTCA |
| Human | BRD7 | Chromatin Readers | AACGTTCATTCTGGGCTTCC |

| | | | |
|---|---|---|---|
| Human | BRD7 | Chromatin Readers | AGGAGAGAAGCAGATTCCAG |
| Human | BRD7 | Chromatin Readers | AAGGAGAGAAGCAGATTCCA |
| Human | BRD7 | Chromatin Readers | AAAGGAGAGAAGCAGATTCC |
| Human | BRD7 | Chromatin Readers | TCTCTACATACTCCTCGTAG |
| Human | BRD7 | Chromatin Readers | AAACGATCATGACAAACACA |
| Human | BRD7 | Chromatin Readers | GCAAAGAAGCTGTTGCACTC |
| Human | GINS4 | Control | ACCAGACACAGATGAGCAGA |
| Human | GINS4 | Control | ACAGCTCCAGATGCAACCAG |
| Human | GINS4 | Control | GAGGGACTACGTGATTGACC |
| Human | GINS4 | Control | TACGTGATTGACCTGGAGAA |
| Human | GINS4 | Control | TTGAAAGATTGGAGCAGGCC |
| Human | GINS4 | Control | GCTCATTGAAAGATTGGAGC |
| Human | GINS4 | Control | ACGTCCTCAGCAGCTACTTG |
| Human | GINS4 | Control | GACACCGCAAGTAGCTGCTG |
| Human | GINS4 | Control | GAAGGTCAGCATCCACCAAA |
| Human | GINS4 | Control | CAGCATCCACCAAATGGAGA |
| Human | GINS4 | Control | AGTAGCTGCTGAGGACGTAG |
| Human | GINS4 | Control | CGCCATGAACTCTCTGGCAA |
| Human | GINS4 | Control | TGTGTTCGCCATGAACTCTC |
| Human | GINS4 | Control | GGCCTTTGCCAGAGAGTTCA |
| Human | CTR9 | Transcription Elongation | GCGCTGGAATACTACAAGCA |
| Human | CTR9 | Transcription Elongation | TTGGAAGCAGCACGTATAGA |
| Human | CTR9 | Transcription Elongation | CATATGGATTGCTTTGGCGC |
| Human | CTR9 | Transcription Elongation | AGCCTGCTTCTGCCTACTTG |
| Human | CTR9 | Transcription Elongation | ACCCTCAAGTAGGCAGAAGC |
| Human | CTR9 | Transcription Elongation | GCCTGCTTCTGCCTACTTGA |
| Human | CTR9 | Transcription Elongation | AAATGCAAGCTTTACCAAGA |
| Human | CTR9 | Transcription Elongation | TTCAACAAGAAGGATTACAG |
| Human | CTR9 | Transcription Elongation | AATGCAAGCTTTACCAAGAA |
| Human | CTR9 | Transcription Elongation | CAAGCTTTACCAAGAAGGGC |
| Human | CTR9 | Transcription Elongation | TTGCATTTCCTTCAACAAGA |
| Human | CTR9 | Transcription Elongation | TCTGGCATTCAGCAGAGCCC |
| Human | CTR9 | Transcription Elongation | CTTTGTGAAACTTAACAAAC |
| Human | CTR9 | Transcription Elongation | TCTAGAACTCAACAATAAAG |
| Human | CTR9 | Transcription Elongation | TATAGGCTCTGGAAAGAAGC |
| Human | CTR9 | Transcription Elongation | GGTCATAATCTTCCTGAACA |
| Human | CTR9 | Transcription Elongation | AGCTAGATCATTCCATGTTC |
| Human | CTR9 | Transcription Elongation | GGACAAATGTATATTTATCG |
| Human | CTR9 | Transcription Elongation | TGCATCTCAGTGCTTTGAGA |
| Human | CTR9 | Transcription Elongation | TTAGAACAGACTGATATACA |
| Human | CTR9 | Transcription Elongation | GATTCGTGTTGCTGTTCCAT |
| Human | CTR9 | Transcription Elongation | CAGGGTGCCCTTTCAGCCTA |

| | | | |
|---|---|---|---|
| Human | CTR9 | Transcription Elongation | TTGCTGTTCCATAGGCTGAA |
| Human | CTR9 | Transcription Elongation | CTTAGAACAGACTGATATAC |
| Human | CTR9 | Transcription Elongation | CCTGGCGAGATTATATGACG |
| Human | CTR9 | Transcription Elongation | CCACGTCATATAATCTCGCC |
| Human | CTR9 | Transcription Elongation | TAATCTCGCCAGGCTATATG |
| Human | CTR9 | Transcription Elongation | ACACATCGCCTCATATAGCC |
| Human | CTR9 | Transcription Elongation | GTTTCCCTTATCTCTAGCCA |
| Human | CTR9 | Transcription Elongation | TGAGGCTTCAGATTGGTTTA |
| Human | CTR9 | Transcription Elongation | GGAGCCATGGCTAGAGATAA |
| Human | CTR9 | Transcription Elongation | CTATTTGCGCCTAGGAGCCA |
| Human | CTR9 | Transcription Elongation | CTCTCAAACTTCTTCTGCCC |
| Human | CTR9 | Transcription Elongation | CTGGGCAGAAGAAGTTTGAG |
| Human | CTR9 | Transcription Elongation | TTTGATTGGCAATCTTCATT |
| Human | CTR9 | Transcription Elongation | GCGTCATCAAGATCGTGCTC |
| Human | CTR9 | Transcription Elongation | GCCCACCCGAGATCGAGAAA |
| Human | CTR9 | Transcription Elongation | CAATGGCATAGGAGCTGTTT |
| Human | CTR9 | Transcription Elongation | GAACTTAGCACACATCTATG |
| Human | CTR9 | Transcription Elongation | TTTGCTGAAGGCTAGACATG |
| Human | CTR9 | Transcription Elongation | GCCCGGGCCCTCTTCAAGTG |
| Human | CTR9 | Transcription Elongation | TAACTTGCCACACTTGAAGA |
| Human | CTR9 | Transcription Elongation | AAGTTGTACTCTATTTGGCC |
| Human | CTR9 | Transcription Elongation | ATGCAAACAGACTTTGCTGA |
| Human | CTR9 | Transcription Elongation | GGACGGCGTTCTAAGAAGGG |
| Human | CTR9 | Transcription Elongation | AAGAGGAGAAGACATCCAAA |
| Human | CTR9 | Transcription Elongation | CGTCTGCCTCCATCAATGAA |
| Human | CTR9 | Transcription Elongation | GGTGGACGGCGTTCTAAGAA |
| Human | CTR9 | Transcription Elongation | TTGCTGATGAAGGACATCCC |
| Human | CTR9 | Transcription Elongation | GGAAATGAATCGGAACCAGA |
| Human | CTR9 | Transcription Elongation | TGGAAATGAATCGGAACCAG |
| Human | KDM3A | Histone Methylation | ATCAGATCATCAAACCTGGA |
| Human | KDM3A | Histone Methylation | TATAAGTTATGAACCTGATG |
| Human | KDM3A | Histone Methylation | TTTGATCATGAATAGGATCG |
| Human | KDM3A | Histone Methylation | GTACTCGGGCAGTGGAATGT |
| Human | KDM3A | Histone Methylation | CATACTCCAAACCCACACCG |
| Human | KDM3A | Histone Methylation | ACTGGGTGTGTCCTCGGTGT |
| Human | KDM3A | Histone Methylation | CACTGGGTGTGTCCTCGGTG |
| Human | KDM3A | Histone Methylation | CACCCAGTGCAGGTTGAAGA |
| Human | KDM3A | Histone Methylation | ACCGAGGACACACCCAGTGC |
| Human | GTF2H3 | Transcription Elongation | CATCACGGCATCTATGCATT |
| Human | GTF2H3 | Transcription Elongation | ATAGATGCCGTGATGGTGCT |
| Human | GTF2H3 | Transcription Elongation | CGAATTTCCCAGCACCATCA |
| Human | GTF2H3 | Transcription Elongation | CATAGATGCCGTGATGGTGC |

| | | | |
|---|---|---|---|
| Human | GTF2H3 | Transcription Elongation | TCATTCTATGAATGTAGCAA |
| Human | GTF2H3 | Transcription Elongation | CATTCTATGAATGTAGCAAA |
| Human | GTF2H3 | Transcription Elongation | ATGAATGTAGCAAAGGGCTT |
| Human | GTF2H3 | Transcription Elongation | GTAGCAAAGGGCTTTGGCCA |
| Human | GTF2H3 | Transcription Elongation | TGTAGCAAAGGGCTTTGGCC |
| Human | PHF3 | Chromatin Readers | CTCAAATTTGAAGGTACCAG |
| Human | PHF3 | Chromatin Readers | CTTCTAAAGAGTTAGCTGCT |
| Human | PHF3 | Chromatin Readers | TCTTTAGAAGCTAGTTCTTC |
| Human | PHF3 | Chromatin Readers | CATGGCAACAGGTTTATGGT |
| Human | PHF3 | Chromatin Readers | ACCATAAACCTGTTGCCATG |
| Human | PHF3 | Chromatin Readers | TCAAGCACAGCAGATGGGCG |
| Human | PHF3 | Chromatin Readers | CTTTCTCAAGCACAGCAGAT |
| Human | PHF3 | Chromatin Readers | CCTGGGAGATGAACAACAGG |
| Human | PHF3 | Chromatin Readers | AATCCAAACATGCTTTGTGG |
| Human | PHF3 | Chromatin Readers | AAATCCAAACATGCTTTGTG |
| Human | PHF3 | Chromatin Readers | TGAGCAAAGCCAAAGCCTGG |
| Human | PHF3 | Chromatin Readers | CTGGAAATGGAAATCCAGGT |
| Human | HLTF | Chromatin Remodeling | ATAGTCCATTACATAGCATA |
| Human | HLTF | Chromatin Remodeling | GTCCATTACATAGCATAAGG |
| Human | HLTF | Chromatin Remodeling | AAAGGGAGCTTGTGCAGTGG |
| Human | HLTF | Chromatin Remodeling | GCCGGTTCTCTAATACGATC |
| Human | HLTF | Chromatin Remodeling | TCAGCAGCAGGATTCCAGGC |
| Human | HLTF | Chromatin Remodeling | TTTGTGTTTACTCGTTTGGA |
| Human | HLTF | Chromatin Remodeling | GAAATACCACTTAAAGCCTC |
| Human | HLTF | Chromatin Remodeling | ACAAATCCAGAGGCTTTAAG |
| Human | HLTF | Chromatin Remodeling | CAAATTGGCACAAATTGAAG |
| Human | HLTF | Chromatin Remodeling | GGTTGGACTACGCTATTACA |
| Human | HLTF | Chromatin Remodeling | GGTGTTTCAATAGCCTCAGC |
| Human | HLTF | Chromatin Remodeling | TGAATGACTTGGCAAATACA |
| Human | HLTF | Chromatin Remodeling | GCTGCTCATTCTGAATGACT |
| Human | HLTF | Chromatin Remodeling | CTGAATGACTTGGCAAATAC |
| Human | HLTF | Chromatin Remodeling | CAGGAACTGTTAAAGAATCC |
| Human | HLTF | Chromatin Remodeling | TGTTTCATTTCGTCAGCATT |
| Human | HLTF | Chromatin Remodeling | TGTTCTGATTTCATTAATTT |
| Human | CHAF1A | DNA and RNA Modifications | TTTGCAATCCATGGCTGTGG |
| Human | CHAF1A | DNA and RNA Modifications | AGTCATCATTGATTTGACAG |
| Human | CHAF1A | DNA and RNA Modifications | TTTAGACCGAAACTTGTCAA |
| Human | CHAF1A | DNA and RNA Modifications | TGGGACGAGTTCCTGGCTAA |
| Human | CHAF1A | DNA and RNA Modifications | TAAGGTCCGCCAGAAACTGA |
| Human | CHAF1A | DNA and RNA Modifications | GACCTTATGGTTCTCAGGGT |
| Human | CHAF1A | DNA and RNA Modifications | CCTTGGCCTTCAGTTTCTGG |
| Human | CHAF1A | DNA and RNA Modifications | GGGACGAGTTCCTGGCTAAG |

| | | | |
|---|---|---|---|
| Human | CHAF1A | DNA and RNA Modifications | GGAGGGCGACTGTATGATCG |
| Human | CHAF1A | DNA and RNA Modifications | AGAACTCAGTGTATGAGAAG |
| Human | CHAF1A | DNA and RNA Modifications | CCGATGTCACATAGCTCCAC |
| Human | CHAF1A | DNA and RNA Modifications | CTGTATGATCGTGGATGTCC |
| Human | CHAF1A | DNA and RNA Modifications | GCAACCACACCGGCAGCCCG |
| Human | CHAF1A | DNA and RNA Modifications | AGCCCTGGAGGCTAAACTTG |
| Human | CHAF1A | DNA and RNA Modifications | GGAGAGACAGGAAGCCCTGG |
| Human | CHAF1A | DNA and RNA Modifications | CAGAGAAGGCCGAAATCACG |
| Human | CHAF1A | DNA and RNA Modifications | CAAAGAGATCAGGAGCGTCT |
| Human | CHAF1A | DNA and RNA Modifications | AGCTCAAGTTACGTGCAGAA |
| Human | RPS6KA5 | Other Histone Modifications | CCTCGAACATCTCCACAAGT |
| Human | RPS6KA5 | Other Histone Modifications | ATGACAAGGCAGTTGACTGG |
| Human | RPS6KA5 | Other Histone Modifications | CCAACTTGTGGAGATGTTCG |
| Human | RPS6KA5 | Other Histone Modifications | ATGACTGGCTTAAATGGTGC |
| Human | RPS6KA5 | Other Histone Modifications | AGGGCTGCGGGAGAATAAGT |
| Human | RPS6KA5 | Other Histone Modifications | TAGGATGGAAGGAGCAACAA |
| Human | RPS6KA5 | Other Histone Modifications | AGTTCTGAGAAGCTGTTTCA |
| Human | RPS6KA5 | Other Histone Modifications | TCTAATTCATCTCGAATGAC |
| Human | RPS6KA5 | Other Histone Modifications | GTAAGCCACATGCATGATGT |
| Human | RPS6KA5 | Other Histone Modifications | GCATAATGAAGGGTGAAGCA |
| Human | RPS6KA5 | Other Histone Modifications | GTATTTGTTAAAGGCGTGGA |
| Human | RPS6KA5 | Other Histone Modifications | TGCTTTCACACAGGTATGCA |
| Human | RPS6KA5 | Other Histone Modifications | GTGGAAGGTTGCTTTCACAC |
| Human | RPS6KA5 | Other Histone Modifications | CCGGAAGATCCTAGAATATC |
| Human | RPS6KA5 | Other Histone Modifications | ACAGGTATGCACGGCAGCTC |
| Human | EHMT2 | Histone Methylation | TTGTACCTGTCCGTGAAGCA |
| Human | EHMT2 | Histone Methylation | GAAGGTGATCCTGATGCTGT |
| Human | EHMT2 | Histone Methylation | CCAGGTTGTCCAACAGCATC |
| Human | EHMT2 | Histone Methylation | CCTGATGCTGTTGGACAACC |
| Human | EHMT2 | Histone Methylation | CTCCAGCCTGCAGCAGCACA |
| Human | EHMT2 | Histone Methylation | CAGCACATGGCAGATCTCCA |
| Human | EHMT2 | Histone Methylation | CTGCATGCAGCCGCCCAGAA |
| Human | EHMT2 | Histone Methylation | AGCCGCCCAGAAGGGCTCCG |
| Human | EHMT2 | Histone Methylation | CTCCACGGAGCCCTTCTGGG |
| Human | EHMT2 | Histone Methylation | TGTAACGGGCTACCTCCAGG |
| Human | EHMT2 | Histone Methylation | CCTGGAGGTAGCCCGTTACA |
| Human | EHMT2 | Histone Methylation | CGTTACATGGTGCAGCGTGG |
| Human | EHMT2 | Histone Methylation | GCCCGTTACATGGTGCAGCG |
| Human | EHMT2 | Histone Methylation | GCTGCTGAGCACAGGACAGG |
| Human | EHMT2 | Histone Methylation | TGCAGAGCACAAGCACATCG |
| Human | EHMT2 | Histone Methylation | TGATCCGCATGCTACTGACG |
| Human | EHMT2 | Histone Methylation | GATCCGCATGCTACTGACGC |

| | | | |
|---|---|---|---|
| Human | EHMT2 | Histone Methylation | CAGAAGGACTTCGGCGATGG |
| Human | EHMT2 | Histone Methylation | GCGCGCATTCAGAAGGACTT |
| Human | EHMT2 | Histone Methylation | ATTCAGAAGGACTTCGGCGA |
| Human | EHMT2 | Histone Methylation | GGTCACAGCGCGCATTCAGA |
| Human | EHMT2 | Histone Methylation | CCGTGAAGGAGGCCCAGTGC |
| Human | EHMT2 | Histone Methylation | GGAATAACAGCACGCAGTCA |
| Human | EHMT2 | Histone Methylation | GCTGTTATTCCTGTCACGTG |
| Human | EHMT2 | Histone Methylation | GTGCTGTTATTCCTGTCACG |
| Human | EHMT2 | Histone Methylation | TGCTGTTATTCCTGTCACGT |
| Human | EHMT2 | Histone Methylation | CCGAGCTGCGATGTGCAGGG |
| Human | EHMT2 | Histone Methylation | GGTGCTGGTATGACAAGGAT |
| Human | EHMT2 | Histone Methylation | CGGTGCTGGTATGACAAGGA |
| Human | EHMT2 | Histone Methylation | CTTAGACAACAAGGATGGAG |
| Human | EHMT2 | Histone Methylation | CTTCATCTGCGAGTATGTCG |
| Human | EHMT2 | Histone Methylation | TTCCCGAGACATCCGGACTG |
| Human | EHMT2 | Histone Methylation | TTCGACTTAGACAACAAGGA |
| Human | EHMT2 | Histone Methylation | GAACTGAAGAAGGCGATGCG |
| Human | EHMT2 | Histone Methylation | TGCACTTCTCAGAGCCACAT |
| Human | EHMT2 | Histone Methylation | CTTCTCAGAGCCACATTGGC |
| Human | EHMT2 | Histone Methylation | GGCGGCGGGAGCTGCAGCGG |
| Human | EHMT2 | Histone Methylation | GGAGGAAGAGGAAGAAGAAG |
| Human | EHMT2 | Histone Methylation | AGAAGAAGAAGAGGAAGAGG |
| Human | EHMT2 | Histone Methylation | AGAAGAAGAGGAAGAGGAGG |
| Human | EHMT2 | Histone Methylation | AGAGGAAGAGGAGGAGGAAG |
| Human | EHMT2 | Histone Methylation | AGAGGAGGAAGAAGAAGAAG |
| Human | CENPJ | Chromatin Remodeling | AAACAGCTGAAGGAAGCAGA |
| Human | CENPJ | Chromatin Remodeling | AGGCCCATTAAAGCTGCTAT |
| Human | CENPJ | Chromatin Remodeling | GATCGAGCATTGTCACCAGG |
| Human | CENPJ | Chromatin Remodeling | ATTGTCAGAGTACAACGTGA |
| Human | CENPJ | Chromatin Remodeling | CAGTCATCCTGATGGAAAGG |
| Human | CENPJ | Chromatin Remodeling | TGGGATCGAGCATTGTCACC |
| Human | CENPJ | Chromatin Remodeling | GGGACAAGCTGCCTCTCCCA |
| Human | HAUS3 | Control | ACGTAATAAATGTCAATTGA |
| Human | HAUS3 | Control | TAACTAGCCACAAATCTCTG |
| Human | HAUS3 | Control | TAAGATCAGTAATGAACTTC |
| Human | HAUS3 | Control | TTCAAGTTTCATAATCTCAC |
| Human | HAUS3 | Control | AATTGAATTAAGAAAGCATC |
| Human | HAUS3 | Control | AAATTGAATTAAGAAAGCAT |
| Human | HAUS3 | Control | TCTAACTCATGGAAACCTTG |
| Human | HAUS3 | Control | AAAGAATTGTTTCTAACTCA |
| Human | HAUS3 | Control | TTCTTTAGTACAAGATCAGT |
| Human | HAUS3 | Control | ATTTCTCAGCCACTTCCTCA |

| | | | |
|---|---|---|---|
| Human | HAUS3 | Control | TCATGGAAACCTTGAGGAAG |
| Human | KMT2A | Histone Methylation | GCAGCCATTAATTCAGATGG |
| Human | KMT2A | Histone Methylation | AGGGAATTACACATCTGTGT |
| Human | KMT2A | Histone Methylation | ACACAGATGTGTAATTCCCT |
| Human | KMT2A | Histone Methylation | TCAAGAGGAAGATGGACCAA |
| Human | KMT2A | Histone Methylation | TAGATCTAGAAGGAGTCAAG |
| Human | KMT2A | Histone Methylation | TCATAGACCCAATCATCATG |
| Human | KMT2A | Histone Methylation | TATTGGATATCAGTGTTCCA |
| Human | KMT2A | Histone Methylation | CTGGAACACTGATATCCAAT |
| Human | KMT2A | Histone Methylation | AAAGAGCTTATCTTCACAGT |
| Human | KMT2A | Histone Methylation | TGTGGTGCTCCAGTATACCC |
| Human | KMT2A | Histone Methylation | CAGAAAGTATTGAAGATGCC |
| Human | KMT2A | Histone Methylation | GATCTGAAAGCCATCATCAC |
| Human | KMT2A | Histone Methylation | TGAGGAACACAACTGCATCA |
| Human | KMT2A | Histone Methylation | TTGCAGGTGTTAACGGTTTG |
| Human | KMT2A | Histone Methylation | TTTGAAATTTCCAGTGATGA |
| Human | KMT2A | Histone Methylation | GCCATGCGTAAGATCTACCG |
| Human | KMT2A | Histone Methylation | GCGTAAGATCTACCGAGGAG |
| Human | KMT2A | Histone Methylation | GTTCCGAATTGATGACTCAG |
| Human | KMT2A | Histone Methylation | TCCTCGGTAGATCTTACGCA |
| Human | KMT2A | Histone Methylation | ACTGTGGCGCCAAGAAATGC |
| Human | KMT2A | Histone Methylation | TTCTTGGCGCCACAGTTGCA |
| Human | KMT2A | Histone Methylation | TTTCTTGGCGCCACAGTTGC |
| Human | KMT2A | Histone Methylation | GACGTCGATCGAGGCGGTGT |
| Human | KMT2A | Histone Methylation | GTACAAACACCACAGTCCTC |
| Human | KMT2A | Histone Methylation | GGACGTCGATCGAGGCGGTG |
| Human | KMT2A | Histone Methylation | ATATTGCGACCACCAAACTT |
| Human | KMT2A | Histone Methylation | TTAGATAAGCCCAAGTTTGG |
| Human | KMT2A | Histone Methylation | GGAAGGGCTCACAACAGACT |
| Human | KMT2A | Histone Methylation | AAACTCTACATGCCCACTAC |
| Human | KMT2A | Histone Methylation | GGAGAACGAGCGCCCTCTGG |
| Human | KMT2A | Histone Methylation | AGCTATCACCCTGAGTGCCT |
| Human | KMT2A | Histone Methylation | TCAGGCTACAAAGCAGCTGC |
| Human | KMT2A | Histone Methylation | CGTTTGTGGAAGGCAACATC |
| Human | KMT2A | Histone Methylation | TCTTCTTTGTGGGTTTGGTG |
| Human | KMT2A | Histone Methylation | AATTCTGTCACGTTTGTGGA |
| Human | KMT2A | Histone Methylation | GAGAGTAAGATGATGCAATG |
| Human | KMT2A | Histone Methylation | ATGTGGAAAGTGTGATCGCT |
| Human | KMT2A | Histone Methylation | TCATAACATTTGTCACAGAG |
| Human | KMT2A | Histone Methylation | CAAGTGTAGGCCACACTTTC |
| Human | KMT2A | Histone Methylation | AATGTGGAAAGTGTGATCGC |
| Human | KMT2A | Histone Methylation | GATGACAGTGCTAATGATGC |

| | | | |
|---|---|---|---|
| Human | KMT2A | Histone Methylation | GCTGGTCGTTTACTATATAT |
| Human | KMT2A | Histone Methylation | TGTGCGTTATGTTTGACTTA |
| Human | KMT2A | Histone Methylation | CACATGTAAATTGTGCTTTG |
| Human | KMT2A | Histone Methylation | TGTGAGACAGCAACCCACGG |
| Human | KMT2A | Histone Methylation | ACACATGAAGTGATAGTTGC |
| Human | KMT2A | Histone Methylation | GAAAGACACAGTTCTTGGCT |
| Human | KMT2A | Histone Methylation | GGCCGGGACGCGTTGCCGCG |
| Human | KMT2A | Histone Methylation | GGGCCGCCACCGCCGACCGG |
| Human | KMT2A | Histone Methylation | AGGGCCGGGACGCGTTGCCG |
| Human | KMT2A | Histone Methylation | CGATGACGAAGACGAAGACG |
| Human | KMT2A | Histone Methylation | CTGCAGGTCTCGGCCGCCAT |
| Human | KMT2A | Histone Methylation | CCATCGGCACCAACCTGCGC |
| Human | KMT2A | Histone Methylation | GCGCAGGTTGGTGCCGATGG |
| Human | KMT2A | Histone Methylation | CACCAACCTGCGCCGGTTCC |
| Human | KMT2A | Histone Methylation | GCACCAACCTGCGCCGGTTC |
| Human | KMT2A | Histone Methylation | GTCCTTCTTCATTCGGCAAA |
| Human | KMT2A | Histone Methylation | TGGTCAAGTCCTTCTTCATT |
| Human | KMT2A | Histone Methylation | GAAGGACTTGACCATGCTGT |
| Human | ELOA | Transcription Elongation | AGACATTCTTGCGGAGACTG |
| Human | ELOA | Transcription Elongation | GTCTACTGTAATAGGCAGGG |
| Human | ELOA | Transcription Elongation | AATGTCTACTGTAATAGGCA |
| Human | ELOA | Transcription Elongation | TAGACATTCTTGCGGAGACT |
| Human | ELOA | Transcription Elongation | TGATCAGCTGTATCGCATAG |
| Human | ELOA | Transcription Elongation | CGGGTTCAAGAACAGAGTAT |
| Human | ELOA | Transcription Elongation | TTGAAGAAACAGATCAATTA |
| Human | ELOA | Transcription Elongation | GATTCAATCTTTGAAGTGGG |
| Human | ELOA | Transcription Elongation | ATCGATTCAATCTTTGAAGT |
| Human | ELOA | Transcription Elongation | CATCGATTCAATCTTTGAAG |
| Human | ING3 | Chromatin Readers | TGGCTGCGTTGGATTGACAG |
| Human | ING3 | Chromatin Readers | ACCAAGATTGCCCTATAGAA |
| Human | ING3 | Chromatin Readers | ACCATTCTATAGGGCAATCT |
| Human | ING3 | Chromatin Readers | TGGTTCCATTATGGCTGCGT |
| Human | ING3 | Chromatin Readers | TCAATCCAACGCAGCCATAA |
| Human | ING3 | Chromatin Readers | ACACGATGATAAGGAAGAAG |
| Human | ING3 | Chromatin Readers | CGATGATAAGGAAGAAGAGG |
| Human | ING3 | Chromatin Readers | TGATAAGGAAGAAGAGGAGG |
| Human | CDK6 | Transcription Elongation | CCACAGCGTGACGACCACTG |
| Human | CDK6 | Transcription Elongation | CCTCAGTGGTCGTCACGCTG |
| Human | CDK6 | Transcription Elongation | ATCAAACAACCTGACCACGT |
| Human | CDK6 | Transcription Elongation | CAAACAACCTGACCACGTTG |
| Human | NAE1 | Control | GGCAACATGATTACCCACAG |
| Human | NAE1 | Control | AAGAAAGATGCTGCCGCTGT |

| | | | |
|---|---|---|---|
| Human | NAE1 | Control | AAAGAAAGATGCTGCCGCTG |
| Human | NAE1 | Control | AAGTTGAAGTCTTGTCTCAC |
| Human | TWISTNB | Transcription Elongation | TGATGCAGATGACACTCCAA |
| Human | TWISTNB | Transcription Elongation | TAATGCGAATGGCATCTGGG |
| Human | TWISTNB | Transcription Elongation | TTTGGTGAGCATTTCAAAGG |
| Human | SPC25 | Chromatin Remodeling | TCTTAGTCCCGCCATCTGAC |
| Human | SPC25 | Chromatin Remodeling | AGTACTGACTGCAAATATCC |
| Human | SPC25 | Chromatin Remodeling | TCTGCTAGGCCCTCAAGATG |
| Human | SPC25 | Chromatin Remodeling | CTGCTAGGCCCTCAAGATGA |
| Human | SPC25 | Chromatin Remodeling | TGCTAGGCCCTCAAGATGAG |
| Human | TAF7 | Transcription Elongation | AAGCAGATTGACAACATGAA |
| Human | TAF7 | Transcription Elongation | TGAATCAGATGAACAGCACC |
| Human | TAF7 | Transcription Elongation | ATATGCCTCTACTGTGAGAA |
| Human | TAF7 | Transcription Elongation | AATATGCCTCTACTGTGAGA |
| Human | TAF7 | Transcription Elongation | TTTCAAGCTTTCCATAACAC |
| Human | TAF7 | Transcription Elongation | AGCTTTCCATAACACAGGGC |
| Human | TAF7 | Transcription Elongation | TTCAAGCTTTCCATAACACA |
| Human | CDK8 | Transcription Elongation | GAAACTTGATTATATGCCAG |
| Human | CDK8 | Transcription Elongation | TAAGCAGCTTCTGAAGCAAG |
| Human | CDK8 | Transcription Elongation | AGCCAAGAGGAAAGATGGGA |
| Human | CDK8 | Transcription Elongation | AAGTGAAGCTGAGCAGCGAG |
| Human | TAF1 | Histone Acetylation | CATCTTCATCATAATCTGAG |
| Human | TAF1 | Histone Acetylation | TCATAATCTGAGTGGCAAAG |
| Human | TAF1 | Histone Acetylation | TGATGAATGTAAGAAGCACT |
| Human | TAF1 | Histone Acetylation | CTTGGTAAATGATGAAGGGT |
| Human | TAF1 | Histone Acetylation | GAAGATACCAGCAGACGATG |
| Human | TAF1 | Histone Acetylation | ACTCTTCCCGAGATGGGTAG |
| Human | TAF1 | Histone Acetylation | GTAAACGCCTCTACCCATCT |
| Human | TAF1 | Histone Acetylation | CCACACTCCAGTCAATGCAA |
| Human | TAF1 | Histone Acetylation | TGTCCTATGAGGGAGACGGT |
| Human | TAF1 | Histone Acetylation | TTGGTATCATACAAATCAGG |
| Human | TAF1 | Histone Acetylation | TGGGTATGAGGTATCAGAGG |
| Human | TAF1 | Histone Acetylation | GCGCTGACTTCAAACGCACA |
| Human | TAF1 | Histone Acetylation | GAAGGGCAAGTGTCTGCTAG |
| Human | TAF1 | Histone Acetylation | AAAGTCTCGAATATGCGTAT |
| Human | TAF1 | Histone Acetylation | GACCTCATAAGTCCATCCAC |
| Human | TAF1 | Histone Acetylation | ATGGACGACAGCGTCACCAT |
| Human | TAF1 | Histone Acetylation | ATTGATGATAGACTCCAAGA |
| Human | TAF1 | Histone Acetylation | CAAATAGTCACAGTGAACAG |
| Human | TAF1 | Histone Acetylation | CCGGCGCCGCACAGACCCTA |
| Human | TAF1 | Histone Acetylation | CATCTTTGACCTACAGAACA |
| Human | TAF1 | Histone Acetylation | GCATCAAGAGCGTTACAAAG |

| | | | |
|---|---|---|---|
| Human | AFF4 | Transcription Elongation | CCAGAAGCTGATGCATCCAG |
| Human | AFF4 | Transcription Elongation | GTATGAATAGCCGTTAGAGG |
| Human | AFF4 | Transcription Elongation | GACTTCCAGTAGCTCCAAGG |
| Human | AFF4 | Transcription Elongation | AACAGCAACAAGGAGACGAG |
| Human | AFF4 | Transcription Elongation | TGCAGATGGAGGACAGTTAG |
| Human | AFF4 | Transcription Elongation | CTCCAAGGAGGTTAAGGAAA |
| Human | AFF4 | Transcription Elongation | TCAAAGCAGAAGAAGACCGA |
| Human | AFF4 | Transcription Elongation | TGCGTATGAAAGAACGGGAA |
| Human | AFF4 | Transcription Elongation | TGTGCTGCGTATGAAAGAAC |
| Human | AFF4 | Transcription Elongation | ATGTGCTGCGTATGAAAGAA |
| Human | AFF4 | Transcription Elongation | CTTTCATACGCAGCACATTC |
| Human | KAT7 | Histone Acetylation | CACATTTGGCCATGTGCCGG |
| Human | KAT7 | Histone Acetylation | GGCACATGGCCAAATGTGTG |
| Human | KAT7 | Histone Acetylation | TGTGTGTGGAAACACCCACC |
| Human | KAT7 | Histone Acetylation | AATATGAAGTGTCCTACACC |
| Human | KAT7 | Histone Acetylation | GCTGAGAGGTTATGATACAG |
| Human | KAT7 | Histone Acetylation | AACTCTCTAGGACACCTTAC |
| Human | KAT7 | Histone Acetylation | TGTCCTAGAGAGTTACAGCC |
| Human | KAT7 | Histone Acetylation | GGGTGACTCGAGCAGATCGT |
| Human | KAT7 | Histone Acetylation | TTGAGAACTCTGGCTTAGCC |
| Human | KAT7 | Histone Acetylation | GGTGACTCGAGCAGATCGTC |
| Human | KAT7 | Histone Acetylation | ACAGACAGTTCAGAAAGTGA |
| Human | KAT7 | Histone Acetylation | GGCTTAGCCTGGCTGAGGAG |
| Human | WEE1 | Other Histone Modifications | CAGAATGAATATTGTAATGG |
| Human | WEE1 | Other Histone Modifications | CAATGGCATGAAATCAGACA |
| Human | WEE1 | Other Histone Modifications | GAAAGCACTTGTGGTATCCG |
| Human | LMNA | DNA and RNA Modifications | GAAAGCGCGCAATACCAAGA |
| Human | LMNA | DNA and RNA Modifications | GCGCGCAATACCAAGAAGGA |
| Human | LMNA | DNA and RNA Modifications | AGCGCGCAATACCAAGAAGG |
| Human | LMNA | DNA and RNA Modifications | TCCAATGAGGACCAGTCCAT |
| Human | LMNA | DNA and RNA Modifications | GTCCAATGAGGACCAGTCCA |
| Human | LMNA | DNA and RNA Modifications | TGCAGCCGGGTGATGCGGGT |
| Human | LMNA | DNA and RNA Modifications | GTGATGCGGGTGGGCGACAG |
| Human | LMNA | DNA and RNA Modifications | GCGGCGCGCCACCCGCAGCG |
| Human | LMNA | DNA and RNA Modifications | TGTCGCCCACCCGCATCACC |
| Human | LMNA | DNA and RNA Modifications | CAGCGGCGCGCCACCCGCAG |
| Human | LMNA | DNA and RNA Modifications | GACCTGCTCCATCACCACCA |
| Human | LMNA | DNA and RNA Modifications | CACTCGCAGCTACCGCAGTG |
| Human | LMNA | DNA and RNA Modifications | CGAGCGCAGGTTGTACTCAG |
| Human | LMNA | DNA and RNA Modifications | AGCGCAGGTTGTACTCAGCG |
| Human | LMNA | DNA and RNA Modifications | GCGAGTGACCGTGACACTGG |
| Human | GTF2E1 | Transcription Elongation | GAAGTAGTAGTTATGGCGAG |

| | | | |
|---|---|---|---|
| Human | GTF2E1 | Transcription Elongation | GGGTAGAGACTGCTGCAGAC |
| Human | GTF2E1 | Transcription Elongation | CATATCCTCCTCTTTCACAC |
| Human | GTF2E1 | Transcription Elongation | GTTTATCAAATGCAGAATGA |
| Human | GTF2E1 | Transcription Elongation | CAGCGGAAAGTTCCTGTCAT |
| Human | GTF2E1 | Transcription Elongation | GTGAAAGTACTACTACAGAC |
| Human | GTF2E1 | Transcription Elongation | CAGCTCTTTGATCCTATGAC |
| Human | GTF2E1 | Transcription Elongation | TCTCAATTCTTCTTCTCATG |
| Human | GTF2E1 | Transcription Elongation | AAAGGAAGCATATATAGCAA |
| Human | GTF2E1 | Transcription Elongation | AAGGAAGCATATATAGCAAT |
| Human | BPTF | Chromatin Readers | TTGCCATTTACGAGGTACTG |
| Human | BPTF | Chromatin Readers | TACGAGGTACTGCGGAACTT |
| Human | BPTF | Chromatin Readers | GTTCCGCAGTACCTCGTAAA |
| Human | BPTF | Chromatin Readers | GAATGTCATTGCCATTTACG |
| Human | BPTF | Chromatin Readers | CCAAGAGCAGTGCACACTCA |
| Human | BPTF | Chromatin Readers | ACCTATGGACCTTGCCACCA |
| Human | BPTF | Chromatin Readers | TTGTACTCTTTCTTCCATGG |
| Human | BPTF | Chromatin Readers | TCCATGGTGGCAAGGTCCAT |
| Human | BPTF | Chromatin Readers | ATGTGTGAAGCCACCTCTTG |
| Human | BPTF | Chromatin Readers | TAGGGTTTGTCACAAACTTG |
| Human | BPTF | Chromatin Readers | GTTTGTGACAAACCCTACAA |
| Human | BPTF | Chromatin Readers | TGTGAAGCCACCTCTTGAGG |
| Human | BPTF | Chromatin Readers | ACCTCTTGAGGAGGTGCCAG |
| Human | BPTF | Chromatin Readers | CGTTGGCATCTTGCAAAGTG |
| Human | BPTF | Chromatin Readers | TGGTACCATGGGCGCTGCGT |
| Human | BPTF | Chromatin Readers | AGATGCCAACGCAGCGCCCA |
| Human | BPTF | Chromatin Readers | TTCCAGATTAGGCTGGATGG |
| Human | BPTF | Chromatin Readers | GTTCTTCTGGCACATCCTGG |
| Human | BPTF | Chromatin Readers | AAGAGACTAGTCCCTCTGAA |
| Human | BPTF | Chromatin Readers | GGTTCTTCTGGCACATCCTG |
| Human | BPTF | Chromatin Readers | TGGTAAGAACTTTGGACCCA |
| Human | BPTF | Chromatin Readers | ATTCAAAGCAAATGAATTGG |
| Human | BPTF | Chromatin Readers | AGTCAGTCTCAGAGTAGATA |
| Human | BPTF | Chromatin Readers | AATGGAACGGTTCTGTCCAT |
| Human | BPTF | Chromatin Readers | ATCAATTGAAAGAGAAGAAA |
| Human | BPTF | Chromatin Readers | GCAACAGGCTAAGAAACGAC |
| Human | BPTF | Chromatin Readers | CAGGGCATTTGCTGAGAGAG |
| Human | BPTF | Chromatin Readers | TGAGAGAGTGGAGAAAGAAA |
| Human | BPTF | Chromatin Readers | GGCACAAGCAGTTGAGCAAC |
| Human | BPTF | Chromatin Readers | ACGGAAGCGTGAAGAGAGTG |
| Human | BPTF | Chromatin Readers | CCAGGATTCTGAGGACGACG |
| Human | BPTF | Chromatin Readers | CGAGGAGACCCAGGATTCTG |
| Human | BPTF | Chromatin Readers | GGATTCTGAGGACGACGAGG |

| | | | |
|---|---|---|---|
| Human | BPTF | Chromatin Readers | CGGCGACGCCGAGGAGACCC |
| Human | BPTF | Chromatin Readers | CGATGACTCCGATTATCCGG |
| Human | BPTF | Chromatin Readers | CTCCGATTATCCGGAGGAGA |
| Human | BPTF | Chromatin Readers | TTCCATCTCCTCCGGATAAT |
| Human | BPTF | Chromatin Readers | TCGTCGTCTTCCATCTCCTC |
| Human | BPTF | Chromatin Readers | TGTAACCAGTGTGCCTCCCG |
| Human | BPTF | Chromatin Readers | GATGTCCACACTGCCGCCTG |
| Human | BPTF | Chromatin Readers | AACCAGTGTGCCTCCCGTGG |
| Human | BPTF | Chromatin Readers | TGCTGTCGTGGAAACAGTGG |
| Human | BPTF | Chromatin Readers | TGTCGTGGAAACAGTGGTGG |
| Human | BPTF | Chromatin Readers | CGTGGAAACAGTGGTGGTGG |
| Human | BPTF | Chromatin Readers | GGTGGTGGTGCTGGCTGTGG |
| Human | BPTF | Chromatin Readers | GTGGTGGCTGTTGTTGCCAA |
| Human | BPTF | Chromatin Readers | GAGGAGCTGGTGTCACTGGG |
| Human | BPTF | Chromatin Readers | TGGAGGAGCTGGTGTCACTG |
| Human | BPTF | Chromatin Readers | GGAGGAGCTGGTGTCACTGG |
| Human | BPTF | Chromatin Readers | CTGGAGGAGCTGGTGTCACT |
| Human | BPTF | Chromatin Readers | ATTCGAGAACTTCTGCACAC |
| Human | DDX39B | Transcription Elongation | TTGAACAGCTCGACATGCGT |
| Human | DDX39B | Transcription Elongation | TTGGATCTTGCATGAACTTG |
| Human | DDX39B | Transcription Elongation | TCCATTCCCAGAATGGCCTG |
| Human | DDX39B | Transcription Elongation | ACTGCCCGCATATCGTCGTG |
| Human | DDX39B | Transcription Elongation | TGTCCTTGAGTTCAACCAGG |
| Human | DDX39B | Transcription Elongation | TGACACCTACCTGCATCGGG |
| Human | DDX39B | Transcription Elongation | ACTTCACAAAGATCACCACC |
| Human | DDX39B | Transcription Elongation | CGAGAAAGCCTCTCCTCCTG |
| Human | DDX39B | Transcription Elongation | TCTGGCCACCCGATGCAGGT |
| Human | DDX39B | Transcription Elongation | CTCCGGGCCATTGTCGACTG |
| Human | DDX39B | Transcription Elongation | AGCCACAGTCGACAATGGCC |
| Human | DDX39B | Transcription Elongation | GCTCAAGCCAGAGTTGCTCC |
| Human | DDX39B | Transcription Elongation | TGCTCAAGCCAGAGTTGCTC |
| Human | GTF2E2 | Transcription Elongation | ATGAAGACACGGCATCAGCG |
| Human | GTF2E2 | Transcription Elongation | TTGTGAATTACATGAAGACA |
| Human | GTF2E2 | Transcription Elongation | TTGGACTCAAGCAGAAACAA |
| Human | GTF2E2 | Transcription Elongation | CTGATCTAAGAGCCTAAGTA |
| Human | GTF2E2 | Transcription Elongation | GCTGATCTAAGAGCCTAAGT |
| Human | FOXP4 | DNA and RNA Modifications | GAGACCCTGTGTGAAGACCT |
| Human | FOXP4 | DNA and RNA Modifications | GGTGTTTGATAAACTGGCCC |
| Human | FOXP4 | DNA and RNA Modifications | CTGGCCCAGGTCTTCACACA |
| Human | FOXP4 | DNA and RNA Modifications | TGTTGAGGTGTTTGATAAAC |
| Human | FOXP4 | DNA and RNA Modifications | ACTGGCCCAGGTCTTCACAC |
| Human | FOXP4 | DNA and RNA Modifications | GCGCACGGCGTTCTTCCAGG |

| | | | |
|---|---|---|---|
| Human | FOXP4 | DNA and RNA Modifications | GGGTTTCCAGGATGGCCTGG |
| Human | FOXP4 | DNA and RNA Modifications | GTCCGCGTGGAGAACGTCAA |
| Human | FOXP4 | DNA and RNA Modifications | AGCTCGCCAAGGAGAGCGAG |
| Human | FOXP4 | DNA and RNA Modifications | GCTGGAGATCCAGCTCGCCA |
| Human | FOXP4 | DNA and RNA Modifications | GCTCGCTCTCCTTGGCGAGC |
| Human | FOXP4 | DNA and RNA Modifications | CGAGCGGCTGCAGGCCATGA |
| Human | FOXP4 | DNA and RNA Modifications | CTGCAGCCGCTCGCTCTCCT |
| Human | FOXP4 | DNA and RNA Modifications | CGCAGCCCAAAGAGGCACTG |
| Human | FOXP4 | DNA and RNA Modifications | CTGTAGCTGCTGGAGCATGA |
| Human | FOXP4 | DNA and RNA Modifications | CCGCAGCCCAAAGAGGCACT |
| Human | CCAR2 | Transcription Elongation | AAGCCCATCCTCCTTCAGCG |
| Human | CCAR2 | Transcription Elongation | AAGCAACAGTTCTGGGTCCA |
| Human | CCAR2 | Transcription Elongation | ATGGCAGCCTGATTAACGTG |
| Human | METTL3 | DNA and RNA Modifications | TTCACATGGAACTGCCCTAT |
| Human | METTL3 | DNA and RNA Modifications | CAGTTCCATGTGAATATCCC |
| Human | METTL3 | DNA and RNA Modifications | ATTCACATGGAACTGCCCTA |
| Human | METTL3 | DNA and RNA Modifications | ATCATTCGGACAGGCCGTAC |
| Human | METTL3 | DNA and RNA Modifications | GGACAGGCCGTACAGGTCAC |
| Human | METTL3 | DNA and RNA Modifications | GGTTCAACCAGTGACCTGTA |
| Human | USP7 | Other Histone Modifications | TGCATGGCAAGACCATGACG |
| Human | USP7 | Other Histone Modifications | AGGATCGGTCACTTCACTCC |
| Human | USP7 | Other Histone Modifications | CTGAATCTGATTCCACGTCA |
| Human | USP7 | Other Histone Modifications | AGTTAACAAAGTCATTTGGG |
| Human | USP7 | Other Histone Modifications | GGCTTACAGGAAGCAGAGAA |
| Human | USP7 | Other Histone Modifications | ACATGGAGGATGACACCAGT |
| Human | USP7 | Other Histone Modifications | TGATGGACACAACACCGCGG |
| Human | USP7 | Other Histone Modifications | TGACATCATAGTATTTCAGA |
| Human | USP7 | Other Histone Modifications | CAGTGAATTACCCACCGCAA |
| Human | TDRD12 | Chromatin Readers | GGAAGAAGGACAGGTGTGTG |
| Human | TDRD12 | Chromatin Readers | GGTGTGTGTGGTCTATTGTG |
| Human | TDRD12 | Chromatin Readers | GGGCACACGATGACTGCGAG |
| Human | TDRD12 | Chromatin Readers | AAACTGTCAACACTGGTAGG |
| Human | TDRD12 | Chromatin Readers | TCATCGATGAAGGCAGGACG |
| Human | TDRD12 | Chromatin Readers | CTTGTAGAGTTCATCGATGA |
| Human | PHF7 | Chromatin Readers | GGCTATGCCTTCGAGAACCT |
| Human | PHF7 | Chromatin Readers | GCTATGCCTTCGAGAACCTG |
| Human | PHF7 | Chromatin Readers | CGAAGGCATAGCCAGCAAAC |
| Human | PHF7 | Chromatin Readers | AGAAACTTCCATCTGCCTTG |
| Human | PHF7 | Chromatin Readers | AAGGCAGATGGAAGTTTCTG |
| Human | PHF7 | Chromatin Readers | TCTGCCTTGTGGCCAAGAAA |
| Human | PHF7 | Chromatin Readers | AGCTGCTATCAACTGCCAGA |
| Human | PHF7 | Chromatin Readers | ATTTCTGTATGCACTTGCGG |

| | | | |
|---|---|---|---|
| Human | PHF7 | Chromatin Readers | CATATTTCTGTATGCACTTG |
| Human | PHF7 | Chromatin Readers | AGAAATGCTTTGCTGATGTG |
| Human | PHF7 | Chromatin Readers | GAAATGCTTTGCTGATGTGT |
| Human | PHF7 | Chromatin Readers | TAGATGGCTTGACTACAACA |
| Human | BAZ1B | Other Histone Modifications | AAGCAGTCAGCTAACACACA |
| Human | BAZ1B | Other Histone Modifications | ATCATGACCAAATATGCTGT |
| Human | BAZ1B | Other Histone Modifications | CTTCATACTCTTCCCGGGTG |
| Human | BAZ1B | Other Histone Modifications | GCGGGCTTCATACTCTTCCC |
| Human | BAZ1B | Other Histone Modifications | AGTATGAAGCCCGCTTGGAA |
| Human | BAZ1B | Other Histone Modifications | GCCTTGAGTGCAGATAAGGG |
| Human | BAZ1B | Other Histone Modifications | GAAGCCTTGAGTGCAGATAA |
| Human | BAZ1B | Other Histone Modifications | GGATGACCAACACCCTGTTA |
| Human | BAZ1B | Other Histone Modifications | GCCACCCTTATCTGCACTCA |
| Human | BAZ1B | Other Histone Modifications | CTGGTCACAGGCTCCCTGAA |
| Human | BAZ1B | Other Histone Modifications | TGCAGAACAAATGTTCCTGT |
| Human | BAZ1B | Other Histone Modifications | GCAGTTGTAAACCTCAGCAT |
| Human | BAZ1B | Other Histone Modifications | GCACTGTCTGAAAGTCCATG |
| Human | BAZ1B | Other Histone Modifications | AGTTTGTCGAAAGAAAGGTG |
| Human | BAZ1B | Other Histone Modifications | GCCCTCTATGAAGTACCAGA |
| Human | BAZ1B | Other Histone Modifications | TCTGGTACTTCATAGAGGGC |
| Human | BAZ1B | Other Histone Modifications | ACCATCTGGTACTTCATAGA |
| Human | BAZ1B | Other Histone Modifications | ATGAAGTACCAGATGGTGAG |
| Human | BAZ1B | Other Histone Modifications | GTGTGACTTCGAGGTTGGGA |
| Human | BAZ1B | Other Histone Modifications | AATTCATCTTCTACGACCCA |
| Human | BAZ1B | Other Histone Modifications | AGTTTCACATCATATTTGTG |
| Human | BAZ1B | Other Histone Modifications | CGGCATAATGCATTACGAGC |
| Human | BAZ1B | Other Histone Modifications | CTTCTATGTCTGAAGAACAA |
| Human | BAZ1B | Other Histone Modifications | TTCTTTCTCTCTTCGTTCTT |
| Human | BAZ1B | Other Histone Modifications | GATGTCTGCAGAGTTGTGGA |
| Human | BAZ1B | Other Histone Modifications | GGAACGGCTTGCTGTGTTGA |
| Human | BAZ1B | Other Histone Modifications | AGCAGATGTCTGCAGAGTTG |
| Human | BAZ1B | Other Histone Modifications | CTGCAGAGTTGTGGAAGGAA |
| Human | BAZ1B | Other Histone Modifications | TAAGAAGGAACGGGAAATCC |
| Human | BAZ1B | Other Histone Modifications | GCTGAGAAAGCTTTCCAGGA |
| Human | BAZ1B | Other Histone Modifications | GTATTCGTTCTTCTTCAGCT |
| Human | BAZ1B | Other Histone Modifications | AAGAGAAATGAAAGTGAAAC |
| Human | BAZ1B | Other Histone Modifications | AAGCTGAAGAAGAACGAATA |
| Human | BAZ1B | Other Histone Modifications | GGAGGAAGAAGATTATGAGG |
| Human | BAZ1B | Other Histone Modifications | AGATGATGAGAGTGATGAAG |
| Human | BAZ1B | Other Histone Modifications | TGATGAGAGTGATGAAGAGG |
| Human | BAZ1B | Other Histone Modifications | GAAGAAGATTATGAGGTGGC |
| Human | BAZ1B | Other Histone Modifications | GGAGGAGGAAGAAGATTATG |

| | | | |
|---|---|---|---|
| Human | BAZ1B | Other Histone Modifications | GCCCACCTGCCAAGAAACCA |
| Human | BAZ1B | Other Histone Modifications | TTTCTTTGTAGTATGCAATG |
| Human | SUV39H1 | Histone Methylation | GGATCTTCTTGTAATCGCAC |
| Human | SUV39H1 | Histone Methylation | CCAGGTAATATTCCTGTTCG |
| Human | SUV39H1 | Histone Methylation | ACACACTTGAGATTCTGCCG |
| Human | SUV39H1 | Histone Methylation | GATATCCACGCCATTTCACC |
| Human | SUV39H1 | Histone Methylation | TCCAGACTCAGAGAGCACCT |
| Human | SUV39H1 | Histone Methylation | AAGTTTGCCTACAATGACCA |
| Human | SUV39H1 | Histone Methylation | CGAGTGCCAGGACTGTCTGT |
| Human | SUV39H1 | Histone Methylation | CTATGACTGCCCAAATCGTG |
| Human | SUV39H1 | Histone Methylation | TGGGTGCCCACAGACAGTCC |
| Human | SUV39H1 | Histone Methylation | GGGAGAGATCATTACCTCAG |
| Human | SUV39H1 | Histone Methylation | AGCTTCGTCATGGAGTACGT |
| Human | SUV39H1 | Histone Methylation | CTGCCCTCGGTATCTCTAAG |
| Human | SUV39H1 | Histone Methylation | TCTTAGAGATACCGAGGGCA |
| Human | SUV39H1 | Histone Methylation | ACTCGACTTCAAAGTCATAG |
| Human | SUV39H1 | Histone Methylation | GGTTCCTCTTAGAGATACCG |
| Human | SUV39H1 | Histone Methylation | CTTTGATTACAACATGCAAG |
| Human | SUV39H1 | Histone Methylation | CACTTGCATGTTGTAATCAA |
| Human | SUV39H1 | Histone Methylation | TAGAAAGGGAGCTGCTCCGG |
| Human | PRDM7 | Histone Methylation | ATGGGCAGAAATGGGAGACT |
| Human | PRDM7 | Histone Methylation | ACACATGAAAGCTGGTCGAG |
| Human | PRDM7 | Histone Methylation | CATTTCCATATACTTCACCA |
| Human | PRDM7 | Histone Methylation | CTTCGGTGACACATGAAAGC |
| Human | PRDM7 | Histone Methylation | ATTCCTGGCTAATCACCAAG |
| Human | PRDM7 | Histone Methylation | GTTCACATACCTCATCCAGT |
| Human | PRDM7 | Histone Methylation | TATTCCTGGCTAATCACCAA |
| Human | PRDM7 | Histone Methylation | ATATTCCTGGCTAATCACCA |
| Human | PRMT1 | Histone Methylation | CATACACGTTCTCCCACCAG |
| Human | PRMT1 | Histone Methylation | AGCCAACAAGTTAGACCACG |
| Human | PRMT1 | Histone Methylation | CCTTGATGATGGTCACCACG |
| Human | PRMT1 | Histone Methylation | ACTCGATCCCGATGACCTTG |
| Human | PRMT1 | Histone Methylation | AAGACTACAAGATCCACTGG |
| Human | MPHOSPH8 | Chromatin Readers | CTGGACATGAAGACCGAGGG |
| Human | MPHOSPH8 | Chromatin Readers | ATCCTGGACATGAAGACCGA |
| Human | MPHOSPH8 | Chromatin Readers | TCCTGGACATGAAGACCGAG |
| Human | MPHOSPH8 | Chromatin Readers | ATACATCGGATGATGATACC |
| Human | MPHOSPH8 | Chromatin Readers | GAGGTCGTCCTGCCCTCCGG |
| Human | MPHOSPH8 | Chromatin Readers | CTGCGACTCCTCATCACAAA |
| Human | MPHOSPH8 | Chromatin Readers | TCTTCTCTGCAGCATGAATG |
| Human | MPHOSPH8 | Chromatin Readers | CTCTGCAGCATGAATGAGGG |
| Human | MPHOSPH8 | Chromatin Readers | TGCAGCATGAATGAGGGCGG |

| | | | |
|---|---|---|---|
| Human | MPHOSPH8 | Chromatin Readers | TAGACTGCTTCGCAAAGTGC |
| Human | MPHOSPH8 | Chromatin Readers | AGACTGCTTCGCAAAGTGCA |
| Human | MPHOSPH8 | Chromatin Readers | TGAAGCCAAAGAAGAACTAA |
| Human | MPHOSPH8 | Chromatin Readers | TCCAAGGAGCTCTCCAGGTC |
| Human | DDOST | Control | AGGACCCAGCACGTGCACAC |
| Human | DDOST | Control | ACTTCCGGTGTGCACGTGCT |
| Human | DDOST | Control | GTGCTGGGTCCTTCGGCAGG |
| Human | DDOST | Control | CTTCCACTCAGGTATCCGTG |
| Human | DDOST | Control | TATCACCCAGTATCCACATG |
| Human | DDOST | Control | TGGCCGCACGGATACCTGAG |
| Human | DDOST | Control | TGGCCAGCATACGCTCATCG |
| Human | DDOST | Control | ACCTTCCTGAAGAAGAAAGG |
| Human | DDOST | Control | GCGCCTTCTCCATGATGCTG |
| Human | DDOST | Control | AGCGCCTTCTCCATGATGCT |
| Human | DDOST | Control | CAGCATCATGGAGAAGGCGC |
| Human | DDOST | Control | CAGCGCCTTCTCCATGATGC |
| Human | SETDB2 | Histone Methylation | AAGGAGTCTACGAAACGTGG |
| Human | SETDB2 | Histone Methylation | AGTTACACTCTGTCTCAAGC |
| Human | SETDB2 | Histone Methylation | CTGCATTCATAAATGCCAGT |
| Human | SETDB2 | Histone Methylation | TAATCCCAAGGAGCTTACTG |
| Human | SETDB2 | Histone Methylation | CATTCACTAGAATCATTGCT |
| Human | ARID1B | Chromatin Remodeling | CTGGCAACCAACCTAAACGT |
| Human | ARID1B | Chromatin Remodeling | GTCTGCGTCAAAGAGATCGG |
| Human | ARID1B | Chromatin Remodeling | CTTTGACGCAGACGTAGAGT |
| Human | ARID1B | Chromatin Remodeling | CGTCTGCGTCAAAGAGATCG |
| Human | ARID1B | Chromatin Remodeling | TAGAGTCGGAACAGGTCCAG |
| Human | ARID1B | Chromatin Remodeling | GCCCGTACATGCCGTCCATA |
| Human | ARID1B | Chromatin Remodeling | GCCATATGGACGGCATGTAC |
| Human | ARID1B | Chromatin Remodeling | CGCCATATGGACGGCATGTA |
| Human | ARID1B | Chromatin Remodeling | GGCACCCACAGCGCCAAGAG |
| Human | ARID1B | Chromatin Remodeling | CTCCGAGGCGGCTCTCAAGG |
| Human | ARID1B | Chromatin Remodeling | TCCGAGGCGGCTCTCAAGGA |
| Human | ARID1B | Chromatin Remodeling | AGCCGCCGCTCTTGGCGCTG |
| Human | ARID1B | Chromatin Remodeling | GGAGGAGGAGGACAGCGCGG |
| Human | ARID1B | Chromatin Remodeling | GCATCCTCTTCCTCCTCGTC |
| Human | ARID1B | Chromatin Remodeling | GGCATCCTCTTCCTCCTCGT |
| Human | ARID1B | Chromatin Remodeling | CTCCTCCGCGGCGGCAGCGG |
| Human | ARID1B | Chromatin Remodeling | CTCCTCCTCCGCGGCGGCAG |
| Human | ARID1B | Chromatin Remodeling | GGATGCCGCCGCTGCCGCCG |
| Human | ARID1B | Chromatin Remodeling | GGTGGAGGTGGTGGGCATGG |
| Human | ARID1B | Chromatin Remodeling | GGTGGTGGGCATGGTGGTGG |
| Human | ARID1B | Chromatin Remodeling | GGCATGGTGGTGGTGGTGGT |

| | | | |
|---|---|---|---|
| Human | ARID1B | Chromatin Remodeling | GGAGGTGGTGGGCATGGTGG |
| Human | ARID1B | Chromatin Remodeling | GGTGGTGGTGGTGGGCATGG |
| Human | ARID1B | Chromatin Remodeling | GCTGCTGCTGCTGCTGCTGC |
| Human | ARID1B | Chromatin Remodeling | GCTGCTGCTGCTGCTGGAAC |
| Human | ARID1B | Chromatin Remodeling | GTAATTATTAAACTCCGGGA |
| Human | ARID1B | Chromatin Remodeling | CCCTGCAGAACTCCCACGAA |
| Human | ARID1B | Chromatin Remodeling | CCGCCATGTTGATCAAAGCA |
| Human | ARID1B | Chromatin Remodeling | CCTTGCTTTGATCAACATGG |
| Human | ARID1B | Chromatin Remodeling | CCCATGATGCGGAGCTACGG |
| Human | ARID1B | Chromatin Remodeling | CCAGAACCAGCACCCGTCGG |
| Human | ARID1B | Chromatin Remodeling | CGCCGTAGCTCCGCATCATG |
| Human | ARID1B | Chromatin Remodeling | GCCAGAACCAGCACCCGTCG |
| Human | ARID1B | Chromatin Remodeling | GAGGTGGCATCTGCTGCTGA |
| Human | ARID1B | Chromatin Remodeling | GTGAGTGGTTACTGCCAGCA |
| Human | ARID1B | Chromatin Remodeling | TGAGGTGGCATCTGCTGCTG |
| Human | ARID1B | Chromatin Remodeling | GACGGCAGATACTGCGCCTG |
| Human | ARID1B | Chromatin Remodeling | CTGCGGCTGGTACCTCTGCT |
| Human | ARID1B | Chromatin Remodeling | GCGGAGGCATCTGACTACCT |
| Human | ARID1B | Chromatin Remodeling | CGGGTCCACGAGCAGCCAAG |
| Human | ARID1B | Chromatin Remodeling | CCGGGTCCACGAGCAGCCAA |
| Human | ARID1B | Chromatin Remodeling | GGAAGCAACCAGTCTCGATC |
| Human | ARID1B | Chromatin Remodeling | CATCTCTCCAGCATCCCGGG |
| Human | ARID1B | Chromatin Remodeling | GTACCATCATATCGTCTGTG |
| Human | ARID1B | Chromatin Remodeling | ATATGATGGTACCCGATCAG |
| Human | ARID1B | Chromatin Remodeling | ACTGCCAAATCACATCTCCA |
| Human | ARID1B | Chromatin Remodeling | ATGTGATTTGGCAGTGACGG |
| Human | ARID1B | Chromatin Remodeling | GGATAAATCATGAGAGCCAG |
| Human | KDM2B | Histone Methylation | TCCACCAGGTCTACCACAGT |
| Human | KDM2B | Histone Methylation | CGGCATGGATCCAACCGGAA |
| Human | KDM2B | Histone Methylation | ACGGCATGGATCCAACCGGA |
| Human | KDM2B | Histone Methylation | TCCGACTGTGGTAGACCTGG |
| Human | KDM2B | Histone Methylation | CTTGGTCAAGCGTCCGACTG |
| Human | KDM2B | Histone Methylation | ACCGCTTATCGCAGCACCAG |
| Human | KDM2B | Histone Methylation | CACCTCCCTGTGCATGACGT |
| Human | KDM2B | Histone Methylation | GGCAGCCCACGTCATGCACA |
| Human | KDM2B | Histone Methylation | ACCGCTGGTGCTGCGATAAG |
| Human | KDM2B | Histone Methylation | GGTGCTGCGATAAGCGGTTG |
| Human | KDM2B | Histone Methylation | TGATAGACTTGCAGTGGTTC |
| Human | KDM2B | Histone Methylation | TGCAGTGGTTCAGGTCAATG |
| Human | KDM2B | Histone Methylation | GCAGTGGTTCAGGTCAATGC |
| Human | KDM2B | Histone Methylation | TCAACCGGCTGCCTGGGCTC |
| Human | KDM2B | Histone Methylation | GGCTCATCAACCGGCTGCCT |

| | | | |
|---|---|---|---|
| Human | KDM2B | Histone Methylation | AGCTGAGCTGGCTCATCAAC |
| Human | KDM2B | Histone Methylation | CAGCTGCTTCTTGGAGATAT |
| Human | KDM2B | Histone Methylation | TCTCCAAGAAGCAGCTGAGC |
| Human | KDM2B | Histone Methylation | CTGCTCATGGATCGCGGTCT |
| Human | KDM2B | Histone Methylation | ACATCCAGGGTCCGGAGCAG |
| Human | KDM2B | Histone Methylation | GCGGACAACTGGAGCTGCAA |
| Human | KDM2B | Histone Methylation | CGGACAACTGGAGCTGCAAA |
| Human | KDM2B | Histone Methylation | GCTCCAGTTGTCCGCTGCTC |
| Human | KDM2B | Histone Methylation | GTGGCGGATGATGAGCCGCA |
| Human | KDM2B | Histone Methylation | GCGGATGATGAGCCGCAGGG |
| Human | KDM2B | Histone Methylation | ACATCACAGATGCCTCCCTG |
| Human | KDM2B | Histone Methylation | TGTGGCGGATGATGAGCCGC |
| Human | KDM2B | Histone Methylation | CTCGGTTAAGGAGTCTCGGG |
| Human | KDM2B | Histone Methylation | TGATAGACTGGTCGGTGACG |
| Human | KDM2B | Histone Methylation | GATCTCGGTTAAGGAGTCTC |
| Human | KDM2B | Histone Methylation | GAGCAGGTTGATAGACTGGT |
| Human | KDM2B | Histone Methylation | TGATCTCGGTTAAGGAGTCT |
| Human | KDM2B | Histone Methylation | CTGTCCTTCTTCAAACGCTG |
| Human | KDM2B | Histone Methylation | GTTTCCACAGCGTTTGAAGA |
| Human | KDM2B | Histone Methylation | CACAGCGTTTGAAGAAGGAC |
| Human | KDM2B | Histone Methylation | CTGGACACTCACAGACATCT |
| Human | KDM2B | Histone Methylation | TGTCTGTGAGTGTCCAGTTT |
| Human | KDM2B | Histone Methylation | ATGTCTGTGAGTGTCCAGTT |
| Human | KDM2B | Histone Methylation | TTTCTTCTACTTGCCCAAAC |
| Human | KDM2B | Histone Methylation | GAACTGCTCACAGCCTTCCT |
| Human | KDM2B | Histone Methylation | GAAGTGGCACTCTCCGCACT |
| Human | KDM2B | Histone Methylation | AGCTCTGCTTCATGCGCCCG |
| Human | KDM2B | Histone Methylation | GACGCGATGCCGCAAGTGCG |
| Human | KDM2B | Histone Methylation | TGAAGCAGAGCTGCATCATG |
| Human | KDM2B | Histone Methylation | CGGAGAGTGCCACTTCTGCA |
| Human | KDM2B | Histone Methylation | TAATCTTAAGGCATCCAGGG |
| Human | KDM2B | Histone Methylation | CTTAAGATTAAGGAGTCAGA |
| Human | KDM2B | Histone Methylation | CCTTAAGATTAAGGAGTCAG |
| Human | KDM2B | Histone Methylation | TGGAGCACTCCATGAGCATG |
| Human | KDM2B | Histone Methylation | CGTGTGCCTTGTGTGTGGCG |
| Human | KDM2B | Histone Methylation | AGCTCAACCACGAGATCCAG |
| Human | KDM2B | Histone Methylation | CAACCACGAGATCCAGAGGA |
| Human | KDM2B | Histone Methylation | TCTCCGTCCTCTGGATCTCG |
| Human | KDM2B | Histone Methylation | CCAGAGGACGGAGAACAGCC |
| Human | KDM2B | Histone Methylation | CCAGGCTGTTCTCCGTCCTC |
| Human | KDM2B | Histone Methylation | CTGTGATCAGCAGCCTCAGG |
| Human | KDM2B | Histone Methylation | TCCTTCTCCTCCTCCTCCTG |

| | | | |
|---|---|---|---|
| Human | KDM2B | Histone Methylation | TCAGCAGCCTCAGGAGGAGG |
| Human | KDM2B | Histone Methylation | GGCCTGTGATCAGCAGCCTC |
| Human | KDM2B | Histone Methylation | CTCCTGAGGCTGCTGATCAC |
| Human | KDM2B | Histone Methylation | CATCTGGATACACTTGGGCG |
| Human | KDM2B | Histone Methylation | TCCATCTGGATACACTTGGG |
| Human | KDM2B | Histone Methylation | CCCGCCCAAGTGTATCCAGA |
| Human | KDM2B | Histone Methylation | CGCTCCATCTGGATACACTT |
| Human | KDM1A | Histone Methylation | CCAACCACAACTGCAGTGTG |
| Human | KDM1A | Histone Methylation | CCGCACACTGCAGTTGTGGT |
| Human | KDM1A | Histone Methylation | AGCACAGTATCACTGTTATA |
| Human | KDM1A | Histone Methylation | GCTCGACAGTTACAAAGTTT |
| Human | KDM1A | Histone Methylation | GAGAGATATACATCACTTGG |
| Human | KDM1A | Histone Methylation | GAGTCGTGGAATCGGCTGTG |
| Human | KDM1A | Histone Methylation | ATTTGGCAACCTTAACAAGG |
| Human | KDM1A | Histone Methylation | TGAGAGATATACATCACTTG |
| Human | KDM1A | Histone Methylation | ACCTCAGCCCAAAGAAACTG |
| Human | KDM1A | Histone Methylation | GAAGGTAGAGTACAGAGAGA |
| Human | KDM1A | Histone Methylation | AATAATACTCATCTTCTGAG |
| Human | KDM1A | Histone Methylation | CAGAGAGATGGATGAAAGCT |
| Human | KDM1A | Histone Methylation | ATACTCATCTTCTGAGAGGT |
| Human | KDM1A | Histone Methylation | CTACCTTCGCCCGCTTGCGC |
| Human | KDM1A | Histone Methylation | CTCCGAGAACGGGTCTGAGG |
| Human | KDM1A | Histone Methylation | CGGCTCCGAGAACGGGTCTG |
| Human | KDM1A | Histone Methylation | AGCGGCAGCAACCGGGACGG |
| Human | KDM1A | Histone Methylation | ACAGCAGGCGGCTCCGAGAA |
| Human | KDM1A | Histone Methylation | TGCAGCGGCAGCAACCGGGA |
| Human | TERF2 | DNA and RNA Modifications | GAAAGCGAGTGGGTCAAGGC |
| Human | TERF2 | DNA and RNA Modifications | GGCTGGAGTGCAGAAATATG |
| Human | TERF2 | DNA and RNA Modifications | AGAAGAAAGCGAGTGGGTCA |
| Human | TERF2 | DNA and RNA Modifications | CGAGCCCTACCTCCTCACGA |
| Human | TERF2 | DNA and RNA Modifications | TTCTCAGCTTCTGAGTTGTG |
| Human | TERF2 | DNA and RNA Modifications | AGGCGGGCGTCCCGCAGTAG |
| Human | TERF2 | DNA and RNA Modifications | CCGGGCCCGCCCGCTACTGC |
| Human | TERF2 | DNA and RNA Modifications | GGGCGTCCCGCAGTAGCGGG |
| Human | TERF2 | DNA and RNA Modifications | GCCGGGCCCGCCCGCTACTG |
| Human | TERF2 | DNA and RNA Modifications | GGCGTCCCGCAGTAGCGGGC |
| Human | CBX7 | Chromatin Readers | TCCCACGTGCTGTACTTTGG |
| Human | CBX7 | Chromatin Readers | CATGGCCTACGAGGAGAAGG |
| Human | CBX7 | Chromatin Readers | GCATCCGGAAGAAGCGCGTG |
| Human | CBX7 | Chromatin Readers | GGTGACGGTGATGGAGTTGG |
| Human | KDM5B | Histone Methylation | TCGCTTTCATCCACAAGATC |
| Human | KDM5B | Histone Methylation | ATCTTGTGGATGAAAGCGAA |

| | | | |
|---|---|---|---|
| Human | KDM5B | Histone Methylation | CAGTCTGCTCGGCTATGGGC |
| Human | KDM5B | Histone Methylation | AGGTGCGGCCGCCGCCGGAT |
| Human | KDM5B | Histone Methylation | ATGCCAGTCTGCTCGGCTAT |
| Human | KDM5B | Histone Methylation | CAGTGGGCTCACATATCAGA |
| Human | KDM5B | Histone Methylation | GCAAAGTACTGGGAGTTACA |
| Human | KDM5B | Histone Methylation | AATAAGTTAGTTGCAGAAGA |
| Human | KDM5B | Histone Methylation | TCAATTAACTACTTGCACTG |
| Human | KDM5B | Histone Methylation | TTTCCAAGAGCCTACCACAG |
| Human | KDM5B | Histone Methylation | TATCAGCAGTAATATGTGCA |
| Human | KDM5B | Histone Methylation | GCAGTGGCAATGATGAAGAC |
| Human | KDM5B | Histone Methylation | GACCGGCTACTGTTGTGTGA |
| Human | KDM5B | Histone Methylation | ATGTTCCCAAGGGAGACTGG |
| Human | KDM5B | Histone Methylation | AGCCATCACACAACAGTAGC |
| Human | KDM5B | Histone Methylation | CCCTTGGGAACATCATGGAG |
| Human | KDM5B | Histone Methylation | ATTGCGGTATAGGTACACGC |
| Human | KDM5B | Histone Methylation | CTCCTTACAAATATAAATTG |
| Human | KDM5B | Histone Methylation | CATGATGCAGGCAAACAAGA |
| Human | KDM5B | Histone Methylation | ACAAATATAAATTGCGGTAT |
| Human | KDM5B | Histone Methylation | TTTATATTTGTAAGGAGGAC |
| Human | KDM5B | Histone Methylation | GCGGTACCCAGTATTTCACA |
| Human | KDM5B | Histone Methylation | TTTCCACACCAGTTGTGTGG |
| Human | KDM5B | Histone Methylation | GTACCGCCACACAACTGGTG |
| Human | KDM5B | Histone Methylation | GGCGGTACCCAGTATTTCAC |
| Human | KDM5B | Histone Methylation | GCAGCCAGAAGGAGATGAGG |
| Human | KDM5B | Histone Methylation | AGAAGGAGATGAGGTGGACT |
| Human | KDM5B | Histone Methylation | GATTGCAGCTGCCATCACAC |
| Human | KDM5B | Histone Methylation | CAGAAGGAGATGAGGTGGAC |
| Human | KDM5B | Histone Methylation | TGGTTTCATCAGGTCTGTGT |
| Human | KDM5B | Histone Methylation | AGCTTGTTGAGAAAGAATTT |
| Human | KDM5B | Histone Methylation | GATAATGCTCCACACACTGT |
| Human | KDM5B | Histone Methylation | AGACATGGCCATTATGATTG |
| Human | KDM5B | Histone Methylation | TTCAACTGTTCAGAAAGACA |
| Human | KDM5B | Histone Methylation | TTTCTCATCCTCAATCATAA |
| Human | KDM5B | Histone Methylation | AGCTTCAATGCATTCATCAT |
| Human | KDM5B | Histone Methylation | GCTTCAATGCATTCATCATA |
| Human | OIP5 | Chromatin Remodeling | TGTAACTCTGGAGAAGACCA |
| Human | OIP5 | Chromatin Remodeling | AGAATACAGATGGAAACCAA |
| Human | SFN | Chromatin Readers | TCTGATCCAGAAGGCCAAGC |
| Human | SFN | Chromatin Readers | CTTGGCCTTCTGGATCAGAC |
| Human | SFN | Chromatin Readers | GACATGGCAGCCTTCATGAA |
| Human | SFN | Chromatin Readers | GCCTATAAGAACGTGGTGGG |
| Human | SFN | Chromatin Readers | GTAGCCTATAAGAACGTGGT |

| | | | |
|---|---|---|---|
| Human | SFN | Chromatin Readers | AGTAGCCTATAAGAACGTGG |
| Human | SFN | Chromatin Readers | CTCAGTAGCCTATAAGAACG |
| Human | SFN | Chromatin Readers | TCTTATAGGCTACTGAGAGC |
| Human | SFN | Chromatin Readers | AAGGTGGAGACTGAGCTCCA |
| Human | SFN | Chromatin Readers | GCGTGAGTACCGGGAGAAGG |
| Human | SFN | Chromatin Readers | GCACGGTGTCGCACACGCCC |
| Human | SFN | Chromatin Readers | GGGCGTGTGCGACACCGTGC |
| Human | SFN | Chromatin Readers | GCTCAGTCTCCACCTTCTCC |
| Human | SFN | Chromatin Readers | GGGTGACTACTACCGCTACC |
| Human | SFN | Chromatin Readers | TCATCTTCAGGTAGAAGACC |
| Human | SFN | Chromatin Readers | GGTCTTCTACCTGAAGATGA |
| Human | SFN | Chromatin Readers | AGTAGTCACCCTTCATCTTC |
| Human | SFN | Chromatin Readers | GTCTTCTACCTGAAGATGAA |
| Human | SFN | Chromatin Readers | AGCGCATCATTGACTCAGCC |
| Human | SFN | Chromatin Readers | CTCAGCCCGGTCAGCCTACC |
| Human | SFN | Chromatin Readers | GTCAGCCTACCAGGAGGCCA |
| Human | SFN | Chromatin Readers | GGCCTCCTGGTAGGCTGACC |
| Human | SFN | Chromatin Readers | AGTTCAGGGCCAGGCCCAGG |
| Human | SFN | Chromatin Readers | GGCCTCGTCGAAAGTGGTCT |
| Human | SFN | Chromatin Readers | AGTGGTCTTGGCCAGAGAGA |
| Human | SFN | Chromatin Readers | TCGCAGCAGCTGCATGATGA |
| Human | SFN | Chromatin Readers | GATGAGGGTGCTGTCTTTGT |
| Human | SFN | Chromatin Readers | CTCGCAGCAGCTGCATGATG |
| Human | JARID2 | Histone Methylation | TCCGCTCATCTACATCGAGT |
| Human | JARID2 | Histone Methylation | CGATGTAGATGAGCGGATCG |
| Human | JARID2 | Histone Methylation | CGAACTTCTCCACCTGAGCG |
| Human | JARID2 | Histone Methylation | CGGCTGATTAATGAGATGGG |
| Human | JARID2 | Histone Methylation | AGTCGTAGGAGAGTAGGTAC |
| Human | JARID2 | Histone Methylation | GACGAGCTCCCGCTCATAGG |
| Human | JARID2 | Histone Methylation | GGACGAGCTCCCGCTCATAG |
| Human | JARID2 | Histone Methylation | TCACTCCAGGCACAGCACCG |
| Human | JARID2 | Histone Methylation | AAGCAACGTCATGATCTCCC |
| Human | JARID2 | Histone Methylation | ACCACCCAGTGGACAAGTAT |
| Human | METTL14 | DNA and RNA Modifications | AGAAATAGATCTAACTTCCG |
| Human | METTL14 | DNA and RNA Modifications | GAAAGAGGTGGCTTTAGAGG |
| Human | METTL14 | DNA and RNA Modifications | TCTAACTTCCGAGGAGAAAG |
| Human | METTL14 | DNA and RNA Modifications | TAAAGCCACCTCTTTCTCCT |
| Human | METTL14 | DNA and RNA Modifications | AACTTCCGAGGAGAAAGAGG |
| Human | METTL14 | DNA and RNA Modifications | GAGGGAGCTCATCAGGCTAA |
| Human | METTL14 | DNA and RNA Modifications | ACCTCAAATGAGCAAGTCAA |
| Human | METTL14 | DNA and RNA Modifications | ATTGGATACTTTAAGAAACA |
| Human | KAT2B | Histone Acetylation | GCTCAAGAGCATCCTCCAGC |

| | | | |
|---|---|---|---|
| Human | KAT2B | Histone Acetylation | TACTGAAGAAGAATTTCTCC |
| Human | KAT2B | Histone Acetylation | CTACAAATGTGCCAATATCC |
| Human | KAT2B | Histone Acetylation | GAAGAATTTCTCCAGGATAT |
| Human | KAT2A | Histone Acetylation | TGTGCTTCGGGTCAAAGACG |
| Human | KAT2A | Histone Acetylation | ACCTCGAATGAGCAGGTCAA |
| Human | KAT2A | Histone Acetylation | GCTTCGGGTCAAAGACGAGG |
| Human | KAT2A | Histone Acetylation | TCATGGTCTTCAGGTCAATG |
| Human | KAT2A | Histone Acetylation | GCTTCCGGGTCACGTAGTAG |
| Human | KAT2A | Histone Acetylation | GATGACCTCGTAGTAGTCAG |
| Human | KAT2A | Histone Acetylation | GAAGTAGAAGAACTTCTCCA |
| Human | KAT2A | Histone Acetylation | GAAGAACTTCTCCAGGGCGC |
| Human | KAT2A | Histone Acetylation | TGAAGTAGAAGAACTTCTCC |
| Human | KAT2A | Histone Acetylation | TCTCCAGGGCGCTGGCACAG |
| Human | KAT2A | Histone Acetylation | CTGCCGCTGTGCCAGCGCCC |
| Human | CLOCK | Histone Acetylation | TCTAGCATTACCAGGAAGCA |
| Human | CLOCK | Histone Acetylation | TCCATCTTTCTAGCATTACC |
| Human | CLOCK | Histone Acetylation | AATGTTCTCATTAAAGAACT |
| Human | CLOCK | Histone Acetylation | TCCTGGTAATGCTAGAAAGA |
| Human | CLOCK | Histone Acetylation | GAACTGGGATCCATGCTTCC |
| Human | CLOCK | Histone Acetylation | CTTTGATCCACAAGATCAGA |
| Human | CLOCK | Histone Acetylation | ACATTTACCATCTGATCTTG |
| Human | CLOCK | Histone Acetylation | TTAATGTTAGAGGCTCTTGA |
| Human | CLOCK | Histone Acetylation | GTTTCTGTTTCTAGATCACA |
| Human | CLOCK | Histone Acetylation | ACAGGGCACCACCCATAATA |
| Human | CLOCK | Histone Acetylation | CATCTAGACATAGTTTAGAA |
| Human | CLOCK | Histone Acetylation | TGAGTACAAACAATAAACTC |
| Human | CLOCK | Histone Acetylation | TATTATAGGTTCCTGACTAA |
| Human | CLOCK | Histone Acetylation | ATTATATCACTTACCATCAG |
| Human | CLOCK | Histone Acetylation | CTTACCATCAGTGGAATTCA |
| Human | CLOCK | Histone Acetylation | TGAAACTGGGACATGCCTTG |
| Human | CLOCK | Histone Acetylation | CGGATACGAGCACTCCACCC |
| Human | CLOCK | Histone Acetylation | TGTTGAGGAAGGGTCTGAGA |
| Human | FTO | DNA and RNA Modifications | ATGATGAATTCTATCAGCAG |
| Human | FTO | DNA and RNA Modifications | AGGAGAAGAAATTCATAATG |
| Human | FTO | DNA and RNA Modifications | TGGCTCAACTGGAAGCACTG |
| Human | FTO | DNA and RNA Modifications | AGTGCTTCCAGTTGAGCCAT |
| Human | FTO | DNA and RNA Modifications | GCACTGTGGAAGAAGATGGA |
| Human | FTO | DNA and RNA Modifications | AGCACTGTGGAAGAAGATGG |
| Human | FTO | DNA and RNA Modifications | CAGTGCTTCCAGTTGAGCCA |
| Human | FTO | DNA and RNA Modifications | TGAGGAGAGAATGGCATGCC |
| Human | FTO | DNA and RNA Modifications | TCACTGCACGCCAGAACCTG |
| Human | FTO | DNA and RNA Modifications | GGCGTGCAGTGAGCGAGGCA |

| | | | |
|---|---|---|---|
| Human | FTO | DNA and RNA Modifications | GTTCTGGCGTGCAGTGAGCG |
| Human | FTO | DNA and RNA Modifications | TGCAGTGAGCGAGGCAAGGA |
| Human | RNF168 | Transcription Elongation | CTTCCAGTCGACCGTCGAAA |
| Human | RNF168 | Transcription Elongation | ACAGCGTGTGGTTACACGGG |
| Human | RNF168 | Transcription Elongation | AGCACGGTTTACACAGCGTG |
| Human | RNF168 | Transcription Elongation | CAGCGTGTGGTTACACGGGA |
| Human | RNF168 | Transcription Elongation | ACTGAGAAGAGAATATGAAG |
| Human | RNF168 | Transcription Elongation | ATTACAACTTCAGAAGGAGG |
| Human | RNF168 | Transcription Elongation | GGCATTACAACTTCAGAAGG |
| Human | RNF168 | Transcription Elongation | ATTGGCATTACAACTTCAGA |
| Human | RNF168 | Transcription Elongation | AGAAGAACAGGACAGGTTAT |
| Human | RNF168 | Transcription Elongation | ATAAACAAGAAGAACAGGAC |
| Human | RNF168 | Transcription Elongation | TGAAGAGGAAATAAGCAAGG |
| Human | RNF168 | Transcription Elongation | AGAGGAAATAAGCAAGGTGG |
| Human | RNF168 | Transcription Elongation | CCAGTGAAGAATACATACAG |
| Human | RNF168 | Transcription Elongation | GGCAGAGCGACGGGCCAGCG |
| Human | RNF168 | Transcription Elongation | AGAATACATACAGAGGTTGT |
| Human | RNF168 | Transcription Elongation | GAGACATAAACAAGAAGAAC |
| Human | G6PD | Control | GCTGTTGAGGCGCTGGTAGG |
| Human | ESCO2 | DNA and RNA Modifications | CTGCATTTCATCTTCAGGGT |
| Human | ESCO2 | DNA and RNA Modifications | ACATGCTGCATTTCATCTTC |
| Human | ESCO2 | DNA and RNA Modifications | CATGCTGCATTTCATCTTCA |
| Human | ELL | Transcription Elongation | TGCTTCTTGGACTTCTTCTT |
| Human | ELL | Transcription Elongation | GTGCTTCTTGGACTTCTTCT |
| Human | ELL | Transcription Elongation | GCACCCGCTTCAGCAGCTGC |
| Human | SIRT4 | Histone Acetylation | TTGACTTTCAGGTCAGCAAA |
| Human | SIRT4 | Histone Acetylation | TGAGCTTTGCGTTGACTTTC |
| Human | SIRT4 | Histone Acetylation | GCTCCACGGATGCATGGACA |
| Human | ZNF326 | Transcription Elongation | TGAACTGAACTATGTAAAGC |
| Human | ZNF326 | Transcription Elongation | ATGGCCCTGGATCTTATGGA |
| Human | ZNF326 | Transcription Elongation | CCATCCCTCCATAAGATCCA |
| Human | ZNF326 | Transcription Elongation | TTCTTGATAAAGGTTCCACT |
| Human | ZNF326 | Transcription Elongation | CAGCCATTTAATAAGCCCAG |
| Human | ZNF326 | Transcription Elongation | GTTCCACTGGGCTTATTAAA |
| Human | ZNF326 | Transcription Elongation | TTTCTTGATAAAGGTTCCAC |
| Human | ZNF326 | Transcription Elongation | CATATGAAGCCTGCACCTGT |
| Human | ZNF326 | Transcription Elongation | CCTCTGTCCTCCATAAACCC |
| Human | ZNF326 | Transcription Elongation | CACCCATATGTCCTCGGCCT |
| Human | ZNF326 | Transcription Elongation | CCTGGGTTTATGGAGGACAG |
| Human | ZNF326 | Transcription Elongation | GAGGGCGAGGGAAATATACA |
| Human | ZNF326 | Transcription Elongation | GGGAAATATACAGGGAGTAG |
| Human | ZNF326 | Transcription Elongation | GAGGGAAATATACAGGGAGT |

| | | | |
|---|---|---|---|
| Human | ZNF326 | Transcription Elongation | TGAACCTATTGAAGAAGAGG |
| Human | ZNF326 | Transcription Elongation | TGCTCAACAGGGAAGTCAGC |
| Human | PHF13 | Chromatin Readers | GCACTCATTACACTCGATCA |
| Human | PHF13 | Chromatin Readers | AGTGTAATGAGTGCCACACC |
| Human | PHF13 | Chromatin Readers | AGAAGAAGAAGAGGAAGCGC |
| Human | PHF13 | Chromatin Readers | AGCTGAAGAAGAAGAAGAAG |
| Human | BRDT | Chromatin Readers | GGAGAATAAATATTATGCGA |
| Human | BRDT | Chromatin Readers | ATGGCCCTTTCAACGTCCTG |
| Human | BRDT | Chromatin Readers | TGTTATTTATATAACAAGCC |
| Human | BRDT | Chromatin Readers | AAATACAATTAAGAAGCGCT |
| Human | BRDT | Chromatin Readers | TTTATAATGGTATAATAATC |
| Human | BRDT | Chromatin Readers | TGACAACGTCATAGTAGTTA |
| Human | BRDT | Chromatin Readers | CCCAAAGCATTAACGTCAAC |
| Human | BRDT | Chromatin Readers | ACTCGCCCAAGTTTATCTCC |
| Human | BRDT | Chromatin Readers | CTCGCCCAAGTTTATCTCCA |
| Human | BRDT | Chromatin Readers | TATCTCTATCTCATCAGGAT |
| Human | BRDT | Chromatin Readers | AAACTCCCTGGAGATAAACT |
| Human | BRDT | Chromatin Readers | TTAGCTTACGGAAAGGTACT |
| Human | BRDT | Chromatin Readers | TAGCTTACGGAAAGGTACTT |
| Human | BRDT | Chromatin Readers | TAAGCGTCTTGCAAAGCTTC |
| Human | PHF21B | Chromatin Readers | GACGGCGCCCAAGGGCGTGT |
| Human | PHF21B | Chromatin Readers | CCCTTGGGCGCCGTCTTGAG |
| Human | PHF21B | Chromatin Readers | ACTGTGCCGCCTGCAAGCGA |
| Human | PHF21B | Chromatin Readers | AAGCAGAAGCTGCTGCAACG |
| Human | PHF21B | Chromatin Readers | GTCAGCAGTGCAGAAATGCC |
| Human | PHF21B | Chromatin Readers | GCACTGCTGACGCCAGCCGC |
| Human | PHF21B | Chromatin Readers | GAACGAGCACCAGCAGCTGG |
| Human | PHF21B | Chromatin Readers | AGCTGGAGGAGCGGGACCGG |
| Human | SKA1 | Chromatin Remodeling | AAGCGGGATTTCATGTACGA |
| Human | SKA1 | Chromatin Remodeling | GAAGAAACGAAGGATACCAA |
| Human | SKA1 | Chromatin Remodeling | GAGTTCCTTGAGTGAATTGT |
| Human | SKA1 | Chromatin Remodeling | ACAAACCAACAATTCACTCA |
| Human | SKA1 | Chromatin Remodeling | ACTTCTAAATAAATTGGAAT |
| Human | SKA1 | Chromatin Remodeling | AAATGAACTTCTAAATAAAT |
| Human | PRDM5 | Histone Methylation | AACTTGGCTGCCATTCAAGA |
| Human | PRDM5 | Histone Methylation | ACGGCCCGCAGAGTGCGAAA |
| Human | PRDM5 | Histone Methylation | GAAGCCAGTTGGAGTGCCGT |
| Human | PRDM5 | Histone Methylation | CTCAGCAAGAATATCCTCAC |
| Human | PRDM5 | Histone Methylation | TTTGCAAAGCCTGCTTAACT |
| Human | PRDM5 | Histone Methylation | GAAGAAATTCCCAGTTAAGC |
| Human | PRDM5 | Histone Methylation | CTTTGCAAAGCCTGCTTAAC |
| Human | PRDM5 | Histone Methylation | TGGAAAGAGGCTGAAGAGCA |

| | | | |
|---|---|---|---|
| Human | PRDM5 | Histone Methylation | AGGCTGACAGCTGTGGAAAG |
| Human | PRDM5 | Histone Methylation | TTCTTCAGCATCAAGCCTAC |
| Human | PRDM5 | Histone Methylation | TTCAAGAGGCTTGATCAAGT |
| Human | PRDM5 | Histone Methylation | TTTGTAATAAGTCTTTCAAG |
| Human | PRDM5 | Histone Methylation | TTTCAAGAGGCTTGATCAAG |
| Human | PRDM5 | Histone Methylation | ATTCTTGTAAACATTTCTGT |
| Human | PRDM5 | Histone Methylation | GATTCTTGTAAACATTTCTG |
| Human | PRDM5 | Histone Methylation | AAGAATGTAAAGCTTTGTTC |
| Human | PRDM5 | Histone Methylation | TAAAGGTAGCATCGCAGTGA |
| Human | PRDM5 | Histone Methylation | ACTGCGATGCTACCTTTAAG |
| Human | PRDM5 | Histone Methylation | CGATGCTACCTTTAAGAGGA |
| Human | PRDM5 | Histone Methylation | TACATTAAATGTTCATGTCC |
| Human | PRDM5 | Histone Methylation | TAATGTATCCTTCCTCTTAA |
| Human | PRDM5 | Histone Methylation | AAGCACTGAAGGTGTAACAA |
| Human | PRDM5 | Histone Methylation | TTATGACTTCTAAGCACTGA |
| Human | PRDM5 | Histone Methylation | GTACACTCAGAGTTCATATC |
| Human | PRDM5 | Histone Methylation | GGCCAGAAATTTGCCAGCAG |
| Human | PRDM5 | Histone Methylation | AACTCTGAGTGTACCACTGC |
| Human | PRDM5 | Histone Methylation | GCAAATTTCTGGCCACAATA |
| Human | PRDM5 | Histone Methylation | TACCACTGCTGGCAAATTTC |
| Human | PRDM5 | Histone Methylation | AAGGCCTTCAGCCAGAAGCG |
| Human | PRDM5 | Histone Methylation | CTTCAGCCAGAAGCGAGGCC |
| Human | PRDM5 | Histone Methylation | GCTCATCCAGGCCTCGCTTC |
| Human | PRDM5 | Histone Methylation | CAGGCCTCGCTTCTGGCTGA |
| Human | PRDM5 | Histone Methylation | TTTGCCATAAGAAGTTTACA |
| Human | PRDM5 | Histone Methylation | TGACTACCTCAAAGTGCACA |
| Human | PRDM5 | Histone Methylation | CATTCCTTGTAAACTTCTTA |
| Human | PRDM6 | Histone Methylation | TCTCTGGGAGATATATGACC |
| Human | PRDM6 | Histone Methylation | AGCCTGTATAGATATCCCTA |
| Human | PRDM6 | Histone Methylation | GCATCTGTAAGAGGATCCGC |
| Human | PRDM6 | Histone Methylation | CATCTGTAAGAGGATCCGCT |
| Human | PRDM6 | Histone Methylation | TTCAGAACTGAGGAACCACG |
| Human | PRDM6 | Histone Methylation | ACGTGGTTCCTCAGTTCTGA |
| Human | PRDM6 | Histone Methylation | GTTCCTCAGTTCTGAAGGCT |
| Human | PRDM6 | Histone Methylation | GGTTCCTCAGTTCTGAAGGC |
| Human | PRDM6 | Histone Methylation | CTGTGGTCGTGCCTTTGCCG |
| Human | PRDM6 | Histone Methylation | ACTGTGGTCGTGCCTTTGCC |
| Human | PRDM6 | Histone Methylation | GATGTGGTTGTTGAGGGTGG |
| Human | PRDM6 | Histone Methylation | GTTGTTGAGGGTGGTGGCCC |
| Human | PRDM6 | Histone Methylation | CCCAGCTGAGCCGACACCAG |
| Human | PRDM6 | Histone Methylation | GTGTGAGAGGAGCTTCACGC |
| Human | PRDM6 | Histone Methylation | CTGGTGTCGGCTCAGCTGGG |

| | | | |
|---|---|---|---|
| Human | PRDM6 | Histone Methylation | GGAGGCTCAGCGCGCTCCGG |
| Human | PRDM6 | Histone Methylation | CGGAGGCTCAGCGCGCTCCG |
| Human | PRDM6 | Histone Methylation | GGAGCGCGCTGAGCCTCCGC |
| Human | PRDM6 | Histone Methylation | GGCGGAGGCTCAGCGCGCTC |
| Human | PRDM6 | Histone Methylation | GGGCGCAGGCTGTCCGGCGG |
| Human | NCBP1 | Transcription Elongation | TACCAATCTCGTCGCACCTG |
| Human | NCBP1 | Transcription Elongation | CAATCTAAATACTCTTCTTG |
| Human | NCBP1 | Transcription Elongation | GCAGTCCATACCTGTAACAT |
| Human | NCBP1 | Transcription Elongation | AAGCTAAAGAGAAACTTGCT |
| Human | NCBP1 | Transcription Elongation | AAGCAGTGACAGGAAAGACG |
| Human | NCBP1 | Transcription Elongation | AACTTGCTAGGCAACACAAA |
| Human | NCBP1 | Transcription Elongation | GAAGCAGTGACAGGAAAGAC |
| Human | NCBP1 | Transcription Elongation | ACAGCGACGAGAACGACGGT |
| Human | NCBP1 | Transcription Elongation | CACAGCGACGAGAACGACGG |
| Human | NCBP1 | Transcription Elongation | ACGGTGGGCAGCCTCACAAA |
| Human | NCBP1 | Transcription Elongation | ATTTCTGAATCTGGGCCCAC |
| Human | NCBP1 | Transcription Elongation | CAAAGATCACCTCAACTATG |
| Human | NCBP1 | Transcription Elongation | CAGTTGCCTGTGCAAGAACT |
| Human | NCBP1 | Transcription Elongation | AGTTGCCTGTGCAAGAACTT |
| Human | NCBP1 | Transcription Elongation | GTTGCCTGTGCAAGAACTTG |
| Human | NCBP1 | Transcription Elongation | ATTAATAAACCTGTCTACAC |
| Human | NCBP1 | Transcription Elongation | GTGTAGACAGGTTTATTAAT |
| Human | NCBP1 | Transcription Elongation | AGAACTTCTCTTACAAACTT |
| Human | NCBP1 | Transcription Elongation | TAAACCGCTGAAATATAACG |
| Human | NCBP1 | Transcription Elongation | GGTATAAGAACTGTATAGAG |
| Human | MCM2 | DNA and RNA Modifications | ATTTGACAAGATGAATGACC |
| Human | MCM2 | DNA and RNA Modifications | GGTCGTAGCGCCCTCCTATG |
| Human | MCM2 | DNA and RNA Modifications | AAACCCAGGTGGCAAGCACA |
| Human | MCM2 | DNA and RNA Modifications | CAGAACCAGGAGGTGAAACC |
| Human | MCM2 | DNA and RNA Modifications | TTTCTGCCAGTCCCAGAACC |
| Human | MCM2 | DNA and RNA Modifications | TGGTTCTGGGACTGGCAGAA |
| Human | MCM2 | DNA and RNA Modifications | CTGCCAGTCCCAGAACCAGG |
| Human | MCM2 | DNA and RNA Modifications | AGGAGCCTGGTTTCACCTCC |
| Human | MCM2 | DNA and RNA Modifications | GCTCCTGTATGACAGCGATG |
| Human | MCM2 | DNA and RNA Modifications | CCTGTATGACAGCGATGAGG |
| Human | MCM2 | DNA and RNA Modifications | CTGGACGCCTATGAGGCCGA |
| Human | MCM2 | DNA and RNA Modifications | ATCATCCAGAGCCAGTCCCT |
| Human | MCM2 | DNA and RNA Modifications | GAGAACCTGGAGGATCTCAA |
| Human | GTF2A2 | Transcription Elongation | CAAACTGAAGTAGAACTTGA |
| Human | GTF2A2 | Transcription Elongation | AAACTGAAGTAGAACTTGAA |
| Human | GTF2A2 | Transcription Elongation | AGTTCTACTTCAGTTTGATA |
| Human | UBR5 | Other Histone Modifications | ATCGCTAGGATTCCCTTCAG |

| | | | |
|---|---|---|---|
| Human | UBR5 | Other Histone Modifications | GCTTCTCCTTCTAGCAAGTG |
| Human | UBR5 | Other Histone Modifications | GGTTGCATTGCTTGTACACG |
| Human | UBR5 | Other Histone Modifications | CTGAGCCTTTGCCAGCACAT |
| Human | UBR5 | Other Histone Modifications | AACAGCCCTTACATGCAATG |
| Human | UBR5 | Other Histone Modifications | CATCTGAACTCTGAGACGCG |
| Human | UBR5 | Other Histone Modifications | TCTGTTTAGAGGAATAGAGT |
| Human | UBR5 | Other Histone Modifications | ACATACTCATATACATTCTG |
| Human | UBR5 | Other Histone Modifications | CCCTTACATGCAATGAGGAA |
| Human | UBR5 | Other Histone Modifications | TGGAGCAGAGCACATTAACC |
| Human | UBR5 | Other Histone Modifications | GTTGTTGTACGGAATGTGCA |
| Human | UBR5 | Other Histone Modifications | GTCATGATTGCAAACTCAAA |
| Human | UBR5 | Other Histone Modifications | AGATGTGGCTGATCAGCCAA |
| Human | UBR5 | Other Histone Modifications | GATGTGGCTGATCAGCCAAG |
| Human | UBR5 | Other Histone Modifications | AAATCCATTTGAAGTGCCTC |
| Human | UBR5 | Other Histone Modifications | AGAGGCACTTCAAATGGATT |
| Human | UBR5 | Other Histone Modifications | GAACCGAGGAGAGAGAGACC |
| Human | UBR5 | Other Histone Modifications | GAGGAGAGAGAGACCGGGAA |
| Human | UBR5 | Other Histone Modifications | TTTCCCGGTCTCTCTCTCCT |
| Human | UBR5 | Other Histone Modifications | GGAACCGAGGAGAGAGAGAC |
| Human | UBR5 | Other Histone Modifications | AGGAGAGAGAGACCGGGAAA |
| Human | BANP | Chromatin Remodeling | CGGATGCCGTAGATGGTGAG |
| Human | BANP | Chromatin Remodeling | AGGCTGTGTCCAACCTCTCG |
| Human | BANP | Chromatin Remodeling | CTGCTTGATTCGGTACCAGT |
| Human | BANP | Chromatin Remodeling | GCGCCATCTTCTCGGCCGTG |
| Human | BANP | Chromatin Remodeling | CGCAGCGTACCCGCATCTCG |
| Human | BANP | Chromatin Remodeling | CCAAGAAGACAGCCACCACG |
| Human | BANP | Chromatin Remodeling | GGGCAGTAGGAGGACGAGTT |
| Human | BANP | Chromatin Remodeling | AGGGCAGTAGGAGGACGAGT |
| Human | BANP | Chromatin Remodeling | TGCTCATCATCGGCTCTGAA |
| Human | BANP | Chromatin Remodeling | TGCGGGAGCTCGCTGGCAGG |
| Human | BANP | Chromatin Remodeling | CAGGTGCAGATCCACCAGAT |
| Human | BANP | Chromatin Remodeling | GAGCTCGCTGGCAGGAGGTG |
| Human | BANP | Chromatin Remodeling | GGCTGCGGGAGCTCGCTGGC |
| Human | BANP | Chromatin Remodeling | AGCCAAATTGCAAGCCCTGG |
| Human | BANP | Chromatin Remodeling | AGGATTTACAAGTAGCCTCC |
| Human | BANP | Chromatin Remodeling | GGATTTACAAGTAGCCTCCA |
| Human | BANP | Chromatin Remodeling | TGAAGCCAAATTGCAAGCCC |
| Human | BANP | Chromatin Remodeling | GAAGACGGACAGGTGCAAGT |
| Human | BANP | Chromatin Remodeling | CAGATCACGCAGGACAGCGA |
| Human | BANP | Chromatin Remodeling | AGGACACCTCCACATCGCCC |
| Human | BANP | Chromatin Remodeling | CATCGCCCAGGTGCCGCAGG |
| Human | BANP | Chromatin Remodeling | CACATCGCCCAGGTGCCGCA |

| | | | |
|---|---|---|---|
| Human | SETD6 | Histone Methylation | AAAGCTATAGGCCATCACAA |
| Human | SETD6 | Histone Methylation | GAAAGCTATAGGCCATCACA |
| Human | SETD6 | Histone Methylation | CTCTGCGAATTGTCTTCGGA |
| Human | SETD6 | Histone Methylation | TGCAGCAGGCACCATCACGG |
| Human | SETD6 | Histone Methylation | GCACCCGATGTTCTGGCCAG |
| Human | SETD6 | Histone Methylation | CCTGCTTCCTGAGCTGGTGC |
| Human | SETD6 | Histone Methylation | CTGTGGCCTGCTTCCTGAGC |
| Human | SETD6 | Histone Methylation | CAGCTCAGGAAGCAGGCCAC |
| Human | SETD6 | Histone Methylation | GCACCTAGTGTACGAGCGCT |
| Human | SETD6 | Histone Methylation | GGCACCTAGTGTACGAGCGC |
| Human | SETD6 | Histone Methylation | AATCCCAGCGCTCGTACACT |
| Human | SETD6 | Histone Methylation | GGAGCTGACCACCACACTAA |
| Human | SETD6 | Histone Methylation | CAAGCCTTACAGGTTCGCTA |
| Human | SETD6 | Histone Methylation | CTGACCATAGCGAACCTGTA |
| Human | NDC80 | Chromatin Remodeling | GTCTGTAGAGAAACATCTTG |
| Human | NDC80 | Chromatin Remodeling | CTTGAGCCCTGTATTTGACA |
| Human | NDC80 | Chromatin Remodeling | TGAAATTAAGTTTAATCCCG |
| Human | NDC80 | Chromatin Remodeling | CTAGAAAGTACTGTTAACCA |
| Human | NDC80 | Chromatin Remodeling | ATGCATGTCAGAAGATCTCT |
| Human | KAT6B | Histone Acetylation | GAAAGACGGACCGCAGTACA |
| Human | KAT6B | Histone Acetylation | AATGGGAGGTTACTGAAAGA |
| Human | KAT6B | Histone Acetylation | TGAAAGACGGACCGCAGTAC |
| Human | KAT6B | Histone Acetylation | GAAGGACTTGAGGAGCCGAA |
| Human | KAT6B | Histone Acetylation | GAGAGCAATTGAAGGACTTG |
| Human | KAT6B | Histone Acetylation | AGAGTGGTGGCAATGTCATG |
| Human | KAT6B | Histone Acetylation | TACCAAATTCAATCACAGAA |
| Human | KAT6B | Histone Acetylation | CATCAACCTCAAATACTGAA |
| Human | KAT6B | Histone Acetylation | CTCTCTTGTCGATCATGTGG |
| Human | KAT6B | Histone Acetylation | AGAGCTTGGCTAACAAGCAA |
| Human | KAT6B | Histone Acetylation | TGCCACAATCTGCACAAGAG |
| Human | KAT6B | Histone Acetylation | TGAATTAACAACAAATGTAA |
| Human | KAT6B | Histone Acetylation | CAAATGTAAAGGCCTTAAGG |
| Human | KAT6B | Histone Acetylation | CATATGAAATCCTCTATCAC |
| Human | KAT6B | Histone Acetylation | TGCAGTGCCTGTAGAGTCCA |
| Human | KAT6B | Histone Acetylation | TGGCAAATCCACATCCCTTT |
| Human | KAT6B | Histone Acetylation | TTTGGCATTCTGGAAAGTGG |
| Human | KAT6B | Histone Acetylation | CCTGATGGTCGCAGATCACG |
| Human | KAT6B | Histone Acetylation | AGCTACGATTCTACTATGGG |
| Human | KAT6B | Histone Acetylation | GCTCGACGATTGCCAACAGT |
| Human | KAT6B | Histone Acetylation | ATAGATTTGTGACTGATGGG |
| Human | KAT6B | Histone Acetylation | CTGTTGGCAATCGTCGAGCG |
| Human | KAT6B | Histone Acetylation | CGGACTTTAACGATGCAAAG |

| | | | |
|---|---|---|---|
| Human | KAT6B | Histone Acetylation | AGAGGAAGAGGATGAAGAGG |
| Human | KAT6B | Histone Acetylation | GGATGAAGAGGAGGAAGAAG |
| Human | KAT6B | Histone Acetylation | AGAAGAGGAAGAGGATGAAG |
| Human | KAT6B | Histone Acetylation | GGAAGAAGAGGAGGAAGAAG |
| Human | KAT6B | Histone Acetylation | CGAGGAGGAGGAAGAAGAGG |
| Human | KAT6B | Histone Acetylation | AAGAAGAGGAAGAAGAGGAA |
| Human | KAT6B | Histone Acetylation | GAAGAAGAGGAAGAAGAGGA |
| Human | KAT6B | Histone Acetylation | AGAGGAAGAAGAGGAAGAAG |
| Human | KAT6B | Histone Acetylation | AGAAGAGGAAGAAGAGGAAG |
| Human | KAT6B | Histone Acetylation | GGAGGAGGAAGAGGAAGAAG |
| Human | MCM6 | DNA and RNA Modifications | CCAATTTCTCAAGCACGTGG |
| Human | MCM6 | DNA and RNA Modifications | GAGCCAATTTCTCAAGCACG |
| Human | MCM6 | DNA and RNA Modifications | AATTTGATAAGATGGACGTG |
| Human | MCM6 | DNA and RNA Modifications | TATTGTGCTTCACCTCAGAA |
| Human | MCM6 | DNA and RNA Modifications | TGTGCTTCACCTCAGAAAGG |
| Human | MCM6 | DNA and RNA Modifications | GCACAATAAGGTTAGAGATT |
| Human | MCM6 | DNA and RNA Modifications | TTCTGAGGTGAAGCACAATA |
| Human | ERCC2 | Transcription Elongation | GCAGGATTGAGTATCGAGCA |
| Human | ERCC2 | Transcription Elongation | AAACGCACGCTGGACGCCAA |
| Human | ERCC2 | Transcription Elongation | CCAGAGAGCATATCCGCTGG |
| Human | ERCC2 | Transcription Elongation | ATACCAGAGAGCATATCCGC |
| Human | ERCC2 | Transcription Elongation | GTGCACAAAGTCGATTCCCT |
| Human | ERCC2 | Transcription Elongation | TGGCCTCCTGGTATGAGCAG |
| Human | ERCC2 | Transcription Elongation | CTTTGTGCACCACTACGGGC |
| Human | ERCC2 | Transcription Elongation | ACTTTGTGCACCACTACGGG |
| Human | DCTN2 | Control | CAACTTCAGTTGTCAGCTCT |
| Human | DCTN2 | Control | GTACCAGCGCCTACTGCATG |
| Human | DCTN2 | Control | GCTCTTGGACCTCATGCAGT |
| Human | DCTN2 | Control | GGACCTCATGCAGTAGGCGC |
| Human | DCTN2 | Control | CAGCTGTCTCCAGCTCTGTC |
| Human | DCTN2 | Control | ACTTTGCCAGCATTGATGAA |
| Human | SMNDC1 | Chromatin Readers | AGATGGACAGTGTTATGAAG |
| Human | SUV39H2 | Histone Methylation | GTGTGACTACAAGGTAGTAA |
| Human | SUV39H2 | Histone Methylation | GGAATACTTGTGTGACTACA |
| Human | SUV39H2 | Histone Methylation | GGCCAGATTCTACAAATACT |
| Human | SUV39H2 | Histone Methylation | AGAATTGCTGAAGCAGTAAC |
| Human | SUV39H2 | Histone Methylation | CATTCATAGATGGGAGTACC |
| Human | SUV39H2 | Histone Methylation | TGCAACTCAAGGTGTCAGTG |
| Human | SUV39H2 | Histone Methylation | TTTCGAACTAGCAATGGACG |
| Human | SUV39H2 | Histone Methylation | TGAGTCTGATGAATTCACAG |
| Human | SUV39H2 | Histone Methylation | TGTGGAGCTGTGACTTGCAG |
| Human | SUV39H2 | Histone Methylation | AAAGGATAGCTCTGCAGAGA |

| | | | |
|---|---|---|---|
| Human | SKA2 | Chromatin Remodeling | ATCTGGATTACATTCAATAC |
| Human | SKA2 | Chromatin Remodeling | GGATTACATTCAATACAGGC |
| Human | SKA2 | Chromatin Remodeling | CCAGAAAGCTGAGTCTGATC |
| Human | SKA2 | Chromatin Remodeling | CCAGATCAGACTCAGCTTTC |
| Human | SKA2 | Chromatin Remodeling | CAACAGCAACTGGTTTAAAG |
| Human | SKA2 | Chromatin Remodeling | AACAGCAACTGGTTTAAAGC |
| Human | SKA2 | Chromatin Remodeling | TCTTTCTGCTCAACAGCAAC |
| Human | DNMT3L | DNA and RNA Modifications | CAGGAACTTGTCCTTACATG |
| Human | DNMT3L | DNA and RNA Modifications | ACACTCGAAGCAGTAGCATC |
| Human | DNMT3L | DNA and RNA Modifications | GGTACCTGTTCCAGTTCCAC |
| Human | DNMT3L | DNA and RNA Modifications | GGAGCCGGTGGAACTGGAAC |
| Human | SOX10 | Transcription Elongation | GCAGCACAAGAAAGACCACC |
| Human | SOX10 | Transcription Elongation | TGCCTGAGCCCACACCATGA |
| Human | SOX10 | Transcription Elongation | GGTCTTTCTTGTGCTGCATA |
| Human | DMAP1 | DNA and RNA Modifications | GCAGTTCAAGAAGCGTTCTG |
| Human | DMAP1 | DNA and RNA Modifications | AACGCTTCTTGAACTGCTGG |
| Human | DMAP1 | DNA and RNA Modifications | CTGTGGAAGACCTGAAGGAG |
| Human | DMAP1 | DNA and RNA Modifications | AGCTGCGCAAGATTGAGGCC |
| Human | DMAP1 | DNA and RNA Modifications | CTCCTCTGCCACCTGCTCTG |
| Human | DMAP1 | DNA and RNA Modifications | CCCAGAGCAGGTGGCAGAGG |
| Human | DMAP1 | DNA and RNA Modifications | TGTGGTAGTACCGCTCCTTC |
| Human | SETD3 | Histone Methylation | TCACTTACGAGGACTACAGG |
| Human | SETD3 | Histone Methylation | GTATGTGCAGATCCGGACTC |
| Human | SETD3 | Histone Methylation | GGGAAGAGTATGTGCAGATC |
| Human | SETD3 | Histone Methylation | ATGGCCATCAAATTGCGCTT |
| Human | SETD3 | Histone Methylation | AAAGAGTGCAGCTGTCAACC |
| Human | SETD3 | Histone Methylation | CTCACCTAAGCGCAATTTGA |
| Human | SETD3 | Histone Methylation | TAAAGAGTGCAGCTGTCAAC |
| Human | SETD3 | Histone Methylation | CTTAGGTGAGAAAGAGATTT |
| Human | EZH2 | Histone Methylation | CGGCTCCACAAGTAAGACAG |
| Human | EZH2 | Histone Methylation | ATTGCTGGCACCATCTGACG |
| Human | EZH2 | Histone Methylation | GTTAACGGTGATCACAGGAT |
| Human | EZH2 | Histone Methylation | TGATGGTTAACGGTGATCAC |
| Human | EZH2 | Histone Methylation | TGTGGTGGATGCAACCCGCA |
| Human | EZH2 | Histone Methylation | CAGTGTTACCAGCATTTGGA |
| Human | EZH2 | Histone Methylation | ACAGTGTTACCAGCATTTGG |
| Human | EZH2 | Histone Methylation | ACACCGAGAATTTGCTTCAG |
| Human | EZH2 | Histone Methylation | CTCCACTCCACATTCTCAGG |
| Human | EAF2 | Transcription Elongation | TGAAGTCATAGCGCACAGTG |
| Human | EAF2 | Transcription Elongation | ATAACTCTGCCAAATATAGA |
| Human | EAF2 | Transcription Elongation | ACCATGATACTGGAGAATGT |
| Human | EAF2 | Transcription Elongation | TTCTTCTGAAGGATACCTTG |

| | | | |
|---|---|---|---|
| Human | EAF2 | Transcription Elongation | GTCTCGAAATCTATTATGAC |
| Human | EAF2 | Transcription Elongation | AATCCTCTACTTCTGATACA |
| Human | EAF2 | Transcription Elongation | ATCCTGTACTGTGTCATGGT |
| Human | SIRT6 | Histone Acetylation | TACGTCCGAGACACAGTCGT |
| Human | SIRT6 | Histone Acetylation | ATAGAGCCGTTGATCCGGGT |
| Human | SIRT6 | Histone Acetylation | TGGTGGGCCGCTCCCGTTTG |
| Human | SIRT6 | Histone Acetylation | CCAAGGAGGAATCTCCCACC |
| Human | SIRT6 | Histone Acetylation | GGAAGGTGTGGGAACTGGCG |
| Human | SIRT6 | Histone Acetylation | GGAGCTGGAGCGGAAGGTGT |
| Human | SIRT6 | Histone Acetylation | AGGAGCTGGAGCGGAAGGTG |
| Human | SIRT6 | Histone Acetylation | GGTGTGGGAACTGGCGAGGC |
| Human | SIRT6 | Histone Acetylation | GGAGCGGAAGGTGTGGGAAC |
| Human | POLE3 | Chromatin Remodeling | AAGCTCTGGAAGCATATAGG |
| Human | POLE3 | Chromatin Remodeling | AGCTCTGGAAGCATATAGGC |
| Human | POLE3 | Chromatin Remodeling | AGAAGAAGAACAGAATGAAG |
| Human | POLE3 | Chromatin Remodeling | GGAAGAGCAAGACAAGAGCA |
| Human | POLE3 | Chromatin Remodeling | GCATATAGGCGGGAGCAGAA |
| Human | BRWD1 | Chromatin Readers | AAGATTTGGTCAACACATAA |
| Human | BRWD1 | Chromatin Readers | TGTGTAGCATTTGATAGGAC |
| Human | BRWD1 | Chromatin Readers | TACAGGTTCAGATGACTGTT |
| Human | BRWD1 | Chromatin Readers | ACAGGACATAGAATCTTTAC |
| Human | BRWD1 | Chromatin Readers | TTTACTGTGTAGCATTTGAT |
| Human | BRWD1 | Chromatin Readers | TGAGAATACAATGATTGCTG |
| Human | BRWD1 | Chromatin Readers | TTCTGCAGAAATTTCAGATA |
| Human | BRWD1 | Chromatin Readers | GAGAATACAATGATTGCTGC |
| Human | BRWD1 | Chromatin Readers | TGGTTTCCACTGGTGCTGAT |
| Human | BRWD1 | Chromatin Readers | CAGTTTAGCCCGATGGCCAA |
| Human | BRWD1 | Chromatin Readers | ATCTTTACAGTTTAGCCCGA |
| Human | BRWD1 | Chromatin Readers | CAAAGGCTCTCAAAGATACA |
| Human | BRWD1 | Chromatin Readers | AACTGTCCCATCAGCACCAG |
| Human | BRWD1 | Chromatin Readers | TACATGATCAGTACTACCTG |
| Human | BRWD1 | Chromatin Readers | GAACAAAGCATTTGAACGCC |
| Human | BRWD1 | Chromatin Readers | AATGGTGATCGGTTCCTAAG |
| Human | BRWD1 | Chromatin Readers | GATTTGAGCAGTTAGAATGG |
| Human | BRWD1 | Chromatin Readers | GTAGCAGAGATGGAACAGCA |
| Human | BRWD1 | Chromatin Readers | GAGATGGAACAGCACGGATT |
| Human | BRWD1 | Chromatin Readers | AAAGTGTGGAATTCTTACAC |
| Human | BRWD1 | Chromatin Readers | TGTAAGAATTCCACACTTTG |
| Human | BRWD1 | Chromatin Readers | ATGATCATGTCCTCAAAGTG |
| Human | BRWD1 | Chromatin Readers | CTAAAGTAACAATGATAGCT |
| Human | BRWD1 | Chromatin Readers | CAAGCTATCATTGTTACTTT |
| Human | BRWD1 | Chromatin Readers | AACATAATTCTGGAATCAAA |

| | | | |
|---|---|---|---|
| Human | BRWD1 | Chromatin Readers | TGATGGCAGCATATTTATAT |
| Human | BRWD1 | Chromatin Readers | TCCAGAATTATGTTATCTGC |
| Human | BRWD1 | Chromatin Readers | ATGTTATCTGCAGGACATGA |
| Human | BRWD1 | Chromatin Readers | TCCTGCAGATAACATAATTC |
| Human | BRWD1 | Chromatin Readers | TTGCCTGTACAGATTCTCAT |
| Human | BRWD1 | Chromatin Readers | GTGCCCATGAGAATCTGTAC |
| Human | BRWD1 | Chromatin Readers | CGAATTGTGTAAAGATCGGT |
| Human | BRWD1 | Chromatin Readers | CATTCGAATTGTGTAAAGAT |
| Human | BRWD1 | Chromatin Readers | GAGGCTGTCTGCGTTAGTTT |
| Human | BRWD1 | Chromatin Readers | GTAAGGGAAACTCTAGATGC |
| Human | BRWD1 | Chromatin Readers | TCTATAATATCTCTGTAGTC |
| Human | BRWD1 | Chromatin Readers | GGATATTCAACCAAATCAAC |
| Human | BRWD1 | Chromatin Readers | AAATAAGGCAGATAATCTCA |
| Human | TAF11 | Transcription Elongation | GGAGGTGGTAGAAGAAGCAC |
| Human | TAF11 | Transcription Elongation | ATATGAGGGAAGCCGTTAGA |
| Human | SETD7 | Histone Methylation | CTTCGTAAGTGTAAACTCCC |
| Human | SETD7 | Histone Methylation | GGAGTTTACACTTACGAAGA |
| Human | SETD7 | Histone Methylation | GAGTTTACACTTACGAAGAT |
| Human | SETD7 | Histone Methylation | GATGATGCCTTGCAGGGCCA |
| Human | SETD7 | Histone Methylation | GTAAACTCCCTGGCCCTGCA |
| Human | SETD7 | Histone Methylation | TGCTGGATATATTACCCAGA |
| Human | SETD7 | Histone Methylation | TATAAAGATAACATTCGTCA |
| Human | SETD7 | Histone Methylation | ACATTCGTCATGGAGTGTGC |
| Human | SETD7 | Histone Methylation | ACACCAAGAGGTTGACAGCA |
| Human | SETD7 | Histone Methylation | CACACCAAGAGGTTGACAGC |
| Human | SETD7 | Histone Methylation | GGAAGGCCTTCAGCTCCACC |
| Human | TDRD5 | Chromatin Readers | CGATACCCAAGGATTCGGAG |
| Human | TDRD5 | Chromatin Readers | CAAAGATGGTTTGAGCCCAC |
| Human | TDRD5 | Chromatin Readers | CCATAGTGGACCGATACCCA |
| Human | TDRD5 | Chromatin Readers | GTATCGGTCCACTATGGAGC |
| Human | TDRD5 | Chromatin Readers | TTACAGTACCACCTGCACCG |
| Human | TDRD5 | Chromatin Readers | GAACAGGAGATAACGCCAAC |
| Human | TDRD5 | Chromatin Readers | AACTCACTCTTCACAACAGC |
| Human | TDRD5 | Chromatin Readers | CACAACAGCTGGAAGAATAG |
| Human | TDRD5 | Chromatin Readers | CATCAACAAAGAACCTGCTC |
| Human | TDRD5 | Chromatin Readers | TCATTCCAATACATGCAATA |
| Human | TDRD5 | Chromatin Readers | TGTTGTATCTAAGTTCCCAG |
| Human | TDRD5 | Chromatin Readers | GTCCTTTCCTGAACTCAACA |
| Human | TDRD5 | Chromatin Readers | TTAGAAATAAACAAACCCTC |
| Human | TDRD5 | Chromatin Readers | CTCCATGTTGAGTTCAGGAA |
| Human | TDRD5 | Chromatin Readers | GCAGCACTTGAGGAACCTCA |
| Human | TDRD5 | Chromatin Readers | GAAGTGTTCTACCCAGACTT |

| | | | |
|---|---|---|---|
| Human | TDRD5 | Chromatin Readers | CTGTTGTGTAAGGATTTCTG |
| Human | TDRD5 | Chromatin Readers | CGGGACATCTCTGTTGTGTA |
| Human | TDRD5 | Chromatin Readers | AGCAGCACTTGAGGAACCTC |
| Human | TDRD5 | Chromatin Readers | AGCGTATACAGGAATGTCTG |
| Human | TDRD5 | Chromatin Readers | TATACAGGAATGTCTGCGGA |
| Human | TRIM28 | Chromatin Readers | TTCTCCCTGGACCAGCCCGG |
| Human | TRIM28 | Chromatin Readers | CAACAAGTTAACTGAGGACA |
| Human | TRIM28 | Chromatin Readers | GTACACGCTCACATTTCCGC |
| Human | TRIM28 | Chromatin Readers | CTTCGAGACGCGCATGAACG |
| Human | TRIM28 | Chromatin Readers | GGGCTGGTCCAGGGAGAAGG |
| Human | TRIM28 | Chromatin Readers | GGCGGGCGACGGCACCGTGG |
| Human | TRIM28 | Chromatin Readers | TAGTGCCTGCTTAGGGCCCG |
| Human | TRIM28 | Chromatin Readers | CTACAGGCCGAGTGCAAACA |
| Human | TRIM28 | Chromatin Readers | AGAGCGCCTGCGACCCGAGA |
| Human | TRIM28 | Chromatin Readers | CCTGTGTAGAGGCGCACCAG |
| Human | TRIM28 | Chromatin Readers | CCCAGCCACCAGCTACTGTG |
| Human | TRIM28 | Chromatin Readers | GCACTCCACACAGTAGCTGG |
| Human | TRIM28 | Chromatin Readers | TCTACACAGGTCTCACACAG |
| Human | TRIM28 | Chromatin Readers | CTGCCAGCTCAATGCCCACA |
| Human | TRIM28 | Chromatin Readers | GGGCATTGAGCTGGCAGTCT |
| Human | TRIM28 | Chromatin Readers | GGCCCTGCAGGATGTACCAG |
| Human | TRIM28 | Chromatin Readers | CTTCTGGCAGACACGGCAAA |
| Human | TRIM28 | Chromatin Readers | ATGCCCAGAAGGTGACTGAG |
| Human | TRIM28 | Chromatin Readers | GATGCCCAGAAGGTGACTGA |
| Human | TRIM28 | Chromatin Readers | GCTGGTCAATGATGCCCAGA |
| Human | TRIM28 | Chromatin Readers | TGATGCCCAGAAGGTGACTG |
| Human | TRIM28 | Chromatin Readers | CCTCTAAGAACTGGTACTGG |
| Human | TRIM28 | Chromatin Readers | CCTGGGAAGACCTTGAAGAC |
| Human | TRIM28 | Chromatin Readers | ACGAGTGAGCCTTGAACGCC |
| Human | TRIM28 | Chromatin Readers | GGTGAGCGGCCTTATGCGCA |
| Human | TRIM28 | Chromatin Readers | GGGAAGACCTTGAAGACGGG |
| Human | TRIM28 | Chromatin Readers | GCGGCGTCGTCGCCCGCGGG |
| Human | TRIM28 | Chromatin Readers | GGCCGAGGCTGCGGCCGAAG |
| Human | TRIM28 | Chromatin Readers | AGCCGCGGCGTCGTCGCCCG |
| Human | TRIM33 | Chromatin Readers | CTTGAAGATCAAACGGACAT |
| Human | TRIM33 | Chromatin Readers | TTTGGTATCGAAGCAGGAAC |
| Human | TRIM33 | Chromatin Readers | ATCCGGGATTTGGTAGTGTT |
| Human | TRIM33 | Chromatin Readers | TTCCCAACACTACCAAATCC |
| Human | TRIM33 | Chromatin Readers | TAGTAGTTTGGTATCGAAGC |
| Human | TRIM33 | Chromatin Readers | CTGCAAGCTCTGCTGACACA |
| Human | TRIM33 | Chromatin Readers | GAGCTTGCAGAGCCGGCGTG |
| Human | TRIM33 | Chromatin Readers | AGGCAGAAGGAGTGAAGACA |

| | | | |
|---|---|---|---|
| Human | TRIM33 | Chromatin Readers | GCTTGGGCTCCGCCTCACGC |
| Human | TRIM33 | Chromatin Readers | CAGGCAGAAGGAGTGAAGAC |
| Human | TRIM33 | Chromatin Readers | TTTGTGTAGAATGTGGAGAG |
| Human | TRIM33 | Chromatin Readers | TTACTAAAGATCACTTGATC |
| Human | TRIM33 | Chromatin Readers | GAAGACAATGCAAGTGCAGT |
| Human | TRIM33 | Chromatin Readers | TATTGGAACACAAAGAACAT |
| Human | TRIM33 | Chromatin Readers | AACTGTTCTTGTTTGTGTAC |
| Human | TRIM33 | Chromatin Readers | ATGTAGAGACTGTCAGCTAT |
| Human | TRIM33 | Chromatin Readers | ACTGTTCTTGTTTGTGTACA |
| Human | TRIM33 | Chromatin Readers | ACACTACTTAGCTTTCCAAG |
| Human | TRIM33 | Chromatin Readers | ACTACTTAGCTTTCCAAGTG |
| Human | TRIM33 | Chromatin Readers | CACTACTTAGCTTTCCAAGT |
| Human | TRIM33 | Chromatin Readers | GCTCTGCTGACACACGGCGC |
| Human | TRIM33 | Chromatin Readers | ACAGCTAGAGAATGTTACAA |
| Human | TRIM33 | Chromatin Readers | CAGCTACTCAGGTGCAGAAT |
| Human | TRIM33 | Chromatin Readers | GGGTGCAATTGAGAATCTAC |
| Human | TRIM33 | Chromatin Readers | GAAGACGTTCACATTTCCTT |
| Human | SSRP1 | DNA and RNA Modifications | CACAGCATGTATGCAGACAT |
| Human | SSRP1 | DNA and RNA Modifications | CATCACGGATCTTTCCAAGA |
| Human | SSRP1 | DNA and RNA Modifications | GAAAGAGGAGTGGGATCGCA |
| Human | SSRP1 | DNA and RNA Modifications | AAGATCAAGTCAGACCATCC |
| Human | SSRP1 | DNA and RNA Modifications | CAAAGAGAAGAAAGAGGAGT |
| Human | SSRP1 | DNA and RNA Modifications | ACCGATGAGTCATTCAACCC |
| Human | SSRP1 | DNA and RNA Modifications | AGCATGATGCCTACTTGGAG |
| Human | SSRP1 | DNA and RNA Modifications | TCTCCAAGTAGGCATCATGC |
| Human | SSRP1 | DNA and RNA Modifications | GGAGGAAGGCAAGATCCGGG |
| Human | SSRP1 | DNA and RNA Modifications | ACCTGGGTTGAATGACTCAT |
| Human | SSRP1 | DNA and RNA Modifications | GCCAGCTCCTCCAGTAATGA |
| Human | SSRP1 | DNA and RNA Modifications | TGCCAGCTCCTCCAGTAATG |
| Human | SSRP1 | DNA and RNA Modifications | ACCCTCATTACTGGAGGAGC |
| Human | SSRP1 | DNA and RNA Modifications | ATTACTGGAGGAGCTGGCAG |
| Human | SSRP1 | DNA and RNA Modifications | CTTGCGGTCCTTGGCCATCT |
| Human | SSRP1 | DNA and RNA Modifications | GAAGAAAGTAAAGGTAAAGA |
| Human | SSRP1 | DNA and RNA Modifications | AAAGAAGAAGAAGAAAGTAA |
| Human | SSRP1 | DNA and RNA Modifications | GCTCTCGCTTAGCTGCCTTG |
| Human | SSRP1 | DNA and RNA Modifications | CATCATCCAAGTCGTCCTCA |
| Human | SSRP1 | DNA and RNA Modifications | GCTGAGTCCTCTGAGCTGGG |
| Human | SSRP1 | DNA and RNA Modifications | CGACTGAGGTTGAGCCGTCA |
| Human | SSRP1 | DNA and RNA Modifications | TCGACTGAGGTTGAGCCGTC |
| Human | SSRP1 | DNA and RNA Modifications | GATGTTCTTTGTGATCAGCC |
| Human | SSRP1 | DNA and RNA Modifications | GGCTGATCACAAAGAACATC |
| Human | SIRT1 | Histone Acetylation | TCTGGTTTCATGATAGCAAG |

| | | | |
|---|---|---|---|
| Human | SIRT1 | Histone Acetylation | ATAGCCTTGTCAGATAAGGA |
| Human | SIRT1 | Histone Acetylation | AAAGTATATGGACCTATCCG |
| Human | SIRT1 | Histone Acetylation | AGTGTCATATCATCCAACTC |
| Human | SIRT1 | Histone Acetylation | GTCATATCATCCAACTCAGG |
| Human | SIRT1 | Histone Acetylation | AGTTGGATGATATGACACTG |
| Human | SIRT1 | Histone Acetylation | GAGTCTTCTGAAACATGAAG |
| Human | SIRT1 | Histone Acetylation | GAAACATGAAGAGGTGTGGG |
| Human | SIRT1 | Histone Acetylation | TCTGAAACATGAAGAGGTGT |
| Human | SIRT1 | Histone Acetylation | TTCTGAAACATGAAGAGGTG |
| Human | CENPB | Chromatin Remodeling | GAAGTCACGGATCATCCAGG |
| Human | CENPB | Chromatin Remodeling | GAGAATCCGGACCTGCGCAA |
| Human | CENPB | Chromatin Remodeling | GGAGAAGTCACGGATCATCC |
| Human | CENPB | Chromatin Remodeling | GAGGCGACAGCTGACGTTCC |
| Human | CENPB | Chromatin Remodeling | TGACGTTCCGGGAGAAGTCA |
| Human | CENPB | Chromatin Remodeling | GTCTCCCTACGACAAGCTCG |
| Human | CENPB | Chromatin Remodeling | CAAGCCCTCGAGCTTGTCGT |
| Human | CENPB | Chromatin Remodeling | TTGAGGATGATGCCCTTGAC |
| Human | CENPB | Chromatin Remodeling | ACGATCTGGTTCATGTGACC |
| Human | CENPB | Chromatin Remodeling | GAACCAGATCGTGTTCCAAG |
| Human | CENPB | Chromatin Remodeling | CCAGATCGTGTTCCAAGTGG |
| Human | CENPB | Chromatin Remodeling | TGTTGATAGTGATGAAGAAG |
| Human | CENPB | Chromatin Remodeling | TGATAGTGATGAAGAAGAGG |
| Human | CENPB | Chromatin Remodeling | GAAGAGGAGGTGGAGGAGGA |
| Human | CENPB | Chromatin Remodeling | AGACGACGATGATGATGAGG |
| Human | CENPB | Chromatin Remodeling | TGAAGACGACGATGATGATG |
| Human | CENPB | Chromatin Remodeling | GACGATGATGATGAGGAGGA |
| Human | CENPB | Chromatin Remodeling | TGATTCAGACAGTGAGGAAG |
| Human | CENPB | Chromatin Remodeling | CAGTGAGGAAGAGGACGATG |
| Human | PYGO1 | Chromatin Readers | TCGTTTGTACAAATTCCACA |
| Human | PYGO1 | Chromatin Readers | TGGTTTCATCGGATCTGTAC |
| Human | PYGO1 | Chromatin Readers | CTTGTCAGAAATGGTTTCAT |
| Human | PYGO1 | Chromatin Readers | CAGATGCTTCTGCAGTTAAG |
| Human | PYGO1 | Chromatin Readers | TGAGTAAGGACCACAGTATG |
| Human | PYGO1 | Chromatin Readers | CTCCAAAGCCAGGATAACCA |
| Human | PYGO1 | Chromatin Readers | TGGTCAGAGTTTGGATTCGG |
| Human | PYGO1 | Chromatin Readers | TCATCCATATTCAAGTGAGG |
| Human | PYGO1 | Chromatin Readers | TCATCCACCTCACTTGAATA |
| Human | RNF8 | Transcription Elongation | GGTTTCGAGAAATCATCAGG |
| Human | RNF8 | Transcription Elongation | GTGGTTTCGAGAAATCATCA |
| Human | RNF8 | Transcription Elongation | ATAATTGTCCATTGGCCCTC |
| Human | RNF8 | Transcription Elongation | TGGTTTCGAGAAATCATCAG |
| Human | RNF8 | Transcription Elongation | ATTTGGTGTCACATACCAAC |

| | | | |
|---|---|---|---|
| Human | RNF8 | Transcription Elongation | GTATCAATGAATGGATGAAG |
| Human | RNF8 | Transcription Elongation | CATCCATTCATTGATACAGT |
| Human | RNF8 | Transcription Elongation | ACAGTAGGAGCAGAAACTGT |
| Human | RNF8 | Transcription Elongation | TTGTTCAGAATACTTCATTG |
| Human | RNF8 | Transcription Elongation | GCTCCTACTGTATCAATGAA |
| Human | RNF8 | Transcription Elongation | GAAGAGGAACAGCATCTTCA |
| Human | RNF8 | Transcription Elongation | CAGGGTTTGGAGATAGCCCA |
| Human | RNF8 | Transcription Elongation | AAAGGACCTGAAGCAACAGC |
| Human | RNF8 | Transcription Elongation | GAAAGTTGTGCAAATGGAGC |
| Human | RNF8 | Transcription Elongation | GGAACAGCATCTTCAGGGTT |
| Human | RNF8 | Transcription Elongation | AGAGCAGACCAAGGAAGAGA |
| Human | RNF8 | Transcription Elongation | CATCTTCTCCTTCTCTTCCT |
| Human | RNF8 | Transcription Elongation | GGAGAAGATGCAAGCACAGA |
| Human | RNF8 | Transcription Elongation | ACGACGAATTGTTCTCATTA |
| Human | RNF8 | Transcription Elongation | AACGACGAATTGTTCTCATT |
| Human | LBR | Chromatin Readers | CCAAATGGCTTCAGAATCAG |
| Human | LBR | Chromatin Readers | AGGCCGACATTAAGGAAGCA |
| Human | LBR | Chromatin Readers | GTGAAGTGGTAAGAGGTCGA |
| Human | LBR | Chromatin Readers | AACATGATGAGAAACACACC |
| Human | LBR | Chromatin Readers | AGCAACAGGAAGAGGAACAC |
| Human | LBR | Chromatin Readers | ACAGGAAGAGGAACACAGGC |
| Human | LBR | Chromatin Readers | GGTGTGTTTCTCATCATGTT |
| Human | LBR | Chromatin Readers | TTCCGATGACTGCAGATGTC |
| Human | LBR | Chromatin Readers | ATCGGAACATCTCTCTTCCA |
| Human | LBR | Chromatin Readers | ATCCTGACATCTGCAGTCAT |
| Human | LBR | Chromatin Readers | CATCGGAACATCTCTCTTCC |
| Human | LBR | Chromatin Readers | CCACCCATCCAATCAATCCG |
| Human | LBR | Chromatin Readers | TGGATGGGTGGTTATTAACT |
| Human | LBR | Chromatin Readers | AACCACCCATCCAATCAATC |
| Human | LBR | Chromatin Readers | ACCACCCATCCAATCAATCC |
| Human | LBR | Chromatin Readers | CGATGGATTTGGATTCATGC |
| Human | LBR | Chromatin Readers | TTTGGATTCATGCTGGCTTT |
| Human | LBR | Chromatin Readers | ATTATTGTTCTGAAACTTTG |
| Human | LBR | Chromatin Readers | ACAATAATTAGAGAAGCCAT |
| Human | LBR | Chromatin Readers | TGTAGAAATAAGGCAGAATG |
| Human | LBR | Chromatin Readers | AAATAAATTATGTAGAAATA |
| Human | KDM4A | Histone Methylation | AGCTCCTTGGGATTCAATGT |
| Human | KDM4A | Histone Methylation | TTGAATCCCAAGGAGCTCAT |
| Human | KDM4A | Histone Methylation | ATTGCCTACATTGAATCCCA |
| Human | KDM4A | Histone Methylation | TCATCGGGCAGGGCTAGCCA |
| Human | KDM4A | Histone Methylation | CTTTGACAAGGTGACTCAAG |
| Human | KDM4A | Histone Methylation | TGCCGTACTCAATCCACCGA |

| | | | |
|---|---|---|---|
| Human | KDM4A | Histone Methylation | AGGATAAAGATTGTCGCTGA |
| Human | KDM4A | Histone Methylation | TCCTGGCTCACTATGTCCTC |
| Human | KDM4A | Histone Methylation | TCCTGAGGACATAGTGAGCC |
| Human | KDM4A | Histone Methylation | GACTTCATAGAAGGTCTCGG |
| Human | KDM4A | Histone Methylation | CCCTATCCAAATGTACCAGG |
| Human | KDM4A | Histone Methylation | TACCAGGTGGAGTTTGAGGA |
| Human | KDM4A | Histone Methylation | CTGGTACATTTGGATAGGGT |
| Human | KDM4A | Histone Methylation | AGCCATCCTCAAACTCCACC |
| Human | KDM4A | Histone Methylation | CCACCCTATCCAAATGTACC |
| Human | KDM4A | Histone Methylation | GGTCCATGCCAGTTGCTATG |
| Human | KDM4A | Histone Methylation | CATAGCAACTGGCATGGACC |
| Human | KDM4A | Histone Methylation | CTTGCAGGAAACGAGTATGC |
| Human | KDM4A | Histone Methylation | GTGCCATCCTCCTCAAGATA |
| Human | KDM4A | Histone Methylation | GAGAGCAAATGATGACAGGT |
| Human | KDM4A | Histone Methylation | AGAGAGCAAATGATGACAGG |
| Human | KDM4A | Histone Methylation | TGCTGTTTATGCTCATTACG |
| Human | KDM4A | Histone Methylation | TTTATGCTCATTACGAGGAG |
| Human | KDM4A | Histone Methylation | GTCAGGCTGCATCATCACAC |
| Human | KDM4A | Histone Methylation | TGATGATGCAGCCTGACGAC |
| Human | KDM4A | Histone Methylation | AATGTATCTTCTGTAAGAAG |
| Human | KDM4A | Histone Methylation | CATCATCACACCGGCAGCCT |
| Human | KDM4A | Histone Methylation | CACACTCCTGCAGCACAAGT |
| Human | KDM4A | Histone Methylation | GCAGCACAAGTGGGTGATGG |
| Human | KDM4A | Histone Methylation | AGCACAAGTGGGTGATGGCG |
| Human | KDM4A | Histone Methylation | CCTGCAGCACAAGTGGGTGA |
| Human | PRDM14 | Histone Methylation | ATGTGGGAGATCTTTGAAGA |
| Human | PRDM14 | Histone Methylation | AGAACCAAGAGCTCCTTGTG |
| Human | PRDM14 | Histone Methylation | GAAGCGGGCACAGTTGACAT |
| Human | PRDM14 | Histone Methylation | CTTATCTCTGTAATATTTGT |
| Human | PRDM14 | Histone Methylation | GAAGAATGTGGATCCGAAGC |
| Human | PRDM14 | Histone Methylation | GATCGTTTGCAGAGAGAACA |
| Human | PRDM14 | Histone Methylation | AAGCCGGTCCCGCTTCTCAA |
| Human | PRDM14 | Histone Methylation | CAAACGATCCTTTGAGAAGC |
| Human | PRDM14 | Histone Methylation | GCAAACGATCCTTTGAGAAG |
| Human | PRDM14 | Histone Methylation | AGTGTGTGTATTGTACTAAG |
| Human | PRDM14 | Histone Methylation | GTGTGTGCGGAGTATGCTGG |
| Human | PRDM14 | Histone Methylation | GCATACTCCGCACACACATC |
| Human | PRDM14 | Histone Methylation | GATGTGTGTGCGGAGTATGC |
| Human | PRDM14 | Histone Methylation | CTGCCCATGACAGCCATGTC |
| Human | PRDM14 | Histone Methylation | CATGGCTGTCATGGGCAGCA |
| Human | PRDM14 | Histone Methylation | ATGGCTGTCATGGGCAGCAT |
| Human | PRDM14 | Histone Methylation | CAGATCCAGAGCTGTCTGGG |

| | | | |
|---|---|---|---|
| Human | PRDM14 | Histone Methylation | TCAGATCCAGAGCTGTCTGG |
| Human | PRDM14 | Histone Methylation | ATCAGATCCAGAGCTGTCTG |
| Human | PRDM14 | Histone Methylation | CCTGCACTTCGTTCTGTACG |
| Human | PRKAA1 | Other Histone Modifications | TGGTATAGTTTGATAATATG |
| Human | PRKAA1 | Other Histone Modifications | ATCTCAGGCAGATTGTATGC |
| Human | PRKAA1 | Other Histone Modifications | TCATATTATCAAACTATACC |
| Human | PRKAA1 | Other Histone Modifications | ACCTTCGGCAAAGTGAAGAT |
| Human | PRKAA1 | Other Histone Modifications | GCCTGAAACCTCATCCAGAA |
| Human | PRKAA1 | Other Histone Modifications | GTAATGAACAGTTTATATAG |
| Human | PRKAA1 | Other Histone Modifications | CAGATGGAGGACTAGAGGCG |
| Human | ZMYND11 | Chromatin Readers | ATTCAAGAGAAAGTGAATGA |
| Human | ZMYND11 | Chromatin Readers | ATTCACTTTCTCTTGAATGG |
| Human | ZMYND11 | Chromatin Readers | AAGGAATCCAGGCCCTCTGG |
| Human | ZMYND11 | Chromatin Readers | CAGAAGGAATCCAGGCCCTC |
| Human | ZMYND11 | Chromatin Readers | GAATCCAGGCCCTCTGGTGG |
| Human | ZMYND11 | Chromatin Readers | GCCACCACCACCAGAGGGCC |
| Human | ZMYND11 | Chromatin Readers | TTAGAGACAGCAGTAGTCCC |
| Human | ZMYND11 | Chromatin Readers | TCACATATCAACACCTCTCC |
| Human | ZMYND11 | Chromatin Readers | GAACTCATCAGACAAACACT |
| Human | ZMYND11 | Chromatin Readers | TGAATGCCATTTGCCTGGAG |
| Human | ZMYND11 | Chromatin Readers | GTTCCAGCAGCAGTGGTACA |
| Human | ZMYND11 | Chromatin Readers | CCAGCAGGAGCACTGGCACG |
| Human | ZMYND11 | Chromatin Readers | CTGCTCCATCAAGTGCCAGC |
| Human | ZMYND11 | Chromatin Readers | CTGGCACTTGATGGAGCAGT |
| Human | ZMYND11 | Chromatin Readers | AGGATGTGTTCCAGCAGCAG |
| Human | ZMYND11 | Chromatin Readers | GCAGTGGTGCTACAACTGTG |
| Human | ZMYND11 | Chromatin Readers | AAAGCTGTAGCCAACATGCA |
| Human | ZMYND11 | Chromatin Readers | GTGGTGCTACAACTGTGAGG |
| Human | ZMYND11 | Chromatin Readers | TAAAGACACATGTCATGAGC |
| Human | ZMYND11 | Chromatin Readers | ATGCAGCTCCAGCTCATCAC |
| Human | RSF1 | Chromatin Readers | TCTCCGTTGCCGACGTCCGG |
| Human | RSF1 | Chromatin Readers | AGCAGACTACGGCGAAGTTG |
| Human | RSF1 | Chromatin Readers | AACTCAGGCAGGTCTAGCAG |
| Human | RSF1 | Chromatin Readers | CCGTTGCCGACGTCCGGCGG |
| Human | RSF1 | Chromatin Readers | ACTTCTCCGTTGCCGACGTC |
| Human | RSF1 | Chromatin Readers | ACAGAAGAATTAGCTCAGGA |
| Human | RSF1 | Chromatin Readers | TCTGGGATGATCATCAGAGG |
| Human | RSF1 | Chromatin Readers | CTGGGATGATCATCAGAGGA |
| Human | RSF1 | Chromatin Readers | CCATCTGGGATGATCATCAG |
| Human | RSF1 | Chromatin Readers | TCACACAGAAGAATTAGCTC |
| Human | RSF1 | Chromatin Readers | AAAGAACGCTTGGTGTATGT |
| Human | RSF1 | Chromatin Readers | GGAAGAAACACCTAAACAAG |

| | | | |
|---|---|---|---|
| Human | RSF1 | Chromatin Readers | CCTGAAAGGATCGAATTTGG |
| Human | RSF1 | Chromatin Readers | TGTCAAACCCATTAAAGTTG |
| Human | RSF1 | Chromatin Readers | GTCTCCTCATCCTCTAAGCT |
| Human | RSF1 | Chromatin Readers | TCTGCCTCATTGGCCGAGAG |
| Human | CDK4 | Transcription Elongation | GTACCAGAGTGTAACAACCA |
| Human | SETDB1 | Histone Methylation | CGGAACTGTATCCCATTTGC |
| Human | SETDB1 | Histone Methylation | TGGTCGCCAAATACAAAGAT |
| Human | SETDB1 | Histone Methylation | ACAAAGATGGGAATCAGGTC |
| Human | SETDB1 | Histone Methylation | TGCTTCCTATGTCACACAGT |
| Human | SETDB1 | Histone Methylation | GGATCTATCGAGGCTCTACA |
| Human | SETDB1 | Histone Methylation | GTGGTGGAAGTCCCGAGTTG |
| Human | SETDB1 | Histone Methylation | CCAGCTTATCAAGACTGAGT |
| Human | SETDB1 | Histone Methylation | GTTATCTATAAGACACCTTG |
| Human | SETDB1 | Histone Methylation | GCGTCGAGTTAACCGCAAGA |
| Human | SETDB1 | Histone Methylation | TCTTGCGGTTAACTCGACGC |
| Human | SETDB1 | Histone Methylation | ATAGTAACGGGACCAGCAGA |
| Human | SETDB1 | Histone Methylation | AGAAGAGTGTCTACCCACAG |
| Human | SETDB1 | Histone Methylation | ACTCTTCTAGTCTCTTGTAC |
| Human | SETDB1 | Histone Methylation | CTGGATAGTTAGTTGATGGC |
| Human | SETDB1 | Histone Methylation | ACAGTGTCAGGACATCATCA |
| Human | SETDB1 | Histone Methylation | TGGTCCAATGAAGCGTCAAG |
| Human | SETDB1 | Histone Methylation | TGCATACTAGTCTTACTAGG |
| Human | SETDB1 | Histone Methylation | CTATTCAAGACACAGAACAA |
| Human | SETDB1 | Histone Methylation | ACTTCGGCATTTCATCGATG |
| Human | SETDB1 | Histone Methylation | GCTGCAACAGGCAGTGGTTG |
| Human | SETDB1 | Histone Methylation | TGAGGAACTGGGTATCTCTA |
| Human | SETDB1 | Histone Methylation | CAGGCAGTGGTTGAGGAACT |
| Human | SETDB1 | Histone Methylation | CATACTCCTCTATGAAGTCA |
| Human | SETDB1 | Histone Methylation | CTCCTCTATGAAGTCACGGC |
| Human | DIMT1 | Control | AGGAATGGGATGGTCTAGTA |
| Human | DIMT1 | Control | GACTTCATCAGATTGCTACA |
| Human | SMYD3 | Histone Methylation | TGATTGAGCAAAGAGATACT |
| Human | SMYD3 | Histone Methylation | ACTGAAGGCACACTGGAAGT |
| Human | SMYD3 | Histone Methylation | ATGGATGCCTGCATCAACCT |
| Human | SMYD3 | Histone Methylation | GTTCTATGGTACTCGGACCA |
| Human | RAD54B | Chromatin Remodeling | ATCTTCTAATCTGTGACGAG |
| Human | RAD54B | Chromatin Remodeling | TGATCTTCTAATCTGTGACG |
| Human | RAD54B | Chromatin Remodeling | ATTACTGGCTTGCCTCCATA |
| Human | RAD54B | Chromatin Remodeling | AGACATTGCCTGAATGTCAG |
| Human | RAD54B | Chromatin Remodeling | CAGTAAGCAAGGTCTTTGTG |
| Human | ELP1 | Transcription Elongation | CTCTCATAGCATCGCAGACA |
| Human | ELP1 | Transcription Elongation | CGATGCTATGAGAGCAGTCA |

| | | | |
|---|---|---|---|
| Human | ELP1 | Transcription Elongation | GGATATCAGCATTGCTTATG |
| Human | ELP1 | Transcription Elongation | CAGGATATCAGCATTGCTTA |
| Human | ELP1 | Transcription Elongation | CATAAGCAATGCTGATATCC |
| Human | ELP1 | Transcription Elongation | AGGATATCAGCATTGCTTAT |
| Human | ELP1 | Transcription Elongation | TTCCTGCCAGAGTTCTGCCG |
| Human | ELP1 | Transcription Elongation | GGCCTCGGCAGAACTCTGGC |
| Human | ELP1 | Transcription Elongation | GGTGGGCCTCGGCAGAACTC |
| Human | ELP1 | Transcription Elongation | TTCTGCCGAGGCCCACCAGC |
| Human | ELP1 | Transcription Elongation | ACAAGGAAGGGAATTACAGA |
| Human | MAPK3 | Other Histone Modifications | ACAAATCTTAAGGTCGCAGG |
| Human | MAPK3 | Other Histone Modifications | GAGATCATGCTGAACTCCAA |
| Human | TAF10 | Transcription Elongation | AATATCTGAGATGAATTTCT |
| Human | TAF10 | Transcription Elongation | GATGAATTTCTGGGCAGCTA |
| Human | TAF10 | Transcription Elongation | CAATATCTGAGATGAATTTC |
| Human | ASXL1 | Chromatin Readers | GCACAATACACAGAGCTTTG |
| Human | ASXL1 | Chromatin Readers | TTCTGTCACGATGACTGTAT |
| Human | ASXL1 | Chromatin Readers | CTTTCAGGCTGCACGCACAC |
| Human | ASXL1 | Chromatin Readers | GCAGCCTTGGCACATGATCA |
| Human | ASXL1 | Chromatin Readers | GAAAGTTGAATCCGGATGGG |
| Human | ASXL1 | Chromatin Readers | TTTCACGTATCAAACCACCC |
| Human | ASXL1 | Chromatin Readers | GGCTGACCTTTAACCACCCA |
| Human | ASXL1 | Chromatin Readers | TCTGCCACCTCCCTCATCGG |
| Human | ASXL1 | Chromatin Readers | TGTGGCCACCCTGAGCCCAG |
| Human | ASXL1 | Chromatin Readers | GGTGATGGTGGTGAGGCCTG |
| Human | KDM2A | Histone Methylation | CCAGTCAATGAAATCCACCG |
| Human | KDM2A | Histone Methylation | CCAGCCTGAGGGAATGACGA |
| Human | KDM2A | Histone Methylation | TCAATGAAATCCACCGTGGA |
| Human | KDM2A | Histone Methylation | GGTGCAGAGGCCCTCCACGG |
| Human | KDM2A | Histone Methylation | ACAGCATGAATCCAGCCTGA |
| Human | KDM2A | Histone Methylation | TACATTCACAAAGTTCTCTG |
| Human | KDM2A | Histone Methylation | GGTGCTGCGACAAGAGACTT |
| Human | KDM2A | Histone Methylation | TGTGCAAGACGTGGTATAAA |
| Human | KDM2A | Histone Methylation | GTTCTCTGCGGCTGAGGTAG |
| Human | KDM2A | Histone Methylation | TGGCTCGTCAATAGGCTGCC |
| Human | KDM2A | Histone Methylation | AACTGACATGGCTCGTCAAT |
| Human | KDM2A | Histone Methylation | AGATCAAGGGTCCTGAGAAG |
| Human | KDM2A | Histone Methylation | GATCAAGGGTCCTGAGAAGG |
| Human | KDM2A | Histone Methylation | GAGGGCAGAGACTGCAGACC |
| Human | KDM2A | Histone Methylation | AGGTCGAGTCGAGACAGGAG |
| Human | KDM2A | Histone Methylation | TGGAGGACTGATCTGTAAGG |
| Human | KDM2A | Histone Methylation | CTCTGTGAGAGAGTAGCGAG |
| Human | KDM2A | Histone Methylation | GACAGCAGTGAGTAGATTGG |

| | | | |
|---|---|---|---|
| Human | KDM2A | Histone Methylation | CCAATCTACTCACTGCTGTC |
| Human | KDM2A | Histone Methylation | GATTGGAGGACTGATCTGTA |
| Human | KDM2A | Histone Methylation | AATGCGCCGTAGGTAGATCA |
| Human | KDM2A | Histone Methylation | TGACGTTGGCAATGCGCCGT |
| Human | KDM2A | Histone Methylation | GCCGTAGGTAGATCAGGGTC |
| Human | KDM2A | Histone Methylation | ACCAGACCCTGATCTACCTA |
| Human | KDM2A | Histone Methylation | CAATGCGCCGTAGGTAGATC |
| Human | KDM2A | Histone Methylation | GTCTGAGATGAAGTGCTCGC |
| Human | KDM2A | Histone Methylation | CAGGCAGTAGAGGCTGTTGA |
| Human | KDM2A | Histone Methylation | ACCACACTCTCCTTGCACAC |
| Human | KDM2A | Histone Methylation | AGAGACATGAAGAAGTTTGG |
| Human | KDM2A | Histone Methylation | CAGGACTGCTTCATGCGTCC |
| Human | KDM2A | Histone Methylation | GAGACATGAAGAAGTTTGGG |
| Human | KDM2A | Histone Methylation | GCAGAGACATGAAGAAGTTT |
| Human | KDM2A | Histone Methylation | GCCTCCAGATGGACGGAGAG |
| Human | KDM2A | Histone Methylation | CCTGGCTGCCTCCAGATGGA |
| Human | KDM2A | Histone Methylation | CTGCCTCCAGATGGACGGAG |
| Human | KDM2A | Histone Methylation | TCTGATCCACCTCTCCACAG |
| Human | KDM2A | Histone Methylation | TGAGATGTGTTGGTATGTGT |
| Human | SUPT6H | Transcription Elongation | CCAGAAGAACGAGTGAAGGT |
| Human | SUPT6H | Transcription Elongation | TGTCACTGAGGAATTTGGTG |
| Human | SUPT6H | Transcription Elongation | TTGTCACTGAGGAATTTGGT |
| Human | SUPT6H | Transcription Elongation | GATTATCCGACCAAGCAGCA |
| Human | SUPT6H | Transcription Elongation | CTCACTGTTGATCCACAGAG |
| Human | SUPT6H | Transcription Elongation | ATTATCCGACCAAGCAGCAA |
| Human | SUPT6H | Transcription Elongation | TCTACCAGCATGTGGATGTG |
| Human | SUPT6H | Transcription Elongation | GCTTCCTGGATACACAGACG |
| Human | SUPT6H | Transcription Elongation | GGAAGAGGAGGATGATGATG |
| Human | SUPT6H | Transcription Elongation | TGAAGAAGGTGAAGGTGACG |
| Human | SUPT6H | Transcription Elongation | TCCCTGAACTAAGCGAGGTG |
| Human | SUPT6H | Transcription Elongation | AGCCATGCAGCCATCGACTG |
| Human | SUPT6H | Transcription Elongation | ATCGCCAGCAATGGACATGG |
| Human | SUPT6H | Transcription Elongation | GGCATCGCCAGCAATGGACA |
| Human | SUPT6H | Transcription Elongation | CATCGCCAGCAATGGACATG |
| Human | SIRT2 | Histone Acetylation | TATGCAGTCAGACTTCCTGA |
| Human | SIRT2 | Histone Acetylation | GCAGTCAGACTTCCTGAAGG |
| Human | SIRT2 | Histone Acetylation | CAAGGAGAAAGCTGGCCAGT |
| Human | SIRT2 | Histone Acetylation | GCTGGAATCTCCACATCCGC |
| Human | SIRT2 | Histone Acetylation | GCTGGAGGACCTTGTCCGGA |
| Human | SIRT2 | Histone Acetylation | TGGCGTGCTCCCTCCGGACA |
| Human | SIRT2 | Histone Acetylation | AGCTGGAGGACCTTGTCCGG |
| Human | SIRT2 | Histone Acetylation | AGGAGCTGGAGGACCTTGTC |

| | | | |
|---|---|---|---|
| Human | KDM6A | Histone Methylation | ACTGTAAACTGTAGTACCTC |
| Human | KDM6A | Histone Methylation | AAGCATTATCTGCATACCAG |
| Human | KDM6A | Histone Methylation | TGGTATGCAGATAATGCTTT |
| Human | KDM6A | Histone Methylation | TTGGATAATCTTCCAATAAG |
| Human | KDM6A | Histone Methylation | AGGTCACTTCAACCTCTTAT |
| Human | KDM6A | Histone Methylation | CCATTATAATGCATTTCAGT |
| Human | KDM6A | Histone Methylation | GGCAATTAAAGCATTTCAGG |
| Human | KDM6A | Histone Methylation | TCCATTATAATGCATTTCAG |
| Human | KDM6A | Histone Methylation | GTGGGCAATTAAAGCATTTC |
| Human | KDM6A | Histone Methylation | TGCAGAATGATATTTCCTCT |
| Human | KDM6A | Histone Methylation | CCCACTTATATGAAACCCAG |
| Human | KDM6A | Histone Methylation | TCCTCTGGGTTTCATATAAG |
| Human | KDM6A | Histone Methylation | CCTCTGGGTTTCATATAAGT |
| Human | KDM6A | Histone Methylation | TTGCAGAATGATATTTCCTC |
| Human | KDM6A | Histone Methylation | CCTGGGAGATAAAGCCACCA |
| Human | KDM6A | Histone Methylation | CATCACACTGTAGATCTCCT |
| Human | KDM6A | Histone Methylation | GCATCACACTGTAGATCTCC |
| Human | KDM6A | Histone Methylation | GATCTGCTTCCAAGGACTTT |
| Human | KDM6A | Histone Methylation | TACTTGAATAGCACCTTCCG |
| Human | KDM6A | Histone Methylation | AGTCCTGGTATTTCCTCGGA |
| Human | KDM6A | Histone Methylation | GGAAGGTGCTATTCAAGTAT |
| Human | KDM6A | Histone Methylation | GCACCTTCCGAGGAAATACC |
| Human | KDM6A | Histone Methylation | AGACTGCCTGTAAGATATAA |
| Human | KDM6A | Histone Methylation | ATAGGCCTGTAAAGCATCCA |
| Human | KDM6A | Histone Methylation | CAATTGTACAGCACAAATAT |
| Human | KDM6A | Histone Methylation | TAGGCCTGTAAAGCATCCAT |
| Human | KDM6A | Histone Methylation | TCAGCCCATGGATGCTTTAC |
| Human | KDM6A | Histone Methylation | TAGCATTTAATGGCATCCTG |
| Human | KDM6A | Histone Methylation | ATTTAATGGCATCCTGAGGC |
| Human | KDM6A | Histone Methylation | TGAATCCTGCAACCAGCCTC |
| Human | KDM6A | Histone Methylation | GGCATCCTGAGGCTGGTTGC |
| Human | KDM6A | Histone Methylation | CCAATATGGTATGACCAACA |
| Human | KDM6A | Histone Methylation | AACTATACATGAAAGTTCCA |
| Human | KDM6A | Histone Methylation | GCTGGTGCAACAACATTGCT |
| Human | KDM6A | Histone Methylation | TTATACTGGCAGGCTGTAAG |
| Human | KDM6A | Histone Methylation | GGAGCTCTTAAGTTAGACCT |
| Human | KDM6A | Histone Methylation | AACTATCTAACTCCACTCAG |
| Human | KDM6A | Histone Methylation | CATCGTCAGATAATTCTGGG |
| Human | KDM6A | Histone Methylation | TGGCAGTGGAACGGTACGAA |
| Human | KDM6A | Histone Methylation | GAAGCAATGTCAGACATTGA |
| Human | KDM6A | Histone Methylation | TGAAGCAATGTCAGACATTG |
| Human | HSPBAP1 | Histone Methylation | AGGTACAAGGAAGGAAACGA |

| | | | |
|---|---|---|---|
| Human | HSPBAP1 | Histone Methylation | GAACAAAGAGAACCTGTCCT |
| Human | HSPBAP1 | Histone Methylation | GGAACAAAGAGAACCTGTCC |
| Human | HSPBAP1 | Histone Methylation | GTCAGCATAAGCCCAGAACT |
| Human | HSPBAP1 | Histone Methylation | CATAAGCCCAGAACTTGGAA |
| Human | HSPBAP1 | Histone Methylation | ATGGTCCAGAAATACTAGAC |
| Human | HSPBAP1 | Histone Methylation | TGTGACCAGTCTAGTATTTC |
| Human | HSPBAP1 | Histone Methylation | ACTAGACTGGTCACAGTTCC |
| Human | CENPT | Chromatin Remodeling | CAGGTTGAGGGATCTGGTGA |
| Human | CENPT | Chromatin Remodeling | TCCTCCAACACAATGTTTGG |
| Human | CSNK2B | Other Histone Modifications | GATCAATCCATAAAGCATCT |
| Human | CSNK2B | Other Histone Modifications | CAGGCAGCCGAGATGCTTTA |
| Human | CSNK2B | Other Histone Modifications | TCTCGGCTGCCTGCTCAATC |
| Human | PARP9 | Other Histone Modifications | ATGTACTGGAAAGCTGCAGA |
| Human | PARP9 | Other Histone Modifications | TTATCCCATTCCATCCACCG |
| Human | PARP9 | Other Histone Modifications | TGGATGGAATGGGATAAACA |
| Human | PARP9 | Other Histone Modifications | GCTGGTGAGATAGCTGTCAC |
| Human | PARP9 | Other Histone Modifications | CTATCCGGGTTAGTTTGCAA |
| Human | PARP9 | Other Histone Modifications | CTGCCGTCTGCCATTCAATG |
| Human | PARP9 | Other Histone Modifications | TGATATTACAGTTGGACCTG |
| Human | PARP9 | Other Histone Modifications | ACAGTTTCAACGGTCCCAGT |
| Human | PARP9 | Other Histone Modifications | TTCCATGTTTCCAGTCCCAA |
| Human | PARP9 | Other Histone Modifications | TGGTTTGCAGGTTCTAAAGG |
| Human | PARP9 | Other Histone Modifications | GTATTTGTGGACATGCACCC |
| Human | PARP9 | Other Histone Modifications | GTGCATGTCCACAAATACTG |
| Human | PARP9 | Other Histone Modifications | TCAGTTCTGAAATGGCAAAG |
| Human | SP140 | Chromatin Readers | TCTCGGAGGCCCATAAGGAA |
| Human | SP140 | Chromatin Readers | AATACATCACTCTTGTCACT |
| Human | SP140 | Chromatin Readers | AGATTGCAAGTGCAATAACA |
| Human | SP140 | Chromatin Readers | TTCTCCATCAGCCATCGTAG |
| Human | SP140 | Chromatin Readers | GGTGAAGTGTATACAGACTG |
| Human | SP140 | Chromatin Readers | CATTCTCCATCAGCCATCGT |
| Human | SP140 | Chromatin Readers | CTTTGATTTCAAATTCCGTG |
| Human | SP140 | Chromatin Readers | CATAAGAAGAAATTGCAGCA |
| Human | SP140 | Chromatin Readers | TCCCGCCTGTGGAAGCTGAG |
| Human | SP140 | Chromatin Readers | GGACTGTCACATCCCGCCTG |
| Human | SP140 | Chromatin Readers | TTGTTCAAGAGTCTTCCATG |
| Human | MORF4L1 | Chromatin Readers | CACAGATAATAAGGAGTATG |
| Human | MORF4L1 | Chromatin Readers | TCACGATTTCCTAAAGTACC |
| Human | MORF4L1 | Chromatin Readers | ACAAATAATCTCAGGAGATG |
| Human | MORF4L1 | Chromatin Readers | CAATTCGTACAAATAATCTC |
| Human | MORF4L1 | Chromatin Readers | ACTTCATACATTACAGTGGT |
| Human | MORF4L1 | Chromatin Readers | TTTCACTTGTTTGTCCTTTA |

| | | | |
|---|---|---|---|
| Human | MORF4L1 | Chromatin Readers | TTCAACAGTAGGATCTACCC |
| Human | MORF4L1 | Chromatin Readers | TTATGAAGCAAAGTGTGTAA |
| Human | MORF4L1 | Chromatin Readers | CACTTTGCTTCATAAAGAAG |
| Human | MORF4L1 | Chromatin Readers | TTCTCGCTGTTTCTGCAAAT |
| Human | ASXL3 | Chromatin Readers | TTCTGCCATGACGACTGCAT |
| Human | ASXL3 | Chromatin Readers | AAGGACCTATGCAGTCGTCA |
| Human | ASXL3 | Chromatin Readers | AAAGCCATGATTGTGTGCAA |
| Human | ASXL3 | Chromatin Readers | GATTGTGTGCAAAGGCTGTG |
| Human | ASXL3 | Chromatin Readers | ACACAATCATGGCTTTCAGC |
| Human | ASXL3 | Chromatin Readers | GTTAAAGGTAAGTCAGACAT |
| Human | ASXL3 | Chromatin Readers | AGGCCCGGGCCCAACAAGCT |
| Human | ASXL3 | Chromatin Readers | CCCTCGCAGATATCAAGGCC |
| Human | ASXL3 | Chromatin Readers | CCCGGGCCTTGATATCTGCG |
| Human | ASXL3 | Chromatin Readers | GGCCCGGGCCCAACAAGCTC |
| Human | ASXL3 | Chromatin Readers | CCTCGCAGATATCAAGGCCC |
| Human | ASXL3 | Chromatin Readers | GTCAACAGACATTAGGACAG |
| Human | ASXL3 | Chromatin Readers | CCCATTTCAGCCACTACAGA |
| Human | ASXL3 | Chromatin Readers | TCCCATTTCAGCCACTACAG |
| Human | ASXL3 | Chromatin Readers | CAGTGCTGCAAACAGATACA |
| Human | ASXL3 | Chromatin Readers | TTTGGTATAATGGCTATGCA |
| Human | ASXL3 | Chromatin Readers | GGGACTTCTGCATTAGGGAG |
| Human | ASXL3 | Chromatin Readers | TCTGCATTAGGGAGAGGTGG |
| Human | ASXL3 | Chromatin Readers | CAGATGGGACTTCTGCATTA |
| Human | ASXL3 | Chromatin Readers | ACTTCTGCATTAGGGAGAGG |
| Human | TRIM24 | Chromatin Readers | TTGATGGTTGACAAATCCAT |
| Human | TRIM24 | Chromatin Readers | CTGGATGTTCTGGTGGCACA |
| Human | TRIM24 | Chromatin Readers | GCCACCAGAACATCCAGAGC |
| Human | TRIM24 | Chromatin Readers | TGGCAGAAAGAGTGCAGGCA |
| Human | TRIM24 | Chromatin Readers | CCACCAGAACATCCAGAGCC |
| Human | TRIM24 | Chromatin Readers | CTGGCAGAAAGAGTGCAGGC |
| Human | TRIM24 | Chromatin Readers | TCAGGTATGTACAAGCTGTG |
| Human | TRIM24 | Chromatin Readers | TTTGTGTAGAGTGTGTTGAA |
| Human | TRIM24 | Chromatin Readers | GTGTATCAGAGCTCATCAGA |
| Human | TRIM24 | Chromatin Readers | CGTGTATCAGAGCTCATCAG |
| Human | TRIM24 | Chromatin Readers | CCTCTTTCTGTCTGACAGTG |
| Human | TRIM24 | Chromatin Readers | GTGCAAATCCACTCTCCACT |
| Human | TRIM24 | Chromatin Readers | ACTGCTATTTAAGAGCAGGG |
| Human | TRIM24 | Chromatin Readers | TCTGACTGCTATTTAAGAGC |
| Human | TRIM24 | Chromatin Readers | CTGACTGCTATTTAAGAGCA |
| Human | TRIM24 | Chromatin Readers | CAGTCAGAGCTCTACTTCTG |
| Human | TRIM24 | Chromatin Readers | AGGGAGGTGAGTATGCTTCT |
| Human | TRIM24 | Chromatin Readers | TCAGTTAGAGAGCCTTGCAA |

| | | | |
|---|---|---|---|
| Human | TRIM24 | Chromatin Readers | GTTGCATAAGTTTCATGCGA |
| Human | TRIM24 | Chromatin Readers | TCATGCGATGGTCCTTTGCA |
| Human | TRIM24 | Chromatin Readers | CAACAACAACAGGAAGTGGC |
| Human | TRIM24 | Chromatin Readers | AGTTGCTATATTTACACTGA |
| Human | TRIM24 | Chromatin Readers | CTATAGGTGTTAACTTAACA |
| Human | UBE2B | Other Histone Modifications | GGCTCATGCGGGATTTCAAG |
| Human | UBE2B | Other Histone Modifications | AATCCCGCATGAGCCTCCTC |
| Human | UBE2B | Other Histone Modifications | ATCCCGCATGAGCCTCCTCC |
| Human | TCF19 | Chromatin Readers | GCTCGGCATGCAGTTCGGCG |
| Human | TCF19 | Chromatin Readers | CATTATTGACCAAAGTACCT |
| Human | TCF19 | Chromatin Readers | CCACAGCAGCCAAGGTACTT |
| Human | TCF19 | Chromatin Readers | TGACCCTCCAGTCATCACCC |
| Human | TCF19 | Chromatin Readers | CTGTGACGTCTGGTTCCATG |
| Human | TCF19 | Chromatin Readers | ACAGCCATCACACTGAACCC |
| Human | TCF19 | Chromatin Readers | CTGGATGCTGCAGCCAACAC |
| Human | TCF19 | Chromatin Readers | TGCAGCCAACACAGGCCACA |
| Human | TCF19 | Chromatin Readers | ACTCCCACTGGAAATCGACG |
| Human | TCF19 | Chromatin Readers | CGGCCACGTCGATTTCCAGT |
| Human | TCF19 | Chromatin Readers | ACGGCCACGTCGATTTCCAG |
| Human | TCF19 | Chromatin Readers | ATTTCCAGTGGGAGTCAGTG |
| Human | FEN1 | DNA and RNA Modifications | TTGCCATCAAAGACATACAC |
| Human | FEN1 | DNA and RNA Modifications | TCTCCATCATGCGAATGGTG |
| Human | FEN1 | DNA and RNA Modifications | GATGTGCTGCAGAATGAGGA |
| Human | FEN1 | DNA and RNA Modifications | GCCGTTCTCCATCATGCGAA |
| Human | FEN1 | DNA and RNA Modifications | AGTCTATGCTGCGGCTACCG |
| Human | FEN1 | DNA and RNA Modifications | TTATCTTGATGCACCCAGTG |
| Human | FEN1 | DNA and RNA Modifications | CAAGATAAGGGATGCCCATG |
| Human | FEN1 | DNA and RNA Modifications | TGGCCAAACGCAGTGAGCGG |
| Human | FEN1 | DNA and RNA Modifications | GGCCAAACGCAGTGAGCGGC |
| Human | FEN1 | DNA and RNA Modifications | AGCTGGCCAAACGCAGTGAG |
| Human | FEN1 | DNA and RNA Modifications | AGCCCGCCGCTCACTGCGTT |
| Human | FEN1 | DNA and RNA Modifications | ACGCAGTGAGCGGCGGGCTG |
| Human | DNA2 | DNA and RNA Modifications | GCTTCGGAAATGGAGAAGAG |
| Human | DNA2 | DNA and RNA Modifications | GCAATAATACCAATATCAGA |
| Human | GTF2A1 | Transcription Elongation | ATCACAGATCTGTATAATTT |
| Human | HDAC9 | Histone Acetylation | TTGCTGTGAAACCATTTGCT |
| Human | HDAC9 | Histone Acetylation | TTGATCCAATGATGTGTGGT |
| Human | HDAC9 | Histone Acetylation | GAAACCATTTGCTCGGCATG |
| Human | HDAC9 | Histone Acetylation | CTGAAGAATCCACAGCCATG |
| Human | HDAC9 | Histone Acetylation | CATCACCTAGGAGTATCCTG |
| Human | HDAC9 | Histone Acetylation | GTAGATCTGGATGTTCACCA |
| Human | TRDMT1 | DNA and RNA Modifications | CTCCATCAGTACCTGACCAG |

| | | | |
|---|---|---|---|
| Human | TRDMT1 | DNA and RNA Modifications | ACTCCATCAGTACCTGACCA |
| Human | TRDMT1 | DNA and RNA Modifications | TACAGCTTTCTCTCAGCGCG |
| Human | TRDMT1 | DNA and RNA Modifications | ATATCCTTTGGTAAAGCACA |
| Human | DNMT3A | Chromatin Readers | TGGGTCATGTGGTTCGGAGA |
| Human | DNMT3A | Chromatin Readers | CTCCGAACCACATGACCCAG |
| Human | DNMT3A | Chromatin Readers | AAGGCACCCGCTGGGTCATG |
| Human | DNMT3A | Chromatin Readers | CACCAAGACACAATGCGGCC |
| Human | DNMT3A | Chromatin Readers | GCGGGCACAAGGGTACCTAC |
| Human | DNMT3A | Chromatin Readers | GGTAACATTGAGGCTCCCAC |
| Human | DNMT3A | Chromatin Readers | TTCCGCACATGAGCACCTCA |
| Human | DNMT3A | Chromatin Readers | GGTGTTCCAGGGTAACATTG |
| Human | DNMT3A | Chromatin Readers | TAGCCAAGTTCAGCAAAGTG |
| Human | DNMT3A | Chromatin Readers | GCTCGCAAGGGCCTCTACGA |
| Human | DNMT3A | Chromatin Readers | CAGAAGCATATCCAGGAGTG |
| Human | DNMT3A | Chromatin Readers | ACGGCCTGTTCATACCGGGA |
| Human | DNMT3A | Chromatin Readers | CACAGAAGCATATCCAGGAG |
| Human | DNMT3A | Chromatin Readers | GACGAGCCAGAGTACGAGGA |
| Human | DNMT3A | Chromatin Readers | CATCTACGAGGTCCTGCAGG |
| Human | DNMT3A | Chromatin Readers | GCTTGTTGTACGTGGCCTGG |
| Human | DNMT3A | Chromatin Readers | AGAACAAGCCCATGATTGAA |
| Human | KDM4D | Histone Methylation | TCCCAAGGTGCACACAGAGC |
| Human | KDM4D | Histone Methylation | AGATAATTCCACCCAAAGAA |
| Human | KDM4D | Histone Methylation | TGATAAATATATTGCTTACA |
| Human | KDM4D | Histone Methylation | AGGTGCACACAGAGCTGGCT |
| Human | KDM4D | Histone Methylation | GCCAGCTCTGTGTGCACCTT |
| Human | KDM4D | Histone Methylation | CTTCAATCGCATAACTCAGG |
| Human | CENPM | Chromatin Remodeling | CCAGCTTCACCACGGTGTGC |
| Human | CENPM | Chromatin Remodeling | CCGGCACACCGTGGTGAAGC |
| Human | CENPM | Chromatin Remodeling | CACCAGGCGCTGCGCCATGG |
| Human | CENPM | Chromatin Remodeling | GGCCACCATGGCGCAGCGCC |
| Human | CENPM | Chromatin Remodeling | GCGCACCAGGCGCTGCGCCA |
| Human | YTHDF2 | DNA and RNA Modifications | AAATATTATACTTAATGGAA |
| Human | YTHDF2 | DNA and RNA Modifications | CAGTGGACACTTCTGTGGCG |
| Human | YTHDF2 | DNA and RNA Modifications | ACACTATGAGAAACGCCAAG |
| Human | YTHDF2 | DNA and RNA Modifications | ACCGAAGCTTCTCCAACACT |
| Human | YTHDF2 | DNA and RNA Modifications | TTTGACTGGAATCTGAAACA |
| Human | MCM3 | DNA and RNA Modifications | GGTCATACCTGCCGTAGACA |
| Human | MCM3 | DNA and RNA Modifications | GTCATGCCCATGGATACTTG |
| Human | MCM3 | DNA and RNA Modifications | CATCAATGCAAACCACGCCT |
| Human | MCM3 | DNA and RNA Modifications | CAGAAGCTGAGACTTGGCAA |
| Human | ADRM1 | Transcription Elongation | GTGACTCCGGATAAGCGGAA |
| Human | ADRM1 | Transcription Elongation | TTTCCGCTTATCCGGAGTCA |

| | | | |
|---|---|---|---|
| Human | ADRM1 | Transcription Elongation | TGACTCCGGATAAGCGGAAA |
| Human | ADRM1 | Transcription Elongation | GGCCGCCAACAAGGGCGATG |
| Human | ADRM1 | Transcription Elongation | TGGATCTCATCCGCGGTCTG |
| Human | ADRM1 | Transcription Elongation | GCTTCCCTACTTGCCATCTG |
| Human | ADRM1 | Transcription Elongation | GCAGAAAGAGGGCGACACGA |
| Human | ADRM1 | Transcription Elongation | TCTGCGCTAGGCGGTGAGGG |
| Human | ADRM1 | Transcription Elongation | CCGGCCTTGGAGGACTGGGT |
| Human | ADRM1 | Transcription Elongation | CCAGCCCACCCAGTCCTCCA |
| Human | ADRM1 | Transcription Elongation | CACGAACTCTCTGCGCTAGG |
| Human | ADRM1 | Transcription Elongation | GCCGGCCTTGGAGGACTGGG |
| Human | ADRM1 | Transcription Elongation | CGACTGGCTCCGGGAGCTGG |
| Human | ADRM1 | Transcription Elongation | CTCCCGGAGCCAGTCGGCAG |
| Human | ADRM1 | Transcription Elongation | GCAGCTCCTCCTCCAGCTCC |
| Human | ADRM1 | Transcription Elongation | CATTCCCGGAACTGGGCGCG |
| Human | SMYD4 | Histone Methylation | AGGTCCTGTGTGTTGTATGG |
| Human | SMYD4 | Histone Methylation | GACACTTAATTATGGCCTAG |
| Human | FOXO1 | DNA and RNA Modifications | CACGAATGAACTTGCTGTGT |
| Human | FOXO1 | DNA and RNA Modifications | CGCTGTCGCAGATCTACGAG |
| Human | FOXO1 | DNA and RNA Modifications | GGGTGACAGCAACAGCTCGG |
| Human | FOXO1 | DNA and RNA Modifications | TAAGGGTGACAGCAACAGCT |
| Human | FOXO1 | DNA and RNA Modifications | CAAGGCCATCGAGAGCTCGG |
| Human | FOXO1 | DNA and RNA Modifications | GACAATAAGTCGAGTTATGG |
| Human | FOXO1 | DNA and RNA Modifications | CGGTTCATACCCGAGGTGTG |
| Human | FOXO1 | DNA and RNA Modifications | TTATGACACCAGTTGATCCT |
| Human | TET1 | DNA and RNA Modifications | GTAAGTGCATTCACCACAGT |
| Human | TET1 | DNA and RNA Modifications | TGCCAGCAGAAGACCAACTG |
| Human | TET1 | DNA and RNA Modifications | CACCACAGTTGGTCTTCTGC |
| Human | TET1 | DNA and RNA Modifications | GGTCCTAGCAAATCAGACAG |
| Human | TET1 | DNA and RNA Modifications | GTCCTAGCAAATCAGACAGA |
| Human | TET1 | DNA and RNA Modifications | GAAGAGGAGGTACTGACCAT |
| Human | TET1 | DNA and RNA Modifications | TACTTTAATGGCTGTAAGTT |
| Human | RNF17 | Chromatin Readers | GGAAGTTGAAGTTAAATATG |
| Human | RNF17 | Chromatin Readers | TATCGGATTGCCTGGACATC |
| Human | RNF17 | Chromatin Readers | ATCTTCAAATTTAGCTACAC |
| Human | RNF17 | Chromatin Readers | AAGAGGCCAGATCATCAGAA |
| Human | RNF17 | Chromatin Readers | GAGGTCTTGCTGTATGATGT |
| Human | RNF17 | Chromatin Readers | GGTTACAGACACATTGGTAG |
| Human | RNF17 | Chromatin Readers | TAAGAGGATCCGATACTCTG |
| Human | RNF17 | Chromatin Readers | GTATCGTGGCAAGGTGATGG |
| Human | RNF17 | Chromatin Readers | TCCGATACTCTGTGGTATCG |
| Human | RNF17 | Chromatin Readers | ATAGGGTAAAGATGGCACTG |
| Human | RNF17 | Chromatin Readers | TCATCATATTCTGCAAGGCA |

| | | | |
|---|---|---|---|
| Human | RNF17 | Chromatin Readers | AATTGTACTAAGATAGATAA |
| Human | RNF17 | Chromatin Readers | AGCCATCATCATATTCTGCA |
| Human | RNF17 | Chromatin Readers | TAAAGGATTAAATTCTTTAA |
| Human | RNF17 | Chromatin Readers | TTAGTACAATTTGTTGATTA |
| Human | RNF17 | Chromatin Readers | CACCATTGTGAACTTCAATG |
| Human | RNF17 | Chromatin Readers | TTGAAGTTCACAATGGTGAC |
| Human | RNF17 | Chromatin Readers | GTCCACATTGAAGTTCACAA |
| Human | RNF17 | Chromatin Readers | ACAATGGTGACCGGATGATC |
| Human | RNF17 | Chromatin Readers | GGATACCCTCCTTCCACACC |
| Human | MLLT3 | Chromatin Readers | TGCAGGTGAAGCTGGAGCTG |
| Human | MLLT3 | Chromatin Readers | CAGCTCCAGCTTCACCTGCA |
| Human | MLLT3 | Chromatin Readers | GTGTGCCGTGCAGGTGAAGC |
| Human | MLLT3 | Chromatin Readers | GTGCAGGTGAAGCTGGAGCT |
| Human | MLLT3 | Chromatin Readers | CGTGCAGGTGAAGCTGGAGC |
| Human | MLLT3 | Chromatin Readers | AACTGGTAGAGCTTCACAGA |
| Human | KDM5C | Histone Methylation | ATGCCCGATTTCTCTGCGAT |
| Human | KDM5C | Histone Methylation | AGATCCGCCCACCCGCGGAC |
| Human | KDM5C | Histone Methylation | GCCCAGCTAGGCTCGAACAC |
| Human | KDM5C | Histone Methylation | AAGCTATCTGCAAGGACCGT |
| Human | KDM5C | Histone Methylation | TCCATTAACTACCTCCACTG |
| Human | KDM5C | Histone Methylation | TGCGGACAACTGGCACACCA |
| Human | KDM5C | Histone Methylation | GCGGACAACTGGCACACCAT |
| Human | KDM5C | Histone Methylation | GGCACCGCCAGACACCCTTG |
| Human | KDM5C | Histone Methylation | CATCACAGCCATCACACAGC |
| Human | KDM5C | Histone Methylation | CCAAGGTATACCGATACCGC |
| Human | KDM5C | Histone Methylation | CCTGCGGTATCGGTATACCT |
| Human | KDM5C | Histone Methylation | CTTGCAGAGATCATTGATGT |
| Human | KDM5C | Histone Methylation | AGAGATCATTGATGTGGGAA |
| Human | KDM5C | Histone Methylation | CCAGTAGCCGGCAGTACCTG |
| Human | KDM5C | Histone Methylation | CTCGCCTCCTCAGCTCTCCG |
| Human | KDM5C | Histone Methylation | GCACTGACACACACCGCCCA |
| Human | KDM5C | Histone Methylation | GTCAGGACTGGTTCCATGGG |
| Human | KDM5C | Histone Methylation | ATTCCCACCAGGCCAGCAGT |
| Human | KDM5C | Histone Methylation | TTCCCACCAGGCCAGCAGTG |
| Human | KDM5C | Histone Methylation | TTGTGGAGAAGGAGTTCTGG |
| Human | KDM5C | Histone Methylation | AACTTGTGGAGAAGGAGTTC |
| Human | KDM5C | Histone Methylation | AGAAGAGCGGCGTCTACGAA |
| Human | KDM5C | Histone Methylation | TCATCATGGTGCAAGAAGAG |
| Human | KDM5C | Histone Methylation | GCATAAGGAGATGTTCATCA |
| Human | KDM5C | Histone Methylation | CCTGGCGGCAGCTGTGCATA |
| Human | KDM5C | Histone Methylation | CCTTATGCACAGCTGCCGCC |
| Human | KDM5C | Histone Methylation | TGCCATGCTGCATAAGCTGA |

| | | | |
|---|---|---|---|
| Human | KDM5C | Histone Methylation | TTCAGCTTATGCAGCATGGC |
| Human | ALKBH5 | DNA and RNA Modifications | GGACACCACCTCGTCAATGC |
| Human | ALKBH5 | DNA and RNA Modifications | GAGGCGCGCAAGGTGAAGAG |
| Human | ALKBH5 | DNA and RNA Modifications | GGGACACCACCTCGTCAATG |
| Human | ALKBH5 | DNA and RNA Modifications | GATCGAGGCCCGCATTGACG |
| Human | ALKBH5 | DNA and RNA Modifications | GGACGAGTGCGCCAAGATCG |
| Human | ALKBH5 | DNA and RNA Modifications | TGCGGCTGCGGCGGCTACGG |
| Human | ALKBH5 | DNA and RNA Modifications | TGCGGCGGCTACGGCGGCTG |
| Human | ALKBH5 | DNA and RNA Modifications | GGCTGCGGCTGCGGCGGCTA |
| Human | ALKBH5 | DNA and RNA Modifications | GGCTACGGCGGCTGCGGCAG |
| Human | ALKBH5 | DNA and RNA Modifications | TACGGCGGCTGCGGCAGCGG |
| Human | GUK1 | Control | TCAGCGTGTCCCATACCACG |
| Human | GUK1 | Control | CCTGTATGGCACGAGCAAGG |
| Human | GUK1 | Control | GAACCTGTATGGCACGAGCA |
| Human | EZH1 | Histone Methylation | GGAAACGATTCTGACAGTCT |
| Human | EZH1 | Histone Methylation | TTGGTAGTTGTACACTTGTG |
| Human | EZH1 | Histone Methylation | TCTCGCACTGCCAGATAGCA |
| Human | EZH1 | Histone Methylation | GACTGTCAGAATCGTTTCCC |
| Human | EZH1 | Histone Methylation | TCCCAACTGTTATGCCAAAG |
| Human | EZH1 | Histone Methylation | GCCAAAGTGGTCATGGTGAA |
| Human | EZH1 | Histone Methylation | TGAGGCTGATCGACGCGGAA |
| Human | EZH1 | Histone Methylation | TGGCAAAGATCCCAATCCGA |
| Human | EZH1 | Histone Methylation | CTGTTATGCCAAAGTGGTCA |
| Human | KDM7A | Histone Methylation | ACAGCATGGATCCACCCTGT |
| Human | KDM7A | Histone Methylation | CCTTATTTGTTCCTACAGGG |
| Human | KDM7A | Histone Methylation | ATACCTTATTTGTTCCTACA |
| Human | KDM7A | Histone Methylation | CTACTCCAACACAGCTGCCG |
| Human | KDM7A | Histone Methylation | GATCGAGTGCGATATCTGCA |
| Human | KDM7A | Histone Methylation | ATGAAGCGGTTCACGTCGTA |
| Human | KDM7A | Histone Methylation | TATCGCACTCGATCATGAAG |
| Human | KDM7A | Histone Methylation | TGGTTCCACGGCAGCTGTGT |
| Human | KDM7A | Histone Methylation | GACTACACAGAAATTGATGA |
| Human | KDM7A | Histone Methylation | AGCCGCCGCGGCAGCCGTGT |
| Human | KDM7A | Histone Methylation | GCCGGGAGCCGCCACCGACA |
| Human | KDM7A | Histone Methylation | CGCCGCGGCAGCCGTGTCGG |
| Human | KDM7A | Histone Methylation | CGCCACCGACACGGCTGCCG |
| Human | KDM7A | Histone Methylation | CACCGACACGGCTGCCGCGG |
| Human | KDM7A | Histone Methylation | GCAGCTCAGGTGTTATGAGA |
| Human | KDM7A | Histone Methylation | TTTGGCTACAAACCAACATA |
| Human | KDM7A | Histone Methylation | GTGCTTTCACTCCCTGTACT |
| Human | KDM7A | Histone Methylation | CTTTAGAAAGTTCTTTAATA |
| Human | ASH1L | Histone Methylation | GCGAGCAATGCTGTAACCAG |

| | | | |
|---|---|---|---|
| Human | ASH1L | Histone Methylation | GCCACATGGGCAAGTGTTGG |
| Human | ASH1L | Histone Methylation | CAGCAAAGATCATTCTATTG |
| Human | ASH1L | Histone Methylation | TACAGCATTGCTCGCCACAT |
| Human | ASH1L | Histone Methylation | CGCCACATGGGCAAGTGTTG |
| Human | ASH1L | Histone Methylation | TCTAGAACGATTTCGAGCTG |
| Human | ASH1L | Histone Methylation | GTGATTGACAGTTACCGCAT |
| Human | ASH1L | Histone Methylation | ATCGGGCCTCATTTCCCATG |
| Human | ASH1L | Histone Methylation | TGTGGCTTTGAGAAATGTCG |
| Human | ASH1L | Histone Methylation | GGAATGCTGAGAAGTACTAT |
| Human | ASH1L | Histone Methylation | CACTGGTTACTATAAGACAG |
| Human | ASH1L | Histone Methylation | TATTGTAAAGGGAGACCCAA |
| Human | ASH1L | Histone Methylation | CACACCACTCTCCATCCCGT |
| Human | ASH1L | Histone Methylation | AATGTGAGATAACAGTCGAT |
| Human | ASH1L | Histone Methylation | GTCGATAGGACTGACGGACC |
| Human | ASH1L | Histone Methylation | GGACATTTCCGAGGTCGACC |
| Human | ASH1L | Histone Methylation | TGAGACCTTCATCCTTGTAG |
| Human | ASH1L | Histone Methylation | AGTGTGACAAGTGCATGGTA |
| Human | ASH1L | Histone Methylation | TATGGGAGTGAACTCAGATG |
| Human | ASH1L | Histone Methylation | ATACCATGCACTTGTCACAC |
| Human | ASH1L | Histone Methylation | TGTGGCCTCTACAAGGATGA |
| Human | ASH1L | Histone Methylation | AAATGTCGAGGAATCATCGG |
| Human | ASH1L | Histone Methylation | TGGTAGCTTCGTAACCAGAA |
| Human | ASH1L | Histone Methylation | GTAGCTTCGTAACCAGAAAG |
| Human | ASH1L | Histone Methylation | AAACAGCAACTTTGTAAGTG |
| Human | ASH1L | Histone Methylation | GAGAAATGTCGAGGAATCAT |
| Human | ASH1L | Histone Methylation | TAGGATTAGGTTACTATGGA |
| Human | ASH1L | Histone Methylation | TGGAGGAGGTGGATAAAGAG |
| Human | ASH1L | Histone Methylation | GTGGTGAAAGAAGGAGATGG |
| Human | ASH1L | Histone Methylation | CCATAGTAACCTAATCCTAT |
| Human | ASH1L | Histone Methylation | AATCCTCCAAGGGATAAGCT |
| Human | ASH1L | Histone Methylation | AGTCACTGCCTTCTAACGAA |
| Human | ASH1L | Histone Methylation | GCACAAATTGTAGCCCTACC |
| Human | ASH1L | Histone Methylation | ACTAATCTTGGACTTCGAGA |
| Human | ASH1L | Histone Methylation | ATTGTAGCCCTACCCGGAAA |
| Human | ASH1L | Histone Methylation | ATCTTCAAAGAAATTTGTGA |
| Human | ASH1L | Histone Methylation | TTTGAAGATCTGGGCTAGGC |
| Human | ASH1L | Histone Methylation | ACAAATTTCTTTGAAGATCT |
| Human | ASH1L | Histone Methylation | CTTTGAAGATCTGGGCTAGG |
| Human | ASH1L | Histone Methylation | TTTCTTTGAAGATCTGGGCT |
| Human | ASH1L | Histone Methylation | CTATTACAATGCCCGGCATG |
| Human | ASH1L | Histone Methylation | AGATGTTTGTCGTCTACGAA |
| Human | ASH1L | Histone Methylation | GAAAGGCCTATTACAATGCC |

| | | | |
|---|---|---|---|
| Human | ASH1L | Histone Methylation | CTGGGCTGATGCCTCATGCC |
| Human | ASH1L | Histone Methylation | TCTGGGCTGATGCCTCATGC |
| Human | PRMT7 | Histone Methylation | CTGAAGTCTATGCTGAACAT |
| Human | PRMT7 | Histone Methylation | CGAGGTGACTGTAGGTCCAG |
| Human | PRMT7 | Histone Methylation | GAGGTGACTGTAGGTCCAGA |
| Human | PRMT7 | Histone Methylation | TGGAGTTCAGGGACCTGTGG |
| Human | CENPC | Chromatin Remodeling | AAACCTTTGGAGTACTGGCG |
| Human | CENPC | Chromatin Remodeling | AGGTTTCAAACGTGTTCTCT |
| Human | CENPC | Chromatin Remodeling | TCTCCTCGCCAGTACTCCAA |
| Human | CENPC | Chromatin Remodeling | GAGAACACGTTTGAAACCTT |
| Human | CENPC | Chromatin Remodeling | GTTTGAAACCTTTGGAGTAC |
| Human | CENPC | Chromatin Remodeling | ATGTTATAATAGTTACCTGA |
| Human | CENPC | Chromatin Remodeling | ACCTTATATATTAAGTACTG |
| Human | CENPC | Chromatin Remodeling | ACACCTTATATATTAAGTAC |
| Human | CENPC | Chromatin Remodeling | CACCTTATATATTAAGTACT |
| Human | CENPC | Chromatin Remodeling | ATCAGTCATCTAAGAATATT |
| Human | CECR2 | Chromatin Readers | GAATTGTCGAAAGTATAATG |
| Human | CECR2 | Chromatin Readers | AATAATCTGATAATAGTTAG |
| Human | CECR2 | Chromatin Readers | TGGAGGTTTATACTGTACCA |
| Human | CECR2 | Chromatin Readers | TATGTATAAAGTTCTAGACG |
| Human | CECR2 | Chromatin Readers | ATTTAGAGAGGTGTTTCCAT |
| Human | CECR2 | Chromatin Readers | AGATGTCTGATAATTTAGAG |
| Human | CECR2 | Chromatin Readers | TTTAGAGAGGTGTTTCCATC |
| Human | CECR2 | Chromatin Readers | CTCTAAATTATCAGACATCT |
| Human | SIRT5 | Histone Acetylation | CTTCAGGCCACAATATAGCT |
| Human | SIRT5 | Histone Acetylation | CAGGCAAATCTGGTTTCGTG |
| Human | SIRT5 | Histone Acetylation | AGCTGGGAAATCAATCGACT |
| Human | SIRT5 | Histone Acetylation | TGGTTTCGTGTGGACGCTGG |
| Human | SIRT5 | Histone Acetylation | CAGCCTGCCTCTTCACACCG |
| Human | SIRT5 | Histone Acetylation | AGCTACCATGGATCTCCAGA |
| Human | SP100 | Chromatin Readers | AAAGACATTTCCATTCCTCG |
| Human | SP100 | Chromatin Readers | GCAGCCTGTCATCTACACCC |
| Human | SP100 | Chromatin Readers | TTGTGATGAGATCACGATCA |
| Human | SP100 | Chromatin Readers | TGGAAGCTAAGTATACGCTG |
| Human | SP100 | Chromatin Readers | AAGCTAAGTATACGCTGCGG |
| Human | SP100 | Chromatin Readers | TATAAGGAGCGATTCAAACA |
| Human | SP100 | Chromatin Readers | TCTGAACTTCCTGTGACCTG |
| Human | SP100 | Chromatin Readers | TACCTTAGTTGACCCTTGTG |
| Human | SP100 | Chromatin Readers | TCTTCTTATGCTCCTCACAA |
| Human | SP100 | Chromatin Readers | CTCCTCACAAGGGTCAACTA |
| Human | SP100 | Chromatin Readers | ATTTGAAGATATGGCAAAGG |
| Human | SP100 | Chromatin Readers | TAAAGAAGTGCTCAGAGACA |

| | | | |
|---|---|---|---|
| Human | SP100 | Chromatin Readers | ACATCATCAATGGACAGGCC |
| Human | SP100 | Chromatin Readers | ATGTTGTGAAGAAACTGGCA |
| Human | SP100 | Chromatin Readers | AAACTGCTTGTCAGCTGCAG |
| Human | SP100 | Chromatin Readers | TCACAACATCATCAATGGAC |
| Human | SP100 | Chromatin Readers | ATGGGCTCTGGTACAATCTG |
| Human | SP100 | Chromatin Readers | GCAGCTCATCCCTAAGAAGA |
| Human | SP100 | Chromatin Readers | CTGAAACGTCTCTTCCTGCT |
| Human | SP100 | Chromatin Readers | CCAGATTGTACCAGAGCCCA |
| Human | SP100 | Chromatin Readers | TTCCTGTTCTGTGCGACTGG |
| Human | SP100 | Chromatin Readers | TAATTCCTGTTCTGTGCGAC |
| Human | SP100 | Chromatin Readers | AAGTAAATGGTCTCCAAAGA |
| Human | SP100 | Chromatin Readers | AAAGTAAATGGTCTCCAAAG |
| Human | SP100 | Chromatin Readers | AGGCATAGATCTAAAGTAAA |
| Human | BRD9 | Chromatin Readers | ATAATGCAATGACATACAAT |
| Human | BRD9 | Chromatin Readers | GGAGCAATTGCATCCGTGAC |
| Human | BRD9 | Chromatin Readers | GTCACGGATGCAATTGCTCC |
| Human | BRD9 | Chromatin Readers | GGAGAAGCATCTGGACGATG |
| Human | BRD9 | Chromatin Readers | TGAGGAAAGAAGGAAGCGAA |
| Human | BRD9 | Chromatin Readers | GAAGAAGAAGAAGTCCGAGA |
| Human | BRD9 | Chromatin Readers | ATCTGGACGATGAGGAAAGA |
| Human | BRD9 | Chromatin Readers | GTCCGAGAAGGAGAAGCATC |
| Human | BRD9 | Chromatin Readers | GCTGGCGGAGGAAGTGTTCC |
| Human | BRD9 | Chromatin Readers | GCGAAGAAGATCCTTCACGC |
| Human | SHH | Transcription Elongation | TCCTCGCTGCTGGTATGCTC |
| Human | SHH | Transcription Elongation | ATACCAGCAGCGAGGAGACG |
| Human | SHH | Transcription Elongation | CTCCTCGCTGCTGGTATGCT |
| Human | SHH | Transcription Elongation | AGTCCTCGTCTCCTCGCTGC |
| Human | SHH | Transcription Elongation | GTTGAACGCTTTGGCCATCT |
| Human | SHH | Transcription Elongation | TAAGGACAAGTTGAACGCTT |
| Human | ZCWPW1 | Chromatin Readers | CTATCATGCCTGGCCACCAG |
| Human | ZCWPW1 | Chromatin Readers | CACATGGTACTTAGACGGCA |
| Human | ZCWPW1 | Chromatin Readers | TCTATCATGCCTGGCCACCA |
| Human | ZCWPW1 | Chromatin Readers | TAGACGGCAGGGAATCAAGA |
| Human | ZCWPW1 | Chromatin Readers | TAATCGCTGTGATATTCCTG |
| Human | ZCWPW1 | Chromatin Readers | CACAGTTTGGGAAGGAACAC |
| Human | ZCWPW1 | Chromatin Readers | AATTATCTGGGAGAACTGAG |
| Human | ZCWPW1 | Chromatin Readers | TCCCAAACTGTGGGAAATGG |
| Human | ZCWPW1 | Chromatin Readers | CAGTGTTCCTTCCCAAACTG |
| Human | ZCWPW1 | Chromatin Readers | TCTTACCAATGCAGAATTTG |
| Human | ZCWPW1 | Chromatin Readers | GCAAGCAGAGAAGAAAGAAA |
| Human | SETD2 | Histone Methylation | GCTCAAGGTGAAATAGCATG |
| Human | SETD2 | Histone Methylation | TCAAGGTGAAATAGCATGTG |

| | | | |
|---|---|---|---|
| Human | SETD2 | Histone Methylation | GGGATTATTGTTCCAATAGA |
| Human | SETD2 | Histone Methylation | CTCAAGGTGAAATAGCATGT |
| Human | SETD2 | Histone Methylation | TTCTTCTCGGTGTCCAAATG |
| Human | SETD2 | Histone Methylation | TCTAGGACAAAGGTGTTCGA |
| Human | SETD2 | Histone Methylation | GACTGTGAACGGACAACTGA |
| Human | SETD2 | Histone Methylation | GGACTGTGAACGGACAACTG |
| Human | SETD2 | Histone Methylation | GGACAAAGGTGTTCGAAGGA |
| Human | SETD2 | Histone Methylation | TGCGGATCAGCCAATTGCCG |
| Human | SETD2 | Histone Methylation | CTGCGGATCAGCCAATTGCC |
| Human | SETD2 | Histone Methylation | TCTGCGGATCAGCCAATTGC |
| Human | SETD2 | Histone Methylation | TGATCCGCAGAAACATTTCT |
| Human | SETD2 | Histone Methylation | CTGATCCGCAGAAACATTTC |
| Human | SETD2 | Histone Methylation | TGATCACAAGGCAGACTCAG |
| Human | SETD2 | Histone Methylation | AGACAGCTCGAGATCCAGAA |
| Human | SETD2 | Histone Methylation | GATCACAAGGCAGACTCAGT |
| Human | SETD2 | Histone Methylation | GAGTCTGCCTTGTGATCACA |
| Human | SETD2 | Histone Methylation | ATTACTACCATGTGATCACA |
| Human | SETD2 | Histone Methylation | CCTGAAGTCATCCATGACAC |
| Human | SETD2 | Histone Methylation | GAAGCCCAGAAATGTTTCTG |
| Human | SETD2 | Histone Methylation | AAGTCATCCATGACACAGGA |
| Human | SETD2 | Histone Methylation | GGAAGGAATGTGCCAAGCAA |
| Human | SETD2 | Histone Methylation | CCAACCAGTGCTGAACCTGG |
| Human | SETD2 | Histone Methylation | CCTGCATGGGATAACCTGGT |
| Human | SETD2 | Histone Methylation | AGGGTTGCTGGGATCCACAT |
| Human | SETD2 | Histone Methylation | GCTGTACACTGACAGACTGT |
| Human | SETD2 | Histone Methylation | TTCTGTAGAATGTCCCACCA |
| Human | SETD2 | Histone Methylation | GAACACACTGGGTCCATGCT |
| Human | SETD2 | Histone Methylation | CTGGCTCGCAAGCTGACTCA |
| Human | SETD2 | Histone Methylation | GGCTGTTTACAAACCCAAAG |
| Human | SETD2 | Histone Methylation | CAGGCACTGGACGATGAACT |
| Human | SETD2 | Histone Methylation | GAAGTACATGCAGAAGTTTG |
| Human | SETD2 | Histone Methylation | TTGAGCAAGAGGTGGCTCAA |
| Human | SETD2 | Histone Methylation | TGAGCAAGAGGTGGCTCAAC |
| Human | SETD2 | Histone Methylation | GAAGTTGTTTGAGCAAGAGG |
| Human | SETD2 | Histone Methylation | CCGGAAGTTGTTTGAGCAAG |
| Human | SETD2 | Histone Methylation | CCTCTTGCTCAAACAACTTC |
| Human | SETD2 | Histone Methylation | ACTGCTACATCTACTGGTAA |
| Human | SETD2 | Histone Methylation | ACATCTACTGGTAAGGGTAC |
| Human | SETD2 | Histone Methylation | TGTTATTGGTGTATGTGGCA |
| Human | SETD2 | Histone Methylation | AGCACCAGTAACAGAGCCAG |
| Human | SETD2 | Histone Methylation | GGTAATGGTGCTGCATGAGT |
| Human | SETD2 | Histone Methylation | ACTCTGATCGTCGCTACCAT |

| | | | |
|---|---|---|---|
| Human | SETD2 | Histone Methylation | GCGCGTCCTCTCTCGATAAG |
| Human | SETD2 | Histone Methylation | GATCAGAAAGAGAGCGACGG |
| Human | SETD2 | Histone Methylation | AGCGCGTCCTCTCTCGATAA |
| Human | UTY | Histone Methylation | AGAGGTTGAAGTGACCTAAT |
| Human | UTY | Histone Methylation | TTGAATAATCTTCCAACAAG |
| Human | UTY | Histone Methylation | AGGTCACTTCAACCTCTTGT |
| Human | UTY | Histone Methylation | TCAGAGATATTACAGTTTAC |
| Human | UTY | Histone Methylation | CTACTACAATGCATTTCATT |
| Human | UTY | Histone Methylation | TCTACTACAATGCATTTCAT |
| Human | UTY | Histone Methylation | CCCATTTGTATGAAACCCAG |
| Human | UTY | Histone Methylation | TGCAGAATGATACTTCCTCT |
| Human | UTY | Histone Methylation | TTGCAGAATGATACTTCCTC |
| Human | UTY | Histone Methylation | GAAGTATCATTCTGCAAAGG |
| Human | UTY | Histone Methylation | GAGGAAGTATCATTCTGCAA |
| Human | UTY | Histone Methylation | CATCATAATATGGATCTAGT |
| Human | UTY | Histone Methylation | AATAGCATAGCTTTCCTTTG |
| Human | UTY | Histone Methylation | AGTAGGAGACAAAGCCACAA |
| Human | UTY | Histone Methylation | GGAAGGTGTTATTCAAGTAT |
| Human | UTY | Histone Methylation | AGATTGCCTGTAAGATATAA |
| Human | UTY | Histone Methylation | GAAGGTGTTATTCAAGTATT |
| Human | UTY | Histone Methylation | TCAGCCTATGGATGCTTTAC |
| Human | UTY | Histone Methylation | ATATATTTGTGCTGTACAAT |
| Human | UTY | Histone Methylation | TATGCCTGTAAAGCATCCAT |
| Human | UTY | Histone Methylation | AGGATTCATAGAGAGTACCT |
| Human | UTY | Histone Methylation | GGCATCTTGAGGTTGATTGC |
| Human | UTY | Histone Methylation | GAGAGTACCTAGGTCCATCC |
| Human | UTY | Histone Methylation | TAGCATTTAATGGCATCTTG |
| Human | UTY | Histone Methylation | TGCATTTAGGTAGCATTTAA |
| Human | UTY | Histone Methylation | CCAGAATGGTATGCCCAACA |
| Human | UTY | Histone Methylation | AACTGTATATGAAAGTTCCA |
| Human | UTY | Histone Methylation | GCTGAATAAATCTATACACA |
| Human | UTY | Histone Methylation | ATTCCTGGAAGGAGGAAGCC |
| Human | UTY | Histone Methylation | AGCCTGGTATTGTGCGTATT |
| Human | UTY | Histone Methylation | ATACCAGGCTTCCTCCTTCC |
| Human | UTY | Histone Methylation | TGGCAGTGGAACGGTATGAA |
| Human | ZBTB38 | DNA and RNA Modifications | AGCTGCCAGAACATTGCTAT |
| Human | ZBTB38 | DNA and RNA Modifications | AAGCTGCCAGAACATTGCTA |
| Human | ZBTB38 | DNA and RNA Modifications | GGCTGGAAATACAGATTGTA |
| Human | ZBTB38 | DNA and RNA Modifications | CTTTGAGATCGTCCAGCTCC |
| Human | ZBTB38 | DNA and RNA Modifications | GATCGTCCAGCTCCAGGACG |
| Human | ZBTB38 | DNA and RNA Modifications | GAGCAGAGTGCTGCTGTCAA |
| Human | ZBTB38 | DNA and RNA Modifications | AGTGCTGCTGTCAAAGGCTT |

| | | | |
|---|---|---|---|
| Human | ZBTB38 | DNA and RNA Modifications | AACAATTCACCACCCTGAAC |
| Human | ZBTB38 | DNA and RNA Modifications | CCGATCCAACCTGTTCAGGG |
| Human | ZBTB38 | DNA and RNA Modifications | CCACCCTGAACAGGTTGGAT |
| Human | ZBTB38 | DNA and RNA Modifications | ATGCCGATCCAACCTGTTCA |
| Human | ZBTB38 | DNA and RNA Modifications | ATTCACCACCCTGAACAGGT |
| Human | ZBTB38 | DNA and RNA Modifications | ATGTCTTCGGAGGCTAGATA |
| Human | ZBTB38 | DNA and RNA Modifications | CATTGGCTGAGTACAGGACA |
| Human | ZBTB38 | DNA and RNA Modifications | TATTTGCATTGGCTGAGTAC |
| Human | ZBTB38 | DNA and RNA Modifications | CTGCAACAAAGTATTTGCAT |
| Human | ZBTB38 | DNA and RNA Modifications | CAAATACTTTGTTGCAGTAA |
| Human | ZBTB38 | DNA and RNA Modifications | ACTTTGTTGCAGTAATGGCA |
| Human | ZBTB38 | DNA and RNA Modifications | GGAAGGGCTCTGGAACTGCT |
| Human | ZBTB38 | DNA and RNA Modifications | CTGCTTGGCGCAGAGCTCGC |
| Human | ZBTB38 | DNA and RNA Modifications | CTACAATTGTACACCAGAGG |
| Human | ZBTB38 | DNA and RNA Modifications | CAGCTACAATTGTACACCAG |
| Human | ZBTB38 | DNA and RNA Modifications | TTTGCCAGACACGGACCACA |
| Human | ZBTB38 | DNA and RNA Modifications | CATTAACAAATTTAACCATG |
| Human | BRIX1 | Control | TTCAGATCTTTAGTACACCA |
| Human | YWHAG | Chromatin Readers | AGCAACTGGTGCAGAAAGCC |
| Human | YWHAG | Chromatin Readers | ACTGGTGCAGAAAGCCCGGC |
| Human | YWHAG | Chromatin Readers | TGGCCTACAAGAACGTTGTG |
| Human | YWHAG | Chromatin Readers | TGTGGCCTACAAGAACGTTG |
| Human | YWHAG | Chromatin Readers | GGAACGAAACCTTCTGTCTG |
| Human | YWHAG | Chromatin Readers | GTACCGGGAGAAGATAGAGA |
| Human | YWHAG | Chromatin Readers | CTATCTTCTCCCGGTACGCA |
| Human | YWHAG | Chromatin Readers | ACTCCTTCTCTATCTTCTCC |
| Human | YWHAG | Chromatin Readers | GAAGATAGAGAAGGAGTTGG |
| Human | YWHAG | Chromatin Readers | GGAGAAGATAGAGAAGGAGT |
| Human | YWHAG | Chromatin Readers | GTTCTACCTGAAGATGAAAG |
| Human | YWHAG | Chromatin Readers | TGTTCTACCTGAAGATGAAA |
| Human | YWHAG | Chromatin Readers | GTGTTCTACCTGAAGATGAA |
| Human | YWHAG | Chromatin Readers | GATCTCGTGGGCTTCGCTGT |
| Human | YWHAG | Chromatin Readers | GGCTTCGCTGTAGGCCTTCT |
| Human | YWHAG | Chromatin Readers | TTCGCTGTAGGCCTTCTCGG |
| Human | YWHAG | Chromatin Readers | GGTGGTGGAGTCCTCCGAGA |
| Human | YWHAG | Chromatin Readers | AGACGGAGTAGTTAAGAGCC |
| Human | YWHAG | Chromatin Readers | AGTTAAGAGCCAGGCCTAAT |
| Human | YWHAG | Chromatin Readers | CGAACGCGGTCTTGGCCAAG |
| Human | YWHAG | Chromatin Readers | GAGGAGCTGCATGATGAGCG |
| Human | MECOM | Histone Methylation | AGATCTCCCATCCATAACTG |
| Human | MECOM | Histone Methylation | TTCGAGAGTCAAATATGCCT |
| Human | MECOM | Histone Methylation | GATGCCAGTCAACCAGATGT |

| | | | |
|---|---|---|---|
| Human | MECOM | Histone Methylation | GTAGTTGCAGACATTGCGCC |
| Human | MECOM | Histone Methylation | TGACCAGCTCTTTGAATCTA |
| Human | MECOM | Histone Methylation | CAGCCTTAGATTCAAAGAGC |
| Human | MECOM | Histone Methylation | TTCTCCAGGCTTTGCAAATC |
| Human | MECOM | Histone Methylation | AGTGTCCCAAGGCATTTAAC |
| Human | MECOM | Histone Methylation | GGACTTCCAGTTAAATGCCT |
| Human | MECOM | Histone Methylation | GACTTCCAGTTAAATGCCTT |
| Human | MECOM | Histone Methylation | CGGACCCTAGCAACCTTCAG |
| Human | MECOM | Histone Methylation | TGTGCCGCTGAAGGTTGCTA |
| Human | MECOM | Histone Methylation | ATGTGCCGCTGAAGGTTGCT |
| Human | MECOM | Histone Methylation | TTGTTTGAGGCCCGACGAAG |
| Human | MECOM | Histone Methylation | CTTCCTTAAATAAACACAGG |
| Human | MECOM | Histone Methylation | CGTCTTCCTTAAATAAACAC |
| Human | MECOM | Histone Methylation | AATCTTGCCACAGTATCTGC |
| Human | MECOM | Histone Methylation | CGTGTTAGGTTTGCAGACCT |
| Human | MECOM | Histone Methylation | CAAGGTCTGCAAACCTAACA |
| Human | MECOM | Histone Methylation | GCATATCTTCTAACTTGCAA |
| Human | MECOM | Histone Methylation | CACTTATGTGATAGGTGTTT |
| Human | MECOM | Histone Methylation | TGTGAAGCCCTTTATCTGTG |
| Human | MECOM | Histone Methylation | AGCGATATTGCCGTTCTTCG |
| Human | MECOM | Histone Methylation | CCAAGAGATACACACGATCC |
| Human | MECOM | Histone Methylation | TGCTGTCACATACTGAAGAG |
| Human | MECOM | Histone Methylation | GTGACAAGTAATTTACATGA |
| Human | MECOM | Histone Methylation | AGATGAGGTGTTGTTAGATG |
| Human | MECOM | Histone Methylation | TTTCTTCATAGTCATCCTCA |
| Human | MECOM | Histone Methylation | TTTACATGAAGGAAACCCTG |
| Human | RNF20 | Other Histone Modifications | TGGCAGAAATGTTGGATCAG |
| Human | RNF20 | Other Histone Modifications | CTCACGAAGTTCATCTTCAA |
| Human | RNF20 | Other Histone Modifications | CTAGAACGGGTGAATTCCAA |
| Human | RNF20 | Other Histone Modifications | CTTGGAGAACTCCTGAGACA |
| Human | RNF20 | Other Histone Modifications | GCACAATCACTATCAATGCT |
| Human | RNF20 | Other Histone Modifications | AACAAGACACGCCTGCGTAG |
| Human | RNF20 | Other Histone Modifications | CAATTTCAAACGAGCCCAGG |
| Human | RNF20 | Other Histone Modifications | ACAAGCAGGCCCTATAAACA |
| Human | RNF20 | Other Histone Modifications | GGAAGAGATTAAGGATTACA |
| Human | HJURP | DNA and RNA Modifications | GCATGCGCCTCTGGAAGCGG |
| Human | HJURP | DNA and RNA Modifications | TCAGCCGCTGCATGCGCCTC |
| Human | HJURP | DNA and RNA Modifications | GCTGCATGCGCCTCTGGAAG |
| Human | HJURP | DNA and RNA Modifications | GCTTCCAGAGGCGCATGCAG |
| Human | HJURP | DNA and RNA Modifications | CCAGTCGCCGCCGCTTCCAG |
| Human | MeCP2 | DNA and RNA Modifications | GGACACGGAAGCTTAAGCAA |
| Human | MeCP2 | DNA and RNA Modifications | CCCTGCCTGAAGGCTGGACA |

| | | | |
|---|---|---|---|
| Human | MeCP2 | DNA and RNA Modifications | GTACGGTCTCCTGCACAGAT |
| Human | MeCP2 | DNA and RNA Modifications | GTCTTCTATCCGATCTGTGC |
| Human | MeCP2 | DNA and RNA Modifications | TCGGATAGAAGACTCCTTCA |
| Human | MeCP2 | DNA and RNA Modifications | GCGGCTGCCACCACACTCCC |
| Human | MeCP2 | DNA and RNA Modifications | GGGTGGGAGCAGTGGCACGG |
| Human | MeCP2 | DNA and RNA Modifications | ACCTGAGCCCGAGAGCTCCG |
| Human | MeCP2 | DNA and RNA Modifications | TCGGAGCTCTCGGGCTCAGG |
| Human | MeCP2 | DNA and RNA Modifications | TCCTCGGAGCTCTCGGGCTC |
| Human | CENPE | Chromatin Remodeling | GATTCTTCTCTGCTGTTCAG |
| Human | CENPE | Chromatin Remodeling | AACAGCAGAGAAGAATCACT |
| Human | CENPE | Chromatin Remodeling | TGCTACCATATCCAAAGAGT |
| Human | CENPE | Chromatin Remodeling | AAGTTTAGGCCTTTGTCCAG |
| Human | CENPE | Chromatin Remodeling | ATTCCGTTCCTTGCATTGAG |
| Human | CENPE | Chromatin Remodeling | TAATCAGCATCTTTCCAATG |
| Human | CENPE | Chromatin Remodeling | AGAAATTGTAGCTAAACACC |
| Human | CENPE | Chromatin Remodeling | TAAACACCTGGAAACTGAAG |
| Human | CCNH | Transcription Elongation | TTTGCATCTGAATTTGCGGT |
| Human | CCNH | Transcription Elongation | TCTGCAAATACTATGAGAAA |
| Human | CCNH | Transcription Elongation | GAAGGCACTTGAACAGATAC |
| Human | CCNH | Transcription Elongation | TCGATGAGGAAGCCCTCAAA |
| Human | CCNH | Transcription Elongation | AAATGGCAGTCAGGGCAATT |
| Human | CCNH | Transcription Elongation | AATGGCAGTCAGGGCAATTT |
| Human | CCNH | Transcription Elongation | TGCTGTTCTGAAACAGAAGT |
| Human | MNAT1 | Transcription Elongation | AGCTTTAGAAGTGGAACGAC |
| Human | MNAT1 | Transcription Elongation | ACTGGAAGAAGCTTTAGAAG |
| Human | MNAT1 | Transcription Elongation | ATTCACCATCAGCTTCAAGG |
| Human | MNAT1 | Transcription Elongation | AAGCTGATGGTGAATGTGTG |
| Human | MNAT1 | Transcription Elongation | TCACCATCAGCTTCAAGGAG |
| Human | RAD51 | Chromatin Remodeling | TGTTGCCTATGCGCCAAAGA |
| Human | RAD51 | Chromatin Remodeling | AAGGAGCTAATAAATATTAA |
| Human | RAD51 | Chromatin Remodeling | GAAGGAGCTAATAAATATTA |
| Human | RAD51 | Chromatin Remodeling | ATATTTATTAGCTCCTTCTT |
| Human | RAD51 | Chromatin Remodeling | TAGCTCCTTCTTTGGCGCAT |
| Human | RAD51 | Chromatin Remodeling | AGAACAGACTACTCGGGTCG |
| Human | RAD51 | Chromatin Remodeling | CAGAAGCATCCGCAGAAACC |
| Human | RAD51 | Chromatin Remodeling | GTAGTCTGTTCTGTAAAGGG |
| Human | RAD51 | Chromatin Remodeling | TCATGGCTGATGCTTGATAA |
| Human | DAPK3 | Other Histone Modifications | TGCTCGTCAAAGATCCCAAG |
| Human | DAPK3 | Other Histone Modifications | ATCACCTATATCCTCCTGAG |
| Human | DAPK3 | Other Histone Modifications | GTCCTGCACGAAGCGCATCT |
| Human | DAPK3 | Other Histone Modifications | CTGCACGAAGCGCATCTCGG |
| Human | DAPK3 | Other Histone Modifications | CTCCGAGATGCGCTTCGTGC |

| | | | |
|---|---|---|---|
| Human | MBD4 | DNA and RNA Modifications | GTGTACTTTATCAGCCCACA |
| Human | MBD4 | DNA and RNA Modifications | ATCCACATGGGACAGACTTA |
| Human | MBD4 | DNA and RNA Modifications | TAAGTCTGTCCCATGTGGAT |
| Human | MBD4 | DNA and RNA Modifications | GGGAAAGAGTTGTGAAGCAA |
| Human | MBD4 | DNA and RNA Modifications | AAATGGCAATACCTGTGCTT |
| Human | MBD4 | DNA and RNA Modifications | TTCATCTGAGAACTTGACAA |
| Human | MBD4 | DNA and RNA Modifications | TAAATATCATGACTGGCTTT |
| Human | MBD4 | DNA and RNA Modifications | AATTAAATAAATATCATGAC |
| Human | MBD4 | DNA and RNA Modifications | ATAAATATCATGACTGGCTT |
| Human | CHD6 | Chromatin Readers | ACTGTCCAGGAGGTTCACCC |
| Human | CHD6 | Chromatin Readers | ACATTGTAAATGGGCCACAA |
| Human | CHD6 | Chromatin Readers | AGTCGAACGGAGGTTCTCCT |
| Human | CHD6 | Chromatin Readers | TAGAACAGCTCCAAGTCGAA |
| Human | CHD6 | Chromatin Readers | ACCATATGAAGAAAGCACGT |
| Human | CHD6 | Chromatin Readers | AGAAAGCACGTGGGAGCTAG |
| Human | CHD6 | Chromatin Readers | ACACCAAGGATGCAGAAACA |
| Human | CHD6 | Chromatin Readers | TACCATATGAAGAAAGCACG |
| Human | CHD6 | Chromatin Readers | GTAGTGAGCACCACTTCACC |
| Human | CHD6 | Chromatin Readers | ATGGTGTACAGAGACGCCCA |
| Human | CHD6 | Chromatin Readers | GAGAAGCACTTTATGTTCCA |
| Human | CHD6 | Chromatin Readers | ACTTGAAGACTCCTGAAAGG |
| Human | CHD6 | Chromatin Readers | AACTTGAAGACTCCTGAAAG |
| Human | CHD6 | Chromatin Readers | AGGAACTGCAAACTTCTGGA |
| Human | CHD6 | Chromatin Readers | GTCAAACATCTCGCGCTCGT |
| Human | CHD6 | Chromatin Readers | TCTTCAGGACATCAACCGAA |
| Human | CHD6 | Chromatin Readers | TGGCTTACAGAACCGGTCGA |
| Human | CHD6 | Chromatin Readers | TTCCCGTTTGGACAAGAAGT |
| Human | CHD6 | Chromatin Readers | CATCCGACTTCTTGTCCAAA |
| Human | CHD6 | Chromatin Readers | GTTCCGCATCATTTCCCGTT |
| Human | CHD6 | Chromatin Readers | TGTCCAAACGGGAAATGATG |
| Human | CHD6 | Chromatin Readers | CTGAAATTAAGCATGTGGAG |
| Human | CHD6 | Chromatin Readers | AGACAGCCAGAAACGACGCT |
| Human | CHD6 | Chromatin Readers | GACATCCCATAATGGGATGG |
| Human | BRPF1 | Chromatin Readers | AGTCAGGTACCTCAGACAGA |
| Human | BRPF1 | Chromatin Readers | CTGCCTCAAGTATAACGCCA |
| Human | BRPF1 | Chromatin Readers | TAGTCAGGTACCTCAGACAG |
| Human | BRPF1 | Chromatin Readers | TGTCCTTGGCGTTATACTTG |
| Human | BRPF1 | Chromatin Readers | TGAGGCAGTTGCTGACGATG |
| Human | BRPF1 | Chromatin Readers | TGGTGGAACATACCTTCTCG |
| Human | BRPF1 | Chromatin Readers | TTGGATCAATGATCAGAGCT |
| Human | BRPF1 | Chromatin Readers | ACAACAAACGAACCTGGCAG |
| Human | BRPF1 | Chromatin Readers | CTCGGCATTTGGCCCACACG |

| | | | |
|---|---|---|---|
| Human | BRPF1 | Chromatin Readers | AAGGTCTACAAGAGTTACAG |
| Human | BRPF1 | Chromatin Readers | TGTAACTCTTGTAGACCTTT |
| Human | BRPF1 | Chromatin Readers | CCCGGTGGAGACCTGCCGAA |
| Human | BRPF1 | Chromatin Readers | ACCTTTCGGCAGGTCTCCAC |
| Human | BRPF1 | Chromatin Readers | CCTTTCGGCAGGTCTCCACC |
| Human | BRPF1 | Chromatin Readers | ACTGGCCCTCAGGGATATAG |
| Human | BRPF1 | Chromatin Readers | GGTTGCACATGTCACAGAAG |
| Human | BRPF1 | Chromatin Readers | CTTCTGTGACATGTGCAACC |
| Human | BRPF1 | Chromatin Readers | CCTTCAAGCAGACAGATGAC |
| Human | BRPF1 | Chromatin Readers | TGACGGGCGCTGGGCCCATG |
| Human | BRPF1 | Chromatin Readers | CCACAAGGCACACACCACAT |
| Human | BRPF1 | Chromatin Readers | GTGCGGGAGACAGGCGCCAA |
| Human | BRPF1 | Chromatin Readers | GACACTGAAAGAGGTGCCGT |
| Human | BRPF1 | Chromatin Readers | TGGTGCTCGCAGAAACATTG |
| Human | BRPF1 | Chromatin Readers | GTACATCGAGAAGTCTGCAG |
| Human | BRPF1 | Chromatin Readers | ATGTACCGGTAATAGGAAGT |
| Human | BRPF1 | Chromatin Readers | CCCTGAGAATGGCAGCAACA |
| Human | BRPF1 | Chromatin Readers | CGGTAATAGGAAGTTGGCCG |
| Human | BRPF1 | Chromatin Readers | CCTCGGCTGCATCTTCAGTG |
| Human | BRPF1 | Chromatin Readers | CCACACTGAAGATGCAGCCG |
| Human | BRPF1 | Chromatin Readers | CCAAGTTGGGAGAGATTCTG |
| Human | BRPF1 | Chromatin Readers | TGGCTGGTAAGTCCATTGGG |
| Human | BRPF1 | Chromatin Readers | CCAATGGACTTACCAGCCAA |
| Human | BRPF1 | Chromatin Readers | TTGGCTGGTAAGTCCATTGG |
| Human | BRPF1 | Chromatin Readers | ATTGGCTGGTAAGTCCATTG |
| Human | BRPF1 | Chromatin Readers | AGCATGGTAAGGAAGAGTCT |
| Human | BRPF1 | Chromatin Readers | AGTACAGATCGCCTACCACA |
| Human | BRPF1 | Chromatin Readers | AGGCGATCTGTACTGACTTG |
| Human | BRPF1 | Chromatin Readers | CAGTACAGATCGCCTACCAC |
| Human | BRPF1 | Chromatin Readers | TGCGGTGCTGCAGAGCCCTG |
| Human | BRPF1 | Chromatin Readers | GTGCTGCAGAGCCCTGTGGT |
| Human | PRMT2 | Histone Methylation | ACTGAGCTGGGTCTCATCGG |
| Human | PRMT2 | Histone Methylation | GGTGGCAGCGTAGTCCGCGA |
| Human | PRMT2 | Histone Methylation | GCGGCCTAGAGCGGTGTACG |
| Human | PRMT2 | Histone Methylation | TTTCTGATCTAGAGACCCTG |
| Human | PRMT2 | Histone Methylation | TCTGATCTAGAGACCCTGAG |
| Human | PRMT2 | Histone Methylation | TCTGCTTCCAGTGTGTGGTG |
| Human | PRMT2 | Histone Methylation | CTAAAGATTTCAGAGCGCTG |
| Human | PRMT2 | Histone Methylation | TGTCCCAGAAGTGAATCGCA |
| Human | PRMT2 | Histone Methylation | AGAGCCTGCTGAGTGCAGTG |
| Human | ARID4A | Chromatin Readers | GGATGTGACAATATTGATAG |
| Human | ARID4A | Chromatin Readers | GCTTATAGAAAGTATCTCTA |

| | | | |
|---|---|---|---|
| Human | ARID4A | Chromatin Readers | AGACTGGTTTATCATCAGGG |
| Human | ARID4A | Chromatin Readers | AAGCAAATTTATATGGACCT |
| Human | ARID4A | Chromatin Readers | TCATGGCAAACCAAGGTTTC |
| Human | ARID4A | Chromatin Readers | CAAAGATTAAGACTGTGAAA |
| Human | ARID4A | Chromatin Readers | GCAAGTTGACAGATGCTAGT |
| Human | CBX2 | Chromatin Readers | GTCCTCCAAACATAACAGCT |
| Human | CBX2 | Chromatin Readers | CTGAGCAAGCGGCTCCGCAA |
| Human | CBX2 | Chromatin Readers | GAAGAAGGAACATGAGAAGG |
| Human | CBX2 | Chromatin Readers | GAGTACCTGGTCAAGTGGCG |
| Human | CBX2 | Chromatin Readers | GGTACTCCAGCTTGCCCTTG |
| Human | CBX2 | Chromatin Readers | AGCTTCCTTGGCCGGCCTCT |
| Human | CBX2 | Chromatin Readers | GCTGAGCAGCGTGGGCGAGC |
| Human | CBX2 | Chromatin Readers | AGCATCGGGTTCCTTGAGCT |
| Human | CBX2 | Chromatin Readers | CAGCCGGCGCTCCAAGCTCA |
| Human | CBX2 | Chromatin Readers | CTGCTGGACTTGGATTTGGA |
| Human | CBX2 | Chromatin Readers | TTGGATTTGGAGGGAGCATC |
| Human | CBX2 | Chromatin Readers | CTTGGATTTGGAGGGAGCAT |
| Human | TRIM66 | Chromatin Readers | GGTAATAATGCCGGGCCAGG |
| Human | TRIM66 | Chromatin Readers | TGGTAATAATGCCGGGCCAG |
| Human | TRIM66 | Chromatin Readers | GATAATCTGGTAATAATGCC |
| Human | TRIM66 | Chromatin Readers | GCTGTGCAGCTCTTGCACAG |
| Human | TRIM66 | Chromatin Readers | TCTGCACCTACTGCAATCGC |
| Human | TRIM66 | Chromatin Readers | GCACAGCCAGCGATTGCAGT |
| Human | TRIM66 | Chromatin Readers | CAGGAACTGCTCTGAGTGCA |
| Human | TRIM66 | Chromatin Readers | TGGTGGAACACAAAGAACAC |
| Human | TRIM66 | Chromatin Readers | TTGCCATAGCTGCCTAGTGG |
| Human | TRIM66 | Chromatin Readers | GTATTGTCCTCTACACACAC |
| Human | TRIM66 | Chromatin Readers | TGTGTGTAGAGGACAATACA |
| Human | TRIM66 | Chromatin Readers | CACTTGCCATAGCTGCCTAG |
| Human | TRIM66 | Chromatin Readers | AGCCTTGCTCAGCTTCCCAG |
| Human | TRIM66 | Chromatin Readers | GCCTTGCTCAGCTTCCCAGG |
| Human | TRIM66 | Chromatin Readers | ACAGTAACTCTCCGCCATTG |
| Human | TRIM66 | Chromatin Readers | GGCTAATAGAGGAATTAGAG |
| Human | TRIM66 | Chromatin Readers | ACAGGCCAATGGGCTAATAG |
| Human | TRIM66 | Chromatin Readers | GGGCTAATAGAGGAATTAGA |
| Human | TRIM66 | Chromatin Readers | GGATTACTAATGAGAGAAAG |
| Human | TRIM66 | Chromatin Readers | GAAGCATCAGCATAGGAAGG |
| Human | TRIM66 | Chromatin Readers | CTGACTAGAAGCCAGATGCT |
| Human | TRIM66 | Chromatin Readers | GGCAGATCAACTACATCGTG |
| Human | TRIM66 | Chromatin Readers | ATCTTCAGCAAACCATTGTG |
| Human | TRIM66 | Chromatin Readers | CCATGTTCTGAGAACAGGCA |
| Human | GTF2B | Transcription Elongation | CATATTTCTTTAAATGTCCT |

| | | | |
|---|---|---|---|
| Human | GTF2B | Transcription Elongation | ATATTGCCTGTAGACAAGAA |
| Human | GTF2B | Transcription Elongation | TATGAACAGAAGAGCCTGAA |
| Human | GTF2B | Transcription Elongation | ATATGAACAGAAGAGCCTGA |
| Human | GTF2B | Transcription Elongation | GTTCGATCAACTATATTTCG |
| Human | GTF2B | Transcription Elongation | GGCAGCGGCAGCTATTTACA |
| Human | GTF2B | Transcription Elongation | TCTTCCTAAACAAGTACAGA |
| Human | GTF2B | Transcription Elongation | AGCTATTTACATGGCCTCAC |
| Human | GTF2B | Transcription Elongation | TTCAGGACAGATCATATCAC |
| Human | GTF2B | Transcription Elongation | TTAGTGGAGGACTACAGAGC |
| Human | GTF2B | Transcription Elongation | ATTCAAAGAAATCACTACCA |
| Human | CHD5 | Chromatin Readers | AGTTGCGATACATCACCGTG |
| Human | CHD5 | Chromatin Readers | CTACCAAAGAAAGAACGACA |
| Human | CHD5 | Chromatin Readers | CTGGGTGAAGGAGCTACAGC |
| Human | CHD5 | Chromatin Readers | TGACAAAGAACTCTCTCTCA |
| Human | CHD5 | Chromatin Readers | AGTTCTTTGTCAAGTGGGCA |
| Human | CHD5 | Chromatin Readers | CCATAGCTTTGACAAGAAGG |
| Human | CHD5 | Chromatin Readers | GAACCATAGCTTTGACAAGA |
| Human | CHD5 | Chromatin Readers | CTCCCAGCATCAGCTCCCTG |
| Human | CHD5 | Chromatin Readers | GCCCTACGACCAGTGCACCT |
| Human | CHD5 | Chromatin Readers | TCTACGTGGTCACCTACACG |
| Human | CHD5 | Chromatin Readers | AATGGTGATGAGCTCATAGG |
| Human | CHD5 | Chromatin Readers | GAGGGAGTAAAGGAACACGA |
| Human | CHD5 | Chromatin Readers | GAGGACAACGCCATTCGGAG |
| Human | CHD5 | Chromatin Readers | AAGGCCTGGATGTCATTGTG |
| Human | CHD5 | Chromatin Readers | CACTGTCATCATCTACGACT |
| Human | CHD5 | Chromatin Readers | GATGATGACAGTGTCCGCCG |
| Human | CHD5 | Chromatin Readers | CTGGGTCCAGGCATACGAGA |
| Human | CHD5 | Chromatin Readers | GTCCAGGCATACGAGATGGT |
| Human | CHD5 | Chromatin Readers | CCTGTGCGACACCTGCCCGA |
| Human | CHD5 | Chromatin Readers | GAGCTGGAGAAGGCTCCCGA |
| Human | CHD5 | Chromatin Readers | CATACGAGATGGTAGGCCCT |
| Human | CHD5 | Chromatin Readers | CCGCTGCCCGAGATCCCAAA |
| Human | CHD5 | Chromatin Readers | GCGGGTTGAGGCAATGCAGG |
| Human | CHD5 | Chromatin Readers | CCGAGATCCCAAACGGTGAA |
| Human | CHD5 | Chromatin Readers | TGCCGCGTGTGCAAGGACGG |
| Human | CHD5 | Chromatin Readers | GAGGCAATGCAGGTGGTAGG |
| Human | CHD5 | Chromatin Readers | GAAGAGAAGTCGGAGAGTGA |
| Human | CHD5 | Chromatin Readers | TGAGCTATCAGAGAATGAAG |
| Human | CHD5 | Chromatin Readers | GGAAGCGGAAGAAGAAAGAG |
| Human | CHD5 | Chromatin Readers | GGGAAGCGGAAGAAGAAAGA |
| Human | HR | Histone Methylation | GGCACACCATCTGGAGAAAG |
| Human | HR | Histone Methylation | GTGGTGTGTCGGCATGCACC |

| | | | |
|---|---|---|---|
| Human | HR | Histone Methylation | AAAGGAAGTCTTTCTGTGCC |
| Human | HR | Histone Methylation | GGGTGTTGAAGAGTCCATGG |
| Human | HR | Histone Methylation | TGAAGAGTCCATGGTGGCAA |
| Human | HR | Histone Methylation | ATGGACTCTTCAACACCCAC |
| Human | FXR2 | Chromatin Readers | CTTCTACAAGGGCTTTGTGA |
| Human | FXR2 | Chromatin Readers | CTTCACAAAGCCCTTGTAGA |
| Human | FXR2 | Chromatin Readers | GCCAATGAACAGGAACCTTG |
| Human | FXR2 | Chromatin Readers | ATGAACAGGAACCTTGTGGC |
| Human | FXR2 | Chromatin Readers | TTGTTACCCTGGAGCGACTT |
| Human | FXR2 | Chromatin Readers | CTACAATGAAATTGTTACCC |
| Human | FXR2 | Chromatin Readers | TTATTCTCGAGCCAATGAAC |
| Human | FXR2 | Chromatin Readers | TCCCAATTGCCAGTCCCATC |
| Human | FXR2 | Chromatin Readers | GGACTGGCAATTGGGACTCA |
| Human | FXR2 | Chromatin Readers | GCGAGAGGACCTGATGGGAC |
| Human | FXR2 | Chromatin Readers | ACAGTGCGAGAGGACCTGAT |
| Human | FXR2 | Chromatin Readers | AACCTGGTTGGCAAAGTGAT |
| Human | FXR2 | Chromatin Readers | GAAAGTGATCCAGGAGATTG |
| Human | FXR2 | Chromatin Readers | CAAGTGCCCAGGAACCTGGT |
| Human | FXR2 | Chromatin Readers | ACTTTGCCAACCAGGTTCCT |
| Human | FXR2 | Chromatin Readers | TTCCAATCACTTTGCCAACC |
| Human | POLR1B | Transcription Elongation | TTGCGCATGGCAGACCAAGA |
| Human | POLR1B | Transcription Elongation | GCGCATGGCAGACCAAGAAG |
| Human | POLR1B | Transcription Elongation | TGCGCATGGCAGACCAAGAA |
| Human | PHF10 | Chromatin Readers | GATGGCCACTATTCTCACAT |
| Human | PHF10 | Chromatin Readers | ATTGTCATATCCAGGCAAGA |
| Human | PHF10 | Chromatin Readers | CTATGATTAAGACCTACCCA |
| Human | PHF10 | Chromatin Readers | TTCTTGCCTGGATATGACAA |
| Human | PHF10 | Chromatin Readers | TTCCATACACTGCCATGGGT |
| Human | PHF10 | Chromatin Readers | GACCTGATGGAATAGCACCA |
| Human | PHF10 | Chromatin Readers | TCACAAATCCAGCGACCTGA |
| Human | PHF10 | Chromatin Readers | TTCTGTGATATGTGTGACAG |
| Human | PHF10 | Chromatin Readers | ATGTTATCCAGGTACCTCAA |
| Human | PHF10 | Chromatin Readers | CTTGCAGACACATGTTATCC |
| Human | PHF10 | Chromatin Readers | CATGTTATCCAGGTACCTCA |
| Human | PHF10 | Chromatin Readers | CCTCTGGCAATGTATCTGAA |
| Human | PHF10 | Chromatin Readers | CTCTGGCAATGTATCTGAAG |
| Human | PHF10 | Chromatin Readers | TCACCATCACTGTCTAGAGC |
| Human | PHF10 | Chromatin Readers | TCCTCTGGCAATGTATCTGA |
| Human | HDAC10 | Histone Acetylation | GCTACACAGGATCCTCGTCG |
| Human | HDAC10 | Histone Acetylation | GTGCCCGGGAGGCCTCACCA |
| Human | HDAC10 | Histone Acetylation | GGGTCAAACTCAAAGGCCAG |
| Human | HDAC10 | Histone Acetylation | CGCAGTGAAAGGTACTCGGG |

| | | | |
|---|---|---|---|
| Human | HDAC10 | Histone Acetylation | CAGCTCAGGGTCAAACTCAA |
| Human | SFMBT2 | Chromatin Readers | CTGGTGTGACGTAGTCATCG |
| Human | SFMBT2 | Chromatin Readers | CCCGCACGTGGTAATGATCG |
| Human | SFMBT2 | Chromatin Readers | CTCGCAGAGGACCTTCCAGG |
| Human | SFMBT2 | Chromatin Readers | CTCCTGGAAGGTCCTCTGCG |
| Human | SFMBT2 | Chromatin Readers | TCCTGGAAGGTCCTCTGCGA |
| Human | SFMBT2 | Chromatin Readers | GATTACAGACTTCGACCAGT |
| Human | SFMBT2 | Chromatin Readers | TGATCCTTAAACACTTCCAT |
| Human | SFMBT2 | Chromatin Readers | TGTGCCATGCAGATTCTTTG |
| Human | SFMBT2 | Chromatin Readers | GATGTATTTCGGAAGCAGAA |
| Human | SFMBT2 | Chromatin Readers | AGATTATCACAAGCAGCATG |
| Human | SFMBT2 | Chromatin Readers | TGATGTATTTCGGAAGCAGA |
| Human | SFMBT2 | Chromatin Readers | GCACTCCTGCACCGTCGGAA |
| Human | SFMBT2 | Chromatin Readers | CTGAAATATCTTGGCCAAGG |
| Human | SFMBT2 | Chromatin Readers | TCCATGCACTCCTGCACCGT |
| Human | SFMBT2 | Chromatin Readers | CCTGCTTCTGACCCTTCCGA |
| Human | HDGFRP2 | Chromatin Readers | TGCCAGGATCGACGACATCG |
| Human | HDGFRP2 | Chromatin Readers | GCGATGTCGTCGATCCTGGC |
| Human | HDGFRP2 | Chromatin Readers | CGATCCTGGCAGGCCAGTGA |
| Human | HDGFRP2 | Chromatin Readers | ATCCGCGATGTCGTCGATCC |
| Human | HDGFRP2 | Chromatin Readers | ACACACGAAACAGCCTTCCT |
| Human | HDGFRP2 | Chromatin Readers | GGTCCTCGGCCCAAAGATCG |
| Human | HDGFRP2 | Chromatin Readers | TCGCCGTTACAAAGCGAACA |
| Human | HDGFRP2 | Chromatin Readers | AGGCAGCAGAAGTCTATACC |
| Human | HDGFRP2 | Chromatin Readers | CCTCGGCCCAAAGATCGAGG |
| Human | HDGFRP2 | Chromatin Readers | TCTATACCCGGCTCAAGTCG |
| Human | HDGFRP2 | Chromatin Readers | GACGCCTGCGCTAAAGATGT |
| Human | HDGFRP2 | Chromatin Readers | GACAAGAGTAGCGACAACAG |
| Human | HDGFRP2 | Chromatin Readers | GAGACCGACATCTTTAGCGC |
| Human | HDGFRP2 | Chromatin Readers | GATGAGCCCGTCAAGAAGCG |
| Human | HDGFRP2 | Chromatin Readers | ACGATGAGCCCGTCAAGAAG |
| Human | CARM1 | Histone Methylation | TCGGCTCCATAATGACCGTG |
| Human | CARM1 | Histone Methylation | TGATTCCTTTAGCCAACACG |
| Human | CARM1 | Histone Methylation | CCTGATTCCTTTAGCCAACA |
| Human | CDK2 | Other Histone Modifications | AACAGATAGCTCTTGATGAG |
| Human | CDK2 | Other Histone Modifications | TCGTACTTACACCCATGAGG |
| Human | CDK2 | Other Histone Modifications | CCTCAGTCTCAGTGTCCAGG |
| Human | CDK2 | Other Histone Modifications | GGAACAGATAGCTCTTGATG |
| Human | CDK2 | Other Histone Modifications | CGCCTGGACACTGAGACTGA |
| Human | CCDC101 | Chromatin Readers | CCATGCCACCAACAAGTATG |
| Human | CCDC101 | Chromatin Readers | TGTGGCTCAGAGATACGTGG |
| Human | CCDC101 | Chromatin Readers | CGGTAGAAGCAGGTAGTCTG |

| | | | |
|---|---|---|---|
| Human | CCDC101 | Chromatin Readers | CAATGTGGCTCAGAGATACG |
| Human | CCDC101 | Chromatin Readers | TATCTCTGAGCCACATTGAG |
| Human | CCDC101 | Chromatin Readers | TATTACCGGACAAAGCTGCG |
| Human | CCDC101 | Chromatin Readers | ACATCCAGAAGACCCATGAG |
| Human | CCDC101 | Chromatin Readers | AAACCCAGGAAGAGCGTTCG |
| Human | CCDC101 | Chromatin Readers | AGAGGCCACGCAGCTTTGTC |
| Human | CCDC101 | Chromatin Readers | AACCGCCAAGGCCGATGCAG |
| Human | CCDC101 | Chromatin Readers | GATGACCCTGCTGCAGCAGT |
| Human | YWHAZ | Chromatin Readers | GCTGGTTCAGAAGGCCAAAC |
| Human | YWHAZ | Chromatin Readers | AGCCCGTAGGTCATCTTGGA |
| Human | YWHAZ | Chromatin Readers | GACCCTCCAAGATGACCTAC |
| Human | YWHAZ | Chromatin Readers | GAGCCCGTAGGTCATCTTGG |
| Human | YWHAZ | Chromatin Readers | CGACCCTCCAAGATGACCTA |
| Human | YWHAZ | Chromatin Readers | AGCTGTCTTTGCAAGAGAGC |
| Human | PHF1 | Histone Methylation | GGTGTGTCTGGTCCAGTTTG |
| Human | PHF1 | Histone Methylation | ACTGATGGGCTGCTATACTT |
| Human | PHF1 | Histone Methylation | GGCAGTCCTGGTGATAAGCA |
| Human | PHF1 | Histone Methylation | CAGAGCGACAGACACAACAG |
| Human | PHF1 | Histone Methylation | GTGTCGCCATGCTTATCACC |
| Human | PHF1 | Histone Methylation | TGTCTGTCGCTCTGAGACTG |
| Human | PHF1 | Histone Methylation | ACTTCTCACAGCTGACCAGC |
| Human | PHF1 | Histone Methylation | AGGCCTCATGGAACCACTGC |
| Human | PHF1 | Histone Methylation | GAGGTGATACAGGACAAGAT |
| Human | PHF1 | Histone Methylation | TGAGGTGATACAGGACAAGA |
| Human | ELL2 | Transcription Elongation | TCTAGCTACAGTCTCCATCC |
| Human | ELL2 | Transcription Elongation | CAGAGCTTTGCATGCCAGGA |
| Human | ELL2 | Transcription Elongation | AGTACAGAGCTTTGCATGCC |
| Human | ELL2 | Transcription Elongation | TGAAGTCATCCTTATAATTC |
| Human | ELL2 | Transcription Elongation | ACAAGCTGGCTCACATCAAA |
| Human | ELL2 | Transcription Elongation | TGAATATCTTCATAACAAGC |
| Human | ELOB | Transcription Elongation | AGCGGCCTCCTGACGAGCAG |
| Human | ELOB | Transcription Elongation | TCGTCGAGGGCATCCTCAAG |
| Human | ELOB | Transcription Elongation | GAACTGAAGCGCATCGTCGA |
| Human | ELOB | Transcription Elongation | CGAACTGAAGCGCATCGTCG |
| Human | ELOB | Transcription Elongation | CTTGGCGTCCGTGAAGATGG |
| Human | HDGFL1 | Chromatin Readers | AGTTAAAGGGCTATGCCCAC |
| Human | HDGFL1 | Chromatin Readers | ACTCAGGAAGGCCGTCTCGT |
| Human | HDGFL1 | Chromatin Readers | GAACGGACTCTCCTCGTCGA |
| Human | HDGFL1 | Chromatin Readers | GGAGGAAGAAGTCGCGGACG |
| Human | HDGFL1 | Chromatin Readers | GGAGAGTCCGTTCCTCGTGG |
| Human | HDGFL1 | Chromatin Readers | CGAGGAGAGTCCGTTCCTCG |
| Human | ELP2 | Transcription Elongation | GAGTCCTGAACTGGAGCTCT |

| | | | |
|---|---|---|---|
| Human | ELP2 | Transcription Elongation | TGATAATCAAGTGATTCACT |
| Human | ELP2 | Transcription Elongation | CCACTGTATGCAATTGACTC |
| Human | ELP2 | Transcription Elongation | CTGTATGCAATTGACTCGGG |
| Human | ELP2 | Transcription Elongation | TCCACTGTATGCAATTGACT |
| Human | ELP2 | Transcription Elongation | CTGATAATCAAGTGATTCAC |
| Human | ELP2 | Transcription Elongation | CAGATTCTGCTGTTCGACTC |
| Human | ELP2 | Transcription Elongation | ACGATCAGTGTACATAATGC |
| Human | ELP2 | Transcription Elongation | ACAGCATGCACCGCATAAAC |
| Human | ELP2 | Transcription Elongation | TGCATGCTGTTTACCAGAGG |
| Human | ELP2 | Transcription Elongation | CGGTGCATGCTGTTTACCAG |
| Human | ELP2 | Transcription Elongation | GATGTACCAATATTAGCATG |
| Human | ELP2 | Transcription Elongation | AATATTGGTACATCAGTATT |
| Human | ELP2 | Transcription Elongation | TCATTGCCACATGCTAATAT |
| Human | ELP2 | Transcription Elongation | ATTGGATTAGAGGAGTGGAA |
| Human | ELP2 | Transcription Elongation | TGAGGATTGGATTAGAGGAG |
| Human | ELP2 | Transcription Elongation | ACAGCTTCCATATTCTTATC |
| Human | ELP2 | Transcription Elongation | AAGATTGCCTGATAAGAATA |
| Human | ELP2 | Transcription Elongation | ACTGGGTAAATGCAGTTCAC |
| Human | ELP2 | Transcription Elongation | GAAGCAGATAATAATCTCAC |
| Human | ELP2 | Transcription Elongation | TGGGCTCCAGATGAAGAGTC |
| Human | ELP2 | Transcription Elongation | GAAAGCATGAGCAATGATCA |
| Human | ELP2 | Transcription Elongation | TAACTGTATTCTGTTTCCAA |
| Human | ELP2 | Transcription Elongation | ATCATTGCTCATGCTTTCCA |
| Human | ELP2 | Transcription Elongation | GATTGCCAGTTCAATGAAGA |
| Human | ELP2 | Transcription Elongation | TGGTGTCCAAGACCTAGTCT |
| Human | ELP2 | Transcription Elongation | ATGGTGTCCAAGACCTAGTC |
| Human | ELP2 | Transcription Elongation | GACCTAGTCTGGGATCCAGA |
| Human | ELP2 | Transcription Elongation | CTCCTTCTGGATCCCAGACT |
| Human | ELP2 | Transcription Elongation | TGGGTATGACCTGAAATGTT |
| Human | ELP2 | Transcription Elongation | AATCGGTTTCAGTTTGTATC |
| Human | ELP2 | Transcription Elongation | AATGTTTGGCAATGATTAAT |
| Human | ELP2 | Transcription Elongation | CTTAGCTGCCTTACAAGCTG |
| Human | ELP2 | Transcription Elongation | GAACACTACATCTTGGAAAC |
| Human | ELP2 | Transcription Elongation | AGCATGCAGCTATCATTCTT |
| Human | ELP2 | Transcription Elongation | TTCTTTGGAACACTACATCT |
| Human | ELP2 | Transcription Elongation | TCTGCTTGCCTCAGCTTGTA |
| Human | ELP2 | Transcription Elongation | CTGTTTCCAGAGATCGAACC |
| Human | ELP2 | Transcription Elongation | GAGATCGAACCTGGTCATTG |
| Human | ELP2 | Transcription Elongation | CAATGACCAGGTTCGATCTC |
| Human | ELP2 | Transcription Elongation | ACAGCAAGTATTTCTTCACT |
| Human | ELP2 | Transcription Elongation | CCACCCTTCTCAACGATACG |
| Human | ELP2 | Transcription Elongation | CCACGTATCGTTGAGAAGGG |

| | | | |
|---|---|---|---|
| Human | ELP2 | Transcription Elongation | GCAACCACGTATCGTTGAGA |
| Human | ELP2 | Transcription Elongation | CAACCACGTATCGTTGAGAA |
| Human | ELP2 | Transcription Elongation | GTTGCAGTAGGATTGGAGTG |
| Human | ELP2 | Transcription Elongation | TGGTTACACTTTGCAAGCTG |
| Human | ELP2 | Transcription Elongation | AGGAAGCAGAAGGTGCTGAG |
| Human | ELP2 | Transcription Elongation | ACTGAACAGAAGGAAGCAGA |
| Human | ELP2 | Transcription Elongation | TTATGCTGGAAGAATTGCAG |
| Human | SHPRH | Chromatin Readers | CTCTCGCCATAAGAAGTGCA |
| Human | SHPRH | Chromatin Readers | TGCAGGGCACTTTCAGTAGC |
| Human | SHPRH | Chromatin Readers | GGGTCTTGGAAAGACAGTGG |
| Human | SHPRH | Chromatin Readers | CGAGAGATTGTTACATCTGA |
| Human | SHPRH | Chromatin Readers | AGTATCATCACACGTGTAGG |
| Human | SHPRH | Chromatin Readers | GTCAGTATCATCACACGTGT |
| Human | SHPRH | Chromatin Readers | GTATCATCACACGTGTAGGA |
| Human | SHPRH | Chromatin Readers | GGTCTCGTGAAACAGATCAA |
| Human | SHPRH | Chromatin Readers | AGAACTGACATACTTATAGA |
| Human | SHPRH | Chromatin Readers | ATTAATCGATGGTGTATCAG |
| Human | SHPRH | Chromatin Readers | AAATGGCCCAGCGTTTGAGT |
| Human | SHPRH | Chromatin Readers | ATTAATCCCACTCAAACGCT |
| Human | SHPRH | Chromatin Readers | GTGAGGATGGGCGTCGCCTA |
| Human | SHPRH | Chromatin Readers | ATGAGATCAACAGGCATGTG |
| Human | SHPRH | Chromatin Readers | AATTCGGTGCACCCTCCCTA |
| Human | SHPRH | Chromatin Readers | AGCTCTTACTGACAACAACA |
| Human | SHPRH | Chromatin Readers | ATGAGCTTCAGGCCATAGGG |
| Human | SHPRH | Chromatin Readers | CACACTTTGCATGTTGCCAC |
| Human | SHPRH | Chromatin Readers | CACTTCAGGCATTGAACACG |
| Human | SHPRH | Chromatin Readers | AATGCCTGAAGTGTCACCTG |
| Human | SHPRH | Chromatin Readers | GTTGCCACAGGTGACACTTC |
| Human | SHPRH | Chromatin Readers | AATTATTGAACAATACAGCG |
| Human | SHPRH | Chromatin Readers | ATTACAGAAACAGTGACCAC |
| Human | SHPRH | Chromatin Readers | CAGCTAGTTTGTGCTCTCAA |
| Human | SHPRH | Chromatin Readers | ACATCTTTGCAGAAGAAATG |
| Human | SHPRH | Chromatin Readers | TGTGCTCTCAATGGCTTAGC |
| Human | SHPRH | Chromatin Readers | TCTGTACAATTCTGCTGCCA |
| Human | SHPRH | Chromatin Readers | TCATGCTACCCATAACTTGA |
| Human | SHPRH | Chromatin Readers | GTAGTGCTCTCGCAGCTGTT |
| Human | SHPRH | Chromatin Readers | TCATGGATGGTCTGCTGCAC |
| Human | SHPRH | Chromatin Readers | ATGCCAGAAGCTAGTAAGAG |
| Human | SHPRH | Chromatin Readers | GTTCTTACTTACAACACAAA |
| Human | SHPRH | Chromatin Readers | GTTGCTTCATGAATATTGGA |
| Human | SHPRH | Chromatin Readers | ATGAATATTGGATGGCTCTG |
| Human | SHPRH | Chromatin Readers | TCTCTTTGAAGCATGGAAGA |

| | | | |
|---|---|---|---|
| Human | SHPRH | Chromatin Readers | CAATGGATCTCTTTGAAGCA |
| Human | SHPRH | Chromatin Readers | TGATGAAGGAAGCACTTCAA |
| Human | TDRD3 | Chromatin Readers | AGGCAAGCTCTTATGGATAA |
| Human | TDRD3 | Chromatin Readers | GCTTCAGTAAGGAAGCATCG |
| Human | TDRD3 | Chromatin Readers | GAAAGCTCTGAAGCACATAA |
| Human | TDRD3 | Chromatin Readers | AGCATCGAGGCAAGCTCTTA |
| Human | TDRD3 | Chromatin Readers | CTGAAGCACATAACGGAAAT |
| Human | TDRD3 | Chromatin Readers | TGACTACGGAAACTATGAAG |
| Human | TDRD3 | Chromatin Readers | TATCAAGCCCATTCAAACAG |
| Human | TDRD3 | Chromatin Readers | TGTCATACCCGAAGAATGGA |
| Human | TDRD3 | Chromatin Readers | GTTGGCCGAGTGGATCGTCT |
| Human | TDRD3 | Chromatin Readers | AAACTGTTGGGTTGGCCGAG |
| Human | RECQL5 | Transcription Elongation | TGGCGGTCAGAGCCACACAA |
| Human | RECQL5 | Transcription Elongation | GTACCTTTAGGGTTAGCAAG |
| Human | RECQL5 | Transcription Elongation | TGCTTTGATTCAGGACCAAG |
| Human | RECQL5 | Transcription Elongation | TCCTGAATCAAAGCAATGAG |
| Human | RECQL5 | Transcription Elongation | CCACCACCAAGTAAGACAGC |
| Human | RECQL5 | Transcription Elongation | ATGGGCGACAAACCTGACAT |
| Human | RECQL5 | Transcription Elongation | ACGCCAAGGCTTACCATGCA |
| Human | RECQL5 | Transcription Elongation | TAATGGTTGCAACAATTACA |
| Human | RECQL5 | Transcription Elongation | ATGTCAGGTTTGTCGCCCAT |
| Human | RECQL5 | Transcription Elongation | CCGGAGGTCAGCTTCATCAG |
| Human | RECQL5 | Transcription Elongation | CAGCGGTACGTGTTGACTGG |
| Human | RECQL5 | Transcription Elongation | GACATTCCGGAACGCCAAGG |
| Human | RECQL5 | Transcription Elongation | CCAGGCTGACTGTGAAGGCA |
| Human | RECQL5 | Transcription Elongation | CGGAACTGATGGAGACAACT |
| Human | RECQL5 | Transcription Elongation | GATGGAGACAACTCGGATCA |
| Human | RECQL5 | Transcription Elongation | CTGCCCGTTCCAGACGGCCA |
| Human | RECQL5 | Transcription Elongation | CCAGACGGCCACGGAACTGA |
| Human | RECQL5 | Transcription Elongation | AGTTGTCTCCATCAGTTCCG |
| Human | RECQL5 | Transcription Elongation | GGCATCAAAGGCCATGATAG |
| Human | RECQL5 | Transcription Elongation | TATCATGGCCTTTGATGCCC |
| Human | RECQL5 | Transcription Elongation | GGCCTCCAGCCGCCTCCGCA |
| Human | RECQL5 | Transcription Elongation | GGCCGTGCGGAGGCGGCTGG |
| Human | RECQL5 | Transcription Elongation | GCGGAGGCGGCTGGAGGCCT |
| Human | PHF19 | Chromatin Readers | TGGAAGGACATACAGCATGC |
| Human | PHF19 | Chromatin Readers | ACAGATGGCCTGTACTACCT |
| Human | PHF19 | Chromatin Readers | AGGGCCAGTATGTGCTGTGC |
| Human | PHF19 | Chromatin Readers | TGACCCTCTTGATCTTCCCG |
| Human | PHF19 | Chromatin Readers | CAGATGAGGATCTCATTCAG |
| Human | PHF19 | Chromatin Readers | ATGCAGCGTCGGCAGAACCA |
| Human | PHF19 | Chromatin Readers | CCAAGTGCAACATCTGCCTA |

| | | | |
|---|---|---|---|
| Human | PHF19 | Chromatin Readers | GGCACTGCTGGTGGTAACCC |
| Human | PHF19 | Chromatin Readers | TCTGCCTAGGGAAGACATCA |
| Human | PHF19 | Chromatin Readers | GGCCCGGAGAATGGTACCTG |
| Human | PHF19 | Chromatin Readers | ATCCGCAGGTACCATTCTCC |
| Human | PHF19 | Chromatin Readers | GTGCAGGCAGTGGTTCCACG |
| Human | PHF19 | Chromatin Readers | AACCGGTCTCCAAACATCAT |
| Human | PRDM1 | Histone Methylation | TTGGAGGATCTATTCCAGAG |
| Human | PRDM1 | Histone Methylation | TTTGGAGGATCTATTCCAGA |
| Human | PRDM1 | Histone Methylation | TCCAATAACCTCTTCACTGT |
| Human | PRDM1 | Histone Methylation | GGACCTTCAGATTGGAGAGC |
| Human | PRDM1 | Histone Methylation | AACGTTTGCGCCAAGACTTT |
| Human | PRDM1 | Histone Methylation | GGAGAGCTGGCCGAAAGTCT |
| Human | PRDM1 | Histone Methylation | CGGCCAGCTCTCCAATCTGA |
| Human | PRDM1 | Histone Methylation | TAAATCTCTTGTGGCAGACC |
| Human | PRDM1 | Histone Methylation | TGGTGCTGCTAAATCTCTTG |
| Human | PRDM1 | Histone Methylation | CAGGTGGGTCTTGAGATTGC |
| Human | PRDM1 | Histone Methylation | CAGTTTCAGGTGCACAAACT |
| Human | PRDM1 | Histone Methylation | GTGCACAAACTGGGTGAACT |
| Human | PRDM1 | Histone Methylation | CAAACTGGGTGAACTTGGCA |
| Human | PRDM1 | Histone Methylation | GCAGTTTCAGGTGCACAAAC |
| Human | PRDM1 | Histone Methylation | ACAAACTGGGTGAACTTGGC |
| Human | PRDM1 | Histone Methylation | GTAGGGAAGGGTCTTGTAGC |
| Human | PRDM1 | Histone Methylation | TACCCGCTGAAGAAGCAGAA |
| Human | PRDM1 | Histone Methylation | CAAAGCTACCTCAGCAGCGA |
| Human | PRDM1 | Histone Methylation | TTCATGGCTTCGTCGCTGCT |
| Human | POLR2I | Transcription Elongation | GGCACGCGTAGAGCAGAATG |
| Human | POLR2I | Transcription Elongation | GCATTCTGCTCTACGCGTGC |
| Human | POLR2I | Transcription Elongation | GGTTCTCCTTGTCTTCCTTG |
| Human | POLR2I | Transcription Elongation | GTAAAGGCGCATGGCGTCCT |
| Human | EP300 | Histone Acetylation | TGAATCTGCAAACAATCGAG |
| Human | EP300 | Histone Acetylation | CCAAGCCATATTTCCTACGC |
| Human | EP300 | Histone Acetylation | GTGGCACGAAGATATTACTC |
| Human | EP300 | Histone Acetylation | CCGGCGTAGGAAATATGGCT |
| Human | EP300 | Histone Acetylation | AGGGTCCACAGGTTGACGAA |
| Human | EP300 | Histone Acetylation | TCACAATATCAAAGTAATCA |
| Human | EP300 | Histone Acetylation | ATATCATCGACATACTGCCA |
| Human | EP300 | Histone Acetylation | TAATGGTAGAAAGATCCATG |
| Human | EP300 | Histone Acetylation | AGAAATTAGGTTACACAACA |
| Human | EP300 | Histone Acetylation | GAAAGCACAGATGTGACCAA |
| Human | EP300 | Histone Acetylation | AAAGCAAGGTTTGTGGACAG |
| Human | EP300 | Histone Acetylation | TGAAAGCACAGATGTGACCA |
| Human | EP300 | Histone Acetylation | GACCATCCATCAGATCGCAG |

| | | | |
|---|---|---|---|
| Human | EP300 | Histone Acetylation | GTCAGGCAAGTCTTGCCAAG |
| Human | EP300 | Histone Acetylation | AAGATGCACAGTGTGCCACT |
| Human | EP300 | Histone Acetylation | ATTCTTCATTGTGCGACAGT |
| Human | EP300 | Histone Acetylation | CATTCTTCATTGTGCGACAG |
| Human | EP300 | Histone Acetylation | CAATGAATGCAAGCACCATG |
| Human | EP300 | Histone Acetylation | AGCACCATGTGGAGACACGC |
| Human | EP300 | Histone Acetylation | CTGGCACTGTACTGTCTGTG |
| Human | EP300 | Histone Acetylation | ATGGTGCTTGCATTCATTGC |
| Human | EP300 | Histone Acetylation | CGGGTTGTGCAGCATACCAA |
| Human | EP300 | Histone Acetylation | TGCAGCGCTGGATACTCAGG |
| Human | EP300 | Histone Acetylation | AGCATACCAAGGGTTGCAAA |
| Human | EP300 | Histone Acetylation | ACAGAATTGGGACTAACCAA |
| Human | EP300 | Histone Acetylation | GAATTGGGACTAACCAATGG |
| Human | EP300 | Histone Acetylation | GCTTGACTTCTCCCAACATG |
| Human | EP300 | Histone Acetylation | TGGTGGCCCAGGTCAAGTCA |
| Human | EP300 | Histone Acetylation | TCAGGCGGCGAGAATCGCCT |
| Human | EP300 | Histone Acetylation | TGCTCTCATCATCTAAGCCA |
| Human | EP300 | Histone Acetylation | ATGTTTAGGCAGAACGGCAC |
| Human | EP300 | Histone Acetylation | TGTTTAGGCAGAACGGCACC |
| Human | EP300 | Histone Acetylation | TGATGAGAGCAACAACCAGC |
| Human | EP300 | Histone Acetylation | GCATGGTCAACCTTTGAACA |
| Human | EP300 | Histone Acetylation | AAGATGTCTCGGAATTGTGA |
| Human | EP300 | Histone Acetylation | TGGCTGCATTCATCAAGCAG |
| Human | EP300 | Histone Acetylation | GGTATCAGCCCACTCAAACC |
| Human | EP300 | Histone Acetylation | GCTGTCGTCTCAAGATGTCT |
| Human | EP300 | Histone Acetylation | GGTGGGTAGCCAACTCCTTG |
| Human | EP300 | Histone Acetylation | CACATGCAACAGATGCAACA |
| Human | EP300 | Histone Acetylation | TTGCTGGAACTGGTTATGGT |
| Human | EP300 | Histone Acetylation | GGCACTGATGCCAACTTTGG |
| Human | EP300 | Histone Acetylation | ACAGGCACTGATGCCAACTT |
| Human | EP300 | Histone Acetylation | ATACTGCTCCAAGCTCTCTG |
| Human | EP300 | Histone Acetylation | TTCAAAGACCTCAGAGAGCT |
| Human | LEO1 | Transcription Elongation | TTCTGAGCGTTCTGACCATG |
| Human | LEO1 | Transcription Elongation | CATAAATCAGAATCTGCAAG |
| Human | LEO1 | Transcription Elongation | AGGCCTCAGATGAAGAACAC |
| Human | LEO1 | Transcription Elongation | TGTGCTGATCTACATCTGAG |
| Human | AARS2 | Control | GGCTGCGGCGGGCCATTCGA |
| Human | AARS2 | Control | CAGTGGCAGCTGCAGCCCGG |
| Human | AARS2 | Control | GAATGGCCCGCCGCAGCCTC |
| Human | AARS2 | Control | CAGCTGCAGCCCGGAGGCTG |
| Human | AARS2 | Control | AATGGCCCGCCGCAGCCTCC |
| Human | NUF2 | Chromatin Remodeling | TTGGCAATATAAATCCTCTG |

| | | | |
|---|---|---|---|
| Human | NUF2 | Chromatin Remodeling | AAATGCCAGACAAGAAGTGG |
| Human | NUF2 | Chromatin Remodeling | ATACAAACGCACAGTAATTG |
| Human | NUF2 | Chromatin Remodeling | CTTCTCATCTATCTTAGCAT |
| Human | NUF2 | Chromatin Remodeling | CAAGTTTAGAAATATTTCCT |
| Human | NUF2 | Chromatin Remodeling | AGAAGACAGCTGAACTGAAG |
| Human | SMARCC1 | Chromatin Readers | AGATGTGTGTGCTGTGATGA |
| Human | SMARCC1 | Chromatin Readers | GTGCTAGTTAAATACTCTTG |
| Human | SMARCC1 | Chromatin Readers | TGCTAGTTAAATACTCTTGG |
| Human | SMARCC1 | Chromatin Readers | TTATTCCTAGTTATGCATCA |
| Human | SMARCC1 | Chromatin Readers | TCAAACCATGATGCATAACT |
| Human | SMARCC1 | Chromatin Readers | CAGGGCCTCCAGGAGTAGAA |
| Human | SMARCC1 | Chromatin Readers | GGAGACCCTTCTACTCCTGG |
| Human | SMARCC1 | Chromatin Readers | AAGAAGCTCTAGAACAACAG |
| Human | SMARCC1 | Chromatin Readers | AGGCTGCTGACCATCAGGGT |
| Human | SMARCC1 | Chromatin Readers | GAAGCTTGAAGGAGCTGAAG |
| Human | SMARCC1 | Chromatin Readers | ACAGGATAGTGAAGTGAGTG |
| Human | SMARCC1 | Chromatin Readers | CTGGTGGTGCATCAGAGGGT |
| Human | SMARCC1 | Chromatin Readers | CATGCCGTTAGGTGCTGTCA |
| Human | SMARCC1 | Chromatin Readers | GTTGATGAGGCATCATGCCA |
| Human | SMARCC1 | Chromatin Readers | CTGCAACAGTGGGAATCATG |
| Human | SMARCD2 | Chromatin Remodeling | GTGCTTGATGTAAAGCCACA |
| Human | SMARCD2 | Chromatin Remodeling | GTCGCTGGTACTGCGCCGCG |
| Human | SMARCD2 | Chromatin Remodeling | AGGTCGCTGGTACTGCGCCG |
| Human | SMARCD2 | Chromatin Remodeling | GCGGCGCAGTACCAGCGACC |
| Human | SMARCD2 | Chromatin Remodeling | CGGAATGGATCCATCATGGT |
| Human | SMARCD2 | Chromatin Remodeling | TCGGAATGGATCCATCATGG |
| Human | BRD1 | Chromatin Readers | GGAAACGGTTAGAAGCTCAA |
| Human | BRD1 | Chromatin Readers | GCTGCGCAAATATCCTGGCG |
| Human | BRD1 | Chromatin Readers | TGGTACCTCCTTCAGACTCA |
| Human | BRD1 | Chromatin Readers | GGGCTGCGCAAATATCCTGG |
| Human | BRD1 | Chromatin Readers | GTCGATGATCAGTGCCGGGT |
| Human | BRD1 | Chromatin Readers | TGATCAGTGCCGGGTAGGAG |
| Human | BRD1 | Chromatin Readers | CGTGCACCAGGAGTGCTACG |
| Human | BRD1 | Chromatin Readers | GTGCTGCATCTGCATGGACG |
| Human | BRD1 | Chromatin Readers | GTCCATGCAGATGCAGCACA |
| Human | BRD1 | Chromatin Readers | GTCACGTGGTGTGTGCCCTG |
| Human | BRD1 | Chromatin Readers | AAAGACAGATGACGACCGCT |
| Human | BRD1 | Chromatin Readers | AGCGGTCTTTCTGACGGAGA |
| Human | BRD1 | Chromatin Readers | CCACGCCCTTCTGCTTACAG |
| Human | BRD1 | Chromatin Readers | GAGCCCGTGAAGGAACTGAC |
| Human | BRD1 | Chromatin Readers | ACAGTAAGCGGTCTTTCTGA |
| Human | BRD1 | Chromatin Readers | CCTTCTGGGCACACGTCACA |

| | | | |
|---|---|---|---|
| Human | BRD1 | Chromatin Readers | GGTGCTGCTGCGCTCAGTGC |
| Human | BRD1 | Chromatin Readers | GTGAGGCTGCGCGATCAGGG |
| Human | BRD1 | Chromatin Readers | CTGATCGCGCAGCCTCACCG |
| Human | BRD1 | Chromatin Readers | GCGGTGAGGCTGCGCGATCA |
| Human | BRD1 | Chromatin Readers | CGCGGTGAGGCTGCGCGATC |
| Human | BRD1 | Chromatin Readers | TCTGAGGCAGGCCCGGCGCG |
| Human | BRD1 | Chromatin Readers | AAGCGATCCGCACGGCCTTC |
| Human | MIS18A | Chromatin Remodeling | TACAGGAAACACAGCGAAGC |
| Human | MIS18A | Chromatin Remodeling | TTTCAATGGCTTCAACACTG |
| Human | L3MBTL4 | Chromatin Readers | TCCTACTGGATGAATGTCAG |
| Human | L3MBTL4 | Chromatin Readers | GAAAGCACACAGAATACCGA |
| Human | L3MBTL4 | Chromatin Readers | ATCCTACTGGATGAATGTCA |
| Human | L3MBTL4 | Chromatin Readers | ACCTGAAATTCTTTAGACAT |
| Human | L3MBTL4 | Chromatin Readers | AGCCCTTATGTCCAGCCAGT |
| Human | L3MBTL4 | Chromatin Readers | ATCTTCAACAATATCTGCTA |
| Human | L3MBTL4 | Chromatin Readers | TATGGTCGCCACACACACCA |
| Human | L3MBTL4 | Chromatin Readers | GTCGCCACACACACCAAGGA |
| Human | L3MBTL4 | Chromatin Readers | GTAGCAACACGAATTAACCT |
| Human | L3MBTL4 | Chromatin Readers | TAGCAACACGAATTAACCTG |
| Human | L3MBTL4 | Chromatin Readers | TTTAAGAAAGAGCAGATCGA |
| Human | L3MBTL4 | Chromatin Readers | AGTTGTAAATCTTCAGTGCT |
| Human | L3MBTL4 | Chromatin Readers | AAAGTGATGAAGATCAAACT |
| Human | L3MBTL4 | Chromatin Readers | CTTCCTGCTTCTGACACAGA |
| Human | L3MBTL4 | Chromatin Readers | CGTCTGTGTCAGAAGCAGGA |
| Human | L3MBTL4 | Chromatin Readers | GTAGGACAGACAGCTTGACC |
| Human | L3MBTL4 | Chromatin Readers | AATAACGTGGACCACGGATA |
| Human | L3MBTL4 | Chromatin Readers | GTCCCGAATAACGTGGACCA |
| Human | TCEA2 | Transcription Elongation | GTGACCAAGAAGAGCGCGGA |
| Human | TCEA2 | Transcription Elongation | GGTGACCAAGAAGAGCGCGG |
| Human | TCEA2 | Transcription Elongation | AGATTGCGCGGATCGCCCGG |
| Human | TCEA2 | Transcription Elongation | GATGGTGACCAAGAAGAGCG |
| Human | TCEA2 | Transcription Elongation | CTGCTCCAGTCCACCCGAGT |
| Human | TCEA2 | Transcription Elongation | GCAGACGGACCATGACCACG |
| Human | TCEA2 | Transcription Elongation | ATCACTGGCCATCTCCTCTG |
| Human | TCEA2 | Transcription Elongation | AAAGAACTGCACCTACACAC |
| Human | TCEA2 | Transcription Elongation | TTTGTTGTCTGCAACGAGTG |
| Human | TCEA2 | Transcription Elongation | CTCGTTGCAGACAACAAAGG |
| Human | TCEA2 | Transcription Elongation | CAGACAACAAAGGTGGTCAT |
| Human | TCEA2 | Transcription Elongation | GCGGGTCTGCACCTGTGTGT |
| Human | FMR1 | Chromatin Readers | TTTCTACAAGGCATTTGTAA |
| Human | FMR1 | Chromatin Readers | TGCCTTGTAGAAAGCGCCAT |
| Human | FMR1 | Chromatin Readers | CTCCAATGGCGCTTTCTACA |

| | | | |
|---|---|---|---|
| Human | FMR1 | Chromatin Readers | TTAGCTAACCACCAACAGCA |
| Human | FMR1 | Chromatin Readers | AGCTAAAGTGAGGATGATAA |
| Human | FMR1 | Chromatin Readers | ATCGTAAGAGAAGATCTGAT |
| Human | FMR1 | Chromatin Readers | GGTCTAGCTATTGGTACTCA |
| Human | FMR1 | Chromatin Readers | GATCTGATGGGTCTAGCTAT |
| Human | FMR1 | Chromatin Readers | TATCGTAAGAGAAGATCTGA |
| Human | FMR1 | Chromatin Readers | CAGGAGATTGTGGACAAGTC |
| Human | FMR1 | Chromatin Readers | AACTTAGTAGGCAAAGTAAT |
| Human | FMR1 | Chromatin Readers | AAAGCTGATTCAGGAGATTG |
| Human | FMR1 | Chromatin Readers | CAAGTTCCAAGGAACTTAGT |
| Human | FMR1 | Chromatin Readers | ACTTTGCCTACTAAGTTCCT |
| Human | FMR1 | Chromatin Readers | GCAAATGTGTGCCAAAGAGG |
| Human | FMR1 | Chromatin Readers | ACGGCAAATGTGTGCCAAAG |
| Human | FMR1 | Chromatin Readers | TCTTTATTATAACCTACAGG |
| Human | FMR1 | Chromatin Readers | TATTATAACCTACAGGAGGT |
| Human | FMR1 | Chromatin Readers | CAGCATATGTGCTCGCTTTG |
| Human | FMR1 | Chromatin Readers | TGCTGATTGACATGCACTTT |
| Human | FMR1 | Chromatin Readers | ACCACCAAATCGTACAGATA |
| Human | FMR1 | Chromatin Readers | GTAATCTCTCCAAACGCAAC |
| Human | FMR1 | Chromatin Readers | GATGGTCAAGGAATGGGTCG |
| Human | FMR1 | Chromatin Readers | GCAGTCAACTCTGATCTGAA |
| Human | FMR1 | Chromatin Readers | TTACAGAATACCTCCAGTGA |
| Human | FMR1 | Chromatin Readers | CAGCCGACTACCTTCACTGG |
| Human | PPP2R1A | Control | GAAGCTGAACGTCCTCATTG |
| Human | PPP2R1A | Control | TCGTCTATGAGCACCGCGAT |
| Human | PPP2R1A | Control | GAAGCTGTCCACCATCGCCT |
| Human | PPP2R1A | Control | GCTGGGAACCTTCACTACCC |
| Human | PPP2R1A | Control | GGGAACCTTCACTACCCTGG |
| Human | PPP2R1A | Control | GGAACCTTCACTACCCTGGT |
| Human | PPP2R1A | Control | TAGATGGTATCTGTAAGGAA |
| Human | PPP2R1A | Control | ACCTTCACTACCCTGGTGGG |
| Human | PPP2R1A | Control | ACTGTGGCCAGCGACTCCAG |
| Human | PPP2R1A | Control | CACCACTGTCTCCTCCACTG |
| Human | PPP2R1A | Control | CTCGTGTGAGATGGCCCGTA |
| Human | PPP2R1A | Control | CAAGGCAGTGGAGTCCTTAC |
| Human | PPP2R1A | Control | GGAGACAGTGGTGCGGGACA |
| Human | PPP2R1A | Control | TGTGCCGCTAGTGAAGCGGC |
| Human | PPP2R1A | Control | ACTTTGTGCCGCTAGTGAAG |
| Human | PPP2R1A | Control | CTGGTTCACCTCCCGCACCT |
| Human | PPP2R1A | Control | ACCTCCCGCACCTCGGCCTG |
| Human | PPP2R1A | Control | CTAGTGAAGCGGCTGGCGGG |
| Human | PPP2R1A | Control | CTGCGGCCCGCCGCACCATG |

| | | | |
|---|---|---|---|
| Human | PPP2R1A | Control | TGTCATCTGAGCACAGGTTC |
| Human | PPP2R1A | Control | TGCGGCCCGCCGCACCATGG |
| Human | PPP2R1A | Control | CTGACGAGCAGGACTCGGTG |
| Human | PPP2R1A | Control | CACCGAGTCCTGCTCGTCAG |
| Human | PPP2R1A | Control | GGCCTCTGACGAGCAGGACT |
| Human | PPP2R1A | Control | GAGGCCAGGTTGGAGAACAT |
| Human | PPP2R1A | Control | AGGCCAGGTTGGAGAACATG |
| Human | PPP2R1A | Control | ACAGAGCTCCAGAAAGCAGT |
| Human | PPP2R1A | Control | CACAGAGCTCCAGAAAGCAG |
| Human | PPP2R1A | Control | ACTTGTCAGCCACCATGTAG |
| Human | PPP2R1A | Control | CTGGCGCGTCCGCTACATGG |
| Human | PPP2R1A | Control | GTCCTGGCGCGTCCGCTACA |
| Human | PPP2R1A | Control | GAACTCTTTGACCTTGTGGG |
| Human | PPP2R1A | Control | GAACCTGATGAAAGACTGTG |
| Human | PPP2R1A | Control | GATGAAAGACTGTGAGGCCG |
| Human | PPP2R1A | Control | AAGACTGTGAGGCCGAGGTG |
| Human | PPP2R1A | Control | GACACCAGCTCCTTGATGCA |
| Human | PPP2R1A | Control | CTTGACATGTTGGTTGGCAT |
| Human | PPP2R1A | Control | AGAGAGACCCATGATGACTG |
| Human | PPP2R1A | Control | GGACACCAGCTCCTTGATGC |
| Human | PPP2R1A | Control | CCAACATGTCAAGTCTGCCC |
| Human | PPP2R1A | Control | GCTGAAGGATGAGTGCCCTG |
| Human | PPP2R1A | Control | TAACCTGGACTGTGTGAACG |
| Human | PPP2R1A | Control | TCACCTCGTTCACACAGTCC |
| Human | PPP2R1A | Control | TGATGTTCAGCCGTACCTCA |
| Human | PPP2R1A | Control | AGGATGAGTGCCCTGAGGTA |
| Human | PPP2R1A | Control | TGGCTGAGGACGCCAAGTGG |
| Human | PPP2R1A | Control | AGCTGGCTGAGGACGCCAAG |
| Human | PPP2R1A | Control | GTCCTCAGCCAGCTCCACAA |
| Human | PPP2R1A | Control | AGGACGCCAAGTGGCGGGTG |
| Human | PPP2R1A | Control | TCATGTATATGCCATCCGCG |
| Human | PPP2R1A | Control | ATATACATGATCCACAAGCC |
| Human | PPP2R1A | Control | CACAAGCCAGGCCATGCACA |
| Human | PPP2R1A | Control | CAGCAACCTGAAGAAGCTAG |
| Human | PPP2R1A | Control | GAAACTTAACTCCTTGTGCA |
| Human | PPP2R1A | Control | CTGCATCAATGTGCTGTCTG |
| Human | PPP2R1A | Control | AGAGCGTAGTCATGCGGTGC |
| Human | PPP2R1A | Control | GGCCACATTGAAGCGGACAT |
| Human | PPP2R1A | Control | CCAGCCATGCGCAGAACCGT |
| Human | PPP2R1A | Control | TGCCAATGTCCGCTTCAATG |
| Human | PPP2R1A | Control | GAGACTTGGCCACATTGAAG |
| Human | PPP2R1A | Control | GCTACCCACGGTTCTGCGCA |

| | | | |
|---|---|---|---|
| Human | PPP2R1A | Control | GGACGTCAAATACTTTGCCC |
| Human | PPP2R1A | Control | CAGAACAGTCAGAGCCTCCT |
| Human | PPP2R1A | Control | GACGTCCACATCCTGGTCCT |
| Human | PPP2R1A | Control | CATCCTAGAGAAGCTGACCC |
| Human | PPP2R1A | Control | ACAGAACAGTCAGAGCCTCC |
| Human | PPP2R1A | Control | TCTCCCGACAGTCAGCTGAG |
| Human | PPP2R1A | Control | CGTGGGTAGCATGTGCTTGG |
| Human | PPP2R1A | Control | CACCAAGCACATGCTACCCA |
| Human | PPP2R1A | Control | AAGGTGCTGTTGTCCAGGAT |
| Human | PPP2R1A | Control | CCACTCCCAGCTGTCCAGCC |
| Human | HMGA1 | DNA and RNA Modifications | ACAAGGGTGCTGCCAAGACC |
| Human | HMGA1 | DNA and RNA Modifications | GGGCATCTCGCAGGAGTCCT |
| Human | HMGA1 | DNA and RNA Modifications | GGAAGAGGAGGGCATCTCGC |
| Human | JMJD6 | Histone Methylation | GAGACTGTCTTTGTACCAGG |
| Human | JMJD6 | Histone Methylation | ATCACAAACCACCTGTAAGG |
| Human | JMJD6 | Histone Methylation | TCTTTGTACCAGGAGGCTGG |
| Human | JMJD6 | Histone Methylation | TGAGTCGGAGTCTGACGAAC |
| Human | JMJD6 | Histone Methylation | GGAGTCTGACGAACTGGAGC |
| Human | JMJD6 | Histone Methylation | GTCTGACGAACTGGAGCTGG |
| Human | JMJD6 | Histone Methylation | GGAGGAGCTGGAAGAGTCGC |
| Human | JMJD6 | Histone Methylation | CGAACTGGAGCTGGAGGAGC |
| Human | JMJD6 | Histone Methylation | GAGCAAGAAGCGCATCCGCG |
| Human | JMJD6 | Histone Methylation | AGCGCATCCGCGAGGCCAAG |
| Human | JMJD6 | Histone Methylation | AGTTATCAAGGAAATGGTAT |
| Human | JMJD6 | Histone Methylation | GTTGGCAGTCCTCGCAGACT |
| Human | CHD1 | Chromatin Readers | GCAGATGGTGACCCAAATGC |
| Human | CHD1 | Chromatin Readers | CAGTATTTAATTAAATGGAA |
| Human | CHD1 | Chromatin Readers | ATTAAATACTGAATCTCTCC |
| Human | CHD1 | Chromatin Readers | AGCTGCACTCTGAGTATGGA |
| Human | CHD1 | Chromatin Readers | GATTATTACTGCAAATGGCA |
| Human | CHD1 | Chromatin Readers | TCCATACTCAGAGTGCAGCT |
| Human | CHD1 | Chromatin Readers | TGCCATTTGCAGTAATAATC |
| Human | CHD1 | Chromatin Readers | ATCCTGATTATTACTGCAAA |
| Human | CHD1 | Chromatin Readers | ACATGATAAGAACTCATGAA |
| Human | CHD1 | Chromatin Readers | ATGCCCAATTTAGACCTCCA |
| Human | CHD1 | Chromatin Readers | AAAGATAAGGCATTCCTTGG |
| Human | CHD1 | Chromatin Readers | TCTTTGAGGGAATTCTGTAG |
| Human | CHD1 | Chromatin Readers | CCTTTGCCAGGAAGTAAGAG |
| Human | CHD1 | Chromatin Readers | ATCCATCTAATCTTTGAAAG |
| Human | CHD1 | Chromatin Readers | ATTTATCGTCTAGTTACAAA |
| Human | CHD1 | Chromatin Readers | TGATCCATCTAATCTTTGAA |
| Human | CHD1 | Chromatin Readers | ATCGTTCTCACTCAGATCAT |

| | | | |
|---|---|---|---|
| Human | CHD1 | Chromatin Readers | TAGTGGAGAATCAAGCCAGT |
| Human | CHD1 | Chromatin Readers | ACCCAGAATCATCATCCGAC |
| Human | CHD1 | Chromatin Readers | CTGACTGCCTGATTCAGATC |
| Human | CHD1 | Chromatin Readers | GAAAGTGTTAGAAACAGTAG |
| Human | CHD1 | Chromatin Readers | TCTGACTCCGGATCTGAATC |
| Human | CHD1 | Chromatin Readers | ATCGGAATCTTCACTGCTAG |
| Human | CHD1 | Chromatin Readers | GATCATCTGTAAGTTCTTGC |
| Human | CHD1 | Chromatin Readers | CCAATTCTCCCAGTCGTCTA |
| Human | CHD1 | Chromatin Readers | CTGGGAGAATTGGTACATAA |
| Human | CHD1 | Chromatin Readers | CCTTAGACGACTGGGAGAAT |
| Human | CHD1 | Chromatin Readers | AAGGTATTAATTCTTCTTCA |
| Human | CHD1 | Chromatin Readers | AGGTATTAATTCTTCTTCAT |
| Human | CHD1 | Chromatin Readers | GATGATTCCAATTTGTTAAT |
| Human | CHD1 | Chromatin Readers | TTTGATGAGGTAGTCTGCAC |
| Human | CHD1 | Chromatin Readers | ATTTGATGAGGTAGTCTGCA |
| Human | CHD1 | Chromatin Readers | CTCTACTAAGTAATTTGATG |
| Human | CHD2 | Chromatin Readers | GAAGCTAATGGCGACCCTAG |
| Human | CHD2 | Chromatin Readers | TACAGTAGTAGATGCTCCAG |
| Human | CHD2 | Chromatin Readers | AAGAGGACGAAATCAAACAA |
| Human | CHD2 | Chromatin Readers | GTATATGCGATTGAAGCTAA |
| Human | CHD2 | Chromatin Readers | CCCAGCTACACTCTGAATAG |
| Human | CHD2 | Chromatin Readers | GCCGGCTTCCGACTATGAGC |
| Human | CHD2 | Chromatin Readers | TCCAGCTCATAGTCGGAAGC |
| Human | CHD2 | Chromatin Readers | TGATGAGCAGAAATACGATA |
| Human | CHD2 | Chromatin Readers | GCACAGTCTTATCTTTCAAG |
| Human | CHD2 | Chromatin Readers | GATGAGCAGAAATACGATAC |
| Human | CHD2 | Chromatin Readers | TTGAAAGATAAGACTGTGCT |
| Human | CHD2 | Chromatin Readers | GAACCATCCAGACGCTGGAA |
| Human | CHD2 | Chromatin Readers | TGATGGAACCATCCAGACGC |
| Human | CHD2 | Chromatin Readers | TGGACCACTTCAATGCAGAT |
| Human | CHD2 | Chromatin Readers | TTTACCGCTTAGTTACAAAG |
| Human | CHD2 | Chromatin Readers | TGAATCTTCTGAGAGTCAGT |
| Human | CHD2 | Chromatin Readers | GAATCTGAGAGCGAATCAGC |
| Human | CHD2 | Chromatin Readers | AGTCAGTCGGAAAGTGAGCA |
| Human | CHD2 | Chromatin Readers | TTCAGACTCAGGCAGTCAGT |
| Human | CHD2 | Chromatin Readers | GAAGCTTCGGGTTCAGACTC |
| Human | CHD2 | Chromatin Readers | TTAAGGAAGAGGCAAGTAGC |
| Human | CHD2 | Chromatin Readers | ATTAAGGAAGAGGCAAGTAG |
| Human | CHD2 | Chromatin Readers | GCAAGTAGCGGGTCTGAGAG |
| Human | CHD2 | Chromatin Readers | CAAGTAGCGGGTCTGAGAGT |
| Human | CHD2 | Chromatin Readers | TCATCTATATCCATTTCCTG |
| Human | CHD2 | Chromatin Readers | GCTAGAAGAGCGTCCTCACA |

| | | | |
|---|---|---|---|
| Human | CHD2 | Chromatin Readers | CTGGGATGAGATCATTCCAG |
| Human | CHD2 | Chromatin Readers | ATCTCATCCCAGTCCTTGTG |
| Human | CHD2 | Chromatin Readers | GATCTCTTCAAAGAACTGGA |
| Human | CXXC1 | Chromatin Readers | CTGCAATGAGTGGTTCCATG |
| Human | CXXC1 | Chromatin Readers | GGTGTGACAACTGCAATGAG |
| Human | CXXC1 | Chromatin Readers | CCATCCGGGAGTGGTACTGT |
| Human | CXXC1 | Chromatin Readers | CCGACAGTACCACTCCCGGA |
| Human | CXXC1 | Chromatin Readers | CATCCGGGAGTGGTACTGTC |
| Human | CXXC1 | Chromatin Readers | TGTCGGCGCACTGAGGACTG |
| Human | CXXC1 | Chromatin Readers | CCTCACACTCACCACACATG |
| Human | CXXC1 | Chromatin Readers | AAACGGTCAGCCCGCATGTG |
| Human | CXXC1 | Chromatin Readers | TGAGGCATGTCGGCGCACTG |
| Human | CXXC1 | Chromatin Readers | CTGCTTGGCACGTAGAATGA |
| Human | CXXC1 | Chromatin Readers | CATTTCCTGAAGGCGAGTGC |
| Human | CXXC1 | Chromatin Readers | ATCGGCGTTCCATTTCCTGA |
| Human | CXXC1 | Chromatin Readers | CCATTTCCTGAAGGCGAGTG |
| Human | CXXC1 | Chromatin Readers | ACGCCGATTCCATGAGCTTG |
| Human | CXXC1 | Chromatin Readers | AGTGGCGTCATCAACAGTCA |
| Human | CXXC1 | Chromatin Readers | TCTTCTGTGTCGCTCATCCA |
| Human | CXXC1 | Chromatin Readers | CTGCACAGAAGTCCTGATAC |
| Human | CXXC1 | Chromatin Readers | CGCTCATCCAGGGCAGGCCA |
| Human | CXXC1 | Chromatin Readers | TGTATCAGGACTTCTGTGCA |
| Human | CXXC1 | Chromatin Readers | GAAGTCTGAGAAGAAGAAGG |
| Human | CXXC1 | Chromatin Readers | GAAGAAGTCTGAGAAGAAGA |
| Human | CXXC1 | Chromatin Readers | AAGTGAAGCATGTGAAGCGT |
| Human | CXXC1 | Chromatin Readers | TCGGCAGAAGCAGAAGCACA |
| Human | PRMT3 | Histone Methylation | TCTATATGCTTAATACCACA |
| Human | PRMT3 | Histone Methylation | AGGGTCACGGTGAGAGAACG |
| Human | PRMT3 | Histone Methylation | TAGAGAACACGACCCTGTTG |
| Human | PRMT3 | Histone Methylation | ATTATTCAACGTGAGGGTCA |
| Human | PRMT3 | Histone Methylation | AGGTGAAGCCTTGAAAGGAA |
| Human | PRMT3 | Histone Methylation | AACCTGTTACAGAACAGGCA |
| Human | PRMT3 | Histone Methylation | TGAATAACCTGTTACAGAAC |
| Human | PRMT3 | Histone Methylation | TAACCTGTTACAGAACAGGC |
| Human | NCOA1 | Histone Acetylation | TGACACACTGGCATCAAGCA |
| Human | NCOA1 | Histone Acetylation | GCTGTCAATGTCACTAATGT |
| Human | NCOA1 | Histone Acetylation | CTACAGCATACTGCACGTGG |
| Human | NCOA1 | Histone Acetylation | TCTACAGCATACTGCACGTG |
| Human | NCOA1 | Histone Acetylation | CTACTTAGGTTACAATCAGG |
| Human | NCOA1 | Histone Acetylation | GTCTACAGCATACTGCACGT |
| Human | NCOA1 | Histone Acetylation | ATCATCGACAGGGAGCACAG |
| Human | NCOA1 | Histone Acetylation | TCATCGACAGGGAGCACAGT |

| | | | |
|---|---|---|---|
| Human | NCOA1 | Histone Acetylation | ATTGACAAACTTGTTCAGGG |
| Human | NCOA1 | Histone Acetylation | GAATTGACAAACTTGTTCAG |
| Human | NCOA1 | Histone Acetylation | AATCTATTTAGCCTGGAAGT |
| Human | NCOA1 | Histone Acetylation | AATTGACAAACTTGTTCAGG |
| Human | NCOA1 | Histone Acetylation | GTCAGCTGTAAACTGGCTCT |
| Human | NCOA1 | Histone Acetylation | TCTGACAACAGCTCTAGTGA |
| Human | NCOA1 | Histone Acetylation | AGCATGCCGTAACTGCTGTT |
| Human | NCOA1 | Histone Acetylation | ACCTCCTGAAGAGTTAGCAG |
| Human | NCOA1 | Histone Acetylation | AAAGATGTCCTGTCTTGCAC |
| Human | NCOA1 | Histone Acetylation | GGTTGTATATTTAATCTGCT |
| Human | NCOA1 | Histone Acetylation | AAATGTTGGCACAACGTCAG |
| Human | NCOA1 | Histone Acetylation | CAACTACAGCAGCGATTACA |
| Human | NCOA1 | Histone Acetylation | GCTGTATTAGCTGCCTCTGT |
| Human | JAK2 | Other Histone Modifications | CATGCCGCCACTGAGCAAAG |
| Human | JAK2 | Other Histone Modifications | TGTCTTACCTCTTTGCTCAG |
| Human | JAK2 | Other Histone Modifications | ATGAAGAGTACAACCTCAGT |
| Human | JAK2 | Other Histone Modifications | TGTACTTCGATGCAGTCCTA |
| Human | JAK2 | Other Histone Modifications | AAGAAAGCAGGTAATCAGAC |
| Human | JAK2 | Other Histone Modifications | ACATACATTCCCATGAATAA |
| Human | JAK2 | Other Histone Modifications | AGTTGCTAAACAGTTGGCAT |
| Human | JAK2 | Other Histone Modifications | ATACGCATAAATTCCGCTGG |
| Human | JAK2 | Other Histone Modifications | TACACATCTCAGATATGCAA |
| Human | JAK2 | Other Histone Modifications | TGATCATATAGATCTCATCT |
| Human | JAK2 | Other Histone Modifications | GAGTGTGCTACAGTGCTGGT |
| Human | JAK2 | Other Histone Modifications | ATCATACGCATAAATTCCGC |
| Human | JAK2 | Other Histone Modifications | CCTTACGATGACAGAAATGG |
| Human | JAK2 | Other Histone Modifications | CTTACGATGACAGAAATGGA |
| Human | JAK2 | Other Histone Modifications | ATTCTGATATATAGAAGAGG |
| Human | JAK2 | Other Histone Modifications | ACCATTCTGATATATAGAAG |
| Human | JAK2 | Other Histone Modifications | CCTCCATTTCTGTCATCGTA |
| Human | TCEA1 | Transcription Elongation | TGTGGACTGCAGTAATTCCA |
| Human | TCEA1 | Transcription Elongation | GTGCAGAAGAAGAACGCGGC |
| Human | TCEA1 | Transcription Elongation | TGCAGTAATTCCAGGGTCAT |
| Human | TCEA1 | Transcription Elongation | TCGCAAGCAGAGTACAGATG |
| Human | TCEA1 | Transcription Elongation | TTACTGCAGTCCACAAGAAT |
| Human | TCEA1 | Transcription Elongation | TTGCTGCAGCTCTTCGAACA |
| Human | TCEA1 | Transcription Elongation | ATTCTAGCAAATAAGTCAGG |
| Human | TCEA1 | Transcription Elongation | TGCTGCAGCTCTTCGAACAG |
| Human | TCEA1 | Transcription Elongation | TAGAATGACAGCAGAGGAAA |
| Human | TCEA1 | Transcription Elongation | CAGACAACAAATGTTGTCAT |
| Human | TCEA1 | Transcription Elongation | TCATTGGTTCATCAGCACTA |
| Human | TCEA1 | Transcription Elongation | TTTGTTGTCTGTAATGAATG |

| | | | |
|---|---|---|---|
| Human | TCEA1 | Transcription Elongation | GAAGAATTGCACTTACACAC |
| Human | TCEA1 | Transcription Elongation | CATTGGTTCATCAGCACTAC |
| Human | RAI1 | Chromatin Readers | TCGCAAACATGAGTGCAGCA |
| Human | RAI1 | Chromatin Readers | CAAACATGAGTGCAGCAAGG |
| Human | RAI1 | Chromatin Readers | TGCATGAGGCCTGTGCCGTG |
| Human | RAI1 | Chromatin Readers | CCAGGAGCACTGGGTGCATG |
| Human | RAI1 | Chromatin Readers | TTTGCGACCCTTGTCGGCTG |
| Human | RAI1 | Chromatin Readers | GCTGGCACACGGGTAGTGGT |
| Human | RAI1 | Chromatin Readers | GAATATGCAACCTGCATCGC |
| Human | RAI1 | Chromatin Readers | TCCTTTGTGGCAGCACCCAA |
| Human | RAI1 | Chromatin Readers | TTCCAGCTGCCAAGAAGCCG |
| Human | RAI1 | Chromatin Readers | GAAGTCGGCCTTCATGGCGC |
| Human | RAI1 | Chromatin Readers | CGTAGTGCTGATACTTGGCG |
| Human | RAI1 | Chromatin Readers | TGCAGGGAGTGAGTCCGAAA |
| Human | RAI1 | Chromatin Readers | GAGGAAAGTGGGTACCCTGG |
| Human | RAI1 | Chromatin Readers | TCGCCAAGTATCAGCACTAC |
| Human | RAI1 | Chromatin Readers | CTGTGCAACTCTTATAGGAG |
| Human | RAI1 | Chromatin Readers | GGACGAGGATGAGGAAGAGG |
| Human | RAI1 | Chromatin Readers | GGATGAGGAAGAGGAGGCAG |
| Human | PAF1 | Transcription Elongation | GGAAGCTCAGGAGGCACGGA |
| Human | PAF1 | Transcription Elongation | GAAGGAACTGGAAGCTCAGG |
| Human | PAF1 | Transcription Elongation | AACTGGAAGCTCAGGAGGCA |
| Human | PAF1 | Transcription Elongation | GGGCAGCGAGAGTGAACGGG |
| Human | PAF1 | Transcription Elongation | TGAACGGGAGGAAGGTGACA |
| Human | PAF1 | Transcription Elongation | GGCAGTGAAGATGAGCACTC |
| Human | PAF1 | Transcription Elongation | TGGTGAGGACGAGAGCAGCG |
| Human | PAF1 | Transcription Elongation | GACGAGGCCAGTGACAAGAG |
| Human | PAF1 | Transcription Elongation | TCTTCTCAATGGCTGTGATC |
| Human | TRIM37 | Other Histone Modifications | GATGGAAATGGAGTTGTGCG |
| Human | TRIM37 | Other Histone Modifications | GCACAACTCCATTTCCATCT |
| Human | TRIM37 | Other Histone Modifications | AAAGTTTACCCAGATGGAAA |
| Human | TRIM37 | Other Histone Modifications | GTTTCAGCTGTATTAGGCGC |
| Human | TRIM37 | Other Histone Modifications | ACAGCTGAAACAACACAGTT |
| Human | TRIM37 | Other Histone Modifications | CAACACAGTTTGGAGCAATG |
| Human | TRIM37 | Other Histone Modifications | TGTGTTGTTTCAGCTGTATT |
| Human | TRIM37 | Other Histone Modifications | GTTTGGAGCAATGAGGACAC |
| Human | TRIM37 | Other Histone Modifications | AGGAGGTGGAGCACCAGTTG |
| Human | TRIM37 | Other Histone Modifications | GCTTAGTTCAAGAAGTGGAA |
| Human | TRIM37 | Other Histone Modifications | TTTGGAATCCTTACTTCAGG |
| Human | TRIM37 | Other Histone Modifications | AATAAGCTTATAACACTGAT |
| Human | TRIM37 | Other Histone Modifications | AAGAGGTAGCCAAACTTCGT |
| Human | TRIM37 | Other Histone Modifications | CAATAGTAAGTCTCTCTTTA |

| | | | |
|---|---|---|---|
| Human | TRIM37 | Other Histone Modifications | CTTTAAGGTTGTTTATTTGT |
| Human | TAF5 | Transcription Elongation | ACGGCAGAGCAAACTCCGCG |
| Human | TAF5 | Transcription Elongation | GCACGGCCAGTAGAGTCTGT |
| Human | TAF5 | Transcription Elongation | CTCACGGCGCAGCGCCTCTT |
| Human | TAF5 | Transcription Elongation | CGCGGAGTTTGCTCTGCCGT |
| Human | TAF5 | Transcription Elongation | CTGCCGTAGGAACTGCAGCA |
| Human | TAF5 | Transcription Elongation | TTACCAGGGTCTCACTGCAG |
| Human | TAF5 | Transcription Elongation | CATCCACTGCAGTGAGACCC |
| Human | TAF5 | Transcription Elongation | TTCAACTGTCAGAGTGTGGT |
| Human | TAF5 | Transcription Elongation | CAGATTCAACTGTCAGAGTG |
| Human | TAF5 | Transcription Elongation | GATGATTCTAGTCTGATTGC |
| Human | TAF5 | Transcription Elongation | GAGCCAGCTTCAGTCCGGAT |
| Human | TAF5 | Transcription Elongation | GAAAGCAGATAGTTCCTATC |
| Human | TAF5 | Transcription Elongation | AACAGTTCCGTCCTCTGAAG |
| Human | TAF5 | Transcription Elongation | AGGACGGAACTGTTAGATTG |
| Human | TAF5 | Transcription Elongation | CTGCTTTCCTCTTCAGAGGA |
| Human | TAF5 | Transcription Elongation | TGACCGAGTAGCTCGGCTCT |
| Human | TAF5 | Transcription Elongation | AGAGCCGAGCTACTCGGTCA |
| Human | TAF5 | Transcription Elongation | TAGCCCAGAGCCGAGCTACT |
| Human | TAF5 | Transcription Elongation | CAACATAATTAGAATTTGGA |
| Human | TAF5 | Transcription Elongation | AGACAGAACTGTGCGGCTCT |
| Human | TAF5 | Transcription Elongation | ATTAGAATTTGGATGGAATC |
| Human | TAF5 | Transcription Elongation | CGGCTCTGGGACGTCCTGAA |
| Human | TAF5 | Transcription Elongation | GCTCTGCAGACAGAACTGTG |
| Human | TAF5 | Transcription Elongation | GTAGCCAGGAATCTCCCATT |
| Human | TAF5 | Transcription Elongation | TGTAGCCAGGAATCTCCCAT |
| Human | TAF5 | Transcription Elongation | CTGGCTACAGGAGCAACAGA |
| Human | TAF5 | Transcription Elongation | GGATATTGGACATGGTTTGA |
| Human | TAF5 | Transcription Elongation | CTTCTTTGGGATATTGGACA |
| Human | TAF5 | Transcription Elongation | CTGATACAGTCTGTTCACTT |
| Human | TAF5 | Transcription Elongation | GGATAATACAGTTCGATTAT |
| Human | TAF5 | Transcription Elongation | TCACTTAGGTTTAGTAGAGA |
| Human | TAF5 | Transcription Elongation | TGGATAATACAGTTCGATTA |
| Human | TAF5 | Transcription Elongation | AAGTACATGTGCACAAACAG |
| Human | TAF5 | Transcription Elongation | TCTGTTTGTGCACATGTACT |
| Human | FOXO4 | DNA and RNA Modifications | CCTGCACAGCAAGTTCATCA |
| Human | FOXO4 | DNA and RNA Modifications | CACTTGCCCAGATCTACGAG |
| Human | FOXO4 | DNA and RNA Modifications | AAGTGTCAGTCGCTTCTCCG |
| Human | FOXO4 | DNA and RNA Modifications | GTACCCTACTTCAAGGACAA |
| Human | PRDM16 | Histone Methylation | CGGATGGGAGCAAATACTGA |
| Human | PRDM16 | Histone Methylation | ACAAAGATCTCCGAAGACCT |
| Human | PRDM16 | Histone Methylation | AAAGGAGACAGACTTCGGAT |

| | | | |
|---|---|---|---|
| Human | PRDM16 | Histone Methylation | GGAGCAAATACTGACGGACG |
| Human | PRDM16 | Histone Methylation | CGAACTCTTCCAGTCCAAGC |
| Human | PRDM16 | Histone Methylation | TCCAGTCCAAGCTGGACCTG |
| Human | PRDM16 | Histone Methylation | ATGGCGCCGCAGGTCCAGCT |
| Human | PRDM16 | Histone Methylation | GCCGCAGGTCCAGCTTGGAC |
| Human | PRDM16 | Histone Methylation | GTACAGCCTGGAGCAGCACA |
| Human | PRDM16 | Histone Methylation | CTGCTCCAGGCTGTACTTGT |
| Human | PRDM16 | Histone Methylation | TGCTCCAGGCTGTACTTGTT |
| Human | PRDM16 | Histone Methylation | GCTCCAGGCTGTACTTGTTG |
| Human | PRDM16 | Histone Methylation | AGTTGAAGGCCTTGGGACAC |
| Human | PRDM16 | Histone Methylation | GAGGTTGGACTTCCAGTTGA |
| Human | PRDM16 | Histone Methylation | AGTGTCCCAAGGCCTTCAAC |
| Human | PRDM16 | Histone Methylation | GGACTTCCAGTTGAAGGCCT |
| Human | PRDM16 | Histone Methylation | GACTTCCAGTTGAAGGCCTT |
| Human | PRDM16 | Histone Methylation | AAACTGCGTGAAGGTGTTCA |
| Human | PRDM16 | Histone Methylation | TGTGCCGCTGAAGGTTGCTG |
| Human | PRDM16 | Histone Methylation | GATGTGCCGCTGAAGGTTGC |
| Human | PRDM16 | Histone Methylation | CGAAGGTCTTCCCGCAGTCG |
| Human | PRDM16 | Histone Methylation | AAGACCTTCGCCACGTCCTC |
| Human | PRDM16 | Histone Methylation | GAGGCCGGAGGACGTGGCGA |
| Human | PRDM16 | Histone Methylation | CTTGTGCTGCTTGAGGCCGG |
| Human | PRDM16 | Histone Methylation | CTGCTTGAGGCCGGAGGACG |
| Human | PRDM16 | Histone Methylation | AGAACTGCGTGTAGGACTTG |
| Human | PRDM16 | Histone Methylation | CGCAGTTCTCCAACCTGTGC |
| Human | PRDM16 | Histone Methylation | CAGGTTGGAGAACTGCGTGT |
| Human | PRDM16 | Histone Methylation | CCAACCTGTGCCGGCACAAG |
| Human | PRDM16 | Histone Methylation | CCTCCTCCCTCAACAAGCAC |
| Human | PRDM16 | Histone Methylation | AGGAGGTAGTGCTGAACATC |
| Human | PRDM16 | Histone Methylation | CCGGTGCTTGTTGAGGGAGG |
| Human | PRDM16 | Histone Methylation | CCAATCTCACCAGACACCTG |
| Human | PRDM16 | Histone Methylation | TCTGGTGAGATTGGCTGATC |
| Human | PRDM16 | Histone Methylation | CTGGTGAGATTGGCTGATCT |
| Human | PRDM16 | Histone Methylation | TGGTGAGATTGGCTGATCTG |
| Human | PRDM16 | Histone Methylation | GCATCTCTTCGAACCTCCAG |
| Human | PRDM16 | Histone Methylation | CGAACCTCCAGCGGCACGTC |
| Human | PRDM16 | Histone Methylation | GAGGTTCGAAGAGATGCTGA |
| Human | PRDM16 | Histone Methylation | TCGAAGAGATGCTGAAGGAG |
| Human | PRDM16 | Histone Methylation | TGGTCTGCTGCCCGAAGCAG |
| Human | PRDM16 | Histone Methylation | CTTCGGGCAGCAGACCAACC |
| Human | PRDM16 | Histone Methylation | GGCAGCAGACCAACCTGGAC |
| Human | PRDM16 | Histone Methylation | CGAAGCAGCGGTTGCACAGG |
| Human | PRDM16 | Histone Methylation | CTTGAGGTGCCGGTCCAGGT |

| | | | |
|---|---|---|---|
| Human | PRDM16 | Histone Methylation | GATACACGTGCAGGTACTGT |
| Human | PRDM16 | Histone Methylation | TCGCAGTACTTACACCTGTA |
| Human | PRDM16 | Histone Methylation | AACGCGAACAGAGAAACGGG |
| Human | PRDM16 | Histone Methylation | CCACACCCGAAGGTGTGCTG |
| Human | PRDM16 | Histone Methylation | GCGGACATGCAGATCGTGGA |
| Human | PRDM16 | Histone Methylation | GCACCAGTGAGCCAGCACCC |
| Human | PRDM16 | Histone Methylation | CTCTTGAGCAGCGATGTGGG |
| Human | PRDM16 | Histone Methylation | GGGCTCTTGAGCAGCGATGT |
| Human | PRDM16 | Histone Methylation | TTTCCCTCCAGGCCGCACCC |
| Human | SMYD2 | Histone Methylation | TGGAGTGTCAGAAAGAAGAT |
| Human | SMYD2 | Histone Methylation | GTGCATATCTGACCATGTCT |
| Human | SMYD2 | Histone Methylation | ATATCTGACCATGTCTCGGA |
| Human | SMYD2 | Histone Methylation | CACATGATGTACCAGGCCAT |
| Human | SMYD2 | Histone Methylation | CATGTTGCACATGATGTACC |
| Human | SMYD2 | Histone Methylation | GCACATGATGTACCAGGCCA |
| Human | SMYD2 | Histone Methylation | CCCTAGCTTCAACCACATGG |
| Human | SMYD2 | Histone Methylation | TCTCCCTAGCTTCAACCACA |
| Human | SMYD2 | Histone Methylation | GCCTCCATGTGGTTGAAGCT |
| Human | SMYD2 | Histone Methylation | CCTCCATGTGGTTGAAGCTA |
| Human | SMYD2 | Histone Methylation | GAAGAAGGCCATTGCAATCA |
| Human | SMYD2 | Histone Methylation | GATTGCAATGGCCTTCTTCA |
| Human | SMYD2 | Histone Methylation | TGATTGCAATGGCCTTCTTC |
| Human | KDM5D | Histone Methylation | GCTGAATTCCAAGACCCGCT |
| Human | KDM5D | Histone Methylation | AAATCCGCCCACCCGCGGAT |
| Human | KDM5D | Histone Methylation | TTCGTAATGTGATCGTAGCA |
| Human | KDM5D | Histone Methylation | AAGCCATCTGCAAGGATCGT |
| Human | KDM5D | Histone Methylation | CCACCGACGATCCTTGCAGA |
| Human | KDM5D | Histone Methylation | CCTTAGTAAGATTGTGATTG |
| Human | KDM5D | Histone Methylation | TCTATTAACTATCTGCATTG |
| Human | KDM5D | Histone Methylation | ACCATGCCCACGTACAGCCA |
| Human | KDM5D | Histone Methylation | CCCAACACTTTGATGTCCCA |
| Human | KDM5D | Histone Methylation | AGAACAGACTGATCTAGCAC |
| Human | KDM5D | Histone Methylation | AAGGTGCCCTGGCTGTACGT |
| Human | KDM5D | Histone Methylation | GTGGTAACAAGCAGAAGATG |
| Human | KDM5D | Histone Methylation | CCAAGGTGTACCGATACCGG |
| Human | KDM5D | Histone Methylation | CCTCCGGTATCGGTACACCT |
| Human | KDM5D | Histone Methylation | GCCGACAGTACCTCCGGTAT |
| Human | KDM5D | Histone Methylation | CAGGGCTGACAAGAAGCACG |
| Human | KDM5D | Histone Methylation | ACCGATACCGGAGGTACTGT |
| Human | KDM5D | Histone Methylation | CATTCCCACCAGGCTAGCAG |
| Human | KDM5D | Histone Methylation | GCACTGACACACACTGCCCA |
| Human | KDM5D | Histone Methylation | CATCTCCACTGCTAGCCTGG |

| | | | |
|---|---|---|---|
| Human | KDM5D | Histone Methylation | AGTGGAGATGAAGTGAGACT |
| Human | KDM5D | Histone Methylation | GACTGGGCTTTGGAGAGGTG |
| Human | RIOX1 | Histone Methylation | ACGTACCAGCGCAATACCTG |
| Human | RIOX1 | Histone Methylation | AACTTCAGTCAGGACGACCT |
| Human | RIOX1 | Histone Methylation | GAGCGGCCCGCCTCTTTGAG |
| Human | RIOX1 | Histone Methylation | ATTGCAGCCTGCACTGCCAG |
| Human | RIOX1 | Histone Methylation | CTTAGAGGCCATACTGCCTC |
| Human | RIOX1 | Histone Methylation | CTGCACTGCCAGAGGCAGTA |
| Human | RIOX1 | Histone Methylation | CATACTGCCTCTGGCAGTGC |
| Human | RIOX1 | Histone Methylation | CCGCACCTTCTCCATGAAAG |
| Human | RIOX1 | Histone Methylation | AAGAACCGCTTTCATGGAGA |
| Human | RIOX1 | Histone Methylation | CATGGAGAAGGTGCGGGTCT |
| Human | RIOX1 | Histone Methylation | CGCTTTCATGGAGAAGGTGC |
| Human | RIOX1 | Histone Methylation | CCGCTTTCATGGAGAAGGTG |
| Human | RIOX1 | Histone Methylation | GAAGTCTTTGGCTCGCTGGT |
| Human | RIOX1 | Histone Methylation | TCAGGATGGGATAGCTCGGC |
| Human | RIOX1 | Histone Methylation | TGCTTCAGGATGGGATAGCT |
| Human | RIOX1 | Histone Methylation | CTATCCCATCCTGAAGCATA |
| Human | RIOX1 | Histone Methylation | ACGTGGTTGCCAAGGACAGC |
| Human | RPS6KA4 | Other Histone Modifications | CCTGGACTATGTGAGCGGCG |
| Human | RPS6KA4 | Other Histone Modifications | CTCATCCTGGACTATGTGAG |
| Human | RPS6KA4 | Other Histone Modifications | CTGGAACACCTGCACAAGCT |
| Human | RPS6KA4 | Other Histone Modifications | CGCAGGCTGAGGTGTCTCGA |
| Human | RPS6KA4 | Other Histone Modifications | GGGCACGGGAGCCTACGGCA |
| Human | RPS6KA4 | Other Histone Modifications | AGTATCCCTGAAAGATTCGG |
| Human | RPS6KA4 | Other Histone Modifications | GAGTATCCCTGAAAGATTCG |
| Human | RPS6KA4 | Other Histone Modifications | ACTTTGCGGAGGAATTCACT |
| Human | RPS6KA4 | Other Histone Modifications | CAGCCAGAGCCACCCAATCG |
| Human | RPS6KA4 | Other Histone Modifications | TCCAGCTCTGAGCGGATTTG |
| Human | RPS6KA4 | Other Histone Modifications | AAGCCAAGGAGCTGGTCCGA |
| Human | RPS6KA4 | Other Histone Modifications | GGCCCGGGTTGGCTGGTGCA |
| Human | MAPK9 | Other Histone Modifications | GCACTGACCATATTGATCAG |
| Human | MAPK9 | Other Histone Modifications | GAACGTTGATATCTGGTCAG |
| Human | MAPK9 | Other Histone Modifications | ATGCAGCACAAACAATCCCT |
| Human | POLR2F | Transcription Elongation | GCGCTCGGGTGCCCAGCACG |
| Human | POLR2F | Transcription Elongation | CGCTCGGGTGCCCAGCACGC |
| Human | MAPK8 | Other Histone Modifications | GAAGGGACTATATTGATCAG |
| Human | MAPK8 | Other Histone Modifications | ACTTAAAGCCAGTCAGGCAA |
| Human | MAPK8 | Other Histone Modifications | ACGTGGATTTATGGTCTGTG |
| Human | MAPK8 | Other Histone Modifications | GTGGAAAGAATTGATATATA |
| Human | PRDM8 | Histone Methylation | CCTCGTCTTTGGCAATCCTG |
| Human | PRDM8 | Histone Methylation | GTTCCTCAGAAGGTCTCATG |

| | | | |
|---|---|---|---|
| Human | PRDM8 | Histone Methylation | GGTCCAATCGGCCAGAGATA |
| Human | PRDM8 | Histone Methylation | GTCGCAGGTGGTGTAAACGC |
| Human | PRDM8 | Histone Methylation | CATATAGGGAAGTATGGCTC |
| Human | PRDM8 | Histone Methylation | GAAACGCAGATGCGCCACAT |
| Human | PRDM8 | Histone Methylation | AACGCAGATGCGCCACATAG |
| Human | PRDM8 | Histone Methylation | AAACGCAGATGCGCCACATA |
| Human | PRDM8 | Histone Methylation | CCGACCTGGTGTACCATATG |
| Human | PRDM8 | Histone Methylation | CCTCATATGGTACACCAGGT |
| Human | PRDM8 | Histone Methylation | ACACCAGGTCGGAGGTCATG |
| Human | PRDM8 | Histone Methylation | CAGGTCGGAGGTCATGCGGA |
| Human | PRDM8 | Histone Methylation | CATCTGCAATGAGTCCTTCA |
| Human | PRDM8 | Histone Methylation | TGAAGGACTCATTGCAGATG |
| Human | PRDM8 | Histone Methylation | CCTGAAGGACTCATTGCAGA |
| Human | PRDM8 | Histone Methylation | TCATGTGCCTGGAGAGGTGG |
| Human | PRDM8 | Histone Methylation | GAGGTGGTGGCGCTCCCTGA |
| Human | PRDM8 | Histone Methylation | AGGAAAGAGGAAATTCCCGG |
| Human | PRDM8 | Histone Methylation | CCCGACGGCATCGCCACGGG |
| Human | PRDM8 | Histone Methylation | CGGGACTTGGCGGCGCGAAG |
| Human | PRDM8 | Histone Methylation | GGAGCTCTTGGGCCAGAAGG |
| Human | PRDM8 | Histone Methylation | CTCCGCTGCCGCTGCAGCCG |
| Human | PRDM8 | Histone Methylation | CGCTGCAGCCGCGGCTGCGG |
| Human | PRDM8 | Histone Methylation | TGCCGCTGCAGCCGCGGCTG |
| Human | PRDM8 | Histone Methylation | GCAGCGGCAGCGGAGCTCTT |
| Human | SMARCD1 | Chromatin Remodeling | GAGACTCAAAGATCTGCTGC |
| Human | SMARCD1 | Chromatin Remodeling | TTAATGATGATAGGTTCTGG |
| Human | SMARCD1 | Chromatin Remodeling | CAACATACCTGGTCTCGGAT |
| Human | SMARCD1 | Chromatin Remodeling | GAGCGGTACAGCCCTTGACC |
| Human | SMARCD1 | Chromatin Remodeling | GGAGCGGCCTATCCGAGACC |
| Human | SMARCD1 | Chromatin Remodeling | CCTGGCAACATACCTGGTCT |
| Human | SMARCD1 | Chromatin Remodeling | GGCAGCCGAATGACACCTCA |
| Human | SMARCD1 | Chromatin Remodeling | CCTCAAGACAATGACTGATG |
| Human | SMARCD1 | Chromatin Remodeling | ACCATCTGGTAGAATGGCAC |
| Human | SMARCD1 | Chromatin Remodeling | CCACATCAGTCATTGTCTTG |
| Human | SMARCD1 | Chromatin Remodeling | GCTGGTAATCCAGCATCAGT |
| Human | SMARCD1 | Chromatin Remodeling | GCACCTTGGAGTAGAAGTAT |
| Human | SMARCD1 | Chromatin Remodeling | CAAGAATTAGAGCAAGCCCT |
| Human | SMARCD1 | Chromatin Remodeling | TGATGTGGTGGGTAACCCAG |
| Human | SMARCD1 | Chromatin Remodeling | TTGTCGTCTCTGCTGCACCT |
| Human | SMARCD1 | Chromatin Remodeling | GTGCCGATACTTCTACTCCA |
| Human | TAF13 | Transcription Elongation | TATCACTGAAATGACTCACA |
| Human | TAF13 | Transcription Elongation | GAATTGCGATGTATGATGTA |
| Human | TAF13 | Transcription Elongation | ATATCCACTGACTCAGTATA |

| | | | |
|---|---|---|---|
| Human | ICE1 | Transcription Elongation | ATTCAAATTCTGCTGCAGAG |
| Human | ICE1 | Transcription Elongation | TAAGGTGAAGAAGCTGCAAG |
| Human | ICE1 | Transcription Elongation | AATTTAAGAAGACACAGGAA |
| Human | ICE1 | Transcription Elongation | AAAGGAATTTAAGAAGACAC |
| Human | ICE1 | Transcription Elongation | GGCTGCTGTCAAGCAAACTC |
| Human | ICE1 | Transcription Elongation | ATGAAGACGGTAAGGAACCC |
| Human | ICE1 | Transcription Elongation | CCCGGGTGAGGTGATGACAT |
| Human | ICE1 | Transcription Elongation | CCGATGTCATCACCTCACCC |
| Human | ICE1 | Transcription Elongation | GTGAGGTGATGACATCGGAG |
| Human | ICE1 | Transcription Elongation | TGAGGTGATGACATCGGAGG |
| Human | ICE1 | Transcription Elongation | GGACTAGAACTCGATATCAG |
| Human | ICE1 | Transcription Elongation | AGGAAGTGCGTGCTTCGGCG |
| Human | ICE1 | Transcription Elongation | GTGGCCGCACAGAACTGCAG |
| Human | ICE1 | Transcription Elongation | CTTGGAGGTGTAGCTATCAG |
| Human | ICE1 | Transcription Elongation | GCTATCAGAGGAGCAAGCAG |
| Human | PRDM12 | Histone Methylation | CATGTGGGAGGTGTTCAATG |
| Human | PRDM12 | Histone Methylation | CATTGAACACCTCCCACATG |
| Human | PRDM12 | Histone Methylation | TGGGAGGTGTTCAATGAGGA |
| Human | PRDM12 | Histone Methylation | GTCAGGTGGGATCATCTCAA |
| Human | PRDM12 | Histone Methylation | TGCTGCGCGAGTTGAAGCCG |
| Human | PRDM12 | Histone Methylation | TGCGCGAGTTGAAGCCGCGG |
| Human | PRDM12 | Histone Methylation | TGGACGACTGGCTGAAGCGG |
| Human | PRDM12 | Histone Methylation | GGTTGCGCAGCGTGGACGAC |
| Human | PRDM12 | Histone Methylation | GCGTGGACGACTGGCTGAAG |
| Human | PRDM12 | Histone Methylation | TGAAGCGGCGGTTGCAGAAG |
| Human | PRDM12 | Histone Methylation | CAGGCCGGCCAGCTGCGAGT |
| Human | PRDM12 | Histone Methylation | CCAGAGCGCCTACTCGCAGC |
| Human | PRDM12 | Histone Methylation | CCAGCTGCGAGTAGGCGCTC |
| Human | PRDM12 | Histone Methylation | AGCGCCTACTCGCAGCTGGC |
| Human | PRDM12 | Histone Methylation | AGTAGGCGCTCTGGCACACC |
| Human | PRDM12 | Histone Methylation | GGCGGCGGCGGCGGCGAGCG |
| Human | PRDM12 | Histone Methylation | GCGGCGGCGGCGGCGAGCGC |
| Human | PRDM13 | Histone Methylation | GCTATTGCAGACTTACCCGG |
| Human | PRDM13 | Histone Methylation | GGAGTGGATAGGGTTAATCC |
| Human | PRDM13 | Histone Methylation | CGGTAGAAGATCTGTCCTCC |
| Human | PRDM13 | Histone Methylation | TAACCCTATCCACTCCAGAG |
| Human | PRDM13 | Histone Methylation | CAGGTGTGCCTTAAGGCTGT |
| Human | PRDM13 | Histone Methylation | AGGTGTGCCTTAAGGCTGTT |
| Human | PRDM13 | Histone Methylation | AGCTGTACTCGCGCAAGTAT |
| Human | PRDM13 | Histone Methylation | ATGGGCTCAAGATCCACATG |
| Human | PRDM13 | Histone Methylation | AGTACAGCTTGCCACAGTAG |
| Human | PRDM13 | Histone Methylation | AAGCTGTACTCGCGCAAGTA |

| | | | |
|---|---|---|---|
| Human | PRDM13 | Histone Methylation | GCAATCTCAACAAGCACATC |
| Human | PRDM13 | Histone Methylation | TGTGCTTGTTGAGATTGCTG |
| Human | PRDM13 | Histone Methylation | ATGTGCTTGTTGAGATTGCT |
| Human | PRDM13 | Histone Methylation | CAAGGTACTTGTGCGCCGCC |
| Human | PRDM13 | Histone Methylation | GCAAGGTACTTGTGCGCCGC |
| Human | PRDM13 | Histone Methylation | TGACATGTCGCTCCAGGTCC |
| Human | PRDM13 | Histone Methylation | CATGTCGCTCCAGGTCCCGG |
| Human | PRDM13 | Histone Methylation | ACTTGTGCGCCGCCGGGACC |
| Human | PRDM13 | Histone Methylation | GGTGGTGCGCGTGGTGGTGA |
| Human | PRDM9 | Histone Methylation | TATAATGCACTGATTACTAT |
| Human | PRDM9 | Histone Methylation | ATGGGCAGAGATGGGAGACT |
| Human | PRDM9 | Histone Methylation | AAGGAAGAATGGGCAGAGAT |
| Human | PRDM9 | Histone Methylation | CTCGCTATAGGAATGTGAAA |
| Human | PRDM9 | Histone Methylation | CAAGGAAGAATGGGCAGAGA |
| Human | PRDM9 | Histone Methylation | ACTCCTGGCTGATCACCAAG |
| Human | PRDM9 | Histone Methylation | ATACTCCTGGCTGATCACCA |
| Human | PRDM9 | Histone Methylation | TACTCCTGGCTGATCACCAA |
| Human | PRDM9 | Histone Methylation | TGGGCTTGGAGTATGGAATG |
| Human | PRDM9 | Histone Methylation | GAGGCATCTGATCTGCCGCT |
| Human | PRDM9 | Histone Methylation | GAGAAATTTCTGACTTGAAA |
| Human | PRDM9 | Histone Methylation | GCCAGACAGCATGAGGGACA |
| Human | PRDM9 | Histone Methylation | CAGATGTTATTACACACCAA |
| Human | PRDM9 | Histone Methylation | CACACCTCCTCATTCACCAG |
| Human | PRDM9 | Histone Methylation | CAGTCCTCCTCACTCACCAG |
| Human | PRDM9 | Histone Methylation | CACACCTCCTCAGACACCAG |
| Human | PRDM9 | Histone Methylation | CAAACCTCCTCAGTCACCAG |
| Human | ZWILCH | Chromatin Remodeling | TACAGGAATTGTCTTATATG |
| Human | ZWILCH | Chromatin Remodeling | ACTGAAATTTCTTCTTGTTT |
| Human | ZWILCH | Chromatin Remodeling | CTAATATTTCTAGAATATGG |
| Human | NUP62 | Control | AACAGCGACTCTTGCTTCGG |
| Human | NUP62 | Control | GGCAAGTGAGAACAGGCCGG |
| Human | NUP62 | Control | ACAGCGACTCTTGCTTCGGG |
| Human | NUP62 | Control | CCAAACGTGAACCCGCCTGT |
| Human | NUP62 | Control | GAACAGCGACTCTTGCTTCG |
| Human | NUP62 | Control | CTCCATTCTCGATCAGCGTG |
| Human | NUP62 | Control | CGATGCACAGCTCAAGCGCA |
| Human | NUP62 | Control | CACCAGCCTGCACCGCGAGG |
| Human | NUP62 | Control | CCTTGACCAACTCCTCCAGT |
| Human | NUP62 | Control | TGCCACGTTGCCCTTCACTC |
| Human | NUP62 | Control | TAAATTCAAGGCAAAGCCGG |
| Human | NUP62 | Control | CGGCAGCTGCTGTATTGCTG |
| Human | YTHDF1 | DNA and RNA Modifications | CTTGTCCTGAGACCAGACCC |

| | | | |
|---|---|---|---|
| Human | YTHDF1 | DNA and RNA Modifications | ATGTACCCAATAACCAGCTC |
| Human | YTHDF1 | DNA and RNA Modifications | TGGAGTACTTAATGGAGCGG |
| Human | YTHDF1 | DNA and RNA Modifications | CAAACCGGTCACAAACTCCC |
| Human | YTHDF1 | DNA and RNA Modifications | TCCATGTTATGCTTTATGGG |
| Human | YTHDF1 | DNA and RNA Modifications | GTTATGCTTTATGGGTGGAG |
| Human | YTHDF1 | DNA and RNA Modifications | TCACTACGAGAAGCGCCAGG |
| Human | YTHDF1 | DNA and RNA Modifications | CTACGAGAAGCGCCAGGAGG |
| Human | YTHDF1 | DNA and RNA Modifications | CGAGAAGCGCCAGGAGGAGG |
| Human | YTHDF1 | DNA and RNA Modifications | GAAGCGCCAGGAGGAGGAGG |
| Human | BRPF3 | Chromatin Readers | GGAACCTCACTCAAGTTGAC |
| Human | BRPF3 | Chromatin Readers | GGAATTCATATCCAAGCCAA |
| Human | BRPF3 | Chromatin Readers | CTCCAAGGTGCGGTACAGGT |
| Human | BRPF3 | Chromatin Readers | TGGGATCGATGATCAAGGCA |
| Human | BRPF3 | Chromatin Readers | CACCTGGCAGTGGCTTCCAA |
| Human | BRPF3 | Chromatin Readers | CATCTTGGGATCGATGATCA |
| Human | BRPF3 | Chromatin Readers | ATCGATGATCAAGGCAGGGT |
| Human | BRPF3 | Chromatin Readers | TGATCAAGGCAGGGTAGGAG |
| Human | BRPF3 | Chromatin Readers | GGCGTCCCATACATCCCTGA |
| Human | BRPF3 | Chromatin Readers | CTGCAACCTGGCTGTACACC |
| Human | BRPF3 | Chromatin Readers | AGCACTCCTGGTGTACAGCC |
| Human | BRPF3 | Chromatin Readers | TATTGTGACATTCATCATCC |
| Human | BRPF3 | Chromatin Readers | TGAAGGCGCCACCCTTATTG |
| Human | BRPF3 | Chromatin Readers | CCAGATGGCACACACCACAT |
| Human | BRPF3 | Chromatin Readers | GCCTTCAAACAGACCAGTGA |
| Human | BRPF3 | Chromatin Readers | AGTCTTGCGCACTGTAAAGA |
| Human | BRPF3 | Chromatin Readers | ATGCGCGAAACCAGCCTCAA |
| Human | BRPF3 | Chromatin Readers | GTTCACCTTATGGCACTGGA |
| Human | BRPF3 | Chromatin Readers | GCGCAAGACTGCCTACTGTG |
| Human | BRPF3 | Chromatin Readers | TCCATGTGACATGTGCACAG |
| Human | BRPF3 | Chromatin Readers | TGGCGATGAGGAAGGGCTGA |
| Human | BRPF3 | Chromatin Readers | GAAGGAGGGTGATGGAGAGG |
| Human | BRPF3 | Chromatin Readers | CGATGAGGAAGGGCTGAAGG |
| Human | BRPF3 | Chromatin Readers | GGAGGAAGAAGAGGAAGAGG |
| Human | BRPF3 | Chromatin Readers | GAAGGGCTGAAGGAGGGTGA |
| Human | BRPF3 | Chromatin Readers | GTGGATCATCGCACGGTCAT |
| Human | ACTL6B | Chromatin Remodeling | GCAGGGCATTGGTGTCGATG |
| Human | ACTL6B | Chromatin Remodeling | TCCATCCCGAGGCACGTGCA |
| Human | ACTL6B | Chromatin Remodeling | GCCCTGCACGTGCCTCGGGA |
| Human | ACTL6B | Chromatin Remodeling | CCAATGCCCTGCACGTGCCT |
| Human | ACTL6B | Chromatin Remodeling | GCACGTGCCTCGGGATGGAG |
| Human | PCID2 | Transcription Elongation | GTAACCAGAGCTGTGAGCGA |
| Human | PCID2 | Transcription Elongation | ATATACATGGGACACGTCAA |

| | | | |
|---|---|---|---|
| Human | PCID2 | Transcription Elongation | AGTAACCAGAGCTGTGAGCG |
| Human | PCID2 | Transcription Elongation | AGAACGGTGACAATGCTCAA |
| Human | SMYD5 | Histone Methylation | CAGTACATCACTTGGCAATG |
| Human | SMYD5 | Histone Methylation | CGTTTCGTGAGCAGCGCCAA |
| Human | SMYD5 | Histone Methylation | GTCCTTCGCCTGCTTCACTG |
| Human | SMYD5 | Histone Methylation | ATGAGGAAGCAGTCAGCCAG |
| Human | SMYD5 | Histone Methylation | AGTGAAGCAGGCGAAGGACA |
| Human | SMYD5 | Histone Methylation | ACCCAATGTGACCTCAGAAG |
| Human | SMYD5 | Histone Methylation | AGCCAGAAGATGCAGAGCTG |
| Human | SMYD5 | Histone Methylation | GAGCCAGAAGATGCAGAGCT |
| Human | SMYD5 | Histone Methylation | TGTGACCTCAGAAGAGGAAG |
| Human | SMYD5 | Histone Methylation | CTCTTCCTCTTCCTCTTCTG |
| Human | CHD7 | Chromatin Readers | AAACGATTTAAGGCAAAGCA |
| Human | CHD7 | Chromatin Readers | AAAGCAGAAGGAATCTGGAG |
| Human | CHD7 | Chromatin Readers | CTCTTATCTTCATTGTCAGT |
| Human | CHD7 | Chromatin Readers | TCTCTTATCTTCATTGTCAG |
| Human | CHD7 | Chromatin Readers | CATAGATCAAGCAAAGATCG |
| Human | CHD7 | Chromatin Readers | GCACGTAGCACAGATGACCG |
| Human | CHD7 | Chromatin Readers | AGTGAGTCACAGGCTCTCCC |
| Human | CHD7 | Chromatin Readers | GTTTGAGAAACTAATGTCCA |
| Human | CHD7 | Chromatin Readers | TTCCTTATGAAGACAGCACG |
| Human | CHD7 | Chromatin Readers | CCCTTTATCACTCGACCCTG |
| Human | CHD7 | Chromatin Readers | ATGGGAGTCAAGCTAGTCGT |
| Human | CHD7 | Chromatin Readers | TCAATGACTACACAGCGCCA |
| Human | CHD7 | Chromatin Readers | GACATCTAGCCTGAGCCTGG |
| Human | CHD7 | Chromatin Readers | ATGGGTACCGTCTTTGAATG |
| Human | CHD7 | Chromatin Readers | AAAGACGGTACCCATATGAA |
| Human | CHD7 | Chromatin Readers | TACCCATATGAAAGGATCGA |
| Human | CHD7 | Chromatin Readers | TGCCAAGAGGCGAAATCTCA |
| Human | CHD7 | Chromatin Readers | CTCCATGAGATTTCGCCTCT |
| Human | CHD7 | Chromatin Readers | GAGGCGAAATCTCATGGAGA |
| Human | CHD7 | Chromatin Readers | AGGCCGAAAGAGCTGCCAAG |
| Human | CHD7 | Chromatin Readers | GGGCAGCTATATGGCACGTG |
| Human | CHD7 | Chromatin Readers | ATGGGCAGCTATATGGCACG |
| Human | CHD7 | Chromatin Readers | GTGCCATGGGAATCGGACAG |
| Human | CHD7 | Chromatin Readers | GGAGAGGCATAAGTTCCCTG |
| Human | CHD7 | Chromatin Readers | ATTGCTTTCATGGGTGACAT |
| Human | CHD7 | Chromatin Readers | AAAGCAATGAGTAATCCAGC |
| Human | CHD7 | Chromatin Readers | GAAGGAACCGAAAGAACCCA |
| Human | CHD7 | Chromatin Readers | TGGGATCCTCTGCTACAATG |
| Human | CHD7 | Chromatin Readers | AAACAACCACATTGTAGCAG |
| Human | CHD7 | Chromatin Readers | TTCAGAGAAAGGTTCCGAAG |

| | | | |
|---|---|---|---|
| Human | CHD7 | Chromatin Readers | GAACTGAAAGGTGTTGAGGT |
| Human | CHD7 | Chromatin Readers | GCTGGAGGATGACGATAAGT |
| Human | CHD7 | Chromatin Readers | AAATGAACTGAAAGGTGTTG |
| Human | CHD7 | Chromatin Readers | AGAAACCGATGGCAGCGGGA |
| Human | NCOA3 | Histone Acetylation | ATTGTCAATATCACTAAGAT |
| Human | NCOA3 | Histone Acetylation | ATACCTGCAATATAAGCAAG |
| Human | NCOA3 | Histone Acetylation | GCAATATAAGCAAGAGGACC |
| Human | NCOA3 | Histone Acetylation | ATTCCTGAACTTGTCAATCA |
| Human | NCOA3 | Histone Acetylation | TGGCCCAACAAGATCATCCA |
| Human | NCOA3 | Histone Acetylation | TGAACTTGTCAATCAGGGAC |
| Human | NCOA3 | Histone Acetylation | GTGGCCCAACAAGATCATCC |
| Human | NCOA3 | Histone Acetylation | TCCAGGGAATTCCTAAGAAG |
| Human | NCOA3 | Histone Acetylation | GTCATCACTTCCGACAACAG |
| Human | NCOA3 | Histone Acetylation | TGGACAGACATACCCAGCAC |
| Human | NCOA3 | Histone Acetylation | TCATCACTTCCGACAACAGA |
| Human | NCOA3 | Histone Acetylation | CCCTGCTGGGAGCTCACCTG |
| Human | NCOA3 | Histone Acetylation | TTTACAACTTGAATCTAGGG |
| Human | NCOA3 | Histone Acetylation | ACCTGTTCTTCTGATGACCG |
| Human | NCOA3 | Histone Acetylation | GTATTGCCAGAAGATGCCAT |
| Human | NCOA3 | Histone Acetylation | AATGGCAGCAGTGGTCACAT |
| Human | NCOA3 | Histone Acetylation | GAATGGCAGCAGTGGTCACA |
| Human | PHF20 | Chromatin Readers | CGAGGAATCAGCTTTGAAGT |
| Human | PHF20 | Chromatin Readers | ACGAGGAATCAGCTTTGAAG |
| Human | PHF20 | Chromatin Readers | GGAACCATCGTTATGATGAG |
| Human | PHF20 | Chromatin Readers | GACAATCAGACCAGCAAGCA |
| Human | PHF20 | Chromatin Readers | TTCCAAAGATCAGAATATTG |
| Human | PHF20 | Chromatin Readers | CTTGTTAACAGCAGTGACTT |
| Human | PHF20 | Chromatin Readers | TCCAAAGATCAGAATATTGT |
| Human | PHF20 | Chromatin Readers | TGAATTTCAAATAAATGAGC |
| Human | PHF20 | Chromatin Readers | CATGTGATAGTGCAACAGTT |
| Human | PHF20 | Chromatin Readers | AGTGCAACAGTTTGGCTTTG |
| Human | PHF20 | Chromatin Readers | AGTGTGAAGAGTGCCAGTGC |
| Human | PHF20 | Chromatin Readers | ACATAACAGGTGTATTTCTC |
| Human | PHF20 | Chromatin Readers | GTGCCAGTGCTGGCAGCATG |
| Human | PHF20 | Chromatin Readers | AAGTCTTCCCAAGAACGAGA |
| Human | PHF20 | Chromatin Readers | AAAGTTAATCTGTTCTGAAA |
| Human | PHF20 | Chromatin Readers | CCCAAGCAGCCTGATAAAGA |
| Human | PHF20 | Chromatin Readers | CCTTCTTTATCAGGCTGCTT |
| Human | PHF20 | Chromatin Readers | GAATAAAGAGAAGAAATTCA |
| Human | FOXP3 | DNA and RNA Modifications | GTAGGGTTGGAACACCTGCT |
| Human | FOXP3 | DNA and RNA Modifications | TGTAGGGTTGGAACACCTGC |
| Human | FOXP3 | DNA and RNA Modifications | GGACTTCCTCAAGCACTGCC |

| | | | |
|---|---|---|---|
| Human | FOXP3 | DNA and RNA Modifications | CAGTGCTTGAGGAAGTCCTC |
| Human | FOXP3 | DNA and RNA Modifications | GAAGGTCTTCGAAGAGCCAG |
| Human | FOXP3 | DNA and RNA Modifications | CAAGTGGCCCGGATGTGAGA |
| Human | FOXP3 | DNA and RNA Modifications | TCGAAGACCTTCTCACATCC |
| Human | FOXP3 | DNA and RNA Modifications | GCGGATGGCGTTCTTCCAGG |
| Human | FOXP3 | DNA and RNA Modifications | GAGCCTCCAGGATGGCCCAG |
| Human | FOXP3 | DNA and RNA Modifications | GATGGCCCAGCGGATGAGCG |
| Human | FOXP3 | DNA and RNA Modifications | CCTCCCACCTGGGATCAACG |
| Human | FOXP3 | DNA and RNA Modifications | TGGCCACGTTGATCCCAGGT |
| Human | FOXP3 | DNA and RNA Modifications | CCACGTTGATCCCAGGTGGG |
| Human | FOXP3 | DNA and RNA Modifications | CTGGCCACGTTGATCCCAGG |
| Human | FOXP3 | DNA and RNA Modifications | CAACGGTGGATGCCCACGCC |
| Human | FOXP3 | DNA and RNA Modifications | TCCTACCCACTGCTGGCAAA |
| Human | FOXP3 | DNA and RNA Modifications | GGAGAAGCTGAGTGCCATGC |
| Human | FOXP3 | DNA and RNA Modifications | GCAGCAGCTGGTGCTGGAGA |
| Human | FOXP3 | DNA and RNA Modifications | TCTGGAGCAGCAGCTGGTGC |
| Human | FOXP3 | DNA and RNA Modifications | CCCAGAGTTCCTCCACAACA |
| Human | FOXP3 | DNA and RNA Modifications | ACAGCCTGTTTGCTGTCCGG |
| Human | FOXP3 | DNA and RNA Modifications | GGACAGCAAACAGGCTGTCA |
| Human | FOXP3 | DNA and RNA Modifications | TGAAGTAGTCCATGTTGTGG |
| Human | FOXP3 | DNA and RNA Modifications | GCTGTCCGGAGGCACCTGTG |
| Human | ERCC3 | Transcription Elongation | AGGGACTTTCCAGCACCGCA |
| Human | ERCC3 | Transcription Elongation | TGCAGCAGTCACACCAACCA |
| Human | ERCC3 | Transcription Elongation | ACCAGCCAAGATGTTCCGAA |
| Human | ERCC3 | Transcription Elongation | AGGCCGAGCGAGTCATGGAG |
| Human | ERCC3 | Transcription Elongation | CAGTCAAACCCAGCTTACAG |
| Human | ERCC3 | Transcription Elongation | ACCATCTTCATATCCAAGGT |
| Human | ERCC3 | Transcription Elongation | ACCGCGACAAGAAGAAATCC |
| Human | ERCC3 | Transcription Elongation | TCCTGGATTTCTTCTTGTCG |
| Human | UBR3 | Other Histone Modifications | TTACCGATGCATTGATCAAA |
| Human | UBR3 | Other Histone Modifications | GAACATGTTGAAGTCGTGTC |
| Human | UBR3 | Other Histone Modifications | GCAGCGGTAGGCCACGAAGT |
| Human | UBR3 | Other Histone Modifications | GGAATCATTACGGAATGACC |
| Human | UBR3 | Other Histone Modifications | GGTGTTTATGTACAAACCTG |
| Human | UBR3 | Other Histone Modifications | CTTGGCAGTATCAATTGGCT |
| Human | UBR3 | Other Histone Modifications | AATCTTACATGGAATCATTA |
| Human | UBR3 | Other Histone Modifications | GCAGTATCAATTGGCTGGGA |
| Human | UBR3 | Other Histone Modifications | GAGGCAGCAGAAATTGCTTG |
| Human | UBR3 | Other Histone Modifications | GGCGACAGAAGGCTAGAGAG |
| Human | UBR3 | Other Histone Modifications | CAAAGAAGAAAGGCGACAGA |
| Human | UBR3 | Other Histone Modifications | AAACATTGGACAAAGAAGAA |
| Human | UBR3 | Other Histone Modifications | AAGGCACTTACACCACTCCA |

| | | | |
|---|---|---|---|
| Human | UBR3 | Other Histone Modifications | CGGCTACGACGAGTTCTGCG |
| Human | UBR3 | Other Histone Modifications | GTGGTGTAAGTGCCTTCTGG |
| Human | UBR3 | Other Histone Modifications | AGCCGGCCGGACAACCGCGC |
| Human | UBR3 | Other Histone Modifications | ACGAGTTCTGCGCGGCGGTG |
| Human | UBR3 | Other Histone Modifications | AGCAGGATCTTTAGGAACCT |
| Human | UBR3 | Other Histone Modifications | CCACACACAAGGCAAACAGC |
| Human | UBR3 | Other Histone Modifications | GCTTGCAGCAAAGTCCTTTC |
| Human | CBX3 | Chromatin Readers | TACAGATGCTGACAATACTT |
| Human | CBX3 | Chromatin Readers | CTAGATCGACGTGTAGTGAA |
| Human | CBX3 | Chromatin Readers | GAATATTTCCTGAAGTGGAA |
| Human | CBX3 | Chromatin Readers | AAGTGGAATATTTCCTGAAG |
| Human | CBX3 | Chromatin Readers | GAAATGGAAAGATTCAGATG |
| Human | CBX3 | Chromatin Readers | AGACTTGGTGCTGGCGAAAG |
| Human | CBX3 | Chromatin Readers | ATTGGTGCCACAGACAGCAG |
| Human | CBX3 | Chromatin Readers | AGATGAGGCAGACTTGGTGC |
| Human | BTBD18 | Transcription Elongation | GCCTGAAGATCAGCACACTT |
| Human | BTBD18 | Transcription Elongation | CCTCACCTTCTGCCTGCAGA |
| Human | BTBD18 | Transcription Elongation | CCTTCTGCAGGCAGAAGGTG |
| Human | BTBD18 | Transcription Elongation | GAGCGGGAGAGGCCAGCTCA |
| Human | BTBD18 | Transcription Elongation | CAGCACACTTAGGAAGCTGG |
| Human | CDYL2 | Chromatin Readers | GAAAGGCTACGGGAGCACCG |
| Human | CDYL2 | Chromatin Readers | GAGTATCTTATCCGATGGAA |
| Human | CDYL2 | Chromatin Readers | TTATCCGATGGAAAGGCTAC |
| Human | CDYL2 | Chromatin Readers | ACGGGAGCACCGAGGACACG |
| Human | CDYL2 | Chromatin Readers | ACTCCTCACAGTGCAAGAGG |
| Human | CENPO | Chromatin Remodeling | GTTTACGGGATCTGCTCACT |
| Human | CENPO | Chromatin Remodeling | TTTACGGGATCTGCTCACTT |
| Human | CENPO | Chromatin Remodeling | CAGCTCTTCAGACTGTTTAC |
| Human | CENPO | Chromatin Remodeling | GCAGCTCTTCAGACTGTTTA |
| Human | CENPO | Chromatin Remodeling | GATCAATGAGCAAGAAGCAT |
| Human | CENPO | Chromatin Remodeling | AAATGTGAAAGCCATTCTGC |
| Human | CENPO | Chromatin Remodeling | AGAAGCATTGGAAGAGAAAT |
| Human | PKN1 | Other Histone Modifications | AGGGTGCTGAGAACCTGCGG |
| Human | PKN1 | Other Histone Modifications | AGGAGGGTGCTGAGAACCTG |
| Human | PKN1 | Other Histone Modifications | GCTGCGGCCCAGGTCAGTGG |
| Human | PKN1 | Other Histone Modifications | GCTGCGGCGGGAAATCCGCA |
| Human | PKN1 | Other Histone Modifications | GCAATGGCAGCACCAAGGAC |
| Human | PKN1 | Other Histone Modifications | TCATCCGCATGCAACTCCGC |
| Human | PKN1 | Other Histone Modifications | GCGCCCGGCGGAGTTGCATG |
| Human | PKN1 | Other Histone Modifications | ATGATCCAGACCTACAGCAA |
| Human | PKN1 | Other Histone Modifications | GCTTCTCCAGGCCCGCCACG |
| Human | PKN1 | Other Histone Modifications | GCTGCGGCACCACTTCCGAG |

| | | | |
|---|---|---|---|
| Human | PKN1 | Other Histone Modifications | CCCAAGGGAGTCCTGACCTG |
| Human | PKN1 | Other Histone Modifications | ACAGAATCCAACCAGAAGCT |
| Human | PKN1 | Other Histone Modifications | ACCTCCCAGAGACCATCCCG |
| Human | PKN1 | Other Histone Modifications | AAGCACAGTGCTGACTTCAC |
| Human | PKN1 | Other Histone Modifications | CTGGGACCAGAGCTTCACTC |
| Human | PKN1 | Other Histone Modifications | TCTGGAGCTGGAAAGGGCAC |
| Human | PKN1 | Other Histone Modifications | CGAACACAAGATCGTCTACA |
| Human | PKN1 | Other Histone Modifications | TCATGAGAAGGCTGCTTCGG |
| Human | PKN1 | Other Histone Modifications | GTGTGAGAAGCGGATATTGG |
| Human | PKN1 | Other Histone Modifications | CGAAGTTGCTGACGTCGGTG |
| Human | PKN1 | Other Histone Modifications | TCGGTGCGGCCGGACAGCGT |
| Human | PKN1 | Other Histone Modifications | AACTTCGACGAGGAGTTCAC |
| Human | PKN1 | Other Histone Modifications | GACAGCGTGGGCACAAAGGG |
| Human | MINA | Histone Methylation | ACCTACCAGAACAATTCATG |
| Human | MINA | Histone Methylation | AGCTCATCCTTAAATCTCTG |
| Human | MINA | Histone Methylation | ATCAAGCGGACACTCCTGCG |
| Human | MINA | Histone Methylation | GGCACGAGAGTACAGCGTGG |
| Human | MINA | Histone Methylation | GACGATTAAGTGGCTTCCTG |
| Human | MINA | Histone Methylation | GCCGGTCTGCAAGTGTCCTC |
| Human | MINA | Histone Methylation | TCCTGAGGACACTTGCAGAC |
| Human | ARID4B | Chromatin Readers | GGATTTGATAATATTGAAAG |
| Human | ARID4B | Chromatin Readers | CCTGCAGCTGAATTTAAGAC |
| Human | ARID4B | Chromatin Readers | AAATTTCGATATCCAAGTAC |
| Human | ARID4B | Chromatin Readers | CTGCAGCTGAATTTAAGACA |
| Human | ARID4B | Chromatin Readers | AGACTTGTACACAAACTTGG |
| Human | ARID4B | Chromatin Readers | CAGAAATGATCTTATTTCAA |
| Human | ARID4B | Chromatin Readers | AATAAGATCATTTCTGTTGT |
| Human | ARID4B | Chromatin Readers | ACGCTTTCTCCTCCGATCAA |
| Human | ARID4B | Chromatin Readers | CTGAAGTAGCTTCCATTGAT |
| Human | ARID4B | Chromatin Readers | CAAAGATGATGACACAACTA |
| Human | ARID4B | Chromatin Readers | ACAAAGATGATGACACAACT |
| Human | ARID4B | Chromatin Readers | GCTGTTATGGAACTGGATGA |
| Human | ARID4B | Chromatin Readers | ACTGGATGAAGGTGAAGAGG |
| Human | ARID4B | Chromatin Readers | TAACATAACAGCAGCTGTTA |
| Human | ARID4B | Chromatin Readers | GGAACTGGATGAAGGTGAAG |
| Human | ARID4B | Chromatin Readers | AACAGCAGCTGTTATGGAAC |
| Human | TET2 | DNA and RNA Modifications | GGAGAAAGAAGCAATTGTGA |
| Human | TET2 | DNA and RNA Modifications | GAAAGAAGCAATTGTGATGG |
| Human | TET2 | DNA and RNA Modifications | GAACCTGTGGAAGAGGAGGA |
| Human | TET2 | DNA and RNA Modifications | AGAAGCAATTGTGATGGTGG |
| Human | TET2 | DNA and RNA Modifications | TGAGGAACCTGTGGAAGAGG |
| Human | TET2 | DNA and RNA Modifications | CTGCGCAACTTGCTCAGCAA |

| | | | |
|---|---|---|---|
| Human | TET2 | DNA and RNA Modifications | ATGAATCAAGGGCAGTCCCA |
| Human | TET2 | DNA and RNA Modifications | CAAGTGCTGTTTCAACACTG |
| Human | TET2 | DNA and RNA Modifications | TGTGGTGGCTGCTTCTGTAG |
| Human | TET2 | DNA and RNA Modifications | TGGCTGCTTCTGTAGAGGCT |
| Human | TET2 | DNA and RNA Modifications | CGGAGCTTACCGAGACGCTG |
| Human | TET2 | DNA and RNA Modifications | TCTGGCTGACAAACTCTACT |
| Human | TET2 | DNA and RNA Modifications | ATATGTTGGTGCCATAAGAG |
| Human | TET2 | DNA and RNA Modifications | TTGGTGCCATAAGAGTGGAC |
| Human | PSMA4 | Control | AACTAAGGCTCATTGCTCAA |
| Human | PSMA4 | Control | ATGTTCTGACTAATGAACTA |
| Human | AXIN1 | Transcription Elongation | AGACTTAAGGAAGGAGGGAT |
| Human | AXIN1 | Transcription Elongation | ATAAATATCAGACTTAAGGA |
| Human | AXIN1 | Transcription Elongation | AAACCAGAAGTCCAGCAAGT |
| Human | AXIN1 | Transcription Elongation | CCCTGCTGGATGACCAAGAT |
| Human | AXIN1 | Transcription Elongation | TGAGCGACGAGTTTGACTGT |
| Human | AXIN1 | Transcription Elongation | CTCCTCAAAGACGGGCAGGA |
| Human | AXIN1 | Transcription Elongation | TGTGGCGTACTACTTCTGCG |
| Human | AXIN1 | Transcription Elongation | GTTGTGGCGTACTACTTCTG |
| Human | AXIN1 | Transcription Elongation | AGTTCAGGTATGGATCCTGG |
| Human | AXIN1 | Transcription Elongation | TTCGCTGTACCGTCTACTGG |
| Human | AXIN1 | Transcription Elongation | TCCAGTAGACGGTACAGCGA |
| Human | AXIN1 | Transcription Elongation | GAAGGCAGAGAGTTCAGGTA |
| Human | AXIN1 | Transcription Elongation | GCTGAAGCGCGTGCGCATGG |
| Human | AXIN1 | Transcription Elongation | GCGCATGGAGGAGGAAGGTG |
| Human | AXIN1 | Transcription Elongation | CCTTCGGCACCCGGTACGTG |
| Human | AXIN1 | Transcription Elongation | AGCACGTACAGCGTGTGCTG |
| Human | AXIN1 | Transcription Elongation | CGGGCTCCGGGATGCACACG |
| Human | AXIN1 | Transcription Elongation | CACAGCCCATGTCCACACAG |
| Human | AXIN1 | Transcription Elongation | GGTTCTCCTCGTGTGCATCC |
| Human | AXIN1 | Transcription Elongation | GGACCGGCCACGGGTCTTCG |
| Human | AXIN1 | Transcription Elongation | GAAGCACGTACCCAAGTCAG |
| Human | AXIN1 | Transcription Elongation | AGTGATGGCCTCGCCCACAG |
| Human | AXIN1 | Transcription Elongation | AGACAGAGACAAGATCGCAG |
| Human | AXIN1 | Transcription Elongation | GCCGTGTGACAGCATCGTTG |
| Human | AXIN1 | Transcription Elongation | GCCACAACGATGCTGTCACA |
| Human | AXIN1 | Transcription Elongation | GGAGAGCTCCATGTCCGACA |
| Human | AXIN1 | Transcription Elongation | GGTACCAGCCGTGTCGGACA |
| Human | NELFA | Transcription Elongation | GGAGCTGCTGCAGAAGTCCA |
| Human | PYGO2 | Chromatin Readers | CACTCCGACAGGCACCACAT |
| Human | PYGO2 | Chromatin Readers | ATCGTTCACCTCACTCCGAC |
| Human | PYGO2 | Chromatin Readers | GCAGGAGGCCTCACACAGAA |
| Human | PYGO2 | Chromatin Readers | ATGTGGTGCCTGTCGGAGTG |

| | | | |
|---|---|---|---|
| Human | PYGO2 | Chromatin Readers | CGAAGGAAGTCAAATACTCA |
| Human | PYGO2 | Chromatin Readers | GCAGGGCCCATAGGATTGGG |
| Human | PRDM15 | Histone Methylation | TCCTTCTGGAACACCTTCAG |
| Human | PRDM15 | Histone Methylation | GTGCTTCGACACCTCCAACG |
| Human | PRDM15 | Histone Methylation | TTGGAGGTGTCGAAGCACAC |
| Human | PRDM15 | Histone Methylation | CATTGAGGAGCTGCAGCTCC |
| Human | PRDM15 | Histone Methylation | GAGGAGCTGCAGCTCCAGGA |
| Human | PRDM15 | Histone Methylation | GAGCTGCAGCTCCAGGAAGG |
| Human | PRDM15 | Histone Methylation | AGTGTGTTCTGCCACCTTCC |
| Human | PRDM15 | Histone Methylation | AGAACAGCAGCAACCTGAGC |
| Human | PRDM15 | Histone Methylation | TGCTCAGGTTGCTGCTGTTC |
| Human | PRDM15 | Histone Methylation | GCTGCTTTAGGCTCTCTTTG |
| Human | PRDM15 | Histone Methylation | GTGCTTGGAGAGGTTGCTGT |
| Human | PRDM15 | Histone Methylation | CTTGGAGAGGTTGCTGTTGG |
| Human | PRDM15 | Histone Methylation | CAGCAAGATGTTCTACCGCA |
| Human | PRDM15 | Histone Methylation | GGTCCAGCATGACGTCCTTG |
| Human | PRDM15 | Histone Methylation | CTACCGCAAGGACGTCATGC |
| Human | PRDM15 | Histone Methylation | CATCTTGCTGCAGACCTCAC |
| Human | PRDM15 | Histone Methylation | AGAACACCTTGCCACACACC |
| Human | PRDM15 | Histone Methylation | GTGGCAAGGTGTTCTCCTGC |
| Human | PRDM15 | Histone Methylation | GAGAACACCTTGCCACACAC |
| Human | PRDM15 | Histone Methylation | GGTGCTTGTTCATATTGCTC |
| Human | PRDM15 | Histone Methylation | CTTGTTCATATTGCTCCGGC |
| Human | PRDM15 | Histone Methylation | CGGGCGCAAGTTCTTCCGCG |
| Human | PRDM15 | Histone Methylation | GGTCCCTGAGCACATCCACG |
| Human | PRDM15 | Histone Methylation | CTTGCGCCCGCAGATCTCGC |
| Human | PRDM15 | Histone Methylation | TCTTCCGCGTGGATGTGCTC |
| Human | PRDM15 | Histone Methylation | CTTCCGCGTGGATGTGCTCA |
| Human | PRDM15 | Histone Methylation | AAGCGGTGCCAGCTCACCTT |
| Human | PRDM15 | Histone Methylation | GGAGTACCTGAAGCACATCA |
| Human | PRDM15 | Histone Methylation | GGTGCCAGCTCACCTTCGGC |
| Human | PRDM15 | Histone Methylation | GCCAGCTCACCTTCGGCCGG |
| Human | PRDM15 | Histone Methylation | GTGCCAGCTCACCTTCGGCC |
| Human | PRDM15 | Histone Methylation | TGGCCTTCAGTGCAAAGCGC |
| Human | PRDM15 | Histone Methylation | GGCCACCTACCACGCCCACA |
| Human | PRDM15 | Histone Methylation | CAACCGGCGCTTTGCACTGA |
| Human | PRDM15 | Histone Methylation | GGGCGGATCTTCAACAGCAT |
| Human | PRDM15 | Histone Methylation | ATCCGCCCGCAGATCTCGCA |
| Human | PRDM15 | Histone Methylation | GGCGGATCTTCAACAGCATC |
| Human | PRDM15 | Histone Methylation | GATCCGCCCGCAGATCTCGC |
| Human | PRDM15 | Histone Methylation | CTTCAACAGCATCGGGAACC |
| Human | PRDM15 | Histone Methylation | TGGGAAGTCCTTTGCCAGGA |

| | | | |
|---|---|---|---|
| Human | PRDM15 | Histone Methylation | TGCCAGGAAGGACATGCTGA |
| Human | PRDM15 | Histone Methylation | GGACTTCCCACACTGCTCGC |
| Human | PRDM15 | Histone Methylation | CAGCATGTCCTTCCTGGCAA |
| Human | PRDM15 | Histone Methylation | AGTGTGGGAAGTCCTTTGCC |
| Human | PRDM15 | Histone Methylation | CTTCATGCCTTTCCCACACT |
| Human | PRDM15 | Histone Methylation | GTGGTGGCGCAGCGCGTGCT |
| Human | PRDM15 | Histone Methylation | TGACCTTCTGCGCGAACCTG |
| Human | PRDM15 | Histone Methylation | CCACCGCAGGTTCGCGCAGA |
| Human | PRDM15 | Histone Methylation | CCTTCTGCGCGAACCTGCGG |
| Human | PRDM15 | Histone Methylation | AGTGCAAGGAGTGCCACCGC |
| Human | PRDM15 | Histone Methylation | ATTCAGCGAGAGGAACACCA |
| Human | PRDM15 | Histone Methylation | GTGGGAAGACATTCAGCGAG |
| Human | PRDM15 | Histone Methylation | CACGTACTTCTTGTCGCACA |
| Human | PRDM15 | Histone Methylation | GTGCTTCTGCAGCATGTACT |
| Human | PRDM15 | Histone Methylation | CCTCCATGAGCCGACACATG |
| Human | PRDM15 | Histone Methylation | CCGCATGTGTCGGCTCATGG |
| Human | PRDM15 | Histone Methylation | CGGGACCAAGGTGTCCACCA |
| Human | PRDM15 | Histone Methylation | ACACCTTGGTCCCGCACAGC |
| Human | PRDM15 | Histone Methylation | GCGGGACCAAGGTGTCCACC |
| Human | MBD5 | Chromatin Readers | GTGCTTTATGTCAGTCCCAG |
| Human | MBD5 | Chromatin Readers | ACTGATGGAACATGCAAGTG |
| Human | MBD5 | Chromatin Readers | TGCATGTTCCATCAGTAAGC |
| Human | MBD5 | Chromatin Readers | CAAGATAACAAAGACCCACT |
| Human | MBD5 | Chromatin Readers | TGCTTTATGTCAGTCCCAGT |
| Human | MBD5 | Chromatin Readers | GCTCCACCTTCCCTTCCTCG |
| Human | MBD5 | Chromatin Readers | AAGTTGAAGACACTAACAGA |
| Human | MBD5 | Chromatin Readers | ACATCTGAGAAAGATCCACT |
| Human | MBD5 | Chromatin Readers | GGCATGCTTACATTCATCAT |
| Human | MBD5 | Chromatin Readers | TCATCATAGGTACCTGAGAG |
| Human | MBD5 | Chromatin Readers | TGGTTGGATTAATGATAACA |
| Human | MBD5 | Chromatin Readers | TTCTGAAGAGCACAAGAAGG |
| Human | CCNT2 | Transcription Elongation | TCACATTTAGTATCCAGCAG |
| Human | CCNT2 | Transcription Elongation | ACCATAATGCTACAAACTCT |
| Human | CCNT2 | Transcription Elongation | CTGCGATTGTTTATATGCAC |
| Human | CCNT2 | Transcription Elongation | GCCGAGCCGCCGCTGCGGAG |
| Human | CCNT2 | Transcription Elongation | CCGCCTCCACTCCGCAGCGG |
| Human | CCNT2 | Transcription Elongation | GGTTGTAAGATGCAGACTGT |
| Human | CCNT2 | Transcription Elongation | AATAATTCTAGAGTAACTGT |
| Human | CCNT2 | Transcription Elongation | GGAGGGCTAATCAGGCAGCT |
| Human | CCNT2 | Transcription Elongation | TATCAACTGATGGAAAGCAT |
| Human | CCNT2 | Transcription Elongation | TATATACTGCTTAACAGAGG |
| Human | CCNT2 | Transcription Elongation | ACAGGACTCCTCAGAACAGT |

| | | | |
|---|---|---|---|
| Human | CCNT2 | Transcription Elongation | GAATACCACCCACTGTTCTG |
| Human | JADE2 | Chromatin Readers | GGCATGCTACGGGATCCTCA |
| Human | JADE2 | Chromatin Readers | GTGTGCATCAGGCATGCTAC |
| Human | JADE2 | Chromatin Readers | GGGATCCTCAAGGTGCCCAC |
| Human | JADE2 | Chromatin Readers | TGTGTGCATCAGGCATGCTA |
| Human | JADE2 | Chromatin Readers | GGGCGAGGATGGCAACGAGA |
| Human | JADE2 | Chromatin Readers | CACTAGAAGTGGGACCAAGT |
| Human | JADE2 | Chromatin Readers | CCACTAGAAGTGGGACCAAG |
| Human | JADE2 | Chromatin Readers | CCTTGAAGCCCACTAGAAGT |
| Human | JADE2 | Chromatin Readers | TCAAGGCTCCTCCTCGCTTG |
| Human | JADE2 | Chromatin Readers | ACAGCTGACATGCACCCACT |
| Human | JADE2 | Chromatin Readers | CGCAGGAAGGCATGGAACAC |
| Human | JADE2 | Chromatin Readers | GGTCAAAGGCGCATGTGACA |
| Human | JADE2 | Chromatin Readers | TGGAACGCTGTGACGCAGGA |
| Human | JADE2 | Chromatin Readers | AGGCATGGAACACTGGATGC |
| Human | JADE2 | Chromatin Readers | CGGGTTACCTTGCTCTCGCG |
| Human | JADE2 | Chromatin Readers | AAGTGAGAAGATGTCCGACG |
| Human | ELP3 | Histone Acetylation | CTCTCCTTACCAATTCAACC |
| Human | ELP3 | Histone Acetylation | CCTACTAAATTTCAGCATCA |
| Human | NCBP2 | Transcription Elongation | GTACATAAATGGGACGCGTC |
| Human | NCBP2 | Transcription Elongation | CCGAATCATTCGCACAGACT |
| Human | FOXP2 | DNA and RNA Modifications | GAAAGCATTTGTGAAGATTT |
| Human | FOXP2 | DNA and RNA Modifications | TCACAAATGCTTTCACAGCC |
| Human | FOXP2 | DNA and RNA Modifications | TTATGCAACTCTCATAAGGC |
| Human | FOXP2 | DNA and RNA Modifications | CTATCATGGAGTCATCTGAC |
| Human | FOXP2 | DNA and RNA Modifications | AGAGTTGCATAAGTAAATGG |
| Human | FOXP2 | DNA and RNA Modifications | AAATTTACAGCTGGTTTACA |
| Human | FOXP2 | DNA and RNA Modifications | TTACTTATGCAACTCTCATA |
| Human | FOXP2 | DNA and RNA Modifications | GCAGGACTTAAGCCAGCTTG |
| Human | FOXP2 | DNA and RNA Modifications | ACTGCAGGTGCCTGTGTCAG |
| Human | FOXP2 | DNA and RNA Modifications | TGATAAACAGAGACCACTGC |
| Human | FOXP2 | DNA and RNA Modifications | ACTGACACAGGCACCTGCAG |
| Human | FOXP2 | DNA and RNA Modifications | GGACTCATCTCCATTCCACC |
| Human | MTF2 | Chromatin Readers | AGCCATCTGACCATCTAGCT |
| Human | MTF2 | Chromatin Readers | AATGTCCTTCCAGAGAACCC |
| Human | MTF2 | Chromatin Readers | GACATTCAAACAGGAGCCAC |
| Human | MTF2 | Chromatin Readers | CTCTGGAAGGACATTCAAAC |
| Human | MTF2 | Chromatin Readers | GTCCTAGCTAGATGGTCAGA |
| Human | MTF2 | Chromatin Readers | ATGGTTATATGTGACAAGTG |
| Human | MTF2 | Chromatin Readers | GACACAACTGATGATATCCT |
| Human | MTF2 | Chromatin Readers | TTCATCTGAATCAATCACAC |
| Human | MTF2 | Chromatin Readers | TCTTCAAATACCAGTCTCCA |

| | | | |
|---|---|---|---|
| Human | MTF2 | Chromatin Readers | AGTGCTGCAAATGTAAGCAG |
| Human | MTF2 | Chromatin Readers | ATCTTCAAATACCAGTCTCC |
| Human | MTF2 | Chromatin Readers | AATCTGTCTCCAAATAGCAT |
| Human | MTF2 | Chromatin Readers | ATTGTGGAGGCCCTGGAGAC |
| Human | MTF2 | Chromatin Readers | CACTTAGGTTGTAAAGGCAT |
| Human | MTF2 | Chromatin Readers | AAGATATGAGCATGTTCTGG |
| Human | MTF2 | Chromatin Readers | TGAAAGATATGAGCATGTTC |
| Human | YWHAB | Chromatin Readers | TGATATGGCTGCAGCCATGA |
| Human | YWHAB | Chromatin Readers | TGACACGCCAGGAAGAGCGG |
| Human | YWHAB | Chromatin Readers | GGCGCCTACCACATTCTTGT |
| Human | YWHAB | Chromatin Readers | AGATGACACGCCAGGAAGAG |
| Human | YWHAB | Chromatin Readers | GTTGCCTACAAGAATGTGGT |
| Human | YWHAB | Chromatin Readers | GTTCTGCCTCTATCTTCTCA |
| Human | YWHAB | Chromatin Readers | GGACATCTGCAATGATGTTC |
| Human | YWHAB | Chromatin Readers | GAAGATAGAGGCAGAACTGC |
| Human | YWHAB | Chromatin Readers | CACTGTGTCGAACTCCCAGC |
| Human | YWHAB | Chromatin Readers | GAACTCCCAGCAGGCTTACC |
| Human | YWHAB | Chromatin Readers | TGCTTCCTGGTAAGCCTGCT |
| Human | YWHAB | Chromatin Readers | ATGCTTCCTGGTAAGCCTGC |
| Human | YWHAB | Chromatin Readers | TCTGATCATGCAGTTACTTA |
| Human | YWHAB | Chromatin Readers | CTCTGATCATGCAGTTACTT |
| Human | HDAC3 | Histone Acetylation | CATACATGTCACCTGTGCCA |
| Human | HDAC3 | Histone Acetylation | CACTGGAGCACAATGCACGT |
| Human | HDAC3 | Histone Acetylation | CCAGAGAGTCAGCTCCACAC |
| Human | HDAC3 | Histone Acetylation | ATGTCTCATATGTCCAGCAG |
| Human | HDAC3 | Histone Acetylation | ACGTGCATTGTGCTCCAGTG |
| Human | HDAC3 | Histone Acetylation | CAAAGATTGTCTGGCGGATC |
| Human | SETD5 | Histone Methylation | TGCGACACATGATTGCAGAT |
| Human | SETD5 | Histone Methylation | CATGTTACGACAGCAATTTG |
| Human | SETD5 | Histone Methylation | CTCTTATAATAGAGTATCGT |
| Human | SETD5 | Histone Methylation | ACTCTTATAATAGAGTATCG |
| Human | SETD5 | Histone Methylation | AATGACGACCTAGAGTTCTG |
| Human | SETD5 | Histone Methylation | TGTGATGTTACTCTCCCATG |
| Human | SART3 | Other Histone Modifications | CAGCCAGAGCTCTTCAGTCA |
| Human | SART3 | Other Histone Modifications | CTTGACTGAAGAGCTCTGGC |
| Human | SART3 | Other Histone Modifications | AGATTTCACTCATCTTCTGG |
| Human | SART3 | Other Histone Modifications | CTGAAGAGCTCTGGCTGGAG |
| Human | SART3 | Other Histone Modifications | AGCCAGAGCTCTTCAGTCAA |
| Human | SART3 | Other Histone Modifications | CCTTCACGGCTTTCTCAAAG |
| Human | SART3 | Other Histone Modifications | CCATACTCTAGCCAAATGTT |
| Human | SART3 | Other Histone Modifications | ATTACATTTGTCCTAACATT |
| Human | SART3 | Other Histone Modifications | AGGACAAATGTAATCCTTCA |

| | | | |
|---|---|---|---|
| Human | SART3 | Other Histone Modifications | CCTCTTTGAGAAAGCCGTGA |
| Human | SART3 | Other Histone Modifications | ACACGGAGCGAACTTTCTCA |
| Human | SART3 | Other Histone Modifications | GTTGGTTTACATATGACCAA |
| Human | SART3 | Other Histone Modifications | CTCTCGGTAAGCCTCCCAGA |
| Human | SART3 | Other Histone Modifications | ACTCTCGGTAAGCCTCCCAG |
| Human | SART3 | Other Histone Modifications | TGACCAAAGGACTCGCCCTC |
| Human | SART3 | Other Histone Modifications | GATCCCACTCTATGATATGG |
| Human | SART3 | Other Histone Modifications | TGGCCTCCATATCATAGAGT |
| Human | SART3 | Other Histone Modifications | GTGGCCTCCATATCATAGAG |
| Human | SART3 | Other Histone Modifications | GGCGATCCCACTCTATGATA |
| Human | SART3 | Other Histone Modifications | TGTAACGGATCCATAAGTCT |
| Human | SART3 | Other Histone Modifications | CTGTAACGGATCCATAAGTC |
| Human | SART3 | Other Histone Modifications | GGATCCATAAGTCTGGGACA |
| Human | SART3 | Other Histone Modifications | GGACAAGGCAGTTCTCGACC |
| Human | SART3 | Other Histone Modifications | GGACAGTTGCCTTATGGAGT |
| Human | SART3 | Other Histone Modifications | TCCATAAGGCAACTGTCCAG |
| Human | SART3 | Other Histone Modifications | CTTATGGAGTCGGTACCTCT |
| Human | SART3 | Other Histone Modifications | CAAGAGGTACCGACTCCATA |
| Human | SART3 | Other Histone Modifications | GGCATTCAAAGCTTTCTCGA |
| Human | SART3 | Other Histone Modifications | CCAAATCTCCACATAATCAG |
| Human | SART3 | Other Histone Modifications | CATCCAGGCCACTGATTATG |
| Human | SART3 | Other Histone Modifications | TGATTATGTGGAGATTTGGC |
| Human | SART3 | Other Histone Modifications | TCTCCACATAATCAGTGGCC |
| Human | SART3 | Other Histone Modifications | GTGGCTAGAGTATTACAACC |
| Human | SART3 | Other Histone Modifications | GCAGAAAGCTCGGGAACTCT |
| Human | SART3 | Other Histone Modifications | GTAATACTCTAGCCACATGT |
| Human | SART3 | Other Histone Modifications | ATGCCAAGTACGCCAACATG |
| Human | SART3 | Other Histone Modifications | ACTCCACGTAGCAGTAACCT |
| Human | SART3 | Other Histone Modifications | GACCCATCTTCAGCAACCGT |
| Human | SART3 | Other Histone Modifications | CGACCCATCTTCAGCAACCG |
| Human | SART3 | Other Histone Modifications | ACCAAAGGGCCTGGCCTACG |
| Human | SART3 | Other Histone Modifications | GGAACTAGAAGAAATCTGTA |
| Human | SART3 | Other Histone Modifications | AACCGGGCTGGCAAACCAAA |
| Human | SART3 | Other Histone Modifications | CCTCAGGCTGGTCACCAACC |
| Human | SART3 | Other Histone Modifications | CAAGGAGGAAGGTGTTATCG |
| Human | SART3 | Other Histone Modifications | CGAGTGGGAATATGACGAAG |
| Human | SART3 | Other Histone Modifications | CATGGTCTTGTATGTCGCAG |
| Human | SART3 | Other Histone Modifications | CGCAGACATACGGAGCGAGG |
| Human | SART3 | Other Histone Modifications | GCCGCAGACATACGGAGCGA |
| Human | SART3 | Other Histone Modifications | GAGTCGGTACCTCTTGGCCA |
| Human | SART3 | Other Histone Modifications | GGCCGCCTTTACTCGTGCCT |
| Human | SART3 | Other Histone Modifications | CTCCAAGGCACGAGTAAAGG |

| | | | |
|---|---|---|---|
| Human | SART3 | Other Histone Modifications | ACTGCCGGAAGGCTCTGCAC |
| Human | SART3 | Other Histone Modifications | CTGCCGGAAGGCTCTGCACC |
| Human | SART3 | Other Histone Modifications | CGGCCCGGTGCAGAGCCTTC |
| Human | BRD2 | Chromatin Readers | CTTGTTGTAAATGTAACAGT |
| Human | BRD2 | Chromatin Readers | TGGACATGGGTACTATTAAG |
| Human | BRD2 | Chromatin Readers | TTAATAGTACCCATGTCCAT |
| Human | BRD2 | Chromatin Readers | TGGACCTCAGCACTGTCAAG |
| Human | BRD2 | Chromatin Readers | TCTTCCGCTTGACAGTGCTG |
| Human | BRD2 | Chromatin Readers | CCGTGATTACCGGGATGCAC |
| Human | BRD2 | Chromatin Readers | CCTGTGCATCCCGGTAATCA |
| Human | BRD2 | Chromatin Readers | CTTGGATTATATGCACAACT |
| Human | BRD2 | Chromatin Readers | GCGCTCAAGCTCTCTAAGTG |
| Human | BRD2 | Chromatin Readers | CCATGAGTTACGATGAGAAG |
| Human | BRD2 | Chromatin Readers | TAGCAGTGAGAGCTCCTCTG |
| Human | BRD2 | Chromatin Readers | ACTGCTACTTTCCTCACTGG |
| Human | BRD2 | Chromatin Readers | GAGCTCCTCTGAGGAAGAGG |
| Human | BRD2 | Chromatin Readers | TTCCTCCTCCTCTTCCTCAG |
| Human | BRD2 | Chromatin Readers | CTCACTGCTACTTTCCTCAC |
| Human | BRD2 | Chromatin Readers | GGTGTCTGAAGACGACGACG |
| Human | BRD2 | Chromatin Readers | TGAAGACGACGACGAGGAAG |
| Human | BRD2 | Chromatin Readers | GGAGGAGCTGGAATCTGAGC |
| Human | BRD2 | Chromatin Readers | AGACGACGACGAGGAAGAGG |
| Human | BRD2 | Chromatin Readers | CGACGAGGAAGAGGAGGAGC |
| Human | BRD2 | Chromatin Readers | TGTCCTAATGGCACAAACGC |
| Human | BRD2 | Chromatin Readers | TTTCCAGCGTTTGTGCCATT |
| Human | BRD2 | Chromatin Readers | GGCAATGGCACGAAAGCTAC |
| Human | SCML2 | Chromatin Readers | CAGGTGGAATCTGAGACTGA |
| Human | SCML2 | Chromatin Readers | CCCAACAGGTTGTATGTCTG |
| Human | SCML2 | Chromatin Readers | TGGTTCCTTCTTAAATAATG |
| Human | SCML2 | Chromatin Readers | CTCATCTGTCCTGCGACCAT |
| Human | SCML2 | Chromatin Readers | GTCGCAGGACAGATGAGATA |
| Human | SCML2 | Chromatin Readers | GCAACATGAAATTGATGGGA |
| Human | SCML2 | Chromatin Readers | TAGTAACACAGCTTTAATGC |
| Human | SCML2 | Chromatin Readers | TCAGGCAACATGAAATTGAT |
| Human | SCML2 | Chromatin Readers | GGTGCTTCGCCGGATTGTGC |
| Human | ATAD2 | Chromatin Readers | CATAATCAGGAACCTCATCA |
| Human | ATAD2 | Chromatin Readers | CAACAGGCTTAGTAAACACT |
| Human | ATAD2 | Chromatin Readers | CACTGTAATAAAGCAACCAA |
| Human | ATAD2 | Chromatin Readers | ACATAATCAGGAACCTCATC |
| Human | ATAD2 | Chromatin Readers | AAGATACATTTAGAGAACTG |
| Human | ATAD2 | Chromatin Readers | GCAGCTCTGTGAAGAAATTC |
| Human | ATAD2 | Chromatin Readers | ATCAAGTTCTTCTTTAATTA |

| | | | |
|---|---|---|---|
| Human | ATAD2 | Chromatin Readers | TCTTAAGAAATGTTACACAT |
| Human | ATAD2 | Chromatin Readers | GGCAGTATCTCTTAAAGCAC |
| Human | PARP2 | Other Histone Modifications | GTACGGAGTCCAAAGTCATG |
| Human | PARP2 | Other Histone Modifications | GCCACTGTCAGCTTCCCAAG |
| Human | PARP2 | Other Histone Modifications | CACTGTCAGCTTCCCAAGTG |
| Human | PARP2 | Other Histone Modifications | TCCACTAATCCGGACACAGA |
| Human | PARP2 | Other Histone Modifications | CTCTTATCAGAGGTAGCTCT |
| Human | PARP2 | Other Histone Modifications | CCCATTCAGGGTGACGAAGT |
| Human | PARP2 | Other Histone Modifications | CCAAACATGTAACCTGTGAT |
| Human | PARP2 | Other Histone Modifications | AAGGGTCTCTGAATCTGTGA |
| Human | PARP2 | Other Histone Modifications | AAGAGTTAATAATGGCAACA |
| Human | PARP2 | Other Histone Modifications | AGCAAGATGGTATGCCAGGA |
| Human | ATAT1 | Histone Acetylation | TTTGCTGCTGTAGATCAACA |
| Human | ATAT1 | Histone Acetylation | TTGCTGCTGTAGATCAACAC |
| Human | ATAT1 | Histone Acetylation | CTTCCAGTATATGTTGCAGA |
| Human | ATAT1 | Histone Acetylation | TATGTTGCAGAAGGAGCGAG |
| Human | CTDP1 | Transcription Elongation | TGAGATTCTCGGGACCCAGG |
| Human | CTDP1 | Transcription Elongation | CTGTACGCACACACCATCGC |
| Human | CTDP1 | Transcription Elongation | CGGTTGTGTGAATCAACGTC |
| Human | CTDP1 | Transcription Elongation | ACTTCCCGATAGAGAAGACG |
| Human | CTDP1 | Transcription Elongation | CTTCTCTGTGCCAGCTCGCG |
| Human | CTDP1 | Transcription Elongation | GGCTCAGCACCAGCCGAGTG |
| Human | CTDP1 | Transcription Elongation | GGAGGAGGACTTCGTGCCCT |
| Human | CTDP1 | Transcription Elongation | GAGAGCAGCAGTGAGTCCGA |
| Human | CTDP1 | Transcription Elongation | CGAGAGCAGCAGTGAGTCCG |
| Human | CTDP1 | Transcription Elongation | TCCTCCTCCTCCGCCTCTGA |
| Human | CTDP1 | Transcription Elongation | TGACCACCTCATCTACCTGG |
| Human | CTDP1 | Transcription Elongation | TCTCCTCCAGGTAGATGAGG |
| Human | CTDP1 | Transcription Elongation | CAGCGCCTTGGCCATCTCGT |
| Human | CTDP1 | Transcription Elongation | CGACGAGATGGCCAAGGCGC |
| Human | HDGFRP3 | Chromatin Readers | GGGAGTTCATCAATCCGGGC |
| Human | HDGFRP3 | Chromatin Readers | CTCTGGGAGTTCATCAATCC |
| Human | HDGFRP3 | Chromatin Readers | CGGATTGATGAACTCCCAGA |
| Human | HDGFRP3 | Chromatin Readers | CAATCCGGGCCGGCCAGTGC |
| Human | HDGFRP3 | Chromatin Readers | ATAGGATACTTGTTTGCTGG |
| Human | GTF2F1 | Transcription Elongation | GCGCAGGAAAGACAGCAGCG |
| Human | GTF2F1 | Transcription Elongation | TGAGGAGGAGAAGCCGCCTG |
| Human | GTF2F1 | Transcription Elongation | TGAGCAGAGCGACAGTAGTG |
| Human | GTF2F1 | Transcription Elongation | GAAAGACAGCAGCGAGGAGT |
| Human | GTF2F1 | Transcription Elongation | AAATTGTACCAGTTGTGCAC |
| Human | GTF2F1 | Transcription Elongation | AATTGTACCAGTTGTGCACG |
| Human | ZBTB4 | DNA and RNA Modifications | TTCAAGTCCCTTCTTCAGAG |

| | | | |
|---|---|---|---|
| Human | ZBTB4 | DNA and RNA Modifications | TTGAAGCAGCCAGGACGCTG |
| Human | ZBTB4 | DNA and RNA Modifications | TCAACTTCATCTACAGCGCC |
| Human | ZBTB4 | DNA and RNA Modifications | ACTTGGTGTCTCCGGCTATG |
| Human | ZBTB4 | DNA and RNA Modifications | CAAACGGCTGCAGACCCATG |
| Human | ZBTB4 | DNA and RNA Modifications | AAAGCTTCATCCATCCCAAA |
| Human | ZBTB4 | DNA and RNA Modifications | GGGTCTGCAGCCGTTTGGGA |
| Human | ZBTB4 | DNA and RNA Modifications | CAGACTGGACAGGGTCACGT |
| Human | ZBTB4 | DNA and RNA Modifications | TGTGTCTCTTCAGACTGGAC |
| Human | ZBTB4 | DNA and RNA Modifications | GTGTCTCTTCAGACTGGACA |
| Human | ZBTB4 | DNA and RNA Modifications | ACTTTCTCACAATAGCGGCA |
| Human | ZBTB4 | DNA and RNA Modifications | CACTTTCTCACAATAGCGGC |
| Human | ZBTB4 | DNA and RNA Modifications | TGAGAAAGTGTTTGCTCTGG |
| Human | ZBTB4 | DNA and RNA Modifications | TTGTGAGAAAGTGTTTGCTC |
| Human | ZBTB4 | DNA and RNA Modifications | CAGGTTATAGTAAGTGACAA |
| Human | ZBTB4 | DNA and RNA Modifications | TCACCACCCTGAGAAAGCTG |
| Human | ZBTB4 | DNA and RNA Modifications | CTTCCGCAGCTTTCTCAGGG |
| Human | ZBTB4 | DNA and RNA Modifications | GGTGCTTCCGCAGCTTTCTC |
| Human | ZBTB4 | DNA and RNA Modifications | GTGCTTCCGCAGCTTTCTCA |
| Human | ZBTB4 | DNA and RNA Modifications | CAGCTTTCTCAGGGTGGTGA |
| Human | ZBTB4 | DNA and RNA Modifications | TTGCACACCTTGGCGCAGTG |
| Human | ZBTB4 | DNA and RNA Modifications | TCTTGCACACCTTGGCGCAG |
| Human | ZBTB4 | DNA and RNA Modifications | AGCTGCGGTCTTGCACACCT |
| Human | ZBTB4 | DNA and RNA Modifications | TGCACACCTTGGCGCAGTGG |
| Human | ZBTB4 | DNA and RNA Modifications | GTGGCGGCTCAGGGCAGCTG |
| Human | ZBTB4 | DNA and RNA Modifications | ATGAGGCCCAGTGCCGACGA |
| Human | ZBTB4 | DNA and RNA Modifications | TGAGGCCCAGTGCCGACGAG |
| Human | ZBTB4 | DNA and RNA Modifications | CCGCGCACACATACAGCACG |
| Human | ZBTB4 | DNA and RNA Modifications | GGAGGAATCAAAGGCTGGTG |
| Human | ZBTB4 | DNA and RNA Modifications | AGGAGGAATCAAAGGCTGGT |
| Human | ZBTB4 | DNA and RNA Modifications | GGAGATGGAGGAGAGTGAGG |
| Human | ZBTB4 | DNA and RNA Modifications | GATGAGGAGGAATCAAAGGC |
| Human | ZBTB4 | DNA and RNA Modifications | AGCAGTGTAAGTGATAACCG |
| Human | ZBTB4 | DNA and RNA Modifications | GTGGGACGGCCAGCACAGGA |
| Human | ZBTB4 | DNA and RNA Modifications | GCAGGTCAGTCTCTGAGATG |
| Human | ZBTB4 | DNA and RNA Modifications | AGAGACTGACCTGCGTCCTG |
| Human | ZBTB4 | DNA and RNA Modifications | GCTGAGCGGAGAGGAGATGG |
| Human | SUPT16H | Transcription Elongation | AGCACATCAGAAAGAACTAG |
| Human | SUPT16H | Transcription Elongation | AACTCAATGAAGAAGCAAAG |
| Human | SUPT16H | Transcription Elongation | CCCGAGAACCTCTTCCCAAA |
| Human | SUPT16H | Transcription Elongation | GAGGCGATTGACTGAACAAA |
| Human | SUPT16H | Transcription Elongation | GGACCGAGAAAGTCGTTACG |
| Human | SUPT16H | Transcription Elongation | AGAAGAGTCAGACTATTCTA |

| | | | |
|---|---|---|---|
| Human | SUPT16H | Transcription Elongation | AAAGGATTGGGATGAACTGG |
| Human | SUPT16H | Transcription Elongation | TGAAGAGACTGTACAGCAAT |
| Human | SUPT16H | Transcription Elongation | TGCTGGCTGCTTTCTTCATT |
| Human | SUPT16H | Transcription Elongation | GTATAACGCTGTCATGGACG |
| Human | SUPT16H | Transcription Elongation | TGACGTGTATAACGCTGTCA |
| Human | JMJD7 | Histone Methylation | CCGGCGTGTACAGCTCATAG |
| Human | JMJD7 | Histone Methylation | AGCATCGGGCATCTTTCCCA |
| Human | JMJD7 | Histone Methylation | TGCAGTGACTTCTTTGCACA |
| Human | JMJD7 | Histone Methylation | TTCTCGTAACTCGCTCCGCA |
| Human | JMJD7 | Histone Methylation | AGAATTCCCGGCCGCTGCAA |
| Human | JMJD7 | Histone Methylation | AGGCGGCTTTGGAAGCCGTG |
| Human | JMJD7 | Histone Methylation | GAGCGAGTTACGAGAATTCC |
| Human | JMJD7 | Histone Methylation | GAGAATTCCCGGCCGCTGCA |
| Human | KMT2D | Histone Methylation | GAGGCTCCAATAGATGCGCG |
| Human | KMT2D | Histone Methylation | AGCAGCGACGATTGTTGGTG |
| Human | KMT2D | Histone Methylation | CCTCTATCCCGTGGGCTACG |
| Human | KMT2D | Histone Methylation | AGCGACGATTGTTGGTGCGG |
| Human | KMT2D | Histone Methylation | GCCACTGCCCTCTATCCCGT |
| Human | KMT2D | Histone Methylation | CATAGCTGAATCACTGCCCG |
| Human | KMT2D | Histone Methylation | ATGCGATTCCACACGGCCTG |
| Human | KMT2D | Histone Methylation | GTAATCAAAGTCATCGAGCA |
| Human | KMT2D | Histone Methylation | TGATTCAGCTATGCGAAGCA |
| Human | KMT2D | Histone Methylation | ATCTACGAAGAGCAGAATCG |
| Human | KMT2D | Histone Methylation | AATGGTTAATGTACCTGGCA |
| Human | KMT2D | Histone Methylation | GACTTCGGCCACACAGTTAG |
| Human | KMT2D | Histone Methylation | TGGCACCATCATTCGGAACG |
| Human | KMT2D | Histone Methylation | GAGCCTGGAATTGTCGGAAA |
| Human | KMT2D | Histone Methylation | TTCCAGGCTCCACAGTGGCA |
| Human | KMT2D | Histone Methylation | TCCAGGCTCCACAGTGGCAG |
| Human | KMT2D | Histone Methylation | GACAATTCCAGGCTCCACAG |
| Human | KMT2D | Histone Methylation | ACTGTGGAGCCTGGAATTGT |
| Human | KMT2D | Histone Methylation | CCCACCTAGGAGAACCTGGA |
| Human | KMT2D | Histone Methylation | ATCTGTGATAGGTCCTCACT |
| Human | KMT2D | Histone Methylation | CAGGGAAACCAATCTGTGAT |
| Human | KMT2D | Histone Methylation | AGGACCCTCCAGGTTCTCCT |
| Human | KMT2D | Histone Methylation | ATCACAGATTGGTTTCCCTG |
| Human | KMT2D | Histone Methylation | AGGTGAGCGGCAAGGGATGG |
| Human | KMT2D | Histone Methylation | AACCGCTGGCAGTCGCGCAG |
| Human | KMT2D | Histone Methylation | GGCACCGAGCCTGGTGCAGT |
| Human | KMT2D | Histone Methylation | CATGGATAGGAAGGAACCGC |
| Human | KMT2D | Histone Methylation | CTGTGGGCATCACTATCACG |
| Human | KMT2D | Histone Methylation | CTGCCCGCAAACGTGCTGGC |

| | | | |
|---|---|---|---|
| Human | KMT2D | Histone Methylation | GCTGTGGGCATCACTATCAC |
| Human | KMT2D | Histone Methylation | GTGATAGTGATGCCCACAGC |
| Human | KMT2D | Histone Methylation | GTTTGTGAGACGTGTGACAA |
| Human | KMT2D | Histone Methylation | GCCAAGCCTGCAGGAAACCT |
| Human | KMT2D | Histone Methylation | GCAGGCAGTTCCTCCATGGG |
| Human | KMT2D | Histone Methylation | TGTCAACAGCAAGATCACCA |
| Human | KMT2D | Histone Methylation | AGCACTGCGAACAGGCAAGG |
| Human | KMT2D | Histone Methylation | TCTTGCTGTTGACACAGTAA |
| Human | KMT2D | Histone Methylation | ACCAAGGTGATGCTGCTCAA |
| Human | KMT2D | Histone Methylation | GAGAGCACTGCGAACAGGCA |
| Human | KMT2D | Histone Methylation | CACAGAGCAGCAGGCGTGAG |
| Human | KMT2D | Histone Methylation | AGGGCGGCTGGAAGTGCAAG |
| Human | KMT2D | Histone Methylation | AGTGCCGCCACTGTGAACGG |
| Human | KMT2D | Histone Methylation | CATGCATCCACCGTTCACAG |
| Human | KMT2D | Histone Methylation | TGTGAACGGTGGATGCATGC |
| Human | KMT2D | Histone Methylation | GCATCCACCGTTCACAGTGG |
| Human | KMT2D | Histone Methylation | AGTAGGTCCTCTTCTACGTA |
| Human | KMT2D | Histone Methylation | TGCTGTTTCTGTCATGAGGA |
| Human | KMT2D | Histone Methylation | CACAGTTGAGGTGCACCCAC |
| Human | KMT2D | Histone Methylation | CTGCTGTTTCTGTCATGAGG |
| Human | KMT2D | Histone Methylation | TGGACCAAAGGGCACAGTTG |
| Human | KMT2D | Histone Methylation | CAGGTCCAGGTTCAGCAGAC |
| Human | KMT2D | Histone Methylation | GGCACCAGTTCGCTGGCACA |
| Human | KMT2D | Histone Methylation | TGAAGAACATGCACTTGGCA |
| Human | KMT2D | Histone Methylation | GAACATGCACTTGGCACGGA |
| Human | KMT2D | Histone Methylation | AGGAGATTCTTCAAATGGTG |
| Human | KMT2D | Histone Methylation | ATAGGCGCGATACCTCAGGT |
| Human | KMT2D | Histone Methylation | AGAGGAGACTCTTCAAATGG |
| Human | KMT2D | Histone Methylation | AGATCCAGCGCAACACCCTG |
| Human | KMT2D | Histone Methylation | CAGCTCTGTTTGGCTAAGGC |
| Human | KMT2D | Histone Methylation | TCTCCAGCTCTGTTTGGCTA |
| Human | KMT2D | Histone Methylation | TCAGCAGCTCTCGTAGTCGC |
| Human | KMT2D | Histone Methylation | CGGCCTTAGCCAAACAGAGC |
| Human | KMT2D | Histone Methylation | CCTGTTGGAGCATGAGAAGA |
| Human | KMT2D | Histone Methylation | CCTTCTTCTCATGCTCCAAC |
| Human | KMT2D | Histone Methylation | TGTTGCTGCTGCTGCTGGGC |
| Human | KMT2D | Histone Methylation | AGAAACAACTGGATCAGGTC |
| Human | KMT2D | Histone Methylation | GAGCAAGATCCAGAAACAAC |
| Human | KMT2D | Histone Methylation | CTAATCTCATGGCAGAATAT |
| Human | KMT2D | Histone Methylation | GAAGGAGCACACTAATCTCA |
| Human | KMT2D | Histone Methylation | TCAGGTCCGGAAACAGCAGA |
| Human | KMT2D | Histone Methylation | GCAGCAGCAACGTATGCAGC |

| | | | |
|---|---|---|---|
| Human | KMT2D | Histone Methylation | CTGCTGCTGCTGCAGTTTCT |
| Human | KMT2D | Histone Methylation | GCTGCTGCTGCTGCAGTTTC |
| Human | KMT2D | Histone Methylation | GCTGCAGCTGCCGCTGCATG |
| Human | KMT2D | Histone Methylation | TTGGAAGTGCAAGGCGTGCC |
| Human | KMT2D | Histone Methylation | CTTGGCACACTTTGCATTCA |
| Human | KMT2D | Histone Methylation | GCAAAGTGTGCCAAGCCTGC |
| Human | KMT2D | Histone Methylation | GCTTGGCACACTTTGCATTC |
| Human | KMT2D | Histone Methylation | GCAAGGCGTGCCGGGTGTGC |
| Human | KMT2D | Histone Methylation | CAAACCACTAGGGAAAGCAG |
| Human | KMT2D | Histone Methylation | AATGTGAAGCCAAACCACTA |
| Human | KMT2D | Histone Methylation | TGAAGGAGTGGCGAACACTG |
| Human | KMT2D | Histone Methylation | GCCCGTTCCCGCATCAAACA |
| Human | KMT2D | Histone Methylation | GCTGCGACCCTGTTTGATGC |
| Human | KMT2D | Histone Methylation | GGAAGACGCCGGCCTCGTGG |
| Human | KMT2D | Histone Methylation | TGCTGCGACCCTGTTTGATG |
| Human | KMT2D | Histone Methylation | GCAGTTTCCCAGGAAGACGC |
| Human | KMT2D | Histone Methylation | AACTATTCTGCCATTCACAG |
| Human | KMT2D | Histone Methylation | AGAATAGTTACACACACTGT |
| Human | KMT2D | Histone Methylation | CAGGTCACCAGGCTGGCACA |
| Human | KMT2D | Histone Methylation | TGTGTCCTGTATGCAGTGTG |
| Human | KMT2D | Histone Methylation | GGTGGTGGCAGAACCGACGG |
| Human | KMT2D | Histone Methylation | GTGGTGGCAGAACCGACGGA |
| Human | KMT2D | Histone Methylation | AAATGGCTGTTGATCCCATG |
| Human | KMT2D | Histone Methylation | CGCCGGCTTGTCTACCTCTG |
| Human | KMT2D | Histone Methylation | GCCAAGCTCAGTGCTCGACG |
| Human | KMT2D | Histone Methylation | AAGACTGTTGTTCAATTCAG |
| Human | KMT2D | Histone Methylation | TGATCAATGAAGCTGACCAG |
| Human | KMT2D | Histone Methylation | AGCTGCTGTAAAGAGCCCAT |
| Human | KMT2D | Histone Methylation | TATCATTGATCCCATTGCAA |
| Human | KMT2D | Histone Methylation | CCTGTAGATAAGGCTCCTGG |
| Human | KMT2D | Histone Methylation | CTGTAGATAAGGCTCCTGGT |
| Human | KMT2D | Histone Methylation | GTGGAGGTGTGGGATGGACA |
| Human | KMT2D | Histone Methylation | TTATCTACAGGACCAGTCCT |
| Human | KMT2D | Histone Methylation | TTCGGTGGGAAGTTCAACCA |
| Human | KMT2D | Histone Methylation | TCGGTGGGAAGTTCAACCAA |
| Human | KMT2D | Histone Methylation | TGGCAGGAGAAGGTGCCAAG |
| Human | KMT2D | Histone Methylation | GGCTCAGCCAAGGGTTCGGT |
| Human | KMT2D | Histone Methylation | TGAACTTCCCACCGAACCCT |
| Human | KMT2D | Histone Methylation | ACCGGCGGCTGCGCACCGAA |
| Human | KMT2D | Histone Methylation | TCCATTCGGTGCGCAGCCGC |
| Human | PRDM4 | Histone Methylation | ATCTGGAAGATATACCACAA |
| Human | PRDM4 | Histone Methylation | TGTGCGCAAAGCCAGGAACC |

| | | | |
|---|---|---|---|
| Human | PRDM4 | Histone Methylation | AGAATTCTAGGACACCATTG |
| Human | PRDM4 | Histone Methylation | CTGCTCCCACAATTGACTGA |
| Human | PRDM4 | Histone Methylation | TGGGAGCAGAAGTTGGTGTA |
| Human | PRDM4 | Histone Methylation | AGGTGGGTCCGCAGGTTGCT |
| Human | PRDM4 | Histone Methylation | TTAGTGATCCCAGCAACCTG |
| Human | PRDM4 | Histone Methylation | GAGGTGGGTCCGCAGGTTGC |
| Human | PRDM4 | Histone Methylation | GGTGAAAGACTTGTCACACA |
| Human | PRDM4 | Histone Methylation | TGACAAGTCTTTCACCCAGA |
| Human | PRDM4 | Histone Methylation | TTTCACCCAGAAGGCTCACC |
| Human | PRDM4 | Histone Methylation | GGCTCACCTGGAGTCCCACA |
| Human | PRDM4 | Histone Methylation | GGGACTCCAGGTGAGCCTTC |
| Human | PRDM4 | Histone Methylation | GTGACAAGTTGTTTATGCGG |
| Human | PRDM4 | Histone Methylation | ACTGTGACAAGTTGTTTATG |
| Human | PRDM4 | Histone Methylation | CAAGTTGTTTATGCGGAGGC |
| Human | PRDM4 | Histone Methylation | CAAGAACAGCTTATCACACT |
| Human | PRDM4 | Histone Methylation | AAGAACAGCTTATCACACTT |
| Human | PRDM4 | Histone Methylation | GTGGCGGGTGAGATGGTATT |
| Human | STK31 | Chromatin Readers | GACCCAAACAAGATTTATGG |
| Human | STK31 | Chromatin Readers | ATCCACCATAAATCTTGTTT |
| Human | STK31 | Chromatin Readers | CTGGTGAGGTACATTGACTA |
| Human | STK31 | Chromatin Readers | CTCAAATACATGAACTCTGG |
| Human | STK31 | Chromatin Readers | TCTGTTAAAGGGCTATTCTG |
| Human | STK31 | Chromatin Readers | CCAGGAGCTGCAGCAAGAGA |
| Human | STK31 | Chromatin Readers | GAAAGTAGAGCAGATTGCCC |
| Human | STK31 | Chromatin Readers | CTTCTCTTGCTGCAGCTCCT |
| Human | STK31 | Chromatin Readers | GCAAGAGAAGGCAGCTGCTG |
| Human | ESCO1 | DNA and RNA Modifications | AGATGCTGTGTTTCATCTTC |
| Human | ESCO1 | DNA and RNA Modifications | CTCGCATGATTGAATGCCTA |
| Human | CBX4 | Chromatin Readers | GTCGCCCAAATATAACACGT |
| Human | CBX4 | Chromatin Readers | GGTCGCCCAAATATAACACG |
| Human | CBX4 | Chromatin Readers | GAGAAGAAGCGGATCCGCAA |
| Human | CBX4 | Chromatin Readers | GCTGATCGCCTTCCAGAACA |
| Human | CBX4 | Chromatin Readers | ATATAACACGTGGGAACCGG |
| Human | CBX4 | Chromatin Readers | GTAGGCACCTGCACCACTAG |
| Human | CBX4 | Chromatin Readers | CTGCGTCAAGACGCGCAGCG |
| Human | CBX4 | Chromatin Readers | ATAATTATATTCCCAAAGAA |
| Human | CBX4 | Chromatin Readers | AGTAACGGTGAGGCAGTTCG |
| Human | CBX4 | Chromatin Readers | GAGGCAGTTCGCGGTGACGT |
| Human | CBX4 | Chromatin Readers | GATAATTATATTCCCAAAGA |
| Human | CBX4 | Chromatin Readers | CTGCCTCACCGTTACTTTCA |
| Human | SETMAR | Histone Methylation | CATCATAGTTCTCTCCATGG |
| Human | SETMAR | Histone Methylation | CTTAGAGATATAGGATCTGG |

| | | | |
|---|---|---|---|
| Human | SETMAR | Histone Methylation | GGAGACAGGAGCAAGTGCCA |
| Human | SETMAR | Histone Methylation | TGACCACTGCAGAAACAGAG |
| Human | SETMAR | Histone Methylation | TCCTGTCCGAATTGACTCAA |
| Human | SETMAR | Histone Methylation | TGGAATTTATACCGAAAGGA |
| Human | SETMAR | Histone Methylation | TCCTTTCGGTATAAATTCCA |
| Human | SETMAR | Histone Methylation | GATTTGGCACCACAGTAACA |
| Human | SETMAR | Histone Methylation | CTAACGAACGTACAGTGCAG |
| Human | SETMAR | Histone Methylation | ACGAACGTACAGTGCAGTGG |
| Human | SETMAR | Histone Methylation | GTACGTTCGTTAGCAGTTCC |
| Human | SETMAR | Histone Methylation | ATCAACAATGCATTTGGCCC |
| Human | SETMAR | Histone Methylation | CGCAACATCAACAATGCATT |
| Human | SETMAR | Histone Methylation | ACTCAATGTCAACCATTCTA |
| Human | SETMAR | Histone Methylation | TGATCAGGAGTGTACTGGAA |
| Human | SETMAR | Histone Methylation | ATGATCAGGAGTGTACTGGA |
| Human | SETMAR | Histone Methylation | GGAGTGTACTGGAAGGGCGC |
| Human | SETMAR | Histone Methylation | ACCGGCGACGATCAGCTCAG |
| Human | SETMAR | Histone Methylation | AGTGTGGGTTGTGCAACATG |
| Human | SETMAR | Histone Methylation | TGTTGAGATGCTTAAAGACG |
| Human | SETMAR | Histone Methylation | CATGCGGTCGGGCATTGTCG |
| Human | DOLK | Control | GGCCGGAGATGGGCACTCTC |
| Human | DOLK | Control | GGGCCGGAGATGGGCACTCT |
| Human | DOLK | Control | AGAGTGCCCATCTCCGGCCC |
| Human | DOLK | Control | GCCAGCACCGATCCACTCAG |
| Human | DOLK | Control | GGCGGCAGTAGTGTTTGCAG |
| Human | DOLK | Control | TGGATCGGTGCTGGCAGAGG |
| Human | DOLK | Control | GAGTGGATCGGTGCTGGCAG |
| Human | DOLK | Control | CCACGGCGCACCACGAGTAT |
| Human | DOLK | Control | CCTTCTACGTCCAATACAAG |
| Human | DOLK | Control | AAAGGCATGACCATGGAGGC |
| Human | DOLK | Control | TCCAAGCAAAGGCATGACCA |
| Human | DOLK | Control | TCCATGGTCATGCCTTTGCT |
| Human | DOLK | Control | AAGCAAAGGCATGACCATGG |
| Human | DOLK | Control | TGACCATGGAGGCGGGCAAT |
| Human | DOLK | Control | AGAACGGGTTCCCAGCAGTC |
| Human | DOLK | Control | GCGCCAACACTGATGAGAAG |
| Human | DOLK | Control | CGCCAACACTGATGAGAAGA |
| Human | DOLK | Control | GGCCCTCTTCTCATCAGTGT |
| Human | DOLK | Control | GGCAGCCACTGGCATGGCAG |
| Human | DOLK | Control | GAGGGCCACTGCCATGCCAG |
| Human | DOLK | Control | AAGATGACACAAGTGTTGGT |
| Human | DOLK | Control | GCCCAAGATGACACAAGTGT |
| Human | DOLK | Control | GTCATCTTGGGCTTGGCTGG |

| | | | |
|---|---|---|---|
| Human | DOLK | Control | CACTTGTGTCATCTTGGGCT |
| Human | DOLK | Control | AACCAACACTTGTGTCATCT |
| Human | DOLK | Control | TGAGATAAACGAAGATCAGA |
| Human | DOLK | Control | CGAAGATCAGAAGGACTTCC |
| Human | DOLK | Control | GAGGCACTGCTGGTATTGGG |
| Human | DOLK | Control | TACCAGCAGTGCCTCACCAG |
| Human | DOLK | Control | ATACCAGCAGTGCCTCACCA |
| Human | DOLK | Control | AATACCAGCAGTGCCTCACC |
| Human | DOLK | Control | TGGTGAGGCACTGCTGGTAT |
| Human | DOLK | Control | GACACTGGTGGAAAGTCAGG |
| Human | DOLK | Control | TGACACTGGTGGAAAGTCAG |
| Human | DOLK | Control | CTGACACTGGTGGAAAGTCA |
| Human | DOLK | Control | TCTGACACTGGTGGAAAGTC |
| Human | DOLK | Control | GTAGTAGGGATGGTACTCAT |
| Human | DOLK | Control | GGTAGTAGGGATGGTACTCA |
| Human | DOLK | Control | TCCCTACTACCACCACCAGC |
| Human | DOLK | Control | GCTGGTGGTGGTAGTAGGGA |
| Human | DOLK | Control | TCCTGCTGGTGGTGGTAGTA |
| Human | DOLK | Control | CACACCAAGGCTCAGCACAC |
| Human | DOLK | Control | GGCTCAGCACACAGGTCATG |
| Human | DOLK | Control | TCAGCACACAGGTCATGAGG |
| Human | DOLK | Control | CTGTGTGCTGAGCCTTGGTG |
| Human | DOLK | Control | ATGACCTGTGTGCTGAGCCT |
| Human | DOLK | Control | GAAACTGAAGAAGCCAGAGC |
| Human | DOLK | Control | AAACTGAAGAAGCCAGAGCA |
| Human | DOLK | Control | AACTGAAGAAGCCAGAGCAG |
| Human | DOLK | Control | GCAGAGACCAATAGGCTAGG |
| Human | DOLK | Control | CCAGCAGAGACCAATAGGCT |
| Human | DOLK | Control | GGTGGCCAGCAGAGACCAAT |
| Human | DOLK | Control | CACCACCAGGCAGGCCAAGG |
| Human | DOLK | Control | GGCCACCTTGGCCTGCCTGG |
| Human | DOLK | Control | GGACTCGGAAGATGACCGCT |
| Human | DOLK | Control | TTCCGAGTCCAAGAAGCACC |
| Human | DOLK | Control | GGCCTGGTGCTTCTTGGACT |
| Human | DOLK | Control | GTGGTAGCCACCTACATCCC |
| Human | DOLK | Control | TGTAGGTGGCTACCACAATG |
| Human | DOLK | Control | AGGTGGCTACCACAATGAGG |
| Human | DOLK | Control | GAAGACCGCCAGGCATACAG |
| Human | DOLK | Control | GACCGCCAGGCATACAGTGG |
| Human | DOLK | Control | CCAGGAAGATGAAGACCGCC |
| Human | DOLK | Control | CCTGGCGGTCTTCATCTTCC |
| Human | DOLK | Control | CTCCGTAGAGTGTGACCCAA |

| | | | |
|---|---|---|---|
| Human | DOLK | Control | AGCCTTTGGGTCACACTCTA |
| Human | DOLK | Control | AGATGGGAAGAGACATGCCC |
| Human | DOLK | Control | ACACACATCTACCTGCTCCT |
| Human | DOLK | Control | TGGGCATGTCTCTTCCCATC |
| Human | DOLK | Control | TAGATGTGTGTCAGAATGAG |
| Human | DOLK | Control | GAAGAGACATGCCCAGGAGC |
| Human | DOLK | Control | TGCACACAGAAGGGTAGCCT |
| Human | DOLK | Control | ACACAGAAGGGTAGCCTGGG |
| Human | DOLK | Control | CTGCACACAGAAGGGTAGCC |
| Human | DOLK | Control | CAGGCTACCCTTCTGTGTGC |
| Human | DOLK | Control | AGGCTACCCTTCTGTGTGCA |
| Human | DOLK | Control | CACAGCCAGGACACCGGCAT |
| Human | DOLK | Control | ACAGCCAGGACACCGGCATA |
| Human | DOLK | Control | CAGCCAGGACACCGGCATAG |
| Human | DOLK | Control | CACACCCACAGCCAGGACAC |
| Human | DOLK | Control | TATCACCCACACCCACAGCC |
| Human | DOLK | Control | CTGCGCAAATATAGATGTCA |
| Human | DOLK | Control | ACCTAAACTACAGTTATGCT |
| Human | DOLK | Control | TCCAAGCATAACTGTAGTTT |
| Human | DOLK | Control | CTACAGTTATGCTTGGATTT |
| Human | DOLK | Control | AGCAATATCAGGAGGTAGAG |
| Human | DOLK | Control | TCAGGAGGTAGAGAGGCAGA |
| Human | SKA3 | Chromatin Remodeling | CGCTGGACTGCGAGACGGCC |
| Human | SKA3 | Chromatin Remodeling | CAGCACGCTGGACTGCGAGA |
| Human | SKA3 | Chromatin Remodeling | CGTCTCGCAGTCCAGCGTGC |
| Human | SKA3 | Chromatin Remodeling | CTGCGGGAAGCTGCGGTCTC |
| Human | SKA3 | Chromatin Remodeling | GGAGCTTCTGCGGGAAGCTG |
| Human | SKA3 | Chromatin Remodeling | GAGTCTGAACTTCTGAATGA |
| Human | SKA3 | Chromatin Remodeling | TTCAGAAGTTCAGACTCTAA |
| Human | SKA3 | Chromatin Remodeling | AGAAGGCATTGATTTCATAA |
| Human | PAK2 | Other Histone Modifications | ATTAGGTACAAGGCCCTCAA |
| Human | PAK2 | Other Histone Modifications | TATTAGGTACAAGGCCCTCA |
| Human | PAK2 | Other Histone Modifications | TCATGGAATACCTTGCTGGG |
| Human | PAK2 | Other Histone Modifications | TCACTGGCATGCCAGAACAG |
| Human | PAK2 | Other Histone Modifications | CACTGGCATGCCAGAACAGT |
| Human | PAK2 | Other Histone Modifications | GCTGTTACTGGAGAATTCAC |
| Human | PAK2 | Other Histone Modifications | CTCTAGTTTGGTGATATTGG |
| Human | PAK2 | Other Histone Modifications | TAACAGCATCAAAGCCAACA |
| Human | ZGPAT | Chromatin Readers | CAAGGGTTTGGGCCGACACG |
| Human | ZGPAT | Chromatin Readers | TGGTCCAGCGACTTCCCTCG |
| Human | ZGPAT | Chromatin Readers | ATGCTGTGGTGTTGCCTCGA |
| Human | ZGPAT | Chromatin Readers | CATGCTGTGGTGTTGCCTCG |

| | | | |
|---|---|---|---|
| Human | ZGPAT | Chromatin Readers | AGAACTGCAGGTTCTCCCAT |
| Human | ZGPAT | Chromatin Readers | GAGAACTGCAGGTTCTCCCA |
| Human | ZGPAT | Chromatin Readers | CCGTGCCCGTTCTTCCTGGA |
| Human | ZGPAT | Chromatin Readers | GCCGCTTTAAGGAGAACTGC |
| Human | ZGPAT | Chromatin Readers | GCCGTGCCCGTTCTTCCTGG |
| Human | ZGPAT | Chromatin Readers | CGCAGCACGCATCACCGATG |
| Human | SIRT7 | Histone Acetylation | TTTACATCGTGAACCTGCAG |
| Human | SIRT7 | Histone Acetylation | GCAGAAGCTGGTGCAGCATG |
| Human | SIRT7 | Histone Acetylation | TCAGGATGCGCGACACCTGG |
| Human | SIRT7 | Histone Acetylation | GACCTGGTAACGGAGCTGCA |
| Human | SIRT7 | Histone Acetylation | GGATGCGCGACACCTGGCGG |
| Human | SIRT7 | Histone Acetylation | GCGGGTCCGGAGGTTGCGGG |
| Human | SIRT7 | Histone Acetylation | CGAGAGCGCGGACCTGGTAA |
| Human | KAT6A | Histone Acetylation | TGATAGCCAATCGTAACTGC |
| Human | KAT6A | Histone Acetylation | GATGTGTCTGCATTATTCGG |
| Human | KAT6A | Histone Acetylation | TTGGCTATCAAACGTGCCAT |
| Human | KAT6A | Histone Acetylation | CATCTTTAAGGAGTCTGCCG |
| Human | KAT6A | Histone Acetylation | TTGGAATAAACTGATAAAGC |
| Human | KAT6A | Histone Acetylation | GGCAGCCTTGAGTATTCTTG |
| Human | KAT6A | Histone Acetylation | TAATCACAAATTGGTCACTA |
| Human | KAT6A | Histone Acetylation | ATGGCCACTGTTGCCACAGT |
| Human | KAT6A | Histone Acetylation | CACTCGAACCGTTAGTTCAG |
| Human | KAT6A | Histone Acetylation | TTCGAGTGAAGGCCTTACGG |
| Human | KAT6A | Histone Acetylation | TGTGCCGACTGTGGCAACAG |
| Human | KAT6A | Histone Acetylation | TGACATATCCACATGCCTTT |
| Human | KAT6A | Histone Acetylation | GAATAGCACTTCCTAAGCCT |
| Human | KAT6A | Histone Acetylation | CAAACCCGCTTTATACTGAG |
| Human | KAT6A | Histone Acetylation | TTGGAGTTGAGTGTTAAAGA |
| Human | KAT6A | Histone Acetylation | TGGCCCACAGACAATCAGGA |
| Human | KAT6A | Histone Acetylation | CAGCCATCCTGATTGTCTGT |
| Human | KAT6A | Histone Acetylation | CAACACTAAAGCAACCAACG |
| Human | KAT6A | Histone Acetylation | TATGTCAAATATGTCGACCT |
| Human | KAT6A | Histone Acetylation | AGTCATAGTGTCCAACTCTG |
| Human | KAT6A | Histone Acetylation | GTCCAACTCTGTGGTCTCAG |
| Human | KAT6A | Histone Acetylation | ACAGAGTTGGACACTATGAC |
| Human | KAT6A | Histone Acetylation | CAACTCTGTGGTCTCAGAGG |
| Human | KAT6A | Histone Acetylation | GGAGTCCAGCGCAAACATTC |
| Human | KAT6A | Histone Acetylation | AGTTACGACTCCACGATGGG |
| Human | KAT6A | Histone Acetylation | GGAACATAAGTATCTATGAG |
| Human | KAT6A | Histone Acetylation | TTCATACTACACTGTGACAG |
| Human | KAT6A | Histone Acetylation | TGGTAACTGCTGTTCATCAT |
| Human | KAT6A | Histone Acetylation | TCATGTAGGCAGGGTTACTA |

| | | | |
|---|---|---|---|
| Human | KAT6A | Histone Acetylation | TGTCATTCGATAGCTGTTCA |
| Human | KAT6A | Histone Acetylation | ATGGTAACTGCTGTTCATCA |
| Human | KAT6A | Histone Acetylation | ATAGCTGTTCATGGCATTCA |
| Human | KAT6A | Histone Acetylation | GGAGGAGGAGATGCAAGCAG |
| Human | KAT6A | Histone Acetylation | GAGGCAGAAGAGGCTGAAGA |
| Human | KAT6A | Histone Acetylation | AGAGGCAGAAGAGGCTGAAG |
| Human | KAT6A | Histone Acetylation | GATGCAAGCAGAGGCAGAAG |
| Human | KAT6A | Histone Acetylation | CTCTGCTCCTCTGAGACACG |
| Human | KAT6A | Histone Acetylation | AGAGGAGCAGAGGCAGTCAG |
| Human | KAT6A | Histone Acetylation | GGAGCAGAGGCAGTCAGAGG |
| Human | KAT6A | Histone Acetylation | ATTAGAGGAGCCAGAGCCAG |
| Human | KAT6A | Histone Acetylation | CATGGAATTGACATTATAGG |
| Human | KAT6A | Histone Acetylation | TTGACATTATAGGCGGGAGT |
| Human | KAT6A | Histone Acetylation | CCTATGCAGCCTAACCCTCA |
| Human | KAT6A | Histone Acetylation | ATGGAATTGACATTATAGGC |
| Human | KAT6A | Histone Acetylation | CTGTGCTGTCTGGTTCATGT |
| Human | CREBBP | Histone Acetylation | AGCTCTAAAGGATCGCCGCA |
| Human | CREBBP | Histone Acetylation | ATAATATTCATCCCTGCTGT |
| Human | CREBBP | Histone Acetylation | CTGGCACGAACATGTCACTC |
| Human | CREBBP | Histone Acetylation | TATGCACTAGATGGCTCCGC |
| Human | CREBBP | Histone Acetylation | CCAAGAGCCCTGGCAGTACG |
| Human | CREBBP | Histone Acetylation | ACGTCGTCCACGTACTGCCA |
| Human | CREBBP | Histone Acetylation | TGTCTTGCGATTATAGAGCC |
| Human | CREBBP | Histone Acetylation | TGACATCGTAAAGAATCCCA |
| Human | CREBBP | Histone Acetylation | AGAAATTAGGGTATGTGACA |
| Human | CREBBP | Histone Acetylation | GCAGCCAGTGAAACCACTGA |
| Human | CREBBP | Histone Acetylation | GAAACCACTGAGGGCAGTCA |
| Human | CREBBP | Histone Acetylation | ACACACGCCTCGTGTTTGGA |
| Human | CREBBP | Histone Acetylation | AAGATGCACAATGGGCAACT |
| Human | CREBBP | Histone Acetylation | TTTGTCGTGAAGATGCACAA |
| Human | CREBBP | Histone Acetylation | TGCACAATGGGCAACTTGGC |
| Human | CREBBP | Histone Acetylation | AGTGCCAGCGCGTCTCCACG |
| Human | CREBBP | Histone Acetylation | CAACGAGTGCAAGCACCACG |
| Human | CREBBP | Histone Acetylation | CTGGCACTGCACTGTGTGCG |
| Human | CREBBP | Histone Acetylation | GTGGTGCTTGCACTCGTTGC |
| Human | CREBBP | Histone Acetylation | TGTTATAGCAGTTGATGCAG |
| Human | CREBBP | Histone Acetylation | TGCAAACGCAAGACCAACGG |
| Human | CREBBP | Histone Acetylation | CTGCAAACGCAAGACCAACG |
| Human | CREBBP | Histone Acetylation | CGGGTGGTGCAGCACACCAA |
| Human | CREBBP | Histone Acetylation | GCAGCGAGCAGTTGGCGTTG |
| Human | CREBBP | Histone Acetylation | TTTCTTGGCAGTGCTTGGCG |
| Human | CREBBP | Histone Acetylation | CCGCAAATGACTGGTCACGC |

| | | | |
|---|---|---|---|
| Human | CREBBP | Histone Acetylation | CTGAGACATATTTGGCACGT |
| Human | CREBBP | Histone Acetylation | TCTGTTTGCTGGCTAACTGG |
| Human | CREBBP | Histone Acetylation | CGCGTGACCAGTCATTTGCG |
| Human | CREBBP | Histone Acetylation | GTCTCAGATGCAAACATCAG |
| Human | CREBBP | Histone Acetylation | TCAGCCGGCGTGACTCCTGG |
| Human | CREBBP | Histone Acetylation | AGGCCCAGCTCATGCGCCGG |
| Human | CREBBP | Histone Acetylation | GCCCATAAGATGGTGAAGTG |
| Human | CREBBP | Histone Acetylation | CTGGTGGAGCTGCACACCCA |
| Human | CREBBP | Histone Acetylation | CACTCCTTGCAATCAACGAA |
| Human | CREBBP | Histone Acetylation | TGCAATCAACGAAAGGTTCG |
| Human | CREBBP | Histone Acetylation | CTAGGGTTGGCATGAGGGCC |
| Human | CREBBP | Histone Acetylation | GCTTGCAGAGGTCTTTGAGC |
| Human | CBX8 | Chromatin Readers | GTCGCAGAAGTACAGCACAT |
| Human | CBX8 | Chromatin Readers | CTGAAGCGGCGCATACGGAA |
| Human | CBX8 | Chromatin Readers | GGTCGCAGAAGTACAGCACA |
| Human | CBX8 | Chromatin Readers | GCGCATACGGAAAGGACGCA |
| Human | CBX8 | Chromatin Readers | CTGCGAGCAAGCGAGCATCC |
| Human | INO80B | Chromatin Remodeling | CGGGAGCAAGCGTAGCGGCG |
| Human | INO80B | Chromatin Remodeling | CAAGCGTAGCGGCGCGGATG |
| Human | INO80B | Chromatin Remodeling | ACTGAAGACTACAGAGTGCC |
| Human | INO80B | Chromatin Remodeling | GCAAGCGTAGCGGCGCGGAT |
| Human | INO80B | Chromatin Remodeling | GCGGAAGCGGCGGCTCCAGG |
| Human | INO80B | Chromatin Remodeling | AGGAGCGGGCGCGGAAGCGG |
| Human | INO80B | Chromatin Remodeling | GCGAGGAGCGGGCGCGGAAG |
| Human | INO80B | Chromatin Remodeling | AGCGGCGGCTCCAGGCGGCG |
| Human | INO80B | Chromatin Remodeling | TGAAGCGCGAGGAGCGGGCG |
| Human | CHEK1 | Other Histone Modifications | GACAGAATAGAGCCAGACAT |
| Human | CHEK1 | Other Histone Modifications | GCTCGCTGGAGAATTGCCAT |
| Human | CHEK1 | Other Histone Modifications | AGCAAGTTGAACTTCTCCAT |
| Human | CHEK1 | Other Histone Modifications | TGGGACTTGGTGCAAACCCT |
| Human | CHEK1 | Other Histone Modifications | CAAGTCCCAGTCTTCCACAA |
| Human | CHEK1 | Other Histone Modifications | CTGGGACTTGGTGCAAACCC |
| Human | CHEK1 | Other Histone Modifications | AAGTCCCAGTCTTCCACAAA |
| Human | CHEK1 | Other Histone Modifications | AGTGCCCTTTGTGGAAGACT |
| Human | CHEK1 | Other Histone Modifications | GTTGACTTCCGGCTTTCTAA |
| Human | CHEK1 | Other Histone Modifications | ATCCATCACCCTTAGAAAGC |
| Human | CHEK1 | Other Histone Modifications | TTACTATATCAACAACTGAT |
| Human | CHEK1 | Other Histone Modifications | GCTTTCTAAGGGTGATGGAT |
| Human | CHEK1 | Other Histone Modifications | TTCCGGCTTTCTAAGGGTGA |
| Human | ENY2 | Transcription Elongation | TAGCTCTCAGCAACTCTTTG |
| Human | ENY2 | Transcription Elongation | GTTGAAGGCACACTGTAAAG |
| Human | ENY2 | Transcription Elongation | GTGGCTGAAATCACTCCAAA |

| | | | |
|---|---|---|---|
| Human | ENY2 | Transcription Elongation | GGAATGTTCTTATTCTTTGT |
| Human | ING4 | Chromatin Readers | ACCACTCAATGGAACAATCA |
| Human | ING4 | Chromatin Readers | CATGCAGACCTATGAGATGG |
| Human | ING4 | Chromatin Readers | CCAAAGAACAGAGGACCTGA |
| Human | ING4 | Chromatin Readers | AAAGAAAGGCCGGACTCAAA |
| Human | ING4 | Chromatin Readers | AGAGAAACTTTCAGCTCATG |
| Human | ING4 | Chromatin Readers | GAGAAACTTTCAGCTCATGA |
| Human | ING4 | Chromatin Readers | AAAGTTTCTCTGTAATTCAA |
| Human | ING4 | Chromatin Readers | AAGTTTCTCTGTAATTCAAA |
| Human | SND1 | Chromatin Readers | AAATGCCCAGGGCTCATCAG |
| Human | SND1 | Chromatin Readers | TCATCAAGATGGTCCTCTCA |
| Human | SND1 | Chromatin Readers | GAAATGCCCAGGGCTCATCA |
| Human | SND1 | Chromatin Readers | GTTGCCATTTGGATGAAGGA |
| Human | SND1 | Chromatin Readers | GGAAGGTTTCGCACGCTGTG |
| Human | SND1 | Chromatin Readers | ATGCTATCATCGAGCATGTG |
| Human | SND1 | Chromatin Readers | GGTACCATCCTTCATCCAAA |
| Human | SND1 | Chromatin Readers | GTGCGAAACCTTCCTTCAGG |
| Human | SND1 | Chromatin Readers | TGGAGGAATAAACATTGCTG |
| Human | SND1 | Chromatin Readers | TGGGAAGAAGGTCAATGTGA |
| Human | SND1 | Chromatin Readers | GAAGAAGGTCAATGTGACGG |
| Human | SND1 | Chromatin Readers | ATTCAGAACCTGCATCACCT |
| Human | SND1 | Chromatin Readers | GAGGCCAGAGCTATTAAGAA |
| Human | SND1 | Chromatin Readers | TTCAATGCCTGCAAGCAAGA |
| Human | SND1 | Chromatin Readers | GTGGTGGAATACGTCTTCAG |
| Human | SND1 | Chromatin Readers | GTAGTAGGAGCTGCGTTCGG |
| Human | SND1 | Chromatin Readers | ACTGGTGCTGCAGCGAGAGG |
| Human | SND1 | Chromatin Readers | CACCAGCAGGACGGACAGGT |
| Human | SND1 | Chromatin Readers | CATTGACTACGGCAACAGAG |
| Human | SND1 | Chromatin Readers | AGGCAGGTGATAGGGTACCC |
| Human | SND1 | Chromatin Readers | GAGACTCGACTTTCTCTACT |
| Human | SND1 | Chromatin Readers | ACGCCGTGGACAGCGTAGTT |
| Human | SND1 | Chromatin Readers | CGCTGTCCACGGCGTCCGTG |
| Human | SND1 | Chromatin Readers | GCTGTCCACGGCGTCCGTGC |
| Human | ZNRD1 | Transcription Elongation | GGTGACCGTATCCTGAGCCC |
| Human | ZNRD1 | Transcription Elongation | CTCGGTCCTGCCTCTGCCCG |
| Human | ZNRD1 | Transcription Elongation | CTGAACGCATCTGTCTGGTG |
| Human | ZNRD1 | Transcription Elongation | ATTCCTTCATGACCACATCG |
| Human | ZNRD1 | Transcription Elongation | CAGATGCGTTCAGCCGATGA |
| Human | MIS12 | Chromatin Remodeling | ATAATGTTGGCAGAGATCAT |
| Human | MIS12 | Chromatin Remodeling | ACAGAACATTAGAGACAATG |
| Human | MIS12 | Chromatin Remodeling | AGTGATGCAGGCCGTTGAAC |
| Human | MIS12 | Chromatin Remodeling | CTACCTATTTGAAGTGATGC |

| | | | |
|---|---|---|---|
| Human | MIS12 | Chromatin Remodeling | CGGCCTGCATCACTTCAAAT |
| Human | MIS12 | Chromatin Remodeling | AAGTTTCTTTGCTTCATGAA |
| Human | KDM3B | Histone Methylation | TTCTGCTGTTATCAACCCAT |
| Human | KDM3B | Histone Methylation | TAGATTGTGAACCTGGTGTG |
| Human | KDM3B | Histone Methylation | CGAAAGGTTGGAGAAGAACA |
| Human | KDM3B | Histone Methylation | GGAGGGTGCTCATGATGAAG |
| Human | KDM3B | Histone Methylation | TTTGTCGCAAATGTGGATTT |
| Human | KDM3B | Histone Methylation | CACTGGGTTTGTCGCAAATG |
| Human | KDM3B | Histone Methylation | AACTCTCTTCAACATCCACT |
| Human | KDM3B | Histone Methylation | TTGTCGCAAATGTGGATTTG |
| Human | KDM3B | Histone Methylation | TCTGCAGAGGAGTGAGACTC |
| Human | KDM3B | Histone Methylation | GAGTCAGTTGTAATGAAGAG |
| Human | KDM3B | Histone Methylation | TAAAGATGCCGAATCTGCAG |
| Human | KDM3B | Histone Methylation | AGGAGTCAGTTGTAATGAAG |
| Human | KDM3B | Histone Methylation | CAGAACACCAGATACTAGCT |
| Human | YWHAQ | Chromatin Readers | GCTGATCCAGAAGGCCAAGC |
| Human | YWHAQ | Chromatin Readers | CGACATGGCCACCTGCATGA |
| Human | YWHAQ | Chromatin Readers | GCTAGAGATGACCCTCCAGG |
| Human | YWHAQ | Chromatin Readers | GGAGCGCAACCTGCTCTCCG |
| Human | YWHAQ | Chromatin Readers | CTCCGTGGCCTACAAGAACG |
| Human | YWHAQ | Chromatin Readers | TAAGGACTATCGGGAGAAAG |
| Human | YWHAQ | Chromatin Readers | TGCAGCTGATTAAGGACTAT |
| Human | YWHAQ | Chromatin Readers | GCTGAGATCCATCTGCACCA |
| Human | YWHAQ | Chromatin Readers | GGTGCAGATGGATCTCAGCT |
| Human | YWHAQ | Chromatin Readers | GCAGCTGATTAAGGACTATC |
| Human | YWHAQ | Chromatin Readers | AAATGAAGGGTGATTACTTC |
| Human | YWHAQ | Chromatin Readers | TGCTTATATCAAATGCCTCT |
| Human | YWHAQ | Chromatin Readers | TTCCCAAGGAGCTTACCAAG |
| Human | YWHAQ | Chromatin Readers | CAAACGATAGATAATTCCCA |
| Human | YWHAQ | Chromatin Readers | TGCCTCTTGGTAAGCTCCTT |
| Human | YWHAQ | Chromatin Readers | ATGCCTCTTGGTAAGCTCCT |
| Human | YWHAQ | Chromatin Readers | TCTAAGCAACTGCATGATGA |
| Human | YWHAQ | Chromatin Readers | CTCTAAGCAACTGCATGATG |
| Human | TCF20 | Chromatin Readers | CCCAGAGAATACAACCCTCA |
| Human | TCF20 | Chromatin Readers | TGCTGTCAAGAGGTAGTTCA |
| Human | TCF20 | Chromatin Readers | GGATTTGTAACTCCAGCTCA |
| Human | TCF20 | Chromatin Readers | CAATCTGCATCAATGGCACA |
| Human | TCF20 | Chromatin Readers | AATCTGCATCAATGGCACAC |
| Human | TCF20 | Chromatin Readers | TAGCAAACAATCTGCATCAA |
| Human | TCF20 | Chromatin Readers | TGCAGATTGTTTGCTACATG |
| Human | TCF20 | Chromatin Readers | AAGGGCATGGAATTAAAGCA |
| Human | TCF20 | Chromatin Readers | TTTAATTCCATGCCCTTGTT |

| | | | |
|---|---|---|---|
| Human | TCF20 | Chromatin Readers | AACAGGATTCTTGTAACTTC |
| Human | TCF20 | Chromatin Readers | AGAAGTTACAAGAATCCTGT |
| Human | TCF20 | Chromatin Readers | ACAGGATTCTTGTAACTTCT |
| Human | TCF20 | Chromatin Readers | GGAGGGAGAGAAGGAAGCCT |
| Human | TCF20 | Chromatin Readers | GAGGGAGAGAAGGAAGCCTG |
| Human | TCF20 | Chromatin Readers | AAGCAGAAGACTGATAGTGT |
| Human | TCF20 | Chromatin Readers | GCTGGTTACCAGTTAAGAGT |
| Human | TCF20 | Chromatin Readers | TGCTGGTTACCAGTTAAGAG |
| Human | HDAC4 | Histone Acetylation | GCGGAGTCTGTGACATCCAA |
| Human | HDAC4 | Histone Acetylation | GAAGGGCAAAGAGAGTGCCG |
| Human | HDAC4 | Histone Acetylation | CTTGGCGGCCTTCAGAACGG |
| Human | HDAC4 | Histone Acetylation | GCACGTCCCAGTCCACGATG |
| Human | HDAC4 | Histone Acetylation | CAGGTACCCGAAGCATCTGG |
| Human | HDAC4 | Histone Acetylation | GTGGACTGGGACGTGCACCA |
| Human | HDAC4 | Histone Acetylation | TCCTCATCGCTGAGTTCCAG |
| Human | HDAC4 | Histone Acetylation | GTTGTTGCTTGATGTGCTCG |
| Human | HDAC4 | Histone Acetylation | CGAGCACATCAAGCAACAAC |
| Human | HDAC4 | Histone Acetylation | CAAGCAACAACAGGAGATGC |
| Human | HDAC4 | Histone Acetylation | AGCAGAAGCAGCAGATCCAG |
| Human | HDAC4 | Histone Acetylation | CTGCAGCTGCTGCTCCCGCA |
| Human | HDAC4 | Histone Acetylation | GCGGGAGCAGCAGCTGCAGC |
| Human | HDAC4 | Histone Acetylation | GCTGCAGCTGCTGCTCCCGC |
| Human | YY1 | DNA and RNA Modifications | TCTTTGTGCAGCCTTTATGA |
| Human | YY1 | DNA and RNA Modifications | AAAGATGTTCAGGGATAACT |
| Human | YY1 | DNA and RNA Modifications | AGGCTGCACAAAGATGTTCA |
| Human | YY1 | DNA and RNA Modifications | ATCTTTGTGCAGCCTTTATG |
| Human | YY1 | DNA and RNA Modifications | AAGGCTGCACAAAGATGTTC |
| Human | YY1 | DNA and RNA Modifications | AGTGCACGTTCGAAGGCTGT |
| Human | YY1 | DNA and RNA Modifications | CTTCTTATTACAACCATCGA |
| Human | YY1 | DNA and RNA Modifications | TTCTTATTACAACCATCGAA |
| Human | YY1 | DNA and RNA Modifications | ACCATCGAGACCACAGTGGT |
| Human | YY1 | DNA and RNA Modifications | GGTCACCGACGACCCGACCC |
| Human | YY1 | DNA and RNA Modifications | GACGTGGACTCTGGGACCGT |
| Human | YY1 | DNA and RNA Modifications | TCTGCACAGACGTGGACTCT |
| Human | YY1 | DNA and RNA Modifications | ACAATAGCTTGCCCTCATAA |
| Human | YY1 | DNA and RNA Modifications | CCAGTTGTTTGGGATCTGAG |
| Human | YY1 | DNA and RNA Modifications | GCCTTCGAACGTGCACTGAA |
| Human | YY1 | DNA and RNA Modifications | CCCTTTCAGTGCACGTTCGA |
| Human | YY1 | DNA and RNA Modifications | TCACATCTTAACACATGCTA |
| Human | YY1 | DNA and RNA Modifications | TCTCTCCAGTATGAACCAGT |
| Human | TAF4 | Transcription Elongation | CTCGTGCAGAATTTACTGGA |
| Human | TAF4 | Transcription Elongation | GGTAAGCTCCTCTTCAGGAA |

| | | | |
|---|---|---|---|
| Human | TAF4 | Transcription Elongation | TCCTCTTCAGGAAAGGCACA |
| Human | TAF4 | Transcription Elongation | AGGCGGGTAAGCTCCTCTTC |
| Human | TAF4 | Transcription Elongation | TCTAATATTCTTCTCTGCAA |
| Human | TAF4 | Transcription Elongation | TATCACATGCCACGCAACAA |
| Human | TAF4 | Transcription Elongation | ATTCTGTAGCCTTTGTTGCG |
| Human | SUPT5H | Transcription Elongation | CAAGGATGACATTGCTCAGG |
| Human | SUPT5H | Transcription Elongation | TCCTGGGTCCGCCTCAAGCG |
| Human | SUPT5H | Transcription Elongation | CATTGCTCAGGTGGACTACG |
| Human | SUPT5H | Transcription Elongation | GACAACGTGGAGGTCTGTGA |
| Human | SUPT5H | Transcription Elongation | TCTTGTTGCCATCCACGCTG |
| Human | SUPT5H | Transcription Elongation | GGCAAGATCCTCAGCGTGGA |
| Human | SUPT5H | Transcription Elongation | GGTGAGCTCATCAACCTGCA |
| Human | SUPT5H | Transcription Elongation | GCAGGGCAAGATCCTCAGCG |
| Human | SUPT5H | Transcription Elongation | GGCCAGCAATCACCTTCACG |
| Human | SUPT5H | Transcription Elongation | GCGACACAGGCCTCATTGTG |
| Human | SUPT5H | Transcription Elongation | TCTCCTCCACCCGCACAATG |
| Human | SUPT5H | Transcription Elongation | GGCCGATTCGAGGGCGACAC |
| Human | SUPT5H | Transcription Elongation | GGTGATTGCTGGCCGATTCG |
| Human | SUPT5H | Transcription Elongation | CTACAAAGGCTACATCGGTG |
| Human | SUPT5H | Transcription Elongation | CACAGAGTCCACGGCCCGTG |
| Human | SUPT5H | Transcription Elongation | GGGCCCTACAAAGGCTACAT |
| Human | SUPT5H | Transcription Elongation | ACACCGATGTAGCCTTTGTA |
| Human | SUPT5H | Transcription Elongation | AGACCGTGCGCATCTCCCAG |
| Human | SUPT5H | Transcription Elongation | GACTGTCAAATGTAAGATTG |
| Human | SUPT5H | Transcription Elongation | GTGGCACCAGAGCATGTGAA |
| Human | SUPT5H | Transcription Elongation | GAAGGGCTACATCTACGTGG |
| Human | SUPT5H | Transcription Elongation | CATGTCTGCTGTGATCACGG |
| Human | SUPT5H | Transcription Elongation | ATGTCTGCTGTGATCACGGA |
| Human | SUPT5H | Transcription Elongation | GAGAAGATCAGGTCCCTGGG |
| Human | SUPT5H | Transcription Elongation | AGGTGGTGACTGAGAGCACA |
| Human | SUPT5H | Transcription Elongation | TGAGAAGATCAGGTCCCTGG |
| Human | SUPT5H | Transcription Elongation | GTGCGGCTACCATCCTGGAG |
| Human | SUPT5H | Transcription Elongation | GGGTGCGGCTACCATCCTGG |
| Human | SUPT5H | Transcription Elongation | GGTGCGGCTACCATCCTGGA |
| Human | SUPT5H | Transcription Elongation | GGCTCCCGAACACCCATGTA |
| Human | SUPT5H | Transcription Elongation | GAACTGGTCTGTGTTGTACA |
| Human | SUPT5H | Transcription Elongation | TGGTCTGTGTTGTACATGGC |
| Human | SUPT5H | Transcription Elongation | GGGCTAGCCTGATAGGCCAT |
| Human | SUPT5H | Transcription Elongation | TTGTATTGTGGGTTGACCTG |
| Human | SUPT5H | Transcription Elongation | GTATTGTGGGTTGACCTGTG |
| Human | SUPT5H | Transcription Elongation | CCAGAATGAAGCCTCCATGG |
| Human | SUPT5H | Transcription Elongation | AGAATGAAGCCTCCATGGCG |

| | | | |
|---|---|---|---|
| Human | SUPT5H | Transcription Elongation | AGGCCGAGGTAGACGAAGAG |
| Human | SUPT5H | Transcription Elongation | CCGCCGCTCTTCGTCTACCT |
| Human | SUPT5H | Transcription Elongation | TGTCATAGGACTGTAGCCAA |
| Human | SUPT5H | Transcription Elongation | TCATAGGACTGTAGCCAACG |
| Human | SUPT5H | Transcription Elongation | GGCTACAGTCCTATGACACC |
| Human | SUPT5H | Transcription Elongation | GTCATAGGACTGTAGCCAAC |
| Human | SUPT5H | Transcription Elongation | GAGGGAGCTCCAGGTGTCAT |
| Human | ACTR5 | Chromatin Remodeling | TGCAGGAGCTCAATGCCCGG |
| Human | ACTR5 | Chromatin Remodeling | GGCGGCAGCAGCAATTGCGG |
| Human | ACTR5 | Chromatin Remodeling | AGGAGCTCAATGCCCGGCGG |
| Human | ACTR5 | Chromatin Remodeling | GCTCAATGCCCGGCGGCGGG |
| Human | ACTR5 | Chromatin Remodeling | GGAGCTCAATGCCCGGCGGC |
| Human | ACTR5 | Chromatin Remodeling | GCTCAGTATAGCAGTGGAGC |
| Human | ACTR5 | Chromatin Remodeling | CCAGAAGCTCAGTATAGCAG |
| Human | ACTR5 | Chromatin Remodeling | TATACTGAGCTTCTGGATGT |
| Human | ACTR5 | Chromatin Remodeling | CCACTGCTATACTGAGCTTC |
| Human | PHIP | Chromatin Readers | TATGGGCAACAGATGATGGG |
| Human | PHIP | Chromatin Readers | ACTGGCAGACGGATATTTAC |
| Human | PHIP | Chromatin Readers | AGGTTCGAAGACACCAGACT |
| Human | PHIP | Chromatin Readers | GAGAATACCATGATAGCAGC |
| Human | PHIP | Chromatin Readers | ACAACTTCCAGCTGCTATCA |
| Human | PHIP | Chromatin Readers | CAGTTCTCACCATTGTGCAG |
| Human | PHIP | Chromatin Readers | GAGATATCTATCTTCTACTG |
| Human | PHIP | Chromatin Readers | AGAGATATCTATCTTCTACT |
| Human | PHIP | Chromatin Readers | CTCTTTGAGCCACTGCACAA |
| Human | PHIP | Chromatin Readers | AAGAGATATCTATCTTCTAC |
| Human | PHIP | Chromatin Readers | GGTGGAATGTTTCTGGCGAC |
| Human | PHIP | Chromatin Readers | AAGCACAGATCATATTATTC |
| Human | PHIP | Chromatin Readers | TAGTGCTGGTGGAATGTTTC |
| Human | PHIP | Chromatin Readers | TGGTGGAATGTTTCTGGCGA |
| Human | PHIP | Chromatin Readers | ACTAGTAACAGGTTTGTAAG |
| Human | PHIP | Chromatin Readers | TACAAACCTGTTACTAGTGT |
| Human | PHIP | Chromatin Readers | GGCAATTTAAACGAAGAGAG |
| Human | PHIP | Chromatin Readers | TTGTAAGTGGCAGTCGTGAT |
| Human | PHIP | Chromatin Readers | GAAGGTTACTATGGTAGCTT |
| Human | PHIP | Chromatin Readers | AAAGTTTGGAATTCTTACAC |
| Human | PHIP | Chromatin Readers | ATAACATGACTCTGAAAGTT |
| Human | PHIP | Chromatin Readers | TGAAGGTTACTATGGTAGCT |
| Human | PHIP | Chromatin Readers | ACTCTAGGATCGAACGGGTG |
| Human | PHIP | Chromatin Readers | TGATGGAAACGTGATAGTGT |
| Human | PHIP | Chromatin Readers | ATAGTGTGGGATCTGGCAAG |
| Human | PHIP | Chromatin Readers | AAACGTGATAGTGTGGGATC |

| | | | |
|---|---|---|---|
| Human | PHIP | Chromatin Readers | ATGATGGAAACGTGATAGTG |
| Human | PHIP | Chromatin Readers | TTTGCATGCACAGACTCTCA |
| Human | PHIP | Chromatin Readers | GACTGCAAATGCTCTCCTGA |
| Human | PHIP | Chromatin Readers | CACAGTAGTGGCATATCCAA |
| Human | PHIP | Chromatin Readers | TACTGTGCAATACATGGGAT |
| Human | PHIP | Chromatin Readers | TCCCATGTATTGCACAGTAG |
| Human | PHIP | Chromatin Readers | TGGGAATTATGAGTCACCAA |
| Human | PHIP | Chromatin Readers | AGACATCATTGACACTCCAA |
| Human | PHIP | Chromatin Readers | AGCCTCTAAAGTTTCTCTAA |
| Human | PHIP | Chromatin Readers | CTCTGTAGTCTGGATATTCA |
| Human | PHIP | Chromatin Readers | GGATATTCAAGGAGATCTAC |
| Human | PHIP | Chromatin Readers | TCAATGCTACAATGTTTGCT |
| Human | PHIP | Chromatin Readers | GATACTGGTAAACTGACTGG |
| Human | PHIP | Chromatin Readers | AGCATTGAAACAGACTATCA |
| Human | PHIP | Chromatin Readers | TCAGGGTACTCAAGTTGAAG |
| Human | PHIP | Chromatin Readers | GTTGGCATAAAGTATGAAGT |
| Human | PHIP | Chromatin Readers | TGAACTGCTTTCTGCTTTCT |
| Human | PHIP | Chromatin Readers | GTTCTTCAAAGAAAGCAGAC |
| Human | RNF2 | Other Histone Modifications | TTCTTTGTTGCCACTTCTAA |
| Human | RNF2 | Other Histone Modifications | ATCATCACAGCCCTTAGAAG |
| Human | RNF2 | Other Histone Modifications | TAAACACTCCTTTGTAGTCA |
| Human | RNF2 | Other Histone Modifications | ATTCTTTGTTGCCACTTCTA |
| Human | RNF2 | Other Histone Modifications | GAAGAACACCATGACTACAA |
| Human | RNF2 | Other Histone Modifications | GTTACAACGAACACCTCAGG |
| Human | RNF2 | Other Histone Modifications | GCCATGAACAGACTGCAGCG |
| Human | RNF2 | Other Histone Modifications | CCATCTGTTATTGCCTCCTG |
| Human | RNF2 | Other Histone Modifications | AGCACTCAGTCACAGCATTG |
| Human | RNF2 | Other Histone Modifications | TATCCAAGTATCTGGCTGTG |
| Human | RNF2 | Other Histone Modifications | TAACCTCACAGCCAGATACT |
| Human | CDK1 | Other Histone Modifications | CACATGAGGTAGTAACACTC |
| Human | TDRD10 | Chromatin Readers | TCCTCCTTAGAAATATCCAG |
| Human | TDRD10 | Chromatin Readers | TCCACTGGATATTTCTAAGG |
| Human | TDRD10 | Chromatin Readers | GGCTGCGCAGAGACTGCACA |
| Human | TDRD10 | Chromatin Readers | GACACGCCTGGAACAGGTGT |
| Human | TAF1B | Transcription Elongation | CAGTGTGCTGCTGTCTCATG |
| Human | TAF1B | Transcription Elongation | TCAGTGTGCTGCTGTCTCAT |
| Human | TAF1B | Transcription Elongation | CTCAGTGTGCTGCTGTCTCA |
| Human | TAF1B | Transcription Elongation | AGATTATCTTCTAATACCGT |
| Human | TAF1B | Transcription Elongation | GCAGAAGCCTTAAAGAACCT |
| Human | TAF1B | Transcription Elongation | GCCTTAAAGAACCTTGGAGT |
| Human | TAF1B | Transcription Elongation | AAGTAAGGACAGATAACAGA |
| Human | TAF1B | Transcription Elongation | TAATCTAAGTCATTCAGACT |

| | | | |
|---|---|---|---|
| Human | TAF1B | Transcription Elongation | AGCTATGCATTTCATCAGGG |
| Human | TAF1B | Transcription Elongation | TTAAGCTATGCATTTCATCA |
| Human | TAF1B | Transcription Elongation | AGAGTCTTGGCCTGACTACG |
| Human | MSH6 | Chromatin Readers | TTTGATGACAGCCCAACAAG |
| Human | MSH6 | Chromatin Readers | GGGTGGTTGTAAACCAGACA |
| Human | MSH6 | Chromatin Readers | CACAACCACCTAGAGCAGAG |
| Human | MSH6 | Chromatin Readers | ACAACCACCTAGAGCAGAGA |
| Human | MSH6 | Chromatin Readers | GTAAGCGGCTCCTAAAGCAA |
| Human | MSH6 | Chromatin Readers | CTCTTATGTCAGCAAGAGCT |
| Human | MSH6 | Chromatin Readers | TATCATTGAAGGACTGCATG |
| Human | MSH6 | Chromatin Readers | CATAAATGCTGAAGAACGGA |
| Human | MSH6 | Chromatin Readers | TCATAAATGCTGAAGAACGG |
| Human | MSH6 | Chromatin Readers | ACGGAGGGATGTATCATTGA |
| Human | MSH6 | Chromatin Readers | ATCTCATAAATGCTGAAGAA |
| Human | MSH6 | Chromatin Readers | TGTAGAGTGTATCGCAGTGT |
| Human | PHF6 | Chromatin Readers | TTGTAAGTCAAATAGAGACA |
| Human | PHF6 | Chromatin Readers | TCAAATAGAGACAAGGAATG |
| Human | PHF6 | Chromatin Readers | ACCTTCACAAGGAATTTACA |
| Human | PHF6 | Chromatin Readers | TCACAACCAATTGTTGCTCC |
| Human | PHF6 | Chromatin Readers | GCATATATTTAATGCCAAGA |
| Human | PHF6 | Chromatin Readers | AAACAACATGCACTTATAAT |
| Human | PHF6 | Chromatin Readers | AAGACTTACCATTACCACTG |
| Human | PHF6 | Chromatin Readers | GTAATGGTAAGTCTTAACAC |
| Human | PHF6 | Chromatin Readers | AGTCAGCCTGGTGCTACTAT |
| Human | PHF6 | Chromatin Readers | CACTCTGATAATGAAAGTCT |
| Human | PHF6 | Chromatin Readers | TCTGATAATGAAAGTCTTGG |
| Human | PHF6 | Chromatin Readers | TTCTATTGAAGATGTCCAAA |
| Human | PHF6 | Chromatin Readers | ACAACATCAAGAGCAGAATT |
| Human | PHF6 | Chromatin Readers | CTCTTGATGTTGTTGTGAGC |
| Human | PRC1 | Control | GTAGTCCTCAGCACAGAAAG |
| Human | PRC1 | Control | GAAAGCGCTGCAATTAGAAG |
| Human | PRC1 | Control | CCAAATCGATTTACAAACCG |
| Human | PRC1 | Control | TGAAGGCACGAATTGAATTG |
| Human | PRC1 | Control | TCTTTGAAGGTGTCCAGAAG |
| Human | PRC1 | Control | GCAGAAATTCATGGAGTATG |
| Human | PRC1 | Control | TTCTCGCTGAAGCCCAACAG |
| Human | PRC1 | Control | CACCTATTCTGAGTTTGCGA |
| Human | PRC1 | Control | AGGAGGAAGTCGAGACACGG |
| Human | PRC1 | Control | ACTGGCTTGCTGCCAGAAGG |
| Human | POU4F1 | Transcription Elongation | GCGAACGCCTCGAGCTCGCG |
| Human | POU4F1 | Transcription Elongation | GGCGTGACGCAGGCCGACGT |
| Human | POU4F1 | Transcription Elongation | CACGGACCCGCGCGAGCTCG |

| | | | |
|---|---|---|---|
| Human | POU4F1 | Transcription Elongation | AACCTCAAGATCCCGGGCGT |
| Human | POU4F1 | Transcription Elongation | CCTGCAGGCGTGGCTCGAGG |
| Human | POU4F1 | Transcription Elongation | GCGGCGATCTTCTCGGACGA |
| Human | POU4F1 | Transcription Elongation | CACGGCGAAGTAGGCCTCGA |
| Human | POU4F1 | Transcription Elongation | CGGCGCCCACGACGGCCCGG |
| Human | POU4F1 | Transcription Elongation | GGCGCCCACGACGGCCCGGG |
| Human | POU4F1 | Transcription Elongation | CCGCCATGAACATGCCGTCC |
| Human | POU4F1 | Transcription Elongation | CCCGGACGGCATGTTCATGG |
| Human | POU4F1 | Transcription Elongation | CAGCCCGGACGGCATGTTCA |
| Human | POU4F1 | Transcription Elongation | GGCGGCGGCGCACCACGGCG |
| Human | POU4F1 | Transcription Elongation | GCCGCCATGAACATGCCGTC |
| Human | SATB1 | Chromatin Remodeling | TGTCTTCCGAAATCTACCAG |
| Human | SATB1 | Chromatin Remodeling | CACACGTGCAAATACCGCCT |
| Human | SATB1 | Chromatin Remodeling | TGAAAGAGACCGAATATACC |
| Human | SATB1 | Chromatin Remodeling | AAGACTGGGATGCAGTCTTG |
| Human | SATB1 | Chromatin Remodeling | ACAGCTCGCACAACCATCCC |
| Human | SATB1 | Chromatin Remodeling | GTCTCAAGCACTGTTTGCAA |
| Human | SATB1 | Chromatin Remodeling | CAGAGTTTCATACAAGACGT |
| Human | SATB1 | Chromatin Remodeling | CCAGAGTTTCATACAAGACG |
| Human | SATB1 | Chromatin Remodeling | GTACTTGGGAAGGTCGAGCT |
| Human | SATB1 | Chromatin Remodeling | CCACGTCTTGTATGAAACTC |
| Human | SATB1 | Chromatin Remodeling | GCCAAAGGGCTAATCCAGGT |
| Human | SATB1 | Chromatin Remodeling | CCAGGCCAAAGGGCTAATCC |
| Human | SATB1 | Chromatin Remodeling | GGAGCATGCAGAATTTGTGC |
| Human | SATB1 | Chromatin Remodeling | GCATCTGTCACGTAAGACAG |
| Human | SATB1 | Chromatin Remodeling | AAGACTTGCCTCCCGAACAA |
| Human | SATB1 | Chromatin Remodeling | GACATGATTGGCGCCTTGCT |
| Human | SATB1 | Chromatin Remodeling | AATGTCAAGAATTTGGAAGG |
| Human | SATB1 | Chromatin Remodeling | CTTTCTGAGCTATCCCAGCA |
| Human | SATB1 | Chromatin Remodeling | AGGCGGGCAATCTTGGCTGG |
| Human | SATB1 | Chromatin Remodeling | ATTGCCCGCCTGGAGCAGAA |
| Human | SATB1 | Chromatin Remodeling | TCCAGGCGGGCAATCTTGGC |
| Human | SATB1 | Chromatin Remodeling | ATCTTGGCTGGTGGACCCTT |
| Human | SATB1 | Chromatin Remodeling | TTGCCCGCCTGGAGCAGAAC |
| Human | SATB1 | Chromatin Remodeling | TATACCAGGACGAAAGGGAA |
| Human | SATB1 | Chromatin Remodeling | ACCGAATATACCAGGACGAA |
| Human | SATB1 | Chromatin Remodeling | CCCTTTCGTCCTGGTATATT |
| Human | SATB1 | Chromatin Remodeling | CCGAATATACCAGGACGAAA |
| Human | INO80 | Chromatin Remodeling | AGGAAATGCGGGAGGCCAAG |
| Human | INO80 | Chromatin Remodeling | ATGGGCATACAACTCTGTCT |
| Human | INO80 | Chromatin Remodeling | GCTTGCTCACCAGTGCATGA |
| Human | INO80 | Chromatin Remodeling | ACAGAAGCATCTCCTTGGTG |

| | | | |
|---|---|---|---|
| Human | INO80 | Chromatin Remodeling | GAGGATCTTCCAACGAACAC |
| Human | INO80 | Chromatin Remodeling | GCTGAGAGAGAGAACATTTG |
| Human | INO80 | Chromatin Remodeling | CAGAGCCCAAAGCTCTGCCA |
| Human | INO80 | Chromatin Remodeling | AGGAGAAGAGTGAGATTCAG |
| Human | INO80 | Chromatin Remodeling | AAACCATGTATTCCTCCAGT |
| Human | INO80 | Chromatin Remodeling | AGACCTACTGGAGGAATACA |
| Human | INO80 | Chromatin Remodeling | TGACTCGGCTCAAGTCTCAA |
| Human | INO80 | Chromatin Remodeling | CAATATCAGTACTACAGTGC |
| Human | INO80 | Chromatin Remodeling | GCTAAGTGATGAATCCAGCG |
| Human | INO80 | Chromatin Remodeling | CTCATAGAAAGGGTCATATG |
| Human | INO80 | Chromatin Remodeling | ATATTCACCAAGCTCGGACA |
| Human | INO80 | Chromatin Remodeling | CAGAGACAAATCGATATAGG |
| Human | INO80 | Chromatin Remodeling | GGTAGTTAACATCACACAGG |
| Human | INO80 | Chromatin Remodeling | AAACGAGATATGGGTCATGA |
| Human | INO80 | Chromatin Remodeling | CAAATGACCTTGTCCGAGCT |
| Human | INO80 | Chromatin Remodeling | TTGGCCAATCTATATGAACA |
| Human | INO80 | Chromatin Remodeling | AAGAGTTGGAGAAGAAACTG |
| Human | INO80 | Chromatin Remodeling | TCAATCATTCACGAGACAGG |
| Human | INO80 | Chromatin Remodeling | AATGGCACTGCGGTAACTCG |
| Human | DYNLRB1 | Control | CATGAGGCTGGCATACTGGG |
| Human | DYNLRB1 | Control | GATGAAGCTGTGCATGAGGC |
| Human | DYNLRB1 | Control | TGTGCATGAGGCTGGCATAC |
| Human | DYNLRB1 | Control | GTGCATGAGGCTGGCATACT |
| Human | DIDO1 | Chromatin Readers | TATCTAAAGAGCTTTCCACG |
| Human | DIDO1 | Chromatin Readers | GAGGCCAGCGAGATCTGTGA |
| Human | DIDO1 | Chromatin Readers | GACTCCATCACAGATCTCGC |
| Human | DIDO1 | Chromatin Readers | TTTCCTCACGCAGAACACGA |
| Human | DIDO1 | Chromatin Readers | ATCATAAACCTGTTGTTGTG |
| Human | DIDO1 | Chromatin Readers | TGGTTTCATGGCGATTGTGT |
| Human | DIDO1 | Chromatin Readers | CATGAAACCATTCTTCACAG |
| Human | DIDO1 | Chromatin Readers | ATGGTTTCATGGCGATTGTG |
| Human | DIDO1 | Chromatin Readers | GTGGGCATTTCTGAGGCTCG |
| Human | DIDO1 | Chromatin Readers | CACCGGTGGTTCTGGAAGAG |
| Human | DIDO1 | Chromatin Readers | GCCCGTCACCTCGTTATACG |
| Human | DIDO1 | Chromatin Readers | CTGGAAGTGAAATCGCGAAG |
| Human | DIDO1 | Chromatin Readers | TGGGCCCACGAAATGACCAG |
| Human | DIDO1 | Chromatin Readers | CTTCTAACGATGGGCCACGA |
| Human | DIDO1 | Chromatin Readers | GAGAGCGCGAGCGAGACCGG |
| Human | DIDO1 | Chromatin Readers | TGGAGTCCTTGTCCCGGTGT |
| Human | DIDO1 | Chromatin Readers | CGAGAGCGCGAGGCCGACCG |
| Human | DIDO1 | Chromatin Readers | AGGAGGAGAGAGCGCGAGCG |
| Human | DIDO1 | Chromatin Readers | CCGAAACTGGAGCCGAGAGC |

| | | | |
|---|---|---|---|
| Human | L3MBTL2 | Chromatin Readers | CCTCTGTCATCCAGACAGCA |
| Human | L3MBTL2 | Chromatin Readers | CGTCCCACTCTATGACCAGT |
| Human | L3MBTL2 | Chromatin Readers | ACGTGCTTGAAACAGCTGAC |
| Human | L3MBTL2 | Chromatin Readers | TGATGGCACACCAGCCAATG |
| Human | L3MBTL2 | Chromatin Readers | CAGAGGCGATCCAGTACACC |
| Human | L3MBTL2 | Chromatin Readers | GGATTTCCACATCAAGATGG |
| Human | L3MBTL2 | Chromatin Readers | CGTGGATTTCCACATCAAGA |
| Human | L3MBTL2 | Chromatin Readers | GTCGTGACCAACCCACTGGG |
| Human | L3MBTL2 | Chromatin Readers | AGCCCACCAGCCGTTTCATG |
| Human | L3MBTL2 | Chromatin Readers | GTACGAGCAGTCTACACAGA |
| Human | L3MBTL2 | Chromatin Readers | CTGATGATCTGTGTGGACGG |
| Human | L3MBTL2 | Chromatin Readers | ATCACAGTAGATCTTCCGGA |
| Human | L3MBTL2 | Chromatin Readers | CGAGCAGTCTACACAGAAGG |
| Human | L3MBTL2 | Chromatin Readers | CTGCTCGTACCTTCTTGAAG |
| Human | L3MBTL2 | Chromatin Readers | AACATGGATTGCCCAAACCA |
| Human | L3MBTL2 | Chromatin Readers | GGACAGCGAGTACGACCAGT |
| Human | L3MBTL2 | Chromatin Readers | TGTGGCCACGGTGAAACGAG |
| Human | L3MBTL2 | Chromatin Readers | TGTGTGGTATCGTGGGTACA |
| Human | L3MBTL2 | Chromatin Readers | TGTGAGATGTGTGGTATCGT |
| Human | L3MBTL2 | Chromatin Readers | CCTTCTTCTCCAAGACCAAG |
| Human | L3MBTL2 | Chromatin Readers | GTGTGGTATCGTGGGTACAA |
| Human | L3MBTL2 | Chromatin Readers | CTGTGAGATGTGTGGTATCG |
| Human | ARIH1 | Control | TACCCTAACTCGTATTTCAC |
| Human | ARIH1 | Control | GGCCAGTGAAATACGAGTTA |
| Human | ARIH1 | Control | TATTTCACTGGCCTTGAATG |
| Human | ARIH1 | Control | AGGCCAGTGAAATACGAGTT |
| Human | ARIH1 | Control | AACCTGTTCGCTGCAAATGT |
| Human | ARIH1 | Control | AGTGCAATCGACTGTTAAAG |
| Human | ARIH1 | Control | AACAACATGGTGGCAATCTG |
| Human | ARIH1 | Control | GGATGGTGGTTGTAATCACA |
| Human | ARIH1 | Control | GTCACAATTGAGAAGGATGG |
| Human | ARIH1 | Control | CTCAATTGTGACATGGCATT |
| Human | ARIH1 | Control | CATGGTGGAATGTATCCGGG |
| Human | ARIH1 | Control | ACACATGGTGGAATGTATCC |
| Human | ARIH1 | Control | ATGTATCCGGGAGGTCAACG |
| Human | ARIH1 | Control | GCAGATTCTACAACACATGG |
| Human | ARIH1 | Control | CGAGCAGATTCTACAACACA |
| Human | ARIH1 | Control | TATTCTCAAAGATAATGGAC |
| Human | TDRKH | Chromatin Readers | GAGCCAATATTAAACAGCTG |
| Human | TDRKH | Chromatin Readers | GAGCGAGTGCTGCTTATCAG |
| Human | TDRKH | Chromatin Readers | CTGTGAAACTCATCATTGGC |
| Human | TDRKH | Chromatin Readers | TCGGATTGATGTGGACACAG |

| | | | |
|---|---|---|---|
| Human | TDRKH | Chromatin Readers | GGCAGAATCATAGGGAGAGG |
| Human | TDRKH | Chromatin Readers | GTGGGCAGAATCATAGGGAG |
| Human | TDRKH | Chromatin Readers | GATCTGTGGGCAGAATCATA |
| Human | TDRKH | Chromatin Readers | ATTCTGCCCACAGATCTCTG |
| Human | TDRKH | Chromatin Readers | AGTCACTCCTGAGAGCCCTG |
| Human | TDRKH | Chromatin Readers | TAATGGAGATTGCCCACTGA |
| Human | TDRKH | Chromatin Readers | TTGCCCACTGAAGGACCTCA |
| Human | TDRKH | Chromatin Readers | TGGTTCCTGGTATCGAGCCC |
| Human | TDRKH | Chromatin Readers | CGATACCAGGAACCATTTGT |
| Human | LMNB2 | DNA and RNA Modifications | GCGCGAGGTGAGTGGCATCA |
| Human | LMNB2 | DNA and RNA Modifications | GTGACCACGCGCGAGGTGAG |
| Human | LMNB2 | DNA and RNA Modifications | GATGCCACTCACCTCGCGCG |
| Human | LMNB2 | DNA and RNA Modifications | ATGGCGAGGAAGTGGCCATG |
| Human | LMNB2 | DNA and RNA Modifications | CGGACAAGGATCAGTCTCTG |
| Human | LMNB2 | DNA and RNA Modifications | CCACACCGTGACCATCTGGC |
| Human | LMNB2 | DNA and RNA Modifications | GCCTGCGCTGCTCCCGACGG |
| Human | LMNB2 | DNA and RNA Modifications | TGTCGCCCACGCGCCTGTCG |
| Human | LMNB2 | DNA and RNA Modifications | CTGCAGCCGCGACAGGCGCG |
| Human | LMNB2 | DNA and RNA Modifications | GACAGGCGCGTGGGCGACAG |
| Human | LMNB2 | DNA and RNA Modifications | GCGTGGCGGGCCCGCCCGCG |
| Human | LMNB2 | DNA and RNA Modifications | CTCATTCTCACGCATCACCG |
| Human | ZW10 | Chromatin Remodeling | GATAGAGAGTGAGGTCCGCC |
| Human | ZW10 | Chromatin Remodeling | TTCAGATAGCTTATCCACCT |
| Human | ZW10 | Chromatin Remodeling | CACTGAACAATCGCACAAAG |
| Human | YWHAH | Chromatin Readers | AGCAGCTGCTGCAGCGGGCG |
| Human | YWHAH | Chromatin Readers | GCTGCTGCAGCGGGCGCGGC |
| Human | YWHAH | Chromatin Readers | CGACATGGCCTCCGCTATGA |
| Human | YWHAH | Chromatin Readers | CTCTGTGGCCTACAAGAATG |
| Human | YWHAH | Chromatin Readers | AGATCGAAATCTCCTCTCTG |
| Human | YWHAH | Chromatin Readers | TGCCAGGCGATCTTCCTGGA |
| Human | YWHAH | Chromatin Readers | GGCACCAACCACATTCTTGT |
| Human | YWHAH | Chromatin Readers | AATGCTGCTAATGACCCTCC |
| Human | YWHAH | Chromatin Readers | TTACCGGGAGAAGATTGAGA |
| Human | YWHAH | Chromatin Readers | GGAGAAAGTTAAAGCTTACC |
| Human | YWHAH | Chromatin Readers | GCTCCTTCTCAATCTTCTCC |
| Human | YWHAH | Chromatin Readers | TGGAGAAAGTTAAAGCTTAC |
| Human | YWHAH | Chromatin Readers | GGAGAAGATTGAGAAGGAGC |
| Human | YWHAH | Chromatin Readers | GATTTCAAAGGCTTCCTTGT |
| Human | YWHAH | Chromatin Readers | AGCTTCTGAAGCTGCCTACA |
| Human | YWHAH | Chromatin Readers | ACACGGAGAAGTTGAGGGCC |
| Human | YWHAH | Chromatin Readers | AGTTGAGGGCCAGGCCCAGC |
| Human | YWHAH | Chromatin Readers | CGAAGGCTTGTTTGGCTAAG |

| | | | |
|---|---|---|---|
| Human | YWHAH | Chromatin Readers | GGCTTGTTTGGCTAAGAGGC |
| Human | YWHAH | Chromatin Readers | CAGCAACTGCATGATCAGCG |
| Human | CDYL | Chromatin Readers | GTGGAAAGGCTATGACAGCG |
| Human | CDYL | Chromatin Readers | TGTGGCGTCTGTTGAAGTCG |
| Human | CDYL | Chromatin Readers | ACAGCACCTCGTGAACTGTG |
| Human | CDYL | Chromatin Readers | GGAAGACAGAGTATTTGGTT |
| Human | CDYL | Chromatin Readers | GGGCCCAATAAAGCTCCGAC |
| Human | CDYL | Chromatin Readers | TCCTGGCCCTCTCGGCACCG |
| Human | CDYL | Chromatin Readers | AATCACGAGTGCCTTAGGAG |
| Human | CDYL | Chromatin Readers | AAGCACAACTGGATTGCACG |
| Human | CDYL | Chromatin Readers | ATGTTTCCCAAGATAATGGG |
| Human | CDYL | Chromatin Readers | GTGCAATCCAGTTGTGCTTG |
| Human | CDYL | Chromatin Readers | CCATGTTAAAGTACTTGCAG |
| Human | CDYL | Chromatin Readers | CCTCTGCAAGTACTTTAACA |
| Human | MBTD1 | Chromatin Readers | TCCAGCAATCCAGAAGACTT |
| Human | MBTD1 | Chromatin Readers | CTGGCTGCACACCAACCAAC |
| Human | MBTD1 | Chromatin Readers | ATTGGTTGGTCTCGAAGCAT |
| Human | MBTD1 | Chromatin Readers | GTATTAATGGGCTGTGCATA |
| Human | MBTD1 | Chromatin Readers | CATTAATACATCATATTGGT |
| Human | MBTD1 | Chromatin Readers | ATTAGAAAGGTGCTAGCTGA |
| Human | MBTD1 | Chromatin Readers | TAATTTCACAGAAACCGACA |
| Human | MBTD1 | Chromatin Readers | AAGCAGCAGACGGATCTGAC |
| Human | MBTD1 | Chromatin Readers | TTAATTTCACAGAAACCGAC |
| Human | MBTD1 | Chromatin Readers | CCAAATCACGGATTTCGTGT |
| Human | MBTD1 | Chromatin Readers | ATGAATAATTCGAGTTACTG |
| Human | MBTD1 | Chromatin Readers | GTTAACTGACACCACCCTAC |
| Human | MBTD1 | Chromatin Readers | GACACCACCCTACAGGATAG |
| Human | MBTD1 | Chromatin Readers | ATGGCTACCTGTGAGATGTG |
| Human | MBTD1 | Chromatin Readers | AACCATCCCACACATCTCAC |
| Human | MBTD1 | Chromatin Readers | TACCTGTGAGATGTGTGGGA |
| Human | MBTD1 | Chromatin Readers | TGTGAGATGTGTGGGATGGT |
| Human | MBTD1 | Chromatin Readers | TGGCTACCTGTGAGATGTGT |
| Human | PWWP2B | Chromatin Readers | ACCTTCGCTTCTCGCCAAGA |
| Human | PWWP2B | Chromatin Readers | GGCGAGAAGCGAAGGTCTCG |
| Human | PWWP2B | Chromatin Readers | GCCGTCTTGGCGAGAAGCGA |
| Human | PWWP2B | Chromatin Readers | CCGGGACACGTACAAGCTGT |
| Human | PWWP2B | Chromatin Readers | CCGTACCCTCCCTACTTCGA |
| Human | PWWP2B | Chromatin Readers | GGGCGCCTTCGAAGTAGGGA |
| Human | EPC2 | Chromatin Remodeling | TTGCTGTTTCTGCTGAAGCT |
| Human | EPC2 | Chromatin Remodeling | AGCAGTTAGTTCAGATGCAA |
| Human | EPC2 | Chromatin Remodeling | ATTGCTGTTTCTGCTGAAGC |
| Human | SUV420H2 | Histone Methylation | GCAGTCATGGTTGATGAAGG |

| | | | |
|---|---|---|---|
| Human | SUV420H2 | Histone Methylation | TCATGTACTCAACCCGCAAG |
| Human | SUV420H2 | Histone Methylation | TTTGCAGTCATGGTTGATGA |
| Human | SUV420H2 | Histone Methylation | TCCATGGAGTAGCGCGTGCA |
| Human | SUV420H2 | Histone Methylation | GCGCTATGGGCTGCCTTACG |
| Human | SUV420H2 | Histone Methylation | GTAAGGCAGCCCATAGCGCC |
| Human | SUV420H2 | Histone Methylation | CGCGTCTCCCTGCACCGATG |
| Human | SUV420H2 | Histone Methylation | AGGTCCACACGCACCACGTA |
| Human | SUV420H2 | Histone Methylation | ACTGTGCGAGAACGACGACC |
| Human | SUV420H2 | Histone Methylation | CCTGGCCACCAGCCTCGTCC |
| Human | SUV420H2 | Histone Methylation | CCAGGACGAGGCTGGTGGCC |
| Human | SUV420H2 | Histone Methylation | AAGTTTCCAGCGCTGAGCGC |
| Human | SUV420H2 | Histone Methylation | GCGATAGACGTGGGTCTTGA |
| Human | SUV420H2 | Histone Methylation | AGCGATAGACGTGGGTCTTG |
| Human | ZMYND8 | Chromatin Readers | AGATGTATTCCGCATAGTCA |
| Human | ZMYND8 | Chromatin Readers | CCTTCCTGGCTGATGCAAAG |
| Human | ZMYND8 | Chromatin Readers | AGGGTGCTGTTCCAATGGAA |
| Human | ZMYND8 | Chromatin Readers | GGAATACATCTTCCATCCAA |
| Human | ZMYND8 | Chromatin Readers | ATTCTTTGGACAACATGACA |
| Human | ZMYND8 | Chromatin Readers | TGGACAACATGACAGGGCCT |
| Human | ZMYND8 | Chromatin Readers | AAGCTCTAAGGGATAAAGAC |
| Human | ZMYND8 | Chromatin Readers | TTCAGTTTGGCCCAGACCAA |
| Human | ZMYND8 | Chromatin Readers | GCAATTATTTATTGGAACCC |
| Human | ZMYND8 | Chromatin Readers | GAGACTGACATCGGAACCAG |
| Human | ZMYND8 | Chromatin Readers | GACTGACATCGGAACCAGAG |
| Human | ZMYND8 | Chromatin Readers | GATGGCCTCCTTCTTGCAGT |
| Human | ZMYND8 | Chromatin Readers | AAAGGATGTGGATTGCTACA |
| Human | ZMYND8 | Chromatin Readers | CAGTGCCATGCAAGAGATGG |
| Human | ZMYND8 | Chromatin Readers | TTCAATGGTGAGCATTGTCA |
| Human | ZMYND8 | Chromatin Readers | GCAAGAGATGGAGGTTTACG |
| Human | ZMYND8 | Chromatin Readers | CCTGCGGTACAACATCAACA |
| Human | ZMYND8 | Chromatin Readers | GCCTCCTCCCGAAAGAGTCT |
| Human | ZMYND8 | Chromatin Readers | CCAGCCGAGACTCTTTCGGG |
| Human | ZMYND8 | Chromatin Readers | GCTGGTACTGGTGTTGTGAT |
| Human | ZMYND8 | Chromatin Readers | CGGGAGGAGGCTCTTCGTGC |
| Human | ZMYND8 | Chromatin Readers | AGCAGCAGTGATGAGAAGAG |
| Human | ZMYND8 | Chromatin Readers | TAGCTGAGATTCGCAGGCTG |
| Human | ZMYND8 | Chromatin Readers | GCACAATAGCTGAGATTCGC |
| Human | ZMYND8 | Chromatin Readers | CAACTTAGAGCTGACCATGG |
| Human | ZMYND8 | Chromatin Readers | ACACAACTTAGAGCTGACCA |
| Human | ZMYND8 | Chromatin Readers | GCGGGACCGGCTCATCGCCG |
| Human | ZMYND8 | Chromatin Readers | AGCTGACCATGGCGGAGATG |
| Human | ZMYND8 | Chromatin Readers | CTCCAGCTGCTTCTTCACCT |

| | | | |
|---|---|---|---|
| Human | NFRKB | Chromatin Remodeling | GGCTAGCCTTCCTATGCTAG |
| Human | NFRKB | Chromatin Remodeling | ACCCAGCAGTGGAAGTTGCT |
| Human | NFRKB | Chromatin Remodeling | GTACCAACTCAGCCCAGTTG |
| Human | NFRKB | Chromatin Remodeling | AGAGTGGCTCACGACCCGGG |
| Human | NFRKB | Chromatin Remodeling | TTGTTCACTGCCGATACAGG |
| Human | NFRKB | Chromatin Remodeling | ACACCCGAGCTAGACTTAGC |
| Human | NFRKB | Chromatin Remodeling | GAACCAGCTAAGTCTAGCTC |
| Human | NFRKB | Chromatin Remodeling | AGTCTGGATGCGCACCTGTG |
| Human | NFRKB | Chromatin Remodeling | GCAGCAAGGTCAGGCCACAC |
| Human | NFRKB | Chromatin Remodeling | TTGCAGCAGCTTCAGCAGCA |
| Human | NFRKB | Chromatin Remodeling | GATGGGATGCAGGGACAGTC |
| Human | NFRKB | Chromatin Remodeling | GCAGCTTCAGCAGCAAGGTC |
| Human | NFRKB | Chromatin Remodeling | GTGACTTGAAGCATAACTGC |
| Human | NFRKB | Chromatin Remodeling | CCCAGCAGCAGTATTTCCAT |
| Human | NFRKB | Chromatin Remodeling | CCGATGGAAATACTGCTGCT |
| Human | NFRKB | Chromatin Remodeling | GCCGATGGAAATACTGCTGC |
| Human | NFRKB | Chromatin Remodeling | AATACTGCTGCTGGGAGTTG |
| Human | GLE1 | Control | GACCTTTGTCGGAACTGCGT |
| Human | GLE1 | Control | GTAGCAAAGGCGACCTTTGT |
| Human | GLE1 | Control | ACCTTTGTCGGAACTGCGTA |
| Human | GLE1 | Control | TCCCAGCAGCGACCCTCAGA |
| Human | GLE1 | Control | TCTGTCACACTTAACCCACA |
| Human | GLE1 | Control | CTGTCACACTTAACCCACAG |
| Human | GLE1 | Control | GTCTGTCACACTTAACCCAC |
| Human | GLE1 | Control | ATTTCTCCAAGAACTGCTTG |
| Human | GLE1 | Control | AGGAAGTCAGAAAGCCCTTG |
| Human | GLE1 | Control | TATCACAAGCTCAGGACAGA |
| Human | GLE1 | Control | GAACTGCTTGAGGCGTATGA |
| Human | GLE1 | Control | TGAGAATCTTTGCTCTGTGA |
| Human | GLE1 | Control | GAAGCGGTTTGATGAATGGA |
| Human | GLE1 | Control | TGGCATCTGAACAACTGAAG |
| Human | GLE1 | Control | GCAGCACAAAGCCCTGCTTA |
| Human | GLE1 | Control | GGGACCTCCTGATGAACTTG |
| Human | GLE1 | Control | CCAGAGCCTGCGAAGACAAG |
| Human | GLE1 | Control | GAGCCTGCGAAGACAAGAGG |
| Human | GLE1 | Control | AGCGAGCTCTGCGGGAAATG |
| Human | GLE1 | Control | CTGAGGCTGAGCGAGCTCTG |
| Human | KMT2E | Histone Methylation | GTACACCCAATGCAGAGGTG |
| Human | KMT2E | Histone Methylation | TCAATGATAAGTGCATCAGG |
| Human | KMT2E | Histone Methylation | TTCTATACACAGTATTCCAA |
| Human | KMT2E | Histone Methylation | ATGAGGCTCGATTCATCAGG |
| Human | KMT2E | Histone Methylation | TAGAAATGTGTGTTGATGCA |

| | | | |
|---|---|---|---|
| Human | KMT2E | Histone Methylation | GTTGTGACAAATGCAGCGTT |
| Human | KMT2E | Histone Methylation | TTGGCAACATATTGACTGCA |
| Human | KMT2E | Histone Methylation | GGCAACATATTGACTGCATG |
| Human | KMT2E | Histone Methylation | AGCAGATTGCAGAAAGGAAA |
| Human | KMT2E | Histone Methylation | GAAGAGAACAAGCTTTGGAA |
| Human | KMT2E | Histone Methylation | CAGAGGAGCAGATTGCAGAA |
| Human | KMT2E | Histone Methylation | GTAGCTTGTGAGGGATACGG |
| Human | KMT2E | Histone Methylation | AGAATGAATATGCGACCCAG |
| Human | KMT2E | Histone Methylation | CCAACAAAGTCTGAACAGCA |
| Human | KMT2E | Histone Methylation | GAGAATGAATATGCGACCCA |
| Human | TLR3 | DNA and RNA Modifications | ACAGCATCCCAAAGGGCAAA |
| Human | TLR3 | DNA and RNA Modifications | CATGCACTCTGTTTGCGAAG |
| Human | TLR3 | DNA and RNA Modifications | ACTGGCCAGTTCAGAAAGAA |
| Human | TLR3 | DNA and RNA Modifications | TACTTGCAATTTATGACGAA |
| Human | TLR3 | DNA and RNA Modifications | AATTCATGCCTATAAAGATA |
| Human | CDC73 | Transcription Elongation | ACACTGAAATCTGTAACGGA |
| Human | CDC73 | Transcription Elongation | ACAACGCGGTCCCAGTCTTG |
| Human | CDC73 | Transcription Elongation | GACACTGAAATCTGTAACGG |
| Human | NELFE | Transcription Elongation | CGAAGGCACAGTTTCTGGGT |
| Human | NELFE | Transcription Elongation | GGTCCCGTTGAGCTCAGCAA |
| Human | NELFE | Transcription Elongation | CAGGCCGTTGCTGAGCTCAA |
| Human | NELFE | Transcription Elongation | ACGAAGGCACAGTTTCTGGG |
| Human | NELFE | Transcription Elongation | ACCGGGACAGAGACCGAGAT |
| Human | NELFE | Transcription Elongation | CCGGGACAGAGACCGAGATC |
| Human | NELFE | Transcription Elongation | TCGAGACCGGGAACGGGACA |
| Human | NELFE | Transcription Elongation | CCGCAGCCATGAGAGGAACC |
| Human | NELFE | Transcription Elongation | CTCTGTCTCTGTCTCGATCC |
| Human | LDB1 | Transcription Elongation | AGGACGAGAGGCTCATCACC |
| Human | LDB1 | Transcription Elongation | ACCCAGTTTGACGCAGCCAA |
| Human | LDB1 | Transcription Elongation | GCCGTTGGCTGCGTCAAACT |
| Human | TDRD6 | Chromatin Readers | GGACGAGGGCCGCACCATCA |
| Human | TDRD6 | Chromatin Readers | GAGGGCCGCACCATCACGGC |
| Human | TDRD6 | Chromatin Readers | CCTCGTCCAGCAGGAAGACA |
| Human | TDRD6 | Chromatin Readers | CAGGCACAGCTCGCCGGGCG |
| Human | TDRD6 | Chromatin Readers | TCATGTGGACTATGGAAGGA |
| Human | TDRD6 | Chromatin Readers | ATTCAGGCAGCAAGTACCGA |
| Human | TDRD6 | Chromatin Readers | ATAGTCCACATGAAGCACCT |
| Human | TDRD6 | Chromatin Readers | CAGGTGCTTCATGTGGACTA |
| Human | TDRD6 | Chromatin Readers | CCAATGTCCATCCAGGCCAC |
| Human | TDRD6 | Chromatin Readers | ATGTGGACTGGTATGACGTA |
| Human | TDRD6 | Chromatin Readers | CTTTCCACTTGACACAGCAA |
| Human | TDRD6 | Chromatin Readers | CAAATTGGATGACAAGAGTG |

| | | | |
|---|---|---|---|
| Human | TDRD6 | Chromatin Readers | TTCGCTTAGCCAAACAAGCA |
| Human | TDRD6 | Chromatin Readers | GGTCCAGAGCACTTATTAGT |
| Human | TDRD6 | Chromatin Readers | TCGCTTAGCCAAACAAGCAA |
| Human | TDRD6 | Chromatin Readers | TATTAGTGGGATACAGTCTG |
| Human | TDRD6 | Chromatin Readers | TGCCTTGCCAAGTATACTGA |
| Human | TDRD6 | Chromatin Readers | TTCCATCAGTATACTTGGCA |
| Human | TDRD6 | Chromatin Readers | AGTATACTTGGCAAGGCACA |
| Human | TDRD6 | Chromatin Readers | ACTGATGGAAACTGGTATAG |
| Human | TDRD6 | Chromatin Readers | CCAGTTTCCATCAGTATACT |
| Human | TDRD6 | Chromatin Readers | CTGCACAGAGAGAAGGTCAT |
| Human | TDRD6 | Chromatin Readers | GATACCATAAATTATCTTCT |
| Human | TDRD6 | Chromatin Readers | ATGGTATCGTGCTGTGATCA |
| Human | TDRD6 | Chromatin Readers | TCTCTGTATCTTACTATACA |
| Human | TDRD6 | Chromatin Readers | ATACAAGGATCTCCAATATA |
| Human | TDRD6 | Chromatin Readers | TCTGTCAGGCTTGTGGACTT |
| Human | TDRD6 | Chromatin Readers | GGTCAAGTCTAAGAAACACA |
| Human | TDRD6 | Chromatin Readers | TCTCCTCCATACCATTTGAA |
| Human | TDRD6 | Chromatin Readers | AACCCTGAGAATGTCTGGAA |
| Human | TDRD6 | Chromatin Readers | GAACCTTTCAAATGGTATGG |
| Human | CHD8 | Chromatin Readers | CTTCAGTATATTGTCCAGAA |
| Human | CHD8 | Chromatin Readers | TAGACAAAGTGCTTTCTATG |
| Human | CHD8 | Chromatin Readers | TACTATCTCCCAACTAGAGA |
| Human | CHD8 | Chromatin Readers | GTGAAGAAGGAGCTCCCTTC |
| Human | CHD8 | Chromatin Readers | AGCCCATTCACAATGCAGAT |
| Human | CHD8 | Chromatin Readers | GGAGCCCGTTATTTACTACC |
| Human | CHD8 | Chromatin Readers | TGAGTCTCACAGTATTGACA |
| Human | CHD8 | Chromatin Readers | AATTTAAACGGATTCAGTCA |
| Human | CHD8 | Chromatin Readers | GCCCTATGAGGATAGCACAT |
| Human | CHD8 | Chromatin Readers | ATGGTGCTCTCTGCCCTATG |
| Human | CHD8 | Chromatin Readers | ATGTACTGCAAAGATTCACG |
| Human | CHD8 | Chromatin Readers | GCACATGGACCTGGAACACA |
| Human | CHD8 | Chromatin Readers | CAGTTAGTAATTGTGGACAG |
| Human | CHD8 | Chromatin Readers | GGGAGCGAGAATTTAATACA |
| Human | CHD8 | Chromatin Readers | CGTCAAACTTGTATGCGCCT |
| Human | CHD8 | Chromatin Readers | AGGAATTACGAGTGATGAGG |
| Human | CHD8 | Chromatin Readers | TGTCTTCTTACTGTGTACCC |
| Human | CHD8 | Chromatin Readers | AGGATTATTTAATCCAGAGG |
| Human | CHD8 | Chromatin Readers | AGGCTTGCTGAAGCGGTCAA |
| Human | CHD8 | Chromatin Readers | AGAAACAACAACGATGGACA |
| Human | CHD8 | Chromatin Readers | GGCGGGAGAAACAACAACGA |
| Human | CHD8 | Chromatin Readers | ACGATCCTTGGGCCACTCAG |
| Human | CHD8 | Chromatin Readers | CTCAAGCCAAGAATGCCCAA |

| | | | |
|---|---|---|---|
| Human | CHD8 | Chromatin Readers | CAGCTGGATAGTTACTACCT |
| Human | CHD8 | Chromatin Readers | CCAGGCAAGATAGTGTTACA |
| Human | CHD8 | Chromatin Readers | GTTGTGCTAGGGAGTCTACC |
| Human | CHD8 | Chromatin Readers | AAGACTTTCAGTACCAGTCA |
| Human | CHD8 | Chromatin Readers | AGGGCTATGATAGCTCAGAA |
| Human | CHD8 | Chromatin Readers | GATGATGATGACTTATCTCA |
| Human | CHD8 | Chromatin Readers | TCTGAGTTAGCTGGCATCAT |
| Human | CHD8 | Chromatin Readers | AGATGATGATGACTTATCTC |
| Human | CHD8 | Chromatin Readers | TCACTGGAGTCTGAGTTAGC |
| Human | HDGF | Chromatin Readers | GGCATCTCGTCAATCCGGGC |
| Human | HDGF | Chromatin Readers | GCCCAGGAATGCCGTCTCGT |
| Human | HDGF | Chromatin Readers | CAATCCGGGCCGGCCAGTGT |
| Human | HDGF | Chromatin Readers | GGACTCTCCTAAACGTCCCA |
| Human | HDGF | Chromatin Readers | GGTCATTGATGAGCCAGCCA |
| Human | CSNK2A2 | Other Histone Modifications | AAGCTGGTGTCTTTGACACG |
| Human | CSNK2A2 | Other Histone Modifications | CATACATCTGATAGTCCACG |
| Human | CSNK2A2 | Other Histone Modifications | CAAAGCTGGTGTCTTTGACA |
| Human | CSNK2A2 | Other Histone Modifications | ACAACATTCACGGAAACGCT |
| Human | AFG3L2 | Control | CACCGCTGTTTGCGGCTGTG |
| Human | AFG3L2 | Control | AGGAGCTGCTGTAGGCCGCG |
| Human | AFG3L2 | Control | CAGCAGCTCCTCGTGCCTGG |
| Human | AFG3L2 | Control | GGCCCACGCCGCCAGGCACG |
| Human | AFG3L2 | Control | GATTTGTTACAACTCAAGCA |
| Human | AFG3L2 | Control | TTGCTTGAGTTGTAACAAAT |
| Human | AFG3L2 | Control | CCCAGAACAGAGCAGTCCAG |
| Human | AFG3L2 | Control | TCCTCTGGACTGCTCTGTTC |
| Human | AFG3L2 | Control | CTCTGGACTGCTCTGTTCTG |
| Human | AFG3L2 | Control | CCTCTGGACTGCTCTGTTCT |
| Human | AFG3L2 | Control | AAGGCGATGATGAGCACCGT |
| Human | AFG3L2 | Control | TCTGCTGAGCATGCTGCCTA |
| Human | AFG3L2 | Control | AGTTTCCAGATTCCGTTCAA |
| Human | MBD3 | DNA and RNA Modifications | AAGAAGTGCCCAGAAGGTCG |
| Human | MBD3 | DNA and RNA Modifications | AGAGGGAAGAAGTGCCCAGA |
| Human | MBD3 | DNA and RNA Modifications | CTGCTTCCTGATGTCCTCGT |
| Human | MBD3 | DNA and RNA Modifications | GTCCTCGTCGGTCACCATGA |
| Human | MBD3 | DNA and RNA Modifications | CGACGAGGACATCAGGAAGC |
| Human | MBD3 | DNA and RNA Modifications | TGGTGACCGACGAGGACATC |
| Human | MBD3 | DNA and RNA Modifications | GCAGGTGCGGAAGCGGCTGG |
| Human | ERCC6 | DNA and RNA Modifications | GTTCAAATTACAGGTTTGAG |
| Human | ERCC6 | DNA and RNA Modifications | GCGAAACTGTTTGCAAGCAA |
| Human | ERCC6 | DNA and RNA Modifications | ACTGCTTGAAGATTTGTCGG |
| Human | ERCC6 | DNA and RNA Modifications | TCATTGTATCTCGTAATCAG |

| | | | |
|---|---|---|---|
| Human | ERCC6 | DNA and RNA Modifications | GCTTAGGTGTCAACCTGACG |
| Human | ERCC6 | DNA and RNA Modifications | AGCGTTTAGAAAGTGAAAGC |
| Human | ERCC6 | DNA and RNA Modifications | GCAGGGCAGAAGCTTCCTGC |
| Human | ERCC6 | DNA and RNA Modifications | GAGCGTTTAGAAAGTGAAAG |
| Human | ERCC6 | DNA and RNA Modifications | CTTTCACTTTCTAAACGCTC |
| Human | ERCC6 | DNA and RNA Modifications | AAGTGAAAGCGGGCACCTGC |
| Human | ERCC6 | DNA and RNA Modifications | ATGGAATAAACTGAGACTGC |
| Human | ERCC6 | DNA and RNA Modifications | AACATCATGGTCTGCTCCAA |
| Human | ERCC6 | DNA and RNA Modifications | TCATGGTCTGCTCCAAAGGC |
| Human | ERCC6 | DNA and RNA Modifications | GAGGCAGAGGGAGACTCTGA |
| Human | ERCC6 | DNA and RNA Modifications | AGAGGCAGAGGGAGACTCTG |
| Human | ERCC6 | DNA and RNA Modifications | GGAAGAGGAGGAAGATGACG |
| Human | ERCC6 | DNA and RNA Modifications | GGGATGTCGTACATGACGTG |
| Human | ERCC6 | DNA and RNA Modifications | CGTGGAGAAGGAGTATCGGT |
| Human | ERCC6 | DNA and RNA Modifications | GGTGGACAATGCCATCCATG |
| Human | ERCC6 | DNA and RNA Modifications | CGTCATGTACGACATCCCTA |
| Human | ERCC6 | DNA and RNA Modifications | TACATGACGTGAGGTCATCC |
| Human | FNTB | Control | GCAAGGTGTGGATACTGACC |
| Human | FNTB | Control | CTGTGTCAGAGCCCAGAAGG |
| Human | FNTB | Control | GCATATGTGGGTGCAAGGTG |
| Human | FNTB | Control | GAGCTGTGTCAGAGCCCAGA |
| Human | FNTB | Control | CAGAGCCCAGAAGGTGGCTT |
| Human | FNTB | Control | GGCTCCTTTCTCATGCATGT |
| Human | FNTB | Control | TCTCATGCATGTCGGAGGTG |
| Human | FNTB | Control | TCCTTTCTCATGCATGTCGG |
| Human | FNTB | Control | TACTCCCTGAAGCAACCTGA |
| Human | FNTB | Control | GGAGCCGTCAGGTTGCTTCA |
| Human | FNTB | Control | ACAGAAGGTATAGCCACCAT |
| Human | FNTB | Control | CCAGGGATGGAAGCCCATGG |
| Human | FNTB | Control | GATAGCAAGGTGTCAGAACT |
| Human | FNTB | Control | CACAGAAGGTATAGCCACCA |
| Human | FNTB | Control | AGGTGTCAGAACTGGGAAGG |
| Human | FNTB | Control | TATTACAATGGGTGACAAGC |
| Human | FNTB | Control | CTCCTTCAAATCGCATCTGC |
| Human | FNTB | Control | CATCCACCAGCTTGTTGCAG |
| Human | FNTB | Control | CGCTGCAACAAGCTGGTGGA |
| Human | FNTB | Control | GGGCCGCTGCAACAAGCTGG |
| Human | FNTB | Control | CACACCTGCTACTGCCTGAG |
| Human | FNTB | Control | CATCAGGATGTACTCCTGCA |
| Human | FNTB | Control | TAGAAATCACGCGACTTGCC |
| Human | FNTB | Control | AGGATCCAATAGCAGAGCCA |
| Human | FNTB | Control | TACAGATGTGTGTCAGTTCC |

| | | | |
|---|---|---|---|
| Human | AIRE | Chromatin Readers | TCTCCTGAAACTTGTCCTCG |
| Human | AIRE | Chromatin Readers | CGCGCTGGCTGACCACGACG |
| Human | AIRE | Chromatin Readers | GCACCGCACGGAGATCGCGG |
| Human | AIRE | Chromatin Readers | ACAACCTGGAGCGCTATGGC |
| Human | AIRE | Chromatin Readers | CGTCTCCTGAAACTTGTCCT |
| Human | AIRE | Chromatin Readers | CAGCAGGTGTTTGAGTCAGG |
| Human | AIRE | Chromatin Readers | GCCTAGCCTCACCTCCACCG |
| Human | AIRE | Chromatin Readers | CCTAGCCTCACCTCCACCGG |
| Human | AIRE | Chromatin Readers | AGTTCGAAGACTCCGGCAGT |
| Human | AIRE | Chromatin Readers | AGCACCTCCAGGTCCCACTG |
| Human | AIRE | Chromatin Readers | CAGGCCCGTCCCGGGCCGGG |
| Human | AIRE | Chromatin Readers | TGGCGCTGCCACTTCCCAGC |
| Human | AIRE | Chromatin Readers | GCAGCGCCAGTGGAAGGCAG |
| Human | AIRE | Chromatin Readers | GAAGTGGCAGCGCCAGTGGA |
| Human | THOC1 | Transcription Elongation | GGTTGCCTGGCAAGATCAAG |
| Human | THOC1 | Transcription Elongation | GTTGCCTGGCAAGATCAAGA |
| Human | THOC1 | Transcription Elongation | AACTCCCTCTTGATCTTGCC |
| Human | THOC1 | Transcription Elongation | CCAACAAGCTGGGTGAACAA |
| Human | THOC1 | Transcription Elongation | TGAGTCTTTCATTTCCAAGT |
| Human | THOC1 | Transcription Elongation | CCCGAGGACTTCCTAGGGAA |
| Human | THOC1 | Transcription Elongation | CAGCACCCGAGGACTTCCTA |
| Human | THOC1 | Transcription Elongation | CCTTTCCCTAGGAAGTCCTC |
| Human | THOC1 | Transcription Elongation | ACAGCACCCGAGGACTTCCT |
| Human | THOC1 | Transcription Elongation | TCCTTTCCCTAGGAAGTCCT |
| Human | TSFM | Transcription Elongation | TGACGCAGAAGAGACCCAGC |
| Human | TSFM | Transcription Elongation | GACGCAGAAGAGACCCAGCC |
| Human | TSFM | Transcription Elongation | GGACCGGGAGCTACCCGGCT |
| Human | TSFM | Transcription Elongation | CGGACCGGGAGCTACCCGGC |
| Human | TSFM | Transcription Elongation | AGAGATCTGGCTCCACAAGG |
| Human | TSFM | Transcription Elongation | GGCAGAGATCTGGCTCCACA |
| Human | JADE3 | Chromatin Readers | TGTGTGCATCAGGCCTGCTA |
| Human | JADE3 | Chromatin Readers | GGTCTCCAGACAGTGAAGAA |
| Human | JADE3 | Chromatin Readers | CTTCTTCACTGTCTGGAGAC |
| Human | JADE3 | Chromatin Readers | GACCTTGAGGATGCCATAGC |
| Human | JADE3 | Chromatin Readers | GTGTAACGTCTGTGTGCATC |
| Human | JADE3 | Chromatin Readers | TTTGGACATAACACACATTG |
| Human | JADE3 | Chromatin Readers | GTGTTATGTCCAAAGAAAGG |
| Human | JADE3 | Chromatin Readers | TGTGTGTTATGTCCAAAGAA |
| Human | JADE3 | Chromatin Readers | CACCAAGACAGGGACTAAAT |
| Human | JADE3 | Chromatin Readers | AGCCCATTTAGTCCCTGTCT |
| Human | JADE3 | Chromatin Readers | ATGAAGACCATCCTAGATGA |
| Human | JADE3 | Chromatin Readers | GATGAAGACCATCCTAGATG |

| | | | |
|---|---|---|---|
| Human | JADE3 | Chromatin Readers | TAGGCCGTGCTCAAAGGCAC |
| Human | JADE3 | Chromatin Readers | GTCACCTGTGCCTTTGAGCA |
| Human | JADE3 | Chromatin Readers | CTTGTGCAAGTTGAAGACGG |
| Human | MCM7 | DNA and RNA Modifications | CCGAGACAATGACCTACGGT |
| Human | MCM7 | DNA and RNA Modifications | CCTGTTGTGTACTGGCTGCG |
| Human | MCM7 | DNA and RNA Modifications | TTCATGCCCGTATATTTCTG |
| Human | MCM7 | DNA and RNA Modifications | TGATGTGCTGGGCCAACCGT |
| Human | MCM7 | DNA and RNA Modifications | CATGTCCAGAGGTTCAAACT |
| Human | MCM7 | DNA and RNA Modifications | CTCATGAGCTTCATGTCCAG |
| Human | MCM7 | DNA and RNA Modifications | AGAGGTTCAAACTGGGAGGG |
| Human | MCM7 | DNA and RNA Modifications | CCTATGTGCACCAGCACAGC |
| Human | MCM7 | DNA and RNA Modifications | TGAGACCAGTTCACGGACGG |
| Human | MCM7 | DNA and RNA Modifications | CTACATCACAGCAGCATACG |
| Human | MCM7 | DNA and RNA Modifications | GACGTGCCAGAGCAGTGGAA |
| Human | MCM7 | DNA and RNA Modifications | GTCCGTGAACTGGTCTCAGG |
| Human | MCM7 | DNA and RNA Modifications | GAGCAGCGCTGTGTATCTCG |
| Human | MCM4 | DNA and RNA Modifications | GATGATCCCAGCCTTTGCAA |
| Human | GSG2 | Other Histone Modifications | GAGGACCTGTTTACCGGTGA |
| Human | GSG2 | Other Histone Modifications | ATCCCGTTCCAAGCGCGACA |
| Human | GSG2 | Other Histone Modifications | TGAGCACCGAGACTTACACT |
| Human | GSG2 | Other Histone Modifications | TGAAGTTCAGCATTGTCCTG |
| Human | GSG2 | Other Histone Modifications | GATCACTATAATTCAACCAA |
| Human | GSG2 | Other Histone Modifications | AAAGGCATCTGATGCTGAAA |
| Human | KDM6B | Histone Methylation | TTTCTGTGATCGGCACGGCG |
| Human | KDM6B | Histone Methylation | AGCGCTTTCTGTGATCGGCA |
| Human | KDM6B | Histone Methylation | TGGTACTGATAGGCGGTGAG |
| Human | KDM6B | Histone Methylation | CAAACCATATTATGCTCCAG |
| Human | KDM6B | Histone Methylation | CCAGAACTACTGCCATGTGA |
| Human | KDM6B | Histone Methylation | GAGCATAATATGGTTTGCTG |
| Human | KDM6B | Histone Methylation | CTGGCTGAGCATCTGCCTCC |
| Human | KDM6B | Histone Methylation | GCTGGACGAATCCATTCGCA |
| Human | KDM6B | Histone Methylation | TCTCTAAGGTCACTTCCGGG |
| Human | KDM6B | Histone Methylation | CCAGGGACAGAGGCTGACGG |
| Human | KDM6B | Histone Methylation | TGTCCCTGGAAAGAAGGCTC |
| Human | KDM6B | Histone Methylation | CCCTGGAAAGAAGGCTCGGG |
| Human | KDM6B | Histone Methylation | CTGTCCCTGGAAAGAAGGCT |
| Human | KDM6B | Histone Methylation | CCTCCCGAGCCTTCTTTCCA |
| Human | KDM6B | Histone Methylation | GGCCTGGTACTGTGCGTACT |
| Human | KDM6B | Histone Methylation | CCAGGTGCAACGCGAGAGCC |
| Human | KDM6B | Histone Methylation | CCAGGCTCTCGCGTTGCACC |
| Human | KDM6B | Histone Methylation | AGTGCTTCATGGACTGCAGC |
| Human | KDM6B | Histone Methylation | GCAGTCCATGAAGCACTGCC |

| | | | |
|---|---|---|---|
| Human | KDM6B | Histone Methylation | TTGCACCTGGCAGTGCTTCA |
| Human | GTF2H1 | Transcription Elongation | TGCATTCACATTTAAACGAT |
| Human | GTF2H1 | Transcription Elongation | GAGTCAAGTGATCAGTGCTG |
| Human | GTF2H1 | Transcription Elongation | TGATCACTTGACTCACAACA |
| Human | GTF2H1 | Transcription Elongation | GATCAGTGCTGAGGAATTCT |
| Human | GTF2H1 | Transcription Elongation | TGATCAGTGCTGAGGAATTC |
| Human | GTF2H1 | Transcription Elongation | TCCTTCTCTGTCATGTTGTG |
| Human | GTF2H1 | Transcription Elongation | ACATGACAGAGAAGGAATTC |
| Human | USP15 | Other Histone Modifications | GCGACTATCGACTAGGTACC |
| Human | USP15 | Other Histone Modifications | GTTGGAATAAACTTGTCAGC |
| Human | USP15 | Other Histone Modifications | GGTACCTAGTCGATAGTCGC |
| Human | USP15 | Other Histone Modifications | AGAGCCTTTAAGACACAGGT |
| Human | USP15 | Other Histone Modifications | CAAACCTATGCAGTACAAAG |
| Human | USP15 | Other Histone Modifications | TTACAATTCGGACAATACCA |
| Human | USP15 | Other Histone Modifications | CTGTGTCTTAAAGGCTCTTG |
| Human | USP15 | Other Histone Modifications | AGAGCCAATAGCACGAAAGG |
| Human | USP15 | Other Histone Modifications | GCACGAAAGGTGGTTGAACA |
| Human | USP15 | Other Histone Modifications | CTATCACTAATACCTGTCCT |
| Human | USP15 | Other Histone Modifications | GTATAAACCAGCATCCTGAA |
| Human | USP15 | Other Histone Modifications | TCAAGAGCCAATAGCACGAA |
| Human | PHF21A | Chromatin Readers | AGCAGTCCAAATGATATACA |
| Human | PHF21A | Chromatin Readers | GCAGTCCAAATGATATACAC |
| Human | PHF21A | Chromatin Readers | AAACAATTCCCAAGGGCATG |
| Human | PHF21A | Chromatin Readers | GGACAGATCCACATGCCCTT |
| Human | PHF21A | Chromatin Readers | TCAGGACCAGATGCTGAAGA |
| Human | PHF21A | Chromatin Readers | CAATTCCATAAGTAAATGCA |
| Human | PHF21A | Chromatin Readers | TTCTTTAAGTCAGCATTCTG |
| Human | PHF21A | Chromatin Readers | ACTGGTTGCTCAAATGAAGC |
| Human | PHF21A | Chromatin Readers | AAAGAGTAGTTGAACAGCTA |
| Human | PHF21A | Chromatin Readers | AATTCAGGTTCACCAGAAAC |
| Human | PARP10 | Other Histone Modifications | CGCGCTCCACACGAACGACG |
| Human | PARP10 | Other Histone Modifications | CAGCATCCGCGTCGTTCGTG |
| Human | PARP10 | Other Histone Modifications | GATGGTGCAGTCACCACGCA |
| Human | PARP10 | Other Histone Modifications | CTTCAGGCCCTACCTTGGCG |
| Human | PARP10 | Other Histone Modifications | GGTGTGAGTGTTGCCCTGCG |
| Human | PARP10 | Other Histone Modifications | CCAGAAGTGCCACCAAGTGG |
| Human | PARP10 | Other Histone Modifications | AAGGCCAAACTCTGATCCCA |
| Human | PARP10 | Other Histone Modifications | GCTGGCCCTCCACCGGTCAC |
| Human | PARP10 | Other Histone Modifications | CTCTGCAGCTGGCCCTCCAC |
| Human | PARP10 | Other Histone Modifications | GTGGAGGGCCAGCTGCAGAG |
| Human | PARP10 | Other Histone Modifications | GGAGGAGGCCGCTCTGCAGC |
| Human | PARP10 | Other Histone Modifications | GAGGGTTAGGGCTTGCCGCA |

| | | | |
|---|---|---|---|
| Human | PARP10 | Other Histone Modifications | AGAGGGTTAGGGCTTGCCGC |
| Human | PARP10 | Other Histone Modifications | GGAGGGCCTAGACCTAGACG |
| Human | PARP10 | Other Histone Modifications | AGGCTCCAGTGACCGGTGGA |
| Human | PARP10 | Other Histone Modifications | CTTGAGGCTCCAGTGACCGG |
| Human | PARP10 | Other Histone Modifications | TAGGTCTAGGCCCTCCAGGG |
| Human | POLR2B | Transcription Elongation | TAATGGCGATTGCCAACACC |
| Human | POLR2B | Transcription Elongation | GGACCCATACATATGAATGC |
| Human | POLR2B | Transcription Elongation | GCATTCATATGTATGGGTCC |
| Human | POLR2B | Transcription Elongation | ATATGTATGGGTCCTGGTGT |
| Human | ELOA3 | Transcription Elongation | GCCAGGATGTCTGCGGTCAT |
| Human | ELOA3 | Transcription Elongation | GATGTCTGCGGTCATGGGCA |
| Human | ELOA3 | Transcription Elongation | ATCTGTCTCTCGAGCGAGTG |
| Human | ELOA3 | Transcription Elongation | GAGAGACAGATGAATTATGG |
| Human | ELOA3 | Transcription Elongation | GCGTTGGAGGGTCCCGATGG |
| Human | ELOA3 | Transcription Elongation | CGCGTTGGAGGGTCCCGATG |
| Human | PHRF1 | Chromatin Readers | GCCTGGACTGCATTGTCGAA |
| Human | PHRF1 | Chromatin Readers | TCTCAACGCATTCAGAGACC |
| Human | PHRF1 | Chromatin Readers | CATTCAGAGACCAGGCCGTG |
| Human | PHRF1 | Chromatin Readers | ACCATTCGACAATGCAGTCC |
| Human | PHRF1 | Chromatin Readers | GCAGTCCAGGCAGAAGTAAT |
| Human | PHRF1 | Chromatin Readers | GTGCGGCAGGAGCGACCGTG |
| Human | PHRF1 | Chromatin Readers | GGGCAGAACCACTCGTCCAC |
| Human | PHRF1 | Chromatin Readers | CATGCAGGAGCGTGCTGTGG |
| Human | PHRF1 | Chromatin Readers | CATGAAGAAGCTGCACATGC |
| Human | PHRF1 | Chromatin Readers | GCACATGCAGGAGCGTGCTG |
| Human | PHRF1 | Chromatin Readers | GGAGGGTCCAGCACACACCA |
| Human | PHRF1 | Chromatin Readers | CCGCGCGATTCGAGAGCGAG |
| Human | PHRF1 | Chromatin Readers | GGAAGAGGAAGACAAGAAGA |
| Human | PHRF1 | Chromatin Readers | GAACCGGAACCGGATCTCCA |
| Human | PHRF1 | Chromatin Readers | AGGTCAGCGTCACCATCAGT |
| Human | PHRF1 | Chromatin Readers | GCAGCCGAGAGCGAGCTAAG |
| Human | PHRF1 | Chromatin Readers | TCTGCAGGCACAAGCATCAG |
| Human | POLR2H | Transcription Elongation | CACAATGCAGTCGAGACACT |
| Human | POLR2H | Transcription Elongation | TGAGAGTGAATCTTTCAAGA |
| Human | POLR2H | Transcription Elongation | CGGTCAAACTTCTTGCCCTC |
| Human | ACTR8 | Chromatin Remodeling | TGGCAGATATTGAAGTAATA |
| Human | SMARCA2 | Chromatin Readers | CTGCAAGCTGCAGCGTTTCG |
| Human | SMARCA2 | Chromatin Readers | ACTGCAAGCTGCAGCGTTTC |
| Human | SMARCA2 | Chromatin Readers | GAATATCATCGGTCTGTGGC |
| Human | SMARCA2 | Chromatin Readers | TAAGGAATATCATCGGTCTG |
| Human | SMARCA2 | Chromatin Readers | GGTATTCCTGGTGTTTCTGA |
| Human | SMARCA2 | Chromatin Readers | AAATACTGTTCAGGTATTCC |

| | | | |
|---|---|---|---|
| Human | SMARCA2 | Chromatin Readers | CAGTTAGATAGAGTCGAAAG |
| Human | SMARCA2 | Chromatin Readers | ACATTCTTGCAAAGATTCGG |
| Human | SMARCA2 | Chromatin Readers | TCCAGTTAGATAGAGTCGAA |
| Human | SMARCA2 | Chromatin Readers | AAATACATGATAGTGGACGA |
| Human | SMARCA2 | Chromatin Readers | TGGTGGTGCCATCAAGGCGT |
| Human | SMARCA2 | Chromatin Readers | TGGGCCTGCAGATCCTGATG |
| Human | SMARCA2 | Chromatin Readers | AATCATAAGTACCGGAGCCT |
| Human | SMARCA2 | Chromatin Readers | GGATTCGTAATCATAAGTAC |
| Human | SMARCA2 | Chromatin Readers | CGCTCAGACGTTCAACCTGG |
| Human | SMARCA2 | Chromatin Readers | TGAATTAATTAGGAAGCCAG |
| Human | SMARCA2 | Chromatin Readers | GCTCAGACGTTCAACCTGGA |
| Human | SMARCA2 | Chromatin Readers | AAACACTGACTGTAAGACGA |
| Human | SMARCA2 | Chromatin Readers | AGTCAGTGTTTAAGAGTGCC |
| Human | SMARCA4 | Chromatin Readers | GCCTACAAGATGCTGGCCAG |
| Human | SMARCA4 | Chromatin Readers | GATCATGGCCTACAAGATGC |
| Human | SMARCA4 | Chromatin Readers | CCAGCTCAGAGCTCAGATCA |
| Human | SMARCA4 | Chromatin Readers | CCTACAAGATGCTGGCCAGG |
| Human | SMARCA4 | Chromatin Readers | CGTACCATGCCAACACGGAG |
| Human | SMARCA4 | Chromatin Readers | GGCCACGTACCATGCCAACA |
| Human | SMARCA4 | Chromatin Readers | GGATCGAGAAGGAGCGCATG |
| Human | SMARCA4 | Chromatin Readers | GAATATCACAGATCCGTCAC |
| Human | SMARCA4 | Chromatin Readers | CAGTTGGACAGCGTTGAGAG |
| Human | SMARCA4 | Chromatin Readers | ACTTCCAACGGATCTTGGCG |
| Human | SMARCA4 | Chromatin Readers | GGGCGTACGAGTTTGACAAG |
| Human | SMARCA4 | Chromatin Readers | CTTGTAAGACACCTTCACCA |
| Human | SMARCA4 | Chromatin Readers | ACATCCTCGCCAAGATCCGT |
| Human | SMARCA4 | Chromatin Readers | CAGGCTTGATGGAACCACGA |
| Human | SMARCA4 | Chromatin Readers | GCTTGATGGAACCACGAAGG |
| Human | SMARCA4 | Chromatin Readers | TCGTGGTTCCATCAAGCCTG |
| Human | SMARCA4 | Chromatin Readers | GAAGATTACTTTGCGTATCG |
| Human | SMARCA4 | Chromatin Readers | TGATGGAACCACGAAGGCGG |
| Human | SMARCA4 | Chromatin Readers | TCGGGCAGCTCCTTTCGCGA |
| Human | SMARCA4 | Chromatin Readers | CGTCCTTCTCTAGGTCGTTG |
| Human | SMARCA4 | Chromatin Readers | CAGCCTCAACGACCTAGAGA |
| Human | SMARCA4 | Chromatin Readers | CATCCAGCTGCCCTCGCGAA |
| Human | SMARCA4 | Chromatin Readers | CGTTGAGGCTGCGGTACTTG |
| Human | SMARCA4 | Chromatin Readers | TTGGGCCCTCCCATACCATG |
| Human | SMARCA4 | Chromatin Readers | GCAGGCCTCATGGTATGGGA |
| Human | SMARCA4 | Chromatin Readers | GGCCGAGGAGTTCCGCCCAG |
| Human | SMARCA4 | Chromatin Readers | AGCAGGCCTCATGGTATGGG |
| Human | SMARCA4 | Chromatin Readers | ATGGAGTCCATGCATGAGAA |
| Human | SMARCA4 | Chromatin Readers | CTCGGAGCTACTTCATACCC |

| | | | |
|---|---|---|---|
| Human | SMARCA4 | Chromatin Readers | TCATGGCTGAAGATGAGGAG |
| Human | SMARCA4 | Chromatin Readers | GGAGGAGCCACCTCTAAAGG |
| Human | SMARCA4 | Chromatin Readers | ACCTCTAAAGGAGGAAGACG |
| Human | SMARCA4 | Chromatin Readers | GTCGTCCTTGATGATCCACG |
| Human | SMARCA4 | Chromatin Readers | GCAGTGGCTCAAGGCCATCG |
| Human | SMARCA4 | Chromatin Readers | GAGAGTGAGGAGGAGGAAGA |
| Human | SMARCA4 | Chromatin Readers | GGAGAGTGAGGAGGAGGAAG |
| Human | SMARCA4 | Chromatin Readers | CAAGAAGATGAAGAAGATTG |
| Human | SMARCA4 | Chromatin Readers | TCTTCTTCATCTTCTTGGTG |
| Human | SMARCA4 | Chromatin Readers | CACAATCTTCTTCATCTTCT |
| Human | SMARCA4 | Chromatin Readers | AGTCGGTCTTCACCAGCGTG |
| Human | SMARCA4 | Chromatin Readers | AGACTCCATCGTCTTGCAGT |
| Human | SMARCA4 | Chromatin Readers | GAAGACCGACTGCAAGACGA |
| Human | HDAC1 | Histone Acetylation | CAGATGCAGAGATTCAACGT |
| Human | HDAC1 | Histone Acetylation | AGTCTGTTACTACTACGACG |
| Human | HDAC1 | Histone Acetylation | AACTGCTAAAGTATCACCAG |
| Human | HDAC1 | Histone Acetylation | ACTGCTAAAGTATCACCAGA |
| Human | HDAC1 | Histone Acetylation | GTCGCTGTTTGATCTTCTCC |
| Human | CDK9 | Transcription Elongation | CACGTTTGGCCACACCTCAG |
| Human | CDK9 | Transcription Elongation | CAGAAACAAGATCCTGCATA |
| Human | CDK9 | Transcription Elongation | GATCGGCCAAGGCACCTTCG |
| Human | CDK9 | Transcription Elongation | CCCTGAGGTGTGGCCAAACG |
| Human | ING2 | Chromatin Readers | TCACTTACCTATAAACCAAA |
| Human | ING2 | Chromatin Readers | TGTTACACAAATGCTCGAAT |
| Human | ING2 | Chromatin Readers | CAAATATCAAGAAACGTTAA |
| Human | ING2 | Chromatin Readers | TACACAAATGCTCGAATTGG |
| Human | ING2 | Chromatin Readers | CTAATTAATAGTCAAGAATT |
| Human | CSNK2A1 | Other Histone Modifications | TCAGTTGGTGAGGATAGCCA |
| Human | CSNK2A1 | Other Histone Modifications | ACAATTATGATCAGTTGGTG |
| Human | CSNK2A1 | Other Histone Modifications | CCCAAGATATCATTGAAACG |
| Human | PHF12 | Chromatin Readers | TGTTAAACTACAGTGAGCAT |
| Human | PHF12 | Chromatin Readers | CAGTGAGCATGGGACAACGG |
| Human | PHF12 | Chromatin Readers | CATCGGGACAGGAGCTGACA |
| Human | PHF12 | Chromatin Readers | GGTCACTGTAACTACGTGTC |
| Human | PHF12 | Chromatin Readers | GTCACTGTAACTACGTGTCC |
| Human | PHF12 | Chromatin Readers | GTGGAGGGTTACAGCACTGG |
| Human | PHF12 | Chromatin Readers | TCAGTGGAGGGTTACAGCAC |
| Human | PHF12 | Chromatin Readers | TGCGATAGCTGCAAGGAAGG |
| Human | PHF12 | Chromatin Readers | TGACACATCCACTCTCCAGG |
| Human | PHF12 | Chromatin Readers | GGCAACATTTCTTCACTCAG |
| Human | PHF12 | Chromatin Readers | GTGTAACAGGAGTTGCCGTG |
| Human | PHF12 | Chromatin Readers | AATCCATGTGAAACAGGAGA |

| | | | |
|---|---|---|---|
| Human | PHF12 | Chromatin Readers | CAATAGTCACACTGGATGAG |
| Human | PHF12 | Chromatin Readers | CGGACACATCCATCTGCCCA |
| Human | PHF12 | Chromatin Readers | CAATCCATGTGAAACAGGAG |
| Human | PHF12 | Chromatin Readers | GCTAGAACCTGGTAGTGCAG |
| Human | PHF12 | Chromatin Readers | GTCGACATCATTCTGCTCAG |
| Human | PHF12 | Chromatin Readers | CACTACCAGGTTCTAGCAAG |
| Human | PHF12 | Chromatin Readers | CAGCATGAATTAGATCACAA |
| Human | PHF12 | Chromatin Readers | GGGTCATGTCAATGGACTGG |
| Human | PHF12 | Chromatin Readers | AGCGCCATCGTCAGATGGCA |
| Human | PHF12 | Chromatin Readers | GCGCTGGTGAGCTGAATCGT |
| Human | PHF12 | Chromatin Readers | CAGGAACCAAACTCCTCTGG |
| Human | UHRF2 | Chromatin Readers | CTCCAAGACGTGCACCATTG |
| Human | UHRF2 | Chromatin Readers | TGTGGGCGCTGTTCGACGTG |
| Human | UHRF2 | Chromatin Readers | AACATTAACAAACATGAACA |
| Human | UHRF2 | Chromatin Readers | TTTGCGGATGAAGTCGACCG |
| Human | UHRF2 | Chromatin Readers | ATATGATGGCATTTATAAGG |
| Human | UHRF2 | Chromatin Readers | AACAAACATGAACAGGGCAT |
| Human | UHRF2 | Chromatin Readers | TCTGTGTGATGAATGTAATG |
| Human | UHRF2 | Chromatin Readers | GCTCCTGTCGTGTATGTGGT |
| Human | UHRF2 | Chromatin Readers | TCTGGGACTTTATCCAAAGG |
| Human | UHRF2 | Chromatin Readers | TTTGGATAAAGTCCCAGAAG |
| Human | UHRF2 | Chromatin Readers | GGCAATCTTTACAGACATTG |
| Human | UHRF2 | Chromatin Readers | ACAGCGCTCCTTTAAGGCAC |
| Human | UHRF2 | Chromatin Readers | GTGCCTTAAAGGAGCGCTGT |
| Human | UHRF2 | Chromatin Readers | TTGCCTACAGCGCTCCTTTA |
| Human | UHRF2 | Chromatin Readers | TGGAAGCACTCAGTTGTCAC |
| Human | UHRF2 | Chromatin Readers | TATGATGAATACCCAGAAAG |
| Human | UHRF2 | Chromatin Readers | GTGCTTCAAACCAAGCACCA |
| Human | UHRF2 | Chromatin Readers | ACAGTCTGAATTAGACGTAG |
| Human | UHRF2 | Chromatin Readers | CTGAAGGAATAGAACGGTCA |
| Human | UHRF2 | Chromatin Readers | CTACGTTTACAGTATCCAGC |
| Human | UHRF2 | Chromatin Readers | CGGCTTTCAGTACTAGCTGA |
| Human | FOXA1 | DNA and RNA Modifications | AGTCATTGAAGGACAGCGAG |
| Human | FOXA1 | DNA and RNA Modifications | GTCCGGGTGCAGCGTCCAGT |
| Human | FOXA1 | DNA and RNA Modifications | CATTGAAGGACAGCGAGTGG |
| Human | FOXA1 | DNA and RNA Modifications | CAATGACTGCTTCGTCAAGG |
| Human | FOXA1 | DNA and RNA Modifications | GCTGGTTCTGCCGGTAATAG |
| Human | TBP | Transcription Elongation | ACGCCGAATATAATCCCAAG |
| Human | TBP | Transcription Elongation | GATTACCGCAGCAAACCGCT |
| Human | TBP | Transcription Elongation | TAATCCCAAGCGGTTTGCTG |
| Human | TBP | Transcription Elongation | ATTACCGCAGCAAACCGCTT |
| Human | TBP | Transcription Elongation | ATAACTACTAAATTGTTGGT |

| | | | |
|---|---|---|---|
| Human | TBP | Transcription Elongation | ATTAAACCAGGAAATAACTC |
| Human | TBP | Transcription Elongation | CATAACTACTAAATTGTTGG |
| Human | TBP | Transcription Elongation | GCTCATAACTACTAAATTGT |
| Human | TBP | Transcription Elongation | TCAAGATTCAGAATATGGTG |
| Human | TBP | Transcription Elongation | CTTCAAGATTCAGAATATGG |
| Human | TBP | Transcription Elongation | TTCAAGATTCAGAATATGGT |
| Human | SP110 | Chromatin Readers | CCAGATTGGGATATTCACGC |
| Human | SP110 | Chromatin Readers | ACAATGTCACCAGAAGAGAC |
| Human | SP110 | Chromatin Readers | TTTGATCCCTGTATCCAGAG |
| Human | SP110 | Chromatin Readers | TGTGCACCACTCTGGATACA |
| Human | SP110 | Chromatin Readers | AAATGGCTTGTGTATGGCAT |
| Human | SP110 | Chromatin Readers | CCCTAGGAGAGCTGCTGAAG |
| Human | SP110 | Chromatin Readers | AAACGGAATATACGTTGTGA |
| Human | SP110 | Chromatin Readers | AGTGAAGTGCATTCGGAATG |
| Human | SP110 | Chromatin Readers | TCTCGAATCAGCTGGTCCTG |
| Human | SP110 | Chromatin Readers | CCAGGGTCTTAGATACATGA |
| Human | SP110 | Chromatin Readers | CCATCATGTATCTAAGACCC |
| Human | SP110 | Chromatin Readers | TGGAAGCCAAGAGGATGCTG |
| Human | SP110 | Chromatin Readers | TTGTCCACGAGTCTTCCATG |
| Human | SP110 | Chromatin Readers | TGCGAGGTGTGCTGTCAAGG |
| Human | SP110 | Chromatin Readers | CAGTCCTCATGGAAGACTCG |
| Human | SP110 | Chromatin Readers | GCAACTCCACAGCATCCTCT |
| Human | DNMT3B | Chromatin Readers | CAGTTTGTCTGCAGAGACCT |
| Human | DNMT3B | Chromatin Readers | AGACATAGCCTGTCGCTTGG |
| Human | DNMT3B | Chromatin Readers | CTTCCAAGACACCACCATGG |
| Human | DNMT3B | Chromatin Readers | GGCACTCCACACAGAAACAC |
| Human | DNMT3B | Chromatin Readers | CTGCTGCCGGTGTTTCTGTG |
| Human | DNMT3B | Chromatin Readers | AGGACGCCATGACAGCGCTG |
| Human | DNMT3B | Chromatin Readers | GCCTGTATGAGGGTACAGGC |
| Human | DNMT3B | Chromatin Readers | TCTTCGGCCTCTGATCTCCG |
| Human | DNMT3B | Chromatin Readers | ACACAGGACTTGACAGGCGA |
| Human | DNMT3B | Chromatin Readers | TTCAGAGTATCAGGATGGGA |
| Human | DNMT3B | Chromatin Readers | CAGTCTGCTAAGCTACACAC |
| Human | DNMT3B | Chromatin Readers | CCCTAGCTTTCTCCAGAGCA |
| Human | DNMT3B | Chromatin Readers | ACCATGCTCTGGAGAAAGCT |
| Human | DNMT3B | Chromatin Readers | CCATGCTCTGGAGAAAGCTA |
| Human | DNMT3B | Chromatin Readers | TTTCTCCAGAGCATGGTACA |
| Human | DNMT3B | Chromatin Readers | GGACCAGCTGAAGCCCATGT |
| Human | DNMT3B | Chromatin Readers | AGCTGAAGCCCATGTTGGAG |
| Human | DNMT3B | Chromatin Readers | ACTCCAACATGGGCTTCAGC |
| Human | FOXA3 | DNA and RNA Modifications | CAAACATGTTCCCTGAGCTG |
| Human | FOXA3 | DNA and RNA Modifications | CCTTGAGTGAAATCTACCAG |

| | | | |
|---|---|---|---|
| Human | CHMP2A | Control | TGAAGAGGAGAGTGATGCTG |
| Human | CHMP2A | Control | GGGCCCTGAACCGTGCCATG |
| Human | CHMP2A | Control | GCGACAGAAACTAGAGACCC |
| Human | CHMP2A | Control | GCTACTGCGGCAGAACCAGA |
| Human | CHMP2A | Control | AGCTACTGCGGCAGAACCAG |
| Human | PARP8 | Other Histone Modifications | GTGTGAGCCATGAAATGCAA |
| Human | PARP8 | Other Histone Modifications | AATATTAATACACAAGAAGG |
| Human | PARP8 | Other Histone Modifications | GCATATCATTTGGTTACTCA |
| Human | PARP8 | Other Histone Modifications | AATACACGATTGCAGCTCCA |
| Human | AKAP1 | Chromatin Readers | GCATCCCAGGCAATGCCAAG |
| Human | AKAP1 | Chromatin Readers | AGCATCCCAGGCAATGCCAA |
| Human | AKAP1 | Chromatin Readers | GATGTTGAGAGCCTTCTATG |
| Human | AKAP1 | Chromatin Readers | GAGGTGCCAAAGCACTTAGT |
| Human | AKAP1 | Chromatin Readers | TGCCAAAGCACTTAGTCGGT |
| Human | AKAP1 | Chromatin Readers | AGCCGACCGACTAAGTGCTT |
| Human | AKAP1 | Chromatin Readers | GTCGGCTAATTGGCAAGCAG |
| Human | AKAP1 | Chromatin Readers | GTATCGAATCTCCACTTCGT |
| Human | AKAP1 | Chromatin Readers | GAGATTCGATACGTGGACTA |
| Human | AKAP1 | Chromatin Readers | TGGACTACGGCGGATATAAG |
| Human | SETD1A | Histone Methylation | TATGTGCCGTAAGTACGGTG |
| Human | SETD1A | Histone Methylation | CCAGGTGCTTGCGCGTACGG |
| Human | SETD1A | Histone Methylation | AGTGCTGGTGAAGAGCACAC |
| Human | SETD1A | Histone Methylation | GTGCCGTAAGTACGGTGAGG |
| Human | SETD1A | Histone Methylation | TCAGAACATCCGTCAGATGG |
| Human | SETD1A | Histone Methylation | CGGAGCCGGATCCACGAGTG |
| Human | SETD1A | Histone Methylation | GGGTCAGAACATCCGTCAGA |
| Human | SETD1A | Histone Methylation | CGAGATGGTCATCGAATACG |
| Human | SETD1A | Histone Methylation | TCTCTGTGCCACACAGACAC |
| Human | SETD1A | Histone Methylation | TGTGGCACAGAGAGCTGCCG |
| Human | SETD1A | Histone Methylation | CTCTCTGTGCCACACAGACA |
| Human | SETD1A | Histone Methylation | GTGTGGCACAGAGAGCTGCC |
| Human | SETD1A | Histone Methylation | TGTGTGGCACAGAGAGCTGC |
| Human | SETD1A | Histone Methylation | ATCTGTTCAAACTCACTGCG |
| Human | SETD1A | Histone Methylation | CCTGTATGACATTTGGAACT |
| Human | SETD1A | Histone Methylation | CAGAGGACATGAGTTACCTG |
| Human | SETD1A | Histone Methylation | GTTCCAAATGTCATACAGGA |
| Human | SETD1A | Histone Methylation | TGACATTTGGAACTCGGGCC |
| Human | SETD1A | Histone Methylation | TGGAGGGCGTGGACACTCAG |
| Human | SETD1A | Histone Methylation | TCAGGTTGGTGATTGTGTGA |
| Human | SETD1A | Histone Methylation | CTGGAGGGCGTGGACACTCA |
| Human | SETD1A | Histone Methylation | TTCACACCTCAGTCCTCCCA |
| Human | SETD1A | Histone Methylation | TGAAGTGGTGCTGCTGGAGT |

| | | | |
|---|---|---|---|
| Human | SETD1A | Histone Methylation | TAGCTGTTCTCTGACCGCCG |
| Human | SETD1A | Histone Methylation | CTTCAGCAGCATCTCGATGC |
| Human | SETD1A | Histone Methylation | ATCCTCCTCATCCTTCTCCG |
| Human | SETD1A | Histone Methylation | CTCCTCGGAGAAGGATGAGG |
| Human | SETD1A | Histone Methylation | AGACGACCCTGAACAAGAGA |
| Human | SETD1A | Histone Methylation | GACAGAGGTGTCGGATGGCG |
| Human | SETD1A | Histone Methylation | CTCCTCATCCTTCTCCGAGG |
| Human | SETD1A | Histone Methylation | TCTCTGTATGCTGACTCAGA |
| Human | SETD1A | Histone Methylation | AGAGGATGAAGAGGACGAGG |
| Human | SETD1A | Histone Methylation | GCTGGAAGAGCTGCTGCTCT |
| Human | SETD1A | Histone Methylation | CTCAGAGGATGAAGAGGACG |
| Human | SETD1A | Histone Methylation | CGAGGAGGATGAGGACGAGG |
| Human | SETD1A | Histone Methylation | GAGAAGAGGGACGCTCATCG |
| Human | SETD1A | Histone Methylation | AACAGGAGGCGTCTCCAGCA |
| Human | SETD1A | Histone Methylation | CTCAGCCACCACCACCTCGG |
| Human | SETD1A | Histone Methylation | CCTCAGCCACCACCACCTCG |
| Human | SETD1A | Histone Methylation | CTGCAGCAGCAGCGGGAGGA |
| Human | SETD1A | Histone Methylation | CGCTGCTGCTGCAGTTGCTG |
| Human | SETD1A | Histone Methylation | CGCCTCAGCCACCACCACCT |
| Human | DSN1 | Chromatin Remodeling | AGAACTGGAAACTGGAAAGC |
| Human | DSN1 | Chromatin Remodeling | TTCAGAAAGAGAAGATGCTT |
| Human | DSN1 | Chromatin Remodeling | CTTGCTTCACTACCAGCAGG |
| Human | DSN1 | Chromatin Remodeling | GCTCTTGCTTCACTACCAGC |
| Human | DSN1 | Chromatin Remodeling | TTCTTTAGAACGTCAGACTT |
| Human | DSN1 | Chromatin Remodeling | TTTCTTTAGAACGTCAGACT |
| Human | DSN1 | Chromatin Remodeling | ATATCTCTTTAGCCTCCTGC |
| Human | DSN1 | Chromatin Remodeling | TTGGTGATGGATGAACTGCA |
| Human | DSN1 | Chromatin Remodeling | TGACTGTATGGAGTTGGTGA |
| Human | DSN1 | Chromatin Remodeling | AGTCTTTGACTGTATGGAGT |
| Human | DSN1 | Chromatin Remodeling | CTGGGTACTTTCATCCATAA |
| Human | DSN1 | Chromatin Remodeling | AGGATCAGTGAAACAGCTGC |
| Human | CENPK | Chromatin Remodeling | TGAAGAAATGTGGAAAGATA |
| Human | CENPK | Chromatin Remodeling | TTAGAGAATGTGAAGAAATG |
| Human | CENPK | Chromatin Remodeling | ACTTAGAAAGGGAACAACGG |
| Human | CENPK | Chromatin Remodeling | AAGACTTAGAAAGGGAACAA |
| Human | CENPK | Chromatin Remodeling | GTTAAAGGAAGACTTAGAAA |
| Human | CENPK | Chromatin Remodeling | AGAAAGGGAACAACGGTGGT |
| Human | CENPK | Chromatin Remodeling | TGAAATGGTACTGTCCACTA |
| Human | L3MBTL3 | Chromatin Readers | GTATCCACAAACCTCCGCGA |
| Human | L3MBTL3 | Chromatin Readers | CACAAACCTCCGCGACGGTG |
| Human | L3MBTL3 | Chromatin Readers | TCGCGGAGGTTTGTGGATAC |
| Human | L3MBTL3 | Chromatin Readers | CTCACCGTCGCGGAGGTTTG |

| | | | |
|---|---|---|---|
| Human | L3MBTL3 | Chromatin Readers | GAATATAACAGTGATCCCAT |
| Human | L3MBTL3 | Chromatin Readers | CGTAGCAACACAGATGAATG |
| Human | L3MBTL3 | Chromatin Readers | TACCAACTCGAAAGCCCGAT |
| Human | L3MBTL3 | Chromatin Readers | AATATAACAGTGATCCCATC |
| Human | L3MBTL3 | Chromatin Readers | GTTTATTAGAGTAGCAACTG |
| Human | L3MBTL3 | Chromatin Readers | TTCAAAGTGAAACCTCCTCA |
| Human | L3MBTL3 | Chromatin Readers | GTTGCTACTCTAATAAACAT |
| Human | L3MBTL3 | Chromatin Readers | TTTAAAGATGAACAAATTGA |
| Human | L3MBTL3 | Chromatin Readers | TTACCTGGGTGTGAAGAACA |
| Human | L3MBTL3 | Chromatin Readers | GGACAGTTGGCAGCACTGTG |
| Human | L3MBTL3 | Chromatin Readers | GACAGTTGGCAGCACTGTGA |
| Human | L3MBTL3 | Chromatin Readers | GCCAATCCCTTTACATCCCG |
| Human | POLR1A | Transcription Elongation | ATGCAGTGGAAGAAGAACTC |
| Human | POLR1A | Transcription Elongation | TTCTTCTTCCACTGCATGGC |
| Human | POLR1A | Transcription Elongation | TGAGTTCTTCTTCCACTGCA |
| Human | ASXL2 | Chromatin Readers | AGGAGACGCACAGTTTGGAG |
| Human | ASXL2 | Chromatin Readers | AAAGCCATGATCATGTGCAA |
| Human | ASXL2 | Chromatin Readers | GCAGGAGACGCACAGTTTGG |
| Human | ASXL2 | Chromatin Readers | ACATGATCATGGCTTTCAAG |
| Human | ASXL2 | Chromatin Readers | TACCTGCCAAACCTGCAACA |
| Human | ASXL2 | Chromatin Readers | GCCAAACCTGCAACATGGGA |
| Human | ASXL2 | Chromatin Readers | CCATCACCCACCATTCCAGC |
| Human | ASXL2 | Chromatin Readers | CCCAACTGGTCAAAGCACAG |
| Human | ASXL2 | Chromatin Readers | CCAACTGGTCAAAGCACAGA |
| Human | ASXL2 | Chromatin Readers | CCTCTGTGCTTTGACCAGTT |
| Human | ASXL2 | Chromatin Readers | CCCTCTGTGCTTTGACCAGT |
| Human | ASXL2 | Chromatin Readers | GAAGGGCAGACTGCTAGAGG |
| Human | ASXL2 | Chromatin Readers | AGAGTCAGTGAAACTGGAAA |
| Human | ASXL2 | Chromatin Readers | CAGAGTCAGTGAAACTGGAA |
| Human | SP140L | Chromatin Readers | CCTGGTCCCTGTACAAAGAG |
| Human | SP140L | Chromatin Readers | GGCAAAGTGTATACAGACTG |
| Human | SP140L | Chromatin Readers | TCCTCCATCAGCCGTCGTAG |
| Human | SP140L | Chromatin Readers | GAAATTGGAACAAGGAACCT |
| Human | SP140L | Chromatin Readers | AGTCTGTATACACTTTGCCA |
| Human | SP140L | Chromatin Readers | CTACGACGGCTGATGGAGGA |
| Human | SP140L | Chromatin Readers | TATACTTGTAAGAGGCCCTG |
| Human | SP140L | Chromatin Readers | ATACTATTATTATATTAGAG |
| Human | SP140L | Chromatin Readers | TATATTAGAGAGGCGTGTCA |
| Human | SP140L | Chromatin Readers | CAGGGCCTCTTACAAGTATA |
| Human | SP140L | Chromatin Readers | CCTCATCTTCCAGAACCACA |
| Human | SP140L | Chromatin Readers | GGACTGCCACATCCCACCTG |
| Human | SP140L | Chromatin Readers | TCTTCTCACTTTCCACAGGT |

| | | | |
|---|---|---|---|
| Human | SP140L | Chromatin Readers | CACTTTCCACAGGTGGGATG |
| Human | UBTF | Transcription Elongation | ACAGGAGTTCGAGCGAAACC |
| Human | UBTF | Transcription Elongation | CCAGGTTGCTCATCTCAGGG |
| Human | UBTF | Transcription Elongation | TATTTCATCTTCTTCTTCTC |
| Human | UBTF | Transcription Elongation | GATTCTGTCCAAGAAATACA |
| Human | UBTF | Transcription Elongation | CCACCCTGAGATGAGCAACC |
| Human | UBTF | Transcription Elongation | GCGGCCAGATGCCACTACGA |
| Human | UBTF | Transcription Elongation | GCCAGATGCCACTACGAAGG |
| Human | UBTF | Transcription Elongation | GTCCTTCACCTCCTTCGTAG |
| Human | UBTF | Transcription Elongation | AGAAGGTGTATCTCAAAGTG |
| Human | UBTF | Transcription Elongation | GTGGTACACCCACGAGAAGA |
| Human | UBTF | Transcription Elongation | TACAGCGAGTAGCTGTTCGG |
| Human | UBTF | Transcription Elongation | AGCGAGTAGCTGTTCGGAGG |
| Human | UBTF | Transcription Elongation | GCTGGGCACGTCCTTCATGT |
| Human | UBTF | Transcription Elongation | CAGTACAGCGAGTAGCTGTT |
| Human | UBTF | Transcription Elongation | GAAGAAGAAGGCCAAGTACA |
| Human | UBTF | Transcription Elongation | GGCCAAGTACAAGGCCCGAG |
| Human | UBTF | Transcription Elongation | CGCTCAAGGCTCAGTCGGAG |
| Human | UBTF | Transcription Elongation | TGGCCCGCTTCAAGAATGAC |
| Human | UBTF | Transcription Elongation | GGCCCGCTTCAAGAATGACC |
| Human | UBTF | Transcription Elongation | CTTCAAGAATGACCGGGTGA |
| Human | UBTF | Transcription Elongation | TCACCCGGTCATTCTTGAAG |
| Human | UBTF | Transcription Elongation | GGTCATTCTTGAAGCGGGCC |
| Human | UBTF | Transcription Elongation | TTCTGGTAACCGTTCATGGG |
| Human | UBTF | Transcription Elongation | GCCGCTGAAGGAGCGCATGG |
| Human | UBTF | Transcription Elongation | TGGTGGAGATCGGCAGTCGC |
| Human | UBTF | Transcription Elongation | GATGAAGATAATGAGTCCGA |
| Human | UBTF | Transcription Elongation | CTCCTCTGAGTCCAGCAGCG |
| Human | ZCWPW2 | Chromatin Readers | CAAAGTATTCCTGGCCAACT |
| Human | ZCWPW2 | Chromatin Readers | AGGTTGATCATGATGAACCA |
| Human | ZCWPW2 | Chromatin Readers | ATCAAGTGAGGATTCAGCCA |
| Human | ZCWPW2 | Chromatin Readers | ATGGAGATTGTTATCAAGTG |
| Human | ZCWPW2 | Chromatin Readers | GTGTTCATGAAGCAGTACCA |
| Human | USP22 | Other Histone Modifications | TTTGCTGGAGGCCATGAAAG |
| Human | USP22 | Other Histone Modifications | CACTGCAAAGGTGATGACAA |
| Human | USP22 | Other Histone Modifications | GCCTCCGTACATCAGATCAA |
| Human | USP22 | Other Histone Modifications | CGTACATCAGATCAATGGCC |
| Human | USP22 | Other Histone Modifications | GAAGGCGAAGCGGCACAACC |
| Human | NSMCE1 | Control | TGGAAGTACTTGGCCACGCA |
| Human | NSMCE1 | Control | TGTCACAGCCTCCTCATCCA |
| Human | NSMCE1 | Control | TAAGTGCATCCTGATCCCAC |
| Human | NSMCE1 | Control | AGGTTTCGCAGCTTTGACCC |

| | | | |
|---|---|---|---|
| Human | NSMCE1 | Control | CAGGATGCACTTACCCTGCG |
| Human | NSMCE1 | Control | ACTTATCTACGGTGGCATTG |
| Human | NSMCE1 | Control | TGGGAGACCCATTTATGCGT |
| Human | NSMCE1 | Control | GGATTTGTTTAGAAAGGCTC |
| Human | NSMCE1 | Control | TGAACTGGATTTGTTTAGAA |
| Human | NSMCE1 | Control | TGGAGATGGAGCAATACATC |
| Human | NSMCE1 | Control | GGATGTATTGCTCCATCTCC |
| Human | NSMCE1 | Control | GTATTGCTCCATCTCCAGGA |
| Human | WDR61 | Transcription Elongation | TCTGGAAATGGCGTGATGAG |
| Human | WDR61 | Transcription Elongation | CCATTTGGTCAGTTGCTTGG |
| Human | WDR61 | Transcription Elongation | GAAATGGCGTGATGAGAGGC |
| Human | WDR61 | Transcription Elongation | GCCATTTCCAGACCTTCACC |
| Human | WDR61 | Transcription Elongation | AATATGAGCATCAAGAGAGC |
| Human | WDR61 | Transcription Elongation | AGAGAGCTGGATGCAGCAAT |
| Human | WDR61 | Transcription Elongation | TCATATTCGTCTTTGGGACT |
| Human | WDR61 | Transcription Elongation | GCTGGATGCAGCAATGGGCA |
| Human | WDR61 | Transcription Elongation | TCAGCTGGGAGTGGTGTCTG |
| Human | WDR61 | Transcription Elongation | GCCAAAGTCCAGGCATCCAC |
| Human | WDR61 | Transcription Elongation | ACCTGTGGATGCCTGGACTT |
| Human | WDR61 | Transcription Elongation | TTTCCCGACATGAGTTCCTG |
| Human | WDR61 | Transcription Elongation | GATTCCCAGTATCTGGCCAC |
| Human | WDR61 | Transcription Elongation | GTATTGCATATAGTCCTGAT |
| Human | WDR61 | Transcription Elongation | AGTATTGCATATAGTCCTGA |
| Human | WDR61 | Transcription Elongation | GATGGGAAATACCTAGCCAG |
| Human | WDR61 | Transcription Elongation | AATATTGATGATTCCATCTA |
| Human | WDR61 | Transcription Elongation | CTAGCCAGTGGAGCCATAGA |
| Human | WDR61 | Transcription Elongation | TGAAGCAGTGACAAGGAGCT |
| Human | WDR61 | Transcription Elongation | GGTCAAGGAGCGAATGGGCA |
| Human | WDR61 | Transcription Elongation | CATCATCTGAAGCAGTGACA |
| Human | WDR61 | Transcription Elongation | GACAAGGAGCTGGGAGTCCG |
| Human | WDR61 | Transcription Elongation | GAAACAAAGTGAGTGTCATC |
| Human | WDR61 | Transcription Elongation | AAGTTTGGGATGTTGGAACG |
| Human | WDR61 | Transcription Elongation | TGTGTCTGTTGGAGATGACC |
| Human | JADE1 | Chromatin Readers | GGAATCCTCAAGGTACCAGA |
| Human | JADE1 | Chromatin Readers | TGGAATCCTCAAGGTACCAG |
| Human | JADE1 | Chromatin Readers | GGCCTGTTATGGAATCCTCA |
| Human | JADE1 | Chromatin Readers | TGAGGATTCCATAACAGGCC |
| Human | JADE1 | Chromatin Readers | AGCTGCCCTCTGGTACCTTG |
| Human | JADE1 | Chromatin Readers | CACTTGGTTCCGCTACGGGT |
| Human | JADE1 | Chromatin Readers | CCACTTGGTTCCGCTACGGG |
| Human | JADE1 | Chromatin Readers | CTTCACAGAGCACTGTATAG |
| Human | RAG2 | Chromatin Readers | TGATGTGGATATCAACACTT |

| | | | |
|---|---|---|---|
| Human | RAG2 | Chromatin Readers | ATGCTGCCCTACTTGTGATG |
| Human | UNG | DNA and RNA Modifications | GGGCGTGTCGCTTCCTGGCG |
| Human | UNG | DNA and RNA Modifications | TCACTGATGCAGTTGTGTCC |
| Human | PRDM11 | Histone Methylation | AGCCAACTGGATGAGGTACG |
| Human | PRDM11 | Histone Methylation | GACCACGTACCTCATCCAGT |
| Human | PRDM11 | Histone Methylation | GAAGCCAGCTGATTTGTCCT |
| Human | ZAK | Other Histone Modifications | CTCCCAGAGAACCACACCAT |
| Human | ZAK | Other Histone Modifications | TGTATGGTTATGGAACCGAG |
| Human | ZAK | Other Histone Modifications | GTATTGAAGATCTGTGACTT |
| Human | ZAK | Other Histone Modifications | TATGGATCACATTATGACCT |
| Human | ZAK | Other Histone Modifications | TGTCACAAGTTTCTGACACA |
| Human | ZAK | Other Histone Modifications | GACATGGGCATTGTCTCCAA |
| Human | ZAK | Other Histone Modifications | ACATGGGCATTGTCTCCAAG |
| Human | ZAK | Other Histone Modifications | CGGAAGACGATGTGTATTGT |
| Human | ZAK | Other Histone Modifications | GAAGAAGACCTGAAAGACAT |
| Human | ZAK | Other Histone Modifications | AGAGCGTGATCTCAGCTTTA |
| Human | ZAK | Other Histone Modifications | AAATTGAGGCAACTCTTGAG |
| Human | ZAK | Other Histone Modifications | GGAGTGGAGGTGCGAAATTG |
| Human | FOXP1 | DNA and RNA Modifications | TCGCACACTGCTTCACAGCC |
| Human | FOXP1 | DNA and RNA Modifications | ACGCACTGCATTCTTCCACG |
| Human | FOXP1 | DNA and RNA Modifications | TCCACGTGGCCGCGTTGCGT |
| Human | FOXP1 | DNA and RNA Modifications | GGAATCATGCCTTGAGCAAG |
| Human | FOXP1 | DNA and RNA Modifications | ACAGCCTAAAGAGCAACAGC |
| Human | FOXP1 | DNA and RNA Modifications | CTGCTGCTGTTGAAGCATGA |
| Human | FOXP1 | DNA and RNA Modifications | TCTTCAGGTTCCCGTGTCAG |
| Human | WHSC1 | Histone Methylation | CCAAGGGTAACCCGACACTT |
| Human | WHSC1 | Histone Methylation | CCAAAGTGTCGGGTTACCCT |
| Human | WHSC1 | Histone Methylation | AAACCATGCAAGGCCACCAA |
| Human | WHSC1 | Histone Methylation | TTTGGTGTGGTCCAAAGTGT |
| Human | WHSC1 | Histone Methylation | TTGGTGTGGTCCAAAGTGTC |
| Human | WHSC1 | Histone Methylation | AACTTGGGAACTACAGATGG |
| Human | WHSC1 | Histone Methylation | TGGGAACTACAGATGGTGGC |
| Human | WHSC1 | Histone Methylation | TGAAACTTGGGAACTACAGA |
| Human | WHSC1 | Histone Methylation | AGATTATTACTGGACGCATC |
| Human | WHSC1 | Histone Methylation | GTTCTGGCAGAACTCGCCCG |
| Human | WHSC1 | Histone Methylation | CACTCCGAATCAAAGCCACA |
| Human | WHSC1 | Histone Methylation | GCCACAAGGATTCTCATCTG |
| Human | WHSC1 | Histone Methylation | CCACAAGGATTCTCATCTGT |
| Human | WHSC1 | Histone Methylation | TTCTGGCAGAACTCGCCCGC |
| Human | WHSC1 | Histone Methylation | AAGGACCGTATAATAGACGC |
| Human | WHSC1 | Histone Methylation | CTACATGCTCACTATAGACA |
| Human | WHSC1 | Histone Methylation | CCGTCTGTGACATTCCTGCA |

| | | | |
|---|---|---|---|
| Human | WHSC1 | Histone Methylation | TGTGGAGCCTCCAATTGCAG |
| Human | WHSC1 | Histone Methylation | TGCAATTGGAGGCTCCACAC |
| Human | WHSC1 | Histone Methylation | GCACCCAGATGCTTCAGGTG |
| Human | WHSC1 | Histone Methylation | CAACACCAAGTTTGCCCTGG |
| Human | WHSC1 | Histone Methylation | AGGTGGAAAGCTCCGCAGCA |
| Human | WHSC1 | Histone Methylation | GGGAAAGCCCAAGGCAGGCG |
| Human | WHSC1 | Histone Methylation | TGCCAGCTGTGTGAGAAGCC |
| Human | WHSC1 | Histone Methylation | AGGGTCCTTCACAGAGCAGG |
| Human | WHSC1 | Histone Methylation | GAGGTGGAAAGCTCCGCAGC |
| Human | WHSC1 | Histone Methylation | CGCTGTGTGGTAACTCAGTG |
| Human | WHSC1 | Histone Methylation | GACAGATGTTAAGCGCTGTG |
| Human | WHSC1 | Histone Methylation | CTGACTGTATTTGAGAGCCG |
| Human | WHSC1 | Histone Methylation | GCTTCGTGTGCTCCAAAGGG |
| Human | WHSC1 | Histone Methylation | TGCTTCGTGTGCTCCAAAGG |
| Human | WHSC1 | Histone Methylation | GTGCTTCGTGTGCTCCAAAG |
| Human | WHSC1 | Histone Methylation | CTGAACATCGAGATGCCTGA |
| Human | WHSC1 | Histone Methylation | CGATGTTCAGGCAGTCAGGG |
| Human | WHSC1 | Histone Methylation | GGACATTCCCACTTCCCGAA |
| Human | WHSC1 | Histone Methylation | AGGACATTCCCACTTCCCGA |
| Human | WHSC1 | Histone Methylation | CAAGCGGCCCTTCGGGAAGT |
| Human | WHSC1 | Histone Methylation | GCTGCCCGCCATCACCGCAG |
| Human | WHSC1 | Histone Methylation | CCTGCCTGGGCCTTGGCAAG |
| Human | WHSC1 | Histone Methylation | GAACAATGCCCATTTCCCAC |
| Human | WHSC1 | Histone Methylation | TTGAGGGCCCAGTGGGAAAT |
| Human | WHSC1 | Histone Methylation | AACAATGCCCATTTCCCACT |
| Human | WHSC1 | Histone Methylation | ATTGAGGGCCCAGTGGGAAA |
| Human | SETD4 | Histone Methylation | AGTCAAACATCCCTGCAGGA |
| Human | SETD4 | Histone Methylation | GAGTCAAACATCCCTGCAGG |
| Human | EIF5B | Control | ATTGTAGAATTTGCACAGCA |
| Human | EIF5B | Control | TGAAGGCAGTGAAGGTGATG |
| Human | EIF5B | Control | TGAAGAAAGTGAAGAAGAGG |
| Human | EIF5B | Control | GAAGAGGAGGGAGAAAGTGA |
| Human | EIF5B | Control | AGATGAAGAAAGTGAAGAAG |
| Human | EIF5B | Control | GAAGAAGAGGAGGAAGAGGA |
| Human | FOXO3 | DNA and RNA Modifications | CCGGATGGAGTTCTTCCAGC |
| Human | FOXO3 | DNA and RNA Modifications | ACTGCATAGTCGATTCATGC |
| Human | FOXO3 | DNA and RNA Modifications | GGATCATCAACCCTGATGGG |
| Human | FOXO3 | DNA and RNA Modifications | CCCTCCGGGTGCCAGCACCC |
| Human | FOXO3 | DNA and RNA Modifications | GCACCCGGGCCGAGTCCTCA |
| Human | FOXO3 | DNA and RNA Modifications | TGCTCCTTGAGGACTCGGCC |
| Human | FOXO3 | DNA and RNA Modifications | GCTCCTTGAGGACTCGGCCC |
| Human | FOXO3 | DNA and RNA Modifications | CCCGGGTGCTGGCACCCGGA |

| | | | |
|---|---|---|---|
| Human | FOXO3 | DNA and RNA Modifications | CTGGTTACCATAGTGTGACA |
| Human | FOXO3 | DNA and RNA Modifications | TGACAGAATTCGACAAGGCA |
| Human | FOXO3 | DNA and RNA Modifications | CCATGGAGACAGCCCGCCGC |
| Human | FOXO3 | DNA and RNA Modifications | ATGGAGACAGCCCGCCGCCG |
| Human | FOXO3 | DNA and RNA Modifications | TGGAGACAGCCCGCCGCCGG |
| Human | FOXO3 | DNA and RNA Modifications | GGAGACAGCCCGCCGCCGGG |
| Human | UBR1 | Other Histone Modifications | AGAGACAACCTATTCTTGCA |
| Human | UBR1 | Other Histone Modifications | TCCATACAGAGTACACATGT |
| Human | UBR1 | Other Histone Modifications | AAGATGCATACTTCTACTGG |
| Human | UBR1 | Other Histone Modifications | CTGTGACTGTGGAGACACAG |
| Human | UBR1 | Other Histone Modifications | TCCAACATGTGTACTCTGTA |
| Human | UBR1 | Other Histone Modifications | GGCAGTAGATTTCTGGACAC |
| Human | UBR1 | Other Histone Modifications | GACACAGGCCGATAATACCA |
| Human | UBR1 | Other Histone Modifications | ATGGATCCAGACTTGGCATA |
| Human | PKM | Other Histone Modifications | CTTTGCCATGAATGTTGGCA |
| Human | PKM | Other Histone Modifications | CCCACAGAAGCCACCGCCGT |
| Human | PKM | Other Histone Modifications | GCAGCACCTGATTGCCCGTG |
| Human | PKM | Other Histone Modifications | ATGAATGTTGGCAAGGCCCG |
| Human | PKM | Other Histone Modifications | CATTCTTGATGGTCTCCGCA |
| Human | PKM | Other Histone Modifications | ACATCCAGGATCTGAAGTTT |
| Human | PKM | Other Histone Modifications | GACATCCAGGATCTGAAGTT |
| Human | PKM | Other Histone Modifications | CAATCATCATCTTCTGAGCA |
| Human | PKM | Other Histone Modifications | CTTGCTCAGAAGATGATGAT |
| Human | PRKCD | Other Histone Modifications | AGGCGCTCAGCACAGAGCGT |
| Human | PRKCD | Other Histone Modifications | TCGATGCCCACATCTATGAG |
| Human | PRKCD | Other Histone Modifications | GAGGCGCTCAGCACAGAGCG |
| Human | PRKCD | Other Histone Modifications | CAGGCCCTGTAGGATCTCAG |
| Human | PRKCD | Other Histone Modifications | TATGCCGCTGAGATAATGTG |
| Human | PRKCD | Other Histone Modifications | AGACTACAGTAACTTTGACC |
| Human | PRKCD | Other Histone Modifications | ACTGGTCCATGGAGTCGATG |
| Human | PRKCD | Other Histone Modifications | TCAAAGTTACTGTAGTCTCT |
| Human | PRKCD | Other Histone Modifications | AGAAGCCAGCGAATGCAGAC |
| Human | PRKCD | Other Histone Modifications | GTGTGCAAAGACTTTGTCTG |
| Human | PRKCD | Other Histone Modifications | TTGCACACAGAACAGAAGGT |
| Human | PRKCD | Other Histone Modifications | GGTGGGTTGCCCAAAGAAGG |
| Human | PRKCD | Other Histone Modifications | TGTGTGCAAAGACTTTGTCT |
| Human | PRKCD | Other Histone Modifications | GGATTAAAGTGTGAAGACTG |
| Human | PRKCD | Other Histone Modifications | GAATGTGCACCATAAATGCC |
| Human | PRKCD | Other Histone Modifications | TGTAGTTGTGAACCTTGAAG |
| Human | PRKCD | Other Histone Modifications | GCACCATAAATGCCGGGAGA |
| Human | PRKCD | Other Histone Modifications | CCCACCTTCTGTGACCACTG |
| Human | NDUFAB1 | Control | CAAAGTCCCGAGCCTCGTCT |

| | | | |
|---|---|---|---|
| Human | NDUFAB1 | Control | GCACCGCTCTCTGCTCCGCG |
| Human | NDUFAB1 | Control | CACAACTGTGTAACTCTACC |
| Human | NDUFAB1 | Control | AAACCCAAATTCGTCTTCCA |
| Human | NDUFAB1 | Control | CTTAGACAGTTTGGACCAAG |
| Human | NDUFAB1 | Control | AATTTACTGAAAGCTTCTCT |
| Human | NDUFAB1 | Control | GGTCCAAACTGTCTAAGCCC |
| Human | UBR2 | Other Histone Modifications | TATAGGATGACAACATCAGG |
| Human | UBR2 | Other Histone Modifications | CAGTCTCTGCAAGAATATGT |
| Human | UBR2 | Other Histone Modifications | CGATATAGGATGACAACATC |
| Human | UBR2 | Other Histone Modifications | TGATACTGAAGCCTGGAAAG |
| Human | UBR2 | Other Histone Modifications | GTGTTCCACAAGACAGATCA |
| Human | UBR2 | Other Histone Modifications | GAACACACACTAGTAGCTGT |
| Human | UBR2 | Other Histone Modifications | TTCAAGCTAAAGAACAGCGA |
| Human | UBR2 | Other Histone Modifications | GGAACACACACTAGTAGCTG |
| Human | UBR2 | Other Histone Modifications | TGTGTTCCACAAGACAGATC |
| Human | UBR2 | Other Histone Modifications | GGGACAAAGACAAAGCTGAG |
| Human | CHD9 | Chromatin Readers | GGCCCACTCACAGTGAAGAT |
| Human | CHD9 | Chromatin Readers | CTCCTATCTTCACTGTGAGT |
| Human | CHD9 | Chromatin Readers | ACTCCTATCTTCACTGTGAG |
| Human | CHD9 | Chromatin Readers | TCTGTATCAATCATCACTCC |
| Human | CHD9 | Chromatin Readers | TGCCATATGAAGATAGTACT |
| Human | CHD9 | Chromatin Readers | GCCATATGAAGATAGTACTT |
| Human | CHD9 | Chromatin Readers | TTGAACAACTGCAAGCTTCA |
| Human | CHD9 | Chromatin Readers | TGTACTTCAGGGATTCACAG |
| Human | CHD9 | Chromatin Readers | ATGTACTTCAGGGATTCACA |
| Human | CHD9 | Chromatin Readers | ACTCATGAATCTGGAACACA |
| Human | CHD9 | Chromatin Readers | AGTGGAGCAATAATCAGGAA |
| Human | CHD9 | Chromatin Readers | AGCAGTTAAAGTCTACAGAC |
| Human | CHD9 | Chromatin Readers | TCTGTAACACAGCTTTATCT |
| Human | CHD9 | Chromatin Readers | GATGAGCACTTTATGACCTC |
| Human | CHD9 | Chromatin Readers | GTGTACCCGAGCTGGTGGGT |
| Human | CHD9 | Chromatin Readers | GATGCCCAACCCACCAGCTC |
| Human | CHD9 | Chromatin Readers | GTTGTCTAATATCACACCAT |
| Human | CHD9 | Chromatin Readers | TGAATGGAAGTACCTGATAC |
| Human | CHD9 | Chromatin Readers | AGAACAAGAAGTCATATTTG |
| Human | CHD9 | Chromatin Readers | AAGTGTAGCAGTCATCAAGA |
| Human | CHD9 | Chromatin Readers | CTTGATGACTGCTACACTTG |
| Human | CHD9 | Chromatin Readers | TGATATATGATTACTGGAAA |
| Human | CHD9 | Chromatin Readers | CACTTGTGGAAGTGTGCTGA |
| Human | CHD9 | Chromatin Readers | AAATACGCAGAAGATATAGA |
| Human | CHD9 | Chromatin Readers | AATACGCAGAAGATATAGAA |
| Human | CHD9 | Chromatin Readers | AAGGATGAAATCCAAGCCAA |

| | | | |
|---|---|---|---|
| Human | CHD9 | Chromatin Readers | TGGCAAACTCATTATTACAT |
| Human | CHD9 | Chromatin Readers | ACTAGAGCCTGATCGAGAGT |
| Human | CHD9 | Chromatin Readers | AACTAGAGCCTGATCGAGAG |
| Human | CHD9 | Chromatin Readers | TTCTGATGAAGATGAAGAGG |
| Human | CHD9 | Chromatin Readers | TCGAGAGTGGGAGCAAGAAG |
| Human | CHD9 | Chromatin Readers | GGAGCAAGAAGAGGATGATG |
| Human | CBX1 | Chromatin Readers | CTCAGATGAGGACAACACAT |
| Human | CBX1 | Chromatin Readers | TCTCAGATGAGGACAACACA |
| Human | CBX1 | Chromatin Readers | GTGGAAGGGATTCTCAGATG |
| Human | CBX1 | Chromatin Readers | GTAGCTCCAATAATCCGCTC |
| Human | CBX1 | Chromatin Readers | AGAAGGATATGACAACCTGT |
| Human | CBX1 | Chromatin Readers | AGCCAATGTCAAGTGCCCAC |
| Human | CBX1 | Chromatin Readers | AACCTGTGGGCACTTGACAT |
| Human | PHF14 | Chromatin Readers | TGTGGCATTACAGTCCATGA |
| Human | PHF14 | Chromatin Readers | TTGTCTGGGAGATAATAGTG |
| Human | PHF14 | Chromatin Readers | ACAGTCCATGAAGGTTGTTA |
| Human | PHF14 | Chromatin Readers | GGAGACAGATGCTGGAAGAT |
| Human | PHF14 | Chromatin Readers | AGGAGACAGATGCTGGAAGA |
| Human | PHF14 | Chromatin Readers | TTAGGACACAGTTCACAGCT |
| Human | PHF14 | Chromatin Readers | TTTGCATTAGCTGTGATGCA |
| Human | PHF14 | Chromatin Readers | CCCTCGCTTTGCTAGAACTG |
| Human | PHF14 | Chromatin Readers | ACCCTCGCTTTGCTAGAACT |
| Human | PHF14 | Chromatin Readers | AAACAGTTATTGGCAGTGCT |
| Human | PHF14 | Chromatin Readers | GTCTGGATCCTCCTCTTACA |
| Human | PHF14 | Chromatin Readers | CTTGGCATCCTTGTAAGAGG |
| Human | PHF14 | Chromatin Readers | GAGGATCCAGACATCCAAGA |
| Human | PHF14 | Chromatin Readers | TGTAAACTACATTACCATCT |
| Human | PHF14 | Chromatin Readers | GAAAGAGAAGGAAAGAGAGA |
| Human | L3MBTL1 | Chromatin Readers | ATAGCCACATACCTCAGCCA |
| Human | L3MBTL1 | Chromatin Readers | CACATACCTCAGCCACGGTG |
| Human | L3MBTL1 | Chromatin Readers | AGCCAGCAGGGTGAATGTCA |
| Human | L3MBTL1 | Chromatin Readers | CAGAATACCCATCAAAGTGC |
| Human | L3MBTL1 | Chromatin Readers | GCAGTCACTCACAACAAGAA |
| Human | L3MBTL1 | Chromatin Readers | TGGCCACGCAGACAAGGGAC |
| Human | L3MBTL1 | Chromatin Readers | GAACCCGTCCCTTGTCTGCG |
| Human | L3MBTL1 | Chromatin Readers | CACACTGGCCACGCAGACAA |
| Human | L3MBTL1 | Chromatin Readers | GAAATATCTGGAAGAAACTG |
| Human | L3MBTL1 | Chromatin Readers | GAACCCAGCCCTGATTCGCG |
| Human | L3MBTL1 | Chromatin Readers | CATGACTCCAGCCATCAAAG |
| Human | L3MBTL1 | Chromatin Readers | CAAAGTGGATCTTTATCCGA |
| Human | L3MBTL1 | Chromatin Readers | CCTGGTCAATATGAAGCTGG |
| Human | L3MBTL1 | Chromatin Readers | AAAGCCGAAGACCTCATCGA |

| | | | |
|---|---|---|---|
| Human | L3MBTL1 | Chromatin Readers | GAAGACTCTAGTCTGGACTG |
| Human | L3MBTL1 | Chromatin Readers | AGCACGCCTCTTCAAAGACG |
| Human | L3MBTL1 | Chromatin Readers | GAAGATGATCTGAGCACACA |
| Human | L3MBTL1 | Chromatin Readers | AGCCGTCGCAACCAGGAGTG |
| Human | L3MBTL1 | Chromatin Readers | CTGTGAACTTGCCTGTGACA |
| Human | L3MBTL1 | Chromatin Readers | GAGCCGTCGCAACCAGGAGT |
| Human | L3MBTL1 | Chromatin Readers | TGCTCCACTCCTCACTCTCT |
| Human | L3MBTL1 | Chromatin Readers | GCTCCACTCCTCACTCTCTG |
| Human | L3MBTL1 | Chromatin Readers | CTGCTCCACTCCTCACTCTC |
| Human | BAZ2B | Chromatin Readers | GTATTCCATTGGAATATGGC |
| Human | BAZ2B | Chromatin Readers | CTCTGCCAGCCATATTCCAA |
| Human | BAZ2B | Chromatin Readers | TGAGATACTTTATTACTTCA |
| Human | BAZ2B | Chromatin Readers | TGGAATAATGGATATCTCAA |
| Human | BAZ2B | Chromatin Readers | AAGTACAAGACTTGCTTGTG |
| Human | BAZ2B | Chromatin Readers | TCTTGAAGAACACTCAGGTT |
| Human | BAZ2B | Chromatin Readers | GATCACATACAGCAGCTGAG |
| Human | BAZ2B | Chromatin Readers | AAGCCTGACATCTAGAGCAA |
| Human | BAZ2B | Chromatin Readers | TTAATAACTTTCTTATAACC |
| Human | BAZ2B | Chromatin Readers | GTAAACTTGAAACTTGTTCC |
| Human | BAZ2B | Chromatin Readers | ACTTAGTTTCTCTCTAATTG |
| Human | BAZ2B | Chromatin Readers | GGAACAAGTTTCAAGTTTAC |
| Human | BAZ2B | Chromatin Readers | TTACTGCCAAATCTGTCGAA |
| Human | BAZ2B | Chromatin Readers | AAGATTACAACAATCCCAGA |
| Human | BAZ2B | Chromatin Readers | AATCTTGGGTCTATGGCAGT |
| Human | BAZ2B | Chromatin Readers | TACTGCCAAATCTGTCGAAA |
| Human | BAZ2B | Chromatin Readers | TATCTCCCTTTCGACAGATT |
| Human | BAZ2B | Chromatin Readers | AACATCAGGAACGAGAGCGA |
| Human | BAZ2B | Chromatin Readers | TGCAAGCTCAAGAAATAGCC |
| Human | BAZ2B | Chromatin Readers | TCGAGCTCAGCAAATTCTAG |
| Human | BAZ2B | Chromatin Readers | GGCAAATGCCAAATTATTGG |
| Human | BAZ2B | Chromatin Readers | AACAACAGAGTCAGTACAGA |
| Human | BAZ2B | Chromatin Readers | GCTCTTCAACACTTGCTGCT |
| Human | BAZ2B | Chromatin Readers | AGCAGCAAGTGTTGAAGAGC |
| Human | BAZ2B | Chromatin Readers | AAATCAAAGATAAGAAACCA |
| Human | BAZ2B | Chromatin Readers | TTGTCAACTCTAGCATCCAA |
| Human | BAZ2B | Chromatin Readers | ACTACTGCTTCAAGTTCCAT |
| Human | BAZ2B | Chromatin Readers | TCATCAGACACCTCAAGTGA |
| Human | BAZ2B | Chromatin Readers | AATTACTATCTGATTCTGAT |
| Human | BAZ2B | Chromatin Readers | AGATTCAAATGAGGATGAAG |
| Human | BAZ2B | Chromatin Readers | AATTCAGAATCAGATACAGA |
| Human | BAZ2B | Chromatin Readers | TCTTCAGAATCTTTACTACT |
| Human | BAZ2B | Chromatin Readers | AGATTCTGAAGATTCAAATG |

| | | | |
|---|---|---|---|
| Human | BAZ2B | Chromatin Readers | AGATAGACAACGAGCAAGAG |
| Human | BAZ2B | Chromatin Readers | AGGAATCCAGGATGAGACGT |
| Human | BAZ2B | Chromatin Readers | TCTCTTGCTCGTTGTCTATC |
| Human | BAZ2B | Chromatin Readers | AACGAGCAAGAGAGGAATCC |
| Human | BAZ2B | Chromatin Readers | ACGATGACAGTGATGACCAA |
| Human | BAZ2B | Chromatin Readers | GACGATGACAGTGATGACCA |
| Human | BAZ2B | Chromatin Readers | CGATGACAGTGATGACCAAG |
| Human | BAZ2B | Chromatin Readers | GACTGATATCTGTGAAGATG |
| Human | BAZ2B | Chromatin Readers | GCAGAGCTGGCCACAATATG |
| Human | BAZ2A | Chromatin Readers | CAAGTACCTGAGCCGCAACG |
| Human | BAZ2A | Chromatin Readers | CTCCGCCACCCATGTTGGAG |
| Human | BAZ2A | Chromatin Readers | GGGAAGACGAACTTCTTCTG |
| Human | BAZ2A | Chromatin Readers | CCAAGCTGTCACCTTGACAC |
| Human | BAZ2A | Chromatin Readers | GCTATGCAGGAACTCCACAA |
| Human | BAZ2A | Chromatin Readers | CCAAAGATGTGCCTAGCCTG |
| Human | BAZ2A | Chromatin Readers | CAAAGATGTGCCTAGCCTGG |
| Human | BAZ2A | Chromatin Readers | CCTGTGTCAAGGTGACAGCT |
| Human | BAZ2A | Chromatin Readers | GGGAGGGTACACCAGCTCAG |
| Human | BAZ2A | Chromatin Readers | GGTACCCACTCACCAAACGT |
| Human | BAZ2A | Chromatin Readers | CGGTACCCACTCACCAAACG |
| Human | BAZ2A | Chromatin Readers | GGCAGTTGTCAAATACCAGG |
| Human | BAZ2A | Chromatin Readers | GGGTCGAGGTCGGCCACCTA |
| Human | BAZ2A | Chromatin Readers | CTGCCATCGTCCCAAGATGG |
| Human | BAZ2A | Chromatin Readers | ACATGTCTAGTCTGCCGGAA |
| Human | BAZ2A | Chromatin Readers | GACATGTCTAGTCTGCCGGA |
| Human | BAZ2A | Chromatin Readers | TTACTGCCATCGTCCCAAGA |
| Human | BAZ2A | Chromatin Readers | CAGCCTCCATCTTGGGACGA |
| Human | BAZ2A | Chromatin Readers | GGCCCAAGAAACATTGAATG |
| Human | BAZ2A | Chromatin Readers | ATGAGGCAGAAGGTTCAACG |
| Human | BAZ2A | Chromatin Readers | AAGGGCAGGCCAGAAATAAG |
| Human | BAZ2A | Chromatin Readers | GAGTGTCAGACTACTATCCA |
| Human | BAZ2A | Chromatin Readers | TGGCTCTGACCCAGTGACAA |
| Human | BAZ2A | Chromatin Readers | GCTCTGACCCAGTGACAAAG |
| Human | BAZ2A | Chromatin Readers | TTCAGCTTCAGTCCCATAAG |
| Human | BAZ2A | Chromatin Readers | CTGGGACAGCCCAAACGGAG |
| Human | BAZ2A | Chromatin Readers | AAATGCCACAGAGTCCCACA |
| Human | BAZ2A | Chromatin Readers | ACCAGAGTGCTTGAGGATCA |
| Human | RPS6KA3 | Other Histone Modifications | TCGGTCTCGAACTTTCAGTG |
| Human | RPS6KA3 | Other Histone Modifications | CCTGCAAACAGATTAGGTGC |
| Human | RPS6KA3 | Other Histone Modifications | TGAAAGTTCGAGACCGAGTT |
| Human | RPS6KA3 | Other Histone Modifications | TGATGTGCATTAGCACTAGG |
| Human | RPS6KA3 | Other Histone Modifications | CATTTAAACCTGCAACGGGC |

| | | | |
|---|---|---|---|
| Human | RPS6KA3 | Other Histone Modifications | TTCAGACACAGCAAAGGACC |
| Human | HDAC11 | Histone Acetylation | CAGGAGGAACCATAATGGCG |
| Human | HDAC11 | Histone Acetylation | ATTGATCTTGATGCCCATCA |
| Human | HDAC11 | Histone Acetylation | GCTATCTTAATGAGCTCAAG |
| Human | HDAC11 | Histone Acetylation | TTGCCAAGCAGGCCATCAGG |
| Human | HDAC11 | Histone Acetylation | GACAGGAGGAACCATAATGG |
| Human | CBX6 | Chromatin Readers | GGCGATCAAGTACAGCACTT |
| Human | CBX6 | Chromatin Readers | GGGCGATCAAGTACAGCACT |
| Human | CBX6 | Chromatin Readers | GGGCGGCCAGAAGTAGCCCA |
| Human | CBX6 | Chromatin Readers | CCTGTCCCTCCCTCCCGAGT |
| Human | CBX6 | Chromatin Readers | AGAGCTGCGGGCGTCGTAGG |
| Human | CBX6 | Chromatin Readers | TGGTGGCTGCCGACTCGGGA |
| Human | ATP6V0B | Control | CCAGAAGGCCACGAAGACCC |
| Human | ATP6V0B | Control | GGCCTTCTGGGCCTGCGCGC |
| Human | ATP6V0B | Control | GGCCAGCGCGCAGGCCCAGA |
| Human | ATP6V0B | Control | TGATGTGGCATGGTTCCTGA |
| Human | ATP6V0B | Control | CGGAGACTTCGCCCTTCATG |
| Human | ATP6V0B | Control | GGAACCATGCCACATCAAAG |
| Human | ATP6V0B | Control | TGAAGGGCGAAGTCTCCGTC |
| Human | ATP6V0B | Control | CCACAGACAGGGAGATAGCT |
| Human | ATP6V0B | Control | CCTAGCTATCTCCCTGTCTG |
| Human | ATP6V0B | Control | CATCATTGGTGGAGGAGTGA |
| Human | ATP6V0B | Control | CTTCACTCCTCCACCAATGA |
| Human | ATP6V0B | Control | GGCCATCTACGGCATCATCA |
| Human | ATP6V0B | Control | TGCCATGATGATGCCGTAGA |
| Human | ATP6V0B | Control | TGTGAGGCTGTGGCCATCTA |
| Human | ATP6V0B | Control | CATCATCTTCTGTGAGGCTG |
| Human | ATP6V0B | Control | GGCCATCGGAACTACCATGC |
| Human | ATP6V0B | Control | ATGGTAGTTCCGATGGCCGA |
| Human | ATP6V0B | Control | AGCCTGCATGGTAGTTCCGA |
| Human | ATP6V0B | Control | TTCCGATGGCCGATGGCCTT |
| Human | ATP6V0B | Control | GGTTAGACAGGCCTACGGTG |
| Human | ATP6V0B | Control | CTCTTCTGTGGAGTCTGCGT |
| Human | ATP6V0B | Control | GGCCTGTCTAACCTCTTCTG |
| Human | ATP6V0B | Control | CCACGCAGACTCCACAGAAG |
| Human | ATP6V0B | Control | CAGCGCCATTGGCCTCTTTG |
| Human | ATP6V0B | Control | GCAGCGCCATTGGCCTCTTT |
| Human | ATP6V0B | Control | GGCAGCGCCATTGGCCTCTT |
| Human | ATP6V0B | Control | GAGATCTTTGGCAGCGCCAT |
| Human | HTATSF1 | Transcription Elongation | AGGCTACAAATTACATGTTG |
| Human | HTATSF1 | Transcription Elongation | CTACAAATTACATGTTGAGG |
| Human | HTATSF1 | Transcription Elongation | TTTGCCTCCAGATATTACAG |

| | | | |
|---|---|---|---|
| Human | HTATSF1 | Transcription Elongation | AAACTTTACAAAGATAATCA |
| Human | HTATSF1 | Transcription Elongation | CTCTTTGATAGGCACCCAGA |
| Human | HTATSF1 | Transcription Elongation | CTGGGTGCCTATCAAAGAGA |
| Human | HTATSF1 | Transcription Elongation | TGATAGGCACCCAGATGGTG |
| Human | HTATSF1 | Transcription Elongation | TCCCGTAAGAGGATCTGAAG |
| Human | HTATSF1 | Transcription Elongation | GAGAAAGATTTGGACGAGGA |
| Human | HTATSF1 | Transcription Elongation | TGTTTGACGATTCTGATGAG |
| Human | HTATSF1 | Transcription Elongation | CCTCAACAAGGAGTCTGAAG |
| Human | TDRD9 | Chromatin Readers | TGTGAAGGTGGTCCTGATGT |
| Human | TDRD9 | Chromatin Readers | CTCCGCCTACTGCAGCATTG |
| Human | TDRD9 | Chromatin Readers | GGTGACCACAATGCTGCAGT |
| Human | TDRD9 | Chromatin Readers | GAATAATCTTTCTGTACCCA |
| Human | TDRD9 | Chromatin Readers | CTCTAGACACTCGTCCAGCA |
| Human | TDRD9 | Chromatin Readers | CTATCCACTCCATTCAAGTG |
| Human | TDRD9 | Chromatin Readers | AGAATAATCTTTCTGTACCC |
| Human | TDRD9 | Chromatin Readers | AGAAATCCTTGTGTACCAGC |
| Human | TDRD9 | Chromatin Readers | GGTGCCAGACAGACCAAGTC |
| Human | TDRD9 | Chromatin Readers | ACATCCAGACTTGGTCTGTC |
| Human | TDRD9 | Chromatin Readers | TGTTTCTGGAAATTCTGCTG |
| Human | TDRD9 | Chromatin Readers | AGACAGACCAAGTCTGGATG |
| Human | SFMBT1 | Chromatin Readers | AAGATTATCTAGAAGAAACA |
| Human | SFMBT1 | Chromatin Readers | TCTGCTCACACCACCCAATG |
| Human | SFMBT1 | Chromatin Readers | CATAGCCATCATAGCGGAGA |
| Human | SFMBT1 | Chromatin Readers | AACAGATCCTGAGACCTACT |
| Human | SFMBT1 | Chromatin Readers | AAGGATTCCTCCCATTACGG |
| Human | SFMBT1 | Chromatin Readers | TAGAGGGCCTCCGTAATGGG |
| Human | SFMBT1 | Chromatin Readers | TTACGGAGGCCCTCTAGCAG |
| Human | SFMBT1 | Chromatin Readers | AGGACTACATGCTCCTATCA |
| Human | SFMBT1 | Chromatin Readers | CTGCTAGAGGGCCTCCGTAA |
| Human | SFMBT1 | Chromatin Readers | TCTTCAGACTCCACTGCACA |
| Human | SFMBT1 | Chromatin Readers | ACACAAAGGATCGCCGTGCG |
| Human | SFMBT1 | Chromatin Readers | AGATGCCAGGACTGTCGGCG |
| Human | SFMBT1 | Chromatin Readers | TCATGTTCAGAAATTAACGG |
| Human | SFMBT1 | Chromatin Readers | CATGTTCAGAAATTAACGGA |
| Human | SFMBT1 | Chromatin Readers | GCTGACTACCTCAAACAGTG |
| Human | SFMBT1 | Chromatin Readers | AACGGAGGGAAGCACCTCTG |
| Human | SFMBT1 | Chromatin Readers | TTGGGCTGGTGTGAAACCAA |
| Human | SFMBT1 | Chromatin Readers | ACTTGATCCTCTCTATGTGA |
| Human | SFMBT1 | Chromatin Readers | TCCATGCATTCTTGAACAGT |
| Human | SFMBT1 | Chromatin Readers | TCCTAGACCAGGAAATTGAT |
| Human | SFMBT1 | Chromatin Readers | TTCCTAGACCAGGAAATTGA |
| Human | SFMBT1 | Chromatin Readers | TGATGGCAAAGTTTGATGGC |

| | | | |
|---|---|---|---|
| Human | TLR7 | DNA and RNA Modifications | TGTCTCTTCAGTGTCCACAT |
| Human | TLR7 | DNA and RNA Modifications | ATCTGTATCTCTCTTTCTCA |
| Human | TLR7 | DNA and RNA Modifications | GAGAAAGAGAGATACAGATA |
| Human | TLR7 | DNA and RNA Modifications | AGGAAAGGGACTGGTTACCA |
| Human | TLR7 | DNA and RNA Modifications | GCAGTGTCTAAAGAACGCCC |
| Human | TLR7 | DNA and RNA Modifications | CCTGGCCACAGACAATCATG |
| Human | CCNT1 | Transcription Elongation | ATTCTTCTGCATGCTTCGCG |
| Human | CCNT1 | Transcription Elongation | TGCATATTGTGACTTCACAT |
| Human | CCNT1 | Transcription Elongation | AAGAATTGGCTGCCCAGAAG |
| Human | CCNT1 | Transcription Elongation | CGCGAAGCATGCAGAAGAAT |
| Human | CCNT1 | Transcription Elongation | GAAGAGGCAACTGGAGAACA |
| Human | CCNT1 | Transcription Elongation | TTGGAATTGGAGGGCATGCG |
| Human | CCNT1 | Transcription Elongation | GATGATGATGATGATTAGAT |
| Human | CCNT1 | Transcription Elongation | TGATGATGATGATGATTAGA |
| Human | CCNT1 | Transcription Elongation | GGGCATAGCTCAGACACAAG |
| Human | CCNT1 | Transcription Elongation | GACCCATTGTAGGAAGTGAA |
| Human | CCNT1 | Transcription Elongation | CCAGGCATCTGACCCATTGT |
| Human | CCNT1 | Transcription Elongation | TGACCCATTGTAGGAAGTGA |
| Human | CCNT1 | Transcription Elongation | GGAGGAGGTTCTGATGGCAG |
| Human | CCNT1 | Transcription Elongation | GGAGGTTCTGATGGCAGAGG |
| Human | CCNT1 | Transcription Elongation | AGTGGTGGAGGAGGTTCTGA |
| Human | CCNT1 | Transcription Elongation | GGTTCTGATGGCAGAGGTGG |
| Human | CCNT1 | Transcription Elongation | ACTTTCTTATCGCCAGCAGG |
| Human | CCNT1 | Transcription Elongation | AGAACTTTCTTATCGCCAGC |
| Human | CCNT1 | Transcription Elongation | GCCAATCTGCTTCAGGACAT |
| Human | CCNT1 | Transcription Elongation | CCAATCTGCTTCAGGACATG |
| Human | CCNT1 | Transcription Elongation | GGCCAATCTGCTTCAGGACA |
| Human | CCNT1 | Transcription Elongation | GCTGCTAGAAACAAGGCTGC |
| Human | CCNT1 | Transcription Elongation | TTTGCAACAAGTTCAAGATC |
| Human | CCNT1 | Transcription Elongation | AGCACAGACTTCTTACTTCA |
| Human | CCNT1 | Transcription Elongation | AGCCAGGTGAATGCAGACAC |
| Human | CCNT1 | Transcription Elongation | GGCCTGTGTCTGCATTCACC |
| Human | CCNT1 | Transcription Elongation | ATGCAGACACAGGCCACCAC |
| Human | KMT2C | Histone Methylation | TATGTATAGAGTGTGGCACA |
| Human | KMT2C | Histone Methylation | TGATAACACTTCCCACAGAA |
| Human | KMT2C | Histone Methylation | ATAACTTATGTCCCTTCTGT |
| Human | KMT2C | Histone Methylation | GACATAAGTTATCCTGCTGT |
| Human | KMT2C | Histone Methylation | GAAGCAGCTGACCAATTGTG |
| Human | KMT2C | Histone Methylation | GATGAGGCTACCCACGCGAA |
| Human | KMT2C | Histone Methylation | TGGAATGCTTGCATCTGCTG |
| Human | KMT2C | Histone Methylation | GGAGCACTCGCTATGCCAAT |
| Human | KMT2C | Histone Methylation | TCCTAAAGCACTCTTCCCTG |

| | | | |
|---|---|---|---|
| Human | KMT2C | Histone Methylation | CATAGCGGAATCACTTCCTG |
| Human | KMT2C | Histone Methylation | CATCTCACCTAAAGGTGTCT |
| Human | KMT2C | Histone Methylation | GTGAGATGTCACTTAGAACC |
| Human | KMT2C | Histone Methylation | GCATAGCGGAATCACTTCCT |
| Human | KMT2C | Histone Methylation | CGAATGGTTGATATACCTTG |
| Human | KMT2C | Histone Methylation | CTTTATGAGTCTCAGAACCG |
| Human | KMT2C | Histone Methylation | TGGCACGGTCTCGGATTCAG |
| Human | KMT2C | Histone Methylation | TGCGGAACATGTACACACCA |
| Human | KMT2C | Histone Methylation | ACTGTGGAGCTGTGAACTGC |
| Human | KMT2C | Histone Methylation | TTCACAGCTCCACAGTGACA |
| Human | KMT2C | Histone Methylation | GAGCTGTGAACTGCCGGAAG |
| Human | KMT2C | Histone Methylation | CTGTAGAATCAAATAAGGCT |
| Human | KMT2C | Histone Methylation | CTCATCACCGTTGTGTGGAG |
| Human | KMT2C | Histone Methylation | TTGATTCTACAGGCACTTGT |
| Human | KMT2C | Histone Methylation | TTGGGCTCATCACCGTTGTG |
| Human | KMT2C | Histone Methylation | TGATTCTACAGGCACTTGTT |
| Human | KMT2C | Histone Methylation | AGCATTTGATAGTGGCTCCA |
| Human | KMT2C | Histone Methylation | ACTGAAATCCTGAAAGGTGC |
| Human | KMT2C | Histone Methylation | GATGTGACTGAAATCCTGAA |
| Human | KMT2C | Histone Methylation | TATCCTTGTGCTGCAGGAGC |
| Human | KMT2C | Histone Methylation | TGGAATGTGCCTGGATATAG |
| Human | KMT2C | Histone Methylation | ATGGAGTAACCGCTATATCC |
| Human | KMT2C | Histone Methylation | ACTGATCTAAGAGGTCTCCC |
| Human | KMT2C | Histone Methylation | ATGAAATCAGTACCAACCAA |
| Human | KMT2C | Histone Methylation | TGCCAGAACTGCAAACAATC |
| Human | KMT2C | Histone Methylation | AATCAGTACCAACCAATGGC |
| Human | KMT2C | Histone Methylation | GTGTGTGATACGTGTGACAA |
| Human | KMT2C | Histone Methylation | GTGCCAGAACTGCAAACAAT |
| Human | KMT2C | Histone Methylation | TTCATTGTAATATGTGCAAA |
| Human | KMT2C | Histone Methylation | CAAACCAACAGATCATGAAC |
| Human | KMT2C | Histone Methylation | GTATCCAGTTCATGATCTGT |
| Human | KMT2C | Histone Methylation | CAGTATTAAGATCACTAAAG |
| Human | KMT2C | Histone Methylation | TAATACTGACACAGTATGGA |
| Human | KMT2C | Histone Methylation | ACTAAAGTGGTTCTTAGCAA |
| Human | KMT2C | Histone Methylation | TGGTTCTTAGCAAAGGTTGG |
| Human | KMT2C | Histone Methylation | AAGTGGTTCTTAGCAAAGGT |
| Human | KMT2C | Histone Methylation | TTGCAGACAGTTCCCAAAGG |
| Human | KMT2C | Histone Methylation | AGACAGTTCCCAAAGGAGGC |
| Human | KMT2C | Histone Methylation | TGTGTGTGAGGCCTGTGGGA |
| Human | KMT2C | Histone Methylation | CACACAGCAGGAGTCTTCCT |
| Human | KMT2C | Histone Methylation | CCATTGCAGACAGTTCCCAA |
| Human | KMT2C | Histone Methylation | TTGTCAGAACTTAAATACTG |

| | | | |
|---|---|---|---|
| Human | KMT2C | Histone Methylation | AATGTAGACAATGTGATAGA |
| Human | KMT2C | Histone Methylation | GTTTCGATAGCAGACTGGAC |
| Human | KMT2C | Histone Methylation | CTTAAATACTGAGGAAGAAG |
| Human | KMT2C | Histone Methylation | TCCACTTGAACTGCGCTCTG |
| Human | KMT2C | Histone Methylation | GATTGACAGATGGACCAGCA |
| Human | KMT2C | Histone Methylation | GAAGGTGATGGATTGACAGA |
| Human | KMT2C | Histone Methylation | GGACCCACAGATCCAAGTCA |
| Human | KMT2C | Histone Methylation | ACCACAGAGCGCAGTTCAAG |
| Human | KMT2C | Histone Methylation | TGTTGGTGCATCGAAATCTG |
| Human | KMT2C | Histone Methylation | CACAAGACGGGTGCCACTAG |
| Human | KMT2C | Histone Methylation | AAATCTGTGGCATCCACTAG |
| Human | KMT2C | Histone Methylation | TGTGTCTTCTGTCACAAGAC |
| Human | KMT2C | Histone Methylation | AAACATGCATTGTGCTTTAA |
| Human | KMT2C | Histone Methylation | GTTCTTGCTGCACAGTGATC |
| Human | KMT2C | Histone Methylation | AGCCACACAGAAACTTGAAC |
| Human | KMT2C | Histone Methylation | CACCTGTTCAAGTTTCTGTG |
| Human | KMT2C | Histone Methylation | CAGTACGTTGTTTAGCTGAA |
| Human | KMT2C | Histone Methylation | AGCTGGGCGTGAATTTCCAG |
| Human | KMT2C | Histone Methylation | AACGTACTGCCAAGAAAGCT |
| Human | KMT2C | Histone Methylation | AGCGTAAGCAGTATGAAGAG |
| Human | KMT2C | Histone Methylation | ACGGATCCAACTCATGCAGG |
| Human | KMT2C | Histone Methylation | ACAACGGATCCAACTCATGC |
| Human | KMT2C | Histone Methylation | GAGAACAGCAAGAGAGACAA |
| Human | KMT2C | Histone Methylation | AAGAGCTTTGCAGCAGAGGA |
| Human | KMT2C | Histone Methylation | TCCGAATCAATACGAGAGCT |
| Human | KMT2C | Histone Methylation | TCCCAGCTCTCGTATTGATT |
| Human | KMT2C | Histone Methylation | CTCCGAATCAATACGAGAGC |
| Human | KMT2C | Histone Methylation | CAGCAGCAACAACAGCAATT |
| Human | KMT2C | Histone Methylation | AATCAGAACATGAACAGGAA |
| Human | KMT2C | Histone Methylation | AAGAACATAAATCTCTGACA |
| Human | KMT2C | Histone Methylation | AAACTACAGGAGACCGGGAT |
| Human | KMT2C | Histone Methylation | GTATCAGGTGGCCTTACCAA |
| Human | KMT2C | Histone Methylation | TTACACCTGTAACAAATCAG |
| Human | KMT2C | Histone Methylation | GGGTCAACAGTAGTAGGCCG |
| Human | KMT2C | Histone Methylation | AATCCAAATCTAAATCCATG |
| Human | KMT2C | Histone Methylation | GACATAGGGCTCCTGACGGA |
| Human | KMT2C | Histone Methylation | GCATATCTAGGTCCTAAAGG |
| Human | KMT2C | Histone Methylation | GTAGCTGATTAGGTAGACCC |
| Human | KMT2C | Histone Methylation | TTAAATTTAGATACAGAGGA |
| Human | KMT2C | Histone Methylation | GGAAACTAATGATCCCAACC |
| Human | KMT2C | Histone Methylation | AAACTTAAATTTAGATACAG |
| Human | KMT2C | Histone Methylation | GGATGTTAAAGACCTTGAGG |

| | | | |
|---|---|---|---|
| Human | KMT2C | Histone Methylation | ACAGATCCAGAACTTGACAT |
| Human | KMT2C | Histone Methylation | TGTGTTATACTGCCATCCTG |
| Human | KMT2C | Histone Methylation | CCATCCTGAGGAATGGCCTG |
| Human | KMT2C | Histone Methylation | GGGAACCTGCTCATCACCAT |
| Human | KMT2C | Histone Methylation | TATACTGCCATCCTGAGGAA |
| Human | KMT2C | Histone Methylation | TGAAGCAAATTGCCAAATTG |
| Human | CBX5 | Chromatin Readers | TTCTGAGGAGCACAATACTT |
| Human | CBX5 | Chromatin Readers | AAGTGGAATATCTACTGAAG |
| Human | CBX5 | Chromatin Readers | GAAGGTGCTAGACAGGCGCG |
| Human | CBX5 | Chromatin Readers | TTTATGAAGAGAGACTGACA |
| Human | CBX5 | Chromatin Readers | CTGTGTCTTTCCATTTCATT |
| Human | CBX5 | Chromatin Readers | AGACACAGATGAAGCTGACC |
| Human | CBX5 | Chromatin Readers | GCATGCATATCCTGAGGATG |
| Human | ARID1A | Chromatin Remodeling | ACTTGCAACCAACCTCAATG |
| Human | ARID1A | Chromatin Remodeling | GGAGATTGGTGGATTGACTC |
| Human | ARID1A | Chromatin Remodeling | TGCTTGATGTGCCCACATTG |
| Human | ARID1A | Chromatin Remodeling | AGAGGTTTCCTACCCACAGC |
| Human | ARID1A | Chromatin Remodeling | CATATGGCCCTCCTGCCAAG |
| Human | ARID1A | Chromatin Remodeling | CGGCCAATGGATGGCACATA |
| Human | ARID1A | Chromatin Remodeling | GGGCCATATGTGCCATCCAT |
| Human | ARID1A | Chromatin Remodeling | AGCAACAGCAGAATTACAAG |
| Human | ARID1A | Chromatin Remodeling | TCCTGCCAAGCGGCACGAAG |
| Human | ARID1A | Chromatin Remodeling | GCAGAATTACAAGCGGCCAA |
| Human | ARID1A | Chromatin Remodeling | AATTACAAGCGGCCAATGGA |
| Human | ARID1A | Chromatin Remodeling | ACAGCGTGCCATACAGCACT |
| Human | THOC5 | Transcription Elongation | CGATCTGACACCGAGCAGGA |
| Human | THOC5 | Transcription Elongation | GGCTCCACCAGATATCAGCA |
| Human | THOC5 | Transcription Elongation | ATAACTCATAGTCTCTGCCA |
| Human | THOC5 | Transcription Elongation | GGAAGGTAAATACTACAGTG |
| Human | PHF5A | Chromatin Readers | ACATATGCGCACCAGAGTGC |
| Human | PHF5A | Chromatin Readers | CATATGCGCACCAGAGTGCA |
| Human | WHSC1L1 | Histone Methylation | TTTAGTATGAACCTCAAGCT |
| Human | WHSC1L1 | Histone Methylation | CAACCAGCCAGAGAGGGCGT |
| Human | WHSC1L1 | Histone Methylation | TCAAATTGGGAAATTACAGA |
| Human | WHSC1L1 | Histone Methylation | TCTGTGCCACTGAACATCCA |
| Human | WHSC1L1 | Histone Methylation | AATTGGGAAATTACAGATGG |
| Human | WHSC1L1 | Histone Methylation | TCTGACAACGATCTCCAGCT |
| Human | WHSC1L1 | Histone Methylation | GCAGTATGAATGCCACCCGC |
| Human | WHSC1L1 | Histone Methylation | CACTCCGATTCCAAGCCACA |
| Human | WHSC1L1 | Histone Methylation | AATTAGTTACACTGTTCTCG |
| Human | WHSC1L1 | Histone Methylation | ATTAGTTACACTGTTCTCGT |
| Human | WHSC1L1 | Histone Methylation | AGTGAATGGAGATGTTCGAG |

| | | | |
|---|---|---|---|
| Human | WHSC1L1 | Histone Methylation | TGCAGTTATCTGCTCCACAG |
| Human | WHSC1L1 | Histone Methylation | TGTGGAGCAGATAACTGCAG |
| Human | WHSC1L1 | Histone Methylation | TTGCAGCACTCTCCCTCACA |
| Human | WHSC1L1 | Histone Methylation | CAGGAAGTGATGCCAATCCC |
| Human | WHSC1L1 | Histone Methylation | GTGCTGCAAACACTTTCACC |
| Human | WHSC1L1 | Histone Methylation | CACTTTCACCTGGAGTGCCT |
| Human | WHSC1L1 | Histone Methylation | GACTCTCTGATTCCTTGTGA |
| Human | WHSC1L1 | Histone Methylation | GATGTGAAGCGTTGTTCTGT |
| Human | WHSC1L1 | Histone Methylation | TTGATTCAAAGATGGCAGTG |
| Human | WHSC1L1 | Histone Methylation | CTATGCTTAGGCATTCCGGG |
| Human | WHSC1L1 | Histone Methylation | AGGCATATGAACTGAACATG |
| Human | WHSC1L1 | Histone Methylation | TTTGTTTCGTTTGTGCCAGA |
| Human | WHSC1L1 | Histone Methylation | GTCCTGAACTATCAGCCCTC |
| Human | WHSC1L1 | Histone Methylation | TCAGCCACCATATGGAAAGT |
| Human | WHSC1L1 | Histone Methylation | CTCAGCCACCATATGGAAAG |
| Human | WHSC1L1 | Histone Methylation | GAGTCAGGTTAAGGCATAGG |
| Human | WHSC1L1 | Histone Methylation | GCTGAGTCAGGTTAAGGCAT |
| Human | WHSC1L1 | Histone Methylation | CACTCCCACTTTCCATATGG |
| Human | WHSC1L1 | Histone Methylation | CTCTTTGTGCTTTCAATTCC |
| Human | RPH3A | Chromatin Readers | GTGAACCGCTGCATACTGTG |
| Human | RPH3A | Chromatin Readers | GGAGCGAATCGGACGCCTGG |
| Human | RPH3A | Chromatin Readers | GGAGAAAGAAATCATCAACA |
| Human | RPH3A | Chromatin Readers | ATCACAGACGGAGATCCTGA |
| Human | RPH3A | Chromatin Readers | CATCACAGACGGAGATCCTG |
| Human | RPH3A | Chromatin Readers | TGCACCATCATTAAGGCCAA |
| Human | RPH3A | Chromatin Readers | AAGTCCAATGATTACATCGG |
| Human | RPH3A | Chromatin Readers | TTCAGCCAGAGCTTGACGAA |
| Human | RPH3A | Chromatin Readers | CTTGAATCCCGAATTCAATG |
| Human | RPH3A | Chromatin Readers | GGCAAGTCCAATGATTACAT |
| Human | RPH3A | Chromatin Readers | TGACATCAAACACAGTGACC |
| Human | RPH3A | Chromatin Readers | CTTTCCAGATCAGAAGCCAG |
| Human | RPH3A | Chromatin Readers | TCCAGATCAGAAGCCAGAGG |
| Human | RPH3A | Chromatin Readers | GAGCCCAGCAGGTTTGAGAC |
| Human | RPH3A | Chromatin Readers | GGAGCCCAGCAGGTTTGAGA |
| Human | KDM8 | Histone Methylation | TAACACGAGCCAGGTTGACG |
| Human | KDM8 | Histone Methylation | ACTTCCTAGTGCAGGTGATG |
| Human | KDM8 | Histone Methylation | AGACCTGAAGTTGGACCTCG |
| Human | KDM8 | Histone Methylation | CGTAGACAAAGACTGGCGCC |
| Human | KDM8 | Histone Methylation | AGAAGCTCAACACGGGCACA |
| Human | PARP6 | Other Histone Modifications | ACTGAATGGATCGGGTCTGG |
| Human | PARP6 | Other Histone Modifications | ATGAACTTCAGCCTGCTGAG |
| Human | PARP6 | Other Histone Modifications | GTGGGACCCATGGAAGGCAA |

| | | | |
|---|---|---|---|
| Human | PARP6 | Other Histone Modifications | GATTTCCAAATATGAGCCCT |
| Human | ING5 | Chromatin Readers | ATCCAGACTGTCCAATTGAG |
| Human | ING5 | Chromatin Readers | AACCACTCAATTGGACAGTC |
| Human | ING5 | Chromatin Readers | GCGAGAGCTGGACCAGAGGA |
| Human | ING5 | Chromatin Readers | GAAAGCAGAGATTGACATCC |
| Human | ING5 | Chromatin Readers | CTTTCTTATCTTCCGTCCTC |
| Human | ING5 | Chromatin Readers | TGATGCGAGAGCTGGACCAG |
| Human | ING5 | Chromatin Readers | CTTCCAGCTGATGCGAGAGC |
| Human | ING5 | Chromatin Readers | CATGCAGACCTACGAGATGG |
| Human | ING5 | Chromatin Readers | GTGTTTATCCACCATCTCGT |
| Human | ING5 | Chromatin Readers | GGCCATGCAGACCTACGAGA |
| Human | ING5 | Chromatin Readers | GTATTCCTTGCACTTGCTGT |
| Human | ING5 | Chromatin Readers | GAACGCCTACAGCAAGTGCA |
| Human | GGPS1 | Control | AGTGAGAACCAAACTTTCAC |
| Human | GGPS1 | Control | ATTAAATGCCTGTGAAAGTT |
| Human | GGPS1 | Control | GCCAAGGAAATACACGTAAT |
| Human | GGPS1 | Control | TTACGTGTATTTCCTTGGCT |
| Human | GGPS1 | Control | TTGGCAGAATTGATGACAGA |
| Human | GGPS1 | Control | GCCAATTACGTGTATTTCCT |
| Human | GGPS1 | Control | AGCCCAAGTGTATTAAGTAG |
| Human | GGPS1 | Control | TAGCATAATCATCCCTAATT |
| Human | GGPS1 | Control | TTTAGCTTCAAGCTCTTTAA |
| Human | GGPS1 | Control | CTTTAGCTTCAAGCTCTTTA |
| Human | YEATS4 | Chromatin Readers | GAAATTACTGAAACAGGATG |
| Human | YEATS4 | Chromatin Readers | TAGTTTACGGTAATGTTGCT |
| Human | YEATS4 | Chromatin Readers | AAGATGGGCACACTCATCAG |
| Human | YEATS4 | Chromatin Readers | TGTGAAACCATATAGAAATG |
| Human | YEATS4 | Chromatin Readers | CTTTGAGATTGCAGAGCTTA |
| Human | YEATS4 | Chromatin Readers | AAATTCTGTTTCATGCTTAT |
| Human | PHF23 | Chromatin Readers | ACTGTCGAAAGCCCTTTGCA |
| Human | PHF23 | Chromatin Readers | TCTTAATCTTAGCACAGGAG |
| Human | PHF23 | Chromatin Readers | CACAGGCTGCACTCAATCAT |
| Human | PHF23 | Chromatin Readers | GGAGGAGGAAGAGGAGGAAG |
| Human | PGK1 | Control | AGCTCGGAAAGCTTCTATTT |
| Human | PGK1 | Control | AAGCAGCAAGAAGTATGCTG |
| Human | PGK1 | Control | CAAAGCTCTCATGGATGAGG |
| Human | PGK1 | Control | AACCAAAGCTCTCATGGATG |
| Human | PGK1 | Control | CACCTCATCCATGAGAGCTT |
| Human | ZBTB33 | DNA and RNA Modifications | AAAGAATATTCTTGTGAGCC |
| Human | ZBTB33 | DNA and RNA Modifications | ATCAGCTCTTCTCTGTTGCT |
| Human | ZBTB33 | DNA and RNA Modifications | CAGCAACAGAGAAGAGCTGA |
| Human | ZBTB33 | DNA and RNA Modifications | ATGTCTGTCTGACAAGCTTG |

| | | | |
|---|---|---|---|
| Human | ZBTB33 | DNA and RNA Modifications | TTTGTGCGATATTCTGCAAG |
| Human | ZBTB33 | DNA and RNA Modifications | ACCTTCTCACAGTAACGGCA |
| Human | ZBTB33 | DNA and RNA Modifications | GAAATACCTTCTCACAGTAA |
| Human | ZBTB33 | DNA and RNA Modifications | GTTGATGAAAGATTTGCCAC |
| Human | ZBTB33 | DNA and RNA Modifications | AAGCATCTCAGGTACAGCGC |
| Human | ZBTB33 | DNA and RNA Modifications | GCTTGGTGTCCCATTGTCAC |
| Human | ZBTB33 | DNA and RNA Modifications | CTGTGATGTTACCGTTATTG |
| Human | ZBTB33 | DNA and RNA Modifications | ATGGCCACGTTGCTCATTCA |
| Human | ZBTB33 | DNA and RNA Modifications | TTTGGGAGGTATCAGCCTGT |
| Human | ZBTB33 | DNA and RNA Modifications | TCAGAACAAGCCAATGACCT |
| Human | ZBTB33 | DNA and RNA Modifications | GATGTAGTGGTCCCTACTGG |
| Human | ZBTB33 | DNA and RNA Modifications | ATTGGGTATAAGGTTGACAC |
| Human | HDAC2 | Histone Acetylation | ATTTAAGCCTATTATCTCAA |
| Human | HDAC2 | Histone Acetylation | CAGATGCAGAGATTTAATGT |
| Human | CDK13 | Transcription Elongation | TGTATAGCTCAGAAGAAAGT |
| Human | CDK13 | Transcription Elongation | AAGTCGGCCGTATACTAACA |
| Human | CDK13 | Transcription Elongation | AATATCCTTCTAAATAATAG |
| Human | CDK13 | Transcription Elongation | GAATTAATAAGCCGAATATG |
| Human | PHF8 | Histone Methylation | CACAGCATGGATCCACCCTG |
| Human | PHF8 | Histone Methylation | TACCAGACTTTCTAACCTTG |
| Human | PHF8 | Histone Methylation | GTCTGGTACCATGTACTCAA |
| Human | PHF8 | Histone Methylation | GTCACACTCGATCATGAAGC |
| Human | PHF8 | Histone Methylation | TGTCACACTCGATCATGAAG |
| Human | PHF8 | Histone Methylation | AACTGCCATGAAACCAGTCC |
| Human | PHF8 | Histone Methylation | GATCGAGTGTGACATGTGCC |
| Human | PHF8 | Histone Methylation | CAGTTGTGTTGGTGTTGAAG |
| Human | PHF8 | Histone Methylation | GCAGTGGGAGCTCCAGCAGT |
| Human | PHF8 | Histone Methylation | CTGCAGGCCTGGTGGACTGG |
| Human | PHF8 | Histone Methylation | GTATGTGGGAAAGCACATCC |
| Human | PHF8 | Histone Methylation | CACATCCTGGACATCTTTCG |
| Human | PHF8 | Histone Methylation | TGGTGGCAAAGCCTTGAACT |
| Human | PHF8 | Histone Methylation | AGTTCAAGGCTTTGCCACCA |
| Human | PHF8 | Histone Methylation | AGGCTTTGCCACCATGGACC |
| Human | PHF8 | Histone Methylation | GCTCATTAAAGATCTGGCCA |
| Human | PHF8 | Histone Methylation | AGCTCATTAAAGATCTGGCC |
| Human | CHP1 | Chromatin Readers | AAATCACTCGCCTCTACAGC |
| Human | CHP1 | Chromatin Readers | TCAGCCGGGAAGATTTCCAG |
| Human | CHP1 | Chromatin Readers | AGAGAATGGGACTCTCAGCC |
| Human | CHP1 | Chromatin Readers | GAGAGAATGGGACTCTCAGC |
| Human | CHP1 | Chromatin Readers | CGGTTCACCAGCCTGGACAA |
| Human | CHP1 | Chromatin Readers | TTTACCTGGTCCTCTCCCTC |
| Human | CHP1 | Chromatin Readers | GAGGACCAGGTAAACTTCCG |

| | | | |
|---|---|---|---|
| Human | CHP1 | Chromatin Readers | CCAAAGTTCGCATGAATCCA |
| Human | CHP1 | Chromatin Readers | GTTACAGGTGCTACGCATGA |
| Human | CHP1 | Chromatin Readers | GCACCTGTAACAGCTCATCA |
| Human | CHP1 | Chromatin Readers | CACCTGTAACAGCTCATCAC |
| Human | CHP1 | Chromatin Readers | CTCCCGTGATGAGCTGTTAC |
| Human | CHP1 | Chromatin Readers | AACAAATTCTGTGAAAGATA |
| Human | CHP1 | Chromatin Readers | TCAGGAGGCTGATCAGGATG |
| Human | CHP1 | Chromatin Readers | CGCAGACAGGACCATTCAGG |
| Human | CHP1 | Chromatin Readers | AGCTGGGCAGCATCGCAGAC |
| Human | CHP1 | Chromatin Readers | AATATCTCAGATGAGCAGCT |
| Human | CHP1 | Chromatin Readers | AAATATCTCAGATGAGCAGC |
| Human | CHP1 | Chromatin Readers | AGAGCTCGAGGAGATCAAGA |
| Human | RTF1 | Transcription Elongation | AAAGGGCTGCAACTACGGCA |
| Human | RTF1 | Transcription Elongation | AAAGAAGCGATGTTCTCTGC |
| Human | RTF1 | Transcription Elongation | ACTCTAAACGGAACACGCGT |
| Human | RTF1 | Transcription Elongation | GGTGGCACCAGAACAAACAA |
| Human | RTF1 | Transcription Elongation | ATATGTTCTTGGTCCGCTGG |
| Human | RTF1 | Transcription Elongation | AGATCAACTGAATGAGCTGG |
| Human | RTF1 | Transcription Elongation | CAGAGGCCCTGGACCGCCAG |
| Human | RTF1 | Transcription Elongation | CCAGCTCATTCAGTTGATCT |
| Human | RTF1 | Transcription Elongation | CTTGGTCCGCTGGCGGTCCA |
| Human | RTF1 | Transcription Elongation | TGACACTTCACCTTCCTCAG |
| Human | RTF1 | Transcription Elongation | CTGACACTTCACCTTCCTCA |
| Human | RTF1 | Transcription Elongation | TCTGACACTTCACCTTCCTC |
| Human | RTF1 | Transcription Elongation | ACGATGATGTTCGTGATGAG |
| Human | RTF1 | Transcription Elongation | TAACATCCCACAACAAGGAA |
| Human | RTF1 | Transcription Elongation | CAAGAAATCTCAAGCCATGG |
| Human | RTF1 | Transcription Elongation | AGACAAGAAATCTCAAGCCA |
| Human | HDAC7 | Histone Acetylation | GGGTTCTGTAGGGAATGCCG |
| Human | HDAC7 | Histone Acetylation | TCAGAGACTGTCTTGCGCAG |
| Human | HDAC7 | Histone Acetylation | CGGCATTCCCTACAGAACCC |
| Human | HDAC7 | Histone Acetylation | TGTTCCCAGAGCAACACCTG |
| Human | HDAC7 | Histone Acetylation | GTTCCCAGAGCAACACCTGA |
| Human | HDAC7 | Histone Acetylation | CCCGGGCACAGTCAGCAGCG |
| Human | HDAC7 | Histone Acetylation | CGAGTCATAGATCAGCCCTG |
| Human | HDAC7 | Histone Acetylation | CTCACGTCCAGGTGATCAAG |
| Human | HDAC7 | Histone Acetylation | GCACGTCCCAGTCTACAATG |
| Human | HDAC7 | Histone Acetylation | GCTAAAGAATGGTTTCGCTG |
| Human | HDAC7 | Histone Acetylation | GTAGACTGGGACGTGCACCA |
| Human | HDAC7 | Histone Acetylation | AGCCAGTGTGAGTGTCTCCG |
| Human | HDAC7 | Histone Acetylation | GCAGATCATTCAACAGCCAT |
| Human | HDAC7 | Histone Acetylation | CGGGTGCACAGTAAATACTG |

| | | | |
|---|---|---|---|
| Human | HDAC7 | Histone Acetylation | CCGAGGGCCTATCTTCAGCC |
| Human | HDAC7 | Histone Acetylation | CACAGAGAGGGACGCCAGTG |
| Human | HDAC7 | Histone Acetylation | GGATCACGGCCTCCAGAGAG |
| Human | HDAC7 | Histone Acetylation | CCTGGCTGAAGATAGGCCCT |
| Human | PRMT8 | Histone Methylation | CATAGACATTCTCCCACCAG |
| Human | PRMT8 | Histone Methylation | CAATGCCTGTTTGATAAAGG |
| Human | PRMT8 | Histone Methylation | GACAAGTGGCTGAAACCTGG |
| Human | PARP11 | Other Histone Modifications | GAAGTGGCACATGTTTCAGC |
| Human | PARP11 | Other Histone Modifications | ACATAGCTCCCGTCTTTGGA |
| Human | PARP11 | Other Histone Modifications | TGGTACCAGCAGTGAATTTG |
| Human | PARP11 | Other Histone Modifications | AGCATTTGTTCATTAATCTG |
| Human | DPF2 | Chromatin Readers | AGTGGTAACTGAGGCCTGGT |
| Human | DPF2 | Chromatin Readers | AAACGTTACAAGAACCGACC |
| Human | DPF2 | Chromatin Readers | GCATAGTGGTAACTGAGGCC |
| Human | DPF2 | Chromatin Readers | GGCAAGATGGATGCCCTGAG |
| Human | DPF2 | Chromatin Readers | CACTGCCGCCATCATCACGG |
| Human | DPF2 | Chromatin Readers | CGGCAGTGAAGACATACCGC |
| Human | DPF2 | Chromatin Readers | CAACTGGTCGTCATTCTCGG |
| Human | DPF2 | Chromatin Readers | GCAGCTCCAACTTCCTTCAG |
| Human | DPF2 | Chromatin Readers | CAGCTCCAACTTCCTTCAGG |
| Human | DPF2 | Chromatin Readers | GAGCAACTGGTCGTCATTCT |
| Human | DPF3 | Chromatin Readers | CAAGCGCTACAAGAACCGAC |
| Human | DPF3 | Chromatin Readers | GCATAGTGGTAGCTGAGCCC |
| Human | DPF3 | Chromatin Readers | AAGCGCTACAAGAACCGACC |
| Human | DPF3 | Chromatin Readers | AGAGCGTCCACAGTCTGCGC |
| Human | DPF3 | Chromatin Readers | ACATGAACAAGAAGAGTGGG |
| Human | DPF3 | Chromatin Readers | CTGGTGTCCTGCGCAGACTG |
| Human | DPF3 | Chromatin Readers | CCAACATGAACAAGAAGAGT |
| Human | DPF3 | Chromatin Readers | CTACTGTGACTTCTGCTTGG |
| Human | DPF3 | Chromatin Readers | TAGCTGGTCATCATTCTCTG |
| Human | DPF3 | Chromatin Readers | CAGCTCCAGCTTCCTTCTGG |
| Human | DPF3 | Chromatin Readers | GCAGCTCCAGCTTCCTTCTG |
| Human | HIF1AN | Histone Methylation | ATGTGACAGACAGAGCCAGG |
| Human | HIF1AN | Histone Methylation | TTTACATCCCAATGTACTGG |
| Human | HIF1AN | Histone Methylation | TGTGAACTTCTGGTATAAGG |
| Human | HIF1AN | Histone Methylation | TATTGAGAATGAGGAGCCTG |
| Human | HIF1AN | Histone Methylation | GAAGCTATAACTGCGCAACT |
| Human | HIF1AN | Histone Methylation | AGGATACACAAGATTTGTGT |
| Human | HIF1AN | Histone Methylation | AGAGGAGCTTATTGAGAATG |
| Human | HIF1AN | Histone Methylation | GGGTCGCTCTGACTCAGACG |
| Human | HIF1AN | Histone Methylation | TGAGTTCGTTGAGAAACTGC |
| Human | HIF1AN | Histone Methylation | AGAAACATTGAGAAGATGCT |

| | | | |
|---|---|---|---|
| Human | HIF1AN | Histone Methylation | CTTCTCAATGTTTCTCATTA |
| Human | CHD4 | Chromatin Readers | CAGCTGCAGTTCAGAAACCC |
| Human | CHD4 | Chromatin Readers | TATGGATGAGCCACCTTCTG |
| Human | CHD4 | Chromatin Readers | GAAGATCCTAATCTGGAAGT |
| Human | CHD4 | Chromatin Readers | AAGATCCTAATCTGGAAGTG |
| Human | CHD4 | Chromatin Readers | AATCACAGGGAGTTAATGAG |
| Human | CHD4 | Chromatin Readers | CCGAATCCTCAACCACAGTG |
| Human | CHD4 | Chromatin Readers | GGCCCTTCTTGTCCACACTG |
| Human | CHD4 | Chromatin Readers | GCAGAGCTATTGGAATCACA |
| Human | CHD4 | Chromatin Readers | GAATCACAGGGAGTTAATGA |
| Human | CHD4 | Chromatin Readers | TGAATTGATCACCATTGACA |
| Human | CHD4 | Chromatin Readers | TACAAGGAGGGTCATTCCAA |
| Human | CHD4 | Chromatin Readers | TGCCTCTTTCTTCATGCGGG |
| Human | CHD4 | Chromatin Readers | CATGTATGTCGTAACCTATG |
| Human | CHD4 | Chromatin Readers | GGCCTCCCGCATGAAGAAAG |
| Human | CHD4 | Chromatin Readers | GAGCACCCGGTGCATTGAAG |
| Human | CHD4 | Chromatin Readers | CAAGATGCTAGACCTGCTAG |
| Human | CHD4 | Chromatin Readers | AAATACGAACGCATCGATGG |
| Human | CHD4 | Chromatin Readers | ATTGACCGCTTCAATGCACC |
| Human | CHD4 | Chromatin Readers | AATGATGCTGACGCATCTAG |
| Human | CHD4 | Chromatin Readers | CAGACCATGTGGTAAGCACG |
| Human | CHD4 | Chromatin Readers | GACATGGAGAAGGCTCCCGA |
| Human | CHD4 | Chromatin Readers | GCAGACCATGTGGTAAGCAC |
| Human | CHD4 | Chromatin Readers | CGGGATCCAGGCAGACCATG |
| Human | CHD4 | Chromatin Readers | GACAGAGCCATTCACCGTTG |
| Human | CHD4 | Chromatin Readers | GGGACAGAGCCATTCACCGT |
| Human | CHD4 | Chromatin Readers | GACTGCCATTGAGTGTACAC |
| Human | CHD4 | Chromatin Readers | TAAACCATGGCTATGCCCGG |
| Human | CHD4 | Chromatin Readers | CTAGCCGGCATTATAAACCA |
| Human | CHD4 | Chromatin Readers | TTATAAACCATGGCTATGCC |
| Human | CHD4 | Chromatin Readers | GGGTCTGACAAACTGGCTGA |
| Human | DPF1 | Chromatin Readers | CACCGTGAATTGTAAACACG |
| Human | DPF1 | Chromatin Readers | CCGTGAATTGTAAACACGAG |
| Human | DPF1 | Chromatin Readers | CGAGGACCTCATCTCCTGTG |
| Human | DPF1 | Chromatin Readers | CAGCCGTGCGGACCTACCGC |
| Human | DPF1 | Chromatin Readers | TGCGGAACCTCCGAGAACGA |
| Human | DPF1 | Chromatin Readers | GCTGGCACCGTCGTTCTCGG |
| Human | DPF1 | Chromatin Readers | CAGCTCCAGCTCCCTTCCGG |
| Human | DPF1 | Chromatin Readers | GAACGACGGTGCCAGCTGGG |
| Human | DPF1 | Chromatin Readers | CGAGAACGACGGTGCCAGCT |
| Human | ELOA2 | Transcription Elongation | CATGACGGCAGACATCCTGG |
| Human | ELOA2 | Transcription Elongation | ACTCTCCGCCTTGCCCATGA |

| | | | |
|---|---|---|---|
| Human | ELOA2 | Transcription Elongation | GTCTGCCGTCATGGGCAAGG |
| Human | ELOA2 | Transcription Elongation | GATGTCTGCCGTCATGGGCA |
| Human | ELOA2 | Transcription Elongation | GCCAGGATGTCTGCCGTCAT |
| Human | ELOA2 | Transcription Elongation | CGCCTTATGATACTTCAAGG |
| Human | ELOA2 | Transcription Elongation | CGGATTGTTTCTAAGCACCT |
| Human | MLLT10 | Chromatin Readers | GCGGTGCATCAAGCTTGCTA |
| Human | MLLT10 | Chromatin Readers | TGCGTTTGCTCAGACGAGAG |
| Human | MLLT10 | Chromatin Readers | GGCATTGTTCAAGTACCCAC |
| Human | MLLT10 | Chromatin Readers | TATACAGGGCACAAACCACA |
| Human | MLLT10 | Chromatin Readers | GCCGATAATGTCCAATACTG |
| Human | MLLT10 | Chromatin Readers | TGCATGACATGTAATAAACA |
| Human | MLLT10 | Chromatin Readers | TAATAAACATGGATGTCGAC |
| Human | MLLT10 | Chromatin Readers | ACATGTCATGCAAGCACCAG |
| Human | MLLT10 | Chromatin Readers | GGACTGCTTTGTGAAGAAGA |
| Human | MLLT10 | Chromatin Readers | GTCTTATTATAACGATCATG |
| Human | MLLT10 | Chromatin Readers | TTGTGCCCTGTATATTCCAG |
| Human | MLLT10 | Chromatin Readers | TTGCTGAATGCAATACACAA |
| Human | MLLT10 | Chromatin Readers | TCCATCATGCCGATACTGCA |
| Human | MLLT10 | Chromatin Readers | CTATTGCCATACATATGAGA |
| Human | RAG1 | Chromatin Remodeling | GGCATCTGAGAATGCAGACC |
| Human | RAG1 | Chromatin Remodeling | GCTTACAGTTGGTCTCCACA |
| Human | RAG1 | Chromatin Remodeling | CTCAGATGCCTCAAAGTCAT |
| Human | RAG1 | Chromatin Remodeling | TCACCACATCTCAAGTCACA |
| Human | RAG1 | Chromatin Remodeling | GTGCAATGAGGAGGTCAGTT |
| Human | RAG1 | Chromatin Remodeling | CAGGAACAAGGTCATGCACA |
| Human | RAG1 | Chromatin Remodeling | GAGTCAGCGACAGAAGATGT |
| Human | RAG1 | Chromatin Remodeling | TCAGCGACAGAAGATGTTGG |
| Human | RAG1 | Chromatin Remodeling | AGCGACAGAAGATGTTGGCG |
| Human | RAG1 | Chromatin Remodeling | AACATCTTCTGTCGCTGACT |
| Human | RAG1 | Chromatin Remodeling | ACATCTGCAACACTGCCCGT |
| Human | RAG1 | Chromatin Remodeling | TCGGGAAGTAAACCTCACAT |
| Human | RAG1 | Chromatin Remodeling | CGGGAAGTAAACCTCACATG |
| Human | PRMT5 | Histone Methylation | CCAGTTCTCTAGCGTCACCA |
| Human | PRMT5 | Histone Methylation | CACCATCAGTACCTGGACAT |
| Human | PRMT5 | Histone Methylation | GTCCATCAGTGGCTGAAGCG |
| Human | PRMT5 | Histone Methylation | GGGTACCCGCATCCAGAACA |
| Human | PRMT5 | Histone Methylation | TGAACTGCACCTCCAACTTG |
| Human | PRMT5 | Histone Methylation | TCCAGAACATGGAAGAGTGA |
| Human | PRMT5 | Histone Methylation | TGGTATATGAGCGGCCTGTG |
| Human | PRMT5 | Histone Methylation | CAAGAAGGTGTGGTATGAGT |
| Human | PRMT5 | Histone Methylation | CCAAGAAGGTGTGGTATGAG |
| Human | PRMT5 | Histone Methylation | ATGGTATATGAGCGGCCTGT |

| | | | |
|---|---|---|---|
| Human | HDAC5 | Histone Acetylation | GTGGACTGGGACATTCACCA |
| Human | HDAC5 | Histone Acetylation | GAATGTCCCAGTCCACGATG |
| Human | HDAC5 | Histone Acetylation | AGCAAGTGCGAGCGGATCCG |
| Human | HDAC5 | Histone Acetylation | CCTTACAGCCTTCAGGACAG |
| Human | HDAC5 | Histone Acetylation | GGCCGAGACTGTGAGCGCCA |
| Human | HDAC5 | Histone Acetylation | GGCCATGGCGCTCACAGTCT |
| Human | HDAC5 | Histone Acetylation | GAGCGCCATGGCCTTGCTGT |
| Human | DHPS | Control | CTGTACAGCGCGCCCGAAGT |
| Human | DHPS | Control | GATGCCCATTCTGGACCAGA |
| Human | DHPS | Control | ATCACCATCTGGTCCAGAAT |
| Human | DHPS | Control | ATTACTGCAAGTTTGAGGAC |
| Human | DHPS | Control | CATTCTGGACCAGATGGTGA |
| Human | DHPS | Control | CATCACCATCTGGTCCAGAA |
| Human | MSL3 | Chromatin Readers | TAATGGTTGGAACAGAAGCT |
| Human | MSL3 | Chromatin Readers | AACAATAACATCAACAATCT |
| Human | MSL3 | Chromatin Readers | GAACAGAAGCTGGGATAGAT |
| Human | MSL3 | Chromatin Readers | GGAACAGAAGCTGGGATAGA |
| Human | MSL3 | Chromatin Readers | AAGCACTAACAGGAGCCAGG |
| Human | MSL3 | Chromatin Readers | GGCATAACGTGATGGTGACG |
| Human | TAF2 | Transcription Elongation | TCTTGTGCTCATGGTGATGG |
| Human | TAF2 | Transcription Elongation | GCTCATGGTGATGGTGGTGA |
| Human | TAF2 | Transcription Elongation | CATGGTGATGGTGGTGATGG |
| Human | CUL1 | Control | ATCTGGGACGACCTCAGAGC |
| Human | CUL1 | Control | AGTGTAGAGCTCCATATATC |
| Human | CUL1 | Control | GAAATTCTACACTCAACAAT |
| Human | CUL1 | Control | TGAAATTCTACACTCAACAA |
| Human | CUL1 | Control | GGCAGAGGCTCGTCTGCTTG |
| Human | CUL1 | Control | GGAGACACACATTCATAATC |
| Human | CUL1 | Control | GCTGCTCTTGATAAGGCTTG |
| Human | CUL1 | Control | CTTTGTGGCTGCTCTTGATA |
| Human | CUL1 | Control | GTATCGAGCCAGCAACTCAG |
| Human | CUL1 | Control | CAGTATCGAGCCAGCAACTC |
| Human | CUL1 | Control | AGACACACTCAATCAAGTGA |
| Human | CUL1 | Control | ATGCGAAGATGCTCGCCAAG |
| Human | CUL1 | Control | TCCAAGTTAAAGCAAGCTTG |
| Human | CUL1 | Control | CCAAGTTAAAGCAAGCTTGC |
| Human | CUL1 | Control | CAGCGCATGTTTCAAGACAT |
| Human | CUL1 | Control | CCCGCAAGCTTGCTTTAACT |
| Human | CUL1 | Control | AAACATGCGCTGAAGTTTAG |
| Human | CUL1 | Control | CCGCAAACTACTGATTCAGG |
| Human | CUL1 | Control | AGACCGCAAACTACTGATTC |
| Human | CUL1 | Control | CCGCCTGAATCAGTAGTTTG |

| | | | |
|---|---|---|---|
| Human | CUL1 | Control | TCAATGCATTTCTTGATCAC |
| Human | INO80E | Chromatin Remodeling | AGCGCAAAGGAAATTACTGA |
| Human | INO80E | Chromatin Remodeling | CGAGCACGAGTGCTTCCAGG |
| Human | INO80E | Chromatin Remodeling | AGCACTCGTGCTCGTAGATG |
| Human | INO80E | Chromatin Remodeling | AGGAGCTGAGGAAAGCGCAA |
| Human | INO80E | Chromatin Remodeling | CGGAAACTCAAGATGGCGGT |
| Human | INO80E | Chromatin Remodeling | ACCGCCATCTTGAGTTTCCG |
| Human | NSD1 | Histone Methylation | AAATGAAAGTTTCCAACCGG |
| Human | NSD1 | Histone Methylation | CTATCGGCAGTACTACGTGG |
| Human | NSD1 | Histone Methylation | AAGTTGGACGATACAGGTGG |
| Human | NSD1 | Histone Methylation | TGGAAGGAACAGCTCGAGGA |
| Human | NSD1 | Histone Methylation | ATGTTGGAAGGAACAGCTCG |
| Human | NSD1 | Histone Methylation | ATTCAGAGTCTATCCCACAG |
| Human | NSD1 | Histone Methylation | GTAGCTTTACAGTTGCAACG |
| Human | NSD1 | Histone Methylation | GTGAATGGAGATACCCGTGT |
| Human | NSD1 | Histone Methylation | GTCCCAAAGGAAACTATGCT |
| Human | NSD1 | Histone Methylation | TTGGCATCTCAGTCAATCCA |
| Human | NSD1 | Histone Methylation | GGATTGACTGAGATGCCAAG |
| Human | NSD1 | Histone Methylation | ATGTGAGGCTCAGTGCTGTG |
| Human | NSD1 | Histone Methylation | TATGTGAGGCTCAGTGCTGT |
| Human | NSD1 | Histone Methylation | AGGTGCCTTCTACCCTTGTG |
| Human | NSD1 | Histone Methylation | CCCTTGTTCTGCATAACAGT |
| Human | NSD1 | Histone Methylation | TGCTTTGTGTGCTCAGAAGG |
| Human | NSD1 | Histone Methylation | CTGAACATTGATATCCCTGA |
| Human | NSD1 | Histone Methylation | GGCAAGAATCACAGCACAGA |
| Human | NSD1 | Histone Methylation | TTGATATCCCTGAAGGAAAC |
| Human | NSD1 | Histone Methylation | ATCTGACCAAGCGACCAGCA |
| Human | NSD1 | Histone Methylation | TCAGATTGAGACAGTCTGCG |
| Human | NSD1 | Histone Methylation | GGACATTCCCATTTCCCTGC |
| Human | NSD1 | Histone Methylation | CAGGGAAATGGGAATGTCCG |
| Human | NSD1 | Histone Methylation | AATCTGACCAAGCGACCAGC |
| Human | NSD1 | Histone Methylation | GAGGGTTTGTCGCTTAGCTG |
| Human | NSD1 | Histone Methylation | ATAAGGTCAGAGACCTCGCT |
| Human | FOXA2 | DNA and RNA Modifications | TCAGCGTCAGCATCTTGTTG |
| Human | FOXA2 | DNA and RNA Modifications | TCGCCCGACAAGCCCGGCAA |
| Human | FOXA2 | DNA and RNA Modifications | CGAACATGTTGCCCGAGTCA |
| Human | FOXA2 | DNA and RNA Modifications | CCGGGCTTGTCGGGCGAGCG |
| Human | FOXA2 | DNA and RNA Modifications | GTTGCCCGAGTCAGGGTGCA |
| Human | FOXA2 | DNA and RNA Modifications | AGCTACTATGCAGAGCCCGA |
| Human | FOXA2 | DNA and RNA Modifications | GCAACATGAACGCCGGCCTG |
| Human | FOXA2 | DNA and RNA Modifications | CTCACGGAGGAGTAGCCCTC |
| Human | ATRIP | Control | GCGAGACTCACTACATCAGA |

| | | | |
|---|---|---|---|
| Human | ATRIP | Control | GAAGAAGTTCTCATTAAGAA |
| Human | ATRIP | Control | GGAGTTCAGACTGCAATGAT |
| Human | ATRIP | Control | TCAGACGGAATCCGTTCTAG |
| Human | ATRIP | Control | TTCCTTGTCACTGAGTGCTT |
| Human | ATRIP | Control | AACCCAAGCACTCAGTGACA |
| Human | ATRIP | Control | ATGCAGAGATGAATGAATTA |
| Human | WAC | Other Histone Modifications | GTCGAACAGAAGTTTCACAA |
| Human | WAC | Other Histone Modifications | TCGAACAGAAGTTTCACAAT |
| Human | WAC | Other Histone Modifications | TCTGAGCATATTAGCTCTTC |
| Human | WAC | Other Histone Modifications | CTGAGCATATTAGCTCTTCT |
| Human | UBR7 | Chromatin Readers | CTTTACTGAGCCCTGAGAGT |
| Human | UBR7 | Chromatin Readers | GCTTGCAGTTATGAATGTCA |
| Human | UBR7 | Chromatin Readers | CAAGCTAAACAAATTCCTGC |
| Human | UBR7 | Chromatin Readers | TGGCACCAAGATGCCTTCCA |
| Human | UBR7 | Chromatin Readers | AGTGCGTAGTCTGTGAAGAC |
| Human | TDRD7 | Chromatin Readers | AGAATGGAGTAGCATTACCC |
| Human | TDRD7 | Chromatin Readers | TCTGAGAAGTGTGCCAGCAG |
| Human | TDRD7 | Chromatin Readers | CTCAGATAGGCTTCTAGTGT |
| Human | TDRD7 | Chromatin Readers | CTGGCCTCATATTTGCACGG |
| Human | TDRD7 | Chromatin Readers | TTCTCCACCGTGCAAATATG |
| Human | TDRD7 | Chromatin Readers | ATACTAAACAAGCATAACAA |
| Human | TDRD7 | Chromatin Readers | ACGGTGGAGAAACCTTGCAG |
| Human | TDRD7 | Chromatin Readers | AAATGCTTTCGGAAGTGCAC |
| Human | TDRD7 | Chromatin Readers | GCTGCATACATCAGAAAGTG |
| Human | TDRD7 | Chromatin Readers | GTACAAAGTGAAATTCCCTG |
| Human | TDRD7 | Chromatin Readers | TACATTTCCTCAAATGCTTT |
| Human | TDRD7 | Chromatin Readers | GGACGCCTGGTTACGGGCAC |
| Human | TDRD7 | Chromatin Readers | GTTGCTGGCCGTAAATGCCG |
| Human | TDRD7 | Chromatin Readers | GATGACCTGTGCCCGTAACC |
| Human | TDRD7 | Chromatin Readers | CCGTAACCAGGCGTCCTCCT |
| Human | TDRD7 | Chromatin Readers | AGCTCAGGGAAATTCCACCT |
| Human | TDRD7 | Chromatin Readers | GATGTTCAGTTCCTGGACTC |
| Human | TDRD7 | Chromatin Readers | GGAACTGAACATCAAGAGCC |
| Human | TDRD7 | Chromatin Readers | AGCGCTCGTATGGTCCACCA |
| Human | TDRD7 | Chromatin Readers | GAAGTGCAACCAGTGGTCTG |
| Human | HDAC6 | Histone Acetylation | GGCCTGACAGTAGTAACACC |
| Human | HDAC6 | Histone Acetylation | TCTTGCTATCAGGTCTACTG |
| Human | HDAC6 | Histone Acetylation | GCTACATCGACCTGTCAGCC |
| Human | HDAC6 | Histone Acetylation | AGTAGTAACACCAGGCTGAC |
| Human | HDAC6 | Histone Acetylation | AGCCTGGTGTTACTACTGTC |
| Human | HDAC6 | Histone Acetylation | ACAGAAACACCGCATCCGGA |
| Human | HDAC6 | Histone Acetylation | CCGGGAAGCTGTCATCCCAG |

| | | | |
|---|---|---|---|
| Human | HDAC6 | Histone Acetylation | TGGCGGCCATCTCACCCTTG |
| Human | HDAC6 | Histone Acetylation | CAATGTGCCTTGGAACCAGG |
| Human | HDAC6 | Histone Acetylation | AATGTGCCTTGGAACCAGGT |
| Human | HDAC6 | Histone Acetylation | GGGACACATATAGCACACTG |
| Human | HDAC6 | Histone Acetylation | ATTATCCTTATCCTAGAGGG |
| Human | HDAC6 | Histone Acetylation | TGTCACAGTGCTGAGTACGT |
| Human | HDAC6 | Histone Acetylation | CAGGGACACATATAGCACAC |
| Human | HDAC6 | Histone Acetylation | TGGTCTCAGTGATTGAGGCC |
| Human | HDAC6 | Histone Acetylation | GGTCTCAGTGATTGAGGCCA |
| Human | BRD4 | Chromatin Readers | TGTTACATCTACAACAAGCC |
| Human | BRD4 | Chromatin Readers | TGATCTTATAGTAATCAGGG |
| Human | BRD4 | Chromatin Readers | TAATGATCTTATAGTAATCA |
| Human | BRD4 | Chromatin Readers | CAACTATTACTGGAATGCTC |
| Human | BRD4 | Chromatin Readers | CACAATCAAGTCTAAACTGG |
| Human | BRD4 | Chromatin Readers | CACCAAACTCCTGAGCATCA |
| Human | BRD4 | Chromatin Readers | CTGAGCATCACGGTACTCAC |
| Human | BRD4 | Chromatin Readers | CCTGAGCATCACGGTACTCA |
| Human | BRD4 | Chromatin Readers | TGATGTCACAGTAGTCGTGT |
| Human | BRD4 | Chromatin Readers | AGTCGATTTCAATCTCGTCG |
| Human | BRD4 | Chromatin Readers | GCGCTCCAGCTCACGCAGTG |
| Human | BRD4 | Chromatin Readers | GTGCAAGCCTATGTCCTATG |
| Human | BRD4 | Chromatin Readers | AAAGTCGATTTCAATCTCGT |
| Human | BRD4 | Chromatin Readers | GCTATGTCACCTCCTGTTTG |
| Human | BRD4 | Chromatin Readers | CAGCAGCAGCGATAGCTCCT |
| Human | BRD4 | Chromatin Readers | GCTATCGCTGCTGCTGTCGC |
| Human | BRD4 | Chromatin Readers | TTCTTTGGTTTGTTCTGCTG |
| Human | BRD4 | Chromatin Readers | GGAGCGCATGAGGAGCCGAG |
| Human | BRD4 | Chromatin Readers | GGAGGTGACCGGTTGAGTAG |
| Human | BRD4 | Chromatin Readers | CAACAGCAATGTGAGCAAGA |
| Human | BRD4 | Chromatin Readers | ACTGGAGCTCTCCGACTCTG |
| Human | BRD4 | Chromatin Readers | GGACGAGAAGCCCTTCATCT |
| Human | BRD4 | Chromatin Readers | GGGCTTCTCGTCCTCAGAGT |
| Human | BRD4 | Chromatin Readers | GTCAGAGGAGCTGGACTCAC |
| Human | BRD4 | Chromatin Readers | AGTCTTAATGGCAGAAGCTC |
| Human | BRD4 | Chromatin Readers | GAGCAGCTCAAGTGCTGCAG |
| Human | BRD4 | Chromatin Readers | CGAACACATCCTGGAGCTTG |
| Human | BRD4 | Chromatin Readers | GAACACATCCTGGAGCTTGC |
| Human | BRD4 | Chromatin Readers | ATCCTGGAGCTTGCGGGCCA |
| Human | BRD4 | Chromatin Readers | GGCCATGGCCCGCAAGCTCC |
| Human | UBE2E1 | Other Histone Modifications | CAAGAGAATTCAGAAGGAGC |
| Human | UBE2E1 | Other Histone Modifications | GAGAATTCAGAAGGAGCTGG |
| Human | UBE2E1 | Other Histone Modifications | CAGCTCCTTCTGAATTCTCT |

| | | | |
|---|---|---|---|
| Human | UBE2E1 | Other Histone Modifications | ATGACAGAATGGCCAGACAG |
| Human | MRPL20 | Control | CTGTAATTCGATTAATCCAG |
| Human | MRPL20 | Control | TGTAATTCGATTAATCCAGA |
| Human | MRPL20 | Control | TTTCACAAAGGCTCGAATCA |
| Human | MRPL20 | Control | TCCTTCGTGTCGCCTCCTAC |
| Human | MRPL20 | Control | GTGTCGCCTCCTACTGGCCA |
| Human | MRPL20 | Control | TGGCTGCCTTGGCCAGTAGG |
| Human | MRPL20 | Control | CTTTGGCTGCCTTGGCCAGT |
| Human | PRKCB | Other Histone Modifications | GTAGGGATCTGACAGGCCAT |
| Human | PRKCB | Other Histone Modifications | GCTGATAAGCAATTATCTCG |
| Human | PRKCB | Other Histone Modifications | CTGATAAGCAATTATCTCGG |
| Human | PRKCB | Other Histone Modifications | GATGAAGAGTTTATCAGTGG |
| Human | PRKCB | Other Histone Modifications | ATGATGAAGAGTTTATCAGT |
| Human | PRKCB | Other Histone Modifications | TAAACTCTTCATCATGAACT |
| Human | PRKCB | Other Histone Modifications | TGATGAAGAGTTTATCAGTG |
| Human | PRKCB | Other Histone Modifications | AGAGTTCACCAGACAGCCTG |
| Human | PRKCB | Other Histone Modifications | CACTGCACCGACTTCATCTG |
| Human | PRKCB | Other Histone Modifications | GGGCTGCTTGAAGAAGCGGG |
| Human | PRKCB | Other Histone Modifications | GCACCACAAAGCAGCAAACT |
| Human | PRKCB | Other Histone Modifications | TGTATGGACTCATCCACCAG |
| Human | PRKCB | Other Histone Modifications | CTGTATGGACTCATCCACCA |
| Human | LMNB1 | DNA and RNA Modifications | TGGCATTCTCAAGTTTGGCA |
| Human | LMNB1 | DNA and RNA Modifications | CGGTCTCGTAGAGCGCCTTG |
| Human | LMNB1 | DNA and RNA Modifications | ATTCCTTCTTAGCATAGTTG |
| Human | LMNB1 | DNA and RNA Modifications | GAACAGGATCAACCAATGGG |
| Human | LMNB1 | DNA and RNA Modifications | TCTGAACAGGATCAACCAAT |
| Human | LMNB1 | DNA and RNA Modifications | GGAGCCGCGACAGGCGCGTG |
| Human | LMNB1 | DNA and RNA Modifications | GCCCATCCGCGGCGGCACGG |
| Human | LMNB1 | DNA and RNA Modifications | CGGATGGGCAGCCGCGCTGG |
| Human | LMNB1 | DNA and RNA Modifications | TGCCCATCCGCGGCGGCACG |
| Human | LMNB1 | DNA and RNA Modifications | ATGCTCGGGATACTGTCACA |
| Human | LMNB1 | DNA and RNA Modifications | CGTAGTGTACGTACAACTAG |
| Human | LMNB1 | DNA and RNA Modifications | TACGTACAACTAGAGGAAAG |
| Human | NSL1 | Chromatin Remodeling | ACCGAAATGCTACAACTGTG |
| Human | NSL1 | Chromatin Remodeling | GCCGCACAGTTGTAGCATTT |
| Human | NSL1 | Chromatin Remodeling | CCGCTCTGCGAGATGCGCAG |
| Human | NSL1 | Chromatin Remodeling | CCACTGCGCATCTCGCAGAG |
| Human | NSL1 | Chromatin Remodeling | CACTGCGCATCTCGCAGAGC |
| Human | NSL1 | Chromatin Remodeling | GCAGTATCCCAGAAAGATCC |
| Human | NSL1 | Chromatin Remodeling | ACACATTCCAGGATCTTTCT |
| Human | NSL1 | Chromatin Remodeling | GACACATTCCAGGATCTTTC |
| Human | NSL1 | Chromatin Remodeling | TAATGCAGGCAAGGACTTCA |

| | | | |
|---|---|---|---|
| Human | DNMT1 | DNA and RNA Modifications | AAAGAAGAATTATCCGAGGA |
| Human | DNMT1 | DNA and RNA Modifications | TAAAGAAGAATTATCCGAGG |
| Human | DNMT1 | DNA and RNA Modifications | CCAAATCTTTGAGCCGCCTG |
| Human | DNMT1 | DNA and RNA Modifications | TTTAGCCAGGTAGCCCTCCT |
| Human | DNMT1 | DNA and RNA Modifications | GCTGTATCTAGCAAGGGTCA |
| Human | DNMT1 | DNA and RNA Modifications | GTGACCCTTGCTAGATACAG |
| Human | DNMT1 | DNA and RNA Modifications | TAGCAAGGGTCACGGCGCTG |
| Human | DNMT1 | DNA and RNA Modifications | TCTTTGCATTATAGGCCTCG |
| Human | DNMT1 | DNA and RNA Modifications | GAGTACACCTGGACGCACTC |
| Human | DNMT1 | DNA and RNA Modifications | GCCAATTCGGTAGGGCTCAG |
| Human | DNMT1 | DNA and RNA Modifications | AAATACTCCGACTACATCAA |
| Human | DNMT1 | DNA and RNA Modifications | CTGAGCCCTACCGAATTGGC |
| Human | DNMT1 | DNA and RNA Modifications | CTGTAAGGACATGAGTGCAT |
| Human | DNMT1 | DNA and RNA Modifications | GTAGTCGCAGTAGCTGAGGA |
| Human | DNMT1 | DNA and RNA Modifications | TGTCAGAGATGCCTGCTTGG |
| Human | DNMT1 | DNA and RNA Modifications | AGGTGCACTGATAGCCCATG |
| Human | DNMT1 | DNA and RNA Modifications | TGTGTCAGCAGCCTGAGTGT |
| Human | DNMT1 | DNA and RNA Modifications | CCGGCGATGTGGCGTCTGTG |
| Human | DNMT1 | DNA and RNA Modifications | GCTTTACATTTCCCACACTC |
| Human | DNMT1 | DNA and RNA Modifications | GCCACATCGCCGGCGCTTAA |
| Human | DNMT1 | DNA and RNA Modifications | AGCAGGCTTGCCAAGAGCGG |
| Human | DNMT1 | DNA and RNA Modifications | AAGATCTAGCTGCCAAACGG |
| Human | DNMT1 | DNA and RNA Modifications | CCAAAGATCTAGCTGCCAAA |
| Human | DNMT1 | DNA and RNA Modifications | CGGTGCTCATGCTTACAACC |
| Human | DNMT1 | DNA and RNA Modifications | GATCAACAAGATCGAGACCA |
| Human | DNMT1 | DNA and RNA Modifications | GATGACCCATCTCTTGAAGG |
| Human | DNMT1 | DNA and RNA Modifications | GCAGCACCCACCAGACGCGG |
| Human | DNMT1 | DNA and RNA Modifications | CAGAATGTATTCGGCAAATG |
| Human | DNMT1 | DNA and RNA Modifications | TATTGGGACACCTCCGCTCT |
| Human | DNMT1 | DNA and RNA Modifications | AGAGCGGAGGTGTCCCAATA |
| Human | DNMT1 | DNA and RNA Modifications | GAGAAGCCGTCAAGACTGAT |
| Human | DNMT1 | DNA and RNA Modifications | GGAGAAGCCGTCAAGACTGA |
| Human | DNMT1 | DNA and RNA Modifications | GAAGAAGAGTTACTATAAGA |
| Human | DNMT1 | DNA and RNA Modifications | AAGAATCGCATCTCTTGGGT |
| Human | DNMT1 | DNA and RNA Modifications | TTTAGCTGAGGCACTCTCTC |
| Human | DNMT1 | DNA and RNA Modifications | AGATCAAGCTGCCCAAGCTG |
| Human | DNMT1 | DNA and RNA Modifications | GGCAGCTTGATCTCTATCTC |
| Human | DNMT1 | DNA and RNA Modifications | GTGCCCGTCTGGCTGAGATG |
| Human | SMN1 | Chromatin Readers | ACCTGTGTTGTGGTTTACAC |
| Human | SMN1 | Chromatin Readers | TCCAGTGTAAACCACAACAC |
| Human | SMN1 | Chromatin Readers | TAAGAGAGAAACCTGTGTTG |
| Human | SMN1 | Chromatin Readers | TCTGCCATTTGGTCAGAAGA |

| | | | |
|---|---|---|---|
| Human | SMN1 | Chromatin Readers | CGCTCTGGCCTGTGCCGCGC |
| Human | SMN1 | Chromatin Readers | GTGCTGTTCCGGCGCGGCAC |
| Human | SMN1 | Chromatin Readers | AGAGCGATGATTCTGACATT |
| Human | SMN1 | Chromatin Readers | ATTAGCTACTTCACAGATTG |
| Human | SMN1 | Chromatin Readers | TCAAGCCCAAATCTGCTCCA |
| Human | SMN1 | Chromatin Readers | CAGTCTTGGCCCTGGCATGG |
| Human | SMN1 | Chromatin Readers | TTATTAGCTACTTCACAGAT |
| Human | SMN1 | Chromatin Readers | TCAGCATCATCAAGAGAATC |
| Human | SMN1 | Chromatin Readers | CTGGACATATGGGAGGTGGT |
| Human | SMN1 | Chromatin Readers | TCAAGAGAATCTGGACATAT |
| Human | SMN1 | Chromatin Readers | TACATGAGTGGCTATCATAC |
| Human | NOC4L | Control | ACAGCCCGTTCAAGGCCAAG |
| Human | NOC4L | Control | GTTCAAGGCCAAGAGGCTGA |
| Human | NOC4L | Control | TCAGCCTCTTGGCCTTGAAC |
| Human | NOC4L | Control | CGTTCAAGGCCAAGAGGCTG |
| Human | NOC4L | Control | GGTGGGAGGAGGACAGGAAG |
| Human | NOC4L | Control | AGGACAGGAAGAGGTCAGCC |
| Human | NOC4L | Control | ACAGGAAGAGGTCAGCCAGG |
| Human | NOC4L | Control | GGACCATGAGCAGGGCCTCA |
| Human | NOC4L | Control | TGAAAGGCAGGACCATGAGC |
| Human | NOC4L | Control | ACCATGAGCAGGGCCTCAGG |
| Human | TAF3 | Chromatin Readers | TGATTCCAACACAGGGCCAG |
| Human | TAF3 | Chromatin Readers | TCGTCAGGCTTGTTACACCC |
| Human | TAF3 | Chromatin Readers | CAGTCGTCACACCCAATCAT |
| Human | TAF3 | Chromatin Readers | GGAAACAAAGTATCCCTGGA |
| Human | TAF3 | Chromatin Readers | AATTGAAAGATGGACTTGTG |
| Human | TAF3 | Chromatin Readers | TTCTTTACCTCCACGGAGGA |
| Human | TAF3 | Chromatin Readers | CAAGCTGCCTTCCTCCGTGG |
| Human | TAF3 | Chromatin Readers | GTCTTCGGCCTGTTCTGCGA |
| Human | TAF3 | Chromatin Readers | GCGCAGGGCTGACGAGCATG |
| Human | TAF3 | Chromatin Readers | GGCGCAGGGCTGACGAGCAT |
| Human | TAF3 | Chromatin Readers | GGCCTGTTCTGCGAGGGCGC |
| Human | TP53BP1 | Chromatin Readers | CTGGTAAGCTGTGTTACACT |
| Human | TP53BP1 | Chromatin Readers | AGCCGCTCCAAACTGCCAGA |
| Human | TP53BP1 | Chromatin Readers | GGCCACAACCAGTGACAAGT |
| Human | TP53BP1 | Chromatin Readers | ACTGGTAAGCTGTGTTACAC |
| Human | TP53BP1 | Chromatin Readers | AAACTGCCAGATGGTCCTAC |
| Human | TP53BP1 | Chromatin Readers | TTCAGCACACTTCAGCACCG |
| Human | TP53BP1 | Chromatin Readers | ACCCACTCTTGTGACACCAC |
| Human | TP53BP1 | Chromatin Readers | TGAATGGTGCTGCTTCACAG |
| Human | TP53BP1 | Chromatin Readers | AGGCTGCACCACCAGTCATG |
| Human | TP53BP1 | Chromatin Readers | TGAGTATTTCAGTGCAGGAG |

| | | | |
|---|---|---|---|
| Human | TP53BP1 | Chromatin Readers | TTCAGTGCAGGAGTGGTGAA |
| Human | TP53BP1 | Chromatin Readers | ACAGACTGAGAGAGCAGTAT |
| Human | TP53BP1 | Chromatin Readers | TATCAGCTTAGACAATTTGG |
| Human | TP53BP1 | Chromatin Readers | AGCAGTAACACCTCTTACAA |
| Human | TP53BP1 | Chromatin Readers | ATATCTGCTGCCTTTGTAAG |
| Human | TP53BP1 | Chromatin Readers | AGCTTAGACAATTTGGTGGA |
| Human | TP53BP1 | Chromatin Readers | AGATATCAGCTTAGACAATT |
| Human | G2E3 | Chromatin Readers | TGAGACATCCAATTAACTCA |
| Human | G2E3 | Chromatin Readers | GATGACTGTCCTAATAAATA |
| Human | G2E3 | Chromatin Readers | AATGATAACTTCGTTTACAT |
| Human | G2E3 | Chromatin Readers | CGAAGTTATCATTTCCCATG |
| Human | G2E3 | Chromatin Readers | ATTCTCTCTGAAGTCCACAT |
| Human | G2E3 | Chromatin Readers | ACAGGTTCAAGCAATAAATG |
| Human | G2E3 | Chromatin Readers | CTTCGTAATATGTTATAACT |
| Human | G2E3 | Chromatin Readers | CTCAATAAATTCCAAGCAAA |
| Human | G2E3 | Chromatin Readers | GTCCTTGTTGTAAGAACGCT |
| Human | G2E3 | Chromatin Readers | CTGCTCCTCATTACGGTCAT |
| Human | G2E3 | Chromatin Readers | TTGCTCCCATGACCGTAATG |
| Human | G2E3 | Chromatin Readers | TAATGCACCTGATAGCAAAT |
| Human | G2E3 | Chromatin Readers | ATAATGCACCTGATAGCAAA |
| Human | SIN3A | DNA and RNA Modifications | ACACACGACTAATCACTCCT |
| Human | SIN3A | DNA and RNA Modifications | CACACGACTAATCACTCCTG |
| Human | SIN3A | DNA and RNA Modifications | TTTAAAGAGACGAGCAACCT |
| Human | SIN3A | DNA and RNA Modifications | ATGGCATGATTAAACTCCAC |
| Human | SIN3A | DNA and RNA Modifications | CACTCCAGCATTGACAGAGC |
| Human | SIN3A | DNA and RNA Modifications | AGAAATGCCAAGGAAGCTGG |
| Human | SIN3A | DNA and RNA Modifications | CAGATCCGCAGACATCCTAC |
| Human | SIN3A | DNA and RNA Modifications | AGTGACAATCTCGTTGGGAA |
| Human | SIN3A | DNA and RNA Modifications | TTGAGAGTATCTATGATGAG |
| Human | SIN3A | DNA and RNA Modifications | GTAGAGTCCTCAGACCACGA |
| Human | SIN3A | DNA and RNA Modifications | GAGGCAAGAGCAGGCTACGG |
| Human | SIN3A | DNA and RNA Modifications | GGATTTGTTCCCAAGCCGAA |
| Human | SIN3A | DNA and RNA Modifications | TCTTCTACATCAACATCCAT |
| Human | SIN3A | DNA and RNA Modifications | GCTACGTCTCAAAGAACCTA |
| Human | SIN3A | DNA and RNA Modifications | TTCTTTGAGACGTAGCTGAA |
| Human | SIN3A | DNA and RNA Modifications | GAATGGGAACGGGAAGTGCT |
| Human | IWS1 | Transcription Elongation | CAGAAAGTGAGGACCCTCCG |
| Human | IWS1 | Transcription Elongation | TGAGTCACTGATACGAGGTT |
| Human | IWS1 | Transcription Elongation | GAGTCACTGATACGAGGTTT |
| Human | IWS1 | Transcription Elongation | TATCAGTGACTCAGAAAGTG |
| Human | IWS1 | Transcription Elongation | AATCAGTGATTCGGAAAGTG |
| Human | IWS1 | Transcription Elongation | CACTTTCCGAATCACTGATT |

| | | | |
|---|---|---|---|
| Human | IWS1 | Transcription Elongation | ACTTTCCGAATCACTGATTC |
| Human | IWS1 | Transcription Elongation | CAGGGAAATTAATCAATGAG |
| Human | IWS1 | Transcription Elongation | GGCAAACTCCTATCTGGTAG |
| Human | IWS1 | Transcription Elongation | GGTCTAACAAGGACATGGCA |
| Human | IWS1 | Transcription Elongation | CAGGAGACCCTGAAGCATAG |
| Human | IWS1 | Transcription Elongation | TCACTGGCCTGGTTCCTTGG |
| Human | IWS1 | Transcription Elongation | ATCACTGGCCTGGTTCCTTG |
| Human | IWS1 | Transcription Elongation | AAGGAACCAGGCCAGTGATT |
| Human | IWS1 | Transcription Elongation | CGAAGAATGAACAGCACTGG |
| Human | IWS1 | Transcription Elongation | AAGCGCAGACGGAACCGCGA |
| Human | IWS1 | Transcription Elongation | CGACGAAGAATGAACAGCAC |
| Human | IWS1 | Transcription Elongation | GTTTCAGGCGACCTCCAAGA |
| Human | IWS1 | Transcription Elongation | ATGAGGATGTCAATCAGCAT |
| Human | IWS1 | Transcription Elongation | GATGCTGTATCAGACAAGTC |
| Human | IWS1 | Transcription Elongation | TCACTCATTCGAGGTTTGGG |
| Human | CENPN | Chromatin Remodeling | CTGTTGCTGAGTTCATCAAG |
| Human | BRD3 | Chromatin Readers | GAGTGCAAGCGAATGTATGC |
| Human | BRD3 | Chromatin Readers | TTCAATTTGATTGCGTCCAC |
| Human | BRD3 | Chromatin Readers | AATGTAACAATTTGTAAACA |
| Human | BRD3 | Chromatin Readers | AGCAATTCGAGAACATCAGC |
| Human | BRD3 | Chromatin Readers | AGTCAATTTCTATCTCGTCG |
| Human | BRD3 | Chromatin Readers | AAAGTCAATTTCTATCTCGT |
| Human | BRD3 | Chromatin Readers | TAGTGCACATCATCCAATCT |
| Human | BRD3 | Chromatin Readers | TTTCATCGTAGCTCATGGGC |
| Human | BRD3 | Chromatin Readers | GGAGCTGCAGGAGCAGCTGA |
| Human | BRD3 | Chromatin Readers | GAAGGAGAAGCACAAAGTGA |
| Human | BRD3 | Chromatin Readers | GGCGGCCAGCTGCTCGTGCA |
| Human | BRD3 | Chromatin Readers | GAAGCACAAAGTGAAGGCCG |
| Human | BRD3 | Chromatin Readers | ACAGGCAGCCAAGTCGAAAG |
| Human | BRD3 | Chromatin Readers | AAGCGTCTGCAGGATGTCAG |
| Human | BRD3 | Chromatin Readers | AGCTAGCTCCTCTTTCGACT |
| Human | BRD3 | Chromatin Readers | AGCGTCTGCAGGATGTCAGC |
| Human | BRD3 | Chromatin Readers | GAAGGCCAACAGCACGACCA |
| Human | BRD3 | Chromatin Readers | GCCGGCAGACAGCTGAAGAA |
| Human | BRD3 | Chromatin Readers | GAAGAAGGCTCCTGCCAAGA |
| Human | BRD3 | Chromatin Readers | CTTCAGCTGTCTGCCGGCCG |
| Human | BRD3 | Chromatin Readers | GGAAGAGAAGAAGGCCAAGG |
| Human | BRD3 | Chromatin Readers | GCCGGGCTTCTCTTTCCGGG |
| Human | BRD3 | Chromatin Readers | GCTGCTGCTCCCAGACTCGG |
| Human | BRD3 | Chromatin Readers | CCCGCCCGGAAAGAGAAGCC |
| Human | BRD3 | Chromatin Readers | GCCCGGCTCAGCACCCTCAG |
| Human | BRD3 | Chromatin Readers | CCCGGCTCAGCACCCTCAGG |

| | | | |
|---|---|---|---|
| Human | BRD3 | Chromatin Readers | GGATGCTGTCGCAGTAGCGT |
| Human | BRD3 | Chromatin Readers | CTACTGCGACAGCATCCTCA |
| Human | BRD3 | Chromatin Readers | GCTACTGCGACAGCATCCTC |
| Human | BRD3 | Chromatin Readers | TCCAGGACGTGTTTGAGATG |
| Human | BRD3 | Chromatin Readers | CAAACACGTCCTGGAGCTTC |
| Human | BRD3 | Chromatin Readers | AAACACGTCCTGGAGCTTCC |
| Human | BRD3 | Chromatin Readers | GGCCATGGCCCGGAAGCTCC |
| Human | BRD3 | Chromatin Readers | GTCCTGGAGCTTCCGGGCCA |
| Human | PARP1 | Other Histone Modifications | GGAAGTCCTCAGACACAACT |
| Human | PARP1 | Other Histone Modifications | GGCCATGATTGAGAAACTCG |
| Human | PARP1 | Other Histone Modifications | AGACAGTGTGCAGGCCAAGG |
| Human | PARP1 | Other Histone Modifications | GTCGATCTCATACTCCACCA |
| Human | PARP1 | Other Histone Modifications | CCTCAGTGAGGTCCAGCAGG |
| Human | PARP1 | Other Histone Modifications | CTGTCTGCATTGTTCAGGAG |
| Human | PARP1 | Other Histone Modifications | ACCTTCAGAAGATGCCCTTG |
| Human | PARP1 | Other Histone Modifications | CAGTTCATACATGTTTCCAA |
| Human | PARP1 | Other Histone Modifications | ACCAAACATGTAGCCTGTCA |
| Human | PARP1 | Other Histone Modifications | TCAGTTCATACATGTTTCCA |
| Human | PARP1 | Other Histone Modifications | CCAAACATGTAGCCTGTCAC |
| Human | PARP1 | Other Histone Modifications | GATTTCTCGGAATTCCTTTG |
| Human | PARP1 | Other Histone Modifications | CATGGGCGACTGCACCATGA |
| Human | PARP1 | Other Histone Modifications | ATCTTGGACCGAGTAGCTGA |
| Human | PARP1 | Other Histone Modifications | TGGCTTATCTGCTGAAGCAG |
| Human | PARP1 | Other Histone Modifications | GCGCTAAGAACAACTCCTGA |
| Human | PARP1 | Other Histone Modifications | TAATTTCATGAAGTGCTCAA |
| Human | MBD6 | DNA and RNA Modifications | GTGCTCTACATCAGTCCAAG |
| Human | MBD6 | DNA and RNA Modifications | ACTTGCAGGTCCCATCGCTG |
| Human | MBD6 | DNA and RNA Modifications | TAGCAATCGAAATACAGTAG |
| Human | KAT5 | Histone Acetylation | ATCTCATTGATGGTGATCTG |
| Human | KAT5 | Histone Acetylation | GGTGAGTTCCTGTGGGTACG |
| Human | KAT5 | Histone Acetylation | AGAAGTACCACGGCTTGAGG |
| Human | KAT5 | Histone Acetylation | CGGTAAATCTCATTGCCTGG |
| Human | KAT5 | Histone Acetylation | GTGTCTGCACTTCACTCCCA |
| Human | KAT5 | Histone Acetylation | GGGAGTGAAGTGCAGACACT |
| Human | KAT5 | Histone Acetylation | TCCCAAGGACTGGAGCAAGA |
| Human | KAT5 | Histone Acetylation | CTCCCAAGGACTGGAGCAAG |
| Human | KAT5 | Histone Acetylation | CCAAGGACTGGAGCAAGAGG |
| Human | KAT5 | Histone Acetylation | AAGCGACGCCTTTCTGACAA |
| Human | KAT5 | Histone Acetylation | GAAGCGACGCCTTTCTGACA |
| Human | KAT5 | Histone Acetylation | CCTTGAGACCCTTGTCAGAA |
| Human | KAT5 | Histone Acetylation | GAAGGTGTCGGAGCTGAAAG |
| Human | KAT5 | Histone Acetylation | CTGGAGTCGCTCCATGAGCT |

| | | | |
|---|---|---|---|
| Human | HNRNPU | Transcription Elongation | TACATGAAGTTCGTATTGGC |
| Human | HNRNPU | Transcription Elongation | GACATACATGAAGTTCGTAT |
| Human | HNRNPU | Transcription Elongation | GGCTGGTCACTAACTACAAG |
| Human | HNRNPU | Transcription Elongation | GAACTCTCGTATGCTAAGAA |
| Human | HNRNPU | Transcription Elongation | TATTATGGATAAGATGATGG |
| Human | HNRNPU | Transcription Elongation | TGCTGCATGTTTAGTAACCC |
| Human | HNRNPU | Transcription Elongation | TAAATTTCCCAAGACACTGG |
| Human | HNRNPU | Transcription Elongation | GTGATGATGATTGGCTTGCC |
| Human | HNRNPU | Transcription Elongation | ATATCCGCCACGATTCCCAG |
| Human | HNRNPU | Transcription Elongation | TATATCCGCCACGATTCCCA |
| Human | HNRNPU | Transcription Elongation | GAAGGAATCTCCGCTCTGGA |
| Human | HNRNPU | Transcription Elongation | CATCTGGTCGCCGTCCAGAG |
| Human | HNRNPU | Transcription Elongation | CGACCAGATGGAGCTAGGAG |
| Human | HNRNPU | Transcription Elongation | CACCGTCACCGCGAACAGCG |
| Human | HNRNPU | Transcription Elongation | TCACCGCGAACAGCGAGGTG |
| Human | HNRNPU | Transcription Elongation | CGTCACCGCGAACAGCGAGG |
| Human | HNRNPU | Transcription Elongation | CTCGCTGTTCGCGGTGACGG |
| Human | HNRNPU | Transcription Elongation | AACTACAACCAGAACTTCAG |
| Human | HNRNPU | Transcription Elongation | TGGCAGCAGGGTCAATTCTG |
| Human | HNRNPU | Transcription Elongation | TGGAGTCAGCATTATCACCA |
| Human | CHD3 | Chromatin Readers | TTATGTATCGAAACTACCAG |
| Human | CHD3 | Chromatin Readers | GATCCTACATTGGCGGTGGG |
| Human | CHD3 | Chromatin Readers | AGTGGGTAGGACTATCCTAC |
| Human | CHD3 | Chromatin Readers | CCGCATCATCAACCACAGTG |
| Human | CHD3 | Chromatin Readers | ATATGACCAGTCCACGTGGG |
| Human | CHD3 | Chromatin Readers | CCACACTGTGGTTGATGATG |
| Human | CHD3 | Chromatin Readers | CTCTACTCACTCTACAAGGA |
| Human | CHD3 | Chromatin Readers | CTACCACAAGACAGGCCCAG |
| Human | CHD3 | Chromatin Readers | TTCTATGTGGTGACATACAC |
| Human | CHD3 | Chromatin Readers | GGTGACATACACGGGTGACA |
| Human | CHD3 | Chromatin Readers | CGATCGGTTTAATGCTCCTG |
| Human | CHD3 | Chromatin Readers | AGGAGCATTAAACCGATCGA |
| Human | CHD3 | Chromatin Readers | GGAAGAGCGAATCACACAAG |
| Human | CHD3 | Chromatin Readers | TGACATCCAGGCCTTTAGCC |
| Human | CHD3 | Chromatin Readers | AAGTATGAGCGCATCGATGG |
| Human | CHD3 | Chromatin Readers | GCTCAGGATCAAGGCAGACG |
| Human | CHD3 | Chromatin Readers | ATCAAGGCAGACGAGGTGGT |
| Human | CHD3 | Chromatin Readers | AGACGAGGTGGTAGGCACGA |
| Human | CHD3 | Chromatin Readers | TGAGCTTGACCGGGCTCCAG |
| Human | CHD3 | Chromatin Readers | GAGCTTGACCGGGCTCCAGA |
| Human | CHD3 | Chromatin Readers | GAGGGTTTAGACAATGAATG |
| Human | CHD3 | Chromatin Readers | TGCCGCGTATGCAAGGACGG |

| | | | |
|---|---|---|---|
| Human | CHD3 | Chromatin Readers | TAGACAATGAATGTGGTAGG |
| Human | CHD3 | Chromatin Readers | GTTTAGACAATGAATGTGGT |
| Human | CHD3 | Chromatin Readers | CCGCGTATGCAAGGACGGCG |
| Human | CHD3 | Chromatin Readers | TCCTCCATGGCTATGCACGG |
| Human | CHD3 | Chromatin Readers | GCCACCGTGCATAGCCATGG |
| Human | CHD3 | Chromatin Readers | CAAGGTTCTGAACCAGCTGG |
| Human | CHD3 | Chromatin Readers | GTTCAGAACCTTGTGCAGGA |
| Human | CHD3 | Chromatin Readers | GAAGAAGAATACGAAGAGGA |
| Human | CHD3 | Chromatin Readers | AGAAGAAGAATACGAAGAGG |
| Human | INTS12 | Chromatin Readers | TGTTTGTAGGCAAATGATGG |
| Human | INTS12 | Chromatin Readers | CATTTGCCTACAAACAACGC |
| Human | INTS12 | Chromatin Readers | GCACAATACCACACCAGGCG |
| Human | INTS12 | Chromatin Readers | CTGGCAACATCATCCAAAGG |
| Human | INTS12 | Chromatin Readers | TAAGCCACTAGTTACTGACG |
| Human | INTS12 | Chromatin Readers | TCAGCTAACCAGAAACCTGT |
| Human | PHF2 | Histone Methylation | TGGCGTAGATCCAGCCTGAG |
| Human | PHF2 | Histone Methylation | CTGGTACCACGTGCTCAAGG |
| Human | PHF2 | Histone Methylation | GTGGCGTAGATCCAGCCTGA |
| Human | PHF2 | Histone Methylation | CATTGTAAAGAAACTGTCAT |
| Human | PHF2 | Histone Methylation | TGAAGCGGGTAACGTCGTAG |
| Human | PHF2 | Histone Methylation | GTTCCACGGCAGCTGTGTTG |
| Human | PHF2 | Histone Methylation | GCAGTGGTATATGTCGATGT |
| Human | PHF2 | Histone Methylation | TTCTCCAGGAGGGATTGAGG |
| Human | PHF2 | Histone Methylation | GTGGCCTTGCCCTTCTTGGG |
| Human | PHF2 | Histone Methylation | GCTCGAAGCCCACACCAAGG |
| Human | PHF2 | Histone Methylation | GGTTCACATGCAGAATGATG |
| Human | PHF2 | Histone Methylation | ACGCTGAAGCTCAAAGATGG |
| Human | PHF2 | Histone Methylation | CACGCCAGCCTCTACCACCC |
| Human | PHF2 | Histone Methylation | AGCAGCCAGGCCTCGCAGGA |
| Human | PHF2 | Histone Methylation | TGGCGAGCTGCCCTCCTGCG |
| Human | PHF2 | Histone Methylation | ACCTCCACCTCCATCTCTGC |
| Human | PHF2 | Histone Methylation | GGTGGTGCCGGCAGAGATGG |
| Human | PHF2 | Histone Methylation | GCAGATGAGAGCATATGAAG |
| Human | PHF2 | Histone Methylation | TGAGAGCATATGAAGTGGAA |
| Human | PHF2 | Histone Methylation | AAACTGCGTGCTGGTACATG |
| Human | PHF2 | Histone Methylation | CACCTGCTGGAGGCATTCAA |
| Human | PHF2 | Histone Methylation | AGCCTTTGAATGCCTCCAGC |
| Human | PHF2 | Histone Methylation | AAGACCTGGCCAAAGAGATC |
| Human | MAPK14 | Other Histone Modifications | AATGTATATACTTTAGACCT |
| Human | MAPK14 | Other Histone Modifications | TCCAGCAAGTCGACAGCCAG |
| Human | MAPK14 | Other Histone Modifications | CTCCAGCAAGTCGACAGCCA |
| Human | MIS18BP1 | Chromatin Remodeling | GGATTGAGCACAACAAACTT |

| | | | |
|---|---|---|---|
| Human | MIS18BP1 | Chromatin Remodeling | GATATCCAAATTATCTCATA |
| Human | MIS18BP1 | Chromatin Remodeling | AACAAACTTAGGACTATATC |
| Human | MIS18BP1 | Chromatin Remodeling | ACAGTAATGTAATTATAGAG |
| Human | MIS18BP1 | Chromatin Remodeling | AATACTGCTATATGTGTAGA |
| Human | MIS18BP1 | Chromatin Remodeling | GATCTCCTGAAGAATGCCAG |
| Human | MIS18BP1 | Chromatin Remodeling | CAAACCTGGTTTCTGGTCAG |
| Human | MIS18BP1 | Chromatin Remodeling | TCTCTTCCAAAGCACAAACC |
| Human | CENPH | Chromatin Remodeling | CAGACAGTCTAGTGTGCTCA |
| Human | CENPH | Chromatin Remodeling | GAAATCACAGCAGGAATCTT |
| Human | CENPH | Chromatin Remodeling | TGAAATCACAGCAGGAATCT |
| Human | TAF12 | Transcription Elongation | AGCTGCATTTAGAGCGCCAG |
| Human | TAF12 | Transcription Elongation | ACTGGCGCTCTAAATGCAGC |
| Human | TAF12 | Transcription Elongation | ACAGGACTTAGTAAGAGAAG |
| Human | KDM4C | Histone Methylation | CCTTTGCAAGACCCGCACGA |
| Human | KDM4C | Histone Methylation | CTTTAGACTCCATGTATGCA |
| Human | KDM4C | Histone Methylation | AGGTGATTCCTCCTAAGGAG |
| Human | KDM4C | Histone Methylation | GTCTAAAGGAGCCCATCGTG |
| Human | KDM4C | Histone Methylation | GACCTTCAGACCCTCCATGG |
| Human | KDM4C | Histone Methylation | CTTTGACAAGATAACCCAGG |
| Human | KDM4C | Histone Methylation | TCCCTTTGACAAGATAACCC |
| Human | KDM4C | Histone Methylation | GGTATAGCATACCAAGACTT |
| Human | KDM4C | Histone Methylation | GTCTTGGTATGCTATACCTC |
| Human | KDM4C | Histone Methylation | CTCCTGGGTTATCTTGTCAA |
| Human | KDM4C | Histone Methylation | CGTGATGACCGTTTGACCCA |
| Human | KDM4C | Histone Methylation | GTATTACAGTTGCAGAGTGA |
| Human | KDM4C | Histone Methylation | TCTCGGCTCACGATATCCTC |
| Human | KDM4C | Histone Methylation | GACATCGCAGACCTTCTATG |
| Human | KDM4C | Histone Methylation | AAACTCAACCTGGTACATGT |
| Human | KDM4C | Histone Methylation | CAAACTCAACCTGGTACATG |
| Human | KDM4C | Histone Methylation | GTCCAAGTCAAGTGGCCCGA |
| Human | KDM4C | Histone Methylation | ATCCATCTTCAAACTCAACC |
| Human | KDM4C | Histone Methylation | AAATATTGCCCACATGTACC |
| Human | KDM4C | Histone Methylation | CCGTACGCAGCACTTTGCAC |
| Human | KDM4C | Histone Methylation | CCATCACAGATCTCATGAGA |
| Human | KDM4C | Histone Methylation | CTGTGCAAAGTGCTGCGTAC |
| Human | KDM4C | Histone Methylation | CCTGTGCAAAGTGCTGCGTA |
| Human | KDM4C | Histone Methylation | CCTTCTCATGAGATCTGTGA |
| Human | KDM4C | Histone Methylation | GCAAACGAAGAACAATAAGT |
| Human | KDM4C | Histone Methylation | CGCAACGGCGCACATGACAT |
| Human | KDM4C | Histone Methylation | TGTCTCTGCAATTTGAGAGG |
| Human | KDM4C | Histone Methylation | TGCTGTCTCTGCAATTTGAG |
| Human | KDM4C | Histone Methylation | CCATGTCATGTGCGCCGTTG |

| | | | |
|---|---|---|---|
| Human | KDM4C | Histone Methylation | GGCACAAGTGACATGGAAGG |
| Human | KDM4C | Histone Methylation | GGAGGCCGGGCAGCGACCGT |
| Human | KDM4C | Histone Methylation | GGCAGCGACCGTAGGAACAC |
| Human | KDM4C | Histone Methylation | CAGCATGGGCACAAGTGACA |
| Human | KDM4C | Histone Methylation | ATGGGCACAAGTGACATGGA |
| Human | EIF2B1 | Control | GGCTGCCATCCGGACGTTGC |
| Human | EHMT1 | Histone Methylation | CAATTCCGTCCACCAGCATG |
| Human | EHMT1 | Histone Methylation | GTGCTCCTCATGCTGGTGGA |
| Human | EHMT1 | Histone Methylation | GAAGGTGCTCCTCATGCTGG |
| Human | EHMT1 | Histone Methylation | TGTACTTCTCCGCCAGGCAA |
| Human | EHMT1 | Histone Methylation | CGCCCGCCTGAACCAGCATG |
| Human | EHMT1 | Histone Methylation | GCGCTCTCCACTGCACGCCG |
| Human | EHMT1 | Histone Methylation | TGCCACATGCTGGTTCAGGC |
| Human | EHMT1 | Histone Methylation | CTGCCACATGCTGGTTCAGG |
| Human | EHMT1 | Histone Methylation | CACGTGTCCAGCCTCTGCCG |
| Human | EHMT1 | Histone Methylation | TTCGGCTGCTTCCATCAACG |
| Human | EHMT1 | Histone Methylation | TGAAGTACCTCATCAAGGCT |
| Human | EHMT1 | Histone Methylation | GAAGTACCTCATCAAGGCTG |
| Human | EHMT1 | Histone Methylation | GGTACTTCACTGCTTCCAGA |
| Human | EHMT1 | Histone Methylation | CAAGAAAGGCCACTACGAAG |
| Human | EHMT1 | Histone Methylation | CCATCTGTCCATTTGAAAGC |
| Human | EHMT1 | Histone Methylation | CCTGCTTTCAAATGGACAGA |
| Human | EHMT1 | Histone Methylation | GTCCATTTGAAAGCAGGTAC |
| Human | EHMT1 | Histone Methylation | GGCCACAGAGTACAAGCACG |
| Human | EHMT1 | Histone Methylation | TGGACAGCAGCAGCTTCACG |
| Human | EHMT1 | Histone Methylation | GTCCACGTGCTTGTACTCTG |
| Human | EHMT1 | Histone Methylation | TACTCTGTGGCCCAGATCAT |
| Human | EHMT1 | Histone Methylation | GTGAAGCTGCTGCTGTCCAA |
| Human | EHMT1 | Histone Methylation | GAACATTTGCCTGCACTGGG |
| Human | EHMT1 | Histone Methylation | CTCGGCTATGTCCACGCAGC |
| Human | EHMT1 | Histone Methylation | GGCGGCGTTCTCCGGCTGCG |
| Human | EHMT1 | Histone Methylation | CGGAGAACGCCGCCCAGTGC |
| Human | EHMT1 | Histone Methylation | GGACATAGCCGAGATCCTGC |
| Human | EHMT1 | Histone Methylation | AGAGGACGACACAGTCGTAG |
| Human | EHMT1 | Histone Methylation | CTGAATCCCGAGAAAGAAAG |
| Human | EHMT1 | Histone Methylation | ACAGTCGTAGCGGTTCTCCC |
| Human | EHMT1 | Histone Methylation | CGCCACTGCACATTGCCGCC |
| Human | EHMT1 | Histone Methylation | TGTCGTCCTCTTTCTTTCTC |
| Human | EHMT1 | Histone Methylation | CGAGCCTCAACTCTCAGGTG |
| Human | EHMT1 | Histone Methylation | GTTGAGGCTCGCACACTGCA |
| Human | EHMT1 | Histone Methylation | GAGCGCTCTGCAGATGAGCA |
| Human | EHMT1 | Histone Methylation | TTGAGGCTCGCACACTGCAG |

| | | | |
|---|---|---|---|
| Human | EHMT1 | Histone Methylation | GCTGGTACGACAAGGATGGC |
| Human | EHMT1 | Histone Methylation | CCTGCCAGAGTTCAACATGG |
| Human | EHMT1 | Histone Methylation | GTGGTTGCATTCGAAGATCA |
| Human | EHMT1 | Histone Methylation | TGGTTGCATTCGAAGATCAA |
| Human | EHMT1 | Histone Methylation | CCCAACATACTCGCAGACAA |
| Human | EHMT1 | Histone Methylation | TGATCTCGACAATAAGGACG |
| Human | EHMT1 | Histone Methylation | TTTGATCTCGACAATAAGGA |
| Human | ARID2 | Chromatin Remodeling | GGCAGTTCCACCACAGCAGA |
| Human | ARID2 | Chromatin Remodeling | TCGCTTTCCTGGACGAGCTG |
| Human | ARID2 | Chromatin Remodeling | CGGCAGTTCCACCACAGCAG |
| Human | ARID2 | Chromatin Remodeling | AAGGCGACCCTCTGCTGTGG |
| Human | ARID2 | Chromatin Remodeling | ACTTGCAGTAAATTAGCTCG |
| Human | ARID2 | Chromatin Remodeling | CAGGCACATATTCATGTGGT |
| Human | ARID2 | Chromatin Remodeling | TGGGCAGGCACATATTCATG |
| Human | ARID2 | Chromatin Remodeling | CTAATTTACTGCAAGTCGAG |
| Human | ARID2 | Chromatin Remodeling | GTGGTTTCAGACACCCTCAC |
| Human | ARID2 | Chromatin Remodeling | GTTCTGAAACCCTACCCTTG |
| Human | ARID2 | Chromatin Remodeling | GATAACATTTGACTGACAGT |
| Human | PHF20L1 | Chromatin Readers | GGAGTCATCGTTATGATGAG |
| Human | PHF20L1 | Chromatin Readers | TGAGATTGGTGCTCGTTTGG |
| Human | PHF20L1 | Chromatin Readers | TTATGATGAGTGGATTTACT |
| Human | PHF20L1 | Chromatin Readers | TAATTGCTTCAATCTTGGCA |
| Human | PHF20L1 | Chromatin Readers | GCACATTAAAGCCATGCCCG |
| Human | PHF20L1 | Chromatin Readers | TTTGTTAATTGCTTCAATCT |
| Human | PHF20L1 | Chromatin Readers | GAGAAGAAGTTCTGGCTCGT |
| Human | PHF20L1 | Chromatin Readers | ATTGAAGCAATTAACAAAGA |
| Human | PHF20L1 | Chromatin Readers | AGTGTGAAGAGTGCTTGTGT |
| Human | PHF20L1 | Chromatin Readers | GGCAACACAGCGTGTGCATG |
| Human | PHF20L1 | Chromatin Readers | TGGCAACACAGCGTGTGCAT |
| Human | PHF20L1 | Chromatin Readers | AACACAAGCACTCTTCACAC |
| Human | PHF20L1 | Chromatin Readers | ATGTATTTGTGAGATGGATG |
| Human | JMJD1C | Histone Methylation | TTTAGCAGCAACTACACCAT |
| Human | JMJD1C | Histone Methylation | AAGGACTCAAGTGAAATACC |
| Human | JMJD1C | Histone Methylation | CAACATTGTTTAACATTCAC |
| Human | JMJD1C | Histone Methylation | AACATTGTTTAACATTCACT |
| Human | SMYD1 | Histone Methylation | ATTTAACAATGGCAATCATG |
| Human | SMYD1 | Histone Methylation | CAATGAGAACATCAGGCTGG |
| Human | SMYD1 | Histone Methylation | GTGGAGGTCTTCACCGCTGA |
| Human | SIRT3 | Histone Acetylation | TTCCAGACTTCAGATCGCCG |
| Human | SIRT3 | Histone Acetylation | CGATGGGCTTGAGAGAGTGT |
| Human | SIRT3 | Histone Acetylation | AGATGCGGGACCTTGTGCAG |
| Human | CENPQ | Chromatin Remodeling | AATTCAGATTCTGGCAAGTG |

| | | | |
|---|---|---|---|
| Human | CENPQ | Chromatin Remodeling | TCAGATTCTGGCAAGTGAGG |
| Human | CENPQ | Chromatin Remodeling | AAGTGAGGTGGAAGAAGAAG |
| Human | PBRM1 | Chromatin Readers | TCAACCAGACTATTATGAAG |
| Human | PBRM1 | Chromatin Readers | TCTCTGTGAGCTCTTCATTA |
| Human | PBRM1 | Chromatin Readers | TCTGGTTGATTTCTTCGCTT |
| Human | PBRM1 | Chromatin Readers | ATTGCCCAGAGGATACAGAA |
| Human | PBRM1 | Chromatin Readers | GCTTCCATTCTGTATCCTCT |
| Human | PBRM1 | Chromatin Readers | TCTGGATATTGCACTTTAGA |
| Human | PBRM1 | Chromatin Readers | GCAATGGTCTTGAGATCTAT |
| Human | PBRM1 | Chromatin Readers | AGCTTCCATTCTGTATCCTC |
| Human | PBRM1 | Chromatin Readers | CGGATCTGTTGTAGTGATAT |
| Human | PBRM1 | Chromatin Readers | TCGGATCTGTTGTAGTGATA |
| Human | PBRM1 | Chromatin Readers | ATTGGGCACATTATAGCGTT |
| Human | PBRM1 | Chromatin Readers | ATCCGCAATGACAAATATGC |
| Human | PBRM1 | Chromatin Readers | TGCCAGGCACTATAATGAGG |
| Human | PBRM1 | Chromatin Readers | AGCTGATGTTCCGGAATGCC |
| Human | PBRM1 | Chromatin Readers | GCCAGGCACTATAATGAGGA |
| Human | PBRM1 | Chromatin Readers | CTCCGGCTCATTGTATGTAC |
| Human | PBRM1 | Chromatin Readers | AGATATTGACTCTATGGTTG |
| Human | PBRM1 | Chromatin Readers | TCAGGCAACTCAGATCTAGA |
| Human | PBRM1 | Chromatin Readers | TCAGAAATATGGCACTGAGG |
| Human | PBRM1 | Chromatin Readers | CTTGAAATAAATCAAGCCGA |
| Human | PBRM1 | Chromatin Readers | GGTGGTTTGTTAGGAAAGTT |
| Human | PBRM1 | Chromatin Readers | ACCAAATGAAACATTCCACC |
| Human | PBRM1 | Chromatin Readers | TGTTGGAGATTACGTCTATG |
| Human | PBRM1 | Chromatin Readers | CTTTCAATACAGACGATATG |
| Human | PBRM1 | Chromatin Readers | GTGTGTGGTCATGTTTGTCA |
| Human | PBRM1 | Chromatin Readers | TTCCGAGAATACATGTCATG |
| Human | PBRM1 | Chromatin Readers | CTTTCCGAGAATACATGTCA |
| Human | PBRM1 | Chromatin Readers | TTTCCGAGAATACATGTCAT |
| Human | PBRM1 | Chromatin Readers | AGAAAGCAGAATATGAAGAG |
| Human | PBRM1 | Chromatin Readers | GAAAGCAGAATATGAAGAGC |
| Human | PBRM1 | Chromatin Readers | TCCTGTTCAGCAGTGAGATG |
| Human | PBRM1 | Chromatin Readers | CCTTGGTTCATCACACCCGG |
| Human | PBRM1 | Chromatin Readers | ACTCCTTGGTTCATCACACC |
| Human | PBRM1 | Chromatin Readers | CCACCGGGTGTGATGAACCA |
| Human | PBRM1 | Chromatin Readers | AGTCCATATGGACAACAGGT |
| Human | PBRM1 | Chromatin Readers | CCTGTTGCAGGCATGATGGG |
| Human | PBRM1 | Chromatin Readers | ATTATAGAGTTCATGGCACA |
| Human | PBRM1 | Chromatin Readers | CAAATCCCAGAGTTTGCAAG |
| Human | PBRM1 | Chromatin Readers | ATAAAGCCGCTTGCAAACTC |
| Human | PBRM1 | Chromatin Readers | TAAAGCCGCTTGCAAACTCT |

| | | | |
|---|---|---|---|
| Human | PBRM1 | Chromatin Readers | TGGCTTCAAGAAGCTGCTCC |
| Human | PBRM1 | Chromatin Readers | TATCCTGGAGAAGTTACTCA |
| Human | PBRM1 | Chromatin Readers | TTATAATGATGCACATATCC |
| Human | PBRM1 | Chromatin Readers | AGAAGTTACTCAAGGAGAAA |
| Human | PBRM1 | Chromatin Readers | TCTCCTTGAGTAACTTCTCC |
| Human | PARP4 | Other Histone Modifications | AAGATTATATGTGTGCACTG |
| Human | PARP4 | Other Histone Modifications | CTGGGTTTGCAATATGAACG |
| Human | PARP4 | Other Histone Modifications | GCACTGAGGATTTAACGAAA |
| Human | PARP4 | Other Histone Modifications | ACGATGTGAGCAAGGCAGAG |
| Human | PARP4 | Other Histone Modifications | CTGCCAGCTAATAAGAGACA |
| Human | PARP4 | Other Histone Modifications | TAGCCTCAACGATGTGAGCA |
| Human | PARP4 | Other Histone Modifications | CTGCCTTGCTCACATCGTTG |
| Human | PARP4 | Other Histone Modifications | TCATCACAGTAAGAGCCCAG |
| Human | PARP4 | Other Histone Modifications | AGTATCAAGTACTCACACCC |
| Human | PARP4 | Other Histone Modifications | ATAAGAACCTTAGCCTTAGG |
| Human | PARP4 | Other Histone Modifications | GCTTATTGTACCCTGCTCGA |
| Human | PARP4 | Other Histone Modifications | TGCTGTAGAACCGATACCGC |
| Human | PARP4 | Other Histone Modifications | CACACCAGGTTATTCGCCTG |
| Human | PARP4 | Other Histone Modifications | GATACCGCAGGCGAATAACC |
| Human | PARP4 | Other Histone Modifications | ACTCCTCAGTCTACAGACAG |
| Human | PARP4 | Other Histone Modifications | AGACTGAGGAGTTCTGTCCA |
| Human | PARP4 | Other Histone Modifications | CATCCTCTGTCTGTAGACTG |
| Human | PARP4 | Other Histone Modifications | CTCAGTCTACAGACAGAGGA |
| Human | TAF4B | Transcription Elongation | GTTAGGCAGAAGTTGTCGTA |
| Human | TAF4B | Transcription Elongation | CTTGATCTCCCAAGCAACAC |
| Human | TAF4B | Transcription Elongation | GTAGTCGTTCCTGTGTTGCT |
| Human | TAF4B | Transcription Elongation | TAGTCGTTCCTGTGTTGCTT |
| Human | TAF4B | Transcription Elongation | GAATCCCAGCTGGTTGTGAA |
| Human | TAF4B | Transcription Elongation | TGCTTGACTTGAACTGCCTG |
| Human | TAF4B | Transcription Elongation | ACAACCAGCTGGGATTCCAC |
| Human | TAF4B | Transcription Elongation | CCAGCTGGTTGTGAAAGGAC |
| Human | TAF4B | Transcription Elongation | CCTGTCCTTTCACAACCAGC |
| Human | TAF4B | Transcription Elongation | GTATTCTCGAGCTCTATACC |
| Human | TAF4B | Transcription Elongation | AATCACGAGAATCTGCCTCA |
| Human | TAF4B | Transcription Elongation | TTCTTTATTAGAACGACTCT |
| Human | TAF4B | Transcription Elongation | AGAAAGAGAAATGTTACTTA |
| Human | TAF4B | Transcription Elongation | AGCCAAAGAGTTACAGCAAT |
| Human | TAF4B | Transcription Elongation | TTCCAATTGCTGTAACTCTT |
| Human | TAF4B | Transcription Elongation | TCAATTGGAGAAGCAGAGAA |
| Human | TAF4B | Transcription Elongation | TGTTGAGGAATGTGAAATTG |
| Human | TAF4B | Transcription Elongation | GTAGTTCAGCAGCCTTCAGG |
| Human | TAF4B | Transcription Elongation | CTAGTAGTTCAGCAGCCTTC |

| | | | |
|---|---|---|---|
| Human | CDY1 | Chromatin Readers | ATACACAGTATTTGGTTCGG |
| Human | CDY1 | Chromatin Readers | TGACAAACAGGATGACACTT |
| Human | CDY1 | Chromatin Readers | ATGACAAACAGGATGACACT |
| Human | CDY1 | Chromatin Readers | GGAATACACAGTATTTGGTT |
| Human | CDY1 | Chromatin Readers | CACAGCAAGCCTTGAAATGG |
| Human | CDY1 | Chromatin Readers | AGTATTCACAAAGTTCTTGA |
| Human | CDY1 | Chromatin Readers | TGATGGTGTCCACCATTTCA |
| Human | CDY1 | Chromatin Readers | GGCCAATGAGAGAGAGTGTG |
| Human | CDY1 | Chromatin Readers | CACCTCACACTCTCTCTCAT |
| Human | CDY1 | Chromatin Readers | AGAGAGAGTGTGAGGTGCTG |
| Human | CDK12 | Transcription Elongation | GAGCTGTGGATGTATTCTTG |
| Human | CDK12 | Transcription Elongation | CAAAGACACAGGAGAACTAG |
| Human | CDK12 | Transcription Elongation | TGGCGGCGGTGGAGGTCCGG |
| Human | CDK12 | Transcription Elongation | AGGTGGCGGCGGTGGAGGTC |
| Human | CDK12 | Transcription Elongation | GGCGGCGGTGGAGGTCCGGG |
| Human | CDK12 | Transcription Elongation | GTGGCGGCGGTGGAGGTCCG |
| Human | CDK12 | Transcription Elongation | GGTGGCGGCGGTGGAGGTCC |
| Human | SUPT4H1 | Transcription Elongation | CACAACCATCATATTCAAAC |
| Human | SUPT4H1 | Transcription Elongation | ATAGACCAGTTTGAATATGA |
| Human | SUPT4H1 | Transcription Elongation | GACCAGCGAACACAGCAAAC |
| Human | SUPT4H1 | Transcription Elongation | AACAGGCCCGCAGATGCCGC |
| Human | SUPT4H1 | Transcription Elongation | AGACGGTGCCGAAGGACCTG |
| Human | SUPT4H1 | Transcription Elongation | CGAAGGACCTGCGGCATCTG |
| Human | SUPT4H1 | Transcription Elongation | CGCAGATGCCGCAGGTCCTT |
| Human | BRWD3 | Chromatin Readers | GTGTAGCATTTGACCGAAGC |
| Human | BRWD3 | Chromatin Readers | TGTGTAGCATTTGACCGAAG |
| Human | BRWD3 | Chromatin Readers | AGATGCATAAGAGAATTCTG |
| Human | BRWD3 | Chromatin Readers | GTGATAAGGTAGTAAGAGTA |
| Human | BRWD3 | Chromatin Readers | CTCTGCTGAAATTTCTGACA |
| Human | BRWD3 | Chromatin Readers | TGCTGCAGGCAGCTGTGATA |
| Human | BRWD3 | Chromatin Readers | CATCAGCACCAGTAGAAGTG |
| Human | BRWD3 | Chromatin Readers | TAGAAGCTGAATGGCCCTGA |
| Human | BRWD3 | Chromatin Readers | AACAGATACCTCACTTCTAC |
| Human | BRWD3 | Chromatin Readers | TTTACTGAGAGATCCAGACC |
| Human | BRWD3 | Chromatin Readers | TGTTCATCTTTCAGTTCTGG |
| Human | BRWD3 | Chromatin Readers | GAACAAGATATCTGGACTCC |
| Human | BRWD3 | Chromatin Readers | GATCTCTCAGTAAATTTCAC |
| Human | BRWD3 | Chromatin Readers | GGTGGTATGTTCATTACAAC |
| Human | BRWD3 | Chromatin Readers | TAGTGGAAGTCGAGATGGAA |
| Human | BRWD3 | Chromatin Readers | GCTGTTCAGTTCTGTAACAA |
| Human | BRWD3 | Chromatin Readers | TGCTGAATTAGAGTCACATA |
| Human | BRWD3 | Chromatin Readers | GAGATGGAACGGCAAGAATT |

| | | | |
|---|---|---|---|
| Human | BRWD3 | Chromatin Readers | TTTGTTAGTGGAAGTCGAGA |
| Human | BRWD3 | Chromatin Readers | TGTGGTATCATAGCGATCCC |
| Human | BRWD3 | Chromatin Readers | TAAACTTAAGGTGACTATGG |
| Human | BRWD3 | Chromatin Readers | ACTGGCAATAATTTGCCATC |
| Human | BRWD3 | Chromatin Readers | GTTCACTGCAGTAATAACTG |
| Human | BRWD3 | Chromatin Readers | CAAAGGATCATACTTTCAGC |
| Human | BRWD3 | Chromatin Readers | TACTTTCAGCAGGTCATGAT |
| Human | BRWD3 | Chromatin Readers | ATACTTTCAGCAGGTCATGA |
| Human | BRWD3 | Chromatin Readers | AAGCCCATCCATTTGATCAA |
| Human | BRWD3 | Chromatin Readers | TGATCCTTTGATCAAATGGA |
| Human | BRWD3 | Chromatin Readers | TTTAACATGATTGAAGGCCA |
| Human | BRWD3 | Chromatin Readers | ATGTCCATGAGAATCTGTGC |
| Human | BRWD3 | Chromatin Readers | TGAAGGCCAAGGCCATGGTG |
| Human | BRWD3 | Chromatin Readers | CTGACCTCAATACCATCAGG |
| Human | BRWD3 | Chromatin Readers | GCTATATTGAACATAATGCC |
| Human | BRWD3 | Chromatin Readers | CCTCTTGCTCCTGATGACCT |
| Human | BRWD3 | Chromatin Readers | CCCAGGTCATCAGGAGCAAG |
| Human | BRWD3 | Chromatin Readers | AGGAAACTATGGTAGTCCTC |
| Human | BRWD3 | Chromatin Readers | AGATGTTATAGATACTCCTG |
| Human | BRWD3 | Chromatin Readers | GCTGATCTTCTTTCTTACCC |
| Human | BRWD3 | Chromatin Readers | TGGAAGACCAGAGGCACAGG |
| Human | PARP3 | Other Histone Modifications | TCTTCACATCCAGGTCCATG |
| Human | PARP3 | Other Histone Modifications | CTTCACATCCAGGTCCATGA |
| Human | PARP3 | Other Histone Modifications | GGACATGCTGCTGGTGCTGG |
| Human | PARP3 | Other Histone Modifications | GGGTCGGATCAGGCTCGGTG |
| Human | PARP3 | Other Histone Modifications | CACCGAGCCTGATCCGACCC |
| Human | PARP3 | Other Histone Modifications | ATCTGGAAAGTAAACCAAGA |
| Human | PARP3 | Other Histone Modifications | TGATGCGGAGCCCACTAGTG |
| Human | PARP3 | Other Histone Modifications | AGGACTTTGAGAAGAAATTT |
| Human | PARP3 | Other Histone Modifications | GGACTTTGAGAAGAAATTTC |
| Human | POLR2G | Transcription Elongation | TTTCGTGCTCTAGGGAGATA |
| Human | POLR2G | Transcription Elongation | CCTTTATCCAGTTAAGTACA |
| Human | POLR2G | Transcription Elongation | CCTTGTACTTAACTGGATAA |
| Human | PRDM2 | Histone Methylation | ACTTTAAAGCCAATCGCGCC |
| Human | PRDM2 | Histone Methylation | TTAAGAATAATGTATACATG |
| Human | PRDM2 | Histone Methylation | AGTGGCCCAGACACCAATCC |
| Human | PRDM2 | Histone Methylation | TGGCTCTGTTGATTTCCAGT |
| Human | PRDM2 | Histone Methylation | CAAGACCCGGATTGGTGTCT |
| Human | PRDM2 | Histone Methylation | GGAAGTTTACAACCAAACAG |
| Human | PRDM2 | Histone Methylation | AGGAAGTTTACAACCAAACA |
| Human | PRDM2 | Histone Methylation | TTCCTTTCACAATGTTGACA |
| Human | PRDM2 | Histone Methylation | AAGGAAGTTTACAACCAAAC |

| | | | |
|---|---|---|---|
| Human | PRDM2 | Histone Methylation | TTGGCACACAGATTAACCGG |
| Human | PRDM2 | Histone Methylation | CCGGTTAATCTGTGTGCCAA |
| Human | PRDM2 | Histone Methylation | TTAACCGGCGGCGACATGAG |
| Human | PRDM2 | Histone Methylation | CCTTTGGCACACAGATTAAC |
| Human | PRDM2 | Histone Methylation | AAGTACTGTGGGAAAGCCTT |
| Human | PRDM2 | Histone Methylation | GAACTCATACTAATATGAGA |
| Human | PRDM2 | Histone Methylation | AGATCTTTAATGGAAAGAAA |
| Human | PRDM2 | Histone Methylation | ATGTTTGGTTAGATCTTTAA |
| Human | PRDM2 | Histone Methylation | AGGATATGTTTGTGCAACTC |
| Human | PRDM2 | Histone Methylation | AATCGAGAGTTCACTTACAT |
| Human | PRDM2 | Histone Methylation | GTGAACTCTCGATTACAGTA |
| Human | PRDM2 | Histone Methylation | TGAACTCTCGATTACAGTAA |
| Human | PRDM2 | Histone Methylation | CTGTGAAAGAGCCAGAAATA |
| Human | PRDM2 | Histone Methylation | AAGAGGTACGGTAGGAAGGA |
| Human | PRDM2 | Histone Methylation | GAAGAGGTACGGTAGGAAGG |
| Human | PRDM2 | Histone Methylation | AGAGGTACGGTAGGAAGGAG |
| Human | PRDM2 | Histone Methylation | GGTGAGGCGGCAGACATCAG |
| Human | PRDM2 | Histone Methylation | TCCCTCTCCCATTCCTCCCG |
| Human | PRDM2 | Histone Methylation | TGATGTCTGCCGCCTCACCC |
| Human | PRDM2 | Histone Methylation | GGGTGAGGCGGCAGACATCA |
| Human | PRDM2 | Histone Methylation | ATGATGAGTTGGAAGACGAG |
| Human | PRDM2 | Histone Methylation | GATGATGAGTTGGAAGACGA |
| Human | PRDM2 | Histone Methylation | TGATGATGAGTTGGAAGACG |
| Human | PRDM2 | Histone Methylation | TGATGAGTTGGAAGACGAGG |
| Human | PRDM2 | Histone Methylation | AGATGATGATGATGATGAGT |
| Human | PRDM2 | Histone Methylation | GGAGGAGGAAGATGAAGAGG |
| Human | PRDM2 | Histone Methylation | AGAGGAGGAGGAAGATGAAG |
| Human | PRDM2 | Histone Methylation | GGGCTGTCCCATTCGTCGGA |
| Human | CCNK | Transcription Elongation | GAGGCCCGGTACCGCCGAGA |
| Human | CCNK | Transcription Elongation | GGTACCGCCGAGAGGGCGCT |
| Human | CCNK | Transcription Elongation | AGATGAACCGAGCGCCCTCT |
| Human | CCNK | Transcription Elongation | CGAGGCCCGGTACCGCCGAG |
| Human | CCNK | Transcription Elongation | ATAAATTATTCCAGTTGCCA |
| Human | CCNK | Transcription Elongation | AATAAATTATTCCAGTTGCC |
| Human | CCNK | Transcription Elongation | CCTGTTGCCTCTTTCTGGCT |
| Human | CCNK | Transcription Elongation | GCCTGTTGCCTCTTTCTGGC |
| Human | CCNK | Transcription Elongation | AGGAGCCTGTTGCCTCTTTC |
| Human | CCNK | Transcription Elongation | GTAGCAGTGATGTATCTCGC |
| Human | BAZ1A | Chromatin Readers | TTTCATGTAAGCGCGAACGA |
| Human | BAZ1A | Chromatin Readers | GTAAGCGCGAACGATGGGTA |
| Human | BAZ1A | Chromatin Readers | TCACAGCACAACTCCACACA |
| Human | BAZ1A | Chromatin Readers | AGTAACCCTAGAAGTATTAG |

| | | | |
|---|---|---|---|
| Human | BAZ1A | Chromatin Readers | ACTTCTAGGGTTACTCCATC |
| Human | BAZ1A | Chromatin Readers | GGAGTTCCTTAATGCATTTG |
| Human | BAZ1A | Chromatin Readers | TGTCATAGTAGTCTGGGACC |
| Human | BAZ1A | Chromatin Readers | CGAATTATATTTAAGGCAAT |
| Human | BAZ1A | Chromatin Readers | TGATGATGTCATAGTAGTCT |
| Human | BAZ1A | Chromatin Readers | AAGCTCAAGACTGTGCCTGA |
| Human | BAZ1A | Chromatin Readers | GAGCTTTGGTCGAACACAGT |
| Human | BAZ1A | Chromatin Readers | TGCAAGATATGTCGAAAGAA |
| Human | BAZ1A | Chromatin Readers | TCAGGCACAGTCTTGAGCTT |
| Human | BAZ1A | Chromatin Readers | AACGGCTTTCGGTGTAGCAG |
| Human | BAZ1A | Chromatin Readers | TGTGAGACAGAAGCCGCCCG |
| Human | BAZ1A | Chromatin Readers | GTGGAAGTCATTAGGAACAA |
| Human | BAZ1A | Chromatin Readers | GGCGGCTTCTGTCTCACAAA |
| Human | BAZ1A | Chromatin Readers | TACATGCGGTCGCGACCCAA |
| Human | BAZ1A | Chromatin Readers | TCCAGTATCGTCTATACATG |
| Human | BAZ1A | Chromatin Readers | ACAAGAAGAAATGAAGTCAC |
| Human | BAZ1A | Chromatin Readers | CAGCTGCCAGAATTCGTAAA |
| Human | BAZ1A | Chromatin Readers | TATATTACGACAGGCAAAGC |
| Human | HDAC8 | Histone Acetylation | GCACCATACTGGCCCGTTTG |
| Human | HDAC8 | Histone Acetylation | TGATGTAGTTGAGGATTTGT |
| Human | HDAC8 | Histone Acetylation | AGTTGAGGATTTGTTGGATT |
| Human | CENPP | Chromatin Remodeling | GTCCTCACACGCTCGCCGCA |
| Human | CENPP | Chromatin Remodeling | GGCCGCGATCTCAGCTTGCA |
| Human | CENPP | Chromatin Remodeling | TCTCTTCCCAGGGCGCCGGT |
| Human | KDM5A | Histone Methylation | CGTCTTTGAGCCGAGTTGGG |
| Human | KDM5A | Histone Methylation | TCAGCTTTATCGGCCGCATC |
| Human | KDM5A | Histone Methylation | AAATTCGGCCGCCCAAGGAC |
| Human | KDM5A | Histone Methylation | CGGCCGCATCCGGCCTTTGG |
| Human | KDM5A | Histone Methylation | AGTCGCTTGGGATATCTGCC |
| Human | KDM5A | Histone Methylation | AAGATTGTTGCCAGCAAAGG |
| Human | KDM5A | Histone Methylation | TCTAAAGTGGGTAGTCGCTT |
| Human | KDM5A | Histone Methylation | AGCAAGATTGTTGCCAGCAA |
| Human | KDM5A | Histone Methylation | ATCTACTCTGAAGATCCCTG |
| Human | KDM5A | Histone Methylation | CATCAACTACTTGCACTGGG |
| Human | KDM5A | Histone Methylation | TCCATCAACTACTTGCACTG |
| Human | KDM5A | Histone Methylation | CACTGATTGGTCCTGTACAC |
| Human | KDM5A | Histone Methylation | CTGGGATTCAAATAACTCGG |
| Human | KDM5A | Histone Methylation | CCATCAACTACTTGCACTGG |
| Human | KDM5A | Histone Methylation | CCACTACCTGATGTGCCCAA |
| Human | KDM5A | Histone Methylation | TTGGGCACATCAGGTAGTGG |
| Human | KDM5A | Histone Methylation | ATGTGCCCAAAGGAGACTGG |
| Human | KDM5A | Histone Methylation | CAGTCTCCTTTGGGCACATC |

| | | | |
|---|---|---|---|
| Human | KDM5A | Histone Methylation | TAGAGACATACAAGCCGCTC |
| Human | KDM5A | Histone Methylation | CATGTTCCTGTAATCCTGAG |
| Human | KDM5A | Histone Methylation | TAAGACATTTCTTCTGCATG |
| Human | KDM5A | Histone Methylation | TTTGCCGCAAGACAGCCAGT |
| Human | KDM5A | Histone Methylation | ACACTGTAGCATAAACCCAC |
| Human | KDM5A | Histone Methylation | ATTTGCCGCAAGACAGCCAG |
| Human | KDM5A | Histone Methylation | TGGGAACTTGATTTAGGAAG |
| Human | KDM5A | Histone Methylation | TAAACCCACTGGCTGTCTTG |
| Human | KDM5A | Histone Methylation | GTAGACTGGGTACAATGTGA |
| Human | KDM5A | Histone Methylation | CAGTCTACCTTGTCCTTGCA |
| Human | KDM5A | Histone Methylation | GACTGGGTACAATGTGATGG |
| Human | KDM5A | Histone Methylation | CTGCAAGGACAAGGTAGACT |
| Human | KDM5A | Histone Methylation | TGGTTTCATCAAGTTTGTGT |
| Human | KDM5A | Histone Methylation | TAGCTCATTATGAAGGAAGG |
| Human | KDM5A | Histone Methylation | AATGAGCTACAAACTACCAA |
| Human | KDM5A | Histone Methylation | TAAACTGTGCAAAGAAGCAG |
| Human | KDM5A | Histone Methylation | ATAAACTGTGCAAAGAAGCA |
| Human | KDM5A | Histone Methylation | ATTAGGTGCAGACAAATCAA |
| Human | KDM5A | Histone Methylation | ACGGAAACGGAAGCTAGAAA |
| Human | KDM5A | Histone Methylation | CTCTTTCACAAGTTCAACTT |
| Human | KDM5A | Histone Methylation | ATCAAAGGAGCTGAATAAAC |
| Human | KDM5A | Histone Methylation | AGTCAATTCTTTGCAGACCA |
| Human | KDM5A | Histone Methylation | TTACACCATATAGCAGAGAA |
| Human | KDM5A | Histone Methylation | TTTACACCATATAGCAGAGA |
| Human | SMARCC2 | Chromatin Readers | TGATGTCTGTGCCATCATGA |
| Human | SMARCC2 | Chromatin Readers | AACACTATTGTAGTCAAACC |
| Human | SMARCC2 | Chromatin Readers | GAAGGCATGGACCCTCATGA |
| Human | SMARCC2 | Chromatin Readers | CAAGACTCCAGAGATCTACC |
| Human | SMARCC2 | Chromatin Readers | TAGAGGTAAGATACTCTTGG |
| Human | SMARCC2 | Chromatin Readers | CAGTGCCTCCAGGAGAAGCA |
| Human | SMARCC2 | Chromatin Readers | CTGTTCTGTCCACTCACGAG |
| Human | SMARCC2 | Chromatin Readers | AGATGCACTCGTCCTGTGTG |
| Human | SMARCC2 | Chromatin Readers | GCATGTGGGAAGCCGCACAC |
| Human | SMARCC2 | Chromatin Readers | CCAGTGCCTCCAGGAGAAGC |
| Human | SMARCC2 | Chromatin Readers | GAGAAGCACTGGAGTATCAG |
| Human | SMARCC2 | Chromatin Readers | GGACCGGGAGCGAGAAGCAC |
| Human | SMARCC2 | Chromatin Readers | ACTCCAGTGCTTCTCGCTCC |
| Human | SMARCC2 | Chromatin Readers | GCTGGAGACTATCATGGACC |
| Human | SMARCC2 | Chromatin Readers | AGCTGGAGACTATCATGGAC |
| Human | SMARCC2 | Chromatin Readers | TGAGGAGAGCGGGAATGACG |
| Human | SMARCC2 | Chromatin Readers | AAGGAGTCCGAGAAGAGTGA |
| Human | SMARCC2 | Chromatin Readers | GCCAGCCCACTGCCAGACCC |

| | | | |
|---|---|---|---|
| Human | AURKB | Other Histone Modifications | GCGAACAGCCACGATCATGG |
| Human | AURKB | Other Histone Modifications | TTAGGTCCACCTTGACGATG |
| Human | AURKB | Other Histone Modifications | CATTGTCTTCCTCCTCAGGG |
| Human | PSIP1 | Chromatin Readers | GGAACTTCGTCTACTCGAGC |
| Human | PSIP1 | Chromatin Readers | TCTACTCGAGCTGGCCAATG |
| Human | PSIP1 | Chromatin Readers | CGTCTACTCGAGCTGGCCAA |
| Human | PSIP1 | Chromatin Readers | CGAGTAGACGAAGTTCCTGA |
| Human | PSIP1 | Chromatin Readers | GAGCAGCAGAATAAAGATGA |
| Human | PSIP1 | Chromatin Readers | ACGCAAGCAAGAGGAACAAA |
| Human | PSIP1 | Chromatin Readers | AAGTGAAGCAAGTTCATCCA |
| Human | PSIP1 | Chromatin Readers | GGATGAACTTGCTTCACTTC |
| Human | SCMH1 | Chromatin Readers | TGTAGGACTGCTTGAAGCAA |
| Human | SCMH1 | Chromatin Readers | AACTACTGTGGCAATACAGG |
| Human | SCMH1 | Chromatin Readers | ACCTGTCAGTCCAACTACTG |
| Human | SCMH1 | Chromatin Readers | TGTCGCTCCCATCAAGGCGC |
| Human | SCMH1 | Chromatin Readers | AAGGCGATGGTGGCTCCTTG |
| Human | SCMH1 | Chromatin Readers | GTGGCTGGGCAAATGAAATG |
| Human | SCMH1 | Chromatin Readers | AAGTTGTGGGAAGGCGATGG |
| Human | SCMH1 | Chromatin Readers | CAAACACGAGATCGATGGCA |
| Human | SCMH1 | Chromatin Readers | TTTCGCAAACACGAGATCGA |
| Human | SCMH1 | Chromatin Readers | TTGCGAAACAGGTCAGCGTG |
| Human | SCMH1 | Chromatin Readers | TGGTAGGAGAGCTTGAGTGC |
| Human | SCMH1 | Chromatin Readers | GTTTGCGAAACAGGTCAGCG |
| Human | PHF11 | Chromatin Readers | CATTATATTCGACATCTTTG |
| Human | PHF11 | Chromatin Readers | ACATTATATTCGACATCTTT |
| Human | PHF11 | Chromatin Readers | TCTCTGATTGTGCAAAGTAT |
| Human | PHF11 | Chromatin Readers | GACATTATATTCGACATCTT |
| Human | PHF11 | Chromatin Readers | CCTCGAACTCCATCAGACTG |
| Human | YTHDF3 | DNA and RNA Modifications | CAATTCAAGGGACACTCAAG |
| Human | YTHDF3 | DNA and RNA Modifications | CAAACCGGTTACCAATTCAA |
| Human | YTHDF3 | DNA and RNA Modifications | GTGGACTATAATGCGTATGC |
| Human | KIAA2026 | Chromatin Readers | AACTTACAGAGAAAGCGTGA |
| Human | KIAA2026 | Chromatin Readers | TGATGATCAATAACTTATTC |
| Human | KIAA2026 | Chromatin Readers | GCCCAAATCACCACTGCCAG |
| Human | KIAA2026 | Chromatin Readers | CTTACATTAGGCCACACCAC |
| Human | KIAA2026 | Chromatin Readers | GTAGTAGTTGTTCCAGCTAC |
| Human | SETD1B | Histone Methylation | CATGGGCAACATTATCCACG |
| Human | SETD1B | Histone Methylation | GACCGTGGCAAAGACCACCT |
| Human | SETD1B | Histone Methylation | CCCAAGACCAAGAAGCACCT |
| Human | SETD1B | Histone Methylation | GTGGATAATGTTGCCCATGA |
| Human | SETD1B | Histone Methylation | GCATGTCTGCGATCACCTGA |
| Human | SETD1B | Histone Methylation | GTCAGGTGATCGCAGACATG |

| | | | |
|---|---|---|---|
| Human | SETD1B | Histone Methylation | GGCAGTTCTCGGAGCCACAG |
| Human | SETD1B | Histone Methylation | TTCTCGGAGCCACAGAGGCA |
| Human | SETD1B | Histone Methylation | GTGGCTCCGAGAACTGCCGG |
| Human | SETD1B | Histone Methylation | CTGTGGCTCCGAGAACTGCC |
| Human | SETD1B | Histone Methylation | TGTGGCTCCGAGAACTGCCG |
| Human | SETD1B | Histone Methylation | GATGGTGGCCGAGGAAAGCA |
| Human | SETD1B | Histone Methylation | TGAGGAGGAGGTAGTGGCCA |
| Human | SETD1B | Histone Methylation | TACCTCCTCCTCATCCTCCG |
| Human | SETD1B | Histone Methylation | GGTAGTGGCCAGGGAAGAGG |
| Human | SETD1B | Histone Methylation | AGAGGAGGAGGAGGAGATGG |
| Human | SETD1B | Histone Methylation | CATCCGCGTCATCATCCTCG |
| Human | SETD1B | Histone Methylation | ATGTTCTGGAGCTCGCTCAG |
| Human | MCM5 | DNA and RNA Modifications | TGATGAAATTCCGGGACGAA |
| Human | RAD54L | Chromatin Remodeling | GGTCTGGTCATATGTGACGA |
| Human | RAD54L | Chromatin Remodeling | TCATATGTGACGAGGGACAC |
| Human | RAD54L | Chromatin Remodeling | GTGCAAGCCAGAAATTGACA |
| Human | RAD54L | Chromatin Remodeling | CGTATAAGTACCTTCGGGCA |
| Human | RAD54L | Chromatin Remodeling | GCTGTCTGCAGGGACCATTG |
| Human | RAD54L | Chromatin Remodeling | TGGTCCCTGCAGACAGCAGG |
| Human | RAD54L | Chromatin Remodeling | TCTACCGCCTGCTGTCTGCA |
| Human | YWHAE | Chromatin Readers | TCTGGTGTACCAGGCGAAGC |
| Human | YWHAE | Chromatin Readers | TCTTATATGCAACAGATAGG |
| Human | YWHAE | Chromatin Readers | GCTGCTGATTATTCTCCAGG |
| Human | YWHAE | Chromatin Readers | GTTGCATATAAGAATGTGAT |
| Human | YWHAE | Chromatin Readers | CATTCTTATATGCAACAGAT |
| Human | YWHAE | Chromatin Readers | AATGCTGCTGATTATTCTCC |
| Human | YWHAE | Chromatin Readers | GATTCGGGAATATCGGCAAA |
| Human | YWHAE | Chromatin Readers | GTTAATCTGTTGTGACATTC |
| Human | YWHAE | Chromatin Readers | TAGCAGCTTTATAAGCCACT |
| Human | NAF1 | Control | CGAGTCCGAATCCGAGTCCG |
| Human | NAF1 | Control | AGAGGAAGAAGACGACGATG |
| Human | NAF1 | Control | GGAGACCTCGGACTCGGATT |
| Human | NAF1 | Control | CGACGATGAGGAAGATGAAG |
| Human | NAF1 | Control | CTCGGACTCGGATTCGGACT |
| Human | NAF1 | Control | GCCCAACCCATGTTTACAGG |
| Human | NAF1 | Control | CCCACCTGTAAACATGGGTT |
| Human | NAF1 | Control | GAGTATGGTAAGTTAAGTAA |
| Human | NAF1 | Control | CCCAACCCATGTTTACAGGT |
| Human | SMARCE1 | Chromatin Remodeling | GAGGTACAGCAGAAAGGTCT |
| Human | SMARCE1 | Chromatin Remodeling | AGCCGCTGATGCCCTACATG |
| Human | SMARCE1 | Chromatin Remodeling | TACCTCATGTAGGGCATCAG |
| Human | SMARCE1 | Chromatin Remodeling | GCAAGATTATTGGTGGCATG |

| | | | |
|---|---|---|---|
| Human | SMARCE1 | Chromatin Remodeling | TGAGGTACAGCAGAAAGGTC |
| Human | SMARCE1 | Chromatin Remodeling | GTGCGGTCTGAAAGTAGAAG |
| Human | SMARCE1 | Chromatin Remodeling | ACTGCGCTCAGCTTGCTCTG |
| Human | SMARCE1 | Chromatin Remodeling | AATGCAGGTCCTCAAACGGC |
| Human | SMARCE1 | Chromatin Remodeling | TACCAGAGGATGCCGTGACC |
| Human | SMARCE1 | Chromatin Remodeling | ACAACAACTACAGGCTGGGA |
| Human | SMARCE1 | Chromatin Remodeling | GGATGCCGTGACCCGGCTGT |
| Human | SMARCE1 | Chromatin Remodeling | CACAAATGCCCAGCACACCA |
| Human | SMARCE1 | Chromatin Remodeling | GAACATTCCGATGGAGACAG |
| Human | SMARCE1 | Chromatin Remodeling | AGAAGGCACGTCTACTCCTG |
| Human | SMARCE1 | Chromatin Remodeling | GAGCAACAGTGCAACAGTGG |
| Human | PARP15 | Other Histone Modifications | TGACAGAAGGCGGGAAACAG |
| Human | PARP15 | Other Histone Modifications | AGAGACTTCTTGGCAGATCA |
| Human | PARP15 | Other Histone Modifications | GGTAACATATTCATGACAAG |
| Human | PARP15 | Other Histone Modifications | TGGAGATGTTCTGTACATCT |
| Human | PARP15 | Other Histone Modifications | TTGCACTTTAAGCATCCACC |
| Human | PARP15 | Other Histone Modifications | GCCATTGGAACAGGAAATGC |
| Human | PARP15 | Other Histone Modifications | TTATTGTAAACTCAACAGCA |
| Human | PARP15 | Other Histone Modifications | ACGCAATAGAGAAGATTGAG |
| Human | PARP15 | Other Histone Modifications | ACTTACTGGAGTCTTCACAA |
| Human | PARP15 | Other Histone Modifications | GTGCCATATGTCAATCAGCA |
| Human | PARP15 | Other Histone Modifications | CTCAATCTTCTCTATTGCGT |
| Human | PARP15 | Other Histone Modifications | GCTTTAATAGAAGTTGTGCT |
| Human | FYN | Other Histone Modifications | CAGCTCGGAAGGAGATTGGT |
| Human | FYN | Other Histone Modifications | ATTTCAAATATTGAACAGCT |
| Human | FYN | Other Histone Modifications | CCCTTTATGACTATGAAGCA |
| Human | FYN | Other Histone Modifications | ACAGCTCGGAAGGAGATTGG |
| Human | FYN | Other Histone Modifications | ATTCGCAAACTTGACAATGG |
| Human | FYN | Other Histone Modifications | GTGGATACTACATTACCACC |
| Human | FYN | Other Histone Modifications | AAGTGAATAGGCACCTTTGG |
| Human | FYN | Other Histone Modifications | ATACCCAGGCATGAACAACC |
| Human | FYN | Other Histone Modifications | CCCAGGCATGAACAACCGGG |
| Human | FYN | Other Histone Modifications | TGTGGACATGGCAGCACAGG |
| Human | PARP14 | Other Histone Modifications | GCCTCATATCCGCTCCAGCG |
| Human | PARP14 | Other Histone Modifications | TCTTCTCAGATACATCCACA |
| Human | PARP14 | Other Histone Modifications | GAGGACCTTAAGCATTATGG |
| Human | PARP14 | Other Histone Modifications | TTCACGTGGTAGCTCCGGAG |
| Human | PARP14 | Other Histone Modifications | CTACTAAGCACGAGATCCAA |
| Human | PARP14 | Other Histone Modifications | CTGTCGCTATGTGCTTCACG |
| Human | PARP14 | Other Histone Modifications | AATTGGACACAGTTGGACAA |
| Human | PARP14 | Other Histone Modifications | CTCATTCAATCTCAAAGCAG |
| Human | PARP14 | Other Histone Modifications | ACTCATTCAATCTCAAAGCA |

| | | | |
|---|---|---|---|
| Human | PARP14 | Other Histone Modifications | TTTGGCATTTCCTGTCCCAA |
| Human | PARP14 | Other Histone Modifications | ACTCACTGATACAATCTGCC |
| Human | PARP14 | Other Histone Modifications | CTCACTGATACAATCTGCCC |
| Human | PARP14 | Other Histone Modifications | GGCGCTGAACAGATAAACTG |
| Human | PARP14 | Other Histone Modifications | ATCTGAAATTAGAGGATGCA |
| Human | PARP14 | Other Histone Modifications | TCAGAATAGAGAAGATTGAG |
| Human | PARP14 | Other Histone Modifications | CTATGCCGGAAAGAATGCTG |
| Human | PARP14 | Other Histone Modifications | ATTTCGATTGACGTGTGGCA |
| Human | PARP14 | Other Histone Modifications | TAGTGATCCTGAGTACAACA |
| Human | PARP14 | Other Histone Modifications | GAATGCTGTGGCATATGGAA |
| Human | NSUN2 | DNA and RNA Modifications | TAACAGCTCACTCACATCAG |
| Human | NSUN2 | DNA and RNA Modifications | TTTCAGACCTTAACGTGACA |
| Human | NSUN2 | DNA and RNA Modifications | CTTAATTTCGAAATTAATGA |
| Human | NSUN2 | DNA and RNA Modifications | CCTTCGGCGGAGCTGACCTG |
| Human | CDC6 | DNA and RNA Modifications | TGCTGAATTGCATGTCCTTG |
| Human | CDC6 | DNA and RNA Modifications | GGTATTAGCAATACCTAAGG |
| Human | CDC6 | DNA and RNA Modifications | AAATCAAAGAGGTCACTCTG |
| Human | AFF2 | Transcription Elongation | TATTCACCCTGCAGAAGACA |
| Human | AFF2 | Transcription Elongation | GGCATTGCCACATTCAGTGA |
| Human | AFF2 | Transcription Elongation | CTTGTCCGCTACGTTCGCCA |
| Human | AFF2 | Transcription Elongation | CCGGATTCTGCAAGACCTCA |
| Human | AFF2 | Transcription Elongation | TTGCAGAATCCGGCTTAAGC |
| Human | AFF3 | Transcription Elongation | GTATACCCATCAACTCTTTG |
| Human | KDM1B | Histone Methylation | AGATTGCAGACCTAATTGTA |
| Human | KDM1B | Histone Methylation | CCTGTTATCAATAGCAAATA |
| Human | KDM1B | Histone Methylation | CTGAGCCCGCCGTCTACCCG |
| Human | KDM1B | Histone Methylation | TGGTGCGGTCGATAGTGTGC |
| Human | CDC6 | DNA and RNA Modifications | GAAGAACAACCATGGCAACT |
| Human | CDC6 | DNA and RNA Modifications | GCTCTTTGATGCTCTTGATC |
| Human | CDC6 | DNA and RNA Modifications | GTAGGCTTCATATAACTGGT |
| Human | CDC6 | DNA and RNA Modifications | CAAGGAAACCCGTTTGACAA |
| Human | CENPF | Chromatin Remodeling | CACCGACGCCAAGAAACGGG |
| Human | CENPF | Chromatin Remodeling | TGTTCTGCAGGTGAGGAGCC |
| Human | CENPF | Chromatin Remodeling | CTCACCTTTCCTGGCCAAGG |
| Human | CENPF | Chromatin Remodeling | ATATCCTCCTTGGCCAGGAA |
| Human | CENPF | Chromatin Remodeling | CCAGATTGTTAATCCACCAA |
| Human | CENPW | Chromatin Remodeling | CAGATGATGAAGCCTGATGC |
| Human | CENPW | Chromatin Remodeling | CTGTACACACACTGGCTGCA |
| Human | CENPW | Chromatin Remodeling | AGGCGCCCGAGGACGCGCCT |
| Human | CENPW | Chromatin Remodeling | CATAGGCTGCCTGAACGGCG |
| Human | KMT5A | Histone Methylation | AGCACTTCAGGAGAGCTTAT |
| Human | KMT5A | Histone Methylation | ACAGCAAGAGGAAACCCTAG |

| | | | |
|---|---|---|---|
| Human | KMT5A | Histone Methylation | GTGGCCTACATAGAGTCGCA |
| Human | JMJD8 | Histone Methylation | GAGTATACCCATCAACTCTT |
| Human | JMJD8 | Histone Methylation | AGTTCCTAATTCAGTACAAC |
| Human | JMJD8 | Histone Methylation | AATCTCCATGTCCCATGAAG |
| Human | JMJD8 | Histone Methylation | CAGTCTCAGAGAGTAGGTCA |
| Human | NARFL | Control | GAGTTGGAACTACGCCTGCA |
| Human | NARFL | Control | TCCACAGCCACTGAGTTGGT |
| Human | NARFL | Control | CTATATGCTACCTTATGATG |
| Human | NARFL | Control | TTGGAAGAGCATATGAACAC |
| Human | GMPPB | Control | TTACCACTATAATATTGACT |
| Human | GMPPB | Control | CATACGTACAGGCTGTCCAG |
| Human | GMPPB | Control | GGCGCCCGAGGACGCGCCTG |
| Human | GMPPB | Control | GGCACGGACTCGGAGTGTGC |
| Human | HMGN1 | Transcription Elongation | GGTACTGAGTTTGAGCCAGA |
| Human | HMGN1 | Transcription Elongation | CCAACGGCTAAACCTCCACC |
| Human | HMGN1 | Transcription Elongation | CCAGTTATTACCCAAGATGG |
| Human | HMGN1 | Transcription Elongation | GAAGCAGCTTTGTGTAGCCA |
| Human | HMGN1 | Transcription Elongation | GATGATGCAGGCTGAAAGAG |
| Human | HAT1 | Histone Acetylation | GCCACCAAGCAGTTCTCCCG |
| Human | HAT1 | Histone Acetylation | GATGACCACCGATAGGAGGT |
| Human | KMT5B | Histone Methylation | TGACAGATTGACCTCATCGA |
| Human | KMT5B | Histone Methylation | CCCGTACGCACCCGGCTGAA |
| Human | KMT5B | Histone Methylation | CTGGACGTGTTCCCGGTGCG |
| Human | KMT5B | Histone Methylation | CGTCCCGAGCTGGTCAATGA |
| Human | KMT5B | Histone Methylation | CTTCATTGACCAGCTCGGGA |
| Human | DOT1L | Histone Methylation | AAGAGTTGATGGGTATACTC |
| Human | DOT1L | Histone Methylation | CTTTAGATGCATATTCAACA |
| Human | DOT1L | Histone Methylation | TCTGGCCGATGCGGGCACTT |
| Human | DOT1L | Histone Methylation | TGTGTGTATCCGGCGGTGCA |
| Human | DOT1L | Histone Methylation | ATCGGCCAGAACTGCAGCAT |
| Human | ING1 | Chromatin Readers | CGACGGGAGGACTTCTCTCA |
| Human | ING1 | Chromatin Readers | GTGCCCACCTGCTTGGCGGA |
| Human | ING1 | Chromatin Readers | GCTGTCTAATGACGGGATGT |
| Human | NARFL | Control | TGAGCTGAGTAGGATCAGAT |
| Human | NARFL | Control | TAAATAGCTTGCAGCTACAT |
| Human | NARFL | Control | AAGAAGATGAAGCCCTACGA |
| Human | NARFL | Control | CAGTATCCATTCTTTATGGC |
| Human | JMJD8 | Histone Methylation | AGTATACCCATCAACTCTTT |
| Human | JMJD8 | Histone Methylation | ACCTCCTTCTCAATGGACGT |
| Human | JMJD8 | Histone Methylation | CAGGTCCCAGGCTCACATTG |
| Human | JMJD8 | Histone Methylation | AAGATGGTGTGTGTATCCGG |
| Human | KDM4B | Histone Methylation | GTACTGCCAGCGCATGCAGG |

| | | | |
|---|---|---|---|
| Human | KDM4B | Histone Methylation | CATACCGCATCGATAAGTCT |
| Human | KDM4B | Histone Methylation | AAATACTCACTAGATCCTCT |
| Human | KDM4B | Histone Methylation | TGGGAGGGAACAATGATCTG |
| Human | KDM4B | Histone Methylation | TCTGCAGAGGAGCCCAAGGT |
| Human | KDM4B | Histone Methylation | TGCGGCTCGGCGACTTGGTG |
| Human | KDM4B | Histone Methylation | GAGTCTGCGGCTCGGCGACT |
| Human | KDM4B | Histone Methylation | GTCTTCAGCAATGATATCGT |
| Human | CENPF | Chromatin Remodeling | CTTTGTGGTGGAATACCACG |
| Human | CENPF | Chromatin Remodeling | AAGGACTTGAAGAAACCTCG |
| Human | CENPF | Chromatin Remodeling | AAGAAGCATTTCTTTCCGCG |
| Human | CENPF | Chromatin Remodeling | GGGTAAACGATTCCAGCAAG |
| Human | CENPF | Chromatin Remodeling | ATGCTGTCGAAGAGTTCCTC |
| Human | GLYR1 | Chromatin Readers | TCTGATGAGTATCTATTACA |
| Human | GLYR1 | Chromatin Readers | GAGAACATGCTACTTAGACA |
| Human | GLYR1 | Chromatin Readers | TAAGCTAAGCAGTGATGAAG |
| Human | GLYR1 | Chromatin Readers | GCAGCCCAACTGCAGAACCT |
| Human | GLYR1 | Chromatin Readers | GAAACATCCGCCAGTACAGG |
| Human | GLYR1 | Chromatin Readers | TCAGTGTCATGTACTCCACA |
| Human | GLYR1 | Chromatin Readers | CCTGCTGCGTTTATAAACCA |
| Human | GLYR1 | Chromatin Readers | CTGGTCCAGTACTCCCAACA |
| Human | GLYR1 | Chromatin Readers | CAGGATGACGGGCCTGACGA |
| Human | GMPPB | Control | GATGGAAATATGTGAAAGTC |
| Human | GMPPB | Control | TCTTTAGAATTACCTGTAAG |
| Human | GMPPB | Control | GAGTTTGTGCGGCGATTCCG |
| Human | ELP4 | Transcription Elongation | AGGTACTGAGTTTGAGCCAG |
| Human | ELP4 | Transcription Elongation | ATCTGACAAGGTTGTGACAC |
| Human | ELP4 | Transcription Elongation | CACTGACAGCGAATCTAATG |
| Human | ELP4 | Transcription Elongation | CCACGCCGCAGAAATGTCAG |
| Human | ELP4 | Transcription Elongation | GAAGCACAAAGCTGATGCAC |
| Human | KDM4B | Histone Methylation | AACATCACGGAGCACGAGGT |
| Human | KDM4B | Histone Methylation | TACTCACTAGATCCTCTGGG |
| Human | KDM4B | Histone Methylation | ATTCCTCACATCATCGTCCG |
| Human | GLYR1 | Chromatin Readers | CTGGCAGGTACATCACCTAC |
| Human | GLYR1 | Chromatin Readers | AGTTAGTATACCACAGTGCG |
| Human | GLYR1 | Chromatin Readers | GTTTCTCAAAGCCAGGTACA |
| Human | GLYR1 | Chromatin Readers | GAAACCTCATCCTCGCACTG |
| Human | GLYR1 | Chromatin Readers | AGTAACCTGGAGTACGCCTG |
| Human | GLYR1 | Chromatin Readers | TTCCTTGGCGGACATTCCAG |
| Human | GLYR1 | Chromatin Readers | GATTTAGGCCTATTAAAGGA |
| Human | GLYR1 | Chromatin Readers | ACTCCTACCACAAAGGTACG |
| Human | GLYR1 | Chromatin Readers | AGGCCGCGTACCTTTGTGGT |
| Human | CENPF | Chromatin Remodeling | TGGGCAAATACCTCTAGCCA |

| | | | |
|---|---|---|---|
| Human | CENPF | Chromatin Remodeling | TGCTAGTGCCTGGATCAAAG |
| Human | CENPF | Chromatin Remodeling | TTCCACTTCCCATGAGACCA |
| Human | SMC1B | Chromatin Remodeling | GAGATCGACGCGAAATACCA |
| Human | SMC1B | Chromatin Remodeling | AGAGAAATGTCTCGCTGATG |
| Human | POLR2E | Transcription Elongation | GGAATCTACAGGAAACGAGA |
| Human | POLR2E | Transcription Elongation | GCAACAGAATCGCAAACTTA |
| Human | POLR2E | Transcription Elongation | TCTCTGCCCAGTCTGAAGAA |
| Human | POLR2E | Transcription Elongation | TGCCGCGGCTTCCACTCCTT |
| Human | POLR2E | Transcription Elongation | TACGCCATCCCACCAGAGCA |
| Human | POLR2E | Transcription Elongation | CATCAAGGTAATCTAGAGCG |
| Human | POLR2E | Transcription Elongation | CATGACGTTTCGCCCAACCA |
| Human | POLR2E | Transcription Elongation | TAAAGACTTCAACAAATACG |
| Human | POLR2E | Transcription Elongation | GCGGCTTCCACTCCTTCGGG |
| Human | POLR2E | Transcription Elongation | CCACATGCCGAAGTACAGGT |
| Human | KNTC1 | Chromatin Remodeling | TCAGTTAAAGGAGCTAAATG |
| Human | KNTC1 | Chromatin Remodeling | AGATTTACTTAAGGGTAGAG |
| Human | KNTC1 | Chromatin Remodeling | CTCCAAGCCCAAGAGGAGAT |
| Human | KNTC1 | Chromatin Remodeling | TTTCTCTCCTGCTTCATCAG |
| Human | POP5 | Control | GTTTCAGCTTGACAGTCTCG |
| Human | POP5 | Control | GTAATTCTAAAGAAGAGCAG |
| Human | KMT5A | Histone Methylation | CCACCTTAAGAAATTAGATG |
| Human | KMT5A | Histone Methylation | CTGGTTGCACAGACAGTACG |
| Human | KMT5A | Histone Methylation | GGGCTCAATCATAAACCCAA |
| Human | KMT5A | Histone Methylation | GACAGTACGTGGGTTCGTTG |
| Human | KMT5A | Histone Methylation | GCCGTGGAAACGTGAGTGAC |
| Human | KMT5B | Histone Methylation | ATTCCACCACAAAGTCACCC |
| Human | KMT5B | Histone Methylation | GCCGTAGTCAATCCACCGCA |
| Human | KMT5B | Histone Methylation | TCCTTCCGGCACGTGCACTG |
| Human | KMT5B | Histone Methylation | ACTTGCCGCAGGCGATCAGG |
| Human | KMT5B | Histone Methylation | CTGCAGGCAGCACTTGCCGC |
| Human | KDM1B | Histone Methylation | TTGATAACAGGAGCAGGCGC |
| Human | KDM1B | Histone Methylation | CTACGGTCGTAAGGTCAGCA |
| Human | KDM1B | Histone Methylation | GAGACCATCCGGTGAGTGCA |
| Human | KDM4B | Histone Methylation | AATCCGCAAGACCATCATGC |
| Human | KDM4B | Histone Methylation | ACCACACTCTGCTAACTCCC |
| Human | KDM4B | Histone Methylation | GGTACAGTGACTAATACCTG |
| Human | KDM4B | Histone Methylation | TTTCCCACAAACTGTTACAG |
| Human | KDM1B | Histone Methylation | GCGAGTGTTACACTTGCGAA |
| Human | KDM1B | Histone Methylation | TGCTCCACATACTCCTGGAA |
| Human | KDM1B | Histone Methylation | ACAGCTTTGGAATCGCAGGT |
| Human | ING1 | Chromatin Readers | TGATCGTGAACATGGTCCAA |
| Human | ING1 | Chromatin Readers | GGTCAACCATCCGACTCCCA |

| | | | |
|---|---|---|---|
| Human | ING1 | Chromatin Readers | CAGCTGCCATGGGAGTCGGA |
| Human | Non Targeting Control | Non Targeting Control | AAAAAGCTTCCGCCTGATGG |
| Human | Non Targeting Control | Non Targeting Control | AACTAGCCCGAGCAGCTTCG |
| Human | Non Targeting Control | Non Targeting Control | AAGTGACGGTGTCATGCGGG |
| Human | Non Targeting Control | Non Targeting Control | AATATTTGGCTCGGCTGCGC |
| Human | Non Targeting Control | Non Targeting Control | AATCGACTCGAACTTCGTGT |
| Human | Non Targeting Control | Non Targeting Control | AATGGGTTGCCGGGAGTAAA |
| Human | Non Targeting Control | Non Targeting Control | ACAGCGCTCTCGTGTACTAT |
| Human | Non Targeting Control | Non Targeting Control | ACCACTAATGAGATTCTTGT |
| Human | Non Targeting Control | Non Targeting Control | ACTAGCCTGTTCGCGAGTAG |
| Human | Non Targeting Control | Non Targeting Control | ACTGCGGAGCGCCCAATATC |
| Human | Non Targeting Control | Non Targeting Control | AGCATTTGCGCGGCAACTGT |
| Human | Non Targeting Control | Non Targeting Control | AGCGCTCTGGTTGCATCCCT |
| Human | Non Targeting Control | Non Targeting Control | AGGGGCAGGGCTATCTTATG |
| Human | Non Targeting Control | Non Targeting Control | AGGTCGGCCGAACATACGGT |
| Human | Non Targeting Control | Non Targeting Control | ATACGAGGCGCTTTTCTTTG |
| Human | Non Targeting Control | Non Targeting Control | ATAGCAGGACGAGGTTCCTT |
| Human | Non Targeting Control | Non Targeting Control | ATCAGGCCTTGTCCGTGATT |
| Human | Non Targeting Control | Non Targeting Control | ATCGATATACCGCCATAAAA |
| Human | Non Targeting Control | Non Targeting Control | ATCTGAGCGTTTTCGGCCGC |
| Human | Non Targeting Control | Non Targeting Control | ATCTTCTCGACGAAAATGCG |
| Human | Non Targeting Control | Non Targeting Control | ATGACATTGCGCGTCTACGG |
| Human | Non Targeting Control | Non Targeting Control | ATGGAAGAGCGTCATGACTT |
| Human | Non Targeting Control | Non Targeting Control | ATTAAACGACACCTTATTCT |
| Human | Non Targeting Control | Non Targeting Control | ATTTATAAACACAGGGTCGC |
| Human | Non Targeting Control | Non Targeting Control | CAATATCGGGTGCTACAGGA |
| Human | Non Targeting Control | Non Targeting Control | CAGCGCCGAAACTCTTTCCG |
| Human | Non Targeting Control | Non Targeting Control | CCAATCTTGAACGTCATGTT |
| Human | Non Targeting Control | Non Targeting Control | CCAGGCTGAAGTTCGTACCT |
| Human | Non Targeting Control | Non Targeting Control | CCAGTGCCCTTTTGTCGCAA |
| Human | Non Targeting Control | Non Targeting Control | CCAGTTGCTCTGGGGGAACA |
| Human | Non Targeting Control | Non Targeting Control | CCATCACCGATCGTGAGCCT |
| Human | Non Targeting Control | Non Targeting Control | CCCGATGGACTATACCGAAC |
| Human | Non Targeting Control | Non Targeting Control | CCGTCTCCGCATCGTCTTTT |
| Human | Non Targeting Control | Non Targeting Control | CCTAGTTCATAAGCTACGCC |
| Human | Non Targeting Control | Non Targeting Control | CGAAGTCTTTCTTAGATGGT |
| Human | Non Targeting Control | Non Targeting Control | CGGCACTAGAAGTTTTTGAA |
| Human | Non Targeting Control | Non Targeting Control | CGGCCGCATCCTGTTATATT |
| Human | Non Targeting Control | Non Targeting Control | CGGGACGTCGCGAAAATGTA |
| Human | Non Targeting Control | Non Targeting Control | CGTAGTAAATATCTAGCTAA |
| Human | Non Targeting Control | Non Targeting Control | CTAACGGACTGCAGAACGGA |
| Human | Non Targeting Control | Non Targeting Control | CTCGGCCCACTTGCGTTATA |

| | | | |
|---|---|---|---|
| Human | Non Targeting Control | Non Targeting Control | CTGAAGGTGTCTGGCAGAGC |
| Human | Non Targeting Control | Non Targeting Control | CTGACGTGTCTGAAATGAGT |
| Human | Non Targeting Control | Non Targeting Control | CTGCCCCAGGCGTAATCCTC |
| Human | Non Targeting Control | Non Targeting Control | CTGGATCGCCCGCAGAAATA |
| Human | Non Targeting Control | Non Targeting Control | CTTAAGGCGAGAAAAATTAG |
| Human | Non Targeting Control | Non Targeting Control | CTTACAATCGTCGGTCCAAT |
| Human | Non Targeting Control | Non Targeting Control | CTTACCCCTATTATAATGAA |
| Human | Non Targeting Control | Non Targeting Control | GAAATGCTATGCTTCGGTTC |
| Human | Non Targeting Control | Non Targeting Control | GAACTCAACCAGAGGGCCAA |
| Human | Non Targeting Control | Non Targeting Control | GAAGACGTGCTGGCGTCACC |
| Human | Non Targeting Control | Non Targeting Control | GACAGTGAAATTAGCTCCCA |
| Human | Non Targeting Control | Non Targeting Control | GACCTTCATTGAAGAAAAGC |
| Human | Non Targeting Control | Non Targeting Control | GACTTCTAGAATATAAAAGA |
| Human | Non Targeting Control | Non Targeting Control | GAGAGTGCGCCTTGATAGTA |
| Human | Non Targeting Control | Non Targeting Control | GAGATGATAACTTAATTTGT |
| Human | Non Targeting Control | Non Targeting Control | GATTCATACTAAACACTCTA |
| Human | Non Targeting Control | Non Targeting Control | GCCAGCGGGGATATGGTGAA |
| Human | Non Targeting Control | Non Targeting Control | GCCCCGCCGCCCTCCCCTCC |
| Human | Non Targeting Control | Non Targeting Control | GCCTAGTCTCGGTAAGAGTG |
| Human | Non Targeting Control | Non Targeting Control | GCCTATCGGCATTCCCACTG |
| Human | Non Targeting Control | Non Targeting Control | GCGGGCAGAACGACCCTGAC |
| Human | Non Targeting Control | Non Targeting Control | GCTCCCATCCATAGTAAAAA |
| Human | Non Targeting Control | Non Targeting Control | GCTGGAGAGACAATTCTACT |
| Human | Non Targeting Control | Non Targeting Control | GGAATCCGGAGCTCATGAGG |
| Human | Non Targeting Control | Non Targeting Control | GGATTAATTCGCTAAATGAT |
| Human | Non Targeting Control | Non Targeting Control | GGATTGAATGGCTAACGCGG |
| Human | Non Targeting Control | Non Targeting Control | GGCCACAAAAACTCGCTAAG |
| Human | Non Targeting Control | Non Targeting Control | GGCTTACGTGGGGGGCAAAA |
| Human | Non Targeting Control | Non Targeting Control | GGGACGCGAAAGAAACCAGT |
| Human | Non Targeting Control | Non Targeting Control | GGGAGGTGGCTTTAGGTTTT |
| Human | Non Targeting Control | Non Targeting Control | GGGCAGAAGTTGCTGTCCTG |
| Human | Non Targeting Control | Non Targeting Control | GGTATTCCGAAGTCCTCCGC |
| Human | Non Targeting Control | Non Targeting Control | GGTCTCACCTGCACCCCGAA |
| Human | Non Targeting Control | Non Targeting Control | GGTGCGGTCTGCTTGTAGAT |
| Human | Non Targeting Control | Non Targeting Control | GTAGGCGCGCCGCTCTCTAC |
| Human | Non Targeting Control | Non Targeting Control | GTATTACTGATATTGGTGGG |
| Human | Non Targeting Control | Non Targeting Control | GTCATCAGCGATTTGACGAG |
| Human | Non Targeting Control | Non Targeting Control | GTCCTCATCCGGTCAGGCTG |
| Human | Non Targeting Control | Non Targeting Control | GTGAACTGCAATCTTATTAT |
| Human | Non Targeting Control | Non Targeting Control | GTTATCCTGTCGAAGTAAAG |
| Human | Non Targeting Control | Non Targeting Control | GTTCATTTCCAAGTCCGCTG |
| Human | Non Targeting Control | Non Targeting Control | GTTGGCATATTGGCCCAGAC |

| | | | |
|---|---|---|---|
| Human | Non Targeting Control | Non Targeting Control | TACGGATCACCAAATCTTAG |
| Human | Non Targeting Control | Non Targeting Control | TACGTCATTAAGAGTTCAAC |
| Human | Non Targeting Control | Non Targeting Control | TAGCTCGAGTCATTTCTCTA |
| Human | Non Targeting Control | Non Targeting Control | TCAGCAAAGGACGAAACAAA |
| Human | Non Targeting Control | Non Targeting Control | TCATGCTTGCTTGGGCAAAA |
| Human | Non Targeting Control | Non Targeting Control | TCGATCGAGGTTGCATTCGG |
| Human | Non Targeting Control | Non Targeting Control | TCGGCATACGGGACACACGC |
| Human | Non Targeting Control | Non Targeting Control | TCGGGCAGTGAGTACAATAC |
| Human | Non Targeting Control | Non Targeting Control | TCGGGGACCACCCACGATCC |
| Human | Non Targeting Control | Non Targeting Control | TGAATCGTAACCTCGCCATT |
| Human | Non Targeting Control | Non Targeting Control | TGAGTCTTACTAGGTCCTGT |
| Human | Non Targeting Control | Non Targeting Control | TGCGGCGTAATGCTTGAAAG |
| Human | Non Targeting Control | Non Targeting Control | TGGCGGCCCAAACTTAACAC |
| Human | Non Targeting Control | Non Targeting Control | TGGTTCCGTAGGTCGGTATA |
| Human | Non Targeting Control | Non Targeting Control | TGTTCTACTTTCGAAGTTAA |
| Human | Non Targeting Control | Non Targeting Control | TTCAATTCACCGAGGGCGCA |
| Human | Non Targeting Control | Non Targeting Control | TTTGGGTTTATCCGCCCCCA |

**TABLE S3**

| Group_1 | Group_2 | DE_gene | p_val | avg_logFC | pct.1 | pct.2 |
|---|---|---|---|---|---|---|
| AFF2 | NTC | LYZ | 4.27E-98 | 0.99809759 | 0.988 | 0.926 |
| AFF2 | NTC | GLO1 | 1.02E-71 | -0.6540925 | 0.998 | 1 |
| AFF2 | NTC | CLEC2B | 5.63E-68 | -0.7556887 | 0.909 | 0.985 |
| AFF2 | NTC | ACTB | 8.59E-51 | 0.27888813 | 0.998 | 1 |
| AFF2 | NTC | BMI1 | 3.29E-50 | -0.4819865 | 0.723 | 0.95 |
| AFF2 | NTC | FTL | 6.27E-45 | 0.27410961 | 1 | 1 |
| AFF2 | NTC | SRGN | 6.22E-43 | 0.45404143 | 1 | 0.997 |
| AFF2 | NTC | LINC02470 | 1.99E-38 | -0.384059 | 0.683 | 0.929 |
| AFF2 | NTC | SH3BGRL3 | 9.18E-36 | 0.32129887 | 0.998 | 0.997 |
| AFF2 | NTC | INPP4B | 1.10E-34 | -0.3678173 | 0.432 | 0.776 |
| AFF2 | NTC | RNF220 | 1.18E-34 | -0.40469 | 0.962 | 0.987 |
| AFF2 | NTC | B2M | 4.91E-34 | 0.32312568 | 0.998 | 0.998 |
| AFF2 | NTC | TMEM163 | 1.14E-33 | -0.3452748 | 0.6 | 0.881 |
| AFF2 | NTC | RNASE3 | 1.21E-33 | 0.48064583 | 0.909 | 0.883 |
| AFF2 | NTC | RETN | 2.94E-32 | 0.59415395 | 0.891 | 0.809 |
| AFF2 | NTC | LGALS1 | 1.39E-30 | 0.4049446 | 0.992 | 0.998 |
| AFF2 | NTC | RNASE2 | 7.12E-30 | 0.4109895 | 0.98 | 0.975 |
| AFF2 | NTC | CD36 | 2.22E-29 | -0.507827 | 0.731 | 0.914 |
| AFF2 | NTC | TLN2 | 1.98E-28 | -0.286916 | 0.451 | 0.794 |
| AFF2 | NTC | CLEC2A | 5.13E-28 | -0.2751867 | 0.372 | 0.726 |
| AFF2 | NTC | S100A4 | 2.24E-27 | 0.33239011 | 0.994 | 1 |
| AFF2 | NTC | CLECL1 | 3.15E-25 | -0.2944038 | 0.778 | 0.946 |
| AFF2 | NTC | ACTR2 | 2.31E-24 | 0.25581783 | 0.99 | 0.997 |
| AFF2 | NTC | GSTP1 | 6.38E-24 | -0.2666467 | 1 | 1 |
| AFF2 | NTC | CDA | 6.53E-24 | 0.28052027 | 0.937 | 0.919 |
| AFF2 | NTC | STXBP2 | 6.74E-23 | 0.41026765 | 0.921 | 0.926 |
| AFF2 | NTC | MEIS1 | 3.59E-22 | -0.3124055 | 0.804 | 0.952 |
| AFF2 | NTC | VIM | 4.44E-22 | 0.53906391 | 0.735 | 0.624 |
| AFF2 | NTC | METTL9 | 9.37E-22 | 0.36804037 | 0.859 | 0.861 |
| AFF2 | NTC | NPC1 | 3.27E-21 | -0.2885592 | 0.495 | 0.774 |
| AFF2 | NTC | FABP5 | 4.26E-21 | -0.2868521 | 0.875 | 0.983 |
| AFF2 | NTC | NME1 | 2.64E-19 | -0.2540496 | 0.885 | 0.974 |
| AFF2 | NTC | C12orf75 | 3.19E-19 | -0.2742737 | 0.723 | 0.919 |
| AFF2 | NTC | S100P | 4.16E-19 | 0.3259584 | 0.947 | 0.964 |
| AFF2 | NTC | CAT | 1.07E-18 | 0.2915952 | 0.887 | 0.883 |
| AFF2 | NTC | NEGR1 | 1.23E-18 | -0.3467419 | 0.507 | 0.771 |
| AFF2 | NTC | H2AFZ | 7.87E-18 | 0.28870888 | 0.99 | 0.995 |
| AFF2 | NTC | SMC4 | 2.33E-17 | -0.2774277 | 0.871 | 0.964 |
| AFF2 | NTC | HSPA8 | 6.22E-17 | -0.2542623 | 0.964 | 0.99 |
| AFF2 | NTC | AC007100.1 | 3.49E-16 | -0.2827883 | 0.493 | 0.726 |
| AFF2 | NTC | CSTA | 6.51E-16 | 0.32860275 | 0.606 | 0.45 |
| AFF2 | NTC | EBNA1BP2 | 6.61E-16 | -0.270402 | 0.752 | 0.926 |

| | | | | | | |
|---|---|---|---|---|---|---|
| AFF2 | NTC | AL355922.1 | 3.77E-15 | 0.30515363 | 0.798 | 0.772 |
| AFF2 | NTC | SLPI | 2.72E-14 | 0.34823806 | 0.485 | 0.32 |
| AFF2 | NTC | CXCR4 | 7.40E-14 | 0.25245044 | 0.949 | 0.941 |
| AFF2 | NTC | SNX10 | 3.18E-12 | 0.26412428 | 0.636 | 0.554 |
| AFF2 | NTC | ZEB2 | 3.46E-12 | -0.2703941 | 0.893 | 0.949 |
| AFF2 | NTC | HCST | 4.01E-12 | 0.28877018 | 0.97 | 0.988 |
| AFF2 | NTC | ANXA1 | 6.56E-12 | 0.29386538 | 0.507 | 0.37 |
| AFF2 | NTC | ETS1 | 1.66E-11 | 0.26139004 | 0.717 | 0.719 |
| AFF2 | NTC | EPDR1 | 9.08E-10 | 0.25503348 | 0.749 | 0.772 |
| AFF2 | NTC | S100A9 | 7.92E-05 | 0.27524134 | 0.412 | 0.351 |
| KMT2A | NTC | RNF220 | 8.55E-59 | -0.5822785 | 0.864 | 0.987 |
| KMT2A | NTC | GLO1 | 2.01E-29 | -0.3371413 | 0.992 | 1 |
| KMT2A | NTC | MEIS1 | 7.01E-28 | -0.324166 | 0.735 | 0.952 |
| KMT2A | NTC | SMC4 | 6.11E-26 | -0.3250363 | 0.891 | 0.964 |
| KMT2A | NTC | PRTN3 | 1.07E-25 | 0.66240917 | 0.924 | 0.891 |
| KMT2A | NTC | ELANE | 1.51E-23 | 0.50047709 | 0.986 | 0.992 |
| KMT2A | NTC | C12orf75 | 1.15E-18 | -0.2691849 | 0.776 | 0.919 |
| KMT2A | NTC | CD36 | 1.30E-17 | -0.3142879 | 0.781 | 0.914 |
| KMT2A | NTC | AZU1 | 1.21E-16 | 0.36685747 | 0.975 | 0.979 |
| KMT2A | NTC | LYZ | 2.59E-16 | 0.36223327 | 0.949 | 0.926 |
| KMT2A | NTC | SRGN | 1.77E-15 | 0.30209141 | 1 | 0.997 |
| KMT2A | NTC | SLPI | 4.11E-13 | 0.29145835 | 0.492 | 0.32 |
| KMT2A | NTC | FAM107B | 2.32E-11 | 0.26392794 | 0.947 | 0.946 |
| CSNK2B | NTC | CSNK2B | 1.06E-22 | -0.2633086 | 0.869 | 0.98 |
| DOT1L | NTC | CLEC2B | 2.57E-120 | -1.3477603 | 0.656 | 0.985 |
| DOT1L | NTC | RNF220 | 1.67E-111 | -1.1120473 | 0.725 | 0.987 |
| DOT1L | NTC | MEIS1 | 6.50E-99 | -0.8222352 | 0.326 | 0.952 |
| DOT1L | NTC | BMI1 | 2.05E-93 | -0.7659432 | 0.462 | 0.95 |
| DOT1L | NTC | LYZ | 3.58E-91 | 1.08823445 | 0.987 | 0.926 |
| DOT1L | NTC | GLO1 | 2.57E-85 | -0.7690928 | 0.996 | 1 |
| DOT1L | NTC | CLEC2A | 1.43E-71 | -0.4562753 | 0.15 | 0.726 |
| DOT1L | NTC | CD36 | 5.89E-70 | -0.8382619 | 0.502 | 0.914 |
| DOT1L | NTC | SRGN | 1.00E-62 | 0.68301348 | 0.998 | 0.997 |
| DOT1L | NTC | TMSB4X | 8.51E-61 | 0.46512985 | 1 | 1 |
| DOT1L | NTC | RNASE3 | 1.84E-53 | 0.5804192 | 0.946 | 0.883 |
| DOT1L | NTC | ACTB | 4.34E-53 | 0.35651807 | 1 | 1 |
| DOT1L | NTC | HOXA9 | 7.52E-51 | -0.3583168 | 0.19 | 0.672 |
| DOT1L | NTC | GSTP1 | 1.36E-50 | -0.4416339 | 0.996 | 1 |
| DOT1L | NTC | PPP3R1 | 9.52E-50 | -0.469858 | 0.775 | 0.967 |
| DOT1L | NTC | SLPI | 1.36E-48 | 0.68040385 | 0.676 | 0.32 |
| DOT1L | NTC | FTL | 1.91E-48 | 0.33224108 | 0.998 | 1 |
| DOT1L | NTC | DNAJC1 | 1.17E-47 | -0.5000946 | 0.56 | 0.899 |
| DOT1L | NTC | HS3ST2 | 8.26E-47 | -0.5329482 | 0.578 | 0.911 |
| DOT1L | NTC | COMMD3 | 6.26E-46 | -0.3250345 | 0.507 | 0.883 |
| DOT1L | NTC | FAM107B | 9.65E-46 | 0.58474477 | 0.967 | 0.946 |

| | | | | | | |
|---|---|---|---|---|---|---|
| DOT1L | NTC | INPP4B | 4.50E-44 | -0.4246242 | 0.375 | 0.776 |
| DOT1L | NTC | MNDA | 2.35E-43 | 0.48471947 | 0.993 | 0.983 |
| DOT1L | NTC | LINC02470 | 1.60E-41 | -0.3945908 | 0.658 | 0.929 |
| DOT1L | NTC | PLEK | 2.22E-41 | -0.374691 | 0.98 | 0.99 |
| DOT1L | NTC | HOXA10 | 1.04E-39 | -0.3568797 | 0.6 | 0.896 |
| DOT1L | NTC | CLECL1 | 6.99E-37 | -0.3869002 | 0.746 | 0.946 |
| DOT1L | NTC | GAS7 | 1.30E-36 | -0.3837277 | 0.549 | 0.876 |
| DOT1L | NTC | EPDR1 | 2.58E-36 | 0.43904699 | 0.848 | 0.772 |
| DOT1L | NTC | C12orf75 | 7.60E-36 | -0.4122772 | 0.692 | 0.919 |
| DOT1L | NTC | TMEM163 | 6.94E-34 | -0.36061 | 0.612 | 0.881 |
| DOT1L | NTC | UBASH3B | 1.51E-33 | -0.3288226 | 0.413 | 0.772 |
| DOT1L | NTC | BCL11A | 5.11E-32 | -0.3122837 | 0.366 | 0.741 |
| DOT1L | NTC | PITPNC1 | 2.60E-31 | -0.3611795 | 0.674 | 0.931 |
| DOT1L | NTC | SMC4 | 5.62E-31 | -0.3839084 | 0.848 | 0.964 |
| DOT1L | NTC | SLC23A2 | 6.90E-31 | -0.2695184 | 0.516 | 0.865 |
| DOT1L | NTC | MS4A3 | 2.52E-30 | 0.40756855 | 0.777 | 0.678 |
| DOT1L | NTC | TNS3 | 3.07E-30 | -0.2601461 | 0.406 | 0.762 |
| DOT1L | NTC | MT.CYB | 3.13E-30 | -0.3256127 | 0.987 | 0.997 |
| DOT1L | NTC | NDUFA8 | 6.71E-30 | -0.2889045 | 0.643 | 0.926 |
| DOT1L | NTC | S100P | 3.09E-29 | 0.35678686 | 0.973 | 0.964 |
| DOT1L | NTC | CHI3L1 | 3.23E-29 | 0.30997559 | 0.469 | 0.186 |
| DOT1L | NTC | STXBP5L | 2.05E-28 | -0.2599552 | 0.393 | 0.729 |
| DOT1L | NTC | CITED2 | 1.53E-27 | 0.39234964 | 0.967 | 0.954 |
| DOT1L | NTC | NAV2 | 4.28E-27 | -0.2912552 | 0.384 | 0.713 |
| DOT1L | NTC | CYBB | 4.89E-26 | 0.31363826 | 0.346 | 0.097 |
| DOT1L | NTC | CMIP | 5.84E-26 | -0.2506227 | 0.547 | 0.842 |
| DOT1L | NTC | SFMBT2 | 2.10E-25 | -0.3816099 | 0.759 | 0.936 |
| DOT1L | NTC | PVT1 | 2.96E-25 | -0.2666614 | 0.507 | 0.809 |
| DOT1L | NTC | SATB1 | 3.86E-25 | -0.2891611 | 0.538 | 0.828 |
| DOT1L | NTC | RETN | 8.44E-25 | 0.56514221 | 0.888 | 0.809 |
| DOT1L | NTC | HSPE1 | 3.98E-24 | -0.2921101 | 0.96 | 0.992 |
| DOT1L | NTC | APOLD1 | 6.62E-24 | -0.2961865 | 0.714 | 0.929 |
| DOT1L | NTC | RNASE2 | 6.66E-24 | 0.3907279 | 0.971 | 0.975 |
| DOT1L | NTC | ARHGDIB | 2.93E-23 | 0.25527784 | 0.996 | 0.99 |
| DOT1L | NTC | TGFBR1 | 3.08E-23 | -0.302697 | 0.81 | 0.952 |
| DOT1L | NTC | FBN1 | 3.64E-23 | 0.35500148 | 0.576 | 0.358 |
| DOT1L | NTC | TLN2 | 1.10E-22 | -0.2607603 | 0.498 | 0.794 |
| DOT1L | NTC | FKBP5 | 4.09E-22 | -0.3019393 | 0.953 | 0.98 |
| DOT1L | NTC | AL355922.1 | 9.18E-22 | 0.44934677 | 0.819 | 0.772 |
| DOT1L | NTC | SSB | 1.61E-21 | -0.2727246 | 0.938 | 0.98 |
| DOT1L | NTC | NRXN3 | 2.62E-21 | -0.2620787 | 0.078 | 0.323 |
| DOT1L | NTC | AL138720.1 | 3.10E-21 | -0.3060947 | 0.589 | 0.845 |
| DOT1L | NTC | PPARD | 3.47E-21 | -0.2582261 | 0.65 | 0.891 |
| DOT1L | NTC | BNC2 | 4.24E-21 | -0.292204 | 0.596 | 0.837 |
| DOT1L | NTC | FABP5 | 4.37E-21 | -0.3079718 | 0.902 | 0.983 |

| | | | | | | |
|---|---|---|---|---|---|---|
| DOT1L | NTC | CDA | 8.30E-21 | 0.27851222 | 0.949 | 0.919 |
| DOT1L | NTC | PRTN3 | 9.84E-21 | 0.57693414 | 0.893 | 0.891 |
| DOT1L | NTC | MGST1 | 1.88E-20 | 0.26054068 | 0.969 | 0.98 |
| DOT1L | NTC | RGL4 | 4.23E-20 | 0.33833739 | 0.732 | 0.602 |
| DOT1L | NTC | COL23A1 | 5.43E-20 | -0.3043106 | 0.877 | 0.969 |
| DOT1L | NTC | CHST11 | 6.95E-20 | -0.2791026 | 0.79 | 0.929 |
| DOT1L | NTC | NPC1 | 1.40E-19 | -0.2739569 | 0.516 | 0.774 |
| DOT1L | NTC | GCA | 1.41E-19 | 0.34510659 | 0.583 | 0.416 |
| DOT1L | NTC | EMB | 3.80E-19 | 0.31309512 | 0.864 | 0.865 |
| DOT1L | NTC | CAB39L | 5.55E-19 | -0.2732914 | 0.19 | 0.455 |
| DOT1L | NTC | PDZD8 | 1.41E-18 | -0.2902613 | 0.897 | 0.975 |
| DOT1L | NTC | NSMAF | 1.54E-18 | -0.2604923 | 0.792 | 0.959 |
| DOT1L | NTC | IPCEF1 | 1.58E-18 | 0.34594294 | 0.674 | 0.51 |
| DOT1L | NTC | ARPC5 | 2.55E-17 | 0.25086925 | 0.987 | 0.987 |
| DOT1L | NTC | ELANE | 6.24E-17 | 0.44125389 | 0.978 | 0.992 |
| DOT1L | NTC | ABCA13 | 1.59E-16 | 0.28818214 | 0.545 | 0.371 |
| DOT1L | NTC | ANXA1 | 1.75E-16 | 0.26458355 | 0.556 | 0.37 |
| DOT1L | NTC | STXBP2 | 4.31E-16 | 0.3633014 | 0.922 | 0.926 |
| DOT1L | NTC | HSPA8 | 1.72E-15 | -0.2514184 | 0.951 | 0.99 |
| DOT1L | NTC | PNP | 2.67E-15 | 0.28794053 | 0.902 | 0.932 |
| DOT1L | NTC | CTSD | 6.32E-15 | 0.25757231 | 0.998 | 0.998 |
| DOT1L | NTC | SNHG5 | 2.09E-14 | 0.27583349 | 0.982 | 0.987 |
| DOT1L | NTC | ROMO1 | 2.68E-14 | 0.27072807 | 0.924 | 0.934 |
| DOT1L | NTC | VIM | 3.31E-14 | 0.38441169 | 0.719 | 0.624 |
| DOT1L | NTC | LCP1 | 1.84E-13 | 0.25356449 | 0.942 | 0.954 |
| DOT1L | NTC | SDK1 | 1.03E-12 | -0.2580348 | 0.924 | 0.96 |
| DOT1L | NTC | NEGR1 | 1.65E-11 | -0.29949 | 0.603 | 0.771 |
| DOT1L | NTC | CSTA | 2.29E-10 | 0.27584334 | 0.558 | 0.45 |
| DOT1L | NTC | TENT5A | 1.26E-09 | 0.28709154 | 0.884 | 0.863 |
| DOT1L | NTC | ALAS1 | 2.17E-09 | 0.36266671 | 0.683 | 0.713 |
| DOT1L | NTC | S100A9 | 2.58E-08 | 0.50252636 | 0.455 | 0.351 |
| BRD4 | NTC | TMSB4X | 6.54E-67 | 0.50504037 | 1 | 1 |
| BRD4 | NTC | RPL36 | 5.59E-49 | -0.271319 | 1 | 1 |
| BRD4 | NTC | RPS17 | 8.03E-47 | -0.262639 | 0.998 | 1 |
| BRD4 | NTC | PLEK | 1.34E-44 | 0.42634974 | 0.998 | 0.99 |
| BRD4 | NTC | NEAT1 | 2.71E-43 | 0.8120989 | 0.987 | 0.972 |
| BRD4 | NTC | RPS5 | 4.22E-40 | -0.2691129 | 1 | 1 |
| BRD4 | NTC | MIF | 4.46E-40 | -0.2531973 | 1 | 0.995 |
| BRD4 | NTC | CTSD | 2.13E-34 | 0.46182877 | 1 | 0.998 |
| BRD4 | NTC | GAL | 4.08E-31 | -0.3457928 | 0.769 | 0.931 |
| BRD4 | NTC | P4HB | 7.20E-30 | 0.26704329 | 1 | 0.997 |
| BRD4 | NTC | LGALS1 | 7.53E-28 | 0.33538974 | 1 | 0.998 |
| BRD4 | NTC | FBL | 7.74E-26 | -0.2725992 | 0.916 | 0.992 |
| BRD4 | NTC | IFT57 | 3.49E-25 | -0.3208542 | 0.591 | 0.814 |
| BRD4 | NTC | TRAPPC6A | 5.45E-25 | 0.2773506 | 0.86 | 0.736 |

| | | | | | | |
|---|---|---|---|---|---|---|
| BRD4 | NTC | MALAT1 | 1.27E-23 | 0.48880781 | 0.977 | 0.969 |
| BRD4 | NTC | ATP8B4 | 1.44E-22 | 0.3821421 | 0.995 | 0.985 |
| BRD4 | NTC | CHSY1 | 5.50E-22 | 0.27021839 | 0.93 | 0.929 |
| BRD4 | NTC | NPC2 | 1.04E-21 | 0.25267498 | 0.952 | 0.929 |
| BRD4 | NTC | EBNA1BP2 | 1.47E-21 | -0.2987967 | 0.819 | 0.926 |
| BRD4 | NTC | PIM1 | 7.27E-21 | 0.32860628 | 0.86 | 0.797 |
| BRD4 | NTC | CLEC2A | 1.47E-20 | 0.32488536 | 0.815 | 0.726 |
| BRD4 | NTC | GSTP1 | 8.54E-20 | 0.36446989 | 1 | 1 |
| BRD4 | NTC | NEGR1 | 1.91E-19 | -0.3345548 | 0.529 | 0.771 |
| BRD4 | NTC | DMXL2 | 3.46E-19 | 0.28638357 | 0.978 | 0.964 |
| BRD4 | NTC | MEIS1 | 2.96E-18 | 0.33345818 | 0.964 | 0.952 |
| BRD4 | NTC | FCER1G | 6.59E-18 | 0.35460912 | 0.919 | 0.894 |
| BRD4 | NTC | LRMDA | 1.37E-17 | 0.28179795 | 0.995 | 0.982 |
| BRD4 | NTC | PPP3R1 | 8.21E-17 | 0.30612596 | 0.98 | 0.967 |
| BRD4 | NTC | LYAR | 9.27E-16 | -0.2518995 | 0.803 | 0.926 |
| BRD4 | NTC | MYC | 4.08E-15 | -0.2722314 | 0.616 | 0.795 |
| BRD4 | NTC | AL034397.3 | 6.10E-14 | 0.25927112 | 0.738 | 0.662 |
| BRD4 | NTC | MYO1G | 1.14E-13 | 0.27271751 | 0.871 | 0.835 |
| BRD4 | NTC | FTL | 1.93E-12 | 0.34709163 | 1 | 1 |
| BRD4 | NTC | TMTC2 | 9.12E-10 | 0.25141155 | 0.604 | 0.498 |
| BRD4 | NTC | FTH1 | 0.00512109 | 0.29895641 | 0.987 | 0.987 |
| KAT7 | NTC | LYZ | 1.75E-40 | 0.56994556 | 0.961 | 0.926 |
| KAT7 | NTC | TMSB4X | 1.09E-35 | 0.29193388 | 1 | 1 |
| KAT7 | NTC | GLO1 | 1.82E-31 | -0.3662492 | 1 | 1 |
| KAT7 | NTC | SH3BGRL3 | 5.72E-28 | 0.27549162 | 0.998 | 0.997 |
| KAT7 | NTC | S100A4 | 1.53E-27 | 0.35074275 | 0.998 | 1 |
| KAT7 | NTC | CLEC2B | 9.12E-22 | -0.3640012 | 0.963 | 0.985 |
| KAT7 | NTC | B2M | 7.52E-21 | 0.2557785 | 0.998 | 0.998 |
| KAT7 | NTC | FCER1G | 4.18E-20 | 0.36365856 | 0.952 | 0.894 |
| KAT7 | NTC | LGALS1 | 2.90E-19 | 0.33108007 | 0.998 | 0.998 |
| KAT7 | NTC | S100A9 | 1.57E-17 | 0.47329104 | 0.543 | 0.351 |
| KAT7 | NTC | RNF220 | 3.01E-17 | -0.2701424 | 0.978 | 0.987 |
| KAT7 | NTC | RNASE2 | 6.78E-17 | 0.31552751 | 0.987 | 0.975 |
| KAT7 | NTC | HCST | 3.65E-14 | 0.32269575 | 0.98 | 0.988 |
| KAT7 | NTC | AL355922.1 | 2.03E-13 | 0.29943233 | 0.824 | 0.772 |
| KAT7 | NTC | STXBP2 | 4.63E-12 | 0.29689732 | 0.943 | 0.926 |
| KAT7 | NTC | RNASE3 | 3.58E-11 | 0.25753959 | 0.919 | 0.883 |
| KAT7 | NTC | RETN | 1.64E-08 | 0.28537069 | 0.857 | 0.809 |
| KAT7 | NTC | VIM | 1.65E-07 | 0.28115617 | 0.67 | 0.624 |
| CSNK2A1 | NTC | GLO1 | 7.93E-32 | -0.3921645 | 1 | 1 |
| CSNK2A1 | NTC | CD36 | 3.68E-22 | -0.4136267 | 0.792 | 0.914 |
| CSNK2A1 | NTC | AL138720.1 | 3.32E-21 | -0.2919672 | 0.631 | 0.845 |
| CSNK2A1 | NTC | CCDC26 | 4.47E-20 | -0.4631248 | 0.671 | 0.847 |
| SGF29 | NTC | S100A4 | 7.30E-72 | 0.75951802 | 1 | 1 |
| SGF29 | NTC | GLO1 | 1.90E-55 | -0.6316105 | 0.991 | 1 |

| | | | | | | |
|---|---|---|---|---|---|---|
| SGF29 | NTC | SH3BGRL3 | 3.95E-54 | 0.51597864 | 1 | 0.997 |
| SGF29 | NTC | RPS2 | 1.02E-51 | -0.3382243 | 1 | 1 |
| SGF29 | NTC | LGALS1 | 5.76E-50 | 0.62165748 | 1 | 0.998 |
| SGF29 | NTC | ACTB | 1.46E-45 | 0.30982195 | 1 | 1 |
| SGF29 | NTC | SF3B6 | 1.57E-44 | -0.4338149 | 0.971 | 0.997 |
| SGF29 | NTC | S100A6 | 3.20E-43 | 0.46563794 | 1 | 0.998 |
| SGF29 | NTC | LYZ | 4.00E-40 | 0.70735831 | 0.968 | 0.926 |
| SGF29 | NTC | HCST | 1.25E-38 | 0.59420663 | 0.997 | 0.988 |
| SGF29 | NTC | VIM | 1.82E-38 | 0.7026344 | 0.832 | 0.624 |
| SGF29 | NTC | ATP5F1E | 4.10E-37 | 0.35856101 | 1 | 0.993 |
| SGF29 | NTC | SRGN | 5.12E-35 | 0.47886347 | 1 | 0.997 |
| SGF29 | NTC | FABP5 | 1.47E-34 | -0.4109485 | 0.838 | 0.983 |
| SGF29 | NTC | CD52 | 2.96E-34 | 0.49720681 | 0.365 | 0.068 |
| SGF29 | NTC | B2M | 1.47E-33 | 0.40717501 | 1 | 0.998 |
| SGF29 | NTC | FCER1G | 4.95E-33 | 0.56634486 | 0.962 | 0.894 |
| SGF29 | NTC | ARPC1B | 8.82E-33 | 0.35984095 | 0.997 | 0.997 |
| SGF29 | NTC | ACTR2 | 8.88E-32 | 0.36740298 | 1 | 0.997 |
| SGF29 | NTC | CLEC2B | 2.36E-31 | -0.5307239 | 0.945 | 0.985 |
| SGF29 | NTC | MYL6 | 1.74E-30 | 0.34274406 | 1 | 0.993 |
| SGF29 | NTC | PEBP1 | 1.88E-30 | -0.2896833 | 0.899 | 0.987 |
| SGF29 | NTC | S100P | 3.09E-30 | 0.49997635 | 0.968 | 0.964 |
| SGF29 | NTC | PLP2 | 9.80E-30 | 0.38113646 | 0.965 | 0.954 |
| SGF29 | NTC | TYROBP | 1.66E-29 | 0.69718772 | 0.559 | 0.264 |
| SGF29 | NTC | NEGR1 | 2.07E-29 | -0.4999304 | 0.417 | 0.771 |
| SGF29 | NTC | LINC02470 | 2.79E-29 | -0.3629359 | 0.658 | 0.929 |
| SGF29 | NTC | NME1 | 3.07E-29 | -0.3704478 | 0.858 | 0.974 |
| SGF29 | NTC | VAMP8 | 6.85E-29 | 0.28110626 | 1 | 0.997 |
| SGF29 | NTC | CAP1 | 9.12E-29 | 0.37119393 | 0.991 | 0.982 |
| SGF29 | NTC | HSPE1 | 1.52E-28 | -0.3637829 | 0.968 | 0.992 |
| SGF29 | NTC | NCF1C | 5.92E-28 | 0.26905574 | 0.339 | 0.076 |
| SGF29 | NTC | PA2G4 | 1.04E-27 | -0.3249298 | 0.928 | 0.99 |
| SGF29 | NTC | FBL | 1.18E-27 | -0.3460306 | 0.838 | 0.992 |
| SGF29 | NTC | PTMA | 3.29E-27 | -0.2526418 | 1 | 1 |
| SGF29 | NTC | LSP1 | 4.40E-27 | 0.3817847 | 0.858 | 0.748 |
| SGF29 | NTC | HSP90AB1 | 8.29E-27 | -0.2910135 | 1 | 0.998 |
| SGF29 | NTC | NPM1 | 8.44E-27 | -0.3456355 | 0.991 | 0.993 |
| SGF29 | NTC | CCND3 | 1.10E-26 | 0.38018931 | 0.971 | 0.969 |
| SGF29 | NTC | HS3ST2 | 1.87E-26 | -0.3698854 | 0.62 | 0.911 |
| SGF29 | NTC | METTL9 | 7.34E-26 | 0.49948981 | 0.89 | 0.861 |
| SGF29 | NTC | SERBP1 | 2.96E-25 | -0.2585673 | 0.997 | 1 |
| SGF29 | NTC | RPL7A | 1.96E-24 | -0.2567286 | 1 | 0.998 |
| SGF29 | NTC | MYO1G | 2.33E-24 | 0.39401312 | 0.87 | 0.835 |
| SGF29 | NTC | LCP1 | 3.55E-24 | 0.35849253 | 0.974 | 0.954 |
| SGF29 | NTC | PYGL | 3.61E-24 | -0.2883804 | 0.745 | 0.941 |
| SGF29 | NTC | TMSB4X | 4.44E-24 | 0.27576899 | 1 | 1 |

| | | | | | | |
|---|---|---|---|---|---|---|
| SGF29 | NTC | C1QBP | 4.90E-24 | -0.2732128 | 0.872 | 0.982 |
| SGF29 | NTC | GRN | 5.81E-24 | 0.3568508 | 0.872 | 0.84 |
| SGF29 | NTC | ODC1 | 6.44E-24 | -0.2868178 | 0.774 | 0.949 |
| SGF29 | NTC | ROMO1 | 1.45E-23 | 0.40817693 | 0.951 | 0.934 |
| SGF29 | NTC | NCL | 1.81E-23 | -0.3348025 | 0.98 | 0.992 |
| SGF29 | NTC | GNAI2 | 2.38E-23 | 0.28857505 | 0.98 | 0.979 |
| SGF29 | NTC | IL2RG | 3.00E-23 | 0.35325617 | 0.881 | 0.807 |
| SGF29 | NTC | EIF5A | 3.21E-23 | -0.2546843 | 0.959 | 0.987 |
| SGF29 | NTC | RETN | 4.15E-23 | 0.62008909 | 0.867 | 0.809 |
| SGF29 | NTC | RNASE2 | 4.15E-23 | 0.37823767 | 0.991 | 0.975 |
| SGF29 | NTC | LYL1 | 4.37E-23 | -0.2597819 | 0.696 | 0.932 |
| SGF29 | NTC | BCL11A | 6.77E-23 | -0.2726033 | 0.4 | 0.741 |
| SGF29 | NTC | PPA1 | 1.01E-22 | -0.2661903 | 0.887 | 0.972 |
| SGF29 | NTC | CDA | 1.18E-22 | 0.34250928 | 0.951 | 0.919 |
| SGF29 | NTC | G6PD | 1.45E-22 | 0.36953024 | 0.774 | 0.691 |
| SGF29 | NTC | ANP32A | 1.51E-22 | -0.262158 | 0.957 | 0.99 |
| SGF29 | NTC | LAMTOR4 | 1.54E-22 | 0.34450169 | 0.983 | 0.98 |
| SGF29 | NTC | HSPD1 | 2.05E-22 | -0.2857169 | 0.977 | 0.995 |
| SGF29 | NTC | CTNNAL1 | 2.14E-22 | -0.2825732 | 0.461 | 0.812 |
| SGF29 | NTC | CMTM6 | 2.36E-22 | 0.3657273 | 0.928 | 0.921 |
| SGF29 | NTC | MCEMP1 | 2.60E-22 | 0.37537666 | 0.771 | 0.662 |
| SGF29 | NTC | DST | 4.29E-22 | -0.2598387 | 0.336 | 0.686 |
| SGF29 | NTC | SLC44A1 | 5.36E-22 | 0.33008075 | 0.852 | 0.781 |
| SGF29 | NTC | IFT57 | 5.80E-22 | -0.3310036 | 0.525 | 0.814 |
| SGF29 | NTC | APOC2 | 6.43E-22 | 0.43747723 | 0.913 | 0.913 |
| SGF29 | NTC | ALOX5AP | 7.63E-22 | 0.38336875 | 0.725 | 0.571 |
| SGF29 | NTC | GAL | 8.37E-22 | -0.3174369 | 0.696 | 0.931 |
| SGF29 | NTC | CSTA | 2.04E-21 | 0.47240406 | 0.652 | 0.45 |
| SGF29 | NTC | EBNA1BP2 | 4.59E-21 | -0.3401834 | 0.719 | 0.926 |
| SGF29 | NTC | PRMT1 | 5.11E-21 | -0.2812461 | 0.751 | 0.936 |
| SGF29 | NTC | BMI1 | 6.05E-21 | -0.3152224 | 0.791 | 0.95 |
| SGF29 | NTC | PGD | 7.50E-21 | 0.33333318 | 0.939 | 0.909 |
| SGF29 | NTC | ITGAL | 8.74E-21 | 0.25658121 | 0.441 | 0.196 |
| SGF29 | NTC | CYBA | 1.12E-20 | 0.30793915 | 0.988 | 0.987 |
| SGF29 | NTC | MYL12A | 1.23E-20 | 0.36114511 | 0.925 | 0.932 |
| SGF29 | NTC | STMN1 | 1.80E-20 | 0.25537272 | 1 | 0.997 |
| SGF29 | NTC | ATP8B4 | 1.82E-20 | 0.35185236 | 0.997 | 0.985 |
| SGF29 | NTC | HSPA8 | 1.87E-20 | -0.3223901 | 0.945 | 0.99 |
| SGF29 | NTC | COX7A2 | 5.03E-20 | 0.29682059 | 0.994 | 0.988 |
| SGF29 | NTC | GMFG | 5.12E-20 | 0.39595026 | 0.965 | 0.974 |
| SGF29 | NTC | LST1 | 8.30E-20 | 0.38654713 | 0.962 | 0.96 |
| SGF29 | NTC | PFDN5 | 1.09E-19 | 0.29874002 | 0.991 | 0.992 |
| SGF29 | NTC | PTPRG | 1.13E-19 | -0.2515281 | 0.446 | 0.756 |
| SGF29 | NTC | EIF4EBP1 | 2.88E-19 | -0.2716219 | 0.936 | 0.993 |
| SGF29 | NTC | OLA1 | 3.97E-19 | -0.2522943 | 0.962 | 0.987 |

| | | | | | | |
|---|---|---|---|---|---|---|
| SGF29 | NTC | TIMP1 | 5.64E-19 | 0.39178731 | 0.916 | 0.937 |
| SGF29 | NTC | SRPK1 | 6.02E-19 | -0.2619503 | 0.983 | 0.992 |
| SGF29 | NTC | MYC | 8.12E-19 | -0.3312172 | 0.51 | 0.795 |
| SGF29 | NTC | ANXA1 | 8.47E-19 | 0.38004096 | 0.577 | 0.37 |
| SGF29 | NTC | DEF6 | 2.10E-18 | 0.33973757 | 0.925 | 0.944 |
| SGF29 | NTC | STXBP2 | 2.15E-18 | 0.39121575 | 0.939 | 0.926 |
| SGF29 | NTC | FTH1 | 7.45E-18 | 0.33310337 | 0.986 | 0.987 |
| SGF29 | NTC | PRDX1 | 9.17E-18 | -0.2506435 | 0.928 | 0.99 |
| SGF29 | NTC | ITGB2 | 1.63E-17 | 0.3211224 | 0.803 | 0.752 |
| SGF29 | NTC | NPC2 | 3.04E-17 | 0.30842781 | 0.922 | 0.929 |
| SGF29 | NTC | AIF1 | 3.17E-17 | 0.35787116 | 0.977 | 0.982 |
| SGF29 | NTC | LAPTM5 | 4.46E-17 | 0.26290851 | 0.954 | 0.954 |
| SGF29 | NTC | CAPG | 4.59E-17 | 0.29701985 | 0.997 | 0.997 |
| SGF29 | NTC | PDCD5 | 5.27E-17 | -0.2545102 | 0.916 | 0.983 |
| SGF29 | NTC | ARPC2 | 5.69E-17 | 0.26727095 | 0.994 | 0.99 |
| SGF29 | NTC | PTK2B | 7.29E-17 | 0.29418824 | 0.893 | 0.85 |
| SGF29 | NTC | NUDC | 1.93E-16 | -0.2523438 | 0.872 | 0.97 |
| SGF29 | NTC | CHMP2A | 1.97E-16 | 0.34785397 | 0.904 | 0.922 |
| SGF29 | NTC | CCL2 | 2.36E-16 | 0.46261458 | 0.452 | 0.238 |
| SGF29 | NTC | SPTLC2 | 3.87E-16 | 0.33302072 | 0.814 | 0.787 |
| SGF29 | NTC | DMXL2 | 4.97E-16 | 0.2977638 | 0.98 | 0.964 |
| SGF29 | NTC | SLPI | 5.39E-16 | 0.41772221 | 0.519 | 0.32 |
| SGF29 | NTC | ARPC5 | 5.42E-16 | 0.2870543 | 0.994 | 0.987 |
| SGF29 | NTC | RCSD1 | 1.09E-15 | 0.26101691 | 0.864 | 0.863 |
| SGF29 | NTC | TPM4 | 1.20E-15 | 0.33152048 | 0.936 | 0.931 |
| SGF29 | NTC | IFI27L2 | 1.37E-15 | 0.32678677 | 0.817 | 0.764 |
| SGF29 | NTC | SDK1 | 1.40E-15 | -0.325591 | 0.89 | 0.96 |
| SGF29 | NTC | SNX10 | 1.45E-15 | 0.3818313 | 0.67 | 0.554 |
| SGF29 | NTC | NPC1 | 1.78E-15 | -0.251629 | 0.51 | 0.774 |
| SGF29 | NTC | CRIP1 | 2.18E-15 | 0.30314356 | 0.977 | 0.982 |
| SGF29 | NTC | NDRG1 | 2.75E-15 | 0.25757548 | 0.748 | 0.62 |
| SGF29 | NTC | CD36 | 2.78E-15 | -0.3903827 | 0.788 | 0.914 |
| SGF29 | NTC | SKAP2 | 4.87E-15 | 0.26936356 | 0.948 | 0.931 |
| SGF29 | NTC | MRPL33 | 7.08E-15 | 0.29551541 | 0.922 | 0.955 |
| SGF29 | NTC | NRG4 | 1.59E-14 | 0.30853127 | 0.487 | 0.309 |
| SGF29 | NTC | S100A10 | 1.90E-14 | 0.25458694 | 0.89 | 0.848 |
| SGF29 | NTC | BLOC1S1 | 2.20E-14 | 0.29736818 | 0.887 | 0.888 |
| SGF29 | NTC | CAT | 2.27E-14 | 0.27477403 | 0.896 | 0.883 |
| SGF29 | NTC | HLA.C | 3.99E-14 | 0.2642193 | 0.939 | 0.962 |
| SGF29 | NTC | CD63 | 4.18E-14 | 0.30369767 | 0.974 | 0.98 |
| SGF29 | NTC | PECAM1 | 5.58E-14 | 0.26207181 | 0.614 | 0.485 |
| SGF29 | NTC | SMC4 | 6.96E-14 | -0.2665561 | 0.878 | 0.964 |
| SGF29 | NTC | NOP56 | 1.13E-13 | -0.2501781 | 0.872 | 0.975 |
| SGF29 | NTC | RNASE3 | 5.46E-13 | 0.30769598 | 0.887 | 0.883 |
| SGF29 | NTC | RAC2 | 6.21E-13 | 0.26309294 | 0.904 | 0.944 |

| | | | | | | |
|---|---|---|---|---|---|---|
| SGF29 | NTC | CD93 | 1.39E-12 | 0.27249949 | 0.983 | 0.97 |
| SGF29 | NTC | TMX4 | 2.11E-12 | 0.32253227 | 0.78 | 0.766 |
| SGF29 | NTC | AC007100.1 | 2.60E-12 | -0.2673516 | 0.484 | 0.726 |
| SGF29 | NTC | SH3PXD2A | 2.95E-12 | 0.29558548 | 0.803 | 0.731 |
| SGF29 | NTC | EMP3 | 4.33E-12 | 0.26063186 | 0.991 | 0.993 |
| SGF29 | NTC | NSUN5 | 7.07E-12 | 0.27894604 | 0.89 | 0.913 |
| SGF29 | NTC | BEX1 | 1.37E-11 | -0.2858745 | 0.901 | 0.965 |
| SGF29 | NTC | EPDR1 | 1.60E-11 | 0.31399058 | 0.751 | 0.772 |
| SGF29 | NTC | ITGA6 | 3.17E-11 | 0.30194028 | 0.896 | 0.911 |
| SGF29 | NTC | CTSS | 3.26E-11 | 0.27303259 | 0.678 | 0.591 |
| SGF29 | NTC | PTPRC | 4.56E-11 | 0.27274505 | 0.82 | 0.764 |
| SGF29 | NTC | AZU1 | 5.87E-11 | 0.26543237 | 0.974 | 0.979 |
| SGF29 | NTC | TPSAB1 | 1.19E-10 | 0.25682155 | 0.571 | 0.469 |
| SGF29 | NTC | SYF2 | 1.82E-10 | 0.25062244 | 0.884 | 0.889 |
| SGF29 | NTC | FAM111A | 2.95E-10 | 0.26710941 | 0.693 | 0.637 |
| SGF29 | NTC | COMMD6 | 3.96E-10 | 0.28406104 | 0.925 | 0.95 |
| SGF29 | NTC | PTPRE | 6.01E-10 | 0.25705829 | 0.67 | 0.609 |
| SGF29 | NTC | LMNA | 6.71E-10 | 0.3025726 | 0.881 | 0.899 |
| SGF29 | NTC | ANXA2 | 8.96E-10 | 0.35166198 | 0.794 | 0.814 |
| SGF29 | NTC | IMMP2L | 1.13E-09 | -0.254573 | 0.635 | 0.82 |
| SGF29 | NTC | LGALS3 | 1.94E-09 | 0.30240968 | 0.632 | 0.558 |
| SGF29 | NTC | MT1F | 2.48E-09 | 0.29377867 | 0.82 | 0.797 |
| SGF29 | NTC | CDC42EP3 | 3.03E-08 | 0.26401487 | 0.681 | 0.67 |
| SGF29 | NTC | ACAP1 | 3.10E-07 | 0.28590361 | 0.646 | 0.589 |
| SGF29 | NTC | C12orf57 | 1.29E-05 | 0.29114969 | 0.664 | 0.662 |
| SGF29 | NTC | PTTG1 | 0.00047754 | 0.29888519 | 0.823 | 0.878 |
| SGF29 | NTC | TNNT1 | 0.001292 | 0.28577885 | 0.438 | 0.398 |
| MLLT1 | NTC | LYZ | 9.95E-119 | 1.21021318 | 0.984 | 0.926 |
| MLLT1 | NTC | GLO1 | 4.03E-110 | -0.9302643 | 0.994 | 1 |
| MLLT1 | NTC | CLEC2B | 2.67E-91 | -0.9395064 | 0.86 | 0.985 |
| MLLT1 | NTC | ACTB | 5.29E-86 | 0.44278466 | 1 | 1 |
| MLLT1 | NTC | METTL9 | 1.36E-73 | 0.64202623 | 0.936 | 0.861 |
| MLLT1 | NTC | SH3BGRL3 | 8.15E-73 | 0.50667999 | 0.996 | 0.997 |
| MLLT1 | NTC | B2M | 1.43E-71 | 0.53713105 | 0.994 | 0.998 |
| MLLT1 | NTC | LINC02470 | 2.35E-70 | -0.5450771 | 0.511 | 0.929 |
| MLLT1 | NTC | GSTP1 | 1.49E-66 | -0.4939538 | 0.996 | 1 |
| MLLT1 | NTC | S100A4 | 9.63E-66 | 0.59851866 | 0.998 | 1 |
| MLLT1 | NTC | BMI1 | 3.30E-61 | -0.5515028 | 0.633 | 0.95 |
| MLLT1 | NTC | LGALS1 | 2.13E-59 | 0.60811826 | 0.998 | 0.998 |
| MLLT1 | NTC | FTL | 2.00E-55 | 0.32069668 | 1 | 1 |
| MLLT1 | NTC | INPP4B | 6.43E-54 | -0.4331507 | 0.313 | 0.776 |
| MLLT1 | NTC | TMSB4X | 6.04E-53 | 0.35375901 | 1 | 1 |
| MLLT1 | NTC | MNDA | 8.93E-52 | 0.5386725 | 0.984 | 0.983 |
| MLLT1 | NTC | TLN2 | 2.45E-51 | -0.400484 | 0.322 | 0.794 |
| MLLT1 | NTC | TMEM163 | 3.65E-51 | -0.4293033 | 0.505 | 0.881 |

| | | | | | | |
|---|---|---|---|---|---|---|
| MLLT1 | NTC | S100A9 | 3.09E-49 | 1.139978 | 0.67 | 0.351 |
| MLLT1 | NTC | CD36 | 3.73E-49 | -0.6590593 | 0.649 | 0.914 |
| MLLT1 | NTC | MEIS1 | 1.82E-48 | -0.4791346 | 0.707 | 0.952 |
| MLLT1 | NTC | STXBP5L | 1.09E-47 | -0.2872435 | 0.254 | 0.729 |
| MLLT1 | NTC | FABP5 | 1.19E-45 | -0.4586089 | 0.81 | 0.983 |
| MLLT1 | NTC | VIM | 1.42E-45 | 0.79988418 | 0.788 | 0.624 |
| MLLT1 | NTC | C12orf75 | 2.26E-45 | -0.4548868 | 0.596 | 0.919 |
| MLLT1 | NTC | RETN | 3.57E-45 | 0.82085983 | 0.887 | 0.809 |
| MLLT1 | NTC | RNF220 | 7.38E-44 | -0.4793989 | 0.95 | 0.987 |
| MLLT1 | NTC | CDA | 3.12E-42 | 0.4226452 | 0.936 | 0.919 |
| MLLT1 | NTC | KAZN | 7.14E-41 | -0.263047 | 0.256 | 0.695 |
| MLLT1 | NTC | HS3ST2 | 8.88E-41 | -0.4578348 | 0.575 | 0.911 |
| MLLT1 | NTC | SRGN | 8.89E-41 | 0.44580836 | 0.998 | 0.997 |
| MLLT1 | NTC | STMN1 | 1.53E-40 | 0.34548434 | 0.998 | 0.997 |
| MLLT1 | NTC | CLECL1 | 2.02E-40 | -0.3728772 | 0.701 | 0.946 |
| MLLT1 | NTC | CLEC2A | 2.17E-40 | -0.3148516 | 0.291 | 0.726 |
| MLLT1 | NTC | LSP1 | 3.49E-40 | 0.44998553 | 0.823 | 0.748 |
| MLLT1 | NTC | NEGR1 | 3.96E-40 | -0.5380998 | 0.381 | 0.771 |
| MLLT1 | NTC | STXBP2 | 1.85E-39 | 0.61685625 | 0.917 | 0.926 |
| MLLT1 | NTC | PRDX1 | 5.01E-39 | -0.3532947 | 0.897 | 0.99 |
| MLLT1 | NTC | NPC1 | 2.18E-38 | -0.3529846 | 0.367 | 0.774 |
| MLLT1 | NTC | GAL | 1.61E-37 | -0.3897156 | 0.641 | 0.931 |
| MLLT1 | NTC | ANXA1 | 2.95E-37 | 0.50998552 | 0.639 | 0.37 |
| MLLT1 | NTC | GNAI2 | 1.42E-36 | 0.33975411 | 0.979 | 0.979 |
| MLLT1 | NTC | PDZD8 | 7.43E-36 | -0.4007424 | 0.841 | 0.975 |
| MLLT1 | NTC | NME1 | 6.67E-35 | -0.3604359 | 0.833 | 0.974 |
| MLLT1 | NTC | EPDR1 | 8.29E-35 | 0.46452941 | 0.812 | 0.772 |
| MLLT1 | NTC | S100A8 | 8.62E-35 | 0.78991974 | 0.334 | 0.056 |
| MLLT1 | NTC | LCP1 | 1.61E-34 | 0.39635048 | 0.959 | 0.954 |
| MLLT1 | NTC | MYL6 | 2.17E-34 | 0.35898245 | 0.994 | 0.993 |
| MLLT1 | NTC | NUDC | 2.48E-34 | -0.3348774 | 0.802 | 0.97 |
| MLLT1 | NTC | PGD | 3.17E-34 | 0.41433542 | 0.911 | 0.909 |
| MLLT1 | NTC | S100P | 4.59E-34 | 0.43398986 | 0.955 | 0.964 |
| MLLT1 | NTC | ACTR2 | 1.51E-33 | 0.32963455 | 0.996 | 0.997 |
| MLLT1 | NTC | HSPE1 | 1.72E-33 | -0.348095 | 0.961 | 0.992 |
| MLLT1 | NTC | C1QBP | 1.94E-33 | -0.3017844 | 0.831 | 0.982 |
| MLLT1 | NTC | CFL1 | 3.83E-33 | 0.25503016 | 0.998 | 0.997 |
| MLLT1 | NTC | CAP1 | 2.66E-32 | 0.36370873 | 0.988 | 0.982 |
| MLLT1 | NTC | S100A11 | 2.98E-32 | 0.32014142 | 0.996 | 0.998 |
| MLLT1 | NTC | COL23A1 | 3.21E-32 | -0.3911191 | 0.823 | 0.969 |
| MLLT1 | NTC | G6PD | 3.58E-32 | 0.37451406 | 0.777 | 0.691 |
| MLLT1 | NTC | CCND3 | 4.46E-32 | 0.38417592 | 0.955 | 0.969 |
| MLLT1 | NTC | EBNA1BP2 | 9.58E-32 | -0.3881048 | 0.668 | 0.926 |
| MLLT1 | NTC | PVT1 | 1.48E-31 | -0.3168335 | 0.452 | 0.809 |
| MLLT1 | NTC | NRXN3 | 2.38E-31 | -0.2933199 | 0.045 | 0.323 |

| | | | | | | |
|---|---|---|---|---|---|---|
| MLLT1 | NTC | GRN | 5.72E-31 | 0.38361399 | 0.839 | 0.84 |
| MLLT1 | NTC | RNASE2 | 9.03E-31 | 0.41161269 | 0.979 | 0.975 |
| MLLT1 | NTC | CAB39L | 1.09E-30 | -0.3027112 | 0.134 | 0.455 |
| MLLT1 | NTC | HSP90AB1 | 1.10E-30 | -0.2802149 | 1 | 0.998 |
| MLLT1 | NTC | VAMP8 | 1.40E-30 | 0.25141494 | 0.988 | 0.997 |
| MLLT1 | NTC | ODC1 | 4.10E-30 | -0.2750863 | 0.74 | 0.949 |
| MLLT1 | NTC | SSB | 4.64E-30 | -0.3151037 | 0.893 | 0.98 |
| MLLT1 | NTC | NDUFA8 | 7.66E-30 | -0.2638931 | 0.617 | 0.926 |
| MLLT1 | NTC | CAT | 2.67E-29 | 0.35167231 | 0.897 | 0.883 |
| MLLT1 | NTC | CMTM6 | 2.98E-29 | 0.34516262 | 0.915 | 0.921 |
| MLLT1 | NTC | ZEB2 | 1.60E-28 | -0.4241276 | 0.835 | 0.949 |
| MLLT1 | NTC | ARPC5 | 2.11E-28 | 0.32998289 | 0.99 | 0.987 |
| MLLT1 | NTC | CCT3 | 7.15E-28 | -0.2933855 | 0.87 | 0.985 |
| MLLT1 | NTC | RNASE3 | 1.72E-27 | 0.46310815 | 0.887 | 0.883 |
| MLLT1 | NTC | NPM1 | 2.84E-27 | -0.3044613 | 0.977 | 0.993 |
| MLLT1 | NTC | CAMK1D | 2.88E-27 | -0.2629419 | 0.355 | 0.716 |
| MLLT1 | NTC | MTHFD1L | 3.20E-27 | -0.2968587 | 0.592 | 0.899 |
| MLLT1 | NTC | MYL12A | 7.21E-27 | 0.35656301 | 0.901 | 0.932 |
| MLLT1 | NTC | HSPD1 | 8.40E-27 | -0.293547 | 0.961 | 0.995 |
| MLLT1 | NTC | FHIT | 1.56E-26 | -0.3367849 | 0.414 | 0.738 |
| MLLT1 | NTC | UBASH3B | 1.64E-26 | -0.2596627 | 0.423 | 0.772 |
| MLLT1 | NTC | AC007100.1 | 3.62E-26 | -0.3393363 | 0.384 | 0.726 |
| MLLT1 | NTC | SRM | 5.20E-26 | -0.2754063 | 0.932 | 0.987 |
| MLLT1 | NTC | NSMAF | 1.95E-25 | -0.2974454 | 0.748 | 0.959 |
| MLLT1 | NTC | ANXA6 | 3.17E-25 | 0.31425782 | 0.936 | 0.954 |
| MLLT1 | NTC | PLEK | 4.33E-25 | -0.2676016 | 0.965 | 0.99 |
| MLLT1 | NTC | CEBPE | 5.83E-25 | 0.3341601 | 0.765 | 0.724 |
| MLLT1 | NTC | HSPA8 | 1.58E-24 | -0.3232674 | 0.918 | 0.99 |
| MLLT1 | NTC | ATP5MC1 | 1.97E-24 | -0.2783561 | 0.858 | 0.988 |
| MLLT1 | NTC | CLEC12A | 4.04E-24 | -0.2739682 | 0.674 | 0.896 |
| MLLT1 | NTC | GPATCH4 | 5.47E-24 | -0.2996227 | 0.594 | 0.87 |
| MLLT1 | NTC | POLR2F | 6.86E-24 | -0.2939931 | 0.674 | 0.939 |
| MLLT1 | NTC | MT.CYB | 1.39E-23 | -0.2640038 | 0.99 | 0.997 |
| MLLT1 | NTC | ETS1 | 1.92E-23 | 0.35399462 | 0.759 | 0.719 |
| MLLT1 | NTC | HLA.C | 1.94E-23 | 0.32420892 | 0.92 | 0.962 |
| MLLT1 | NTC | TKT | 2.41E-23 | 0.29454563 | 0.969 | 0.988 |
| MLLT1 | NTC | RGL4 | 6.51E-23 | 0.37926913 | 0.695 | 0.602 |
| MLLT1 | NTC | H2AFZ | 9.76E-23 | 0.2906111 | 0.986 | 0.995 |
| MLLT1 | NTC | LARS | 1.76E-22 | -0.2699605 | 0.728 | 0.947 |
| MLLT1 | NTC | SLPI | 1.85E-22 | 0.52068345 | 0.522 | 0.32 |
| MLLT1 | NTC | ARPC1B | 1.92E-22 | 0.2769347 | 0.986 | 0.997 |
| MLLT1 | NTC | S100A6 | 2.05E-22 | 0.29406132 | 0.996 | 0.998 |
| MLLT1 | NTC | CDC42EP3 | 1.41E-21 | 0.3484371 | 0.728 | 0.67 |
| MLLT1 | NTC | IMMP2L | 1.56E-21 | -0.3556886 | 0.559 | 0.82 |
| MLLT1 | NTC | PRMT1 | 1.96E-21 | -0.255814 | 0.722 | 0.936 |

| | | | | | | |
|---|---|---|---|---|---|---|
| MLLT1 | NTC | CSTA | 2.39E-21 | 0.45121252 | 0.612 | 0.45 |
| MLLT1 | NTC | PLP2 | 6.67E-21 | 0.30087751 | 0.944 | 0.954 |
| MLLT1 | NTC | SNX10 | 1.35E-20 | 0.40282364 | 0.656 | 0.554 |
| MLLT1 | NTC | PRDX4 | 2.05E-20 | -0.2689086 | 0.798 | 0.949 |
| MLLT1 | NTC | LYAR | 3.24E-20 | -0.2874919 | 0.689 | 0.926 |
| MLLT1 | NTC | HCST | 3.77E-20 | 0.43998166 | 0.961 | 0.988 |
| MLLT1 | NTC | CAPG | 8.21E-20 | 0.27483812 | 0.994 | 0.997 |
| MLLT1 | NTC | RCSD1 | 1.19E-19 | 0.27304996 | 0.829 | 0.863 |
| MLLT1 | NTC | FAM107B | 1.80E-19 | 0.29307575 | 0.932 | 0.946 |
| MLLT1 | NTC | SDK1 | 3.78E-19 | -0.3010584 | 0.878 | 0.96 |
| MLLT1 | NTC | SLC44A1 | 2.40E-18 | 0.31742961 | 0.773 | 0.781 |
| MLLT1 | NTC | MYL12B | 3.94E-18 | 0.26342447 | 0.94 | 0.977 |
| MLLT1 | NTC | SMC4 | 9.32E-18 | -0.2668636 | 0.817 | 0.964 |
| MLLT1 | NTC | PYCARD | 2.24E-17 | 0.26236583 | 0.924 | 0.964 |
| MLLT1 | NTC | MYC | 8.15E-17 | -0.2664451 | 0.542 | 0.795 |
| MLLT1 | NTC | ROMO1 | 3.54E-16 | 0.34372333 | 0.899 | 0.934 |
| MLLT1 | NTC | PSTPIP1 | 2.69E-14 | 0.3760776 | 0.695 | 0.69 |
| MLLT1 | NTC | FTH1 | 3.42E-14 | 0.37727212 | 0.969 | 0.987 |
| MLLT1 | NTC | PRDX5 | 8.36E-14 | 0.32634347 | 0.928 | 0.975 |
| MLLT1 | NTC | AL355922.1 | 1.83E-13 | 0.35408272 | 0.742 | 0.772 |
| MLLT1 | NTC | TYROBP | 1.85E-13 | 0.42267609 | 0.419 | 0.264 |
| MLLT1 | NTC | FCER1G | 2.94E-13 | 0.34983079 | 0.887 | 0.894 |
| MLLT1 | NTC | IPCEF1 | 4.47E-13 | 0.32616767 | 0.61 | 0.51 |
| MLLT1 | NTC | CALM1 | 5.67E-13 | 0.26476341 | 0.963 | 0.98 |
| MLLT1 | NTC | ALOX5AP | 8.80E-13 | 0.32496671 | 0.616 | 0.571 |
| MLLT1 | NTC | NSUN5 | 1.01E-12 | 0.26038925 | 0.87 | 0.913 |
| MLLT1 | NTC | FAM111A | 1.92E-12 | 0.29448217 | 0.662 | 0.637 |
| MLLT1 | NTC | IFI16 | 2.24E-12 | 0.30147451 | 0.598 | 0.513 |
| MLLT1 | NTC | GLRX | 1.22E-11 | 0.26870042 | 0.699 | 0.718 |
| MLLT1 | NTC | WWOX | 1.61E-11 | -0.2725994 | 0.827 | 0.921 |
| MLLT1 | NTC | ANKRD44 | 1.62E-11 | 0.27793435 | 0.843 | 0.881 |
| MLLT1 | NTC | GMFG | 3.84E-11 | 0.26190342 | 0.92 | 0.974 |
| MLLT1 | NTC | PLAC8 | 9.41E-11 | 0.2703052 | 0.412 | 0.287 |
| MLLT1 | NTC | MRPL33 | 1.68E-10 | 0.29643784 | 0.866 | 0.955 |
| MLLT1 | NTC | RALGAPA2 | 5.12E-10 | 0.28715527 | 0.685 | 0.693 |
| MLLT1 | NTC | ANXA2 | 1.12E-09 | 0.33633131 | 0.759 | 0.814 |
| MLLT1 | NTC | CTSS | 2.62E-09 | 0.29218529 | 0.61 | 0.591 |
| MLLT1 | NTC | AFF2 | 2.35E-08 | 0.25411061 | 0.911 | 0.962 |
| MLLT1 | NTC | S100A10 | 2.35E-08 | 0.25039173 | 0.767 | 0.848 |
| MLLT1 | NTC | MGAT4A | 3.36E-08 | 0.25104418 | 0.557 | 0.512 |
| MLLT1 | NTC | IFI30 | 7.54E-07 | 0.31479366 | 0.693 | 0.723 |
| MLLT6 | NTC | CLEC2B | 1.77E-66 | -0.7462976 | 0.949 | 0.985 |
| MLLT6 | NTC | PLEK | 1.92E-25 | -0.2617133 | 0.998 | 0.99 |
| MLLT6 | NTC | LINC02470 | 2.74E-22 | -0.2716585 | 0.802 | 0.929 |
| MLLT6 | NTC | GLO1 | 3.41E-22 | -0.2987578 | 1 | 1 |

| | | | | | | |
|---|---|---|---|---|---|---|
| MLLT6 | NTC | CLEC2A | 1.34E-20 | -0.2512442 | 0.492 | 0.726 |
| MLLT6 | NTC | CD36 | 1.66E-09 | -0.2742475 | 0.864 | 0.914 |
| MLLT10 | NTC | CLEC2B | 1.14E-39 | -0.5303373 | 0.832 | 0.985 |
| MLLT10 | NTC | RNF220 | 1.77E-30 | -0.409829 | 0.91 | 0.987 |
| MLLT10 | NTC | GLO1 | 2.86E-30 | -0.3592327 | 0.991 | 1 |
| MLLT10 | NTC | SRGN | 2.77E-25 | 0.42051563 | 1 | 0.997 |
| MLLT10 | NTC | MEIS1 | 9.70E-23 | -0.2880451 | 0.678 | 0.952 |
| MLLT10 | NTC | HS3ST2 | 1.38E-20 | -0.3197189 | 0.725 | 0.911 |
| MLLT10 | NTC | PRTN3 | 1.92E-20 | 0.69016379 | 0.928 | 0.891 |
| MLLT10 | NTC | INPP4B | 3.82E-19 | -0.2511287 | 0.529 | 0.776 |
| MLLT10 | NTC | RNASE3 | 2.92E-18 | 0.40546153 | 0.942 | 0.883 |
| MLLT10 | NTC | FAM107B | 7.82E-18 | 0.37130957 | 0.968 | 0.946 |
| MLLT10 | NTC | LYZ | 1.68E-17 | 0.43367254 | 0.955 | 0.926 |
| MLLT10 | NTC | C12orf75 | 1.21E-16 | -0.2599525 | 0.755 | 0.919 |
| MLLT10 | NTC | CD36 | 2.20E-16 | -0.3123571 | 0.733 | 0.914 |
| MLLT10 | NTC | ELANE | 3.72E-15 | 0.49546218 | 0.996 | 0.992 |
| MLLT10 | NTC | CTSD | 1.33E-14 | 0.27913838 | 1 | 0.998 |
| MLLT10 | NTC | AZU1 | 1.18E-11 | 0.41299601 | 0.991 | 0.979 |
| MLLT10 | NTC | EMB | 1.86E-10 | 0.26864206 | 0.861 | 0.865 |
| MLLT10 | NTC | SLPI | 1.06E-09 | 0.31469803 | 0.454 | 0.32 |

| p_val_adj |
| --- |
| 1.09E-93 |
| 2.59E-67 |
| 1.43E-63 |
| 2.18E-46 |
| 8.37E-46 |
| 1.59E-40 |
| 1.58E-38 |
| 5.06E-34 |
| 2.33E-31 |
| 2.79E-30 |
| 3.00E-30 |
| 1.25E-29 |
| 2.90E-29 |
| 3.07E-29 |
| 7.48E-28 |
| 3.54E-26 |
| 1.81E-25 |
| 5.63E-25 |
| 5.02E-24 |
| 1.30E-23 |
| 5.70E-23 |
| 7.99E-21 |
| 5.87E-20 |
| 1.62E-19 |
| 1.66E-19 |
| 1.71E-18 |
| 9.12E-18 |
| 1.13E-17 |
| 2.38E-17 |
| 8.31E-17 |
| 1.08E-16 |
| 6.71E-15 |
| 8.12E-15 |
| 1.06E-14 |
| 2.71E-14 |
| 3.13E-14 |
| 2.00E-13 |
| 5.92E-13 |
| 1.58E-12 |
| 8.87E-12 |
| 1.65E-11 |
| 1.68E-11 |

| |
|---|
| 9.59E-11 |
| 6.92E-10 |
| 1.88E-09 |
| 8.07E-08 |
| 8.79E-08 |
| 1.02E-07 |
| 1.67E-07 |
| 4.21E-07 |
| 2.31E-05 |
| 1 |
| 2.17E-54 |
| 5.11E-25 |
| 1.78E-23 |
| 1.55E-21 |
| 2.72E-21 |
| 3.84E-19 |
| 2.92E-14 |
| 3.29E-13 |
| 3.07E-12 |
| 6.58E-12 |
| 4.49E-11 |
| 1.04E-08 |
| 5.89E-07 |
| 2.68E-18 |
| 6.54E-116 |
| 4.25E-107 |
| 1.65E-94 |
| 5.21E-89 |
| 9.09E-87 |
| 6.53E-81 |
| 3.63E-67 |
| 1.50E-65 |
| 2.55E-58 |
| 2.16E-56 |
| 4.67E-49 |
| 1.10E-48 |
| 1.91E-46 |
| 3.45E-46 |
| 2.42E-45 |
| 3.46E-44 |
| 4.85E-44 |
| 2.98E-43 |
| 2.10E-42 |
| 1.59E-41 |
| 2.45E-41 |

1.14E-39
5.97E-39
4.07E-37
5.65E-37
2.64E-35
1.78E-32
3.30E-32
6.55E-32
1.93E-31
1.76E-29
3.85E-29
1.30E-27
6.62E-27
1.43E-26
1.75E-26
6.41E-26
7.81E-26
7.94E-26
1.70E-25
7.86E-25
8.20E-25
5.21E-24
3.88E-23
1.09E-22
1.24E-21
1.48E-21
5.33E-21
7.52E-21
9.81E-21
2.15E-20
1.01E-19
1.68E-19
1.69E-19
7.45E-19
7.82E-19
9.24E-19
2.81E-18
1.04E-17
2.33E-17
4.10E-17
6.66E-17
7.88E-17
8.83E-17
1.08E-16
1.11E-16

| |
|---|
| 2.11E-16 |
| 2.50E-16 |
| 4.78E-16 |
| 1.07E-15 |
| 1.38E-15 |
| 1.77E-15 |
| 3.56E-15 |
| 3.59E-15 |
| 9.66E-15 |
| 1.41E-14 |
| 3.58E-14 |
| 3.92E-14 |
| 4.01E-14 |
| 6.47E-13 |
| 1.59E-12 |
| 4.03E-12 |
| 4.45E-12 |
| 1.10E-11 |
| 4.36E-11 |
| 6.80E-11 |
| 1.61E-10 |
| 5.32E-10 |
| 6.82E-10 |
| 8.42E-10 |
| 4.67E-09 |
| 2.63E-08 |
| 4.20E-07 |
| 5.82E-06 |
| 3.20E-05 |
| 5.52E-05 |
| 0.00065631 |
| 1.66E-62 |
| 1.42E-44 |
| 2.04E-42 |
| 3.41E-40 |
| 6.90E-39 |
| 1.07E-35 |
| 1.13E-35 |
| 5.40E-30 |
| 1.04E-26 |
| 1.83E-25 |
| 1.91E-23 |
| 1.97E-21 |
| 8.87E-21 |
| 1.39E-20 |

| |
|---|
| 3.24E-19 |
| 3.66E-18 |
| 1.40E-17 |
| 2.64E-17 |
| 3.72E-17 |
| 1.85E-16 |
| 3.75E-16 |
| 2.17E-15 |
| 4.85E-15 |
| 8.79E-15 |
| 7.53E-14 |
| 1.67E-13 |
| 3.49E-13 |
| 2.09E-12 |
| 2.35E-11 |
| 1.04E-10 |
| 1.55E-09 |
| 2.91E-09 |
| 4.90E-08 |
| 2.32E-05 |
| 1 |
| 4.44E-36 |
| 2.77E-31 |
| 4.61E-27 |
| 1.45E-23 |
| 3.88E-23 |
| 2.32E-17 |
| 1.91E-16 |
| 1.06E-15 |
| 7.36E-15 |
| 3.99E-13 |
| 7.65E-13 |
| 1.72E-12 |
| 9.26E-10 |
| 5.15E-09 |
| 1.18E-07 |
| 9.10E-07 |
| 0.00041587 |
| 0.00420182 |
| 2.01E-27 |
| 9.36E-18 |
| 8.45E-17 |
| 1.14E-15 |
| 1.86E-67 |
| 4.84E-51 |

1.00E-49
2.59E-47
1.46E-45
3.72E-41
3.99E-40
8.14E-39
1.02E-35
3.17E-34
4.63E-34
1.04E-32
1.30E-30
3.74E-30
7.53E-30
3.74E-29
1.26E-28
2.24E-28
2.26E-27
6.01E-27
4.42E-26
4.77E-26
7.85E-26
2.49E-25
4.22E-25
5.25E-25
7.10E-25
7.80E-25
1.74E-24
2.32E-24
3.85E-24
1.50E-23
2.64E-23
2.99E-23
8.35E-23
1.12E-22
2.11E-22
2.15E-22
2.79E-22
4.74E-22
1.86E-21
7.51E-21
4.99E-20
5.91E-20
9.02E-20
9.17E-20
1.13E-19

| |
|---|
| 1.25E-19 |
| 1.48E-19 |
| 1.64E-19 |
| 3.68E-19 |
| 4.61E-19 |
| 6.05E-19 |
| 7.63E-19 |
| 8.17E-19 |
| 1.05E-18 |
| 1.05E-18 |
| 1.11E-18 |
| 1.72E-18 |
| 2.56E-18 |
| 3.00E-18 |
| 3.68E-18 |
| 3.84E-18 |
| 3.90E-18 |
| 5.20E-18 |
| 5.44E-18 |
| 6.01E-18 |
| 6.60E-18 |
| 1.09E-17 |
| 1.36E-17 |
| 1.48E-17 |
| 1.63E-17 |
| 1.94E-17 |
| 2.13E-17 |
| 5.17E-17 |
| 1.17E-16 |
| 1.30E-16 |
| 1.54E-16 |
| 1.91E-16 |
| 2.22E-16 |
| 2.85E-16 |
| 3.14E-16 |
| 4.57E-16 |
| 4.64E-16 |
| 4.75E-16 |
| 1.28E-15 |
| 1.30E-15 |
| 2.11E-15 |
| 2.77E-15 |
| 2.88E-15 |
| 7.33E-15 |
| 1.01E-14 |

1.43E-14
1.53E-14
2.06E-14
2.15E-14
5.33E-14
5.47E-14
1.89E-13
2.33E-13
4.15E-13
7.72E-13
8.05E-13
1.13E-12
1.17E-12
1.34E-12
1.45E-12
1.85E-12
4.91E-12
5.00E-12
6.00E-12
9.83E-12
1.26E-11
1.37E-11
1.38E-11
2.77E-11
3.04E-11
3.47E-11
3.56E-11
3.68E-11
4.51E-11
5.55E-11
6.99E-11
7.06E-11
1.24E-10
1.80E-10
4.05E-10
4.82E-10
5.60E-10
5.78E-10
1.01E-09
1.06E-09
1.42E-09
1.77E-09
2.87E-09
1.39E-08
1.58E-08

| |
|---|
| 3.53E-08 |
| 5.36E-08 |
| 6.61E-08 |
| 7.49E-08 |
| 1.10E-07 |
| 1.80E-07 |
| 3.48E-07 |
| 4.07E-07 |
| 8.06E-07 |
| 8.27E-07 |
| 1.16E-06 |
| 1.49E-06 |
| 3.03E-06 |
| 4.63E-06 |
| 7.50E-06 |
| 1.01E-05 |
| 1.53E-05 |
| 1.70E-05 |
| 2.28E-05 |
| 2.87E-05 |
| 4.94E-05 |
| 6.31E-05 |
| 0.00076993 |
| 0.0078839 |
| 0.32803067 |
| 1 |
| 1 |
| 2.53E-114 |
| 1.02E-105 |
| 6.77E-87 |
| 1.34E-81 |
| 3.46E-69 |
| 2.07E-68 |
| 3.64E-67 |
| 5.97E-66 |
| 3.78E-62 |
| 2.45E-61 |
| 8.39E-57 |
| 5.40E-55 |
| 5.09E-51 |
| 1.63E-49 |
| 1.54E-48 |
| 2.27E-47 |
| 6.22E-47 |
| 9.28E-47 |

7.86E-45
9.49E-45
4.64E-44
2.77E-43
3.01E-41
3.61E-41
5.73E-41
9.08E-41
1.88E-39
7.93E-38
1.81E-36
2.26E-36
2.26E-36
3.89E-36
5.13E-36
5.52E-36
8.86E-36
1.01E-35
4.70E-35
1.27E-34
5.55E-34
4.08E-33
7.50E-33
3.62E-32
1.89E-31
1.70E-30
2.11E-30
2.19E-30
4.10E-30
5.51E-30
6.31E-30
8.05E-30
1.17E-29
3.83E-29
4.38E-29
4.93E-29
9.74E-29
6.76E-28
7.56E-28
8.17E-28
9.09E-28
1.13E-27
2.43E-27
3.76E-27
6.05E-27

1.45E-26
2.29E-26
2.76E-26
2.79E-26
3.56E-26
1.04E-25
1.18E-25
1.95E-25
6.80E-25
7.57E-25
4.06E-24
5.36E-24
1.82E-23
4.36E-23
7.22E-23
7.32E-23
8.14E-23
1.83E-22
2.13E-22
3.97E-22
4.17E-22
9.21E-22
1.32E-21
4.96E-21
8.06E-21
1.10E-20
1.48E-20
4.00E-20
5.01E-20
1.03E-19
1.39E-19
1.74E-19
3.54E-19
4.88E-19
4.93E-19
6.14E-19
1.66E-18
2.48E-18
4.48E-18
4.71E-18
4.87E-18
5.20E-18
3.58E-17
3.96E-17
4.99E-17

| |
|---|
| 6.08E-17 |
| 1.69E-16 |
| 3.44E-16 |
| 5.21E-16 |
| 8.24E-16 |
| 9.58E-16 |
| 2.09E-15 |
| 3.02E-15 |
| 4.58E-15 |
| 9.60E-15 |
| 6.09E-14 |
| 1.00E-13 |
| 2.37E-13 |
| 5.69E-13 |
| 2.07E-12 |
| 9.01E-12 |
| 6.84E-10 |
| 8.70E-10 |
| 2.12E-09 |
| 4.66E-09 |
| 4.71E-09 |
| 7.48E-09 |
| 1.14E-08 |
| 1.44E-08 |
| 2.24E-08 |
| 2.56E-08 |
| 4.88E-08 |
| 5.69E-08 |
| 3.09E-07 |
| 4.09E-07 |
| 4.11E-07 |
| 9.77E-07 |
| 2.39E-06 |
| 4.26E-06 |
| 1.30E-05 |
| 2.84E-05 |
| 6.66E-05 |
| 0.00059712 |
| 0.00059765 |
| 0.00085454 |
| 0.01915327 |
| 4.50E-62 |
| 4.89E-21 |
| 6.96E-18 |
| 8.66E-18 |

3.40E-16
4.22E-05
2.89E-35
4.49E-26
7.27E-26
7.03E-21
2.47E-18
3.52E-16
4.88E-16
9.72E-15
7.41E-14
1.99E-13
4.27E-13
3.07E-12
5.60E-12
9.45E-11
3.39E-10
3.00E-07
4.73E-06
2.70E-05

**TABLE S4**

| Organism | Gene | Primer sequence |
|---|---|---|
| Human | HOXA7 forward | 5'- TACCCCTGGATGCGGTCTT-3' |
| Human | HOXA9 forward | 5'- CACCAGACGAACAGTGAGGA-3' |
| Human | HOXA10 forward | 5'- ACACTGGAGCTGGAGAAGGA-3' |
| Human | MEIS1 forward | 5'- GATGATTCAAGCCATACAAG-3' |
| Human | GAPDH forward | 5'- GAAGGTGAAGGTCGGAGTC-3' |
| Human | HPRT forward | 5'- TGGAGTCCTATTGACATCGCC-3' |
| Human | HOXA7 reverse | 5'- CAGGTAGCGGTTGAAGTGGAA-3' |
| Human | HOXA9 reverse | 5'- TGGTCAGTAGGCCTTGAGGT-3' |
| Human | HOXA10 reverse | 5'- TCACTTGTCTGTCCGTGAGG-3' |
| Human | MEIS1 reverse | 5'- GGGGTTCCTCCTGAACGAGT-3' |
| Human | GAPDH reverse | 5'- ATGGGATTTCCATTGATGAC-3' |
| Human | HPRT reverse | 5'- AACAACAATCCGCCCAAAGGGAAC-3' |

| Antibody | Catalog Number | Manufacturer |
|---|---|---|
| KAT2A | ab217876 | Abcam |
| Flag M2 | F1804 | Sigma Aldrich |
| Vinculin | V9131 | Sigma Aldrich |
| Histone H3 | ab1791 | Abcam |
| SGF29 | ab204367 | Abcam |
| Goat anti-rabbit | AP307P | MilliporeSigma |
| Goat anti-mouse | IB23002 | ThermoFisher Scientific |
| Normal rabbit IgG | 2729 | Cell Signaling Technology |
| Histone H3K9 ac | ab4441 | Abcam |
| FLAG M2 affinity beads | A2220 | Sigma Aldrich |

| Organism | Gene and sgRNA No. | sgRNA sequence |
|---|---|---|
| Human | AFF2-1 | CTTGTCCGCTACGTTCGCCA |
| Human | AFF2-2 | GAGCTTCATGACCCACCAAG |
| Human | AFF2-3 | GTACTCTAGAGAGTAAGTCA |
| Human | CCDC101-1 | TATCTCTGAGCCACATTGAG |
| Human | CCDC101-3 | CAATGTGGCTCAGAGATACG |
| Human | CSNK2A1-1 | TGAGGATAGCCAAGGTTCTG |
| Human | CSNK2A1-2 | AGTCACATGTGGTGGAATGG |
| Human | CSNK2A1-3 | GATTGATCATGAGCACAGAA |
| Human | CSNK2B-2 | CAGGCAGCCGAGATGCTTTA |
| Human | CSNK2B-3 | TCTCGGCTGCCTGCTCAATC |
| Human | DOT1L-1 | GGGAGCGAATCGCCAACACG |
| Human | DOT1L-2 | GCAGCAGGTCTACAACCACT |
| Human | DOT1L-3 | GTCCACAAACAGGTCGTCGT |
| Human | ENY2-1 | TAGCTCTCAGCAACTCTTTG |

| | | |
|---|---|---|
| Human | ENY2-2 | CAGTTGAAGGCACACTGTAA |
| Human | ENY2-3 | AAAGGCTGGCATGCTGAGCA |
| Human | JADE3-1 | TTTGGACATAACACACATTG |
| Human | JADE3-2 | AGCCCATTTAGTCCCTGTCT |
| Human | JADE3-3 | GTCACCTGTGCCTTTGAGCA |
| Human | KAT7-1 | TCTCATCGTGAGATACATTG |
| Human | KAT7-2 | TGTCCTAGAGAGTTACAGCC |
| Human | KAT7-3 | ACAGACAGTTCAGAAAGTGA |
| Human | KMT2A-1 | GATCTGAAAGCCATCATCAC |
| Human | KMT2A-2 | TCATAACATTTGTCACAGAG |
| Human | KMT2A-3 | CCATCGGCACCAACCTGCGC |
| Human | MLLT1-1 | GAGCTTGTACCGGAACTCCG |
| Human | MLLT1-3 | CAGCCATGGACAATCAGGTG |
| Human | NTC-1 | ATGGAAGAGCGTCATGACTT |
| Human | NTC-2 | CCAGTTGCTCTGGGGGAACA |
| Human | BRD4-1 | CACCAAACTCCTGAGCATCA |
| Human | BRD4-2 | CTGAGCATCACGGTACTCAC |
| Human | MLLT10-1 | ACATGTCATGCAAGCACCAG |
| Human | MLLT10-2 | TCCATCATGCCGATACTGCA |
| Human | MLLT3-1 | CAGCGGAGGTGATTCACTGG |
| Human | MLLT6-1 | GCTGTGCCCACACAAAGACG |
| Human | MLLT6-2 | CGTTGGCAAATTGCACCTCG |
| Human | MLLT3-1 | TGCAGGTGAAGCTGGAGCTG |
| Human | AFF4-1 | GACTTCCAGTAGCTCCAAGG |
| Human | AFF4-2 | ATGTGCTGCGTATGAAAGAA |
| Mouse | Sgf29-1 | AACCGCCAAGACTGATGCGG |
| Mouse | Sgf29-2 | CACAACCGCCAAGACTGATG |
| Mouse | Sgf29-3 | AGAGGCCTCGAAGCTTTGTC |
| Mouse | Sgf29-4 | GTCGAGAAGTGAGCACAACT |
| Mouse | Sgf29-intron-1 | CCCTCTGGTCCTGCTGTGTC |
| Mouse | Sgf29-intron-2 | ACTGTAATCATCTCAAAATT |

| **Experiment** |
|---|
| Immunoblotting |
| Immunoblotting |
| Immunoblotting |
| Immunoblotting |
| Immunoblotting |
| Immunoblotting |
| Immunoblotting |
| ChIP-seq |
| ChIP-seq |
| ChIP-seq |

| **Used in CROP-seq** | **Used in competition assays** |
|---|---|
| ✓ | ✓ |
| | ✓ |
| ✓ | ✓ |
| ✓ | ✓ |
| ✓ | ✓ |
| ✓ | ✓ |
| ✓ | ✓ |
| ✓ | ✓ |
| ✓ | ✓ |
| ✓ | ✓ |
| ✓ | ✓ |
| ✓ | ✓ |
| | ✓ |
| | ✓ |

| | |
|---|---|
| | √ |
| | √ |
| | √ |
| | √ |
| | √ |
| √ | √ |
| √ | √ |
| | √ |
| √ | √ |
| √ | √ |
| | √ |
| √ | √ |
| √ | √ |
| √ | √ |
| √ | √ |
| √ | |
| √ | |
| √ | |
| √ | |
| √ | |
| √ | |
| √ | |
| √ | |
| √ | |
| √ | |
